# TFIIFα interacts with the Topoisomerase VI complex and selectively controls the expression of genes encoding PPR proteins involved in organellar RNA editing in Arabidopsis

**DOI:** 10.1101/2022.04.23.489258

**Authors:** Laura Dimnet, Cécile Vriet, Dorine Achard, Cécile Lecampion, Christian Breuer, Ludivine Soubigou-Taconnat, Keiko Sugimoto, Etienne Delannoy, Christophe Laloi

## Abstract

Communication between organelles and the nucleus is referred to as anterograde (nucleus to organelle) and retrograde (organelle to nucleus) signalling. In plants, the pentatricopeptide repeat (PPR) proteins represent a large family of nuclear-encoded proteins that are required for post-transcriptional control of chloroplast and mitochondria gene expression, and hence play a central role in the nuclear anterograde control of organelle genome expression. How *PPR* gene expression is controlled and regulated by retrograde signals is, however, still unknown. Here, we report a significant role for the general transcription factor TFIIF α-subunit (TFIIFα) in controlling *PPR* gene expression in Arabidopsis. First, we found that TFIIFα interacts with the BIN4 subunit of the Topoisomerase VI (Topo VI). Transcriptome analysis of TFIIF and Topo VI mutant lines then revealed that many PLS-type PPR genes involved in RNA editing are reciprocally controlled by TFIIF and Topo VI. The misexpression of *CLB19* and *DYW1* genes in two allelic *tfIIfα* mutants was associated with editing impairments in their plastid target RNAs *rpoA* and *ndhD*, respectively. Interestingly, we also detected a change in NDH activity in *tfIIfα* plants. We also show that TFIIFα and Topo VI coordinate the expression of NDH subunits encoded by the nuclear and plastid genomes. These results reveal the crucial role of the nuclear TFIIFα and Topo VI complexes in controlling plastid genome expression at multiple levels of regulation, including the particular regulation of PPR gene expression.

## INTRODUCTION

Proteins encoded by plastid and mitochondrial genomes are not sufficient to support the development and the metabolism of organelles, and most of the proteins they contain are nuclear-encoded and synthesized in the cytosol before organellar targeting. Consequently, organellar proteomes from separated genomes require coordinated expression between cellular compartments to maintain organelle homeostasis (Woodson and Chory, 2008). This regulation includes both the anterograde (nucleus-to-organelles) and the retrograde (organelle-to-nucleus) signalling. In a genetic screen designed to identify Arabidopsis *(Arabidopsis thaliana*) genes involved in singlet oxygen (^1^O_2_)-mediated retrograde signalling, a mutant called *constitutive activator of AAA-ATPase 39* (*caa39*) was isolated where ^1^O_2_-responsive genes are constitutively up-regulated under steady-state conditions, and are not further activated under ^1^O_2_–producing conditions (Baruah et al., 2009). This mutant is affected in the gene encoding the A-subunit of Topoisomerase VI (Topo VI) and reveals the involvement of Topo VI in ^1^O_2_ retrograde signalling as well as a putative dual function as a transcriptional activator and repressor, depending on environmental conditions.

Topo VI belongs to the topoisomerase superfamily, a class of enzymes which resolve DNA topological constraints by relaxing supercoils, knots and catenanes in prokaryotic and eukaryotic cells. During DNA processes such as transcription, supercoils usually occur on double-helical structure and if this phenomenon is persistent, it can lead to transcriptional regulation defects as well as DNA breaks that are damaging for gene expression and cell viability (Corbett and Berger, 2003). The structure and mechanism of action of Topo VI have been characterized in Archaea where the complex was originally discovered (Corbett et al., 2007; Graille et al., 2008). With a heterotetrameric A2B2 structure and ATP-dependent double-stranded break activity, Topo VI belongs to the type IIB Topoisomerases. The complex is composed of A-subunits (Topo VIA) involved in DNA binding and cleavage and B-subunits (Topo VIB) that allow ATP fixation and hydrolysis. In contrast to Archaea, plant Topo VI possesses two additional subunits, BIN4/MID (AT5G24630, hereafter called BIN4) and RHL1/HYP7 (AT1G48380, hereafter called RHL1) that interact together and with Topo VIA (Breuer et al., 2007; Sugimoto-Shirasu et al., 2005; Kirik et al., 2007). The function of BIN4 and RHL1 in the Topo VI complex remains unclear. However, given that BIN4 and RHL1 possess sequence similarity to the regulatory C-terminal region of animal Topo IIα(Gadelle et al., 2003) and exhibits stable DNA binding *in vitro*, it has been hypothesized that BIN4 and RHL1 could help the enzyme to hold the substrate DNA during the decatenation reaction (Sugimoto-Shirasu et al., 2005; Breuer et al., 2007). Topo VI knock-out mutant plants, whatever the subunit affected, show a similar pleiotropic phenotype: severe growth inhibition, ploidy decrease, and defective skotomorphogenesis (Yin et al., 2002; Hartung et al., 2002; Sugimoto-Shirasu et al., 2002, 2005; Schrader et al., 2013; Kirik et al., 2007; Breuer et al., 2007). Overexpression of rice *OsTOP6A3* and *OsTOP6B* in Arabidopsis plants confers stress tolerance that coincides with enhanced induction of many stress-responsive genes (Jain et al., 2006). More recently, we reported that Topo VI is a key regulatory factor during the activation of ROS-responsive genes (Šimková et al, 2012). Taken together, these results emphasize the crucial role of Topo VI in plant stress responses. However, how Topo VI controls the expression of specific genes remains obscure.

Here, we reveal that the α-subunit of the general transcription factor TFIIF (TFIIFα) interacts with the BIN4 subunit of the Topo VI complex. RNA-sequencing carried out in two different *tfIIfα* mutants showed a massive repression of genes encoding pentatricopeptide repeat (PPR) proteins involved in organellar RNA editing. Remarkably, these *PPR* genes were inversely affected in the Topo VI mutant *caa39*. In *tfIIfα* mutants, misexpression of two of *PPR* genes, *CLB19* and *DYW1*, was associated with editing impairments in their target RNAs in the plastid, *rpoA* and *ndhD*, respectively. Concurrently, mutations in TFIIFα and Topo VI also affected the expression of NDH subunits encoded by both the nuclear and plastid genomes. Finally, we detected a change in NDH activity as a likely consequence of these defects in *tfIIfα* plants.

## RESULTS

### BIN4 interacts with the General Transcription Factor RAP74/TFIIFα

To determine whether BIN4 may interact with other proteins and hence govern the activity of the Topo VI complex, we performed a yeast two-hybrid screen for Arabidopsis cDNAs encoding proteins that can interact directly with BIN4. The screen was performed under two different stringency conditions (Hybrigenics, Supplemental Table 1). Respectively 36 and 86 putative interactions (positive colonies) were analyzed. A strong interaction was confirmed with the Topo VI subunit RHL1, for which respectively 8 and 19 clones were identified under the two stringency conditions, thereby demonstrating the reliability of the screening procedure (Supplemental Table 1). However, the strongest interactor identified during these screens was not described before and corresponds to the ATRAP74/TFIIFα (AT4G12610, hereafter called TFIIFα), for which respectively 8 and 33 clones were identified under the two stringency conditions (Supplemental Table 1). TFIIFα/RAP74 is the large subunit of the general transcription factor TFIIF, which is needed for accurate transcription initiation and stimulates elongation by RNA polymerase II (Pol II) in metazoa. After transcription termination, the interaction of TFIIF with TFIIF-associated C-terminal domain (CTD) phosphatase 1 (FCP1), which catalyzes the Ser2 and Ser5 dephosphorylation of Pol II CTD, is essential for Pol II recycling at new promoters (Lin et al., 2002; Abbott et al., 2005; Kimura et al., 2002; Archambault et al., 1998; Palancade et al., 2002; Yang et al., 2009; Kumar et al., 2013; Nguyen et al., 2003; Kamada et al., 2003).

**Table 1.**
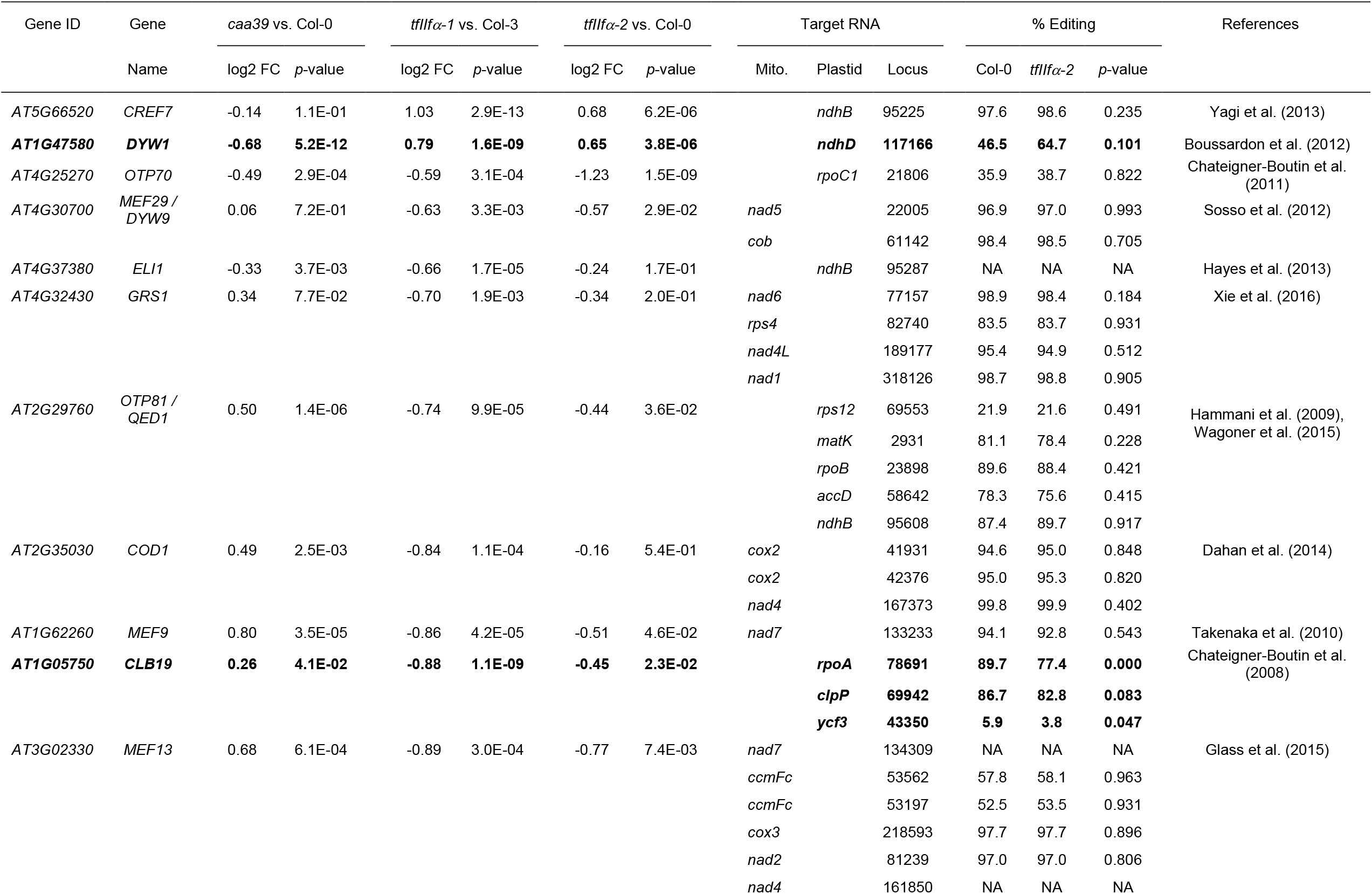
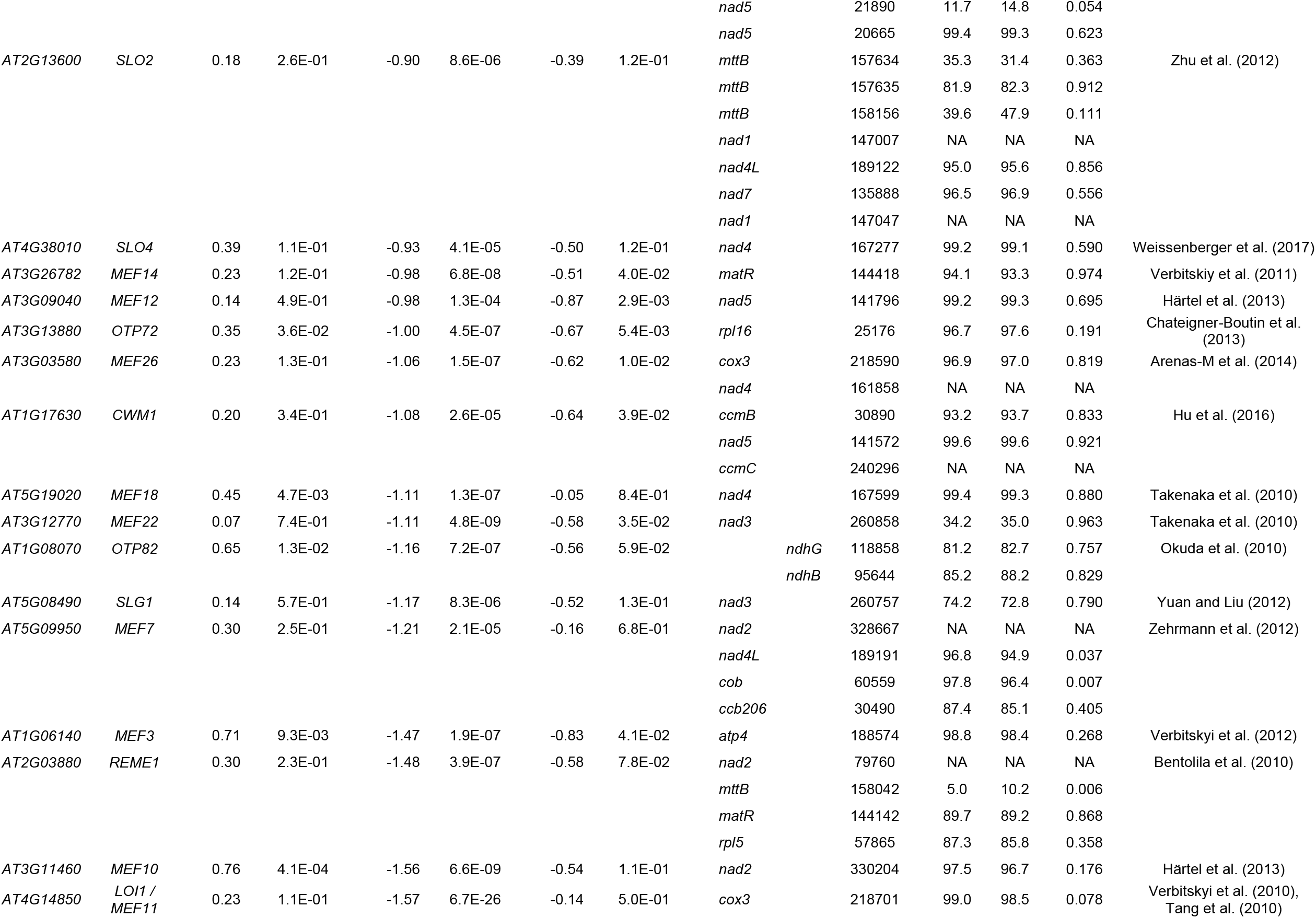
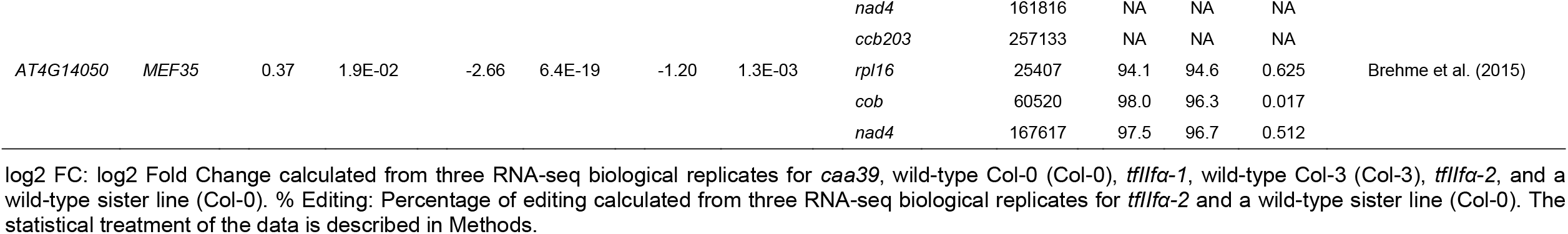
Deregulated PLS-type PPR gene expression related to target site editing in *tfIIfα* mutants.3

We performed an independent yeast two-hybrid assay to further confirm the interaction between BIN4 and TFIIFα and to determine whether TFIIFα could directly interact with other components of the plant Topo VI complex. As shown in Figure 1A, TFIIFα strongly interacted with BIN4, but not directly with other subunits of the Topo VI complex. In order to confirm the interaction between BIN4 and TFIIFα *in planta*, we performed a bimolecular fluorescence complementation (BiFC) assay. The N-terminal and the C-terminal parts of the yellow fluorescent protein (nYFP and cYFP) were fused to TFIIFα and BIN4, and then were transiently co-expressed in agro-transformed *Nicotiana benthamiana* mesophyll cells. We also co-expressed the Topo VI A subunit fused with the cyan fluorescent protein (AtTOP6A-CFP) to visualize the nuclei and localize the interaction of BIN4 and TFIIFα. The BiFC assay revealed reconstituted YFP fluorescence that was specifically localized in the nucleus with a speckled-like distribution. This result confirms the interaction between BIN4 and TFIIFα (Figure 1B). Interestingly, the YFP fluorescence pattern was very similar to AtTOP6A-CFP fluorescence, suggesting that the BIN4-TFIIFα interaction loci co-localise with Topo VI within the nucleus (Figure 1B).

**Figure 1.**
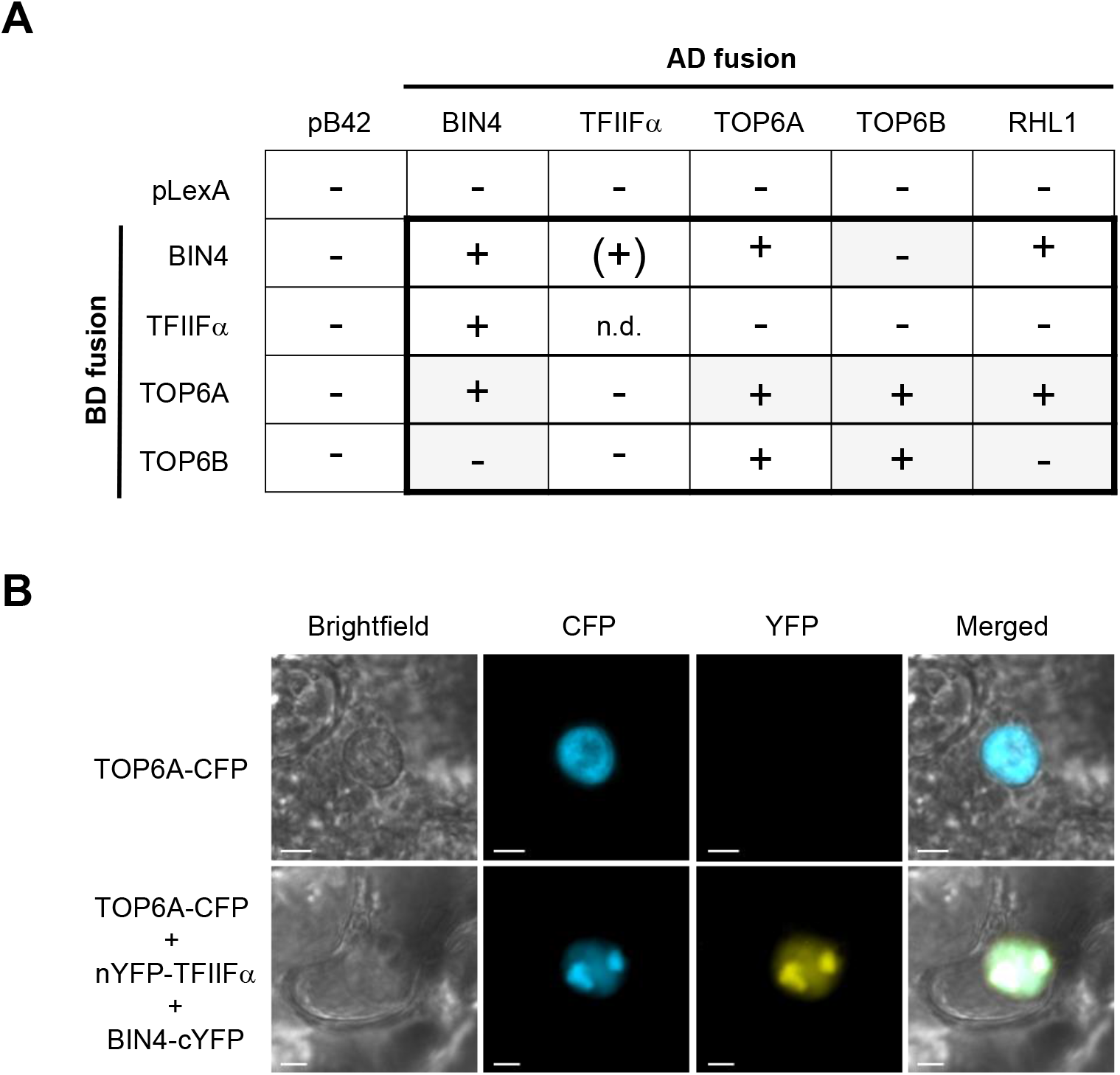
Interaction between BIN4 and TFIIFα. **(A)** Yeast two-hybrid analysis of BIN4, TFIIFα, RHL1, TOP6A and TOP6B protein interactions. The *Arabidopsis thaliana BIN4, TFIIFα, RHL1, TOP6A* and *TOP6B* genes were cloned into pLexA (BD, binding domain fusion) and pB42 (AD, activator domain fusion) vectors, and their protein interactions were detected by induction (+: strong induction; (+): weak induction) or no induction (-) of the lacZ reporter gene. n.d.: not determined. Grey boxes: data from Breuer *et al*. 2007. **(B)** Transiently agro-transformed mesophyll cells from *N. benthamiana* leaves expressing different combinations of BiFC constructs and/or TOP6A fused with CFP, as indicated. Scale bars: 2 µm.

### Two *TFIIFα* transcripts are generated from a single-copy gene in Arabidopsis

The Arabidopsis genome contains one *TFIIFα* gene (*AtRAP74, AT4G12610*) that encodes at least two transcripts, *TFIIFα*.*1* (*AT4G12610*.*1*) and *TFIIFα*.*2* (*AT4G12610*.*2*) (Figure 2A, www.arabidopsis.org). The conserved C-terminal domain of TFIIFα.1, which is required for the interaction with the two FCP1-type CTD-phosphatase proteins CPL3 and CPL4 (Bang et al., 2006; Li et al., 2014) is encoded by exons 9 and 10. Quite unusually, this CTD-phosphatase interaction domain is duplicated in the TFIIFα.2 peptide (encoded by exon 11), whereas part of the last intron (intron 10) constitutes the 3’UTR of *TFIIFα*.*1* transcript (Figure 2A). The presence of a TFIIFα isoform with two CTD-phosphatase interaction domains could be identified only within the Arabidopsis genus. Quantitative RT-PCR analysis with different primer pairs designed to amplify specifically *TFIIFα*.*1, TFIIFα*.*2* or both transcripts revealed that *TFIIFα*.*2* was much less abundant than *TFIIFα*.*1* (Figure 2B). This unequal abundance of the two transcripts was further confirmed by inspection of publically available RNA-seq data (www.araport.org).

**Figure 2.**
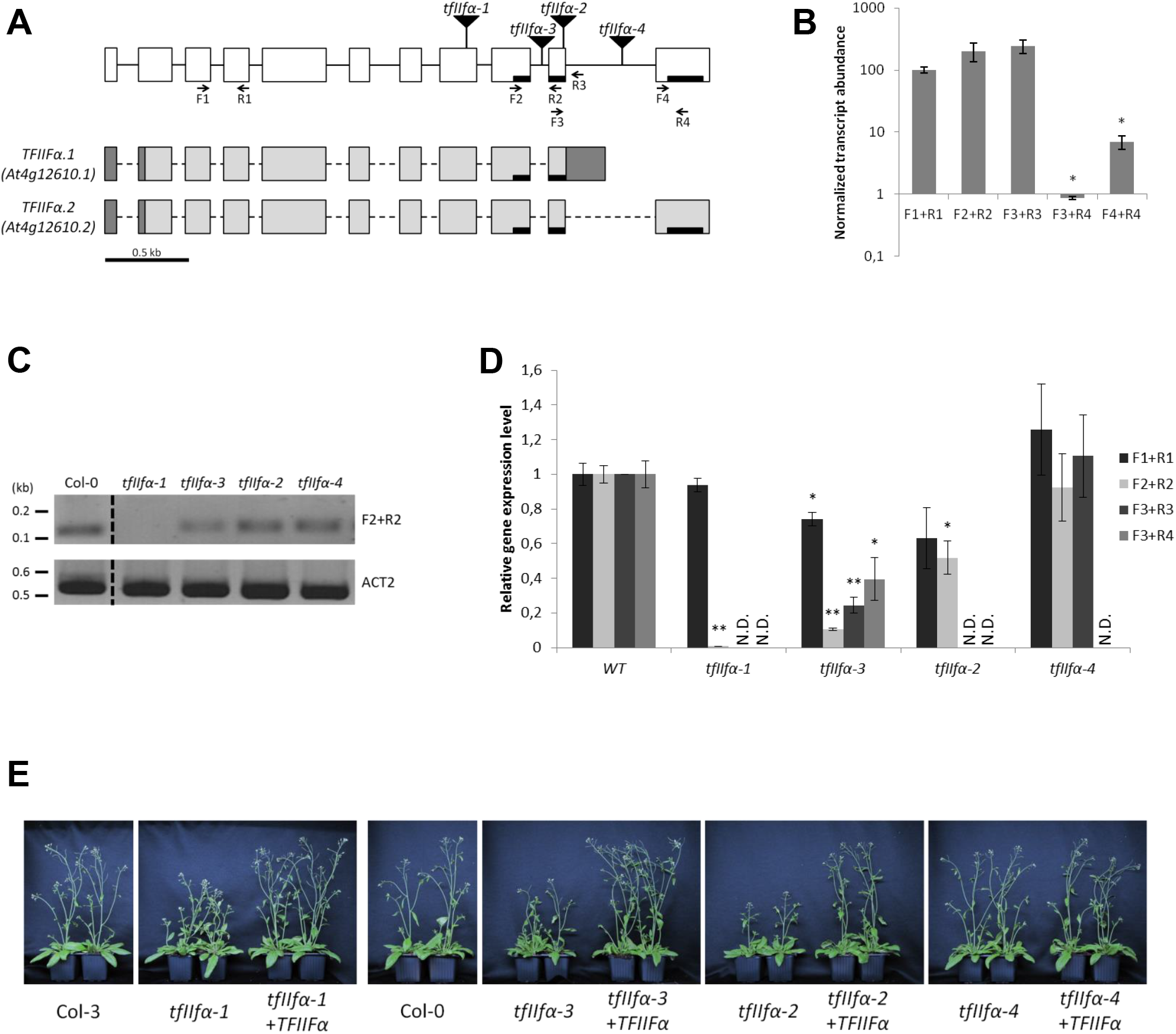
Characterization of *tfIIfα* mutants. **(A)** Schematic structure of the *TFIIFα* (*At4g12610*) gene and the two major *TFIIFα* transcripts. Boxes and lines represent exons and introns, respectively. 5’-UTR and 3’-UTR in *TFIIFα* transcripts are shown as dark grey boxes. Sequences encoding CTD-phosphatase interaction domains are shown as black bars. Locations of the T-DNA insertions and the primers used for PCR are shown as triangles and arrows, respectively. **(B)** RT-qPCR-based analysis of *TFIIFα* transcripts in aerial parts of 6-day-old wild-type plants with different primer pairs. Different primer pairs were designed to amplify specifically TFIIFα.1 (primer pair F3+R3), TFIIFα.2 (primer pairs F3+R4 and F4+R4) or both transcripts (primer pairs F1+R1 and F2+R2). Transcript levels were expressed relative to the levels of transcripts detected with the F1+R1 primer pair. Error bars represent standard deviation from biological triplicates. Significant expression differences between transcripts were estimated with a t-test: * if p-value < 0.01. **(C)** RT-PCR-based *TFIIFα* transcript analysis in the four mutants compared to Col-0 wild-type plants. RT-PCR was performed with the F2+R2 primer pair; *ACTIN 2* (*At3g18780*) was used as control. **(D)** RT-qPCR-based *TFIIFα* transcript level analyses in the four *tfIIfα* relative to corresponding wild-type plants (Col-3 or Col-0). Error bars represent standard deviation from biological duplicates. Significant expression differences between *tfIIfα* and wild-type lines were estimated with a t-test: ** if p-value < 0.01 and * if p-value < 0.05. N.D.: not detected. **(E)** Phenotype of *tfIIfα* mutants and their complemented lines. According to their ecotype, 40-day-old *tfIIfα-1* plants were compared to Col-3 wild-type plants while 38-day-old *tfIIfα-2, -3*, and *-4* lines were compared to Col-0 wild-type plants.

### TFIIFα mutants exhibit growth defects

TFIIF function has been established from a wealth of experiments mostly performed in human and yeast, on the one hand, and using a limited number of model TATA box-containing promoters, on the other hand (Luse, 2012). The role of TFIIF, which is still incompletely understood and seem to differ in yeast and mammals, was nearly completely uninvestigated in plants until recently (Babiychuk et al., 2016). In order to describe the role of TFIIFα in Arabidopsis, we characterized different *TFIIFα* T-DNA insertion mutants. When we started this work, only the SAIL_1171_F02 line (hereafter named *tfIIfα-1*) in the Col-3 ecotype background was available. This line carries a T-DNA in the TFIIFα CDS (exon 8). More recently, T-DNA-sequencing programs have allowed the identification of new *tfIIfα* mutant lines in the Col-0 ecotype (O’Malley et al., 2015). The T-DNA insertion in *tfIIfα-2* (SALKseq_038203) is localized in exon 10 and interrupts the first CTD-phosphatase interaction domain, and the T-DNA insertions in *tfIIfα-3* (SALKseq_038203) and *tfIIfα-4* (SALKseq_095102) are inserted in introns 9 and 10, respectively (Figure 2A).

In order to determine the impact of these insertions on the *TFIIFα* transcript levels, we performed RT-PCR and quantitative RT-PCR with several primer pairs distributed along the *TFIIFα* gene (Figure 2A). This analysis revealed that *tfIIfα-1* could not produce any transcript that would allow the synthesis of a protein containing any CTD-phosphatase interaction domain (Figures 2C and 2D). Similarly, *tfIIfα-2* could barely produce any TFIIFα peptide containing a complete CTD-phosphatase interaction domain (Figures 2D). On the contrary, *TFIIFα*.*1* or *TFIIFα*.*2* native transcripts are still present, although to a lower level, in the *tfIIfα-3* mutant (Figures 2C and 2D). These results suggest that intron 9 splicing was reduced but not completely abolished in *tfIIfα-3* as a consequence of the T-DNA insertion. Finally, the *tfIIfα-4* mutant was the only one with a T-DNA insertion that would impair only the second, and not the first, CTD-phosphatase interaction domain (Figure 2D).

The phenotype of the four *tfIIfα* mutant lines was first assessed under long-day conditions (16 h light / 8 h dark) in soil. Plant growth appeared differentially affected: *tfIIfα-1* was smaller than wild-type Col-3, as were the *tfIIfα-2* and *tfIIfα-3* mutants compared to wild-type Col-0 (Figure 2E). However, growth inhibition during the vegetative stage was slightly less pronounced in *tfIIfα-3* than in *tfIIfα-2* and *tfIIfα-1* plants (Figure 2E and Supplemental Figure 1A). The *tfIIfα-4* mutant displayed a wild-type phenotype (Figure 2E and Supplemental Figure 1A). In all cases, complementation of the various mutant lines with a wild-type copy of *TFIIFα* restored a wild-type phenotype (Figure 2E), and wild-type, or above wild-type, gene expression levels (Supplemental Figure 1B). These phenotypes are consistent with the molecular defects of the different alleles: the degree of impairment of the first CTD-phosphatase interaction domain correlates with the severity of the phenotype. Collectively, these results also suggest that the second CTD-phosphatase interaction domain is dispensable for TFIIFα function, in agreement with its general absence in TFIIFα orthologues.

### TFIIFα defects mainly affect the expression of PLS-type PPR genes that are inversely affected in the Topo VI mutant *caa39*

In order to investigate the role of TFIIFα in plant gene expression, we performed an RNA-seq analysis of *tfIIfα-1* and Col-3 wild-type plants. Strikingly, genes coding for pentatricopeptide repeat (PPR) proteins were strongly enriched in down-regulated genes in *tfIIfα-1*: they represent 20.1% of genes down-regulated more than 2 times (13.8% of genes down-regulated more than 1.5 times), whereas PPR genes account for only 1.4% of total genes in the genome and 2.5% of expressed genes in our RNA-seq analysis (Figure 3A). Conversely, only 0.8% of genes up-regulated more than 2 times (1.3% of genes up-regulated more than 1.5 times) encode PPR genes. PPR proteins are nuclear encoded and targeted to mitochondria or plastids where they perform post-transcriptional functions. They are classified into two major subfamilies: P-type PPR proteins are mostly involved in RNA stabilization, splicing and translation; PLS-type PPR proteins, which are further divided into five subclasses according to their C-terminus (PLS, E1, E2, E+ and DYW subgroups), are primarily involved in RNA editing in organelles. Remarkably, PLS-type PPR genes were strongly over-represented among PPR genes that are constitutively down-regulated in *tfIIfα-1* (79.2% of repressed PPR genes, or 122 out of 154 genes down-regulated more than 1.5 times) (Figure 3B and Supplemental Table 2), whereas P-type PPR genes were mainly up-regulated (80% of induced PPR gene, or 8 out of 10 genes up-regulated more than 1.5 times) (Figure 3B and Supplemental Table 2), showing that *tfIIfα-1* mutation mostly affected the expression of PPR genes involved in RNA editing. However, we also noticed that genes located in a 1.5 Mb region of chromosome IV, from the T-DNA insertion site in *TFIIFα* to the first exon of *At4g15610*, were down-regulated in *tfIIfα-1* (Supplemental Figure 2A). This region contains 411 genes that were all down-regulated except *At4g14690* (*ELIP2*) (Supplemental Figure 2B). As the repressed region precisely follows the T-DNA locus, the T-DNA insertion his likely to be responsible for the translocation of the 1.5 Mb region elsewhere in the *tfIIfα-1* genome, resulting in a global down-regulation. This translocation hypothesis was further supported by DNA-sequencing of the *tfIIfα-1* genome (Supplemental Figure 2C) that also revealed the loss of an approx. 900 bp-long region containing exons 9 and 10 and the 3’ end of exon 8 (Supplemental Figure 2D). Because of this chromosomal rearrangement in *tfIIfα-1*, we investigated PPR gene expression in the three other *tfIIfα* mutants and their complemented lines by RT-qPCR. Five PPR genes were chosen on the basis of their high down-regulation (*At1g03510, At2g36980*, and *At5g47460*) or up-regulation (*At1g47580* also called *DYW1*, and *At2g35130*) in *tfIIfα-1* (Supplemental Table 2). In agreement with RNA-seq data in *tfIIfα-1, At1g03510, At2g36980*, and *At5g47460* were repressed whereas *DYW1* and *At2g35130* were induced in *tfIIfα-2* and *tfIIfα-3* mutants (Figure 3C), though to a lesser extent in the less severe mutant *tfIIfα-3*. In contrast, PPR gene expression levels in *tfIIfα-4* and the complemented lines were very similar to those observed in wild-type plants (Figure 3C).

**Figure 3.**
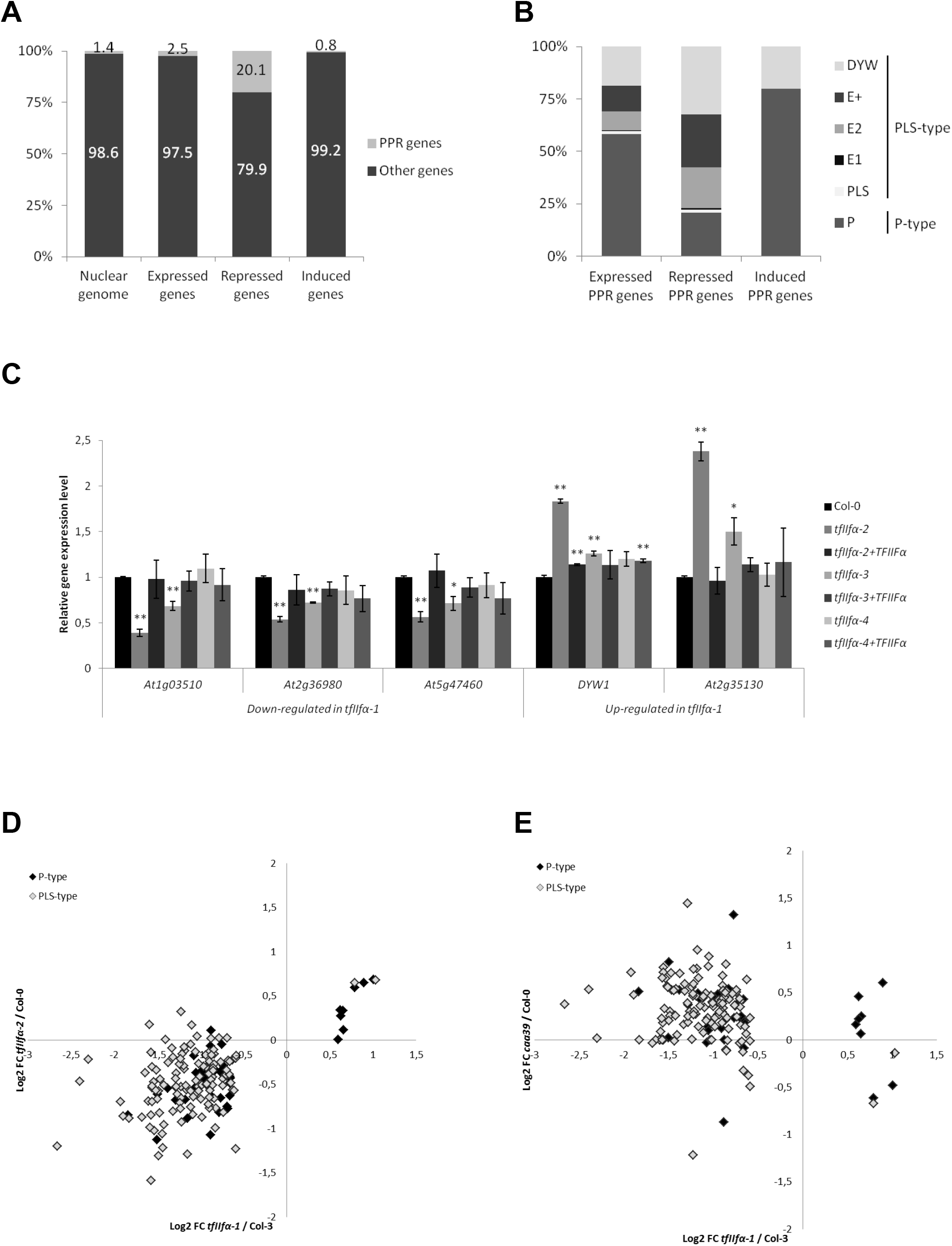
TFIIFα and Topo VI exert opposite control over PPR gene expression. **(A)** Proportion of PPR genes in the Arabidopsis nuclear genome and among expressed, repressed and induced (> 2-fold in RNA-seq, p-value < 0.01) genes in *tfIIfα-1* mutant. **(B)** Distribution of expressed, repressed and induced (> 1.5-fold in *tfIIfα-1*, p-value < 0.01) PPR genes according to their subgroups. **(C)** Expression of five representative PPR genes measured by RT-qPCR in *tfIIfα-2, tfIIfα-3, tfIIfα-4* and their respective complementation lines relative to wild-type Col-0. Error bars represent standard deviation from biological triplicates. Significant expression differences between mutant and wild-type lines were estimated with a *t*-test: ** if p-value < 0.01 and * if p-value < 0.05. **(E)** Scatter-plot comparative analysis of PPR gene expression in *tfIIfα-1, tfIIfα-2* **(D)** and *caa39* **(E)**. PPR genes repressed or induced more than 1.5-fold in *tfIIfα-1* (p-value < 0.01) have been plotted. P-and PLS-type PPR genes are represented by black and grey diamonds, respectively.

We further confirmed globally the PPR gene deregulation by RNA-seq analysis of the *tfIIfα-2* mutant, whose phenotype is similar to *tfIIfα-1*. Among PPR genes which are down- and up-regulated more than 1.5 times in *tfIIfα-1*, we observed the same tendency in *tfIIfα-2* (Figure 3D and Supplemental Table 2). In particular, PLS-type PPR genes down-regulated in *tfIIfα-1* mutants were also massively down-regulated in *tfIIfα-2*. As TFIIFα is a protein interactor of Topo VI, we then asked whether Topo VI might also control the expression of PPR genes, by assessing the expression of PPR genes in *caa39* and the respective Col-0 wild-type plants. Surprisingly, PPR genes down-regulated more than 1.5 times in *tfIIfα-1* are on the contrary mainly up-regulated in *caa39* (Figure 3E and Supplemental Table 2).

### Misexpression of PLS-type PPR genes in *tfIIfα* mutants results in RNA editing defect in organelles

Because the majority of PPR genes that are deregulated in *tfIIfα-1* and *tfIIfα-2* mutants encode PLS-type PPR proteins involved in C to U editing in chloroplasts and mitochondria, we then investigated whether PPR gene misexpression could lead to RNA editing defects in the organelles of *tfIIfα-2* plants. Total RNA-seq analysis detected 693 and 271 edited sites in mitochondrial and plastid RNAs, respectively (Supplemental Dataset 1). Among these, 93 sites (80 mitochondrial and 13 plastid target cytosines; rate 0.05, p-value < 0.05) showed significantly different editing levels between *tfIIfα-2* and wild-type (Supplemental Dataset 1). However, because the majority of these sites are edited by unknown PPR proteins, it was not possible to associate editing defects with PPR deregulation in a global manner. Instead, we examined individual sites that are edited by known PLS-PPR proteins and whose gene expression is affected in *tfIIfα-1* and *tfIIfα-2*. Among the PLS-PPR genes that were deregulated in *tfIIfα* mutants, 28 encoded PLS-PPR proteins with known targets, of which 26 were repressed both in *tfIIfα*.*1* and *tfIIfα*.*2* (Table 1). Most of the cytosines targeted by those 28 PLS-PPR were not differently edited between *tfIIfα-2* and wild-type Col-0 (Table 1), suggesting that the partial repression of those PLS-PPR genes in *tfIIfα-2* was not sufficient to reduce the editing of their targets. However, the editing of the well characterized *rpoA* site 78691, which was significantly reduced in *tfIIfα-2* (89.7% of edition in wild-type, 77.4% in *tfIIfα-2*, p-value = 0.00025, Table 1), was correlated with the reduced expression level of *CLB19* whose protein is required for *rpoA* 78691 editing (Chateigner-Boutin et al., 2008). Interestingly, two other sites that required the PLS-PPR protein CLB19 for their editing, *clpP* 69642 and *ycf3* 43350 (Chateigner-Boutin et al., 2008), were also less efficiently edited in *tfIIfα-2*; however, their reduced editing was not significant in our RNA-seq analysis (Table 1). Therefore, to confirm these editing defects we used Sanger sequencing on wild-type Col-0, *tfIIfα-2* and *tfIIfα-4* mutants, as well as wild-type Col-3 and *tfIIfα-1*. First of all, we confirmed, in independent RT-qPCR experiments, the down-regulation of *CLB19* in *tfIIfα-2*, whereas *CLB19* expression in *tfIIfα-4* was similar to wild-type (Figure 4A). This analysis confirmed the markedly reduced editing of *rpoA* 78691 in *tfIIfα-2* and *tfIIfα-1*, but not in *tfIIfα-4* (Figure 4B and Supplemental Figure 3A). This indicates that the editing defect was genetically linked to TFIIFα mutations that disrupt both CTD-phosphatase interaction domains and which lead to the down-regulation of *CLB19*. The second CTD-phosphatase interaction domain that is missing in *tfIIfα-4* is dispensable for this function. Sanger sequencing also confirmed the reduced editing of *clpP* 69942 in *tfIIfα-2* and *tfIIfα-1* (Supplemental Figure 3A and 3B). *ycf3* 43350 editing levels were too low to allow the validation of editing differences by Sanger sequencing (Supplemental Figure 3A and 3B).

**Figure 4.**
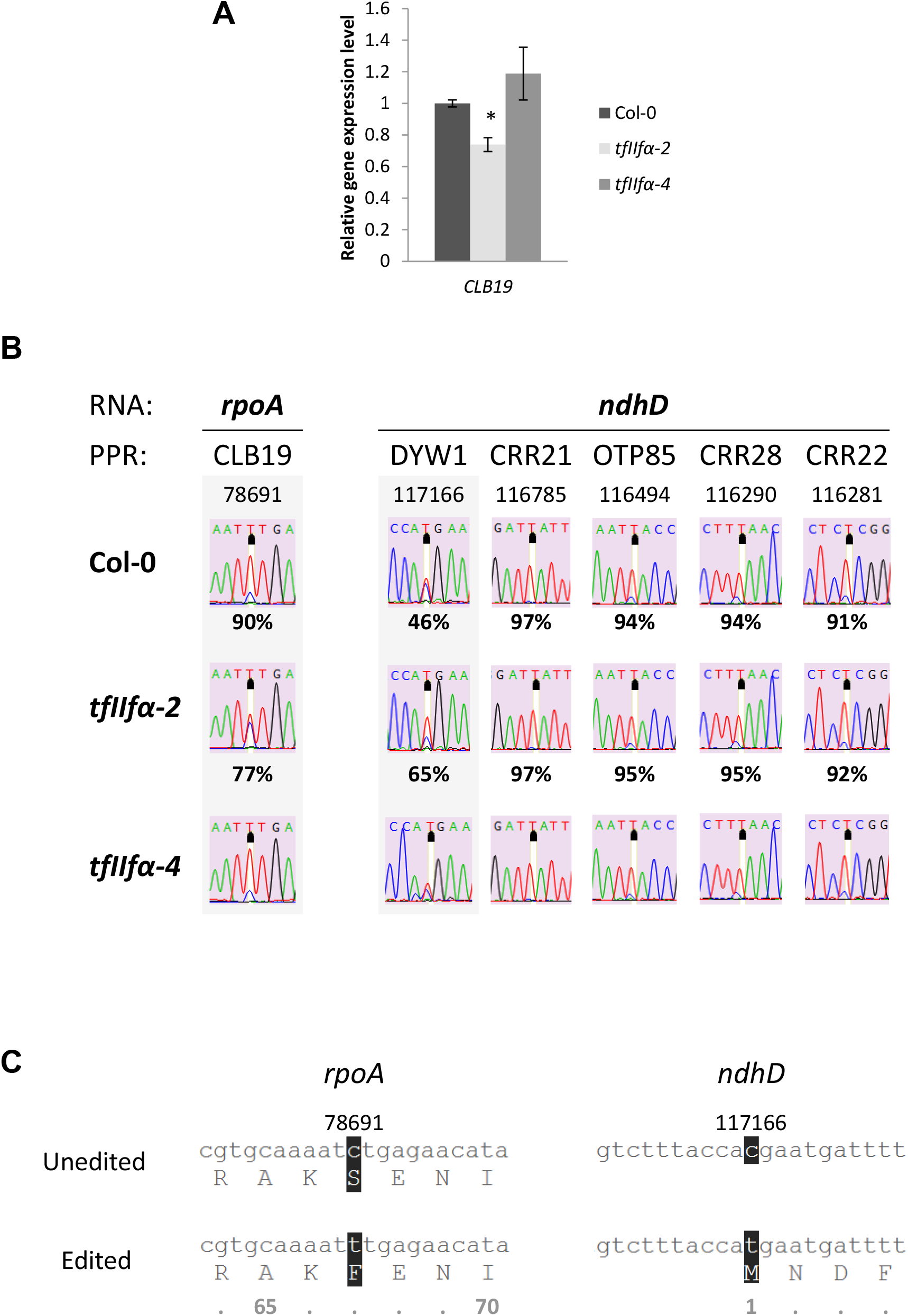
The misexpressions of *CLB19* and *DYW1* PPR genes in *tfIIf*α-2 mutant are correlated with editing level impairment in *rpoA* and *ndhD* RNA. **(A)** Expression of PPR gene *CLB19* measured by RT-qPCR in *tfIIfα-2* and *tfIIfα-4* relative to wild-type Col-0. Error bars represent standard deviation from biological duplicates. Significant expression difference between *tfIIfα* and wild-type lines were estimated with a t-test : * if p-value < 0.05. ***(B)*** *rpoA* and *ndhD* editing levels measured by Sanger sequencing in wild-type Col-0, *tfIIfα-2*, and *tfIIfα-4*. Chromatograms of *rpoA* (78691) and *ndhD* (117166) edited sites targeted by the CLB19 and DYW1 PPR proteins, respectively (grey backdrop). For *ndhD*, editing at the genomic position 117166 is compared with those of four other loci not edited by DYW1. Under each chromatogram is indicated the editing percentage detected in RNA-seq. **(C)** Comparison of nucleic acid and protein sequences of *rpoA* and *ndhD* depending on editing process or not at genomic positions 78691 and 117166, respectively. Numbers under protein sequences refer to amino acid position.

In addition to the three target sites of CLB19, another editing site drew our attention as the editing difference was the highest between *tfIIfα-2* and Col-0: editing of *ndhD* (117166) increased from 46.5% in wild-type to 64.7% in the mutant, even though this difference was not very significant in our RNA-seq statistical analysis (Table 1). However, Sanger sequencing clearly confirmed this increased editing level in *tfIIfα-1* and *tfIIfα-2* mutants (Figure 4B and Supplemental Figure 3A). The gene encoding the PPR protein DYW1 that edits this site (Kotera et al., 2005) was one of the two PPR genes that instead of being down-regulated was up-regulated in *tfIIfα-1* and *tfIIfα-2* (Figure 3C and Table 1). In addition to site 117166 that is processed by DYW1, *ndhD* is also edited at sites 116785, 116494, 116290 and 116281 by CRR21, OTP85, CRR28, and CRR22 PPR proteins, respectively (Okuda et al., 2007; Hammani et al., 2009; Okuda et al., 2009). Unlike *DYW1*, the expression of *CRR21, OTP85, CRR28*, and *CRR22* PPR genes was not markedly affected in *tfIIfα-2* (Supplemental Table 2). According to RNA-seq analysis, their target cytosines were also not differently edited in *tfIIfα* mutants, which was further confirmed by Sanger sequencing (Figure 4B and Supplemental Figure 3A). These results support the idea that the increased editing of *ndhD* (117166) site is a direct consequence of the increased expression of *DYW1* in *tfIIfα* mutants.

### *rpoA* editing defect in *tfIIfα* mutants does not impair PEP function

Editing of *rpoA* (78691) causes the modification of Ser67 to the conserved hydrophobic Phe67 residue, the function of which remains unknown (Figure 4C). Knowing that RpoA, together with RpoB, RpoC1 and RpoC2, is a core subunit of the plastid-encoded RNA polymerase (PEP) (Yu et al., 2014), we then asked whether the reduced *rpoA* (78691) editing in *tfIIfα-1* and *tfIIfα-2* mutants might affect PEP function and hence plastid transcription. Therefore, we specifically analyzed plastid gene expression in *tfIIfα-2* and wild-type Col-0 in our RNA-seq experiment. In a previous report, the requirement of CLB19 for efficient plastid expression was demonstrated by analysing the null mutant *clb19-1* and its complemented line clb19-1c (Chateigner-Boutin et al., 2008). *clb19-1* shows widespread deregulation of plastid gene expression (Figure 5). On the contrary, the plastid gene expression profile in *tfIIfα-2* was very similar to the wild-type in spite of the down-regulation of *CLB19* in this mutant (Figure 5 and Supplemental Table 3). Consequently, *CLB19* down-regulation in *tfIIfα* mutants does not seem to be sufficient to affect PEP function.

**Figure 5.**
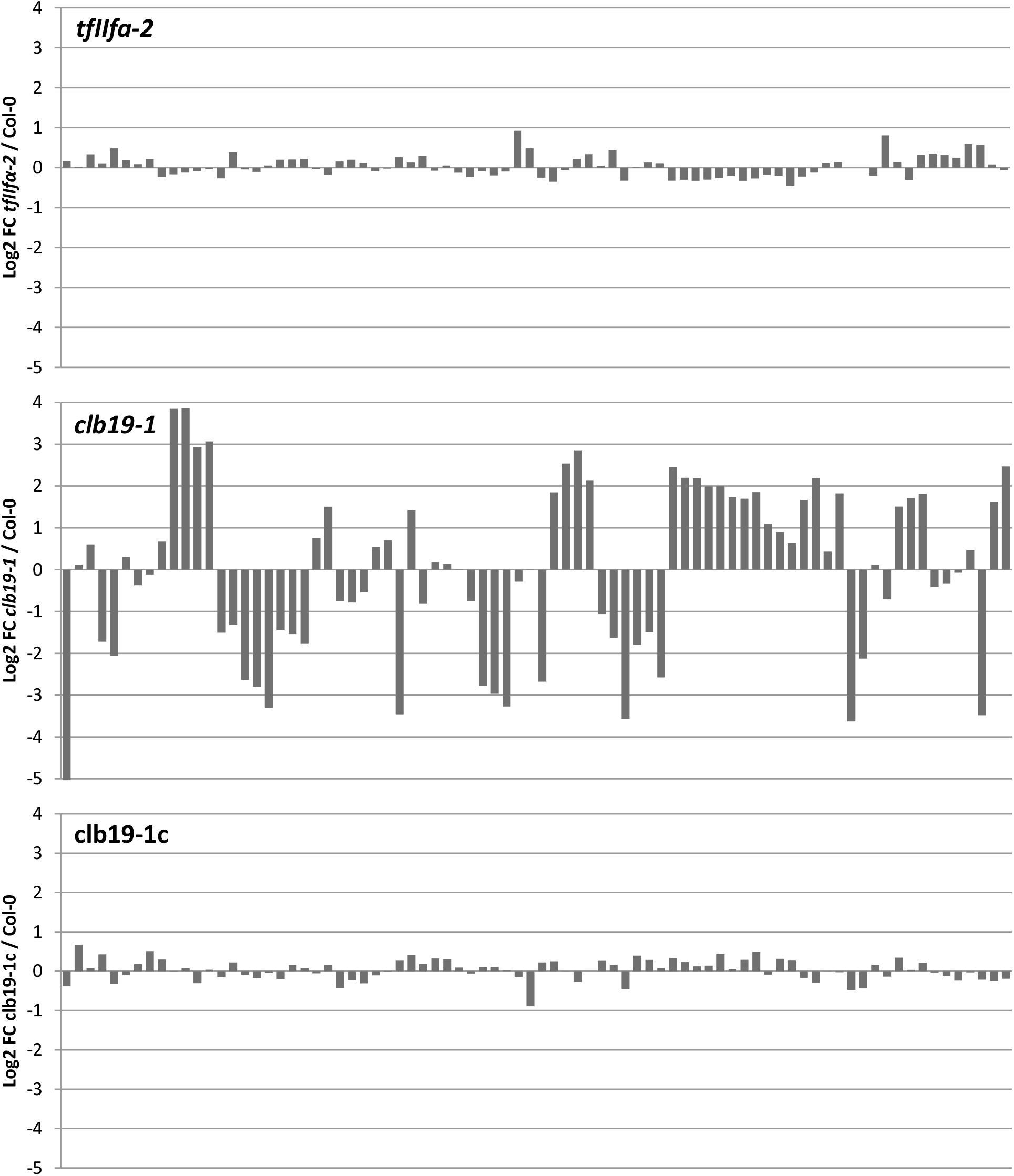
Plastid gene expression depending on *CLB19* defect. Comparison of plastid gene expression between *tfIIfα-2* (*CLB19* repressed), *clb19-1* (KO mutant) and its complemented line clb19-1c (displaying wild-type phenotype). RNA-seq data of *clb19-1* and clb19-1c were from Chateigner-Boutin *et al*. (2008). Genes are sorted from left to right according to their genomic position on the plastid chromosome (see Table S3 for details).

### *tfIIfα* mutation affects the function of the chloroplast NADH dehydrogenase-like (NDH) protein complex at multiple levels

Because editing of *ndhD* (117166) is essential for the introduction of a START codon (Figure 4C), the increased editing efficiency observed in *tfIIfα-2* could be assumed to increase the production of NdhD peptide in this mutant. NdhD is a subunit of the NDH complex involved in cyclic electron flow with PSI (Munekage et al., 2004). The NDH complex is made up of several subunits that are encoded by the nuclear and plastid genomes. Thus, higher levels of NdhD alone are unlikely to be sufficient to enhance NDH activity in *tfIIfα-2*. Therefore, we examined the expression of other NDH subunits in *tfIIfα-2*. As shown in Figure 6A, almost all nuclear and plastid genes that encode NDH subunits were up-regulated in *tfIIfα-2*. We also analyzed NDH nuclear gene expression in *tfIIfα-1* and *caa39* mutant. These NDH genes were mostly up-regulated in both *tfIIfα-1* and *tfIIfα-2* mutants whereas all genes are massively down-regulated in *caa39* (Figure 6B). These results further highlight the genetic link between TFIIFα and Topo VI and the opposite control they exert over PPR proteins and cellular processes whose regulation implicates PPR proteins. They also show that TFIIFα might regulate NDH at multiple levels: firstly, at the transcript level, where TFIIFα participates in the coordination of the expression of subunits encoded by the nuclear and plastid genomes; and secondly at the post-transcriptional/translation level by enhancing NdhD protein production via DYW1 regulation.

**Figure 6.**
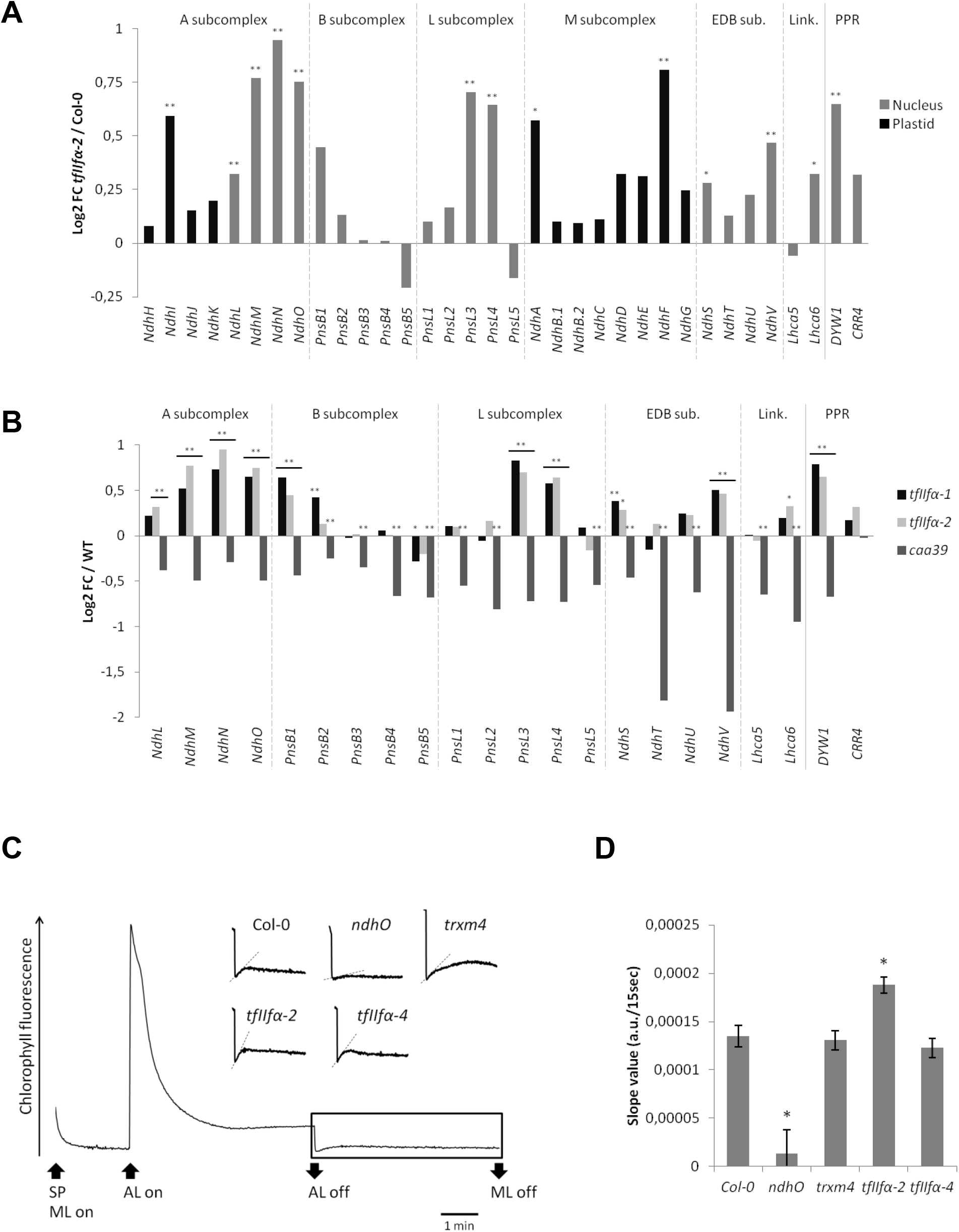
NDH expression and activity are affected by the *TFIIFα* mutation. **(A) (B)** Expression of NDH subunit and PPR genes required for *NDH* transcript editing. **(A)** Expression of nuclear- and plastid-encoded genes in *tfIIfα-2* are expressed relative to wild-type Col-0 **(B)** Expression of nuclear-encoded genes in *tfIIfα-1, tfIIfα-2* and *caa39* mutants are expressed relative their wild-type (Col-3 or Col-0). Significant expression differences between a mutant line and the corresponding wild-type are shown: * if p-value < 0.05 and ** if p-value < 0.01. EDB: electron donor-binding; Link.: linkers. **(C)** Analysis of NDH activity by measuring the chlorophyll fluorescence rise after turning off AL. The bottom curve represents a typical trace of chlorophyll fluorescence in wild-type Col-0. Insets are magnified traces from the boxed area. Slope of the curve is indicated by dash line. SP, saturating pulse; ML, measuring light; AL, actinic light. **(D)** Calculation of the slope of the curve during the first 15 s after AL off. Error bars represent standard deviation from five biological replicates. Significant differences between mutant and wild-type lines were estimated with a *t*-test: * : if p-value < 0.01.

To test whether the increase in NdhD editing and *NDH* gene up-regulation lead to increased NDH activity, we measured NDH activity in *tfIIfα-1, tfIIfα-2* and *tfIIfα-4* leaves respective to their wild-type ecotype, as well as in *ndhO* and *trxm4* mutants as known negative and positive controls for NDH activity (Courteille et al., 2013). NDH activity can be measured *in vivo* as a distinctive transient increase in chlorophyll fluorescence that occurs when actinic light (AL) exposition is suddenly stopped (Shikanai et al., 1998). Here, the fluorescence rise was more pronounced in the three *tfIIfα* mutants as well as in wild-type plants than in the negative control *ndhO* (Figure 6C and Supplemental Figure 4A & B). However, we noticed a different fluorescence induction kinetics for *tfIIfα-1* and *tfIIfα-2*: although the transient increase was not as long as in the positive control *trxm4*, the slope of the curve during the first 15 sec was steeper than in the wild-type Col-3 and Col-0, respectively (Figure 6D and Supplemental Figures 4C). In contrast, the slope value of *tfIIfα-4* and *trxm4* was similar to the wild-type and close to 0 in *ndhO*. Consequently, NDH activity seems to be affected in the *tfIIfα* mutant, even though it does not correspond to a tremendous increase of NDH activity as can be observed in the *trxm4* mutant.

## DISCUSSION

Besides its role in endoreduplication, the plant Topoisomerase VI has been implicated in transcriptional silencing (Kirik et al., 2007) and gene expression control, notably during the response of plants to stresses and phytohormones (Yin et al., 2002; Jain et al., 2008; Mittal et al., 2014; Jain et al., 2006). For instance, the constitutive expression of the rice Topo VIA or Topo VIB subunits enhanced the expression of stress-responsive genes and conferred abiotic stress tolerance to transgenic Arabidopsis plants (Jain et al. 2006). Topo VI has also been proposed to be a key regulatory factor of oxidative stress-responsive genes and eventually of the plant responses to adverse environmental conditions (Šimková et al, 2012). However, whether this control of gene expression is a direct consequence of the participation of Topo VI in the process of transcription, notably by solving topological problems associated with transcription elongation, is unclear. Here, we showed by yeast two hybrid assay and BiFC in *N. benthamiana* leaves that the Arabidopsis Topo VI complex is associated with the general transcription factor TFIIF, via the interaction of its BIN4 subunit with the alpha subunit of TFIIF. Recent experiments on the *tfIIfβ1* mutant revealed plant growth inhibition and development defects in meristematic organization responsible for stem fasciation and inflorescence impairments (Babiychuk et al., 2016). We also observed growth inhibition in *tfIIfα* mutants but no such developmental perturbations. Thanks to four different allelic mutants, we show the essential role of the first CTD-phosphatase interaction domain, whereas the second CTD-phosphatase interaction domain, which is missing in *tfIIfα-4*, seemed to be dispensable for TFIIFα function.

RNA-seq analysis performed in two different allelic mutants, *tfIIfα-1* and *tfIIfα-2*, showed that TFIIFα defects preferentially affected the expression of PPR genes, and particularly led to the repression of the PLS subfamily involved in RNA editing in mitochondria and plastids. In contrast, we observed an opposite regulation in the Topo VIA mutant *caa39* in which the PPR genes down-regulated in *tfIIfα* mutants are mainly up-regulated. Thus, Topo VI appears as a transcriptional repressor of PLS-type PPR genes in contrast to TFIIFα. A similar transcriptional repression by topoisomerases has been reported for the human Topoisomerase I (Topo I) that can interact with the general transcription factor TFIID (Merino et al., 1993). TFIID interaction with Topo I was proposed to block the transcriptional machinery at initiation step and prevent gene expression. Upon transcriptional activator, Topo I and TFIID would dissociate, release the transcription initiation complex and finally allow transcription elongation. In our context, as PPR genes are repressed in *tfIIfα-1* and *tfIIfα-2* mutants, we can suppose that TFIIFα is required for PLS-type PPR gene expression by recognizing PPR promoters and recruiting the transcriptional machinery. Topo VI would act as a transcriptional repressor by interacting with TFIIFα and physically blocking the transcription of PLS-type PPR genes. Additional experiments will be required to confirm this model and unveil whether gene expression control by the Topo VI-TFIIF interaction is directly associated with Pol II. Indeed, even in far better characterized models such as human cells, genome-wide analyses of TFIIF-binding sites have revealed that only 20% of them co-localize with Pol II, supporting a paradigm in which TFIIF may play other roles besides being an accessory protein for Pol II dependent transcription (Gelev et al., 2014).

At first glance consistent with the extensive repression of PLS-type PPR genes in *tfIIfα-1* and *tfIIfα-2*, we observed a broad, but partial, disruption of RNA editing in *tfIIfα-2*. Global editing deficiencies have been reported previously in mutant plants unable to produce the PPR-associated proteins MORF/RIP. Members of the MORF/RIP protein family are required for efficient editing of probably all targeted cytosines in both organelles. Among these, MORF8/RIP1 is the major editing factor as 75% and 20% of mitochondrial and plastid sites, respectively, are affected in *rip1* mutant with an editing defect reaching up to 81% (Bentolila et al., 2013). In *tfIIfα-1* and *tfIIfα-2* mutants, RNA-seq data showed that none of the *MORF*/*RIP* genes are down-regulated, but on the contrary some of them are induced in *tfIIfα-1* and *tfIIfα-2* such as *MORF3*/*RIP3, MORF4*/*RIP4, MORF5*/*RIP5* and *MORF6*/*RIP6*. Expression of the main factor *MORF8*/*RIP1* is not affected by the *tfIIfα* mutation, suggesting that editing defect observed in *tfIIfα-1* and *tfIIfα-2* cannot be attributed to *MORF*/*RIP* deregulation, but instead is a consequence of the control of PPR gene expression by TFIIFα.

Knocking out PLS-type PPR genes often has drastic effects on target RNA editing sites. For instance, the editing of cytosines in *rpl16* (25407), *cob* (60520) and *nad4* (167617), which are targeted by MEF35 and fully edited in wild-type plants, is completely lost in the *mef35-1* K-O mutant (Brehme et al., 2015). In contrast, the more than two fold down-regulation of *MEF35* gene in *tfIIfα-2* had no effect on the editing of these sites. The down-regulation, but not complete repression, of a single PLS-PPR gene appears to be generally not sufficient to disturb the editing efficiency in *tfIIfα-2*. However, one repressed PLS-PPR gene, *CLB19*, was an exception. CLB19 is required for editing *rpoA* at codon 67 (changing Ser to Phe), which encodes one of the core subunits of the plastid-encoded polymerase (PEP). Although we did observe a decreased efficiency of *rpoA* (78691) editing in *tfIIfα-2*, we did not detect any resulting deregulation of plastid gene expression in *tfIIfα-2*. The decreased efficiency of *rpoA* editing is probably not sufficient to impair the PEP activity. Plant chloroplasts possess a second RNA polymerase, the nucleus-encoded polymerase (NEP). NEP is mostly active in young tissues whereas the PEP activity increases with plastid maturation (Yu et al., 2014; Liere et al., 2011). However, the vast majority of plastids genes can be transcribed by either PEP or NEP, therefore we cannot exclude that NEP can compensate for the partial PEP defect in the *tfIIfα* mutant under the conditions tested.

Whereas the broad down-regulation of PLS-type PPR genes only exceptionally led to a significant reduction of target site editing in *tfIIfα-2*, the most striking editing alteration was of the increased editing efficiency that affected *ndhD. NdhD* is edited at the genomic position 117166 thanks to DYW1 interacting with the CRR4 PPR protein (Boussardon et al., 2012). RNA-seq data from *tfIIfα-1* and *tfIIfα-2* revealed the concomitant up-regulation of *DYW1* and *CRR4*, although this was not statistically significant for the latter gene. This editing process is essential for NdhD translation because it allows the start codon formation. Interestingly, almost all transcripts encoding NDH subunits accumulated in *tfIIfα-1* and *tfIIfα-2*, whereas they were massively down-regulated in *caa39*. These results revealed a multileveled control of NDH by TFIIFα: at the transcript level, TFIIFα participates to coordinate the expression of subunits encoded by the nuclear and plastid genomes; at the translation level, TFIIFα participates in NdhD protein production via *DYW1* regulation. Consequently, the chlorophyll fluorescence transient increase that is attributed to NDH activity was slightly more pronounced in *tfIIfα-1* and *tfIIfα-2* than in wild-type plants. Interestingly, in Synechocystis cells exposed to high light stress, the slope of the chlorophyll fluorescence also increases with light intensity, and it was concluded that NDH activity is induced by high light (Chen et al.,2016). Regarding the very moderate difference between *tfIIfα* and wild-type chlorophyll fluorescence patterns, as compared to in *trxm4* and *ndhO* mutants, it remains difficult, however, to firmly conclude on a clear increase of NDH activity in *tfIIfα* mutants.

The *tfIIfα-1* and *tfIIfα-2* allelic mutants represent a very rare case of PPR gene deregulation which translates into an editing defect in a target RNA. They also highlight an anterograde signalling pathway in Arabidopsis: the association between Topo VI and the general transcription factor TFIIF in nucleus controls the expression of nuclear encoded PPR proteins that are involved in cytoplasmic RNA editing for proper organelle function. However, the discovery of the molecular mechanisms that allow TFIIF to specifically regulate the expression of PLS-type PPR genes needs further investigation. Further research is also required to understand the significance of this regulation, under the opposite control of TFIIFα and Topo VI, in response to changing and adverse environmental conditions.

## METHODS

### Cloning and plasmids

Genes were amplified by PCR and cloned into the pDONR207 vector by Gateway BP reaction and subcloned into a destination vector by Gateway LR reaction (ThermoFisher Scientific). The destination vectors were pMDC99 for the complementation analysis (*TFIIFα* gene), pBIFP2 (nYFP) and pBIFP3 (cYFP) for the BiFC experiment (*TFIIFα* and *BIN4* genes, respectively), and pEarleyGate102 (CFP) for the subcellular localization (*TopoVIA* gene) (Curtis and Grossniklaus, 2003; Azimzadeh et al., 2008; Earley et al., 2006). For the transiently expression in *N. benthamiana*, the p19 plasmid was simultaneously used with the other constructs.

### Yeast Two-Hybrid Screen and Assay

The yeast two-hybrid screen was performed by Hybrigenics using the Arabidopsis RP1 library. The full-length *BIN4* cDNA (*AT5G24630*.*3/4*) was used as bait. The yeast two-hybrid assays were performed using Full-length cDNAs of RHL1/HYP7, AtSPO11-3/RHL2/BIN5 and AtTOP6B/RHL3/HYP6/BIN3 were previously cloned into pLexA (DNA-binding domain) and pB42AD (activator domain fusion) vectors of the Matchmaker LexA two-hybrid system (Clontech) (Sugimoto-Shirasu et al., 2005). The BIN4 full-length cDNA was amplified by PCR with primers 5′-TTGCGGCAATTGAGCAGCAGCTCTAGAGAGGGATC-3′ and 5′-GCTCGAGCCTTTCTTGGCTTTTGGC-3′, excised with MfeI and XhoI, and cloned into the EcoRI and XhoI sites of pLexA and pB42AD vectors. Positive interactions were detected by induction of the lacZ reporter gene in yeast EGY48 cells that are pre-transformed with p8op-lacZ reporter plasmids.

### Subcellular localization and BiFC

The cDNA of *TopoVIA* and *BIN4* were amplified without their stop codon while the gDNA of *TFIIFα* was amplified without the start codon using primers described in Supplemental Table 4. They are subsequently cloned by Gateway reactions. After transformation by electroporation and selection, C58C1 *Agrobacterium tumefaciens* was inoculated on LB medium with rifampicin and gentamycin 50 µg/mL each (and if necessary kanamycin or spectinomycin 50 µg/mL) and incubated at 28°C with 200 rpm. Cells were pelleted by centrifugation (4000 rpm, 7 min), suspended in infiltration buffer (10 mM MgCl2, 10 mM MES, pH 5.6, 200 µM acetosyringone) such as OD_600_ was 1.0, and incubated at room temperature for 4 h. Equal volumes of the *A. tumefaciens* suspensions containing interest gene and p19 plasmid were mixed and infiltrated into 5-week-old *Nicotiana benthamiana* leaves. Tobacco leaves were observed 4 d after transformation by epi-fluorescence microscopy (AxioImager Z1 Apotome, Zeiss) allowing the detection of CFP (BP 436/20 nm excitation, FT 455 nm, BP 480/40 nm emission), YFP (BP 500/25 nm excitation, FT 515 nm, BP 535/30 nm emission) and brightfield.

### Plant material, growth conditions and phenotypic characterization

Ecotypes Col-0 and Col-3 were used in this study. T-DNA insertion mutants were obtained from NASC Stock Center. The position of the T-DNA insertion was determined by sequencing PCR products obtained with a gene-specific and a T-DNA left border-specific primers. With respect to the start codon, T-DNA insertions map at +1922 bp in *tfIIfα-1* (SAIL_1171_F02), +2479 bp in *tfIIfα-2* (SALKseq_038203), +2352 bp in *tfIIfα-3* (SALKseq_123141), and +2820 bp in *tfIIfα-4* (SALKseq_095102), respectively. Plant genotypes were confirmed by PCR using primers described in Supplemental Table 4. For complemented lines, the *TFIIFα* gene with its native promoter and terminator was amplified by PCR on genomic Arabidopsis Col-0 DNA and cloned by Gateway reactions. The clone was introduced into *tfIIfα-1, tfIIfα-2, tfIIfα-3* and *tfIIfα-4* homozygous mutant plants. The *ndhO* and *trxm4* mutants were provided by Dominique Rumeau (Courteille et al., 2013).

*Arabidopsis thaliana* plants were grown 6 weeks on soil in a phytotron under 16 h light / 8 h dark photoperiod (80-90 µmol photons m^-2^ s^-1^) and controlled conditions of temperature (22/20°C, day/night) and relative air humidity (55/75%, day/night). For *in vitro* culture, seeds were sterilized with bleach, vernalized at 4°C for 3 days and grown on Murashige and Skoog 1/2 media containing 1% sucrose. Plates were placed in growth chambers under 16 h light / 8 h dark photoperiod (80 µmol photons m^-2^ s^-1^) at 22/20°C (day/night). Seedlings were collected after 6 d growth.

### Chlorophyll Fluorescence Analysis

NDH activity was detected by chlorophyll fluorescence measurements with a DUAL-PAM-100 (DUAL-PAM/F, Walz). A mature leaf was dark-acclimated for 20 min and the transient increase in chlorophyll fluorescence was monitored as previously described (Shikanai et al., 1998). Leaves were exposed to Actinic Light (AL) (250 µmol photons m^-2^ s^-1^) for 5 min. AL was turned off and the subsequent transient rise in fluorescence ascribed to NDH activity was monitored by chlorophyll fluorimetry.

### RNA extraction

RNA extraction was performed from one hundred 6-day-old seedlings for each biological replicate. For expression and editing analysis, the total RNA was extracted using TRI Reagent (MRC) and treated with DNase I (1 U/µL; Thermo Scientific) for 30 min at 37°C according to the manufacturer’ instructions. For RNA-sequencing, total RNA was extracted according to a published protocol (Box et al., 2011). Extracted RNA was purified with RNeasy Plant Mini kit (Qiagen) and treated with DNase I as described above.

### RT-qPCR and Editing Analysis

cDNA was synthesized from 500 ng of total RNA using PrimeScript RT Reagent kit (Perfect Real Time; Takara) with oligodT and random hexamers for quantitative RT-PCR (RT-qPCR) experiments, and with random hexamers only for RNA editing analysis.

RT-qPCR was performed using SYBR Premix Ex Taq II (Tli RNase H Plus, Takara) using a CFX96 Real-Time System (CFX Manager; BioRad). Each reaction was prepared using 0.5 µL of cDNA (25 ng/µL), 7.5 µL of SYBR Green Master mix, and 5 µM forward and reverse primer each, in a total volume of 15 µL. The amplification profile consisted of 95°C for 30 sec and 45 cycles (95°C for 10 s, 60°C for 30 s, and 72°C for 30 s). *PP2A* and *PRF1* were taken as housekeeping genes to normalize the expression of gene of interest.

For editing analysis, cDNA was amplified by PCR before purification using NucleoSpin Gel and PCR Clean-up (Macherey-Nagel). Purified cDNAs were sequenced by GATC Biotech (Sanger sequencing SUPREMERUN) using specific primers. Chromatograms were analyzed with DNA Baser software. Primers used for in RT-qPCR, PCR amplification, and DNA sequencing were listed in Supplemental Table 4.

### RNA-seq library preparations and sequencing

Three independent biological replicates were produced for each line. For each biological repetition, RNA samples were obtained by pooling RNA from more than 100 plants. Aerial parts were collected from plants at 1.00 developmental growth stages (Boyes et al., 2001), cultivated as described above. Total RNA was extracted using RNeasy kit (Qiagen®, Hilden, Germany) according to the supplier’s instructions.

For *tfIIfα-1, caa39* (and the respective Col-3 and Col-0 wild-type) gene expression analysis, RNA-seq experiment was carried out at plateform POPS, transcriptOmic Plateform of the Institute of Plant Sciences -Paris-Saclay, using a IG-CNS Illumina Hiseq2000 to perform paired-end 100bp sequencing, on RNA-seq libraries constructed with the TruSeq_Stranded_mRNA_SamplePrep_Guide_15031047_D protocol (Illumina®, California, U.S.A.). The RNA-seq samples were sequenced in paired-end (PE) with a sizing of 260 bp and a read length of 100 bases. Six samples by lane of Hiseq2000 using individual bar-coded adapters and giving approximately 30 million of PE reads by sample were generated.

For *tfIIfα-2* and Col-0 wild-type gene expression and editome analyses, RNA-seq libraries were generated using TruSeq® Stranded Total RNA (with RiboZero plant) #RS-122-2401 (composed by ref 15032611 / batch 20167353; ref 15032612 / batch 20172414; ref 15032615 / batch 20172978; ref 15035748 / batch 20142725) according to the supplier’s instructions RS-122-9007DOC (Illumina®, California, U.S.A.). Using a NextSeq® 500/550 High Output kit v2 (75 cycles) #FC-404-2005 (composed by ref 15057934 / batch 20157769; ref 15058251 / batch 20166120; ref 15057941 / batch 20158908; ref 1506573 / batch 20169394) and according to the supplier’s instructions 15048776 v02 (Illumina®, California, U.S.A.), the RNA-seq samples were sequenced in single-end (SE) with a sizing of 260 bp and a read length of 75 bases. 8 samples by lane of NextSeq500 using individual bar-coded adapters and giving approximately 40 million of SE reads by sample were generated.

### RNA-seq bioinformatic treatment and analysis

To facilitate comparisons, each RNA-Seq sample followed the same pipeline from trimming to transcript abundance quantification as follows. Read preprocessing criteria included trimming of library adapters and performing quality control checks using FastQC (Version 0.11.5) (Andrews, 2010). The raw data (fastq) were trimmed for Phred Quality Score > 20, read length > 30 bases and sort by Trimming Modified homemade fastx_toolkit-0.0.13.2 software for rRNA sequences. Bowtie2 (version 2.2.9) (Langmead and Salzberg, 2012) was used to align reads against the *Arabidopsis thaliana* transcriptome (TAIR10_cdna_20110103_representative_gene_model_updated) (with --local option). Reads were counted using a command line modified from Pieterse MJ and al. (Pieterse et al., 2013).

Differential expression was performed with SARTools (version 1.5.1) (Varet et al., 2016) using edgeR with default settings except cpmCutoff which was disabled.

For editing analysis, reads were aligned with STAR (version 2.5.3a) (Dobin et al., 2013) against the genome (Araport11_GFF3_genes_transposons.201606) with options --genomeFastaFiles Arabidopsis_thaliana.TAIR10.31.dna.genome.modified.fa, --runMode genomeGenerate, --sjdbOverhang 75. Bam files were sorted by coordinates and indexed. Reads were counted with Htseq-counts (version 0.9.1) and differential expression was performed with edgeR (version 3.12.1). Editing analysis was made as in (Malbert et al., 2018).

### DNA-seq analysis of *tfIIfα-1*

Genomic DNA from a pool of *tfIIfα-1* plants was extracted and purified using the NucleoSpin Plant II Maxi kit (Machery-Nagel, Düren, Germany) with PL1 buffer and polyvinylpolypyrrolidone (PVPP, at half the plant tissue weight). DNA-seq library preparation and sequencing were performed at the Earlham Institute (Norwich, UK). DNA libraries were sequenced with 150 bp paired-end run metrics on an Illumina HiSeq4000 Sequencing System. After checking quality with FASTQC (0.11.5), sequences were first aligned against pCSA110 sequence with bowtie2 (2.2.9). Aligned reads were extracted to a new bam file which was converted to fastq and fasta format for further use. The fastq format was used to align those reads against *Arabidopsis thaliana* genome (TAIR 10). Alignment was visualized in IGV and genes with flanking plasmid borders were identified. Finally, the reads with both Arabidopsis and plasmid sequences were extracted and BLASTn was used to identify the border (right or left) of the plasmid and the position of the insertion in the genome.

### Accession Numbers

Sequence data from this article can be found in the Arabidopsis Genome Initiative or GenBank/EMBL databases under the following accession numbers: *At4g12610* (*TFIIFα*), *At5g24630* (*BIN4*), *At5g02820* (*AtSPO11-3*), *At3g20780* (*AtTOP6B*), *At1g48380* (*RHL1*), *At1g03510, At2g36980, At5g47460, At1g47580* (*DYW1*), *At2g35130, At1g05750* (*CLB19*), *AtCg00740* (*rpoA*), *AtCg00670* (*pclpP*), *AtCg00360* (*ycf3*), *AtCg01050* (*ndhD*), *At5g55740* (*CRR21*), *At2g02980* (*OTP85*), *At1g59720* (*CRR28*), *At1g11290* (*CRR22*). RNA-seq datasets are available at https://www.ncbi.nlm.nih.gov/geo/query/acc.cgi?acc=GSE103924 (reviewer token: slchuqakvvwfpsj)

## Supplemental Data

**Supplemental Figure 1**. Characterization of *TFIIFα* lines compared to wild-type (wt) plants.

**Supplemental Figure 2**. Chromosomal rearrangement in *tfIIfα-1* mutant.

**Supplemental Figure 3**. Effects of *tfIIfα* mutations on editing efficiency.

**Supplemental Figure 4**. NDH activity is affected by the *TFIIFα* mutation.

**Supplemental Table 1**. Yeast two-hybrid screen using the full-length Arabidpsis BIN4 as a bait.

**Supplemental Table 2**. PPR gene expression.

**Supplemental Table 3**. Plastid gene expression from RNA-seq data.

**Supplemental Table 4**. Primer list.

**Supplemental Dataset 1**. Editing level of organellar RNAs in *tfIIfα-2* compared to wild-type Col-0.

## ACKNOWLEDGEMENTS

This work was supported by the French National Research Agency (ANR 2010-JCJC-1205-01 and ANR-14-CE02-0010 to CL). LD was supported by CEA and Région PACA. High-throughput RNA-sequencing was performed at the POPS plateform, supported by the LabEx Saclay Plant Sciences-SPS (ANR-10-LABX-0040-SPS). We are deeply grateful to Mr. Michel Terese for his priceless help with bioinformatics analyses. We also want to express our gratitude to the students who contributed to this work, especially Justine Quillet. Dominique Rumeau and Stefano Caffarri at BIAM are thanked for technical help and fruitful discussion on chlorophyll fluorescence analysis. We thank Ben Field for critical reading of the manuscript.

## AUTHOR CONTRIBUTIONS

L.D., K.S. and C.L. designed the research. L.D., C.V., D.A. and C.B. performed research. L.S.-T. and C.L. performed and analyzed RNA-seq. E.D. contributed new computational tools. L.D., C.L. and C.L. analyzed data. L.D. and C.L. wrote the paper with input from all coauthors.

