## Supplemental Figures & Tables for "TFIIFα interacts with the Topoisomerase VI complex and selectively controls the expression of genes encoding PPR proteins involved in organellar RNA editing in Arabidopsis"

**A**

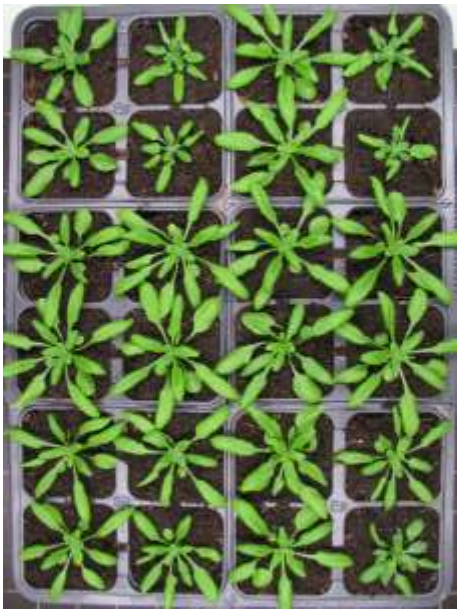

| | | | |
|---|---|---|---|
| wt Col-3 | *tfIIfα-1* | wt Col-0 | *tfIIfα-2* |
| wt Col-3 | *tfIIfα-1* | wt Col-0 | *tfIIfα-2* |
| | | wt Col-0 | *tfIIfα-4* |
| | | wt Col-0 | *tfIIfα-4* |
| | | wt Col-0 | *tfIIfα-3* |
| | | wt Col-0 | *tfIIfα-3* |

**B**

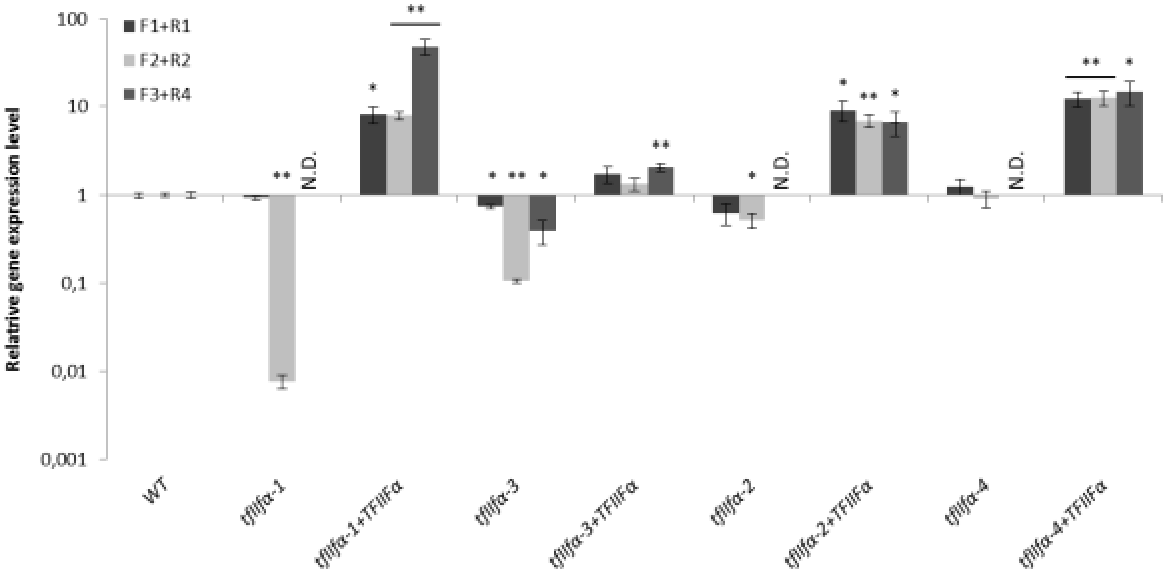


**Figure S1.** Characterization of *TFIIFα* lines compared to wild-type (wt) plants.

**(A)** 4-week-old plants were cultivated in soil under continuous light.

**(B)** RT-qPCR-based *TFIIFα* transcript level analyses in *tfIIfα* mutants and their respective complemented lines relative to corresponding wild-type plants (Col-3 or Col-0). Error bars represent standard deviation from biological triplicates. Significant expression differences between *tfIIfα* and wild-type lines were estimated with a *t*-test: ** if p-value < 0.01 and * if p-value < 0.05. N.D.: not detected.

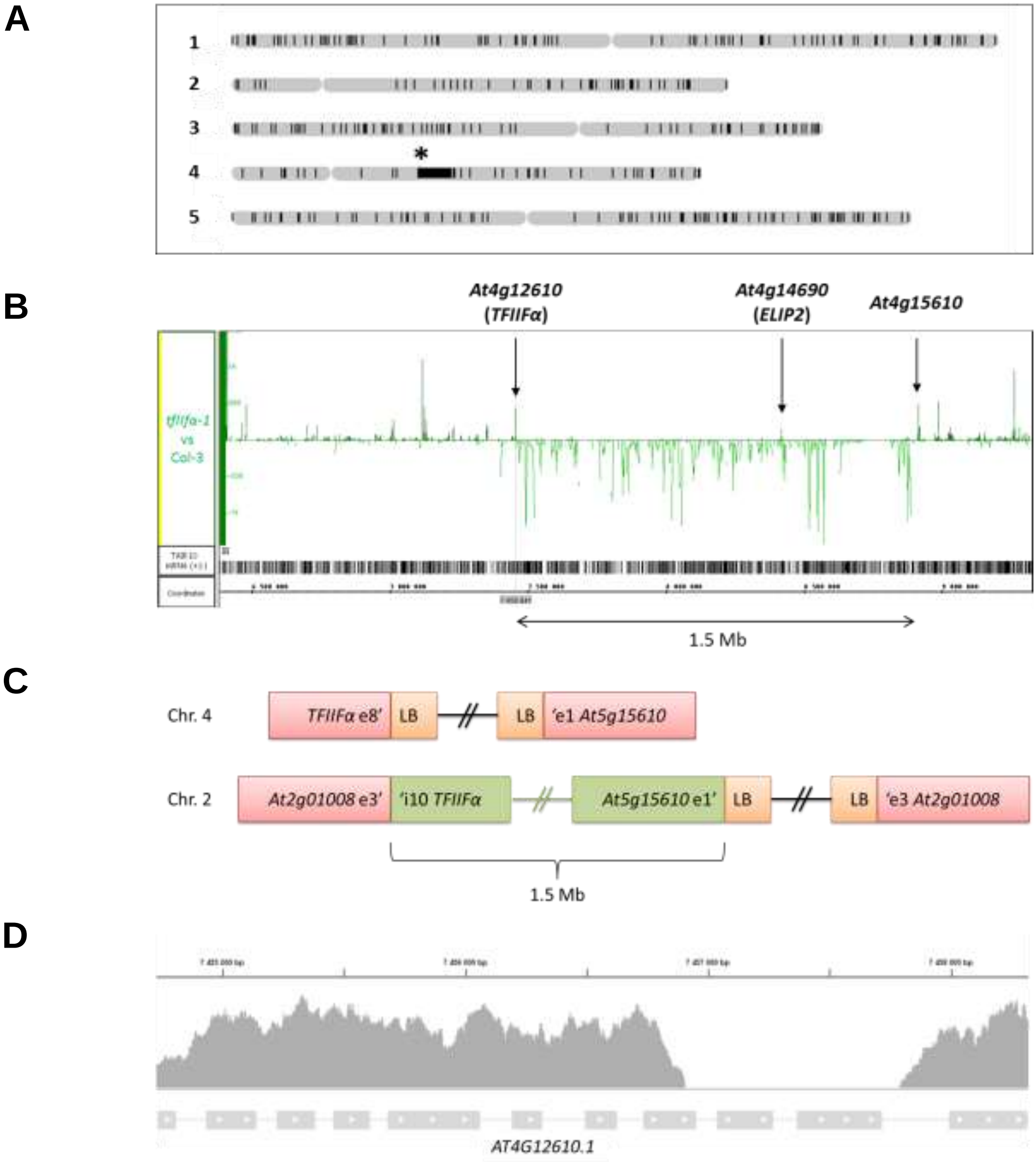


**Figure S2.** Chromosomal rearrangement in *tfIIfα-1* mutant.

**(A)** The localisation of down-regulated genes in *tfIIfα-1* mutant is schematically represented along the five chromosomes (black lines). The *TFIIFα* locus in chromosome 4 is indicated as an asterisc.

**(B)** Differential expression profile (normalized differential read density from RNA-seq) of *tfIIfα-1* mutant compare to wild-type (Col-3) around the T-DNA insertion. Positions of *TFIIFα*, *ELIP2* and *At4g15610* genes are indicated by arrows.

**(C)** Chromosomal rearrangement in *tfIIfα-1* as supported by DNA-seq. The 1.5 Mb region spanning from the T-DNA insertion site in *TFIIFα* exon 8 to the first exon of *At4g15610*, appeared to be translocated to the top of chromosome 2, in *At2g01008*.

**(D)** DNA-seq read density at the *TFIIFα* locus in *tfIIfα-1***.**

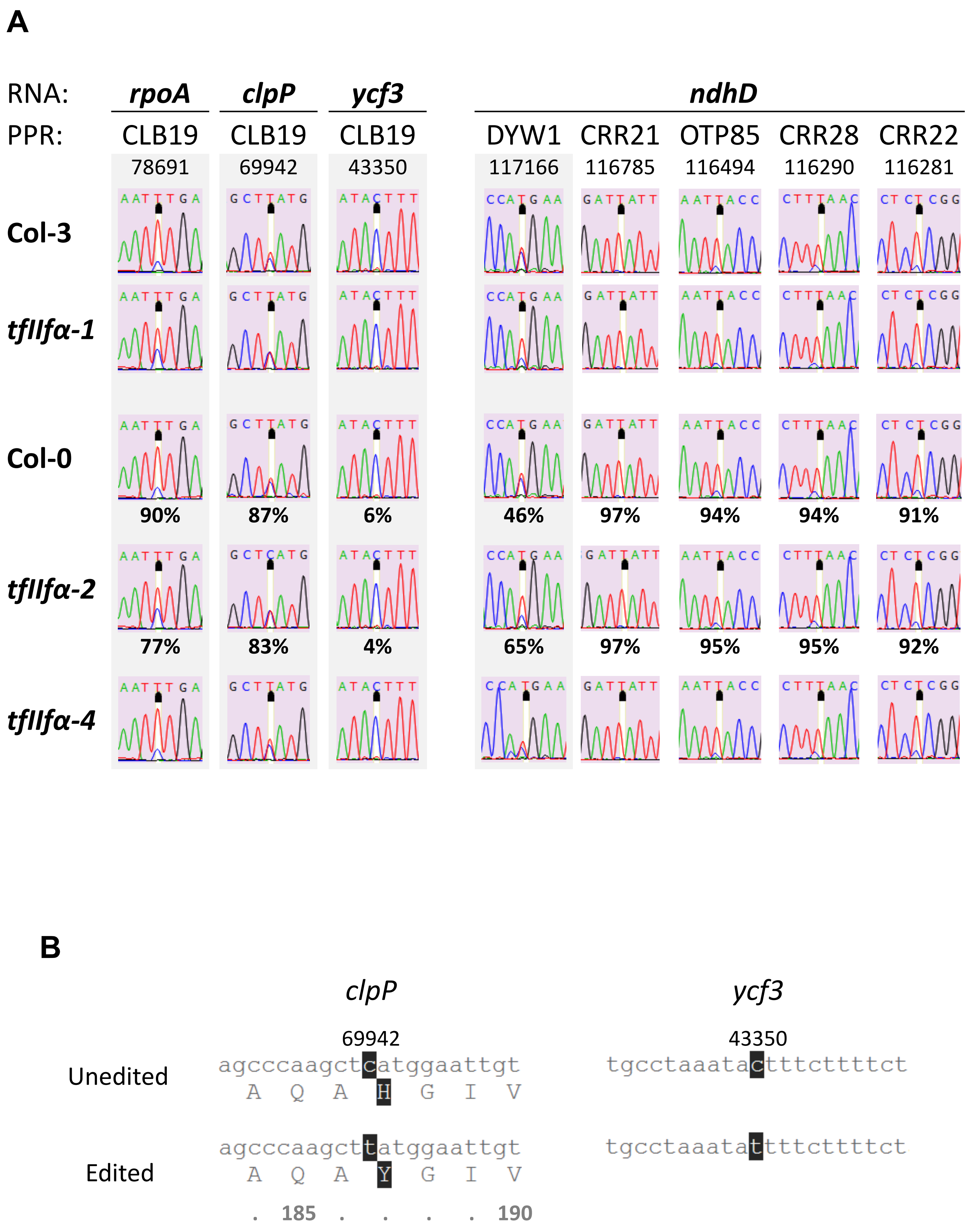
A
RNA:
rpoA
clpP
ycf3
ndhD
PPR:
CLB19
CLB19
CLB19
DYW1
CRR21
OTP85
CRR28
CRR22
78691
69942
43350
117166
116785
116494
116290
116281
Col-3
tfIIfα-1
Col-0
90%
87%
6%
46%
97%
94%
94%
91%
tfIIfα-2
77%
83%
4%
65%
97%
95%
95%
92%
tfIIfα-4
B
clpP
ycf3
69942
43350
Unedited
agcccaagctcatggaattgt
A Q A H G I V
tgcctaaatactttcttttct
Edited
agcccaagcttatggaattgt
A Q A Y G I V
tgcctaaatattttcttttct
. 185 . . . . 190

**Figure S3.** Effects of *tfIIfα* mutations on editing efficiency

**(A)** *rpoA*, *clpP*, *ycf3* and *ndhD* editing levels measured by Sanger sequencing in wild-type (Col-3, Col-0), *tfIIfα-1*, *tfIIfα-2,* and *tfIIfα-4* mutants. Chromatograms of *rpoA* (78691), *clpP* (69942), and *ycf3* (43350) targeted by CLB19, and *ndhD* (117166) processed by DYW1. For *ndhD*, editing at the genomic position 117166 is compared with those of four other loci not edited by DYW1. Under each chromatogram of wild-type Col-0 and *tfIIfα-2* is indicated the editing percentage detected in RNA-seq.
**(B)** Comparison of nucleic acid and protein sequences of *clpP* and *ycf3* depending on editing process or not at genomic positions 69942 and 43350, respectively. Numbers under protein sequences refer to amino acid position.

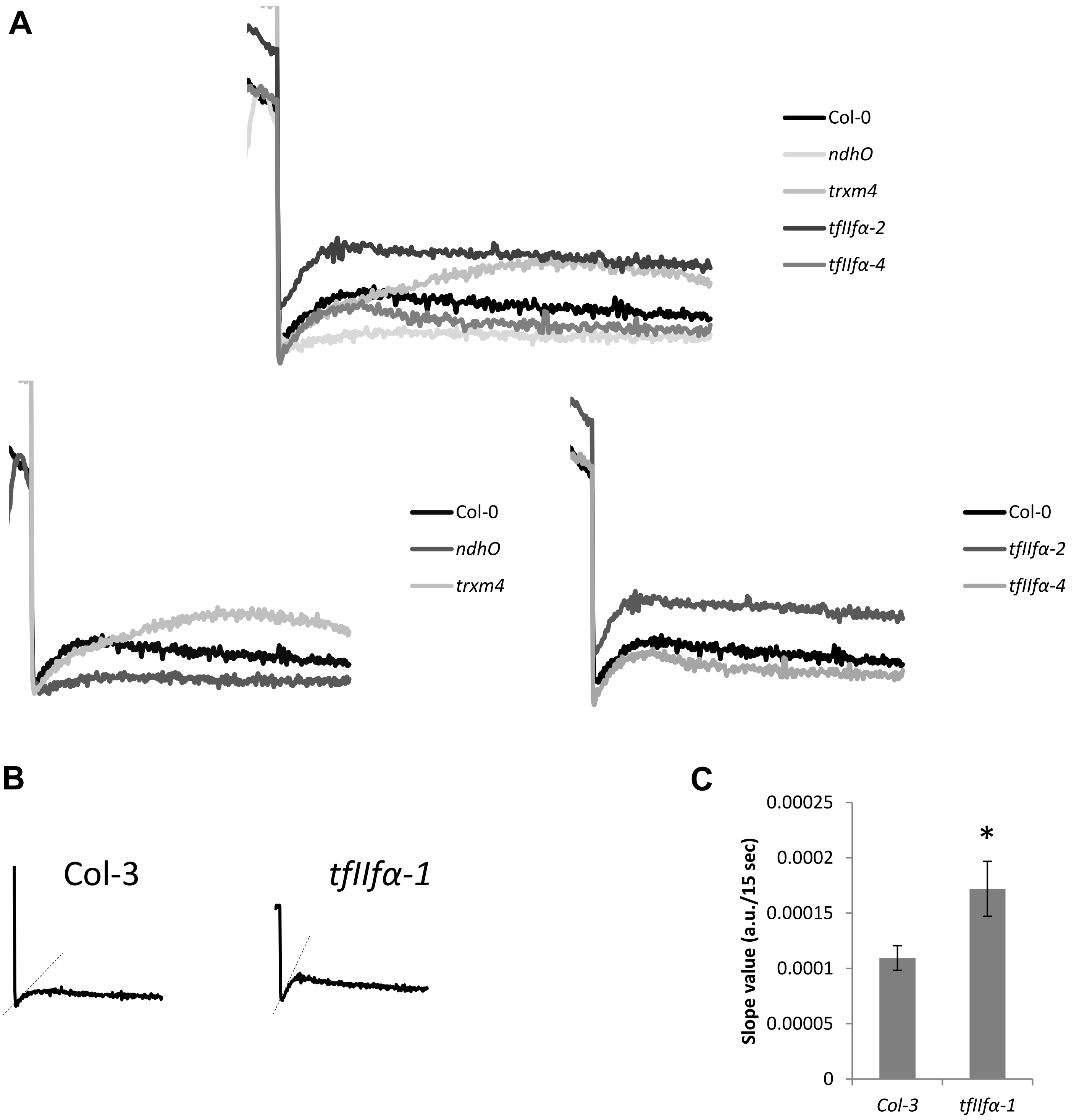


**Figure S4.** NDH activity is affected by the *TFIIFα* mutation.

**(A) (B)** Analysis of NDH activity by measuring chlorophyll fluorescence rise after turning off AL. **(A)** *tfIIfα-2* and *tfIIfα-4* traces are confronted with those of wild-type (Col-0), negative (*ndhO*) and positive (*trxm4*) controls. **(B)** *tfIIfα-1* chlorophyll fluorescence to wild-type Col-3. Slope of the curve is indicated by dash line.
**(C)** Calculation of the slope of the curve during the first 15 sec after AL off. Error bars represent standard deviation from five biological replicates. Significant difference between *tfIIfα-1* and wild-type Col-3 was estimated with a t-test: * if p-value < 0.01.

**Table S1. Yeast two-hybrid screen using the full-length Arabidopsis BIN4 as a bait**

Nature: cDNA
Reference Bait Fragment: BIN4 (2-451) ; hgx1216v1
Prey Library: Arabidopsis thaliana RP1
Vector(s): pB27 (LexA, C-terminal fusion)

Tested Diploids: 47.22 millions (pB27)
**3AT Concentration: 2.0 mM (pB27)**

| Clone Name | Gene ID | Gene Names | Start | Stop | Frame | Orientation | % Id 5p/3p |
|---|---|---|---|---|---|---|---|
| hgx1216v1_pB27A-20 | AT1G48380 | HYP7, HYPOCOTYL 7, RHL1, ROOT HAIRLESS 1 | 3 | No Data | IF | Sense | 98.9 / 97.5 |
| hgx1216v1_pB27A-3 | | | 3 | No Data | IF | Sense | 99.4 / 96.4 |
| hgx1216v1_pB27A-30 | | | 3 | No Data | IF | Sense | 96.3 |
| hgx1216v1_pB27A-25 | | | 3 | No Data | IF | Sense | 98.8 / 91.7 |
| hgx1216v1_pB27A-31 | | | 48 | 691 | IF | Sense | 99.7 / 99.4 |
| hgx1216v1_pB27A-5 | | | 157 | No Data | OOF1 | Sense | 96.3 |
| hgx1216v1_pB27A-12 | | | 183 | 1309 | IF | Sense | 99.1 / 64.4 |
| hgx1216v1_pB27A-1 | | | 183 | No Data | IF | Sense | 95.6 |
| hgx1216v1_pB27A-14 | AT4G12610 | TFIIFα / RAP74 | -85 | 734 | IF | Sense | 99.7 / 97.8 |
| hgx1216v1_pB27A-21 | | | -82 | 994 | IF | Sense | 99.0 / 97.9 |
| hgx1216v1_pB27A-24 | | | -73 | 685 | IF | Sense | 99.0 / 99.1 |
| hgx1216v1_pB27A-19 | | | -67 | 685 | IF | Sense | 99.0 / 99.3 |
| hgx1216v1_pB27A-18 | | | -67 | 685 | IF | Sense | 98.6 / 96.8 |
| hgx1216v1_pB27A-15 | | | -52 | 685 | IF | Sense | 99.4 / 99.0 |
| hgx1216v1_pB27A-27 | | | -22 | 652 | IF | Sense | 99.8 / 98.6 |
| hgx1216v1_pB27A-7 | | | -22 | 652 | IF | Sense | 99.7 / 99.5 |

Tested Diploids: 47.22 millions (pB27) + 44.33 millions (pB27)
**3AT Concentration: 2.0 mM (pB27) + 0.5 mM (pB27)**

| Clone Name | Gene ID | Other Gene Names | Start | Stop | Frame | Orientation | % Id 5p/3p |
|---|---|---|---|---|---|---|---|
| hgx1216v1_pB27B-6 | AT1G48380 | HYP7, HYPOCOTYL 7, RHL1, ROOT HAIRLESS 1 | 2 | No Data | OOF2 | Sense | 91.6 |
| hgx1216v1_pB27A-20 | | | 3 | No Data | IF | Sense | 98.9 / 97.5 |
| hgx1216v1_pB27A-30 | | | 3 | No Data | IF | Sense | 96.3 |
| hgx1216v1_pB27A-3 | | | 3 | No Data | IF | Sense | 99.4 / 96.4 |
| hgx1216v1_pB27B-38 | | | 3 | No Data | IF | Sense | 98.2 |
| hgx1216v1_pB27A-25 | | | 3 | No Data | IF | Sense | 98.8 / 91.7 |
| hgx1216v1_pB27A-31 | | | 48 | 691 | IF | Sense | 99.7 / 99.4 |
| hgx1216v1_pB27B-44 | | | 48 | 691 | IF | Sense | 99.8 / 99.7 |
| hgx1216v1_pB27B-14 | | | 57 | No Data | IF | Sense | 99.6 |
| hgx1216v1_pB27B-29 | | | 57 | No Data | IF | Sense | 93.1 |
| hgx1216v1_pB27A-5 | | | 157 | No Data | OOF1 | Sense | 96.3 |
| hgx1216v1_pB27B-17 | | | 182 | No Data | OOF2 | Sense | 97.1 |
| hgx1216v1_pB27B-47 | | | 183 | 1309 | IF | Sense | 94.1 / 61.6 |
| hgx1216v1_pB27B-12 | | | 183 | 1309 | IF | Sense | 99.5 / 58.1 |
| hgx1216v1_pB27A-12 | | | 183 | 1309 | IF | Sense | 99.1 / 64.4 |
| hgx1216v1_pB27B-24 | | | 183 | 1309 | IF | Sense | 98.7 / 63.8 |
| hgx1216v1_pB27B-13 | | | 183 | 1309 | IF | Sense | 98.8 / 63.8 |
| hgx1216v1_pB27B-31 | | | 183 | 1309 | IF | Sense | 99.3 / 65.0 |
| hgx1216v1_pB27A-1 | | | 183 | No Data | IF | Sense | 95.6 |
| hgx1216v1_pB27A-14 | AT4G12610 | TFIIFα / RAP74 | -85 | 734 | IF | Sense | 99.7 / 97.8 |
| hgx1216v1_pB27B-39 | | | -85 | 734 | IF | Sense | 99.7 / 98.7 |
| hgx1216v1_pB27B-41 | | | -82 | 700 | IF | Sense | 99.7 / 88.7 |
| hgx1216v1_pB27B-16 | | | -82 | 734 | IF | Sense | 96.4 / 93.6 |
| hgx1216v1_pB27B-50 | | | -82 | 700 | IF | Sense | 98.7 / 99.4 |
| hgx1216v1_pB27A-21 | | | -82 | 994 | IF | Sense | 99.0 / 97.9 |
| hgx1216v1_pB27B-11 | | | -82 | 700 | IF | Sense | 99.6 / 98.6 |
| hgx1216v1_pB27B-1 | | | -73 | 520 | IF | Sense | 99.0 / 98.8 |
| hgx1216v1_pB27B-30 | | | -73 | 520 | IF | Sense | 98.1 / 99.2 |
| hgx1216v1_pB27B-37 | | | -73 | 847 | IF | Sense | 99.6 / 98.6 |
| hgx1216v1_pB27B-48 | | | -73 | 847 | IF | Sense | 96.5 / 98.2 |
| hgx1216v1_pB27A-24 | | | -73 | 685 | IF | Sense | 99.0 / 99.1 |
| hgx1216v1_pB27B-46 | | | -72 | No Data | OOF1 | Sense | 97.5 |
| hgx1216v1_pB27B-19 | | | -71 | No Data | OOF2 | Sense | 88.3 |
| hgx1216v1_pB27B-43 | | | -70 | 746 | IF | Sense | 97.0 / 99.2 |
| hgx1216v1_pB27A-19 | | | -67 | 685 | IF | Sense | 99.0 / 99.3 |
| hgx1216v1_pB27A-18 | | | -67 | 685 | IF | Sense | 98.6 / 96.8 |
| hgx1216v1_pB27B-35 | | | -53 | 544 | OOF2 | Sense | 99.8 / 99.4 |
| hgx1216v1_pB27B-3 | | | -53 | 544 | OOF2 | Sense | 99.8 / 100.0 |
| hgx1216v1_pB27B-28 | | | -53 | 544 | OOF2 | Sense | 100.0 / 99.8 |
| hgx1216v1_pB27B-22 | | | -52 | 552 | IF | Sense | 100.0 / 100.0 |
| hgx1216v1_pB27B-26 | | | -52 | 680 | IF | Sense | 99.3 / 99.3 |
| hgx1216v1_pB27A-15 | | | -52 | 685 | IF | Sense | 99.4 / 99.0 |
| hgx1216v1_pB27B-10 | | | -52 | 680 | IF | Sense | 99.4 / 99.4 |
| hgx1216v1_pB27B-5 | | | -23 | 652 | OOF2 | Sense | 97.9 / 83.7 |
| hgx1216v1_pB27B-23 | | | -22 | 652 | IF | Sense | 99.5 / 99.1 |
| hgx1216v1_pB27A-27 | | | -22 | 652 | IF | Sense | 99.8 / 98.6 |
| hgx1216v1_pB27B-18 | | | -22 | 652 | IF | Sense | 99.5 / 92.0 |
| hgx1216v1_pB27A-7 | | | -22 | 652 | IF | Sense | 99.7 / 99.5 |
| hgx1216v1_pB27B-2 | | | -16 | 659 | IF | Sense | 99.8 / 99.5 |
| hgx1216v1_pB27B-51 | | | -16 | 659 | IF | Sense | 88.3 / 87.1 |
| hgx1216v1_pB27B-49 | | | No Data | 728 | ?? | Sense | 99.2 |
| hgx1216v1_pB27B-32 | | | No Data | 846 | ?? | Sense | 99.4 |

**Table S2. Expression of genes deregulated in *tfIIfα-1* .**

| Id | | | | WTCol-0_rep1 | WTCol-0_rep2 | WTCol-0_rep3 | tfIIfα-1_rep1 | tfIIfα-1_rep2 | tfIIfα-1_rep3 | norm.WTCol-( | norm.WTCol-( | norm.WTCol-( | norm.tfIIfα-1_ | norm.tfIIfα-1_ | norm.tfIIfα-1_ | baseMean | WTCol-0 | tfIIfα-1 | FC | log2FoldChan | pvalue |
|---|---|---|---|---|---|---|---|---|---|---|---|---|---|---|---|---|---|---|---|---|---|
| AT2G26020.1 | PDF1.2B | protein_coding | plant defensin 1.2b | 4 | 0 | 0 | 72 | 7 | 4 | 8 | 0 | 0 | 156 | 19 | 11 | 48.22 | 3 | 62 | 42.858 | 5.421 | 7.29E-09 |
| AT1G05680.1 | UGT74E2 | protein_coding | Uridine diphosphate glyc | 17 | 35 | 1 | 261 | 51 | 51 | 36 | 86 | 3 | 566 | 135 | 139 | 232.77 | 42 | 280 | 37.883 | 5.243 | 1.63E-14 |
| AT2G07815.1 | | protein_coding | | 860 | 830 | 848 | 1065 | 971 | 961 | 1808 | 2039 | 2171 | 2310 | 2578 | 2613 | 1127.54 | 2006 | 2500 | 30.646 | 4.938 | 8.80E-07 |
| AT2G04050.1 | DTXL1 | protein_coding | MATE efflux family prote | 21 | 64 | 0 | 189 | 125 | 159 | 44 | 157 | 0 | 410 | 332 | 432 | 254.62 | 67 | 391 | 26.198 | 4.711 | 5.29E-10 |
| AT5G61160.1 | ACT | protein_coding | anthocyanin 5-aromatic a | 2 | 1 | 1 | 76 | 8 | 3 | 4 | 2 | 3 | 165 | 21 | 8 | 36.82 | 3 | 65 | 21.541 | 4.429 | 1.04E-08 |
| AT2G26010.1 | PDF1.3 | protein_coding | plant defensin 1.3 | 4 | 0 | 0 | 64 | 3 | 3 | 8 | 0 | 0 | 139 | 8 | 8 | 43.86 | 3 | 52 | 19.721 | 4.302 | 1.47E-06 |
| AT3G20760.1 | NSE4B | protein_coding | Nse4, component of Smc! | 371 | 230 | 281 | 568 | 441 | 342 | 780 | 565 | 719 | 1232 | 1171 | 930 | 460.93 | 688 | 1111 | 16.957 | 4.084 | 1.45E-09 |
| AT1G18830.1 | SEC31A | protein_coding | Transducin/WD40 repeat | 13 | 19 | 3 | 113 | 200 | 73 | 27 | 47 | 8 | 245 | 531 | 199 | 126.76 | 27 | 325 | 15.279 | 3.933 | 6.58E-20 |
| AT2G18193.1 | | protein_coding | P-loop containing nucleo: | 26 | 67 | 10 | 312 | 155 | 157 | 55 | 165 | 26 | 677 | 411 | 427 | 246.65 | 82 | 505 | 14.748 | 3.882 | 4.70E-17 |
| AT5G44420.1 | PDF1.2A | protein_coding | plant defensin 1.2 | 4 | 0 | 0 | 37 | 4 | 5 | 8 | 0 | 0 | 80 | 11 | 14 | 34.48 | 3 | 35 | 13.68 | 3.774 | 4.91E-08 |
| AT1G03445.1 | BSU1 | protein_coding | Serine/threonine protein | 4 | 3 | 4 | 31 | 16 | 14 | 8 | 7 | 10 | 67 | 42 | 38 | 41.04 | 8 | 49 | 11.876 | 3.57 | 2.34E-27 |
| AT2G07734.1 | | protein_coding | Alpha-L RNA-binding mot | 4531 | 3173 | 3842 | 6049 | 4719 | 4478 | 9524 | 7795 | 9836 | 13121 | 12528 | 12178 | 5415.51 | 9052 | 12609 | 11.452 | 3.518 | 0.00324537 |
| AT1G54020.2 | | protein_coding | GDSL-like Lipase/Acylhyd | 12 | 4 | 2 | 85 | 30 | 17 | 25 | 10 | 5 | 184 | 80 | 46 | 51.68 | 13 | 103 | 11.119 | 3.475 | 1.50E-13 |
| AT1G17960.1 | | protein_coding | Threonyl-tRNA synthetas | 23 | 25 | 18 | 236 | 118 | 131 | 48 | 61 | 46 | 512 | 313 | 356 | 210.2 | 52 | 394 | 11.109 | 3.474 | 5.64E-32 |
| AT3G59190.1 | | protein_coding | F-box/RNI-like superfamil | 15 | 13 | 14 | 42 | 43 | 35 | 32 | 32 | 36 | 91 | 114 | 95 | 59.99 | 33 | 100 | 10.798 | 3.433 | 2.13E-26 |
| AT5G60250.1 | | protein_coding | zinc finger (C3HC4-type R | 11 | 11 | 10 | 226 | 164 | 143 | 23 | 27 | 26 | 490 | 435 | 389 | 169.66 | 25 | 438 | 10.028 | 3.326 | 8.30E-58 |
| AT2G07768.1 | | protein_coding | Cytochrome C assembly p | 258 | 228 | 207 | 386 | 282 | 350 | 542 | 560 | 530 | 837 | 749 | 952 | 347.8 | 544 | 846 | 9.721 | 3.281 | 0.00715478 |
| AT2G07707.1 | YMF19 | protein_coding | Plant mitochondrial ATPa | 6409 | 4821 | 5740 | 6943 | 5821 | 5724 | 13472 | 11844 | 14695 | 15060 | 15453 | 15566 | 7174.51 | 13337 | 15360 | 9.711 | 3.28 | 0.00715922 |
| AT2G37770.2 | AKR4C9 | protein_coding | NAD(P)-linked oxidoredu | 27 | 11 | 2 | 156 | 8 | 6 | 57 | 27 | 5 | 338 | 21 | 16 | 94.66 | 30 | 125 | 9.01 | 3.172 | 9.31E-05 |
| AT2G07811.1 | | pseudogene | | 17910 | 13837 | 14923 | 19663 | 15851 | 16331 | 37648 | 33995 | 38204 | 42651 | 42081 | 44411 | 19916.95 | 36616 | 43048 | 8.4 | 3.07 | 3.08E-05 |
| AT2G04040.1 | DTX1 | protein_coding | MATE efflux family prote | 104 | 93 | 23 | 406 | 121 | 119 | 219 | 228 | 59 | 881 | 321 | 324 | 490.18 | 169 | 509 | 8.235 | 3.042 | 3.36E-13 |
| AT1G02850.4 | BGLU11 | protein_coding | beta glucosidase 11 | 216 | 62 | 17 | 781 | 80 | 61 | 454 | 152 | 44 | 1694 | 212 | 166 | 396.13 | 217 | 691 | 7.798 | 2.963 | 3.78E-05 |
| AT5G58770.1 | | protein_coding | Undecaprenyl pyrophosp | 52 | 19 | 9 | 386 | 105 | 73 | 109 | 47 | 23 | 837 | 279 | 199 | 220.35 | 60 | 438 | 7.329 | 2.874 | 1.73E-10 |
| AT2G15490.1 | UGT73B4 | protein_coding | UDP-glycosyltransferase | 5 | 4 | 3 | 48 | 5 | 3 | 11 | 10 | 8 | 104 | 13 | 8 | 29.76 | 10 | 42 | 7.21 | 2.85 | 3.32E-06 |
| AT1G20390.1 | | transposable_elen | transposable element ger | 327 | 505 | 336 | 1646 | 1451 | 2528 | 687 | 1241 | 860 | 3570 | 3852 | 6875 | 3674.19 | 929 | 4766 | 6.942 | 2.795 | 9.43E-32 |
| AT5G01320.1 | PDC4 | protein_coding | Thiamine pyrophosphate | 6 | 1 | 4 | 39 | 23 | 11 | 13 | 2 | 10 | 85 | 61 | 30 | 31.3 | 8 | 59 | 6.781 | 2.762 | 1.04E-10 |
| AT4G15610.1 | | protein_coding | Uncharacterised protein f | 22 | 17 | 11 | 31 | 12 | 8 | 46 | 42 | 28 | 67 | 32 | 22 | 104.8 | 39 | 40 | 6.682 | 2.74 | 1.42E-26 |
| AT4G22390.1 | | protein_coding | F-box associated ubiquiti | 23 | 23 | 23 | 78 | 52 | 60 | 48 | 57 | 59 | 169 | 138 | 163 | 98.57 | 55 | 157 | 6.394 | 2.677 | 1.41E-30 |
| AT4G25020.1 | | protein_coding | D111/G-patch domain-co | 15 | 17 | 16 | 77 | 57 | 47 | 32 | 42 | 41 | 167 | 151 | 128 | 84.6 | 38 | 149 | 6.018 | 2.589 | 2.03E-32 |
| AT1G01060.1 | LHY | protein_coding | Homeodomain-like super | 175 | 110 | 122 | 881 | 702 | 735 | 368 | 270 | 312 | 1911 | 1864 | 1999 | 713.78 | 317 | 1925 | 5.956 | 2.574 | 8.77E-62 |
| AT1G11070.1 | | protein_coding | | 197 | 135 | 97 | 891 | 421 | 484 | 414 | 332 | 248 | 1933 | 1118 | 1316 | 702.62 | 331 | 1456 | 5.915 | 2.564 | 3.20E-33 |
| AT1G43800.1 | S-ACP-DES6 | protein_coding | Plant stearoyl-acyl-carrie | 5 | 12 | 4 | 30 | 18 | 55 | 11 | 29 | 10 | 65 | 48 | 150 | 45.28 | 17 | 88 | 5.108 | 2.353 | 1.40E-06 |
| AT2G04460.1 | | transposable_elen | transposable element ger | 32 | 7 | 1 | 126 | 14 | 8 | 67 | 17 | 3 | 273 | 37 | 22 | 82.78 | 29 | 111 | 5.088 | 2.347 | 0.00174607 |
| AT4G03295.1 | | small_nucleolar_r | snoRNA | 58 | 22 | 36 | 41 | 25 | 15 | 122 | 54 | 92 | 89 | 66 | 41 | 40.58 | 89 | 65 | 4.927 | 2.301 | 0.00041428 |
| AT4G08112.1 | | transposable_elen | transposable element ger | 33 | 40 | 62 | 46 | 74 | 66 | 69 | 98 | 159 | 100 | 196 | 179 | 69.94 | 109 | 158 | 4.757 | 2.25 | 6.57E-05 |
| AT5G07100.1 | WRKY26 | protein_coding | WRKY DNA-binding prote | 10 | 10 | 10 | 60 | 14 | 33 | 21 | 25 | 26 | 130 | 37 | 90 | 47.27 | 24 | 86 | 4.703 | 2.234 | 1.49E-09 |
| AT4G01920.1 | | protein_coding | Cysteine/Histidine-rich C1 | 29 | 18 | 38 | 79 | 63 | 67 | 61 | 44 | 97 | 171 | 167 | 182 | 89.82 | 67 | 173 | 4.683 | 2.227 | 2.91E-19 |
| AT3G12203.1 | SCPL17 | protein_coding | serine carboxypeptidase- | 9 | 7 | 5 | 25 | 23 | 14 | 19 | 17 | 13 | 54 | 61 | 38 | 36.57 | 16 | 51 | 4.617 | 2.207 | 8.48E-13 |
| AT4G16680.1 | | protein_coding | P-loop containing nucleo: | 59 | 86 | 143 | 338 | 266 | 415 | 124 | 211 | 366 | 733 | 706 | 1129 | 360.97 | 234 | 856 | 4.547 | 2.185 | 3.78E-12 |
| AT3G46660.1 | UGT76E12 | protein_coding | UDP-glucosyl transferase | 1 | 6 | 1 | 13 | 2 | 1 | 2 | 15 | 3 | 28 | 5 | 3 | 22.27 | 7 | 12 | 4.494 | 2.168 | 9.54E-07 |
| AT5G58310.1 | MES18 | protein_coding | methyl esterase 18 | 62 | 49 | 16 | 235 | 206 | 137 | 130 | 120 | 41 | 510 | 547 | 373 | 550.85 | 97 | 477 | 4.475 | 2.162 | 1.59E-18 |
| AT1G60110.1 | JAL17 | protein_coding | Mannose-binding lectin s | 43 | 15 | 17 | 116 | 37 | 67 | 90 | 37 | 44 | 252 | 98 | 182 | 96.31 | 57 | 177 | 4.427 | 2.146 | 7.65E-10 |
| AT5G51440.1 | HSP23.5 | protein_coding | HSP20-like chaperones su | 33 | 12 | 3 | 152 | 66 | 67 | 69 | 29 | 8 | 330 | 175 | 182 | 131.6 | 35 | 229 | 4.384 | 2.132 | 5.51E-09 |
| AT5G54190.1 | PORA | protein_coding | protochlorophyllide oxidc | 19 | 45 | 42 | 80 | 314 | 509 | 40 | 111 | 108 | 174 | 834 | 1384 | 318.64 | 86 | 797 | 4.318 | 2.11 | 7.76E-06 |
| AT1G52100.1 | | protein_coding | Mannose-binding lectin s | 2625 | 2428 | 1759 | 10760 | 7976 | 5990 | 5518 | 5965 | 4503 | 23339 | 21174 | 16289 | 13036.72 | 5329 | 20267 | 4.135 | 2.048 | 5.52E-39 |
| AT2G36750.1 | UGT73C1 | protein_coding | UDP-glucosyl transferase | 72 | 27 | 13 | 233 | 72 | 34 | 151 | 66 | 33 | 505 | 191 | 92 | 161.16 | 83 | 263 | 4.102 | 2.036 | 1.85E-05 |
| AT3G22840.1 | ELIP1 | protein_coding | Chlorophyll A-B binding fa | 75 | 26 | 14 | 353 | 123 | 209 | 158 | 64 | 36 | 766 | 327 | 568 | 266.11 | 86 | 554 | 4.102 | 2.036 | 2.11E-08 |
| AT5G18840.1 | | protein_coding | Major facilitator superfar | 17 | 12 | 5 | 53 | 18 | 10 | 36 | 29 | 13 | 115 | 48 | 27 | 56.08 | 26 | 63 | 4.08 | 2.029 | 3.51E-08 |
| AT3G01600.1 | anac044 | protein_coding | NAC domain containing p | 14 | 18 | 6 | 74 | 57 | 67 | 29 | 44 | 15 | 161 | 151 | 182 | 88.55 | 29 | 165 | 4.068 | 2.024 | 2.62E-16 |
| AT2G27420.1 | | protein_coding | Cysteine proteinases supe | 91 | 51 | 21 | 344 | 214 | 86 | 191 | 125 | 54 | 746 | 568 | 234 | 310.39 | 123 | 516 | 4.062 | 2.022 | 4.44E-09 |
| AT3G23550.1 | LAL5 | protein_coding | MATE efflux family prote | 22 | 12 | 12 | 96 | 15 | 19 | 46 | 29 | 31 | 208 | 40 | 52 | 72.46 | 35 | 100 | 4.021 | 2.008 | 3.43E-06 |
| AT3G18610.1 | NUCL2 | protein_coding | nucleolin like 2 | 31 | 22 | 21 | 99 | 85 | 68 | 65 | 54 | 54 | 215 | 226 | 185 | 121.46 | 58 | 209 | 4.001 | 2.001 | 7.35E-20 |
| AT1G09500.1 | | protein_coding | NAD(P)-binding Rossman | 8 | 12 | 9 | 29 | 9 | 12 | 17 | 29 | 23 | 63 | 24 | 33 | 43.88 | 23 | 40 | 3.917 | 1.97 | 5.51E-10 |
| AT3G49620.1 | DIN11 | protein_coding | 2-oxoglutarate (2OG) anc | 33 | 40 | 47 | 96 | 55 | 110 | 69 | 98 | 120 | 208 | 146 | 299 | 117.84 | 96 | 218 | 3.806 | 1.928 | 4.99E-13 |
| AT3G47580.1 | | protein_coding | Leucine-rich repeat prote | 110 | 114 | 92 | 237 | 176 | 115 | 231 | 280 | 236 | 514 | 467 | 313 | 252.19 | 249 | 431 | 3.754 | 1.909 | 4.18E-26 |
| AT2G39330.1 | JAL23 | protein_coding | jacalin-related lectin 23 | 1 | 6 | 2 | 54 | 50 | 26 | 2 | 15 | 5 | 117 | 133 | 71 | 50.01 | 7 | 107 | 3.706 | 1.89 | 1.54E-07 |
| AT2G17690.1 | SDC | protein_coding | F-box family protein with | 11 | 6 | 5 | 18 | 12 | 15 | 23 | 15 | 13 | 39 | 32 | 41 | 38.61 | 17 | 37 | 3.705 | 1.889 | 8.09E-09 |
| AT5G35940.1 | JAL41 | protein_coding | Mannose-binding lectin s | 31 | 10 | 10 | 189 | 83 | 47 | 65 | 25 | 26 | 410 | 220 | 128 | 132.84 | 39 | 253 | 3.666 | 1.874 | 4.58E-07 |
| AT2G07718.1 | | protein_coding | Cytochrome b/b6 protein | 349 | 190 | 260 | 378 | 274 | 279 | 734 | 467 | 666 | 820 | 727 | 759 | 348.73 | 622 | 769 | 3.641 | 1.864 | 0.00748097 |
| AT5G13210.1 | | protein_coding | Uncharacterised conserve | 18 | 28 | 4 | 34 | 23 | 28 | 38 | 69 | 10 | 74 | 61 | 76 | 56.6 | 39 | 70 | 3.613 | 1.853 | 4.07E-06 |
| AT2G16586.1 | | protein_coding | | 31013 | 30510 | 16987 | 41285 | 31871 | 36193 | 65192 | 74957 | 43488 | 89551 | 84610 | 98425 | 38454.53 | 61212 | 90862 | 3.591 | 1.844 | 2.84E-20 |
| AT5G44980.1 | | protein_coding | F-box/RNI-like/FBD-like d | 30 | 14 | 10 | 94 | 62 | 63 | 63 | 34 | 26 | 204 | 165 | 171 | 96.67 | 41 | 180 | 3.552 | 1.829 | 8.00E-14 |
| AT5G05430.1 | | protein_coding | RNA-binding protein | 22 | 7 | 3 | 34 | 12 | 15 | 46 | 17 | 8 | 74 | 32 | 41 | 40.35 | 24 | 49 | 3.508 | 1.811 | 1.30E-06 |
| AT4G14690.1 | ELIP2 | protein_coding | Chlorophyll A-B binding fa | 124 | 52 | 53 | 653 | 204 | 475 | 261 | 128 | 136 | 1416 | 542 | 1292 | 612.56 | 175 | 1083 | 3.504 | 1.809 | 2.47E-10 |
| AT1G17170.1 | GSTU24 | protein_coding | glutathione S-transferase | 79 | 31 | 9 | 195 | 11 | 19 | 166 | 76 | 23 | 423 | 29 | 52 | 126.07 | 88 | 168 | 3.449 | 1.786 | 0.00897198 |
| AT3G21030.1 | | transposable_elen | transposable element ger | 16 | 13 | 5 | 18 | 15 | 9 | 34 | 32 | 13 | 39 | 40 | 24 | 37.84 | 26 | 34 | 3.402 | 1.766 | 1.30E-10 |
| AT1G08430.1 | ALMT1 | protein_coding | aluminum-activated mala | 36 | 19 | 22 | 68 | 24 | 38 | 76 | 47 | 56 | 147 | 64 | 103 | 78.95 | 60 | 105 | 3.348 | 1.743 | 1.05E-07 |
| AT4G20690.1 | | protein_coding | | 38 | 35 | 19 | 77 | 74 | 60 | 80 | 86 | 49 | 167 | 196 | 163 | 96.09 | 72 | 175 | 3.342 | 1.741 | 4.52E-12 |
| AT2G11270.1 | | protein_coding | citrate synthase-related | 30 | 18 | 30 | 45 | 41 | 41 | 63 | 44 | 77 | 98 | 109 | 111 | 46.77 | 61 | 106 | 3.339 | 1.739 | 5.38E-05 |
| AT5G43450.1 | | protein_coding | 2-oxoglutarate (2OG) anc | 123 | 127 | 181 | 574 | 289 | 335 | 259 | 312 | 463 | 1245 | 767 | 911 | 578.88 | 345 | 974 | 3.31 | 1.727 | 2.61E-16 |
| AT2G36790.1 | UGT73C6 | protein_coding | UDP-glucosyl transferase | 34 | 18 | 17 | 99 | 50 | 38 | 71 | 44 | 44 | 215 | 133 | 103 | 82.56 | 53 | 150 | 3.282 | 1.714 | 3.21E-07 |
| AT5G25230.1 | | protein_coding | Ribosomal protein S5/Elo | 140 | 96 | 94 | 195 | 132 | 127 | 294 | 236 | 241 | 423 | 350 | 345 | 210.32 | 257 | 373 | 3.265 | 1.707 | 4.38E-10 |
| AT4G31520.1 | | protein_coding | SDA1 family protein | 22 | 6 | 7 | 39 | 28 | 27 | 46 | 15 | 18 | 85 | 74 | 73 | 68.67 | 26 | 77 | 3.238 | 1.695 | 1.50E-09 |
| AT5G48780.1 | | protein_coding | disease resistance proteir | 11 | 11 | 27 | 25 | 19 | 23 | 23 | 27 | 69 | 54 | 50 | 63 | 39.02 | 40 | 56 | 3.22 | 1.687 | 1.62E-06 |
| AT2G41480.1 | PER25 | protein_coding | Peroxidase superfamily p | 8 | 3 | 4 | 27 | 17 | 13 | 17 | 7 | 10 | 59 | 45 | 35 | 29.9 | 11 | 46 | 3.204 | 1.68 | 0.0001601 |
| AT3G16360.2 | AHP4 | protein_coding | HPT phosphotransmitter | 28 | 25 | 24 | 74 | 79 | 52 | 59 | 61 | 61 | 161 | 210 | 141 | 137.21 | 60 | 171 | 3.202 | 1.679 | 4.21E-14 |
| AT1G14640.1 | | protein_coding | SWAP (Suppressor-of-Wh | 38 | 42 | 50 | 119 | 118 | 157 | 80 | 103 | 128 | 258 | 313 | 427 | 201.84 | 104 | 333 | 3.181 | 1.669 | 2.94E-16 |
| AT1G49920.1 | | protein_coding | MuDR family transposase | 25 | 16 | 15 | 52 | 36 | 23 | 53 | 39 | 38 | 113 | 96 | 63 | 61.74 | 43 | 91 | 3.174 | 1.666 | 7.27E-09 |
| AT4G33560.1 | | protein_coding | Wound-responsive family | 5 | 7 | 2 | 18 | 6 | 10 | 11 | 17 | 5 | 39 | 16 | 27 | 26.46 | 11 | 27 | 3.143 | 1.652 | 4.54E-06 |
| AT2G34700.1 | | protein_coding | Pollen Ole e 1 allergen ar | 8 | 14 | 17 | 33 | 28 | 30 | 17 | 34 | 44 | 72 | 74 | 82 | 49.99 | 32 | 76 | 3.09 | 1.627 | 1.33E-07 |
| AT5G67090.1 | | protein_coding | Subtilisin-like serine endc | 16 | 14 | 12 | 34 | 34 | 36 | 34 | 34 | 31 | 74 | 90 | 98 | 46.44 | 33 | 87 | 3.085 | 1.625 | 2.98E-06 |
| AT3G27620.1 | AOX1C | protein_coding | alternative oxidase 1C | 19 | 6 | 9 | 41 | 18 | 16 | 40 | 15 | 23 | 89 | 48 | 44 | 40.71 | 26 | 60 | 3.078 | 1.622 | 4.39E-06 |
| AT3G13438.1 | SADHU7-1 | transposable_elen | transposable element ger | 29 | 26 | 26 | 60 | 40 | 72 | 61 | 64 | 67 | 130 | 106 | 196 | 225.52 | 64 | 144 | 3.072 | 1.619 | 2.26E-18 |
| AT4G15620.1 | | protein_coding | Uncharacterised protein f | 179 | 151 | 78 | 336 | 355 | 166 | 376 | 371 | 200 | 729 | 942 | 451 | 911.84 | 316 | 707 | 3.058 | 1.613 | 6.23E-12 |
| AT5G05420.1 | FKBP15-3 | protein_coding | FKBP-like peptidyl-prolyl | 14 | 5 | 7 | 20 | 12 | 13 | 29 | 12 | 18 | 43 | 32 | 35 | 39.9 | 20 | 37 | 3.051 | 1.609 | 2.45E-05 |
| AT1G65370.1 | | protein_coding | TRAF-like family protein | 178 | 146 | 160 | 618 | 494 | 433 | 374 | 359 | 410 | 1341 | 1311 | 1178 | 932.66 | 381 | 1277 | 3.051 | 1.609 | 8.49E-40 |
| AT3G09680.1 | RPS23A | protein_coding | Ribosomal protein S12/S2 | 14 | 4 | 2 | 26 | 19 | 11 | 29 | 10 | 5 | 56 | 50 | 30 | 48.07 | 15 | 45 | 3.039 | 1.604 | 2.67E-06 |
| AT1G05490.1 | CLSY3 | protein_coding | chromatin remodeling 31 | 40 | 26 | 54 | 108 | 102 | 166 | 84 | 64 | 138 | 234 | 271 | 451 | 133.46 | 95 | 319 | 3.021 | 1.595 | 1.65E-08 |
| AT3G21720.1 | ICL | protein_coding | isocitrate lyase | 63 | 52 | 21 | 470 | 348 | 455 | 132 | 128 | 54 | 1019 | 924 | 1237 | 418.37 | 105 | 1060 | 3.018 | 1.594 | 2.68E-10 |
| AT3G44690.1 | | protein_coding | | 162 | 126 | 134 | 297 | 199 | 208 | 341 | 310 | 343 | 644 | 528 | 566 | 305.81 | 331 | 579 | 3.016 | 1.593 | 1.73E-29 |
| AT1G71400.1 | RLP12 | protein_coding | receptor like protein 12 | 121 | 54 | 46 | 141 | 86 | 72 | 254 | 133 | 118 | 306 | 228 | 196 | 189.11 | 168 | 243 | 2.98 | 1.575 | 5.21E-11 |
| AT5G42280.1 | | protein_coding | Cysteine/Histidine-rich C1 | 54 | 82 | 74 | 133 | 100 | 134 | 114 | 201 | 189 | 288 | 265 | 364 | 198.64 | 168 | 306 | 2.98 | 1.575 | 1.50E-13 |
| AT2G31141.1 | | protein_coding | | 10575 | 12205 | 8247 | 12139 | 13174 | 14149 | 22229 | 29985 | 21113 | 26331 | 34974 | 38477 | 14445.82 | 24442 | 33261 | 2.977 | 1.574 | 1.22E-10 |
| AT3G27500.1 | | protein_coding | Cysteine/Histidine-rich C1 | 37 | 12 | 18 | 46 | 29 | 26 | 78 | 29 | 46 | 100 | 77 | 71 | 54.11 | 51 | 83 | 2.942 | 1.557 | 7.93E-06 |
| AT2G39240.1 | | protein_coding | RNA polymerase I specific | 13 | 13 | 9 | 25 | 25 | 13 | 27 | 32 | 23 | 54 | 66 | 35 | 50.1 | 27 | 52 | 2.943 | 1.557 | 9.50E-10 |
| AT5G52320.1 | CYP96A4 | protein_coding | cytochrome P450, family | 63 | 60 | 31 | 168 | 208 | 97 | 132 | 147 | 79 | 364 | 552 | 264 | 229.15 | 119 | 393 | 2.941 | 1.556 | 3.07E-11 |
| AT2G23673.1 | | pseudogene | | 11 | 9 | 10 | 35 | 28 | 10 | 23 | 22 | 26 | 76 | 74 | 27 | 54.41 | 24 | 59 | 2.93 | 1.551 | 5.79E-09 |
| AT4G37730.1 | AtbZIP7 | protein_coding | basic leucine-zipper 7 | 14 | 10 | 14 | 39 | 27 | 26 | 29 | 25 | 36 | 85 | 72 | 71 | 45.81 | 30 | 76 | 2.925 | 1.549 | 9.70E-09 |
| AT1G22150.1 | SULTR1 | protein_coding | sulfate transporter 1;3 | 10 | 8 | 2 | 43 | 19 | 6 | 21 | 20 | 5 | 93 | 50 | 16 | 32.23 | 15 | 53 | 2.921 | 1.546 | 0.0010451 |
| AT3G14210.1 | ESM1 | protein_coding | epithiospecifier modifier | 5630 | 2830 | 2046 | 8640 | 4649 | 5039 | 11835 | 6953 | 5238 | 18741 | 12342 | 13703 | 10478.2 | 8009 | 14929 | 2.914 | 1.543 | 3.41E-11 |
| AT4G02660.1 | | protein_coding | Beige/BEACH domain ;WI | 197 | 132 | 120 | 403 | 337 | 249 | 414 | 324 | 307 | 874 | 895 | 677 | 388.87 | 348 | 815 | 2.914 | 1.543 | 3.43E-17 |
| AT5G59240.1 | RPS8B | protein_coding | Ribosomal protein S8e fa | 69 | 23 | 18 | 77 | 59 | 46 | 145 | 57 | 46 | 167 | 157 | 125 | 141.96 | 83 | 150 | 2.868 | 1.52 | 3.90E-09 |
| AT1G19320.1 | | protein_coding | Pathogenesis-related tha | 1 | 2 | 3 | 3 | 2 | 6 | 2 | 5 | 8 | 7 | 5 | 16 | 18 | 5 | 9 | 2.86 | 1.516 | 1.87E-05 |
| AT5G05400.1 | | protein_coding | LRR and NB-ARC domains | 44 | 19 | 37 | 61 | 67 | 42 | 92 | 47 | 95 | 132 | 178 | 114 | 86.64 | 78 | 141 | 2.853 | 1.513 | 8.53E-08 |
| AT2G32550.1 | | protein_coding | Cell differentiation, Rcd1- | 155 | 97 | 97 | 221 | 147 | 132 | 326 | 238 | 248 | 479 | 390 | 359 | 352.91 | 271 | 409 | 2.842 | 1.507 | 1.40E-18 |
| AT5G06990.1 | | protein_coding | Protein of unknown funct | 10 | 2 | 2 | 19 | 6 | 10 | 21 | 5 | 5 | 41 | 16 | 27 | 26.04 | 10 | 28 | 2.828 | 1.5 | 2.12E-05 |
| AT2G13580.1 | | pseudogene | | 8 | 7 | 3 | 6 | 11 | 13 | 17 | 17 | 8 | 13 | 29 | 35 | 38.63 | 14 | 26 | 2.828 | 1.5 | 2.78E-06 |
| AT3G06900.1 | U4.2 | small_nuclear_rna | U4.2; snRNA | 6269 | 6342 | 4158 | 7131 | 7341 | 4874 | 13178 | 15581 | 10645 | 15468 | 19489 | 13255 | 7311.19 | 13135 | 16071 | 2.825 | 1.498 | 1.36E-07 |
| AT5G05600.1 | | protein_coding | 2-oxoglutarate (2OG) anc | 46 | 29 | 22 | 108 | 30 | 23 | 97 | 71 | 56 | 234 | 80 | 63 | 104 | 75 | 126 | 2.822 | 1.496 | 1.53E-05 |
| AT2G37760.2 | AKR4C8 | protein_coding | NAD(P)-linked oxidoredu | 179 | 56 | 26 | 378 | 69 | 77 | 376 | 138 | 67 | 820 | 183 | 209 | 275.53 | 194 | 404 | 2.783 | 1.477 | 0.00321397 |
| AT5G16980.2 | | protein_coding | Zinc-binding dehydrogena | 111 | 44 | 35 | 222 | 93 | 56 | 233 | 108 | 90 | 482 | 247 | 152 | 242.89 | 144 | 294 | 2.78 | 1.475 | 2.77E-05 |
| AT2G40100.1 | LHCB4.3 | protein_coding | light harvesting complex | 372 | 307 | 252 | 1132 | 769 | 1350 | 782 | 754 | 645 | 2455 | 2042 | 3671 | 2145.78 | 727 | 2723 | 2.777 | 1.473 | 1.44E-15 |
| AT5G06720.1 | PER53 | protein_coding | peroxidase 2 | 25 | 25 | 22 | 110 | 53 | 58 | 53 | 61 | 56 | 239 | 141 | 158 | 148.93 | 57 | 179 | 2.752 | 1.461 | 3.83E-11 |
| AT3G53650.1 | | protein_coding | Histone superfamily prot | 161 | 127 | 84 | 429 | 428 | 360 | 338 | 312 | 215 | 931 | 1136 | 979 | 615.98 | 288 | 1015 | 2.745 | 1.457 | 3.78E-24 |
| AT5G06190.2 | | protein_coding | | 12 | 10 | 13 | 39 | 31 | 26 | 25 | 25 | 33 | 85 | 82 | 71 | 47.32 | 28 | 79 | 2.74 | 1.454 | 7.02E-09 |
| AT4G23600.1 | CORI3 | protein_coding | Tyrosine transaminase fa | 69 | 63 | 37 | 222 | 188 | 106 | 145 | 155 | 95 | 482 | 499 | 288 | 279.38 | 132 | 423 | 2.714 | 1.44 | 1.54E-11 |
| AT3G54510.2 | | protein_coding | Early-responsive to dehyc | 20 | 13 | 38 | 34 | 24 | 49 | 42 | 32 | 97 | 74 | 64 | 133 | 53.26 | 57 | 90 | 2.711 | 1.439 | 0.00042867 |
| AT3G12580.1 | MED37C | protein_coding | heat shock protein 70 | 338 | 172 | 73 | 576 | 510 | 296 | 711 | 423 | 187 | 1249 | 1354 | 805 | 709.4 | 440 | 1136 | 2.712 | 1.439 | 6.38E-06 |
| AT5G13330.1 | ERF113 | protein_coding | related to AP2 6l | 21 | 12 | 11 | 42 | 21 | 47 | 44 | 29 | 28 | 91 | 56 | 128 | 58.32 | 34 | 92 | 2.7 | 1.433 | 2.68E-05 |
| AT4G34135.1 | UGT73B2 | protein_coding | UDP-glucosyltransferase | 50 | 33 | 9 | 168 | 30 | 32 | 105 | 81 | 23 | 364 | 80 | 87 | 189.49 | 70 | 177 | 2.69 | 1.428 | 0.00223205 |
| AT3G09450.1 | | protein_coding | | 67 | 24 | 35 | 123 | 31 | 54 | 141 | 59 | 90 | 267 | 82 | 147 | 92.86 | 97 | 165 | 2.67 | 1.417 | 0.00039479 |
| AT3G16530.1 | | protein_coding | Legume lectin family prot | 527 | 154 | 87 | 1669 | 351 | 248 | 1108 | 378 | 223 | 3620 | 932 | 674 | 1411.15 | 570 | 1742 | 2.664 | 1.414 | 0.0046418 |
| AT4G01870.1 | | protein_coding | tolB protein-related | 295 | 226 | 167 | 877 | 162 | 314 | 620 | 555 | 428 | 1902 | 430 | 854 | 855.86 | 534 | 1062 | 2.651 | 1.406 | 6.24E-05 |
| AT5G23480.1 | | protein_coding | SWIB/MDM2 domain;Plu | 75 | 50 | 29 | 78 | 58 | 37 | 158 | 123 | 74 | 169 | 154 | 101 | 121.23 | 118 | 141 | 2.631 | 1.396 | 2.65E-09 |
| AT3G43580.1 | | protein_coding | Beta-galactosidase relate | 16 | 11 | 16 | 31 | 26 | 11 | 34 | 27 | 41 | 67 | 69 | 30 | 55.26 | 34 | 55 | 2.623 | 1.391 | 9.66E-07 |
| AT5G54075.1 | U3D | small_nucleolar_r | U3D; snoRNA | 11623 | 12295 | 8119 | 17390 | 15327 | 20741 | 24432 | 30207 | 20785 | 37721 | 40690 | 56404 | 17587.69 | 25141 | 44938 | 2.622 | 1.391 | 1.82E-11 |
| AT3G58270.1 | | protein_coding | Arabidopsis phospholipas | 33 | 19 | 23 | 68 | 60 | 53 | 69 | 47 | 59 | 147 | 159 | 144 | 80.4 | 58 | 150 | 2.619 | 1.389 | 6.20E-10 |
| AT4G38780.1 | | protein_coding | Pre-mRNA-processing-spl | 52 | 66 | 45 | 124 | 106 | 84 | 109 | 162 | 115 | 269 | 281 | 228 | 153.51 | 129 | 259 | 2.615 | 1.387 | 7.82E-15 |
| AT1G05560.1 | UGT75B1 | protein_coding | UDP-glucosyltransferase | 234 | 215 | 108 | 798 | 225 | 247 | 492 | 528 | 276 | 1731 | 597 | 672 | 796.49 | 432 | 1000 | 2.614 | 1.386 | 2.46E-06 |
| AT4G34131.1 | UGT73B3 | protein_coding | UDP-glucosyl transferase | 17 | 15 | 7 | 47 | 19 | 12 | 36 | 37 | 18 | 102 | 50 | 33 | 52.95 | 30 | 62 | 2.6 | 1.379 | 2.86E-05 |
| AT5G22300.1 | NIT4 | protein_coding | nitrilase 4 | 79 | 52 | 40 | 161 | 73 | 62 | 166 | 128 | 102 | 349 | 194 | 169 | 225.87 | 132 | 237 | 2.59 | 1.373 | 7.50E-09 |
| AT1G14490.1 | AHL28 | protein_coding | Predicted AT-hook DNA-b | 1 | 5 | 1 | 5 | 3 | 4 | 2 | 12 | 3 | 11 | 8 | 11 | 28.19 | 6 | 10 | 2.586 | 1.371 | 0.00169085 |
| AT2G16140.1 | | transposable_elen | transposable element ger | 157 | 318 | 240 | 220 | 105 | 72 | 330 | 781 | 614 | 477 | 279 | 196 | 227.47 | 575 | 317 | 2.577 | 1.365 | 0.00309977 |
| AT5G62210.1 | | protein_coding | Embryo-specific protein 3 | 37 | 21 | 16 | 53 | 38 | 53 | 78 | 52 | 41 | 115 | 101 | 144 | 130.32 | 57 | 120 | 2.567 | 1.36 | 4.03E-13 |
| AT3G56290.1 | | protein_coding | | 726 | 528 | 359 | 1739 | 1113 | 922 | 1526 | 1297 | 919 | 3772 | 2955 | 2507 | 2257.43 | 1247 | 3078 | 2.566 | 1.359 | 5.52E-15 |
| AT3G13080.1 | ABCC3 | protein_coding | multidrug resistance-assc | 1390 | 744 | 674 | 3107 | 1801 | 1282 | 2922 | 1828 | 1725 | 6739 | 4781 | 3486 | 2490.34 | 2158 | 5002 | 2.557 | 1.355 | 1.16E-09 |
| AT2G29320.1 | | protein_coding | NAD(P)-binding Rossman | 293 | 260 | 219 | 654 | 464 | 368 | 616 | 639 | 561 | 1419 | 1232 | 1001 | 853.6 | 605 | 1217 | 2.512 | 1.329 | 6.11E-23 |
| AT5G08600.2 | | protein_coding | U3 ribonucleoprotein (Ut | 216 | 116 | 64 | 258 | 173 | 145 | 454 | 285 | 164 | 560 | 459 | 394 | 325.66 | 301 | 471 | 2.504 | 1.324 | 3.49E-08 |
| AT4G33070.1 | PDC1 | protein_coding | Thiamine pyrophosphate | 11 | 9 | 8 | 39 | 14 | 37 | 23 | 22 | 20 | 85 | 37 | 101 | 40.28 | 22 | 74 | 2.488 | 1.315 | 0.00312637 |
| AT1G78190.1 | | protein_coding | Trm112p-like protein | 32 | 22 | 11 | 76 | 35 | 54 | 67 | 54 | 28 | 165 | 93 | 147 | 94.07 | 50 | 135 | 2.475 | 1.308 | 1.20E-06 |
| AT5G67160.1 | EPS1 | protein_coding | HXXXD-type acyl-transfer | 42 | 47 | 35 | 131 | 110 | 75 | 88 | 115 | 90 | 284 | 292 | 204 | 160.07 | 98 | 260 | 2.47 | 1.304 | 3.74E-12 |
| AT3G28270.2 | | protein_coding | Protein of unknown funct | 103 | 60 | 50 | 204 | 123 | 61 | 217 | 147 | 128 | 442 | 327 | 166 | 178.27 | 164 | 312 | 2.465 | 1.302 | 1.57E-05 |

| | | | | | | | | | | | | | | | | | | | | | |
|---|---|---|---|---|---|---|---|---|---|---|---|---|---|---|---|---|---|---|---|---|---|
| AT1G74870.1 | | protein_coding | RING/U-box superfamily | 26 | 8 | 13 | 53 | 23 | 18 | 55 | 20 | 33 | 115 | 61 | 49 | 50.52 | 36 | 75 | 2.451 | 1.294 | 0.00011614 |
| AT1G64563.1 | | other_rna | other RNA | 79 | 62 | 78 | 145 | 126 | 93 | 166 | 152 | 200 | 315 | 335 | 253 | 143.5 | 173 | 301 | 2.45 | 1.293 | 6.24E-09 |
| AT2G38230.1 | PDX11 | protein_coding | pyridoxine biosynthesis 1 | 246 | 316 | 199 | 532 | 457 | 695 | 517 | 776 | 509 | 1154 | 1213 | 1890 | 3619.02 | 601 | 1419 | 2.445 | 1.29 | 6.18E-13 |
| AT1G61580.1 | ARP2 | protein_coding | R-protein L3 B | 834 | 635 | 390 | 1975 | 1310 | 1235 | 1753 | 1560 | 998 | 4284 | 3478 | 3359 | 2639.16 | 1437 | 3707 | 2.437 | 1.285 | 3.28E-13 |
| AT4G33330.2 | GUX2 | protein_coding | plant glycogenin-like star | 8 | 10 | 21 | 28 | 13 | 25 | 17 | 25 | 54 | 61 | 35 | 68 | 37.95 | 32 | 55 | 2.426 | 1.279 | 8.36E-05 |
| AT5G57550.1 | XTH25 | protein_coding | xyloglucan endotransgluc | 9 | 9 | 24 | 39 | 11 | 28 | 19 | 22 | 61 | 85 | 29 | 76 | 44 | 34 | 63 | 2.422 | 1.276 | 0.00135059 |
| AT5G64060.1 | anac103 | protein_coding | NAC domain containing p | 40 | 20 | 28 | 95 | 76 | 52 | 84 | 49 | 72 | 206 | 202 | 141 | 106.51 | 68 | 183 | 2.419 | 1.274 | 6.65E-08 |
| AT1G55380.1 | | protein_coding | Cysteine/Histidine-rich C1 | 14 | 13 | 12 | 44 | 24 | 12 | 29 | 32 | 31 | 95 | 64 | 33 | 54.54 | 31 | 64 | 2.415 | 1.272 | 4.46E-05 |
| AT5G09910.1 | | protein_coding | Ras-related small GTP-bir | 114 | 108 | 123 | 198 | 140 | 90 | 240 | 265 | 315 | 429 | 372 | 245 | 158.62 | 273 | 349 | 2.404 | 1.266 | 0.00378759 |
| AT5G52570.1 | BETA-OHASE | protein_coding | beta-carotene hydroxylas | 219 | 176 | 202 | 489 | 304 | 562 | 460 | 432 | 517 | 1061 | 807 | 1528 | 726.54 | 470 | 1132 | 2.4 | 1.263 | 3.07E-12 |
| AT5G15850.1 | COL1 | protein_coding | CONSTANS-like 1 | 245 | 139 | 112 | 533 | 222 | 342 | 515 | 341 | 287 | 1156 | 589 | 930 | 499.28 | 381 | 892 | 2.399 | 1.262 | 1.26E-07 |
| AT5G28630.1 | | protein_coding | glycine-rich protein | 6 | 6 | 1 | 6 | 3 | 2 | 13 | 15 | 3 | 13 | 8 | 5 | 59.5 | 10 | 9 | 2.398 | 1.262 | 2.70E-09 |
| AT5G34871.1 | | other_rna | other RNA | 9 | 17 | 9 | 13 | 6 | 29 | 19 | 42 | 23 | 28 | 16 | 79 | 116.91 | 28 | 41 | 2.397 | 1.261 | 2.76E-06 |
| AT2G17036.1 | | protein_coding | F-box family protein with | 35 | 31 | 46 | 61 | 59 | 76 | 74 | 76 | 118 | 132 | 157 | 207 | 114.72 | 89 | 165 | 2.395 | 1.26 | 5.26E-09 |
| AT5G25140.1 | CYP71B13 | protein_coding | cytochrome P450, family | 88 | 35 | 41 | 156 | 76 | 54 | 185 | 86 | 105 | 338 | 202 | 147 | 168.58 | 125 | 229 | 2.394 | 1.259 | 6.05E-06 |
| AT3G25820.1 | TPS27 | protein_coding | terpene synthase-like seq | 35 | 7 | 12 | 91 | 44 | 17 | 74 | 17 | 31 | 197 | 117 | 46 | 60.52 | 41 | 120 | 2.391 | 1.258 | 0.00771275 |
| AT4G04223.1 | | other_rna | other RNA | 43 | 31 | 36 | 73 | 54 | 47 | 90 | 76 | 92 | 158 | 143 | 128 | 156.88 | 86 | 143 | 2.39 | 1.257 | 2.87E-16 |
| AT3G21020.1 | | transposable_elen | transposable element ger | 22 | 12 | 16 | 23 | 24 | 16 | 46 | 29 | 41 | 50 | 64 | 44 | 51.09 | 39 | 53 | 2.387 | 1.255 | 2.40E-07 |
| AT3G44765.1 | | other_rna | other RNA | 27 | 33 | 34 | 51 | 37 | 26 | 57 | 81 | 87 | 111 | 98 | 71 | 46.37 | 75 | 93 | 2.379 | 1.25 | 0.00333717 |
| AT5G44270.1 | | protein_coding | TPX2 (targeting protein fo | 17 | 13 | 13 | 43 | 25 | 18 | 36 | 32 | 33 | 93 | 66 | 49 | 58.37 | 34 | 69 | 2.367 | 1.243 | 4.64E-06 |
| AT3G23560.1 | ALF5 | protein_coding | MATE efflux family prote | 90 | 89 | 78 | 162 | 132 | 136 | 189 | 219 | 200 | 351 | 350 | 370 | 268 | 203 | 357 | 2.364 | 1.241 | 3.88E-14 |
| AT1G07485.1 | | protein_coding | | 2 | 6 | 4 | 21 | 25 | 10 | 4 | 15 | 10 | 46 | 66 | 27 | 32.55 | 10 | 46 | 2.35 | 1.233 | 0.00048987 |
| AT1G50520.1 | CYP705A27 | protein_coding | cytochrome P450, family | 13 | 14 | 14 | 31 | 26 | 25 | 27 | 34 | 36 | 67 | 69 | 68 | 56.52 | 32 | 68 | 2.341 | 1.227 | 5.59E-08 |
| AT1G78860.1 | | protein_coding | D-mannose binding lectir | 97 | 39 | 23 | 173 | 47 | 66 | 204 | 96 | 59 | 375 | 125 | 179 | 154.81 | 120 | 226 | 2.335 | 1.224 | 0.0007795 |
| AT2G15042.1 | | protein_coding | Leucine-rich repeat (LRR) | 7 | 5 | 5 | 16 | 11 | 4 | 15 | 12 | 13 | 35 | 29 | 11 | 28.01 | 13 | 25 | 2.331 | 1.221 | 0.00337619 |
| AT1G64500.1 | | protein_coding | Glutaredoxin family prote | 29 | 35 | 17 | 55 | 39 | 64 | 61 | 86 | 44 | 119 | 104 | 174 | 290.22 | 64 | 132 | 2.332 | 1.221 | 1.10E-12 |
| AT5G42760.2 | | protein_coding | Leucine carboxyl methylt | 94 | 44 | 40 | 272 | 106 | 47 | 198 | 108 | 102 | 590 | 281 | 128 | 221.15 | 136 | 333 | 2.325 | 1.217 | 0.00075239 |
| AT1G31290.1 | AGO3 | protein_coding | ARGONAUTE 3 | 27 | 26 | 35 | 44 | 34 | 51 | 57 | 64 | 90 | 95 | 90 | 139 | 71.06 | 70 | 108 | 2.32 | 1.214 | 8.91E-07 |
| AT5G28030.2 | DES1 | protein_coding | L-cysteine desulfhydrase | 77 | 37 | 42 | 115 | 80 | 55 | 162 | 91 | 108 | 249 | 212 | 150 | 132.11 | 120 | 204 | 2.316 | 1.212 | 6.47E-08 |
| AT1G56170.1 | NFYC2 | protein_coding | nuclear factor Y, subunit | 34 | 40 | 32 | 72 | 69 | 78 | 71 | 98 | 82 | 156 | 183 | 212 | 153.09 | 84 | 184 | 2.31 | 1.208 | 1.83E-13 |
| AT5G37440.1 | | protein_coding | Chaperone DnaJ-domain | 50 | 33 | 40 | 62 | 58 | 59 | 105 | 81 | 102 | 134 | 154 | 160 | 105.65 | 96 | 149 | 2.308 | 1.207 | 1.22E-10 |
| AT3G28210.1 | SAP12 | protein_coding | zinc finger (AN1-like) fam | 7 | 12 | 6 | 41 | 10 | 12 | 15 | 29 | 15 | 89 | 27 | 33 | 35.39 | 20 | 50 | 2.303 | 1.203 | 0.00200798 |
| AT1G61275.1 | U12 | small_nuclear_rna | U12; snRNA | 256 | 261 | 245 | 343 | 389 | 337 | 538 | 641 | 627 | 744 | 1033 | 916 | 380.4 | 602 | 898 | 2.301 | 1.202 | 0.00030922 |
| AT2G06950.1 | | transposable_elen | transposable element ger | 784 | 429 | 349 | 1666 | 954 | 1010 | 1648 | 1054 | 893 | 3614 | 2533 | 2747 | 1897.59 | 1198 | 2965 | 2.3 | 1.202 | 5.75E-10 |
| AT1G35310.1 | MLP168 | protein_coding | MLP-like protein 168 | 32 | 12 | 5 | 41 | 43 | 17 | 67 | 29 | 13 | 89 | 114 | 46 | 54.28 | 36 | 83 | 2.297 | 1.2 | 0.00631363 |
| AT5G66740.1 | | protein_coding | Protein of unknown funct | 28 | 33 | 14 | 57 | 55 | 34 | 59 | 81 | 36 | 124 | 146 | 92 | 96.34 | 59 | 121 | 2.289 | 1.195 | 2.18E-07 |
| AT4G27300.1 | SD11 | protein_coding | S-locus lectin protein kina | 593 | 281 | 241 | 1133 | 560 | 451 | 1247 | 690 | 617 | 2458 | 1487 | 1226 | 1130.06 | 851 | 1724 | 2.288 | 1.194 | 8.48E-07 |
| AT1G32900.1 | GBSS1 | protein_coding | UDP-Glycosyltransferase | 927 | 363 | 279 | 1547 | 843 | 602 | 1949 | 892 | 714 | 3356 | 2238 | 1637 | 1382.19 | 1185 | 2410 | 2.276 | 1.187 | 3.55E-05 |
| AT2G45290.1 | TKL-2 | protein_coding | Transketolase | 286 | 175 | 155 | 577 | 307 | 329 | 601 | 430 | 397 | 1252 | 815 | 895 | 683.69 | 476 | 987 | 2.274 | 1.186 | 4.45E-11 |
| AT1G25053.1 | | pseudogene | | 10 | 12 | 17 | 24 | 25 | 20 | 21 | 29 | 44 | 52 | 66 | 54 | 40.45 | 31 | 57 | 2.273 | 1.185 | 7.39E-05 |
| AT4G09990.1 | GXM2 | protein_coding | Protein of unknown funct | 9 | 14 | 16 | 19 | 20 | 13 | 19 | 34 | 41 | 41 | 53 | 35 | 39.58 | 31 | 43 | 2.272 | 1.184 | 0.00019363 |
| AT1G10370.1 | GSTU17 | protein_coding | Glutathione S-transferase | 522 | 307 | 199 | 1224 | 880 | 669 | 1097 | 754 | 509 | 2655 | 2336 | 1819 | 1567.22 | 787 | 2270 | 2.27 | 1.183 | 6.82E-09 |
| AT1G68570.1 | NPF3.1 | protein_coding | Major facilitator superfar | 210 | 134 | 70 | 423 | 268 | 211 | 441 | 329 | 179 | 918 | 711 | 574 | 424.88 | 316 | 734 | 2.268 | 1.182 | 5.32E-07 |
| AT3G45730.1 | | protein_coding | | 40 | 42 | 41 | 54 | 41 | 77 | 84 | 103 | 105 | 117 | 109 | 209 | 116.03 | 97 | 145 | 2.259 | 1.176 | 6.99E-07 |
| AT1G08115.1 | | small_nuclear_rna | snRNA | 7156 | 9625 | 6784 | 6747 | 8267 | 10566 | 15042 | 23647 | 17367 | 14635 | 21947 | 28734 | 10171.46 | 18685 | 21772 | 2.251 | 1.17 | 0.00016714 |
| AT4G33550.1 | | protein_coding | Bifunctional inhibitor/lipi | 16 | 18 | 15 | 59 | 92 | 52 | 34 | 44 | 38 | 128 | 244 | 141 | 138.05 | 39 | 171 | 2.249 | 1.169 | 8.16E-08 |
| AT4G30180.1 | | protein_coding | sequence-specific DNA bi | 18 | 11 | 26 | 45 | 71 | 81 | 38 | 27 | 67 | 98 | 188 | 220 | 70.65 | 44 | 169 | 2.241 | 1.164 | 0.00279253 |
| AT3G09480.1 | | protein_coding | Histone superfamily prote | 60 | 45 | 32 | 110 | 95 | 82 | 126 | 111 | 82 | 239 | 252 | 223 | 151.75 | 106 | 238 | 2.236 | 1.161 | 5.05E-10 |
| AT5G61820.1 | | protein_coding | | 525 | 321 | 221 | 1066 | 332 | 285 | 1104 | 789 | 566 | 2312 | 881 | 775 | 990.46 | 820 | 1323 | 2.232 | 1.159 | 0.00013933 |
| AT2G42510.2 | | protein_coding | | 48 | 26 | 34 | 106 | 81 | 59 | 101 | 64 | 87 | 230 | 215 | 160 | 123.69 | 84 | 202 | 2.232 | 1.158 | 9.39E-09 |
| AT3G05936.1 | | protein_coding | | 60 | 47 | 42 | 85 | 70 | 42 | 126 | 115 | 108 | 184 | 186 | 114 | 117.06 | 116 | 161 | 2.222 | 1.152 | 3.55E-06 |
| AT5G03555.1 | NCS1 | protein_coding | permease, cytosine/purin | 406 | 314 | 254 | 753 | 533 | 742 | 853 | 771 | 650 | 1633 | 1415 | 2018 | 1129.68 | 758 | 1689 | 2.22 | 1.151 | 9.28E-16 |
| AT3G46540.1 | | protein_coding | ENTH/VHS family protein | 673 | 428 | 318 | 1072 | 777 | 493 | 1415 | 1052 | 814 | 2325 | 2063 | 1341 | 1430.95 | 1094 | 1910 | 2.213 | 1.146 | 4.51E-09 |
| AT1G53830.1 | PME2 | protein_coding | pectin methylesterase 2 | 14 | 30 | 15 | 80 | 43 | 54 | 29 | 74 | 38 | 174 | 114 | 147 | 74.18 | 47 | 145 | 2.208 | 1.143 | 4.17E-05 |
| AT2G38210.1 | PDX1L4 | protein_coding | putative PDX1-like protei | 14 | 29 | 16 | 35 | 28 | 53 | 29 | 71 | 41 | 76 | 74 | 144 | 242.94 | 47 | 98 | 2.208 | 1.143 | 2.31E-10 |
| AT1G21520.1 | | protein_coding | | 8 | 12 | 9 | 33 | 17 | 20 | 17 | 29 | 23 | 72 | 45 | 54 | 45.23 | 23 | 57 | 2.204 | 1.14 | 3.77E-05 |
| AT1G54575.1 | | protein_coding | | 46 | 16 | 13 | 80 | 29 | 24 | 97 | 39 | 33 | 174 | 77 | 65 | 87.99 | 56 | 105 | 2.2 | 1.137 | 0.00190772 |
| AT2G23680.1 | | protein_coding | Cold acclimation protein | 20 | 9 | 5 | 39 | 17 | 8 | 42 | 22 | 13 | 85 | 45 | 22 | 35.15 | 26 | 51 | 2.198 | 1.136 | 0.00687313 |
| AT1G78090.1 | TPPB | protein_coding | trehalose-6-phosphate ph | 34 | 24 | 32 | 36 | 30 | 27 | 71 | 59 | 82 | 78 | 80 | 73 | 107.97 | 71 | 77 | 2.197 | 1.136 | 1.44E-09 |
| AT3G59750.1 | LECRK58 | protein_coding | Concanavalin A-like lectir | 21 | 21 | 6 | 32 | 33 | 8 | 44 | 52 | 15 | 69 | 88 | 22 | 47.55 | 37 | 60 | 2.196 | 1.135 | 0.00327094 |
| AT5G18270.1 | ANAC087 | protein_coding | Arabidopsis NAC domain | 48 | 40 | 22 | 72 | 61 | 54 | 101 | 98 | 56 | 156 | 162 | 147 | 89.16 | 85 | 155 | 2.197 | 1.135 | 0.00011231 |
| AT3G11402.1 | PRA1A3 | protein_coding | Cysteine/Histidine-rich C1 | 41 | 7 | 5 | 36 | 20 | 15 | 86 | 17 | 13 | 78 | 53 | 41 | 58.43 | 39 | 57 | 2.195 | 1.134 | 0.00530771 |
| AT2G15400.1 | NRPD3B | protein_coding | DNA-directed RNA polym | 15 | 16 | 16 | 32 | 27 | 28 | 32 | 39 | 41 | 69 | 72 | 76 | 54.23 | 37 | 72 | 2.192 | 1.132 | 2.48E-05 |
| AT4G35940.1 | | protein_coding | | 121 | 109 | 132 | 321 | 241 | 280 | 254 | 268 | 338 | 696 | 640 | 761 | 420.49 | 287 | 699 | 2.183 | 1.126 | 6.47E-18 |
| AT3G26320.1 | CYP71B36 | protein_coding | cytochrome P450, family | 52 | 39 | 42 | 71 | 93 | 64 | 109 | 96 | 108 | 154 | 247 | 174 | 121.18 | 104 | 192 | 2.177 | 1.122 | 1.48E-06 |
| AT1G29420.1 | SAUR61 | protein_coding | SAUR-like auxin-responsi | 14 | 10 | 23 | 55 | 68 | 39 | 29 | 25 | 59 | 119 | 181 | 106 | 62.94 | 38 | 135 | 2.171 | 1.118 | 0.0001559 |
| AT4G00975.1 | | other_rna | other RNA | 59 | 135 | 51 | 84 | 63 | 107 | 124 | 332 | 131 | 182 | 167 | 291 | 113.62 | 196 | 213 | 2.164 | 1.114 | 0.00540837 |
| AT1G29720.1 | RFK1 | protein_coding | Leucine-rich repeat transı | 102 | 76 | 73 | 143 | 132 | 100 | 214 | 187 | 187 | 310 | 350 | 272 | 222.87 | 196 | 311 | 2.163 | 1.113 | 8.11E-08 |
| AT4G21730.1 | | pseudogene | | 6 | 14 | 16 | 24 | 58 | 39 | 13 | 34 | 41 | 52 | 154 | 106 | 56.77 | 29 | 104 | 2.16 | 1.111 | 0.00171426 |
| AT5G53902.1 | U3B | small_nucleolar_r | U3B; snoRNA | 11485 | 11939 | 7878 | 16668 | 15069 | 20421 | 24142 | 29332 | 20168 | 36154 | 40005 | 55534 | 17154.03 | 24547 | 43898 | 2.156 | 1.108 | 1.21E-05 |
| AT1G77570.1 | | protein_coding | Winged helix-turn-helix t | 32 | 12 | 24 | 62 | 45 | 41 | 67 | 29 | 61 | 134 | 119 | 111 | 74.79 | 52 | 121 | 2.151 | 1.105 | 2.64E-05 |
| AT5G20710.1 | BGAL7 | protein_coding | beta-galactosidase 7 | 27 | 26 | 24 | 52 | 47 | 30 | 57 | 64 | 61 | 113 | 125 | 82 | 72.18 | 61 | 107 | 2.151 | 1.105 | 1.83E-05 |
| AT3G30720.1 | QQS | protein_coding | qua-quine starch | 13 | 9 | 16 | 13 | 22 | 15 | 27 | 22 | 41 | 28 | 58 | 41 | 56.12 | 30 | 42 | 2.151 | 1.105 | 6.81E-06 |
| AT3G21750.1 | UGT71B1 | protein_coding | UDP-glucosyl transferase | 178 | 113 | 74 | 355 | 179 | 191 | 374 | 278 | 189 | 770 | 475 | 519 | 474.19 | 280 | 588 | 2.149 | 1.104 | 1.07E-08 |
| AT3G16030.1 | CES101 | protein_coding | lectin protein kinase fami | 41 | 32 | 31 | 69 | 68 | 61 | 86 | 79 | 79 | 150 | 181 | 166 | 116.52 | 81 | 166 | 2.146 | 1.102 | 1.00E-10 |
| AT4G35640.1 | SAT4 | protein_coding | serine acetyltransferase 3 | 35 | 9 | 9 | 24 | 36 | 13 | 74 | 22 | 23 | 52 | 96 | 35 | 47.03 | 40 | 61 | 2.137 | 1.096 | 0.00572682 |
| AT1G59406.1 | | protein_coding | GDSL-like Lipase/Acylhyd | 40 | 24 | 7 | 69 | 64 | 49 | 84 | 59 | 18 | 150 | 170 | 133 | 103.2 | 54 | 151 | 2.138 | 1.096 | 1.06E-05 |
| AT3G12110.1 | ACT11 | protein_coding | actin-11 | 696 | 472 | 363 | 1104 | 778 | 613 | 1463 | 1160 | 929 | 2395 | 2065 | 1667 | 1782.64 | 1184 | 2042 | 2.137 | 1.096 | 1.68E-11 |
| AT3G62460.1 | | protein_coding | Putative endonuclease or | 280 | 137 | 150 | 441 | 304 | 303 | 589 | 337 | 384 | 957 | 807 | 824 | 638.24 | 437 | 863 | 2.136 | 1.095 | 2.46E-11 |
| AT2G03760.1 | SOT12 | protein_coding | sulphotransferase 12 | 198 | 209 | 168 | 570 | 300 | 345 | 416 | 513 | 430 | 1236 | 796 | 938 | 603.68 | 453 | 990 | 2.137 | 1.095 | 1.82E-13 |
| AT1G13650.2 | | protein_coding | | 673 | 721 | 353 | 835 | 1035 | 370 | 1415 | 1771 | 904 | 1811 | 2748 | 1006 | 1577.06 | 1363 | 1855 | 2.133 | 1.093 | 8.44E-05 |
| AT3G47540.2 | | protein_coding | Chitinase family protein | 27 | 32 | 15 | 48 | 36 | 28 | 57 | 79 | 38 | 104 | 96 | 76 | 82.47 | 58 | 92 | 2.131 | 1.092 | 3.86E-08 |
| AT4G19090.1 | | protein_coding | Protein of unknown funct | 72 | 63 | 48 | 80 | 57 | 50 | 151 | 155 | 123 | 174 | 151 | 136 | 113.4 | 143 | 154 | 2.131 | 1.092 | 2.85E-08 |
| AT1G10960.1 | FD1 | protein_coding | ferredoxin 1 | 2810 | 2088 | 1582 | 6898 | 3856 | 5187 | 5907 | 5130 | 4050 | 14962 | 10237 | 14106 | 13767.24 | 5029 | 13102 | 2.131 | 1.092 | 2.83E-10 |
| AT3G12320.1 | T2E22.34 | protein_coding | | 300 | 198 | 182 | 595 | 398 | 450 | 631 | 486 | 466 | 1291 | 1057 | 1224 | 608 | 528 | 1191 | 2.129 | 1.09 | 1.65E-14 |
| AT1G15300.1 | | transposable_elen | transposable element ger | 54 | 41 | 40 | 71 | 66 | 41 | 114 | 101 | 102 | 154 | 175 | 111 | 94.45 | 106 | 147 | 2.127 | 1.089 | 1.25E-06 |
| AT2G34460.1 | | protein_coding | NAD(P)-binding Rossman | 2002 | 1530 | 1442 | 3101 | 2423 | 2247 | 4208 | 3759 | 3692 | 6726 | 6433 | 6111 | 5363.84 | 3886 | 6423 | 2.123 | 1.086 | 4.37E-22 |
| AT3G17050.1 | | transposable_elen | transposable element ger | 6 | 5 | 7 | 18 | 7 | 4 | 13 | 12 | 18 | 39 | 19 | 11 | 111.3 | 14 | 23 | 2.122 | 1.085 | 3.13E-05 |
| AT1G25209.1 | | pseudogene | | 11 | 13 | 17 | 22 | 41 | 33 | 23 | 32 | 44 | 48 | 109 | 90 | 49.21 | 33 | 82 | 2.117 | 1.082 | 0.00028008 |
| AT1G58725.1 | | protein_coding | GDSL-like Lipase/Acylhyd | 32 | 25 | 15 | 60 | 54 | 52 | 67 | 61 | 38 | 130 | 143 | 141 | 98.14 | 55 | 138 | 2.116 | 1.082 | 1.93E-06 |
| AT5G24280.1 | GMI1 | protein_coding | gamma-irradiation and m | 178 | 130 | 125 | 358 | 332 | 274 | 374 | 319 | 320 | 777 | 881 | 745 | 403.48 | 338 | 801 | 2.117 | 1.082 | 1.32E-11 |
| AT1G14120.1 | | protein_coding | 2-oxoglutarate (2OG) anc | 162 | 130 | 78 | 419 | 202 | 106 | 341 | 319 | 200 | 909 | 536 | 288 | 490.12 | 287 | 578 | 2.11 | 1.077 | 5.78E-05 |
| AT5G35777.1 | | transposable_elen | transposable element ger | 83 | 89 | 85 | 202 | 189 | 118 | 174 | 219 | 218 | 438 | 502 | 321 | 285.39 | 204 | 420 | 2.101 | 1.071 | 1.51E-10 |
| AT3G54160.1 | | protein_coding | RNI-like superfamily prot | 16 | 11 | 12 | 31 | 28 | 18 | 34 | 27 | 31 | 67 | 74 | 49 | 44.22 | 31 | 63 | 2.087 | 1.062 | 2.87E-05 |
| AT5G13930.1 | CHS | protein_coding | Chalcone and stilbene syr | 1028 | 543 | 273 | 2209 | 1352 | 2036 | 2161 | 1334 | 699 | 4792 | 3589 | 5537 | 2767.73 | 1398 | 4639 | 2.082 | 1.058 | 7.90E-05 |
| AT2G21330.1 | FBA1 | protein_coding | fructose-bisphosphate alc | 8779 | 11432 | 5486 | 17749 | 13846 | 22024 | 18454 | 28086 | 14045 | 38499 | 36758 | 59893 | 40987.1 | 20195 | 45050 | 2.083 | 1.058 | 5.41E-06 |
| AT1G64780.1 | AMT1-2 | protein_coding | ammonium transporter 1 | 168 | 112 | 69 | 294 | 209 | 145 | 353 | 275 | 177 | 638 | 555 | 394 | 423.61 | 268 | 529 | 2.08 | 1.056 | 1.69E-08 |
| AT5G26200.1 | | protein_coding | Mitochondrial substrate | 126 | 85 | 145 | 228 | 117 | 260 | 265 | 209 | 371 | 495 | 311 | 707 | 302 | 282 | 504 | 2.078 | 1.055 | 7.97E-06 |
| AT1G15150.1 | | protein_coding | MATE efflux family prote | 13 | 8 | 3 | 21 | 24 | 12 | 27 | 20 | 8 | 46 | 64 | 33 | 42.96 | 18 | 48 | 2.061 | 1.043 | 4.80E-05 |
| AT1G23740.1 | | protein_coding | Oxidoreductase, zinc-binc | 2329 | 2049 | 1517 | 3795 | 3238 | 3075 | 4896 | 5034 | 3884 | 8232 | 8596 | 8362 | 7469.85 | 4605 | 8397 | 2.058 | 1.041 | 4.03E-17 |
| AT2G28190.1 | CSD2 | protein_coding | copper/zinc superoxide d | 144 | 539 | 234 | 199 | 305 | 346 | 303 | 1324 | 599 | 432 | 810 | 941 | 1865.06 | 742 | 728 | 2.057 | 1.04 | 0.0043019 |
| AT3G61630.1 | CRF6 | protein_coding | cytokinin response factor | 48 | 66 | 47 | 114 | 100 | 158 | 101 | 162 | 120 | 247 | 265 | 430 | 172.13 | 128 | 314 | 2.051 | 1.036 | 6.81E-06 |
| AT5G66520.1 | PCMP-H61 | protein_coding | Tetratricopeptide repeat | 331 | 299 | 249 | 528 | 482 | 323 | 696 | 735 | 637 | 1145 | 1280 | 878 | 796.63 | 689 | 1101 | 2.048 | 1.034 | 2.89E-13 |
| AT5G26920.1 | CBP60G | protein_coding | Cam-binding protein 60-l | 25 | 33 | 28 | 37 | 22 | 28 | 53 | 81 | 72 | 80 | 58 | 76 | 53.66 | 69 | 71 | 2.043 | 1.031 | 0.00023721 |
| AT3G55485.1 | | other_rna | | 385 | 285 | 241 | 439 | 480 | 367 | 809 | 700 | 617 | 952 | 1274 | 998 | 450.33 | 709 | 1075 | 2.042 | 1.03 | 0.00385496 |
| AT3G16430.1 | PBP2 | protein_coding | jacalin-related lectin 31 | 192 | 127 | 91 | 568 | 296 | 286 | 404 | 312 | 233 | 1232 | 786 | 778 | 701.96 | 316 | 932 | 2.042 | 1.03 | 3.99E-09 |
| AT4G00380.1 | FDM2 | protein_coding | XH/XS domain-containing | 144 | 122 | 128 | 128 | 139 | 149 | 303 | 300 | 328 | 278 | 369 | 405 | 266.86 | 310 | 351 | 2.041 | 1.03 | 1.76E-10 |
| AT2G22850.1 | AtbZIP6 | protein_coding | basic leucine-zipper 6 | 23 | 24 | 17 | 50 | 45 | 38 | 48 | 59 | 44 | 108 | 119 | 103 | 68.66 | 50 | 110 | 2.029 | 1.021 | 3.58E-05 |
| AT1G73066.1 | | protein_coding | Leucine-rich repeat famil | 93 | 69 | 69 | 94 | 80 | 60 | 195 | 170 | 177 | 204 | 212 | 163 | 102.04 | 181 | 193 | 2.024 | 1.018 | 0.00040385 |
| AT2G39310.1 | JAL22 | protein_coding | jacalin-related lectin 22 | 1553 | 1396 | 977 | 4399 | 3394 | 2933 | 3265 | 3430 | 2501 | 9542 | 9010 | 7976 | 6166.07 | 3065 | 8843 | 2.026 | 1.018 | 1.42E-12 |
| AT2G46830.1 | CCA1 | protein_coding | circadian clock associatec | 407 | 292 | 289 | 695 | 588 | 514 | 856 | 717 | 740 | 1508 | 1561 | 1398 | 871.14 | 771 | 1489 | 2.025 | 1.018 | 6.88E-15 |
| AT3G46650.1 | | protein_coding | UDP-Glycosyltransferase | 10 | 8 | 2 | 32 | 25 | 10 | 21 | 20 | 5 | 69 | 66 | 27 | 38.46 | 15 | 54 | 2.024 | 1.017 | 0.0031967 |
| AT3G02790.1 | | protein_coding | zinc finger (C2H2 type) fa | 191 | 168 | 163 | 395 | 354 | 479 | 401 | 413 | 417 | 857 | 940 | 1303 | 771.79 | 410 | 1033 | 2.011 | 1.008 | 3.76E-11 |
| AT4G03140.1 | | protein_coding | NAD(P)-binding Rossman | 44 | 21 | 32 | 104 | 68 | 70 | 92 | 52 | 82 | 226 | 181 | 190 | 140.08 | 75 | 199 | 2.01 | 1.007 | 8.64E-07 |
| AT3G60960.1 | | protein_coding | Tetratricopeptide repeat | 186 | 116 | 86 | 220 | 164 | 197 | 391 | 285 | 220 | 477 | 435 | 536 | 336.29 | 299 | 483 | 2.01 | 1.007 | 6.42E-10 |
| AT3G59052.1 | CPuORF18 | protein_coding | conserved peptide upstre | 131 | 116 | 116 | 282 | 223 | 203 | 275 | 285 | 297 | 612 | 592 | 552 | 420.1 | 286 | 585 | 2.009 | 1.007 | 4.19E-15 |
| AT3G61560.2 | RTNLB6 | protein_coding | Reticulon family protein | 129 | 100 | 51 | 191 | 139 | 133 | 271 | 246 | 131 | 414 | 369 | 362 | 315.97 | 216 | 382 | 2.006 | 1.004 | 1.00E-09 |
| AT2G27510.1 | FD3 | protein_coding | ferredoxin 3 | 192 | 170 | 124 | 298 | 214 | 202 | 404 | 418 | 317 | 646 | 568 | 549 | 630.9 | 380 | 588 | 2.005 | 1.004 | 6.14E-10 |
| AT3G61880.2 | CYP78A9 | protein_coding | cytochrome p450 78a9 | 14 | 24 | 13 | 25 | 17 | 12 | 29 | 59 | 33 | 54 | 45 | 33 | 46.48 | 40 | 44 | 2.004 | 1.003 | 0.00039037 |
| AT1G48390.1 | | protein_coding | RNI-like superfamily prot | 22 | 19 | 11 | 26 | 30 | 22 | 46 | 47 | 28 | 56 | 80 | 60 | 46.46 | 40 | 65 | 2.004 | 1.003 | 0.00032947 |
| AT2G15480.1 | UGT73B5 | protein_coding | UDP-glucosyl transferase | 33 | 23 | 28 | 96 | 36 | 37 | 69 | 57 | 72 | 208 | 96 | 101 | 97.41 | 66 | 135 | 2.001 | 1.001 | 4.69E-05 |
| AT1G54040.2 | ESP | protein_coding | epithiospecifier protein | 58 | 89 | 33 | 121 | 140 | 123 | 122 | 219 | 84 | 262 | 372 | 334 | 360.19 | 142 | 323 | 1.999 | 0.999 | 1.79E-07 |
| AT2G24100.1 | | protein_coding | | 177 | 101 | 150 | 301 | 183 | 209 | 372 | 248 | 384 | 653 | 486 | 568 | 363.81 | 335 | 569 | 1.999 | 0.999 | 5.54E-10 |
| AT1G53345.1 | | protein_coding | | 94 | 55 | 56 | 111 | 114 | 67 | 198 | 135 | 143 | 241 | 303 | 182 | 169.37 | 159 | 242 | 1.995 | 0.997 | 1.14E-05 |
| AT5G34800.1 | | transposable_elen | transposable element ger | 24 | 21 | 16 | 24 | 23 | 22 | 50 | 52 | 41 | 52 | 61 | 60 | 42.97 | 48 | 58 | 1.993 | 0.995 | 0.00035352 |
| AT2G29165.1 | | transposable_elen | transposable element ger | 21 | 23 | 30 | 21 | 16 | 16 | 44 | 57 | 77 | 46 | 42 | 44 | 48.93 | 59 | 44 | 1.985 | 0.989 | 0.0055175 |
| AT1G09350.1 | GOLS3 | protein_coding | galactinol synthase 3 | 16 | 17 | 29 | 48 | 54 | 93 | 34 | 42 | 74 | 104 | 143 | 253 | 89.22 | 50 | 167 | 1.984 | 0.989 | 0.00393059 |
| AT1G78290.2 | SRK2C | protein_coding | Protein kinase superfamil | 69 | 50 | 34 | 129 | 86 | 85 | 145 | 123 | 87 | 280 | 228 | 231 | 164.22 | 118 | 246 | 1.974 | 0.981 | 1.69E-06 |
| AT5G45730.1 | | protein_coding | Cysteine/Histidine-rich C1 | 24 | 27 | 27 | 35 | 36 | 39 | 50 | 66 | 69 | 76 | 96 | 106 | 63.15 | 62 | 93 | 1.969 | 0.977 | 0.0001719 |
| AT1G78320.1 | GSTU23 | protein_coding | glutathione S-transferase | 16 | 8 | 3 | 40 | 24 | 29 | 34 | 20 | 8 | 87 | 64 | 79 | 50.14 | 21 | 77 | 1.967 | 0.976 | 0.00014475 |
| AT1G53490.1 | HEI10 | protein_coding | RING/U-box superfamily | 27 | 30 | 28 | 15 | 24 | 26 | 57 | 74 | 72 | 33 | 64 | 71 | 135.59 | 68 | 56 | 1.964 | 0.974 | 6.85E-09 |
| AT1G11300.1 | | protein_coding | protein serine/threonine | 273 | 202 | 220 | 420 | 377 | 279 | 574 | 496 | 563 | 911 | 1001 | 759 | 633.16 | 544 | 890 | 1.958 | 0.969 | 1.64E-14 |
| AT2G37460.1 | | protein_coding | nodulin MtN21 /EamA-lik | 253 | 106 | 60 | 368 | 298 | 121 | 532 | 260 | 154 | 798 | 791 | 329 | 413.81 | 315 | 639 | 1.956 | 0.968 | 0.00432537 |
| AT1G14790.1 | RDR1 | protein_coding | RNA-dependent RNA poly | 103 | 103 | 107 | 159 | 141 | 132 | 217 | 253 | 274 | 345 | 374 | 359 | 208.94 | 248 | 359 | 1.954 | 0.967 | 7.63E-07 |
| AT4G20860.1 | | protein_coding | FAD-binding Berberine fa | 264 | 106 | 142 | 376 | 134 | 144 | 555 | 260 | 364 | 816 | 356 | 392 | 312.81 | 393 | 521 | 1.952 | 0.965 | 0.00087166 |
| AT5G59540.1 | | protein_coding | 2-oxoglutarate (2OG) anc | 1545 | 1461 | 1128 | 3142 | 2534 | 1783 | 3248 | 3589 | 2888 | 6815 | 6727 | 4849 | 4532.12 | 3242 | 6130 | 1.952 | 0.965 | 3.22E-11 |
| AT2G21850.1 | | protein_coding | Cysteine/Histidine-rich C1 | 25 | 12 | 16 | 45 | 24 | 33 | 53 | 29 | 41 | 98 | 64 | 90 | 61.55 | 41 | 84 | 1.948 | 0.962 | 0.00023711 |
| AT5G18290.1 | SIP1-2 | protein_coding | Aquaporin-like superfami | 55 | 61 | 51 | 61 | 68 | 63 | 116 | 150 | 131 | 132 | 181 | 171 | 152.73 | 132 | 161 | 1.949 | 0.962 | 1.73E-08 |
| AT5G38565.1 | | protein_coding | F-box/FBD-like domains c | 23 | 3 | 8 | 54 | 24 | 31 | 48 | 7 | 20 | 117 | 64 | 84 | 66.53 | 25 | 88 | 1.942 | 0.957 | 0.00120795 |
| AT2G23000.1 | SCPL10 | protein_coding | serine carboxypeptidase-l | 636 | 535 | 328 | 1067 | 980 | 532 | 1337 | 1314 | 840 | 2314 | 2602 | 1447 | 1247.21 | 1164 | 2121 | 1.94 | 0.956 | 6.46E-06 |
| AT2G46640.3 | | protein_coding | | 142 | 148 | 107 | 302 | 337 | 270 | 298 | 364 | 274 | 655 | 895 | 734 | 518.63 | 312 | 761 | 1.94 | 0.956 | 7.92E-13 |
| AT3G20100.1 | CYP705A19 | protein_coding | cytochrome P450, family | 129 | 132 | 68 | 185 | 170 | 132 | 271 | 324 | 174 | 401 | 451 | 359 | 341.9 | 256 | 404 | 1.935 | 0.953 | 1.10E-05 |
| AT1G59030.1 | | protein_coding | GDSL-like Lipase/Acylhyd | 21 | 14 | 9 | 56 | 42 | 37 | 44 | 34 | 23 | 121 | 112 | 101 | 75.28 | 34 | 111 | 1.928 | 0.947 | 4.61E-06 |
| AT1G51080.1 | | protein_coding | | 242 | 145 | 159 | 307 | 232 | 220 | 509 | 356 | 407 | 666 | 616 | 598 | 569.9 | 424 | 627 | 1.928 | 0.947 | 1.67E-11 |
| AT2G27402.1 | | protein_coding | | 1441 | 1365 | 1299 | 4500 | 3935 | 3212 | 3029 | 3354 | 3326 | 9761 | 10447 | 8735 | 6868.58 | 3236 | 9648 | 1.926 | 0.945 | 6.97E-15 |

| | | | | | | | | | | | | | | | | | | | | | |
|---|---|---|---|---|---|---|---|---|---|---|---|---|---|---|---|---|---|---|---|---|---|
| AT3G59050.1 | PAO3 | protein_coding | polyamine oxidase 3 | 145 | 125 | 116 | 320 | 235 | 205 | 305 | 307 | 297 | 694 | 624 | 557 | 429.32 | 303 | 625 | 1.913 | 0.936 | 5.93E-16 |
| AT1G72140.1 | NPF5.12 | protein_coding | Major facilitator superfar | 95 | 81 | 29 | 163 | 94 | 60 | 200 | 199 | 74 | 354 | 250 | 163 | 207.38 | 158 | 256 | 1.909 | 0.933 | 0.00105142 |
| AT1G75300.1 | | protein_coding | NmrA-like negative trans | 11 | 10 | 13 | 28 | 34 | 25 | 23 | 25 | 33 | 61 | 90 | 68 | 46.32 | 27 | 73 | 1.909 | 0.933 | 0.00079055 |
| AT5G63090.2 | LOB | protein_coding | Lateral organ boundaries | 9 | 10 | 8 | 10 | 7 | 19 | 19 | 25 | 20 | 22 | 19 | 52 | 32.49 | 21 | 31 | 1.908 | 0.932 | 0.00775081 |
| AT1G79520.2 | MTP9 | protein_coding | Cation efflux family prote | 102 | 80 | 49 | 141 | 142 | 115 | 214 | 197 | 125 | 306 | 377 | 313 | 249.48 | 179 | 332 | 1.907 | 0.931 | 7.78E-09 |
| AT5G17760.1 | | protein_coding | P-loop containing nucleo | 106 | 57 | 41 | 131 | 55 | 72 | 223 | 140 | 105 | 284 | 146 | 196 | 170.91 | 156 | 209 | 1.904 | 0.929 | 0.0001035 |
| AT1G68585.1 | | protein_coding | | 83 | 56 | 53 | 192 | 143 | 76 | 174 | 138 | 136 | 416 | 380 | 207 | 271.29 | 149 | 334 | 1.903 | 0.929 | 1.38E-06 |
| AT4G25700.1 | BETA-OHASE | protein_coding | beta-hydroxylase 1 | 479 | 390 | 364 | 859 | 617 | 811 | 1007 | 958 | 932 | 1863 | 1638 | 2205 | 1619.84 | 966 | 1902 | 1.903 | 0.928 | 1.91E-12 |
| AT5G43530.1 | | protein_coding | Helicase protein with RIN | 20 | 20 | 23 | 48 | 54 | 24 | 42 | 49 | 59 | 104 | 143 | 65 | 57.02 | 50 | 104 | 1.897 | 0.924 | 0.00260854 |
| AT3G16175.1 | | protein_coding | Thioesterase superfamily | 21 | 32 | 18 | 34 | 47 | 25 | 44 | 79 | 46 | 74 | 125 | 68 | 77.83 | 56 | 89 | 1.898 | 0.924 | 0.00021485 |
| AT4G06572.1 | | transposable_elem | transposable element ger | 343 | 183 | 195 | 863 | 546 | 560 | 721 | 450 | 499 | 1872 | 1450 | 1523 | 605.6 | 557 | 1615 | 1.896 | 0.923 | 1.45E-05 |
| AT5G08335.1 | ICMTB | protein_coding | Isoprenylcysteine carboxy | 78 | 61 | 71 | 90 | 72 | 78 | 164 | 150 | 182 | 195 | 191 | 212 | 168.49 | 165 | 199 | 1.896 | 0.923 | 4.80E-07 |
| AT4G15910.1 | ATDI21 | protein_coding | drought-induced 21 | 186 | 159 | 186 | 327 | 309 | 305 | 391 | 391 | 476 | 709 | 820 | 829 | 643.4 | 419 | 786 | 1.896 | 0.923 | 3.92E-12 |
| AT1G24909.1 | | protein_coding | Glutamine amidotransfer | 19 | 21 | 22 | 48 | 30 | 26 | 40 | 52 | 56 | 104 | 80 | 71 | 56.87 | 49 | 85 | 1.891 | 0.919 | 0.00108858 |
| AT3G51895.1 | SULTR3 | protein_coding | sulfate transporter 3;1 | 82 | 90 | 58 | 103 | 116 | 91 | 172 | 221 | 148 | 223 | 308 | 247 | 212.98 | 180 | 259 | 1.891 | 0.919 | 3.27E-08 |
| AT5G44780.1 | MORF4 | protein_coding | | 164 | 107 | 86 | 160 | 160 | 137 | 345 | 263 | 220 | 347 | 425 | 373 | 269.18 | 276 | 382 | 1.89 | 0.918 | 8.16E-08 |
| AT1G15125.1 | | protein_coding | S-adenosyl-L-methionine- | 52 | 43 | 22 | 136 | 27 | 42 | 109 | 106 | 56 | 295 | 72 | 114 | 288.47 | 90 | 160 | 1.888 | 0.917 | 0.00382083 |
| AT1G72520.1 | LOX4 | protein_coding | PLAT/LH2 domain-contai | 53 | 41 | 14 | 77 | 56 | 29 | 111 | 101 | 36 | 167 | 149 | 79 | 93.06 | 83 | 132 | 1.888 | 0.916 | 0.00253123 |
| AT3G16150.1 | | protein_coding | N-terminal nucleophile a | 10 | 11 | 11 | 31 | 17 | 27 | 21 | 27 | 28 | 67 | 45 | 73 | 44.39 | 25 | 62 | 1.886 | 0.915 | 0.00104911 |
| AT5G45475.1 | | other_rna | | 79 | 62 | 76 | 83 | 87 | 80 | 166 | 152 | 195 | 180 | 231 | 218 | 133.4 | 171 | 210 | 1.885 | 0.915 | 8.77E-07 |
| AT5G65850.1 | | protein_coding | F-box and associated inte | 63 | 28 | 28 | 61 | 54 | 45 | 132 | 69 | 72 | 132 | 143 | 122 | 99.01 | 91 | 132 | 1.884 | 0.914 | 6.74E-05 |
| AT2G41120.1 | | protein_coding | | 261 | 164 | 160 | 473 | 302 | 267 | 549 | 403 | 410 | 1026 | 802 | 726 | 627.06 | 454 | 851 | 1.884 | 0.914 | 1.78E-10 |
| AT5G01445.1 | | protein_coding | | 22 | 18 | 13 | 45 | 21 | 19 | 46 | 44 | 33 | 98 | 56 | 52 | 57.98 | 41 | 69 | 1.883 | 0.913 | 0.00247012 |
| AT4G23410.1 | TET5 | protein_coding | tetraspanin5 | 33 | 25 | 38 | 51 | 51 | 59 | 69 | 61 | 97 | 111 | 135 | 160 | 88.16 | 76 | 135 | 1.883 | 0.913 | 0.00012897 |
| AT4G02850.1 | | protein_coding | phenazine biosynthesis Pl | 275 | 347 | 180 | 533 | 717 | 441 | 578 | 853 | 461 | 1156 | 1903 | 1199 | 1220.87 | 631 | 1419 | 1.879 | 0.91 | 5.02E-06 |
| AT3G26460.1 | | protein_coding | Polyketide cyclase/dehyd | 16 | 10 | 9 | 58 | 25 | 26 | 34 | 25 | 23 | 126 | 66 | 71 | 58.84 | 27 | 88 | 1.873 | 0.905 | 0.00175829 |
| AT4G17098.1 | | other_rna | other RNA | 75 | 55 | 63 | 109 | 40 | 32 | 158 | 135 | 161 | 236 | 106 | 87 | 119.08 | 151 | 143 | 1.871 | 0.904 | 0.00170889 |
| AT3G45140.1 | LOX2 | protein_coding | lipoxygenase 2 | 2190 | 3085 | 1119 | 3790 | 4840 | 2472 | 4604 | 7579 | 2865 | 8221 | 12849 | 6722 | 7042.92 | 5016 | 9264 | 1.867 | 0.901 | 0.0005524 |
| AT5G39030.1 | | protein_coding | Protein kinase superfamil | 37 | 40 | 31 | 43 | 40 | 22 | 78 | 98 | 79 | 93 | 106 | 60 | 64.75 | 85 | 86 | 1.867 | 0.901 | 0.0001866 |
| AT5G13880.1 | | protein_coding | | 17 | 25 | 22 | 28 | 33 | 45 | 36 | 61 | 56 | 61 | 88 | 122 | 55.37 | 51 | 90 | 1.867 | 0.9 | 0.00294095 |
| AT3G21805.1 | | small_nucleolar_r | snoRNA | 609 | 433 | 525 | 489 | 384 | 292 | 1280 | 1064 | 1344 | 1061 | 1019 | 794 | 551.92 | 1229 | 958 | 1.863 | 0.898 | 0.00830334 |
| AT3G04000.1 | | protein_coding | NAD(P)-binding Rossman | 64 | 35 | 30 | 101 | 20 | 36 | 135 | 86 | 77 | 219 | 53 | 98 | 117.44 | 99 | 123 | 1.862 | 0.897 | 0.00672151 |
| AT3G05360.1 | AtRLP30 | protein_coding | receptor like protein 30 | 121 | 113 | 93 | 160 | 114 | 83 | 254 | 278 | 238 | 347 | 303 | 226 | 170.3 | 257 | 292 | 1.862 | 0.897 | 4.17E-06 |
| AT2G01860.1 | EMB975 | protein_coding | Tetratricopeptide repeat | 470 | 328 | 415 | 736 | 503 | 566 | 988 | 806 | 1062 | 1596 | 1335 | 1539 | 986.49 | 952 | 1490 | 1.861 | 0.896 | 2.18E-11 |
| AT5G01480.1 | | protein_coding | Cysteine/Histidine-rich C1 | 12 | 13 | 7 | 11 | 17 | 9 | 25 | 32 | 18 | 24 | 45 | 24 | 32.49 | 25 | 31 | 1.858 | 0.894 | 0.00115704 |
| AT5G17050.1 | UGT78D2 | protein_coding | UDP-glucosyl transferase | 188 | 146 | 103 | 392 | 211 | 224 | 395 | 359 | 264 | 850 | 560 | 609 | 590.41 | 339 | 673 | 1.859 | 0.894 | 1.08E-08 |
| AT2G41250.1 | | protein_coding | Haloacid dehalogenase-li | 228 | 199 | 166 | 395 | 310 | 292 | 479 | 489 | 425 | 857 | 823 | 794 | 527.49 | 464 | 825 | 1.857 | 0.893 | 2.26E-14 |
| AT5G04190.1 | PKS4 | protein_coding | phytochrome kinase subs | 87 | 89 | 136 | 131 | 147 | 250 | 183 | 219 | 348 | 284 | 390 | 680 | 241.08 | 250 | 451 | 1.855 | 0.892 | 0.00106378 |
| AT3G28160.1 | | transposable_elem | transposable element ger | 711 | 484 | 477 | 1208 | 766 | 814 | 1495 | 1189 | 1221 | 2620 | 2034 | 2214 | 1415.74 | 1302 | 2289 | 1.856 | 0.892 | 3.86E-10 |
| AT5G61280.1 | | protein_coding | Remorin family protein | 8 | 13 | 13 | 15 | 18 | 13 | 17 | 32 | 33 | 33 | 48 | 35 | 37.92 | 27 | 39 | 1.854 | 0.891 | 0.00257927 |
| AT2G31690.1 | | protein_coding | alpha/beta-Hydrolases su | 13 | 24 | 18 | 47 | 26 | 34 | 27 | 59 | 46 | 102 | 69 | 92 | 60.7 | 44 | 88 | 1.855 | 0.891 | 0.00097698 |
| AT5G16970.1 | P1 | protein_coding | alkenal reductase | 1577 | 907 | 717 | 2200 | 1022 | 937 | 3315 | 2228 | 1836 | 4772 | 2713 | 2548 | 3098.63 | 2460 | 3344 | 1.855 | 0.891 | 6.06E-05 |
| AT5G25130.1 | CYP71B12 | protein_coding | cytochrome P450, family | 95 | 107 | 69 | 190 | 182 | 84 | 200 | 263 | 177 | 412 | 483 | 228 | 226.18 | 213 | 374 | 1.852 | 0.889 | 0.00034745 |
| AT3G25020.1 | AtRLP42 | protein_coding | receptor like protein 42 | 38 | 54 | 56 | 75 | 71 | 70 | 80 | 133 | 143 | 163 | 188 | 190 | 150.31 | 119 | 180 | 1.852 | 0.889 | 1.41E-05 |
| AT3G56070.1 | CYP19-3 | protein_coding | rotamase cyclophilin 2 | 920 | 476 | 388 | 1204 | 855 | 674 | 1934 | 1169 | 993 | 2612 | 2270 | 1833 | 1885.31 | 1365 | 2238 | 1.852 | 0.889 | 6.25E-06 |
| AT3G20370.1 | | protein_coding | TRAF-like family protein | 3154 | 2308 | 1938 | 5844 | 3522 | 2785 | 6630 | 5670 | 4961 | 12676 | 9350 | 7574 | 8916.42 | 5754 | 9867 | 1.852 | 0.889 | 5.98E-08 |
| AT2G27740.1 | | protein_coding | Family of unknown functi | 20 | 23 | 20 | 33 | 36 | 41 | 42 | 57 | 51 | 72 | 96 | 111 | 87.22 | 50 | 93 | 1.847 | 0.885 | 2.50E-05 |
| AT1G15720.1 | TRFL5 | protein_coding | TRF-like 5 | 58 | 39 | 29 | 69 | 38 | 46 | 122 | 96 | 74 | 150 | 101 | 125 | 102.73 | 97 | 125 | 1.844 | 0.883 | 0.00042168 |
| AT3G26960.1 | | protein_coding | Pollen Ole e 1 allergen ar | 101 | 46 | 27 | 174 | 136 | 57 | 212 | 113 | 69 | 377 | 361 | 155 | 224.5 | 131 | 298 | 1.843 | 0.882 | 0.00288096 |
| AT1G77870.1 | MUB5 | protein_coding | membrane-anchored ubi | 25 | 29 | 47 | 51 | 37 | 59 | 53 | 71 | 120 | 111 | 98 | 160 | 88.7 | 81 | 123 | 1.84 | 0.88 | 0.0041501 |
| AT3G14770.1 | SWEET2 | protein_coding | Nodulin MtN3 family pro | 464 | 283 | 305 | 757 | 527 | 534 | 975 | 695 | 781 | 1642 | 1399 | 1452 | 1020 | 817 | 1498 | 1.84 | 0.88 | 3.68E-11 |
| AT4G21800.1 | QQT2 | protein_coding | P-loop containing nucleo | 426 | 294 | 299 | 843 | 654 | 588 | 895 | 722 | 765 | 1829 | 1736 | 1599 | 1176.35 | 794 | 1721 | 1.84 | 0.879 | 1.77E-13 |
| AT2G29090.1 | CYP707A2 | protein_coding | cytochrome P450, family | 180 | 63 | 62 | 271 | 128 | 107 | 378 | 155 | 159 | 588 | 340 | 291 | 286.46 | 231 | 406 | 1.837 | 0.878 | 0.00185255 |
| AT2G28460.1 | | protein_coding | Cysteine/Histidine-rich C1 | 48 | 40 | 34 | 54 | 59 | 45 | 101 | 98 | 87 | 117 | 157 | 122 | 83.94 | 95 | 132 | 1.833 | 0.874 | 4.79E-05 |
| AT1G19397.1 | | protein_coding | | 35 | 44 | 59 | 66 | 43 | 68 | 74 | 108 | 151 | 143 | 114 | 185 | 128.24 | 111 | 147 | 1.83 | 0.872 | 2.66E-05 |
| AT4G35090.1 | CAT2 | protein_coding | catalase 2 | 12315 | 8222 | 7208 | 22711 | 13127 | 12713 | 25887 | 20200 | 18453 | 49262 | 34849 | 34572 | 30183.08 | 21513 | 39561 | 1.83 | 0.871 | 5.62E-07 |
| AT1G52000.1 | JAL5 | protein_coding | Mannose-binding lectin s | 83 | 85 | 55 | 147 | 176 | 105 | 174 | 209 | 141 | 319 | 467 | 286 | 226.04 | 175 | 357 | 1.827 | 0.869 | 3.37E-06 |
| AT1G71528.1 | | other_rna | other RNA | 74 | 46 | 65 | 102 | 93 | 55 | 156 | 113 | 166 | 221 | 247 | 150 | 188.74 | 145 | 206 | 1.827 | 0.869 | 1.72E-06 |
| AT4G35800.1 | NRPB1 | protein_coding | RNA polymerase II large s | 3150 | 2625 | 2656 | 6801 | 5406 | 4999 | 6622 | 6449 | 6800 | 14752 | 14352 | 13594 | 7573.23 | 6624 | 14233 | 1.823 | 0.867 | 4.06E-14 |
| AT5G13370.1 | | protein_coding | Auxin-responsive GH3 fa | 370 | 304 | 201 | 614 | 323 | 356 | 778 | 747 | 515 | 1332 | 857 | 968 | 822.27 | 680 | 1052 | 1.821 | 0.865 | 2.51E-07 |
| AT1G68440.1 | | protein_coding | | 209 | 162 | 143 | 583 | 257 | 474 | 439 | 398 | 366 | 1265 | 682 | 1289 | 644.15 | 401 | 1079 | 1.819 | 0.863 | 1.31E-05 |
| AT5G45472.1 | | other_rna | | 184 | 173 | 191 | 271 | 224 | 217 | 387 | 425 | 489 | 588 | 595 | 590 | 364.79 | 434 | 591 | 1.818 | 0.862 | 1.68E-09 |
| AT2G02010.1 | GAD4 | protein_coding | glutamate decarboxylase | 49 | 44 | 47 | 103 | 55 | 59 | 103 | 108 | 120 | 223 | 146 | 160 | 110.41 | 110 | 176 | 1.816 | 0.861 | 0.00090062 |
| AT1G53700.1 | WAG1 | protein_coding | WAG 1 | 21 | 19 | 7 | 36 | 22 | 13 | 44 | 47 | 18 | 78 | 58 | 35 | 47.74 | 36 | 57 | 1.815 | 0.86 | 0.00970457 |
| AT3G07000.1 | | protein_coding | Cysteine/Histidine-rich C1 | 41 | 18 | 33 | 71 | 36 | 43 | 86 | 44 | 84 | 154 | 96 | 117 | 81.37 | 71 | 122 | 1.812 | 0.858 | 0.00026465 |
| AT5G64950.1 | | protein_coding | Mitochondrial transcripti | 47 | 41 | 32 | 78 | 52 | 57 | 99 | 101 | 82 | 169 | 138 | 155 | 109.47 | 94 | 154 | 1.813 | 0.858 | 0.00015651 |
| AT1G73870.1 | COL7 | protein_coding | B-box type zinc finger pro | 107 | 104 | 97 | 189 | 163 | 184 | 225 | 256 | 248 | 410 | 433 | 500 | 396.53 | 243 | 448 | 1.813 | 0.858 | 6.07E-10 |
| AT1G67520.1 | | protein_coding | lectin protein kinase fami | 28 | 28 | 32 | 52 | 38 | 35 | 59 | 69 | 82 | 113 | 101 | 95 | 75.33 | 70 | 103 | 1.812 | 0.857 | 0.00430131 |
| AT3G10050.1 | OMR1 | protein_coding | L-O-methylthreonine resi | 1008 | 643 | 522 | 1505 | 1021 | 1026 | 2119 | 1580 | 1336 | 3264 | 2711 | 2790 | 2360.95 | 1678 | 2922 | 1.812 | 0.857 | 2.97E-08 |
| AT5G05060.1 | | protein_coding | Cystatin/monellin superfa | 456 | 332 | 328 | 597 | 496 | 502 | 959 | 816 | 840 | 1295 | 1317 | 1365 | 1241.12 | 872 | 1326 | 1.811 | 0.857 | 7.92E-13 |
| AT3G59140.1 | ABCC10 | protein_coding | multidrug resistance-asso | 480 | 311 | 176 | 866 | 518 | 228 | 1009 | 764 | 451 | 1878 | 1375 | 620 | 813.45 | 741 | 1291 | 1.808 | 0.855 | 0.00291657 |
| AT5G66650.1 | | protein_coding | Protein of unknown funct | 46 | 50 | 52 | 58 | 40 | 81 | 97 | 123 | 133 | 126 | 106 | 220 | 106.06 | 118 | 151 | 1.804 | 0.851 | 0.00053731 |
| AT1G71890.1 | SUC5 | protein_coding | Major facilitator superfar | 132 | 84 | 45 | 243 | 105 | 86 | 277 | 206 | 115 | 527 | 279 | 234 | 200.07 | 199 | 347 | 1.803 | 0.85 | 0.00310187 |
| AT2G19650.1 | | protein_coding | Cysteine/Histidine-rich C1 | 222 | 140 | 111 | 329 | 226 | 209 | 467 | 344 | 284 | 714 | 600 | 568 | 397.42 | 365 | 627 | 1.802 | 0.85 | 1.90E-05 |
| AT3G06437.1 | | pseudogene | | 13 | 13 | 9 | 26 | 12 | 17 | 27 | 32 | 23 | 56 | 32 | 46 | 73.89 | 27 | 45 | 1.801 | 0.849 | 0.00397066 |
| AT1G79470.1 | IMPDH | protein_coding | Aldolase-type TIM barrel | 141 | 113 | 100 | 221 | 167 | 160 | 296 | 278 | 256 | 479 | 443 | 435 | 363.51 | 277 | 452 | 1.799 | 0.847 | 4.18E-08 |
| AT3G55240.1 | | protein_coding | Plant protein 1589 of unk | 650 | 331 | 365 | 1614 | 1337 | 1575 | 1366 | 813 | 934 | 3501 | 3549 | 4283 | 1775.58 | 1038 | 3778 | 1.796 | 0.845 | 2.80E-07 |
| AT3G19450.1 | CAD4 | protein_coding | GroES-like zinc-binding al | 1090 | 895 | 580 | 2085 | 1528 | 1242 | 2291 | 2199 | 1485 | 4523 | 4056 | 3378 | 2991.74 | 1992 | 3986 | 1.797 | 0.845 | 4.14E-08 |
| AT1G79245.1 | | pseudogene | | 3753 | 2666 | 2883 | 6826 | 5110 | 4180 | 7889 | 6550 | 7381 | 14806 | 13566 | 11367 | 11987 | 7273 | 13246 | 1.795 | 0.844 | 4.12E-09 |
| AT2G33060.1 | AtRLP27 | protein_coding | receptor like protein 27 | 39 | 43 | 58 | 54 | 73 | 62 | 82 | 106 | 148 | 117 | 194 | 169 | 110.99 | 112 | 160 | 1.794 | 0.843 | 0.00028523 |
| AT5G52810.1 | | protein_coding | NAD(P)-binding Rossman | 108 | 80 | 73 | 201 | 82 | 61 | 227 | 197 | 187 | 436 | 218 | 166 | 237.11 | 204 | 273 | 1.79 | 0.84 | 0.00045362 |
| AT5G44060.1 | | protein_coding | | 59 | 81 | 61 | 185 | 153 | 158 | 124 | 199 | 156 | 401 | 406 | 430 | 440.71 | 160 | 412 | 1.79 | 0.84 | 4.65E-10 |
| AT1G08830.1 | CSD1 | protein_coding | copper/zinc superoxide d | 210 | 458 | 275 | 295 | 428 | 477 | 441 | 1125 | 704 | 640 | 1136 | 1297 | 1192.42 | 757 | 1024 | 1.789 | 0.839 | 0.00233254 |
| AT5G10920.1 | T30N20_190 | protein_coding | L-Aspartase-like family pr | 1212 | 820 | 608 | 1403 | 996 | 865 | 2548 | 2015 | 1557 | 3043 | 2644 | 2352 | 2227.19 | 2040 | 2680 | 1.789 | 0.839 | 1.24E-07 |
| AT1G05385.1 | PSB27-2 | protein_coding | photosystem II 11 kDa pr | 485 | 403 | 361 | 664 | 581 | 444 | 1020 | 990 | 924 | 1440 | 1542 | 1207 | 1275.63 | 978 | 1396 | 1.789 | 0.839 | 1.11E-11 |
| AT5G19875.1 | | protein_coding | | 3 | 2 | 4 | 7 | 6 | 7 | 6 | 5 | 10 | 15 | 16 | 19 | 26.78 | 7 | 17 | 1.787 | 0.837 | 0.00207008 |
| AT4G39830.1 | | protein_coding | Cupredoxin superfamily p | 18 | 14 | 6 | 33 | 17 | 23 | 38 | 34 | 15 | 72 | 45 | 63 | 49.88 | 29 | 60 | 1.786 | 0.837 | 0.00125151 |
| AT3G08690.1 | UBC11 | protein_coding | ubiquitin-conjugating enz | 189 | 166 | 134 | 295 | 244 | 196 | 397 | 408 | 343 | 640 | 648 | 533 | 437.37 | 383 | 607 | 1.785 | 0.836 | 1.32E-11 |
| AT4G39810.1 | NEN4 | protein_coding | Polynucleotidyl transfera | 18 | 14 | 13 | 22 | 16 | 17 | 38 | 34 | 33 | 48 | 42 | 46 | 41.5 | 35 | 45 | 1.784 | 0.835 | 0.0007277 |
| AT2G30140.1 | UGT87A2 | protein_coding | UDP-Glycosyltransferase | 115 | 94 | 64 | 217 | 86 | 97 | 242 | 231 | 164 | 471 | 228 | 264 | 321.24 | 212 | 321 | 1.784 | 0.835 | 4.74E-05 |
| AT1G77330.1 | | protein_coding | 2-oxoglutarate (2OG) and | 30 | 29 | 31 | 44 | 38 | 31 | 63 | 71 | 79 | 95 | 101 | 84 | 67.58 | 71 | 93 | 1.779 | 0.831 | 0.0002204 |
| AT1G23310.1 | GGAT1 | protein_coding | glutamate:glyoxylate ami | 27748 | 24490 | 25179 | 46978 | 34906 | 36868 | 58328 | 60167 | 64460 | 101900 | 92667 | 100260 | 79281.83 | 60985 | 98276 | 1.779 | 0.831 | 2.80E-07 |
| AT1G26560.1 | BGLU40 | protein_coding | beta glucosidase 40 | 314 | 271 | 188 | 523 | 408 | 232 | 660 | 666 | 481 | 1134 | 1083 | 631 | 643.69 | 602 | 949 | 1.776 | 0.829 | 4.57E-06 |
| AT3G51240.1 | F3H | protein_coding | flavanone 3-hydroxylase | 186 | 107 | 65 | 283 | 190 | 160 | 391 | 263 | 166 | 614 | 504 | 435 | 379.93 | 273 | 518 | 1.775 | 0.828 | 0.00018587 |
| AT3G02380.1 | COL2 | protein_coding | CONSTANS-like 2 | 278 | 220 | 141 | 430 | 334 | 245 | 584 | 540 | 361 | 933 | 887 | 666 | 531.48 | 495 | 829 | 1.775 | 0.828 | 1.00E-06 |
| AT2G46790.1 | APRR9 | protein_coding | pseudo-response regulat | 430 | 382 | 376 | 545 | 558 | 589 | 904 | 939 | 963 | 1182 | 1481 | 1602 | 877.22 | 935 | 1422 | 1.775 | 0.828 | 5.94E-07 |
| AT3G30875.1 | | pseudogene | | 36 | 21 | 15 | 47 | 44 | 26 | 76 | 52 | 38 | 102 | 117 | 71 | 57.52 | 55 | 97 | 1.774 | 0.827 | 0.00459508 |
| AT3G01440.1 | PNSL3 | protein_coding | PsbQ-like 1 | 679 | 581 | 413 | 1036 | 876 | 661 | 1427 | 1427 | 1057 | 2247 | 2326 | 1798 | 2018.98 | 1304 | 2124 | 1.773 | 0.827 | 3.26E-09 |
| AT2G41630.1 | TFIIB1 | protein_coding | transcription factor IIB | 635 | 487 | 552 | 1257 | 946 | 963 | 1335 | 1196 | 1413 | 2727 | 2511 | 2619 | 1780.71 | 1315 | 2619 | 1.774 | 0.827 | 1.25E-12 |
| AT1G07180.1 | NDA1 | protein_coding | alternative NAD(P)H deh | 898 | 821 | 741 | 1435 | 1322 | 1345 | 1888 | 2017 | 1897 | 3113 | 3510 | 3658 | 2380.45 | 1934 | 3427 | 1.772 | 0.826 | 4.54E-13 |
| AT4G09012.1 | | protein_coding | Mitochondrial ribosomal | 21 | 11 | 18 | 34 | 20 | 25 | 44 | 27 | 46 | 74 | 53 | 68 | 58.51 | 39 | 65 | 1.771 | 0.824 | 0.00429477 |
| AT5G41360.1 | XPB2 | protein_coding | homolog of Xeroderma p | 336 | 234 | 254 | 540 | 419 | 363 | 706 | 575 | 650 | 1171 | 1112 | 987 | 712.9 | 644 | 1090 | 1.771 | 0.824 | 2.81E-13 |
| AT5G63370.1 | CDKG1 | protein_coding | Protein kinase superfamil | 1191 | 1042 | 1229 | 1997 | 1737 | 1884 | 2504 | 2560 | 3146 | 4332 | 4611 | 5123 | 3179.76 | 2737 | 4689 | 1.769 | 0.823 | 1.42E-10 |
| AT5G14840.1 | | pseudogene | | 198 | 172 | 160 | 192 | 177 | 211 | 416 | 423 | 410 | 416 | 470 | 574 | 725.78 | 416 | 487 | 1.767 | 0.822 | 7.94E-11 |
| AT5G55950.1 | | protein_coding | Nucleotide/sugar transpo | 11 | 16 | 11 | 22 | 24 | 23 | 23 | 39 | 28 | 48 | 64 | 63 | 46.71 | 30 | 58 | 1.765 | 0.82 | 0.00398702 |
| AT2G41290.1 | SSL2 | protein_coding | strictosidine synthase-lik | 53 | 47 | 33 | 57 | 63 | 45 | 111 | 115 | 84 | 124 | 167 | 122 | 126.98 | 103 | 138 | 1.764 | 0.819 | 8.25E-05 |
| AT2G29440.1 | GSTU6 | protein_coding | glutathione S-transferase | 29 | 24 | 26 | 51 | 27 | 31 | 61 | 59 | 67 | 111 | 72 | 84 | 61.85 | 62 | 89 | 1.762 | 0.817 | 0.00127756 |
| AT4G04220.1 | AtRLP46 | protein_coding | receptor like protein 46 | 30 | 38 | 38 | 67 | 64 | 58 | 63 | 93 | 97 | 145 | 170 | 158 | 95.83 | 84 | 158 | 1.762 | 0.817 | 2.04E-05 |
| AT1G06490.1 | CALS7 | protein_coding | glucan synthase-like 7 | 151 | 155 | 150 | 221 | 228 | 217 | 317 | 381 | 384 | 479 | 605 | 590 | 320.9 | 361 | 558 | 1.762 | 0.817 | 7.51E-09 |
| AT2G37240.1 | | protein_coding | Thioredoxin superfamily | 502 | 412 | 305 | 791 | 655 | 566 | 1055 | 1012 | 781 | 1716 | 1739 | 1539 | 1313.38 | 949 | 1665 | 1.757 | 0.813 | 7.77E-10 |
| AT1G06570.1 | PDS1 | protein_coding | phytoene desaturation 1 | 388 | 360 | 581 | 722 | 484 | 930 | 816 | 884 | 1487 | 1566 | 1285 | 2529 | 1260.24 | 1062 | 1793 | 1.756 | 0.812 | 0.00026047 |
| AT2G42650.1 | | protein_coding | Ribosomal protein L1p/L1 | 116 | 76 | 74 | 144 | 100 | 78 | 244 | 187 | 189 | 312 | 265 | 212 | 225.05 | 207 | 263 | 1.755 | 0.811 | 3.37E-07 |
| AT2G17787.1 | | protein_coding | | 105 | 92 | 90 | 195 | 172 | 193 | 221 | 226 | 230 | 423 | 457 | 525 | 308.15 | 226 | 468 | 1.754 | 0.811 | 2.87E-08 |
| AT2G20250.1 | | protein_coding | | 163 | 189 | 135 | 210 | 226 | 151 | 343 | 464 | 346 | 456 | 600 | 411 | 374.12 | 384 | 489 | 1.753 | 0.81 | 3.72E-07 |
| AT1G04220.1 | KCS17 | protein_coding | 3-ketoacyl-CoA synthase | 29 | 26 | 6 | 53 | 74 | 40 | 61 | 64 | 15 | 115 | 196 | 109 | 92.45 | 47 | 140 | 1.753 | 0.809 | 0.00372718 |
| AT2G22100.1 | UBA1B | protein_coding | RNA-binding (RRM/RBD/ | 172 | 111 | 107 | 165 | 153 | 115 | 362 | 273 | 274 | 358 | 406 | 313 | 315.11 | 303 | 359 | 1.752 | 0.809 | 2.44E-08 |
| AT5G24160.1 | SQE6 | protein_coding | squalene monoxygenase | 76 | 38 | 20 | 164 | 96 | 57 | 160 | 93 | 51 | 356 | 255 | 155 | 185.84 | 101 | 255 | 1.751 | 0.808 | 0.00506683 |
| AT5G18770.1 | | protein_coding | F-box/FBD-like domains c | 106 | 93 | 97 | 179 | 149 | 127 | 223 | 228 | 248 | 388 | 396 | 345 | 289.76 | 233 | 376 | 1.751 | 0.808 | 5.92E-09 |
| AT3G30122.1 | | pseudogene | | 69 | 49 | 42 | 51 | 36 | 41 | 145 | 120 | 108 | 111 | 96 | 111 | 102.28 | 124 | 106 | 1.75 | 0.807 | 0.00023565 |
| AT1G68870.1 | ATSOFL2 | protein_coding | SOB five-like 2 | 84 | 63 | 62 | 176 | 128 | 77 | 177 | 155 | 159 | 382 | 340 | 209 | 257.29 | 164 | 310 | 1.749 | 0.807 | 5.58E-06 |
| AT3G05327.1 | | protein_coding | Cyclin family protein | 13 | 11 | 7 | 26 | 14 | 15 | 27 | 27 | 18 | 56 | 37 | 41 | 40.1 | 24 | 45 | 1.748 | 0.806 | 0.00133734 |
| AT5G48775.1 | | other_rna | other RNA | 72 | 37 | 49 | 85 | 74 | 76 | 151 | 91 | 125 | 184 | 196 | 207 | 134.33 | 122 | 196 | 1.748 | 0.806 | 1.20E-05 |
| AT3G05660.1 | AtRLP33 | protein_coding | receptor like protein 33 | 51 | 17 | 32 | 48 | 24 | 39 | 107 | 42 | 82 | 104 | 64 | 106 | 85.07 | 77 | 91 | 1.746 | 0.804 | 0.00543327 |
| AT5G49910.1 | HSP70-7 | protein_coding | chloroplast heat shock pr | 10508 | 6566 | 4072 | 9817 | 7409 | 5371 | 22089 | 16131 | 10425 | 21294 | 19669 | 14606 | 17923.26 | 16215 | 18523 | 1.746 | 0.804 | 0.00015415 |
| AT3G25760.1 | AOC1 | protein_coding | allene oxide cyclase 1 | 71 | 80 | 61 | 169 | 151 | 106 | 149 | 197 | 156 | 367 | 401 | 288 | 404.61 | 167 | 352 | 1.746 | 0.804 | 1.43E-09 |
| AT3G58200.1 | | protein_coding | TRAF-like family protein | 31 | 22 | 18 | 51 | 35 | 30 | 65 | 54 | 46 | 111 | 93 | 82 | 63.24 | 55 | 95 | 1.745 | 0.803 | 0.00287066 |
| AT4G38960.3 | BBX19 | protein_coding | B-box type zinc finger fan | 80 | 81 | 53 | 136 | 99 | 82 | 168 | 199 | 136 | 295 | 263 | 223 | 194.9 | 168 | 260 | 1.745 | 0.803 | 2.03E-05 |
| AT1G31860.1 | HISN2 | protein_coding | histidine biosynthesis bif | 563 | 358 | 304 | 686 | 556 | 452 | 1183 | 880 | 778 | 1488 | 1476 | 1229 | 1207.39 | 947 | 1398 | 1.745 | 0.803 | 4.39E-08 |
| AT2G32480.1 | ARASP | protein_coding | ARABIDOPSIS SERIN PRO | 1736 | 1375 | 1224 | 2455 | 1845 | 1432 | 3649 | 3378 | 3134 | 5325 | 4898 | 3894 | 4691.24 | 3387 | 4706 | 1.743 | 0.802 | 4.45E-10 |
| AT3G01890.1 | | protein_coding | SWIB/MDM2 domain sup | 145 | 130 | 110 | 253 | 188 | 204 | 305 | 319 | 282 | 549 | 499 | 555 | 418.52 | 302 | 534 | 1.744 | 0.802 | 1.81E-10 |
| AT5G03630.1 | ATMDAR2 | protein_coding | Pyridine nucleotide-disul | 972 | 755 | 591 | 1554 | 865 | 857 | 2043 | 1855 | 1513 | 3371 | 2296 | 2331 | 2674.56 | 1804 | 2666 | 1.742 | 0.801 | 2.74E-07 |
| AT3G14900.1 | | protein_coding | | 35 | 22 | 25 | 51 | 30 | 25 | 74 | 54 | 64 | 111 | 80 | 68 | 420.97 | 64 | 86 | 1.743 | 0.801 | 2.23E-07 |
| AT2G13130.1 | | pseudogene | | 24 | 31 | 31 | 46 | 32 | 42 | 50 | 76 | 79 | 100 | 85 | 114 | 71.04 | 68 | 100 | 1.74 | 0.799 | 0.00125542 |
| AT1G74055.1 | | protein_coding | | 8 | 19 | 17 | 25 | 30 | 45 | 17 | 47 | 44 | 54 | 80 | 122 | 65.76 | 36 | 85 | 1.738 | 0.798 | 0.00459085 |
| AT2G43820.1 | UGT74F2 | protein_coding | UDP-glucosyltransferase | 527 | 346 | 349 | 752 | 304 | 458 | 1108 | 850 | 893 | 1631 | 807 | 1246 | 977.93 | 950 | 1228 | 1.739 | 0.798 | 4.53E-05 |
| AT5G10970.1 | | protein_coding | C2H2 and C2HC zinc finge | 30 | 23 | 25 | 47 | 48 | 44 | 63 | 57 | 64 | 102 | 127 | 120 | 93.5 | 61 | 116 | 1.738 | 0.797 | 0.0001267 |
| AT2G41040.1 | | protein_coding | S-adenosyl-L-methionine- | 133 | 123 | 113 | 262 | 180 | 185 | 280 | 302 | 289 | 568 | 478 | 503 | 450.45 | 290 | 516 | 1.736 | 0.796 | 1.58E-11 |
| AT3G14420.1 | GLO1 | protein_coding | Aldolase-type TIM barrel | 15434 | 14677 | 12353 | 23879 | 19577 | 20971 | 32443 | 36059 | 31624 | 51796 | 51972 | 57029 | 51500.29 | 33375 | 53599 | 1.734 | 0.795 | 1.51E-07 |
| AT2G31380.1 | BBX25 | protein_coding | salt tolerance homologue | 429 | 368 | 325 | 851 | 639 | 659 | 902 | 904 | 832 | 1846 | 1696 | 1792 | 1191.02 | 879 | 1778 | 1.736 | 0.795 | 4.60E-09 |
| AT3G12510.1 | | protein_coding | MADS-box family protein | 15 | 26 | 17 | 31 | 40 | 38 | 32 | 64 | 44 | 67 | 106 | 103 | 79.01 | 47 | 92 | 1.734 | 0.794 | 0.002406 |
| AT4G24570.1 | PUMP4 | protein_coding | dicarboxylate carrier 2 | 43 | 28 | 21 | 59 | 38 | 49 | 90 | 69 | 54 | 128 | 101 | 133 | 129.35 | 71 | 121 | 1.734 | 0.794 | 0.00079465 |
| AT1G68600.1 | ALMT5 | protein_coding | Aluminium activated mal | 128 | 138 | 116 | 143 | 140 | 89 | 269 | 339 | 297 | 310 | 372 | 242 | 401.94 | 302 | 308 | 1.732 | 0.793 | 3.30E-06 |
| AT5G64840.1 | ABCF5 | protein_coding | general control non-repre | 5524 | 5007 | 5311 | 9129 | 7790 | 7797 | 11612 | 12301 | 13596 | 19802 | 20681 | 21203 | 13678.45 | 12503 | 20562 | 1.732 | 0.792 | 3.54E-10 |
| AT5G14830.1 | | transposable_elem | transposable element ger | 41 | 50 | 32 | 36 | 45 | 50 | 86 | 123 | 82 | 78 | 119 | 136 | 84.81 | 97 | 111 | 1.73 | 0.791 | 0.00084514 |
| AT1G53160.2 | SPL4 | protein_coding | squamosa promoter bind | 35 | 19 | 32 | 33 | 32 | 26 | 74 | 47 | 82 | 72 | 85 | 71 | 68.79 | 68 | 76 | 1.729 | 0.79 | 0.00204822 |

| | | | | | | | | | | | | | | | | | | | | | |
|---|---|---|---|---|---|---|---|---|---|---|---|---|---|---|---|---|---|---|---|---|---|
| AT2G35130.2 | | protein_coding | Tetratricopeptide repeat | 589 | 514 | 375 | 874 | 731 | 510 | 1238 | 1263 | 960 | 1896 | 1941 | 1387 | 1418.58 | 1154 | 1741 | 1.73 | 0.79 | 7.45E-08 |
| AT1G47580.1 | DYW1 | protein_coding | Pentatricopeptide repeat | 326 | 295 | 290 | 603 | 432 | 334 | 685 | 725 | 742 | 1308 | 1147 | 908 | 836.82 | 717 | 1121 | 1.729 | 0.79 | 1.59E-09 |
| AT3G07720.1 | | protein_coding | Galactose oxidase/kelch r | 150 | 154 | 116 | 261 | 204 | 199 | 315 | 378 | 297 | 566 | 542 | 541 | 502.23 | 330 | 550 | 1.729 | 0.79 | 4.74E-13 |
| AT4G19520.1 | | protein_coding | disease resistance protei | 542 | 430 | 271 | 643 | 529 | 285 | 1139 | 1056 | 694 | 1395 | 1404 | 775 | 820.85 | 963 | 1191 | 1.724 | 0.785 | 8.14E-05 |
| AT3G51450.1 | SSL7 | protein_coding | Calcium-dependent phos | 51 | 46 | 31 | 93 | 73 | 48 | 107 | 113 | 79 | 202 | 194 | 131 | 171.96 | 100 | 176 | 1.723 | 0.785 | 4.28E-06 |
| AT5G27030.2 | TPR3 | protein_coding | TOPLESS-related 3 | 318 | 299 | 246 | 433 | 365 | 316 | 668 | 735 | 630 | 939 | 969 | 859 | 709.33 | 678 | 922 | 1.722 | 0.784 | 3.18E-13 |
| AT3G42806.1 | | transposable_elen | transposable element ger | 467 | 196 | 190 | 299 | 119 | 159 | 982 | 482 | 486 | 649 | 316 | 432 | 344.91 | 650 | 466 | 1.721 | 0.783 | 0.00325887 |
| AT3G09600.1 | RVE8 | protein_coding | Homeodomain-like super | 96 | 54 | 35 | 183 | 141 | 123 | 202 | 133 | 90 | 397 | 374 | 334 | 200.26 | 142 | 368 | 1.721 | 0.783 | 0.0001314 |
| AT2G36307.1 | ZPR4 | other_rna | ZPR4 (LITTLE ZIPPER 4); o | 24 | 16 | 26 | 36 | 42 | 32 | 50 | 39 | 67 | 78 | 112 | 87 | 58.02 | 52 | 92 | 1.719 | 0.782 | 0.0045451 |
| AT3G27200.1 | | protein_coding | Cupredoxin superfamily p | 25 | 49 | 28 | 74 | 64 | 81 | 53 | 120 | 72 | 161 | 170 | 220 | 172.06 | 82 | 184 | 1.719 | 0.782 | 3.39E-05 |
| AT2G23100.1 | | protein_coding | Cysteine/Histidine-rich C1 | 157 | 95 | 112 | 149 | 119 | 132 | 330 | 233 | 287 | 323 | 316 | 359 | 263.02 | 283 | 333 | 1.719 | 0.782 | 9.27E-07 |
| AT1G26210.1 | ATSOFL1 | protein_coding | SOB five-like 1 | 385 | 233 | 187 | 578 | 383 | 265 | 809 | 572 | 479 | 1254 | 1017 | 721 | 948.01 | 620 | 997 | 1.715 | 0.778 | 6.23E-05 |
| AT5G48930.1 | HST | protein_coding | hydroxycinnamoyl-CoA sh | 1959 | 2089 | 1369 | 2850 | 2553 | 1910 | 4118 | 5132 | 3505 | 6182 | 6778 | 5194 | 6064.26 | 4252 | 6051 | 1.715 | 0.778 | 2.71E-07 |
| AT4G05440.1 | EDA35 | protein_coding | temperature sensing prot | 221 | 139 | 116 | 299 | 167 | 151 | 465 | 341 | 297 | 649 | 443 | 411 | 419.97 | 368 | 501 | 1.714 | 0.777 | 3.33E-06 |
| AT1G60960.1 | IRT3 | protein_coding | iron regulated transporte | 178 | 80 | 83 | 293 | 138 | 190 | 374 | 197 | 212 | 636 | 366 | 517 | 347.56 | 261 | 506 | 1.712 | 0.776 | 0.00087647 |
| AT5G18370.1 | | protein_coding | Disease resistance protei | 111 | 86 | 82 | 109 | 83 | 95 | 233 | 211 | 210 | 236 | 220 | 258 | 152.84 | 218 | 238 | 1.712 | 0.775 | 4.10E-05 |
| AT2G01008.1 | | protein_coding | | 7835 | 12170 | 4701 | 11973 | 9383 | 16154 | 16470 | 29899 | 12035 | 25971 | 24910 | 43930 | 12834.68 | 19468 | 31604 | 1.711 | 0.774 | 0.00812745 |
| AT4G27030.1 | FAD4 | protein_coding | fatty acid desaturase A | 437 | 370 | 222 | 876 | 648 | 689 | 919 | 909 | 568 | 1900 | 1720 | 1874 | 1373.46 | 799 | 1831 | 1.71 | 0.774 | 3.99E-07 |
| AT4G17470.1 | | protein_coding | alpha/beta-Hydrolases su | 51 | 24 | 13 | 98 | 63 | 33 | 107 | 59 | 33 | 213 | 167 | 90 | 146.92 | 66 | 157 | 1.708 | 0.773 | 0.00818478 |
| AT1G06620.1 | | protein_coding | 2-oxoglutarate (2OG) anc | 37 | 21 | 15 | 91 | 58 | 32 | 78 | 52 | 38 | 197 | 154 | 87 | 102.96 | 56 | 146 | 1.709 | 0.773 | 0.00288758 |
| AT4G33050.2 | EDA39 | protein_coding | calmodulin-binding famil | 192 | 94 | 92 | 261 | 114 | 131 | 404 | 231 | 236 | 566 | 303 | 356 | 293.3 | 290 | 408 | 1.709 | 0.773 | 0.00078066 |
| AT2G35075.1 | | protein_coding | | 63 | 29 | 35 | 44 | 41 | 24 | 132 | 71 | 90 | 95 | 109 | 65 | 102.06 | 98 | 90 | 1.707 | 0.772 | 0.000202 |
| AT1G72680.1 | CAD1 | protein_coding | cinnamyl-alcohol dehydrc | 244 | 128 | 174 | 301 | 134 | 172 | 513 | 314 | 445 | 653 | 356 | 468 | 391.77 | 424 | 492 | 1.707 | 0.772 | 9.26E-05 |
| AT3G53230.1 | CDC48D | protein_coding | ATPase, AAA-type, CDC48 | 114 | 78 | 50 | 198 | 173 | 133 | 240 | 192 | 128 | 429 | 459 | 362 | 247.63 | 187 | 417 | 1.707 | 0.772 | 5.11E-05 |
| AT5G64940.1 | ATATH13 | protein_coding | ABC2 homolog 13 | 7312 | 4997 | 3897 | 10242 | 8110 | 5921 | 15370 | 12277 | 9977 | 22216 | 21530 | 16102 | 13666.87 | 12541 | 19949 | 1.708 | 0.772 | 2.15E-06 |
| AT5G18790.1 | | protein_coding | Ribosomal protein L33 fa | 158 | 105 | 81 | 152 | 113 | 116 | 332 | 258 | 207 | 330 | 300 | 315 | 301.1 | 266 | 315 | 1.708 | 0.772 | 9.22E-07 |
| AT3G44205.1 | | transposable_elen | transposable element ger | 46 | 38 | 54 | 59 | 46 | 39 | 97 | 93 | 138 | 128 | 122 | 106 | 93.36 | 109 | 119 | 1.707 | 0.771 | 0.00188183 |
| AT3G55110.1 | ABCG18 | protein_coding | ABC-2 type transporter fa | 126 | 111 | 99 | 196 | 197 | 132 | 265 | 273 | 253 | 425 | 523 | 359 | 259.99 | 264 | 436 | 1.706 | 0.771 | 1.98E-06 |
| AT4G28680.2 | TYRDC | protein_coding | L-tyrosine decarboxylase | 80 | 51 | 28 | 74 | 80 | 46 | 168 | 125 | 72 | 161 | 212 | 125 | 133.54 | 122 | 166 | 1.705 | 0.77 | 0.00070552 |
| AT4G35030.3 | | protein_coding | Protein kinase superfamil | 67 | 61 | 45 | 61 | 86 | 64 | 141 | 150 | 115 | 132 | 228 | 174 | 133.65 | 135 | 178 | 1.705 | 0.77 | 0.00010477 |
| AT5G56010.1 | HSP90-3 | protein_coding | heat shock protein 81-3 | 7901 | 5972 | 3826 | 8580 | 7092 | 4900 | 16608 | 14672 | 9795 | 18611 | 18828 | 13325 | 12799.75 | 13692 | 16921 | 1.705 | 0.77 | 1.69E-05 |
| AT4G34590.1 | ATB2 | protein_coding | G-box binding factor 6 | 104 | 89 | 41 | 163 | 107 | 79 | 219 | 219 | 105 | 354 | 284 | 215 | 365.27 | 181 | 284 | 1.704 | 0.769 | 3.52E-05 |
| AT1G52270.1 | | protein_coding | | 25 | 28 | 25 | 66 | 45 | 38 | 53 | 69 | 64 | 143 | 119 | 103 | 134.99 | 62 | 122 | 1.7 | 0.766 | 2.51E-06 |
| AT5G35490.1 | ATMRU1 | protein_coding | mto 1 responding up 1 | 100 | 90 | 79 | 116 | 67 | 50 | 210 | 221 | 202 | 252 | 178 | 136 | 210.65 | 211 | 189 | 1.699 | 0.765 | 0.00014183 |
| AT3G16410.1 | NSP4 | protein_coding | nitrile specifier protein 4 | 667 | 640 | 398 | 1283 | 887 | 1028 | 1402 | 1572 | 1019 | 2783 | 2355 | 2796 | 1602.87 | 1331 | 2645 | 1.7 | 0.765 | 3.42E-06 |
| AT3G08590.1 | | protein_coding | Phosphoglycerate mutase | 1969 | 1463 | 996 | 2822 | 1898 | 1721 | 4139 | 3594 | 2550 | 6121 | 5039 | 4680 | 4025.23 | 3428 | 5280 | 1.699 | 0.764 | 1.04E-06 |
| AT1G09830.1 | PUR2 | protein_coding | Glycinamide ribonucleoti | 568 | 458 | 323 | 713 | 561 | 540 | 1194 | 1125 | 827 | 1547 | 1489 | 1468 | 1505.38 | 1049 | 1501 | 1.698 | 0.764 | 8.95E-08 |
| AT2G35340.1 | MEE29 | protein_coding | helicase domain-containi | 82 | 82 | 110 | 124 | 119 | 102 | 172 | 201 | 282 | 269 | 316 | 277 | 223.72 | 218 | 287 | 1.697 | 0.763 | 1.69E-05 |
| AT3G16400.1 | NSP1 | protein_coding | nitrile specifier protein 1 | 3408 | 3021 | 2179 | 7121 | 4894 | 5162 | 7164 | 7422 | 5578 | 15446 | 12992 | 14038 | 11610.81 | 6721 | 14159 | 1.697 | 0.763 | 4.47E-08 |
| AT2G46670.1 | | protein_coding | CCT motif family protein | 85 | 59 | 63 | 79 | 104 | 126 | 179 | 145 | 161 | 171 | 276 | 343 | 164.78 | 162 | 263 | 1.693 | 0.76 | 0.00080275 |
| AT5G42460.1 | | protein_coding | F-box and associated inte | 87 | 98 | 94 | 179 | 153 | 167 | 183 | 241 | 241 | 388 | 406 | 454 | 255.57 | 222 | 416 | 1.693 | 0.759 | 2.23E-07 |
| AT5G01190.1 | LAC10 | protein_coding | laccase 10 | 20 | 20 | 16 | 28 | 23 | 24 | 42 | 49 | 41 | 61 | 61 | 65 | 65.19 | 44 | 62 | 1.69 | 0.757 | 0.00028484 |
| AT3G24500.1 | MBF1C | protein_coding | multiprotein bridging fac | 103 | 75 | 64 | 101 | 113 | 67 | 217 | 184 | 164 | 219 | 300 | 182 | 200.93 | 188 | 234 | 1.689 | 0.757 | 4.09E-05 |
| AT3G26650.1 | GAPA1 | protein_coding | glyceraldehyde 3-phosph | 37196 | 31344 | 26702 | 58229 | 49546 | 48122 | 78189 | 77006 | 68359 | 126304 | 131533 | 130865 | 104147.44 | 74518 | 129567 | 1.688 | 0.756 | 5.89E-06 |
| AT4G37430.1 | CYP81F1 | protein_coding | cytochrome P450, family | 32 | 21 | 15 | 45 | 26 | 26 | 67 | 52 | 38 | 98 | 69 | 71 | 90.67 | 52 | 79 | 1.687 | 0.755 | 0.00015087 |
| AT1G62510.1 | | protein_coding | Bifunctional inhibitor/lipi | 81 | 133 | 107 | 143 | 116 | 178 | 170 | 327 | 274 | 310 | 308 | 484 | 1003.87 | 257 | 367 | 1.688 | 0.755 | 4.88E-05 |
| AT5G59920.1 | ULI3 | protein_coding | Cysteine/Histidine-rich C1 | 558 | 384 | 275 | 796 | 596 | 458 | 1173 | 943 | 704 | 1727 | 1582 | 1246 | 1015.27 | 940 | 1518 | 1.688 | 0.755 | 4.08E-06 |
| AT3G18870.1 | | protein_coding | Mitochondrial transcripti | 112 | 99 | 68 | 113 | 113 | 111 | 235 | 243 | 174 | 245 | 300 | 302 | 279.79 | 217 | 282 | 1.688 | 0.755 | 1.36E-06 |
| AT2G29420.1 | GSTU7 | protein_coding | glutathione S-transferase | 68 | 48 | 59 | 153 | 42 | 51 | 143 | 118 | 151 | 332 | 112 | 139 | 242.1 | 137 | 194 | 1.687 | 0.754 | 0.00237911 |
| AT5G17980.1 | | protein_coding | C2 calcium/lipid-binding | 71 | 73 | 44 | 149 | 93 | 120 | 149 | 179 | 113 | 323 | 247 | 326 | 190.97 | 147 | 299 | 1.684 | 0.752 | 4.01E-05 |
| AT1G07450.1 | | protein_coding | NAD(P)-binding Rossman | 74 | 52 | 28 | 85 | 58 | 34 | 156 | 128 | 72 | 184 | 154 | 92 | 146.33 | 119 | 143 | 1.683 | 0.751 | 0.00076992 |
| AT3G21010.1 | | transposable_elen | transposable element ger | 44 | 35 | 37 | 55 | 51 | 41 | 92 | 86 | 95 | 119 | 135 | 111 | 97.75 | 91 | 122 | 1.683 | 0.751 | 5.62E-05 |
| AT1G24880.1 | | protein_coding | UDP-3-O-acyl N-acetylgly | 222 | 185 | 219 | 304 | 229 | 306 | 467 | 455 | 561 | 659 | 608 | 832 | 703.05 | 494 | 700 | 1.683 | 0.751 | 7.87E-09 |
| AT1G72120.1 | NPF5.14 | protein_coding | Major facilitator superfar | 19 | 6 | 13 | 22 | 21 | 13 | 40 | 15 | 33 | 48 | 56 | 35 | 42.18 | 29 | 46 | 1.681 | 0.75 | 0.00742224 |
| AT1G55080.1 | MED9 | protein_coding | | 67 | 47 | 55 | 124 | 85 | 78 | 141 | 115 | 141 | 269 | 226 | 212 | 212.5 | 132 | 236 | 1.68 | 0.749 | 1.09E-06 |
| AT5G16200.1 | | protein_coding | 50S ribosomal protein-rel | 17 | 15 | 9 | 16 | 13 | 12 | 36 | 37 | 23 | 35 | 35 | 33 | 51.34 | 32 | 34 | 1.68 | 0.748 | 0.00104879 |
| AT4G24930.1 | | protein_coding | thylakoid lumenal 17.9 kD | 882 | 699 | 692 | 1167 | 930 | 912 | 1854 | 1717 | 1772 | 2531 | 2469 | 2480 | 3453.84 | 1781 | 2493 | 1.679 | 0.748 | 7.35E-12 |
| AT1G74100.1 | SOT16 | protein_coding | sulfotransferase 16 | 800 | 606 | 403 | 1140 | 941 | 620 | 1682 | 1489 | 1032 | 2473 | 2498 | 1686 | 2037.15 | 1401 | 2219 | 1.679 | 0.747 | 9.49E-06 |
| AT2G45920.1 | PUB37 | protein_coding | U-box domain-containing | 97 | 73 | 83 | 153 | 92 | 98 | 204 | 179 | 212 | 332 | 244 | 267 | 246.17 | 198 | 281 | 1.675 | 0.744 | 3.12E-07 |
| AT5G59780.3 | MYB59 | protein_coding | myb domain protein 59 | 182 | 152 | 117 | 251 | 205 | 157 | 383 | 373 | 300 | 544 | 544 | 427 | 511.3 | 352 | 505 | 1.673 | 0.742 | 3.66E-09 |
| AT4G39470.1 | | protein_coding | Tetratricopeptide repeat | 274 | 199 | 175 | 400 | 297 | 259 | 576 | 489 | 448 | 868 | 788 | 704 | 670.15 | 504 | 787 | 1.671 | 0.741 | 3.11E-10 |
| AT4G16870.1 | | transposable_elen | transposable element ger | 360 | 501 | 389 | 469 | 508 | 500 | 757 | 1231 | 996 | 1017 | 1349 | 1360 | 2076.06 | 995 | 1242 | 1.67 | 0.74 | 6.28E-06 |
| AT1G70980.1 | SYNC3 | protein_coding | Class II aminoacyl-tRNA a | 377 | 264 | 268 | 619 | 381 | 457 | 792 | 649 | 686 | 1343 | 1011 | 1243 | 777.26 | 709 | 1199 | 1.671 | 0.74 | 3.65E-08 |
| AT3G57010.1 | SSL8 | protein_coding | Calcium-dependent phos | 51 | 46 | 17 | 79 | 87 | 46 | 107 | 113 | 44 | 171 | 231 | 125 | 117.78 | 88 | 176 | 1.669 | 0.739 | 0.00623456 |
| AT4G36060.1 | BHLH11 | protein_coding | basic helix-loop-helix (bH | 25 | 17 | 24 | 31 | 34 | 25 | 53 | 42 | 61 | 67 | 90 | 68 | 72.54 | 52 | 75 | 1.669 | 0.739 | 0.00194319 |
| AT5G66130.1 | RAD17 | protein_coding | RADIATION SENSITIVE 17 | 83 | 86 | 60 | 107 | 124 | 107 | 174 | 211 | 154 | 232 | 329 | 291 | 214.45 | 180 | 284 | 1.669 | 0.739 | 3.10E-06 |
| AT5G13520.1 | LKHA4 | protein_coding | peptidase M1 family prot | 333 | 284 | 234 | 479 | 442 | 407 | 700 | 698 | 599 | 1039 | 1173 | 1107 | 924.14 | 666 | 1106 | 1.666 | 0.737 | 2.30E-10 |
| AT3G51990.1 | | protein_coding | Protein kinase superfamil | 70 | 77 | 52 | 84 | 74 | 66 | 147 | 189 | 133 | 182 | 196 | 179 | 184.09 | 156 | 186 | 1.665 | 0.736 | 1.63E-06 |
| AT5G05990.1 | | protein_coding | Mitochondrial glycoprote | 266 | 138 | 124 | 351 | 228 | 182 | 559 | 339 | 317 | 761 | 605 | 495 | 499.94 | 405 | 620 | 1.664 | 0.735 | 0.00014465 |
| AT2G23330.1 | | transposable_elen | transposable element ger | 49 | 28 | 46 | 108 | 67 | 87 | 103 | 69 | 118 | 234 | 178 | 237 | 190.2 | 97 | 216 | 1.663 | 0.734 | 6.26E-06 |
| AT1G05140.1 | ARASP2 | protein_coding | Peptidase M50 family pro | 875 | 770 | 703 | 1287 | 1075 | 916 | 1839 | 1892 | 1800 | 2792 | 2854 | 2491 | 2587.11 | 1844 | 2712 | 1.663 | 0.733 | 1.52E-09 |
| AT4G38545.1 | | other_rna | other RNA | 46 | 44 | 50 | 50 | 47 | 62 | 97 | 108 | 128 | 108 | 125 | 169 | 143.19 | 111 | 134 | 1.66 | 0.732 | 9.79E-06 |
| AT2G31902.1 | | other_rna | | 71 | 68 | 65 | 123 | 106 | 103 | 149 | 167 | 166 | 267 | 281 | 280 | 157.92 | 161 | 276 | 1.661 | 0.732 | 5.77E-06 |
| AT5G28335.1 | | transposable_elen | transposable element ger | 17 | 17 | 8 | 15 | 9 | 9 | 36 | 42 | 20 | 33 | 24 | 24 | 46.28 | 33 | 27 | 1.66 | 0.731 | 0.00304775 |
| AT2G43375.1 | | other_rna | other RNA | 878 | 535 | 388 | 835 | 826 | 493 | 1846 | 1314 | 993 | 1811 | 2193 | 1341 | 854.83 | 1384 | 1782 | 1.66 | 0.731 | 0.00126185 |
| AT2G36355.1 | | protein_coding | Family of unknown functi | 27 | 17 | 17 | 19 | 21 | 19 | 57 | 42 | 44 | 41 | 56 | 52 | 68.03 | 48 | 50 | 1.66 | 0.731 | 0.00038441 |
| AT4G34230.1 | CAD5 | protein_coding | cinnamyl alcohol dehydrc | 236 | 188 | 131 | 379 | 246 | 256 | 496 | 462 | 335 | 822 | 653 | 696 | 604.18 | 431 | 724 | 1.66 | 0.731 | 3.06E-07 |
| AT2G43340.1 | | protein_coding | Protein of unknown funct | 265 | 218 | 251 | 477 | 347 | 299 | 557 | 536 | 643 | 1035 | 921 | 813 | 623.99 | 579 | 923 | 1.66 | 0.731 | 3.52E-09 |
| AT3G21870.1 | CYCU1-1 | protein_coding | cyclin p2;1 | 62 | 62 | 69 | 129 | 61 | 106 | 130 | 152 | 177 | 280 | 162 | 288 | 184.48 | 153 | 243 | 1.658 | 0.73 | 9.51E-05 |
| AT4G18490.1 | | protein_coding | | 72 | 76 | 63 | 170 | 131 | 99 | 151 | 187 | 161 | 369 | 348 | 269 | 170.28 | 166 | 329 | 1.659 | 0.73 | 6.88E-05 |
| AT5G58260.1 | ndhN | protein_coding | oxidoreductases, acting o | 702 | 721 | 612 | 1291 | 1177 | 1237 | 1476 | 1771 | 1567 | 2800 | 3125 | 3364 | 2992.36 | 1605 | 3096 | 1.658 | 0.73 | 2.52E-09 |
| AT1G55180.1 | PLDEPSILON | protein_coding | phospholipase D alpha 4 | 43 | 35 | 39 | 79 | 50 | 62 | 90 | 86 | 100 | 171 | 133 | 169 | 92.44 | 92 | 158 | 1.658 | 0.729 | 0.00358528 |
| AT5G39890.1 | PCO2 | protein_coding | Protein of unknown funct | 72 | 62 | 76 | 113 | 83 | 109 | 151 | 152 | 195 | 245 | 220 | 296 | 148.6 | 166 | 254 | 1.657 | 0.729 | 0.00010664 |
| AT2G03710.1 | AGL3 | protein_coding | K-box region and MADS-b | 119 | 92 | 82 | 164 | 161 | 104 | 250 | 226 | 210 | 356 | 427 | 283 | 285.49 | 229 | 355 | 1.657 | 0.729 | 1.53E-06 |
| AT2G28315.1 | | protein_coding | Nucleotide/sugar transpo | 31 | 25 | 33 | 42 | 42 | 35 | 65 | 61 | 84 | 91 | 112 | 95 | 96.48 | 70 | 99 | 1.656 | 0.728 | 0.00036261 |
| AT1G32583.1 | TDL1 | protein_coding | | 67 | 40 | 50 | 73 | 80 | 59 | 141 | 98 | 128 | 158 | 212 | 160 | 126.59 | 122 | 177 | 1.656 | 0.728 | 5.35E-05 |
| AT1G03340.1 | | protein_coding | | 58 | 59 | 35 | 44 | 45 | 39 | 122 | 145 | 90 | 95 | 119 | 106 | 98.87 | 119 | 107 | 1.655 | 0.727 | 0.00324914 |
| AT5G18780.1 | | protein_coding | F-box/RNI-like superfamil | 21 | 11 | 19 | 44 | 40 | 33 | 44 | 27 | 49 | 95 | 106 | 90 | 64.85 | 40 | 97 | 1.656 | 0.727 | 0.00205794 |
| AT1G72360.2 | ERF073 | protein_coding | Integrase-type DNA-bindi | 26 | 28 | 35 | 47 | 26 | 42 | 55 | 69 | 90 | 102 | 69 | 114 | 73.03 | 71 | 95 | 1.654 | 0.726 | 0.003417 |
| AT5G14090.1 | | protein_coding | | 178 | 146 | 108 | 338 | 237 | 190 | 374 | 359 | 276 | 733 | 629 | 517 | 464.07 | 336 | 626 | 1.654 | 0.726 | 1.53E-06 |
| AT3G11900.1 | ANT1 | protein_coding | aromatic and neutral trar | 275 | 261 | 241 | 583 | 371 | 351 | 578 | 641 | 617 | 1265 | 985 | 955 | 763.86 | 612 | 1068 | 1.655 | 0.726 | 2.93E-08 |
| AT3G57470.2 | | protein_coding | Insulinase (Peptidase fam | 1035 | 882 | 955 | 1194 | 1208 | 1156 | 2176 | 2167 | 2445 | 2590 | 3207 | 3144 | 2514.75 | 2263 | 2980 | 1.654 | 0.726 | 3.68E-09 |
| AT1G69730.1 | WAKL9 | protein_coding | Wall-associated kinase fa | 377 | 177 | 169 | 564 | 313 | 206 | 792 | 435 | 433 | 1223 | 831 | 560 | 530.38 | 553 | 871 | 1.653 | 0.725 | 0.00226451 |
| AT2G46450.1 | CNGC12 | protein_coding | cyclic nucleotide-gated ch | 775 | 490 | 349 | 1122 | 682 | 400 | 1629 | 1204 | 893 | 2434 | 1811 | 1088 | 1501.51 | 1242 | 1778 | 1.653 | 0.725 | 0.00130345 |
| AT3G48490.1 | | protein_coding | | 89 | 74 | 64 | 101 | 108 | 93 | 187 | 182 | 164 | 219 | 287 | 253 | 212.64 | 178 | 253 | 1.653 | 0.725 | 5.37E-07 |
| AT5G27660.1 | DEGP14 | protein_coding | Trypsin family protein wit | 1512 | 901 | 945 | 4252 | 2328 | 2172 | 3178 | 2214 | 2419 | 9223 | 6180 | 5907 | 3209.93 | 2604 | 7103 | 1.651 | 0.724 | 0.0041927 |
| AT3G03470.1 | CYP89A9 | protein_coding | cytochrome P450, family | 65 | 48 | 38 | 67 | 41 | 50 | 137 | 118 | 97 | 145 | 109 | 136 | 124.14 | 117 | 130 | 1.651 | 0.723 | 0.00015629 |
| AT3G05650.1 | AtRLP32 | protein_coding | receptor like protein 32 | 13 | 20 | 24 | 44 | 38 | 35 | 27 | 49 | 61 | 95 | 101 | 95 | 67.96 | 46 | 97 | 1.65 | 0.722 | 0.00538273 |
| AT1G09340.1 | CSP41B | protein_coding | chloroplast RNA binding | 16340 | 15813 | 14136 | 19823 | 17432 | 18539 | 34348 | 38850 | 36189 | 42998 | 46278 | 50416 | 47421.38 | 36462 | 46564 | 1.649 | 0.722 | 2.83E-06 |
| AT5G09443.1 | | other_rna | other RNA | 21 | 23 | 8 | 42 | 31 | 44 | 44 | 57 | 20 | 91 | 82 | 120 | 70.32 | 40 | 98 | 1.648 | 0.721 | 0.0033182 |
| AT1G16090.1 | WAKL7 | protein_coding | wall associated kinase-lik | 66 | 85 | 66 | 91 | 83 | 65 | 139 | 209 | 169 | 197 | 220 | 177 | 185.22 | 172 | 198 | 1.649 | 0.721 | 2.82E-05 |
| AT2G35810.1 | | protein_coding | | 194 | 136 | 92 | 268 | 178 | 182 | 408 | 334 | 236 | 581 | 473 | 495 | 408.91 | 326 | 516 | 1.645 | 0.718 | 1.32E-06 |
| AT3G61100.1 | | protein_coding | Putative endonuclease or | 147 | 85 | 88 | 155 | 120 | 123 | 309 | 209 | 225 | 336 | 319 | 334 | 292.66 | 248 | 330 | 1.643 | 0.716 | 3.49E-05 |
| AT2G46140.1 | | protein_coding | Late embryogenesis abun | 7 | 16 | 14 | 15 | 18 | 16 | 15 | 39 | 36 | 33 | 48 | 44 | 46.57 | 30 | 42 | 1.639 | 0.713 | 0.00871693 |
| AT3G23920.1 | BAM1 | protein_coding | beta-amylase 1 | 347 | 419 | 320 | 564 | 502 | 640 | 729 | 1029 | 819 | 1223 | 1333 | 1740 | 1226.6 | 859 | 1432 | 1.639 | 0.713 | 1.48E-06 |
| AT1G68010.2 | HPR | protein_coding | hydroxypyruvate reducta | 8830 | 9373 | 7495 | 14182 | 12274 | 13754 | 18561 | 23028 | 19188 | 30762 | 32585 | 37403 | 29087.57 | 20259 | 33583 | 1.64 | 0.713 | 1.22E-06 |
| AT1G11860.1 | GDCST | protein_coding | Glycine cleavage T-protei | 20857 | 20291 | 16744 | 25942 | 23809 | 25051 | 43843 | 49851 | 42866 | 56271 | 63207 | 68125 | 62101.05 | 45520 | 62534 | 1.638 | 0.712 | 7.12E-06 |
| AT3G13470.1 | CPN60B2 | protein_coding | TCP-1/cpn60 chaperonin | 5019 | 3001 | 1708 | 4306 | 3094 | 2133 | 10550 | 7373 | 4373 | 9340 | 8214 | 5801 | 9240.83 | 7432 | 7785 | 1.637 | 0.711 | 0.00254719 |
| AT1G16030.1 | HSP70-5 | protein_coding | heat shock protein 70B | 39 | 64 | 59 | 47 | 39 | 44 | 82 | 157 | 151 | 102 | 104 | 120 | 124.71 | 130 | 109 | 1.637 | 0.711 | 0.00099636 |
| AT2G30520.1 | RPT2 | protein_coding | Phototropic-responsive N | 6841 | 7185 | 5451 | 11303 | 9213 | 8062 | 14380 | 17652 | 13955 | 24517 | 24458 | 21924 | 15091.38 | 15329 | 23633 | 1.636 | 0.71 | 1.10E-07 |
| AT2G44930.1 | | protein_coding | Plant protein of unknowr | 24 | 27 | 26 | 44 | 39 | 46 | 50 | 66 | 67 | 95 | 104 | 125 | 74.61 | 61 | 108 | 1.634 | 0.708 | 0.00822732 |
| AT5G08610.1 | RH26 | protein_coding | P-loop containing nucleo | 1508 | 1006 | 704 | 1462 | 1149 | 955 | 3170 | 2472 | 1802 | 3171 | 3050 | 2597 | 2887.07 | 2481 | 2939 | 1.634 | 0.708 | 2.95E-05 |
| AT4G38540.1 | | protein_coding | FAD/NAD(P)-binding oxid | 54 | 42 | 34 | 136 | 43 | 50 | 114 | 103 | 87 | 295 | 114 | 136 | 127.36 | 101 | 182 | 1.633 | 0.707 | 0.00704664 |
| AT1G62085.1 | | protein_coding | Mitochondrial transcripti | 147 | 110 | 133 | 188 | 167 | 146 | 309 | 270 | 340 | 408 | 443 | 397 | 308.62 | 306 | 416 | 1.632 | 0.707 | 1.35E-07 |
| AT1G60030.1 | NAT7 | protein_coding | nucleobase-ascorbate tra | 18 | 20 | 10 | 35 | 25 | 18 | 38 | 49 | 26 | 76 | 66 | 49 | 48.6 | 38 | 64 | 1.631 | 0.706 | 0.00426258 |
| AT2G17830.1 | | protein_coding | F-box and associated inte | 81 | 53 | 63 | 78 | 68 | 74 | 170 | 130 | 161 | 169 | 181 | 201 | 129.42 | 154 | 184 | 1.632 | 0.706 | 0.00030066 |
| AT5G27680.1 | RECQSIM | protein_coding | RECQ helicase SIM | 138 | 140 | 126 | 157 | 130 | 116 | 290 | 344 | 323 | 341 | 345 | 315 | 270.38 | 319 | 334 | 1.63 | 0.705 | 3.38E-06 |
| AT3G55010.1 | PUR5 | protein_coding | phosphoribosylformylglyc | 745 | 432 | 374 | 832 | 653 | 575 | 1566 | 1061 | 957 | 1805 | 1734 | 1564 | 1511.57 | 1195 | 1701 | 1.627 | 0.703 | 3.73E-05 |
| AT1G80770.1 | PDE318 | protein_coding | P-loop containing nucleo | 1044 | 680 | 544 | 992 | 800 | 667 | 2195 | 1671 | 1393 | 2152 | 2124 | 1814 | 2138.74 | 1753 | 2030 | 1.628 | 0.703 | 1.60E-06 |
| AT3G23510.1 | | protein_coding | Cyclopropane-fatty-acyl-p | 149 | 86 | 99 | 300 | 161 | 144 | 313 | 211 | 253 | 651 | 427 | 392 | 231.48 | 259 | 490 | 1.627 | 0.702 | 0.00107984 |
| AT1G16445.1 | | protein_coding | S-adenosyl-L-methionine- | 60 | 70 | 52 | 81 | 62 | 57 | 126 | 172 | 133 | 176 | 165 | 155 | 170.48 | 144 | 165 | 1.626 | 0.702 | 1.65E-05 |
| AT4G38440.1 | | protein_coding | | 311 | 273 | 224 | 388 | 309 | 288 | 654 | 671 | 573 | 842 | 820 | 783 | 613.04 | 633 | 815 | 1.626 | 0.702 | 2.02E-09 |
| AT4G27370.1 | VIII-B | protein_coding | P-loop containing nucleo | 251 | 214 | 227 | 249 | 208 | 217 | 528 | 526 | 581 | 540 | 552 | 590 | 507.46 | 545 | 561 | 1.626 | 0.701 | 1.32E-09 |
| AT1G27920.1 | MAP65-8 | protein_coding | microtubule-associated p | 18 | 44 | 30 | 38 | 39 | 28 | 38 | 108 | 77 | 82 | 104 | 76 | 79.9 | 74 | 87 | 1.624 | 0.7 | 0.00508274 |
| AT1G61600.1 | | protein_coding | Protein of unknown funct | 10 | 13 | 20 | 25 | 24 | 14 | 21 | 32 | 51 | 54 | 64 | 38 | 74.44 | 35 | 52 | 1.625 | 0.7 | 0.00164476 |
| AT2G05300.1 | | pseudogene | | 24 | 24 | 22 | 43 | 41 | 28 | 50 | 59 | 56 | 93 | 109 | 76 | 80.49 | 55 | 93 | 1.624 | 0.7 | 0.00029213 |
| AT1G50790.1 | | protein_coding | Plant mobile domain prot | 68 | 60 | 77 | 85 | 91 | 78 | 143 | 147 | 197 | 184 | 242 | 212 | 124.57 | 162 | 213 | 1.624 | 0.699 | 0.00062846 |
| AT2G31560.1 | | protein_coding | Protein of unknown funct | 191 | 141 | 97 | 296 | 248 | 215 | 401 | 346 | 248 | 642 | 658 | 585 | 473.08 | 332 | 628 | 1.624 | 0.699 | 3.27E-06 |
| AT4G34588.1 | CPuORF2 | protein_coding | conserved peptide upstre | 100 | 85 | 41 | 171 | 112 | 78 | 210 | 209 | 105 | 371 | 297 | 212 | 360.86 | 175 | 293 | 1.622 | 0.698 | 0.00022054 |
| AT4G04020.1 | PAP1 | protein_coding | fibrillin | 1507 | 1256 | 826 | 2220 | 1611 | 1396 | 3168 | 3086 | 2115 | 4815 | 4277 | 3796 | 4769 | 2790 | 4296 | 1.622 | 0.698 | 1.59E-06 |
| AT2G36970.1 | UGT86A1 | protein_coding | UDP-Glycosyltransferase | 115 | 84 | 48 | 151 | 62 | 51 | 242 | 206 | 123 | 328 | 165 | 139 | 264.21 | 190 | 211 | 1.621 | 0.697 | 0.00384043 |
| AT5G60030.1 | | protein_coding | | 19 | 18 | 10 | 27 | 25 | 16 | 40 | 44 | 26 | 59 | 66 | 44 | 276.31 | 37 | 56 | 1.621 | 0.697 | 2.28E-05 |
| AT4G27860.1 | MEB1 | protein_coding | vacuolar iron transporter | 614 | 359 | 279 | 1014 | 602 | 461 | 1291 | 882 | 714 | 2199 | 1598 | 1254 | 1404.43 | 962 | 1684 | 1.62 | 0.696 | 0.00047438 |
| AT1G26218.1 | | other_rna | other RNA | 39 | 50 | 31 | 74 | 38 | 50 | 82 | 123 | 79 | 161 | 101 | 136 | 117.87 | 95 | 133 | 1.62 | 0.696 | 0.00031966 |
| AT4G18220.1 | | protein_coding | Drug/metabolite transpo | 165 | 140 | 119 | 256 | 209 | 130 | 347 | 344 | 305 | 555 | 555 | 354 | 454.04 | 332 | 488 | 1.62 | 0.696 | 1.40E-06 |
| AT1G62975.1 | BHLH125 | protein_coding | basic helix-loop-helix (bH | 62 | 71 | 62 | 117 | 86 | 56 | 130 | 174 | 159 | 254 | 228 | 152 | 163.07 | 154 | 211 | 1.618 | 0.695 | 0.00058864 |
| AT5G63320.1 | GTE10 | protein_coding | nuclear protein X1 | 1041 | 818 | 872 | 1206 | 916 | 830 | 2188 | 2010 | 2232 | 2616 | 2432 | 2257 | 2159.78 | 2143 | 2435 | 1.619 | 0.695 | 2.21E-09 |
| AT1G06450.1 | CAF1-1 | protein_coding | Polynucleotidyl transfera | 54 | 62 | 38 | 94 | 80 | 87 | 114 | 152 | 97 | 204 | 212 | 237 | 134.65 | 121 | 218 | 1.617 | 0.694 | 0.00113296 |
| AT3G06350.1 | EMB3004 | protein_coding | dehydroquinate dehydrat | 1214 | 807 | 782 | 1934 | 1275 | 1227 | 2552 | 1983 | 2002 | 4195 | 3385 | 3337 | 3096.64 | 2179 | 3639 | 1.618 | 0.694 | 2.05E-07 |
| AT1G13330.1 | HOP2 | protein_coding | Arabidopsis Hop2 homolo | 52 | 48 | 58 | 99 | 70 | 72 | 109 | 118 | 148 | 215 | 186 | 196 | 137.65 | 125 | 199 | 1.617 | 0.693 | 0.00023848 |
| AT3G61860.1 | RS31 | protein_coding | RNA-binding (RRM/RBD/ | 387 | 365 | 274 | 557 | 428 | 466 | 814 | 897 | 701 | 1208 | 1136 | 1267 | 1135.5 | 804 | 1204 | 1.617 | 0.693 | 1.75E-08 |
| AT5G23060.1 | CAS | protein_coding | calcium sensing receptor | 9770 | 5596 | 4811 | 12171 | 9177 | 7575 | 20537 | 13748 | 12316 | 26400 | 24363 | 20600 | 19922.7 | 15534 | 23788 | 1.616 | 0.692 | 7.26E-05 |
| AT3G57590.1 | | protein_coding | F-box and associated inte | 53 | 49 | 57 | 69 | 59 | 68 | 111 | 120 | 146 | 150 | 157 | 185 | 151.34 | 126 | 164 | 1.616 | 0.692 | 1.83E-05 |
| AT4G19820.1 | | protein_coding | Glycosyl hydrolase family | 38 | 23 | 25 | 38 | 35 | 55 | 80 | 57 | 64 | 82 | 93 | 150 | 86.45 | 67 | 108 | 1.614 | 0.691 | 0.00930395 |
| AT2G26170.1 | CYP711A1 | protein_coding | cytochrome P450, family | 210 | 112 | 111 | 416 | 290 | 189 | 441 | 275 | 284 | 902 | 770 | 514 | 435.53 | 333 | 729 | 1.615 | 0.691 | 0.00033562 |

| | | | | | | | | | | | | | | | | | | | | | |
|---|---|---|---|---|---|---|---|---|---|---|---|---|---|---|---|---|---|---|---|---|---|
| AT5G56380.1 | | protein_coding | F-box/RNI-like/FBD-like d | 198 | 130 | 149 | 291 | 216 | 194 | 416 | 319 | 381 | 631 | 573 | 528 | 514.12 | 372 | 577 | 1.614 | 0.691 | 1.85E-07 |
| AT1G66880.1 | | protein_coding | Protein kinase superfamil | 141 | 114 | 109 | 215 | 160 | 121 | 296 | 280 | 279 | 466 | 425 | 329 | 296.65 | 285 | 407 | 1.613 | 0.69 | 1.93E-06 |
| AT5G40980.1 | | protein_coding | Protein of unknown funct | 23 | 20 | 17 | 42 | 45 | 25 | 48 | 49 | 44 | 91 | 119 | 68 | 73.7 | 47 | 93 | 1.611 | 0.688 | 0.00504102 |
| AT1G56430.1 | NAS4 | protein_coding | nicotianamine synthase 4 | 287 | 280 | 148 | 406 | 409 | 212 | 603 | 688 | 379 | 881 | 1086 | 577 | 767.22 | 557 | 848 | 1.612 | 0.688 | 0.00073678 |
| AT1G33800.1 | GXM3 | protein_coding | Protein of unknown funct | 27 | 30 | 36 | 43 | 45 | 35 | 57 | 74 | 92 | 93 | 119 | 95 | 92.92 | 74 | 102 | 1.61 | 0.687 | 0.00547444 |
| AT5G20110.1 | | protein_coding | Dynein light chain type 1 | 35 | 39 | 50 | 75 | 57 | 84 | 74 | 96 | 128 | 163 | 151 | 228 | 179.83 | 99 | 181 | 1.61 | 0.687 | 0.00010255 |
| AT3G58930.1 | | protein_coding | F-box/RNI-like superfamil | 116 | 74 | 69 | 126 | 95 | 81 | 244 | 182 | 177 | 273 | 252 | 220 | 228.2 | 201 | 248 | 1.61 | 0.687 | 8.13E-06 |
| AT2G04360.1 | | protein_coding | | 750 | 509 | 476 | 1052 | 773 | 671 | 1577 | 1251 | 1219 | 2282 | 2052 | 1825 | 1930.22 | 1349 | 2053 | 1.61 | 0.687 | 3.29E-07 |
| AT2G01140.1 | FBA3 | protein_coding | Aldolase superfamily prot | 902 | 992 | 614 | 1374 | 1027 | 1809 | 1896 | 2437 | 1572 | 2980 | 2726 | 4919 | 3815.26 | 1968 | 3542 | 1.609 | 0.686 | 0.00033917 |
| AT4G24830.1 | | protein_coding | arginosuccinate synthase | 1446 | 1047 | 793 | 1860 | 1329 | 1235 | 3040 | 2572 | 2030 | 4035 | 3528 | 3359 | 3194.32 | 2547 | 3641 | 1.608 | 0.685 | 2.15E-06 |
| AT1G04000.1 | | protein_coding | | 47 | 72 | 43 | 72 | 69 | 100 | 99 | 177 | 110 | 156 | 183 | 272 | 333.31 | 129 | 204 | 1.607 | 0.684 | 0.00045897 |
| AT3G05730.1 | | protein_coding | | 5922 | 6867 | 6672 | 7836 | 8451 | 7641 | 12448 | 16871 | 17081 | 16997 | 22435 | 20779 | 18039.36 | 15467 | 20070 | 1.607 | 0.684 | 0.00029126 |
| AT2G35370.1 | GDH1 | protein_coding | glycine decarboxylase cor | 7565 | 5961 | 5513 | 11445 | 9336 | 8667 | 15902 | 14645 | 14114 | 24825 | 24785 | 23569 | 22084.89 | 14887 | 24393 | 1.607 | 0.684 | 2.80E-07 |
| AT3G26060.2 | PRXQ | protein_coding | Thioredoxin superfamily | | 4528 | 3615 | 3615 | 6569 | 5401 | 5353 | 9518 | 8881 | 9255 | 14249 | 14338 | 14557 | 17860.87 | 9218 | 14381 | 1.606 | 0.684 | 7.06E-08 |
| AT4G38100.1 | CURT1D | protein_coding | | 481 | 257 | 181 | 500 | 367 | 260 | 1011 | 631 | 463 | 1085 | 974 | 707 | 967.68 | 702 | 922 | 1.605 | 0.683 | 0.00108052 |
| AT1G04870.2 | PRMT10 | protein_coding | protein arginine methyltr | 437 | 287 | 278 | 558 | 423 | 396 | 919 | 705 | 712 | 1210 | 1123 | 1077 | 919.64 | 779 | 1137 | 1.605 | 0.683 | 1.91E-07 |
| AT1G32870.1 | NAC13 | protein_coding | NAC domain protein 13 | 88 | 80 | 87 | 196 | 148 | 127 | 185 | 197 | 223 | 425 | 393 | 345 | 232.49 | 202 | 388 | 1.605 | 0.682 | 2.37E-05 |
| AT2G42690.1 | | protein_coding | alpha/beta-Hydrolases su | 1181 | 1086 | 784 | 2220 | 1798 | 1257 | 2483 | 2668 | 2007 | 4815 | 4773 | 3418 | 3061.44 | 2386 | 4335 | 1.604 | 0.682 | 2.08E-06 |
| AT3G01800.1 | | protein_coding | Ribosome recycling facto | 189 | 126 | 82 | 236 | 167 | 140 | 397 | 310 | 210 | 512 | 443 | 381 | 384.35 | 306 | 445 | 1.603 | 0.681 | 0.00011937 |
| AT3G28455.1 | CLE25 | protein_coding | CLAVATA3/ESR-RELATED | 34 | 27 | 36 | 52 | 53 | 34 | 71 | 66 | 92 | 113 | 141 | 92 | 108.13 | 76 | 115 | 1.602 | 0.68 | 0.00022336 |
| AT2G22240.1 | IPS2 | protein_coding | myo-inositol-1-phosphate | 306 | 303 | 258 | 482 | 536 | 357 | 643 | 744 | 660 | 1046 | 1423 | 971 | 747.32 | 682 | 1147 | 1.602 | 0.68 | 2.65E-06 |
| AT3G02190.1 | RPL39B | protein_coding | Ribosomal protein L39 fa | 151 | 141 | 107 | 251 | 182 | 216 | 317 | 346 | 274 | 544 | 483 | 587 | 494.25 | 312 | 538 | 1.602 | 0.68 | 1.10E-06 |
| AT3G50220.1 | IRX15 | protein_coding | Protein of unknown funct | 116 | 93 | 107 | 120 | 94 | 103 | 244 | 228 | 274 | 260 | 250 | 280 | 144.5 | 249 | 263 | 1.601 | 0.679 | 0.00284131 |
| AT3G59068.1 | | other_rna | | 718 | 571 | 481 | 795 | 727 | 674 | 1509 | 1403 | 1231 | 1724 | 1930 | 1833 | 833.06 | 1381 | 1829 | 1.601 | 0.679 | 0.00011458 |
| AT3G52670.1 | | protein_coding | FBD, F-box, Skp2-like and | 45 | 36 | 31 | 54 | 49 | 39 | 95 | 88 | 79 | 117 | 130 | 106 | 99.63 | 87 | 118 | 1.601 | 0.679 | 0.00010973 |
| AT4G28900.1 | | transposable_elen | transposable element ger | 105 | 75 | 60 | 149 | 107 | 110 | 221 | 184 | 154 | 323 | 284 | 299 | 272.49 | 186 | 302 | 1.601 | 0.679 | 1.90E-05 |
| AT5G37400.1 | | protein_coding | Family of unknown functi | 85 | 56 | 59 | 118 | 88 | 78 | 179 | 138 | 151 | 256 | 234 | 212 | 167.56 | 156 | 234 | 1.599 | 0.678 | 4.05E-05 |
| AT1G44575.1 | PSBS | protein_coding | Chlorophyll A-B binding f | 14563 | 16646 | 11578 | 18462 | 16344 | 19175 | 30612 | 40896 | 29640 | 40046 | 43390 | 52145 | 58783.47 | 33716 | 45194 | 1.599 | 0.677 | 0.00012079 |
| AT5G52050.1 | | protein_coding | MATE efflux family prote | 52 | 29 | 38 | 38 | 21 | 36 | 109 | 71 | 97 | 82 | 56 | 98 | 87.44 | 92 | 79 | 1.598 | 0.676 | 0.00172676 |
| AT5G18670.1 | BAM9 | protein_coding | beta-amylase 3 | 402 | 266 | 286 | 575 | 350 | 550 | 845 | 654 | 732 | 1247 | 929 | 1496 | 702.37 | 744 | 1224 | 1.598 | 0.676 | 0.00027383 |
| AT5G09660.4 | PMDH2 | protein_coding | peroxisomal NAD-malate | 9733 | 11493 | 10547 | 15177 | 14122 | 14666 | 20459 | 28236 | 27001 | 32920 | 37491 | 39883 | 34270.39 | 25232 | 36765 | 1.598 | 0.676 | 2.59E-05 |
| AT2G25840.2 | OVA4 | protein_coding | Nucleotidylyl transferase | 544 | 373 | 341 | 595 | 496 | 490 | 1144 | 916 | 873 | 1291 | 1317 | 1333 | 1940.05 | 978 | 1314 | 1.598 | 0.676 | 5.05E-08 |
| AT2G47910.1 | CRR6 | protein_coding | chlororespiratory reducti | 315 | 344 | 199 | 440 | 291 | 507 | 662 | 845 | 509 | 954 | 773 | 1379 | 1256.66 | 672 | 1035 | 1.596 | 0.674 | 0.00028289 |
| AT2G45440.1 | DHDPS2 | protein_coding | dihydrodipicolinate synth | 1195 | 627 | 577 | 1304 | 912 | 766 | 2512 | 1540 | 1477 | 2829 | 2421 | 2083 | 1947.37 | 1843 | 2444 | 1.596 | 0.674 | 0.00016612 |
| AT2G38860.2 | DJ1E | protein_coding | Class I glutamine amidotr | 57 | 63 | 41 | 124 | 73 | 55 | 120 | 155 | 105 | 269 | 194 | 150 | 293.01 | 127 | 204 | 1.595 | 0.674 | 5.52E-05 |
| AT3G46940.1 | DUT | protein_coding | DUTP-PYROPHOSPHATAS | 565 | 424 | 319 | 663 | 539 | 443 | 1188 | 1042 | 817 | 1438 | 1431 | 1205 | 1312.08 | 1016 | 1358 | 1.596 | 0.674 | 4.07E-06 |
| AT1G79610.1 | NHX6 | protein_coding | Na+/H+ antiporter 6 | 171 | 133 | 111 | 276 | 219 | 160 | 359 | 327 | 284 | 599 | 581 | 435 | 393.41 | 323 | 538 | 1.596 | 0.674 | 7.62E-07 |
| AT3G60176.1 | | other_rna | other RNA | 21 | 16 | 10 | 23 | 16 | 23 | 44 | 39 | 26 | 50 | 42 | 63 | 61.03 | 36 | 52 | 1.594 | 0.673 | 0.00860829 |
| AT1G53060.1 | | protein_coding | Legume lectin family prot | 22 | 17 | 23 | 39 | 33 | 33 | 46 | 42 | 59 | 85 | 88 | 90 | 90.78 | 49 | 88 | 1.595 | 0.673 | 0.00077187 |
| AT1G75900.1 | EXL3 | protein_coding | GDSL-like Lipase/Acylhyd | 514 | 516 | 330 | 706 | 557 | 302 | 1080 | 1268 | 845 | 1531 | 1479 | 821 | 1277 | 1064 | 1277 | 1.595 | 0.673 | 0.00052938 |
| AT1G07010.3 | | protein_coding | Calcineurin-like metallo-p | 953 | 588 | 434 | 1208 | 842 | 660 | 2003 | 1445 | 1111 | 2620 | 2235 | 1795 | 1810.63 | 1520 | 2217 | 1.594 | 0.673 | 0.00013475 |
| AT5G04610.1 | | protein_coding | S-adenosyl-L-methionine- | 21 | 19 | 17 | 32 | 20 | 29 | 44 | 47 | 44 | 69 | 53 | 79 | 50.81 | 45 | 67 | 1.593 | 0.672 | 0.00495399 |
| AT4G38530.1 | PLC3 | protein_coding | phospholipase C1 | 22 | 47 | 42 | 72 | 77 | 66 | 46 | 115 | 108 | 156 | 204 | 179 | 117.36 | 90 | 180 | 1.593 | 0.672 | 0.00442226 |
| AT5G42210.1 | | protein_coding | Major facilitator superfar | 34 | 33 | 22 | 56 | 51 | 41 | 71 | 81 | 56 | 121 | 135 | 111 | 90.49 | 69 | 122 | 1.593 | 0.672 | 0.00201695 |
| AT5G67460.1 | | protein_coding | O-Glycosyl hydrolases far | 25 | 24 | 22 | 42 | 26 | 28 | 53 | 59 | 56 | 91 | 69 | 76 | 97.76 | 56 | 79 | 1.592 | 0.671 | 0.00020726 |
| AT2G23180.1 | CYP96A1 | protein_coding | cytochrome P450, family | 195 | 169 | 158 | 362 | 257 | 205 | 410 | 415 | 404 | 785 | 682 | 557 | 407.8 | 410 | 675 | 1.592 | 0.671 | 1.01E-06 |
| AT5G45590.1 | | protein_coding | Ribosomal protein L35 | 241 | 132 | 150 | 293 | 205 | 166 | 507 | 324 | 384 | 636 | 544 | 451 | 496.8 | 405 | 544 | 1.592 | 0.67 | 1.45E-05 |
| AT5G15450.1 | CLPB3 | protein_coding | casein lytic proteinase B3 | 3753 | 2448 | 2293 | 4244 | 3395 | 3005 | 7889 | 6014 | 5870 | 9206 | 9013 | 8172 | 5361.12 | 6591 | 8797 | 1.591 | 0.67 | 3.52E-07 |
| AT1G48630.1 | RACK1B | protein_coding | receptor for activated C k | 1440 | 885 | 529 | 1635 | 1074 | 779 | 3027 | 2174 | 1354 | 3546 | 2851 | 2118 | 3980.11 | 2185 | 2838 | 1.59 | 0.669 | 0.00244244 |
| AT5G27120.1 | NOP5-1 | protein_coding | NOP56-like pre RNA proc | 1834 | 1060 | 825 | 1759 | 1369 | 1029 | 3855 | 2604 | 2112 | 3815 | 3634 | 2798 | 2764.92 | 2857 | 3416 | 1.59 | 0.669 | 0.0006919 |
| AT2G15620.1 | NIR1 | protein_coding | nitrite reductase 1 | 9697 | 6288 | 5632 | 11417 | 6298 | 6087 | 20384 | 15448 | 14418 | 24765 | 16720 | 16553 | 18572.64 | 16750 | 19346 | 1.59 | 0.669 | 0.00011895 |
| AT5G20220.1 | | protein_coding | zinc knuckle (CCHC-type) | 356 | 262 | 231 | 395 | 296 | 231 | 748 | 644 | 591 | 857 | 786 | 628 | 740.42 | 661 | 757 | 1.587 | 0.666 | 8.67E-08 |
| AT3G19270.1 | CYP707A4 | protein_coding | cytochrome P450, family | 23 | 29 | 17 | 21 | 25 | 27 | 48 | 71 | 44 | 46 | 66 | 73 | 61.04 | 54 | 62 | 1.586 | 0.665 | 0.00382924 |
| AT5G64530.1 | ANAC104 | protein_coding | xylem NAC domain 1 | 25 | 25 | 22 | 33 | 35 | 26 | 53 | 61 | 56 | 72 | 93 | 71 | 69.43 | 57 | 79 | 1.585 | 0.665 | 0.0034402 |
| AT5G64870.1 | FLOT3 | protein_coding | SPFH/Band 7/PHB domai | 22 | 18 | 16 | 47 | 26 | 28 | 46 | 44 | 41 | 102 | 69 | 76 | 59.34 | 44 | 82 | 1.585 | 0.664 | 0.00948171 |
| AT5G44530.1 | | protein_coding | Subtilase family protein | 337 | 457 | 339 | 645 | 944 | 458 | 708 | 1123 | 868 | 1399 | 2506 | 1246 | 1104.59 | 900 | 1717 | 1.585 | 0.664 | 0.00188268 |
| AT1G63350.1 | | protein_coding | Disease resistance protei | 79 | 71 | 79 | 106 | 71 | 64 | 166 | 174 | 202 | 230 | 188 | 174 | 143.69 | 181 | 197 | 1.585 | 0.664 | 0.00139277 |
| AT3G24080.1 | | protein_coding | KRR1 family protein | 76 | 75 | 50 | 110 | 76 | 64 | 160 | 184 | 128 | 239 | 202 | 174 | 948.32 | 157 | 205 | 1.585 | 0.664 | 3.78E-07 |
| AT5G19310.1 | CHR23 | protein_coding | Homeotic gene regulator | 370 | 294 | 279 | 385 | 430 | 299 | 778 | 722 | 714 | 835 | 1142 | 813 | 751.38 | 738 | 930 | 1.585 | 0.664 | 2.31E-07 |
| AT1G54080.2 | UBP1A | protein_coding | oligouridylate-binding pr | 1552 | 1248 | 1264 | 2810 | 2046 | 1884 | 3262 | 3066 | 3236 | 6095 | 5432 | 5123 | 4159.59 | 3188 | 5550 | 1.583 | 0.663 | 1.05E-08 |
| AT4G21320.1 | HSA32 | protein_coding | Aldolase-type TIM barrel | 74 | 50 | 54 | 93 | 54 | 66 | 156 | 123 | 138 | 202 | 143 | 179 | 153.32 | 139 | 175 | 1.582 | 0.662 | 6.99E-05 |
| AT3G01060.1 | | protein_coding | | 1724 | 1411 | 881 | 1978 | 1780 | 1278 | 3624 | 3467 | 2255 | 4290 | 4725 | 3475 | 4156.95 | 3115 | 4163 | 1.583 | 0.662 | 4.16E-05 |
| AT4G12600.2 | | protein_coding | Ribosomal protein L7Ae/l | 1434 | 882 | 721 | 1704 | 1189 | 1009 | 3014 | 2167 | 1846 | 3696 | 3157 | 2744 | 2740.21 | 2342 | 3199 | 1.581 | 0.661 | 9.64E-05 |
| AT4G17600.1 | LIL3:1 | protein_coding | Chlorophyll A-B binding f | 2934 | 2335 | 2120 | 3258 | 2655 | 2597 | 6167 | 5737 | 5427 | 7067 | 7048 | 7062 | 8893.38 | 5777 | 7059 | 1.58 | 0.66 | 9.91E-09 |
| AT1G75040.1 | PR5 | protein_coding | pathogenesis-related gen | 41 | 25 | 15 | 36 | 13 | 17 | 86 | 61 | 38 | 78 | 35 | 46 | 90.79 | 62 | 53 | 1.579 | 0.659 | 0.00285112 |
| AT1G72030.1 | | protein_coding | Acyl-CoA N-acyltransfera | 365 | 379 | 252 | 549 | 472 | 358 | 767 | 931 | 645 | 1191 | 1253 | 974 | 1083.02 | 781 | 1139 | 1.579 | 0.659 | 1.67E-06 |
| AT5G24850.1 | CRY3 | protein_coding | cryptochrome 3 | 304 | 238 | 225 | 407 | 283 | 288 | 639 | 585 | 576 | 883 | 751 | 783 | 657.79 | 600 | 806 | 1.579 | 0.659 | 1.01E-08 |
| AT1G08610.1 | | protein_coding | Pentatricopeptide repeat | 244 | 133 | 125 | 207 | 168 | 134 | 513 | 327 | 320 | 449 | 446 | 364 | 381.85 | 387 | 420 | 1.578 | 0.658 | 5.35E-05 |
| AT1G27400.1 | RPL17A | protein_coding | Ribosomal protein L22p/l | 2165 | 1681 | 1222 | 2445 | 1983 | 2058 | 4551 | 4130 | 3128 | 5303 | 5264 | 5597 | 5806.05 | 3936 | 5388 | 1.578 | 0.658 | 6.24E-06 |
| AT5G04700.1 | | protein_coding | Ankyrin repeat family pro | 22 | 30 | 29 | 47 | 41 | 27 | 46 | 74 | 74 | 102 | 109 | 73 | 77.45 | 65 | 95 | 1.577 | 0.657 | 0.00566158 |
| AT5G60720.1 | | protein_coding | Protein of unknown funct | 35 | 42 | 35 | 71 | 46 | 52 | 74 | 103 | 90 | 154 | 122 | 141 | 106.59 | 89 | 139 | 1.577 | 0.657 | 0.00422657 |
| AT1G62570.1 | FMOGS-OX4 | protein_coding | flavin-monooxygenase gl | 83 | 56 | 92 | 134 | 62 | 82 | 174 | 138 | 236 | 291 | 165 | 223 | 167.89 | 183 | 226 | 1.577 | 0.657 | 0.00158614 |
| AT1G02380.1 | | protein_coding | | 61 | 50 | 42 | 87 | 81 | 50 | 128 | 123 | 108 | 189 | 215 | 136 | 144.52 | 120 | 180 | 1.576 | 0.656 | 0.00599156 |
| AT3G51970.1 | ASAT1 | protein_coding | acyl-CoA sterol acyl trans | 24 | 17 | 9 | 48 | 36 | 34 | 50 | 42 | 23 | 104 | 96 | 92 | 74.53 | 38 | 97 | 1.574 | 0.655 | 0.00598934 |
| AT4G26960.1 | | protein_coding | | 72 | 76 | 62 | 69 | 60 | 99 | 151 | 187 | 159 | 150 | 159 | 269 | 226.92 | 166 | 193 | 1.575 | 0.655 | 0.00015365 |
| AT2G29450.1 | GSTU5 | protein_coding | glutathione S-transferase | 1114 | 1125 | 871 | 1483 | 1208 | 881 | 2342 | 2764 | 2230 | 3217 | 3207 | 2396 | 2741.67 | 2445 | 2940 | 1.574 | 0.655 | 5.95E-06 |
| AT1G64860.1 | SIGA | protein_coding | sigma factor A | 1716 | 1536 | 1302 | 2383 | 1981 | 1584 | 3607 | 3774 | 3333 | 5169 | 5259 | 4308 | 4648.37 | 3571 | 4912 | 1.575 | 0.655 | 2.97E-08 |
| AT4G31870.1 | GPX7 | protein_coding | glutathione peroxidase 7 | 95 | 46 | 52 | 91 | 38 | 42 | 200 | 113 | 133 | 197 | 101 | 114 | 174.57 | 149 | 137 | 1.574 | 0.654 | 0.00609172 |
| AT1G15960.1 | NRAMP6 | protein_coding | NRAMP metal ion transp | 189 | 128 | 92 | 302 | 159 | 130 | 397 | 314 | 236 | 655 | 422 | 354 | 435.54 | 316 | 477 | 1.573 | 0.654 | 0.00116417 |
| AT5G06290.1 | 2-Cys | protein_coding | 2-cysteine peroxiredoxin | 1904 | 1862 | 1416 | 2128 | 1815 | 1746 | 4002 | 4575 | 3625 | 4616 | 4818 | 4748 | 9806.62 | 4067 | 4727 | 1.573 | 0.654 | 1.64E-07 |
| AT2G40240.1 | | protein_coding | Tetratricopeptide repeat | 27 | 22 | 14 | 29 | 24 | 22 | 57 | 54 | 36 | 63 | 64 | 60 | 75.56 | 49 | 62 | 1.572 | 0.653 | 0.00060197 |
| AT1G04350.1 | | protein_coding | 2-oxoglutarate (2OG) and | 810 | 774 | 615 | 1551 | 1256 | 1012 | 1703 | 1902 | 1574 | 3364 | 3334 | 2752 | 2424.61 | 1726 | 3150 | 1.572 | 0.653 | 1.74E-07 |
| AT3G10400.1 | SNRNP31 | protein_coding | RNA recognition motif an | 15 | 38 | 23 | 30 | 22 | 29 | 32 | 93 | 59 | 65 | 58 | 79 | 88.9 | 61 | 67 | 1.572 | 0.652 | 0.00590103 |
| AT4G22592.1 | CPuORF27 | protein_coding | conserved peptide upstre | 54 | 78 | 44 | 107 | 77 | 81 | 114 | 192 | 113 | 232 | 204 | 220 | 176.15 | 140 | 219 | 1.571 | 0.652 | 0.00061585 |
| AT1G17050.1 | SPS2 | protein_coding | solanesyl diphosphate syı | 986 | 678 | 646 | 1471 | 974 | 817 | 2073 | 1666 | 1654 | 3191 | 2586 | 2222 | 1926.32 | 1798 | 2666 | 1.572 | 0.652 | 3.43E-06 |
| AT5G11030.2 | ALF4 | protein_coding | aberrant lateral root forn | 198 | 172 | 200 | 241 | 193 | 188 | 416 | 423 | 512 | 523 | 512 | 511 | 443.16 | 450 | 515 | 1.571 | 0.651 | 7.69E-09 |
| AT3G23570.1 | | protein_coding | alpha/beta-Hydrolases su | 293 | 237 | 157 | 543 | 440 | 320 | 616 | 582 | 402 | 1178 | 1168 | 870 | 844.33 | 533 | 1072 | 1.57 | 0.65 | 3.11E-05 |
| AT1G66730.1 | AtLIG6 | protein_coding | DNA LIGASE 6 | 341 | 345 | 433 | 489 | 469 | 497 | 717 | 848 | 1109 | 1061 | 1245 | 1352 | 850.32 | 891 | 1219 | 1.569 | 0.65 | 1.74E-05 |
| AT1G74880.1 | ndhO | protein_coding | NAD(P)H:plastoquinone c | 1103 | 931 | 878 | 1714 | 1518 | 1398 | 2319 | 2287 | 2248 | 3718 | 4030 | 3802 | 3753.4 | 2285 | 3850 | 1.57 | 0.65 | 2.08E-09 |
| AT4G36080.1 | | protein_coding | phosphotransferases, alc | 1766 | 1437 | 1412 | 2356 | 2127 | 1608 | 3712 | 3530 | 3615 | 5110 | 5647 | 4373 | 2694.89 | 3619 | 5043 | 1.568 | 0.649 | 7.32E-07 |
| AT5G47880.1 | ERF1-1 | protein_coding | eukaryotic release factor | 693 | 550 | 551 | 885 | 655 | 686 | 1457 | 1351 | 1411 | 1920 | 1739 | 1866 | 1712.32 | 1406 | 1842 | 1.568 | 0.649 | 4.03E-08 |
| AT1G77450.1 | anac032 | protein_coding | NAC domain containing p | 23 | 16 | 13 | 61 | 30 | 32 | 48 | 39 | 33 | 132 | 80 | 87 | 78.46 | 40 | 100 | 1.567 | 0.648 | 0.0034557 |
| AT2G32940.1 | AGO6 | protein_coding | Argonaute family protein | 76 | 50 | 51 | 76 | 77 | 76 | 160 | 123 | 131 | 165 | 204 | 207 | 144.77 | 138 | 192 | 1.567 | 0.648 | 0.00138233 |
| AT1G75240.1 | ZHD5 | protein_coding | homeobox protein 33 | 251 | 306 | 292 | 384 | 380 | 504 | 528 | 752 | 748 | 833 | 1009 | 1371 | 960.23 | 676 | 1071 | 1.567 | 0.648 | 0.00015821 |
| AT5G11230.1 | | protein_coding | Nucleotide-sugar transpo | 27 | 24 | 23 | 52 | 37 | 29 | 57 | 59 | 59 | 113 | 98 | 79 | 72.63 | 58 | 97 | 1.566 | 0.647 | 0.00580158 |
| AT2G37690.1 | | protein_coding | phosphoribosylaminoimid | 559 | 298 | 289 | 676 | 471 | 399 | 1175 | 732 | 740 | 1466 | 1250 | 1085 | 1036.57 | 882 | 1267 | 1.566 | 0.647 | 0.00019449 |
| AT1G15980.1 | PNSB1 | protein_coding | NDH-dependent cyclic ele | 3734 | 3566 | 3276 | 4943 | 4401 | 4289 | 7849 | 8761 | 8387 | 10722 | 11684 | 11664 | 10473.24 | 8332 | 11357 | 1.565 | 0.646 | 3.12E-08 |
| AT1G62750.1 | CPEFG | protein_coding | Translation elongation fa | 15696 | 13660 | 10239 | 18354 | 15928 | 14102 | 32994 | 33560 | 26212 | 39812 | 42285 | 38350 | 31133 | 30922 | 40149 | 1.562 | 0.643 | 1.11E-05 |
| AT1G08230.2 | GAT1 | protein_coding | Transmembrane amino a | 42 | 38 | 38 | 42 | 46 | 58 | 88 | 93 | 97 | 91 | 122 | 158 | 145.72 | 93 | 124 | 1.561 | 0.642 | 0.0009176 |
| AT5G64900.1 | PEP1 | protein_coding | precursor of peptide 1 | 26 | 18 | 12 | 21 | 16 | 10 | 55 | 44 | 31 | 46 | 42 | 27 | 53.5 | 43 | 38 | 1.56 | 0.641 | 0.00504273 |
| AT1G64105.1 | ANAC027 | protein_coding | NAC domain containing p | 53 | 62 | 28 | 78 | 57 | 51 | 111 | 152 | 72 | 169 | 151 | 139 | 147.31 | 112 | 153 | 1.559 | 0.641 | 0.00176564 |
| AT2G36530.1 | ENO2 | protein_coding | Enolase | 13410 | 8497 | 6753 | 18059 | 10712 | 10923 | 28189 | 20876 | 17288 | 39172 | 28438 | 29704 | 28535.67 | 22118 | 32438 | 1.559 | 0.641 | 0.00054677 |
| AT1G14280.1 | PKS2 | protein_coding | phytochrome kinase subs | 1097 | 962 | 1218 | 1568 | 1431 | 1590 | 2306 | 2363 | 3118 | 3401 | 3799 | 4324 | 2489.09 | 2596 | 3841 | 1.559 | 0.641 | 2.20E-05 |
| AT1G69310.1 | WRKY57 | protein_coding | WRKY DNA-binding prote | 40 | 23 | 23 | 76 | 39 | 52 | 84 | 57 | 59 | 165 | 104 | 141 | 131.33 | 67 | 137 | 1.558 | 0.64 | 0.00348216 |
| AT5G06790.1 | | protein_coding | | 90 | 83 | 66 | 92 | 78 | 46 | 189 | 204 | 169 | 200 | 207 | 125 | 154.56 | 187 | 177 | 1.558 | 0.64 | 0.00045584 |
| AT5G21280.1 | | protein_coding | hydroxyproline-rich glyco | 20 | 14 | 13 | 32 | 21 | 14 | 42 | 34 | 33 | 69 | 56 | 38 | 111.69 | 36 | 54 | 1.558 | 0.639 | 0.00029354 |
| AT2G46290.1 | | protein_coding | Transducin/WD40 repeat | 149 | 100 | 79 | 154 | 118 | 164 | 313 | 246 | 202 | 334 | 313 | 446 | 334.6 | 254 | 364 | 1.554 | 0.636 | 4.81E-05 |
| AT1G25220.2 | ASB1 | protein_coding | anthranilate synthase bet | 380 | 244 | 245 | 544 | 371 | 287 | 799 | 599 | 627 | 1180 | 985 | 780 | 896.34 | 675 | 982 | 1.554 | 0.636 | 3.43E-05 |
| AT3G10500.1 | NAC053 | protein_coding | NAC domain containing p | 111 | 90 | 94 | 192 | 147 | 129 | 233 | 221 | 241 | 416 | 390 | 351 | 268.31 | 232 | 386 | 1.554 | 0.636 | 3.05E-05 |
| AT1G65030.1 | | protein_coding | Transducin/WD40 repeat | 143 | 109 | 70 | 215 | 143 | 120 | 301 | 268 | 179 | 466 | 380 | 326 | 493.77 | 249 | 391 | 1.553 | 0.635 | 2.39E-05 |
| AT2G45510.1 | CYP704A2 | protein_coding | cytochrome P450, family | 105 | 44 | 80 | 189 | 122 | 118 | 221 | 108 | 205 | 410 | 324 | 321 | 211 | 178 | 352 | 1.552 | 0.634 | 0.00156678 |
| AT3G62870.1 | RPL7AB | protein_coding | Ribosomal protein L7Ae/l | 6449 | 4474 | 3144 | 6478 | 4807 | 4857 | 13556 | 10992 | 8049 | 14051 | 12761 | 13208 | 13386.15 | 10866 | 13340 | 1.552 | 0.634 | 0.0001854 |
| AT5G14800.1 | PROC1 | protein_coding | pyrroline-5- carboxylate ( | 769 | 558 | 468 | 785 | 644 | 692 | 1616 | 1371 | 1198 | 1703 | 1710 | 1882 | 1982.78 | 1395 | 1765 | 1.552 | 0.634 | 4.45E-06 |
| AT5G57480.1 | | protein_coding | P-loop containing nucleo | 17 | 19 | 20 | 31 | 24 | 38 | 36 | 47 | 51 | 67 | 64 | 103 | 63.92 | 45 | 78 | 1.551 | 0.633 | 0.00926427 |
| AT4G08870.1 | ARGAH2 | protein_coding | Arginase/deacetylase sup | 637 | 370 | 216 | 744 | 354 | 361 | 1339 | 909 | 553 | 1614 | 940 | 982 | 1139.22 | 934 | 1179 | 1.551 | 0.633 | 0.00810694 |
| AT2G23340.1 | ERF008 | protein_coding | DREB and EAR motif prot | 32 | 28 | 21 | 30 | 25 | 19 | 67 | 69 | 54 | 65 | 66 | 52 | 118.9 | 63 | 61 | 1.551 | 0.633 | 0.0006291 |
| AT5G11090.1 | | protein_coding | serine-rich protein-relate | 330 | 263 | 352 | 499 | 328 | 579 | 694 | 646 | 901 | 1082 | 871 | 1575 | 814.26 | 747 | 1176 | 1.55 | 0.632 | 0.00033647 |
| AT1G33110.1 | | protein_coding | MATE efflux family prote | 401 | 302 | 232 | 594 | 341 | 287 | 843 | 742 | 594 | 1288 | 905 | 780 | 835.04 | 726 | 991 | 1.55 | 0.632 | 0.00027503 |
| AT5G64930.1 | CPR5 | protein_coding | CPR5 protein, putative | 199 | 152 | 202 | 288 | 214 | 259 | 418 | 373 | 517 | 625 | 568 | 704 | 425.78 | 436 | 632 | 1.55 | 0.632 | 9.52E-06 |
| AT5G51730.1 | | protein_coding | RNA-binding (RRM/RBD/ | 228 | 130 | 161 | 268 | 197 | 197 | 479 | 319 | 412 | 581 | 523 | 536 | 475.47 | 403 | 547 | 1.55 | 0.632 | 3.04E-06 |
| AT5G10550.1 | GTE2 | protein_coding | global transcription facto | 1127 | 1061 | 977 | 1201 | 1067 | 1096 | 2369 | 2607 | 2501 | 2605 | 2833 | 2981 | 2707.97 | 2492 | 2806 | 1.549 | 0.631 | 4.43E-08 |
| AT2G37040.1 | PAL1 | protein_coding | PHE ammonia lyase 1 | 1947 | 1616 | 1142 | 2188 | 1611 | 1467 | 4093 | 3970 | 2924 | 4746 | 4277 | 3989 | 3271.41 | 3662 | 4337 | 1.548 | 0.63 | 1.12E-05 |
| AT5G08040.1 | TOM5 | protein_coding | mitochondrial import rec | 692 | 561 | 456 | 887 | 643 | 548 | 1455 | 1378 | 1167 | 1924 | 1707 | 1490 | 1485.98 | 1333 | 1707 | 1.546 | 0.629 | 3.20E-05 |
| AT1G66500.1 | PCFS1 | protein_coding | Pre-mRNA cleavage com | 49 | 36 | 33 | 46 | 36 | 50 | 103 | 88 | 84 | 100 | 96 | 136 | 97.88 | 92 | 111 | 1.546 | 0.628 | 0.00215888 |
| AT3G10880.1 | | protein_coding | | 55 | 41 | 34 | 70 | 64 | 43 | 116 | 101 | 87 | 152 | 170 | 117 | 110.43 | 101 | 146 | 1.545 | 0.628 | 0.00077066 |
| AT5G56030.2 | HSP90-2 | protein_coding | heat shock protein 81-2 | 8019 | 5875 | 4025 | 8203 | 6584 | 4994 | 16857 | 14434 | 10304 | 17793 | 17479 | 13581 | 12987.39 | 13865 | 16284 | 1.545 | 0.628 | 0.00020333 |
| AT5G04140.2 | GLU1 | protein_coding | glutamate synthase 1 | 37447 | 33904 | 34094 | 49152 | 43031 | 40384 | 78716 | 83296 | 87283 | 106615 | 114237 | 109822 | 77561.84 | 83098 | 110225 | 1.545 | 0.628 | 6.88E-05 |
| AT1G48850.1 | EMB1144 | protein_coding | chorismate synthase, put | 1494 | 962 | 775 | 1741 | 1220 | 1091 | 3140 | 2363 | 1984 | 3776 | 3239 | 2967 | 3252.25 | 2496 | 3327 | 1.545 | 0.627 | 3.04E-05 |
| AT1G19940.1 | AtGH9B5 | protein_coding | glycosyl hydrolase 9B5 | 44 | 34 | 40 | 58 | 82 | 46 | 92 | 84 | 102 | 126 | 218 | 125 | 119 | 93 | 156 | 1.544 | 0.626 | 0.00283415 |
| AT3G05020.1 | ACP1 | protein_coding | acyl carrier protein 1 | 307 | 295 | 414 | 578 | 530 | 603 | 645 | 725 | 1060 | 1254 | 1407 | 1640 | 1237.31 | 810 | 1434 | 1.544 | 0.626 | 0.00017206 |
| AT5G38530.1 | TSBtype2 | protein_coding | tryptophan synthase beta | 763 | 497 | 381 | 1160 | 622 | 427 | 1604 | 1221 | 975 | 2516 | 1651 | 1161 | 1494.02 | 1267 | 1776 | 1.542 | 0.625 | 0.0039571 |
| AT4G21170.1 | | protein_coding | Tetratricopeptide repeat | 78 | 66 | 64 | 110 | 61 | 69 | 164 | 162 | 164 | 239 | 162 | 188 | 158.65 | 163 | 196 | 1.542 | 0.625 | 0.00038838 |
| AT1G64200.1 | VHA-E3 | protein_coding | vacuolar H+-ATPase subu | 351 | 251 | 172 | 551 | 362 | 268 | 738 | 617 | 440 | 1195 | 961 | 729 | 1014.07 | 598 | 962 | 1.542 | 0.625 | 0.00029336 |
| AT5G07000.1 | SOT14 | protein_coding | sulfotransferase 2B | 335 | 243 | 304 | 459 | 306 | 458 | 704 | 597 | 778 | 996 | 812 | 1246 | 911.19 | 693 | 1018 | 1.542 | 0.625 | 1.41E-05 |
| AT5G16715.1 | EMB2247 | protein_coding | ATP binding;valine-tRNA | 2064 | 1506 | 1344 | 2418 | 1993 | 1711 | 4339 | 3700 | 3441 | 5245 | 5291 | 4653 | 4194.52 | 3827 | 5063 | 1.542 | 0.625 | 5.33E-07 |
| AT2G26790.1 | | protein_coding | Pentatricopeptide repeat | 62 | 47 | 39 | 73 | 60 | 44 | 130 | 115 | 100 | 158 | 159 | 120 | 109.54 | 115 | 146 | 1.541 | 0.624 | 0.0009919 |
| AT5G16320.1 | FRL1 | protein_coding | FRIGIDA like 1 | 90 | 92 | 62 | 105 | 93 | 83 | 189 | 226 | 159 | 228 | 247 | 226 | 180.14 | 191 | 234 | 1.541 | 0.624 | 0.0007207 |
| AT2G34260.1 | | protein_coding | transducin family protein | 255 | 162 | 143 | 241 | 203 | 153 | 536 | 398 | 366 | 523 | 539 | 416 | 479.29 | 433 | 493 | 1.541 | 0.624 | 4.31E-05 |
| AT1G09780.1 | PGM1 | protein_coding | Phosphoglycerate mutase | 5661 | 2971 | 2814 | 5987 | 3820 | 3538 | 11900 | 7299 | 7204 | 12986 | 10141 | 9621 | 9014.22 | 8801 | 10916 | 1.54 | 0.623 | 0.00038158 |
| AT3G49680.1 | BCAT3 | protein_coding | branched-chain aminotra | 2805 | 2266 | 1704 | 3758 | 3388 | 2421 | 5896 | 5567 | 4362 | 8151 | 8994 | 6584 | 7669.42 | 5275 | 7910 | 1.54 | 0.623 | 1.61E-05 |
| AT2G32530.1 | CSLB3 | protein_coding | cellulose synthase-like B3 | 260 | 145 | 111 | 236 | 134 | 91 | 547 | 356 | 284 | 512 | 356 | 247 | 327.79 | 396 | 372 | 1.538 | 0.621 | 0.00625312 |
| AT1G01720.1 | NAC002 | protein_coding | NAC (No Apical Meristem | 99 | 69 | 83 | 178 | 77 | 117 | 208 | 170 | 212 | 386 | 204 | 318 | 246.25 | 197 | 303 | 1.538 | 0.621 | 0.00089445 |
| AT1G64900.1 | CYP89A2 | protein_coding | cytochrome P450, family | 729 | 566 | 416 | 1037 | 583 | 661 | 1532 | 1391 | 1065 | 2249 | 1548 | 1798 | 1890.9 | 1329 | 1865 | 1.538 | 0.621 | 6.80E-05 |
| AT5G19900.1 | | protein_coding | PRLI-interacting factor, p | 199 | 205 | 202 | 311 | 271 | 266 | 418 | 504 | 517 | 675 | 719 | 723 | 783.4 | 480 | 706 | 1.538 | 0.621 | 1.01E-08 |
| AT4G33220.1 | PME44 | protein_coding | pectin methylesterase 44 | 2978 | 2676 | 1782 | 3692 | 2806 | 2530 | 6260 | 6574 | 4562 | 8008 | 7449 | 6880 | 6370.88 | 5799 | 7446 | 1.537 | 0.62 | 8.02E-06 |

| | | | | | | | | | | | | | | | | | | | | | |
|---|---|---|---|---|---|---|---|---|---|---|---|---|---|---|---|---|---|---|---|---|---|
| AT4G02530.1 | | protein_coding | chloroplast thylakoid lum | 4187 | 3656 | 3369 | 5881 | 5130 | 4632 | 8801 | 8982 | 8625 | 12756 | 13619 | 12596 | 12210.18 | 8803 | 12990 | 1.536 | 0.62 | 1.75E-07 |
| AT1G75100.1 | JAC1 | protein_coding | J-domain protein require | 1316 | 983 | 966 | 1515 | 1154 | 1122 | 2766 | 2415 | 2473 | 3286 | 3064 | 3051 | 2252.81 | 2551 | 3134 | 1.537 | 0.62 | 1.67E-07 |
| AT4G22590.1 | TPPG | protein_coding | Haloacid dehalogenase-li | 69 | 73 | 61 | 126 | 99 | 64 | 145 | 179 | 156 | 273 | 263 | 174 | 192.2 | 160 | 237 | 1.536 | 0.619 | 0.00175498 |
| AT3G22660.1 | | protein_coding | rRNA processing protein- | 269 | 193 | 120 | 236 | 191 | 179 | 565 | 474 | 307 | 512 | 507 | 487 | 777.62 | 449 | 502 | 1.536 | 0.619 | 0.00013695 |
| AT5G67370.1 | | protein_coding | Protein of unknown funct | 1034 | 695 | 530 | 1804 | 1178 | 856 | 2174 | 1707 | 1357 | 3913 | 3127 | 2328 | 2774.64 | 1746 | 3123 | 1.535 | 0.618 | 0.00060729 |
| AT1G79710.1 | | protein_coding | Major facilitator superfar | 131 | 107 | 85 | 198 | 115 | 90 | 275 | 263 | 218 | 429 | 305 | 245 | 262.86 | 252 | 326 | 1.535 | 0.618 | 0.00052313 |
| AT2G06040.1 | | protein_coding | | 159 | 150 | 131 | 195 | 171 | 170 | 334 | 369 | 335 | 423 | 454 | 462 | 341.43 | 346 | 446 | 1.535 | 0.618 | 5.39E-07 |
| AT1G15950.1 | CCR1 | protein_coding | cinnamoyl coa reductase | 937 | 944 | 573 | 1560 | 1125 | 1142 | 1970 | 2319 | 1467 | 3384 | 2987 | 3106 | 3033.29 | 1919 | 3159 | 1.531 | 0.615 | 1.81E-05 |
| AT2G21200.1 | | protein_coding | SAUR-like auxin-responsi | 36 | 22 | 45 | 57 | 49 | 50 | 76 | 54 | 115 | 124 | 130 | 136 | 92.9 | 82 | 130 | 1.531 | 0.614 | 0.00830788 |
| AT4G39550.1 | | protein_coding | Galactose oxidase/kelch r | 35 | 35 | 26 | 61 | 44 | 65 | 74 | 86 | 67 | 132 | 117 | 177 | 105.77 | 76 | 142 | 1.53 | 0.614 | 0.0009524 |
| AT2G40475.1 | | protein_coding | | 332 | 296 | 207 | 437 | 537 | 335 | 698 | 727 | 530 | 948 | 1426 | 911 | 938.35 | 652 | 1095 | 1.531 | 0.614 | 0.00041283 |
| AT1G16280.1 | RH36 | protein_coding | RNA helicase 36 | 259 | 169 | 180 | 283 | 232 | 189 | 544 | 415 | 461 | 614 | 616 | 514 | 541.56 | 473 | 581 | 1.531 | 0.614 | 5.73E-07 |
| AT1G09100.1 | RPT5B | protein_coding | 26S proteasome AAA-ATF | 277 | 209 | 236 | 301 | 275 | 270 | 582 | 513 | 604 | 653 | 730 | 734 | 615.41 | 566 | 706 | 1.531 | 0.614 | 2.72E-08 |
| AT3G56940.1 | CRD1 | protein_coding | dicarboxylate diiron prot | 41316 | 43301 | 36868 | 54009 | 51417 | 52994 | 86849 | 106383 | 94384 | 117151 | 136500 | 144114 | 122275.17 | 95872 | 132588 | 1.53 | 0.613 | 0.00055052 |
| AT3G47780.1 | ABCA7 | protein_coding | ABC2 homolog 6 | 99 | 78 | 63 | 106 | 71 | 53 | 208 | 192 | 161 | 230 | 188 | 144 | 160.19 | 187 | 187 | 1.529 | 0.613 | 0.0005495 |
| AT5G20020.1 | RAN2 | protein_coding | RAS-related GTP-binding | 2301 | 1439 | 1176 | 2947 | 1890 | 1782 | 4837 | 3535 | 3011 | 6392 | 5018 | 4846 | 4982 | 3794 | 5419 | 1.529 | 0.612 | 0.0001705 |
| AT5G65350.1 | HTR11 | protein_coding | histone 3 11 | 13 | 18 | 15 | 31 | 27 | 12 | 27 | 44 | 38 | 67 | 72 | 33 | 53.54 | 36 | 57 | 1.528 | 0.611 | 0.00971378 |
| AT1G69410.1 | ELF5A-3 | protein_coding | eukaryotic elongation fac | 588 | 382 | 334 | 766 | 510 | 474 | 1236 | 939 | 855 | 1662 | 1354 | 1289 | 1281.89 | 1010 | 1435 | 1.528 | 0.611 | 4.39E-05 |
| AT2G30580.1 | DRIP2 | protein_coding | DREB2A-interacting prote | 140 | 100 | 130 | 211 | 145 | 162 | 294 | 246 | 333 | 458 | 385 | 441 | 309.58 | 291 | 428 | 1.527 | 0.611 | 2.64E-05 |
| AT1G07260.1 | UGT71C3 | protein_coding | UDP-glucosyl transferase | 15 | 24 | 18 | 23 | 27 | 30 | 32 | 59 | 46 | 50 | 72 | 82 | 63.42 | 46 | 68 | 1.527 | 0.61 | 0.00499313 |
| AT5G04590.1 | SIR | protein_coding | sulfite reductase | 9120 | 6603 | 4816 | 11357 | 8592 | 6908 | 19171 | 16222 | 12329 | 24634 | 22810 | 18786 | 16879.72 | 15907 | 22077 | 1.526 | 0.61 | 0.0001397 |
| AT3G12140.3 | EML1 | protein_coding | Emsy N Terminus (ENT)/ | 310 | 248 | 281 | 354 | 294 | 330 | 652 | 609 | 719 | 768 | 781 | 897 | 740.24 | 660 | 815 | 1.526 | 0.61 | 3.49E-08 |
| AT1G08300.1 | NVL | protein_coding | no vein-like | 98 | 71 | 93 | 115 | 95 | 82 | 206 | 174 | 238 | 249 | 252 | 223 | 209.31 | 206 | 241 | 1.526 | 0.609 | 0.00265317 |
| AT2G42760.1 | | protein_coding | | 50 | 41 | 36 | 70 | 47 | 44 | 105 | 101 | 92 | 152 | 125 | 120 | 116.67 | 99 | 132 | 1.526 | 0.609 | 0.00029104 |
| AT2G46040.1 | ARID1 | protein_coding | ARID/BRIGHT DNA-bindir | 205 | 107 | 125 | 190 | 164 | 151 | 431 | 263 | 320 | 412 | 435 | 411 | 319.15 | 338 | 419 | 1.523 | 0.607 | 5.15E-05 |
| AT5G19850.1 | | protein_coding | alpha/beta-Hydrolases su | 404 | 261 | 228 | 702 | 405 | 322 | 849 | 641 | 584 | 1523 | 1075 | 876 | 922.1 | 691 | 1158 | 1.522 | 0.606 | 0.00046447 |
| AT5G23810.1 | AAP7 | protein_coding | amino acid permease 7 | 73 | 42 | 51 | 78 | 34 | 39 | 153 | 103 | 131 | 169 | 90 | 106 | 119.77 | 129 | 122 | 1.521 | 0.605 | 0.00750411 |
| AT5G17860.1 | CCX1 | protein_coding | calcium exchanger 7 | 114 | 107 | 107 | 169 | 84 | 160 | 240 | 263 | 274 | 367 | 223 | 435 | 251.27 | 259 | 342 | 1.521 | 0.605 | 0.00108253 |
| AT1G11840.6 | ATGLX1 | protein_coding | glyoxalase I homolog | 2994 | 2157 | 1641 | 4232 | 3622 | 2431 | 6294 | 5299 | 4201 | 9180 | 9616 | 6611 | 6990.28 | 5265 | 8469 | 1.521 | 0.605 | 0.00044287 |
| AT4G38950.1 | | protein_coding | ATP binding microtubule | 536 | 528 | 445 | 585 | 659 | 590 | 1127 | 1297 | 1139 | 1269 | 1749 | 1604 | 1314.29 | 1188 | 1541 | 1.521 | 0.605 | 6.53E-06 |
| AT1G17420.1 | LOX3 | protein_coding | lipoxygenase 3 | 214 | 156 | 92 | 200 | 143 | 121 | 450 | 383 | 236 | 434 | 380 | 329 | 376.78 | 356 | 381 | 1.52 | 0.604 | 0.00072044 |
| AT1G55920.1 | SAT1 | protein_coding | serine acetyltransferase 2 | 420 | 256 | 185 | 604 | 302 | 272 | 883 | 629 | 474 | 1310 | 802 | 740 | 770.36 | 662 | 951 | 1.518 | 0.603 | 0.0039652 |
| AT3G48870.1 | CLPC2 | protein_coding | Clp ATPase | 4256 | 3317 | 2619 | 4620 | 3521 | 3235 | 8946 | 8149 | 6705 | 10021 | 9347 | 8797 | 8385.26 | 7933 | 9388 | 1.519 | 0.603 | 3.33E-06 |
| AT2G29340.1 | | protein_coding | NAD-dependent epimera | 601 | 282 | 340 | 762 | 392 | 434 | 1263 | 693 | 870 | 1653 | 1041 | 1180 | 1072.84 | 942 | 1291 | 1.517 | 0.602 | 0.0016743 |
| AT1G31835.1 | | protein_coding | | 2416 | 2087 | 1998 | 2435 | 2276 | 1894 | 5079 | 5127 | 5115 | 5282 | 6042 | 5151 | 2731.49 | 5107 | 5492 | 1.518 | 0.602 | 0.00036135 |
| AT3G14570.1 | CALS8 | protein_coding | glucan synthase-like 4 | 191 | 177 | 181 | 256 | 278 | 215 | 401 | 435 | 463 | 555 | 738 | 585 | 418.54 | 433 | 626 | 1.518 | 0.602 | 9.38E-06 |
| AT4G34350.1 | ISPH | protein_coding | 4-hydroxy-3-methylbut-2 | 6852 | 6234 | 6438 | 8350 | 6967 | 7834 | 14403 | 15316 | 16482 | 18112 | 18496 | 21304 | 17622.16 | 15400 | 19304 | 1.518 | 0.602 | 5.49E-06 |
| AT3G56170.1 | CAN1 | protein_coding | Ca-2+ dependent nucleas | 103 | 82 | 108 | 155 | 126 | 172 | 217 | 201 | 276 | 336 | 335 | 468 | 294.61 | 231 | 380 | 1.517 | 0.601 | 8.73E-05 |
| AT3G55800.1 | SBPASE | protein_coding | sedoheptulose-bisphosph | 20746 | 18169 | 16409 | 25159 | 22451 | 22261 | 43610 | 44638 | 42008 | 54572 | 59602 | 60537 | 58084.63 | 43419 | 58237 | 1.517 | 0.601 | 6.49E-05 |
| AT5G61220.1 | | protein_coding | LYR family of Fe/S cluster | 220 | 184 | 128 | 270 | 222 | 180 | 462 | 452 | 328 | 586 | 589 | 489 | 529.21 | 414 | 555 | 1.517 | 0.601 | 1.16E-05 |
| AT1G62030.1 | | protein_coding | Cysteine/Histidine-rich C1 | 186 | 191 | 150 | 271 | 249 | 227 | 391 | 469 | 384 | 588 | 661 | 617 | 354.49 | 415 | 622 | 1.517 | 0.601 | 7.24E-06 |
| AT1G06690.1 | | protein_coding | NAD(P)-linked oxidoredu | 799 | 794 | 693 | 943 | 851 | 802 | 1680 | 1951 | 1774 | 2045 | 2259 | 2181 | 2660.3 | 1802 | 2162 | 1.516 | 0.601 | 4.89E-07 |
| AT5G06530.2 | ABCG22 | protein_coding | ABC-2 type transporter fa | 346 | 251 | 236 | 534 | 390 | 475 | 727 | 617 | 604 | 1158 | 1035 | 1292 | 669.4 | 649 | 1162 | 1.516 | 0.6 | 1.72E-05 |
| AT4G10450.1 | RPL9D | protein_coding | Ribosomal protein L6 fam | 1013 | 628 | 531 | 1082 | 768 | 686 | 2129 | 1543 | 1359 | 2347 | 2039 | 1866 | 2031.28 | 1677 | 2084 | 1.515 | 0.599 | 0.00022085 |
| AT4G29410.1 | RPL28C | protein_coding | Ribosomal L28e protein f | 2036 | 1411 | 1200 | 2335 | 1770 | 1645 | 4280 | 3467 | 3072 | 5065 | 4699 | 4473 | 4285.99 | 3606 | 4746 | 1.515 | 0.599 | 2.72E-05 |
| AT2G15980.1 | | protein_coding | Tetratricopeptide repeat | 63 | 31 | 34 | 42 | 34 | 44 | 132 | 76 | 87 | 91 | 90 | 120 | 116.91 | 98 | 100 | 1.513 | 0.598 | 0.00197687 |
| AT1G20140.1 | ASK4 | protein_coding | SKP1-like 4 | 65 | 57 | 50 | 103 | 87 | 117 | 137 | 140 | 128 | 223 | 231 | 318 | 235.48 | 135 | 257 | 1.514 | 0.598 | 0.00142662 |
| AT1G75330.1 | OTC | protein_coding | ornithine carbamoyltrans | 1073 | 700 | 586 | 1333 | 889 | 885 | 2256 | 1720 | 1500 | 2891 | 2360 | 2407 | 2836.35 | 1825 | 2553 | 1.513 | 0.598 | 4.50E-05 |
| AT5G46390.2 | CTPA1 | protein_coding | Peptidase S41 family prot | 603 | 482 | 401 | 758 | 587 | 489 | 1268 | 1184 | 1027 | 1644 | 1558 | 1330 | 1299.55 | 1160 | 1511 | 1.513 | 0.598 | 2.36E-06 |
| AT5G24760.1 | | protein_coding | GroES-like zinc-binding de | 141 | 124 | 119 | 198 | 184 | 176 | 296 | 305 | 305 | 429 | 488 | 479 | 392.37 | 302 | 465 | 1.513 | 0.598 | 1.49E-07 |
| AT3G21755.1 | | other_rna | other RNA | 63 | 91 | 50 | 95 | 104 | 107 | 132 | 224 | 128 | 206 | 276 | 291 | 226.73 | 161 | 258 | 1.512 | 0.597 | 0.00137943 |
| AT5G20000.1 | RPT6B | protein_coding | AAA-type ATPase family p | 432 | 345 | 307 | 566 | 437 | 439 | 908 | 848 | 786 | 1228 | 1160 | 1194 | 1140.9 | 847 | 1194 | 1.513 | 0.597 | 5.76E-08 |
| AT4G26110.1 | NAP1 | protein_coding | nucleosome assembly pro | 1985 | 1171 | 1085 | 2109 | 1526 | 1414 | 4173 | 2877 | 2778 | 4575 | 4051 | 3845 | 3283.02 | 3276 | 4157 | 1.511 | 0.596 | 8.67E-05 |
| AT5G19600.1 | SULTR3 | protein_coding | sulfate transporter 3;5 | 156 | 146 | 125 | 248 | 183 | 202 | 328 | 359 | 320 | 538 | 486 | 549 | 396.72 | 336 | 524 | 1.511 | 0.596 | 6.36E-05 |
| AT1G28680.1 | | protein_coding | HXXXD-type acyl-transfer | 216 | 170 | 178 | 327 | 259 | 275 | 454 | 418 | 456 | 709 | 688 | 748 | 709.12 | 443 | 715 | 1.512 | 0.596 | 6.84E-09 |
| AT1G19920.1 | APS2 | protein_coding | Pseudouridine synthase/a | 2066 | 1386 | 1174 | 2945 | 2290 | 1649 | 4343 | 3405 | 3006 | 6388 | 6079 | 4484 | 4539.53 | 3585 | 5650 | 1.511 | 0.595 | 8.34E-05 |
| AT3G56100.1 | | protein_coding | glycine-rich protein / olec | 181 | 202 | 196 | 237 | 225 | 283 | 380 | 496 | 502 | 514 | 597 | 770 | 545.58 | 459 | 627 | 1.51 | 0.595 | 7.23E-05 |
| AT5G47250.1 | | protein_coding | Plant protein of unknowr | 1253 | 1164 | 1210 | 1898 | 1382 | 1435 | 2634 | 2860 | 3098 | 4117 | 3669 | 3902 | 3091.47 | 2864 | 3896 | 1.511 | 0.595 | 7.19E-07 |
| AT5G59750.2 | RIBA3 | protein_coding | DHBP synthase RibB-like | 653 | 592 | 526 | 1147 | 812 | 768 | 1373 | 1454 | 1347 | 2488 | 2156 | 2089 | 1853.16 | 1391 | 2244 | 1.511 | 0.595 | 2.24E-07 |
| AT1G76450.1 | PPD3 | protein_coding | Photosystem II reaction c | 2361 | 2104 | 2049 | 3087 | 2738 | 2595 | 4963 | 5169 | 5246 | 6696 | 7269 | 7057 | 6526.5 | 5126 | 7007 | 1.51 | 0.595 | 1.06E-07 |
| AT4G02700.1 | SULTR3 | protein_coding | sulfate transporter 3;2 | 123 | 82 | 83 | 174 | 94 | 158 | 259 | 201 | 212 | 377 | 250 | 430 | 241.84 | 224 | 352 | 1.509 | 0.594 | 0.00207171 |
| AT1G25145.1 | LPXC4 | protein_coding | UDP-3-O-acyl N-acetylgly | 55 | 54 | 62 | 73 | 59 | 87 | 116 | 133 | 159 | 158 | 157 | 237 | 238.71 | 136 | 184 | 1.509 | 0.594 | 6.89E-05 |
| AT2G28570.1 | | protein_coding | | 13 | 11 | 13 | 20 | 15 | 16 | 27 | 27 | 33 | 43 | 40 | 44 | 45.19 | 29 | 42 | 1.509 | 0.593 | 0.0066046 |
| AT3G47950.1 | AHA4 | protein_coding | H(+)-ATPase 4 | 50 | 37 | 48 | 46 | 35 | 38 | 105 | 91 | 123 | 100 | 93 | 103 | 92.06 | 106 | 99 | 1.507 | 0.592 | 0.00153903 |
| AT1G64060.1 | RBOHF | protein_coding | respiratory burst oxidase | 269 | 222 | 236 | 456 | 381 | 232 | 565 | 545 | 604 | 989 | 1011 | 631 | 579.6 | 571 | 877 | 1.508 | 0.592 | 6.77E-05 |
| AT3G16470.1 | JAL35 | protein_coding | Mannose-binding lectin s | 707 | 652 | 469 | 1433 | 1382 | 950 | 1486 | 1602 | 1201 | 3108 | 3669 | 2583 | 2287.72 | 1430 | 3120 | 1.506 | 0.591 | 4.76E-05 |
| AT1G15700.1 | ATPC2 | protein_coding | ATPase, F1 complex, gam | 126 | 76 | 117 | 164 | 119 | 111 | 265 | 187 | 300 | 356 | 316 | 302 | 242.65 | 251 | 325 | 1.506 | 0.59 | 0.00033117 |
| AT5G18040.1 | | protein_coding | | 79 | 70 | 49 | 91 | 93 | 79 | 166 | 172 | 125 | 197 | 247 | 215 | 209.8 | 154 | 220 | 1.504 | 0.589 | 0.00054037 |
| AT4G20340.1 | | protein_coding | Transcription factor TFIIE | 156 | 110 | 136 | 217 | 167 | 135 | 328 | 270 | 348 | 471 | 443 | 367 | 380.61 | 315 | 427 | 1.503 | 0.588 | 3.03E-06 |
| AT3G52210.3 | | protein_coding | S-adenosyl-L-methionine- | 158 | 157 | 164 | 263 | 198 | 206 | 332 | 386 | 420 | 570 | 526 | 560 | 470.73 | 379 | 552 | 1.503 | 0.588 | 3.76E-07 |
| AT3G25590.1 | | protein_coding | | 76 | 51 | 47 | 73 | 52 | 53 | 160 | 125 | 120 | 158 | 138 | 144 | 118.01 | 135 | 147 | 1.502 | 0.587 | 0.0016769 |
| AT4G16630.1 | RH28 | protein_coding | DEA(D/H)-box RNA helica | 630 | 413 | 302 | 544 | 485 | 411 | 1324 | 1015 | 773 | 1180 | 1288 | 1118 | 1203.03 | 1037 | 1195 | 1.503 | 0.587 | 0.00031032 |
| AT2G38760.1 | ANN3 | protein_coding | annexin 3 | 135 | 128 | 69 | 150 | 142 | 74 | 284 | 314 | 177 | 325 | 377 | 201 | 288.99 | 258 | 301 | 1.501 | 0.586 | 0.00257819 |
| AT4G30250.1 | | protein_coding | P-loop containing nucleo | 47 | 43 | 33 | 63 | 57 | 29 | 99 | 106 | 84 | 137 | 151 | 79 | 119.8 | 96 | 122 | 1.501 | 0.585 | 0.00799354 |
| AT4G03030.1 | OR23 | protein_coding | Galactose oxidase/kelch r | 218 | 220 | 285 | 345 | 272 | 397 | 458 | 540 | 730 | 748 | 722 | 1080 | 715.58 | 576 | 850 | 1.5 | 0.585 | 0.00027592 |
| AT1G75280.1 | | protein_coding | NmrA-like negative trans | 868 | 829 | 594 | 1354 | 1180 | 825 | 1825 | 2037 | 1521 | 2937 | 3133 | 2244 | 2378.68 | 1794 | 2771 | 1.5 | 0.585 | 3.60E-05 |
| AT4G32320.1 | APX6 | protein_coding | ascorbate peroxidase 6 | 258 | 266 | 217 | 250 | 258 | 287 | 542 | 654 | 556 | 542 | 685 | 780 | 812.46 | 584 | 669 | 1.5 | 0.585 | 9.85E-06 |
| AT4G39540.3 | SK2 | protein_coding | shikimate kinase 2 | 520 | 427 | 499 | 811 | 676 | 604 | 1093 | 1049 | 1277 | 1759 | 1795 | 1643 | 1478.56 | 1140 | 1732 | 1.5 | 0.585 | 2.44E-06 |
| AT2G12490.1 | | transposable_elen | transposable element ger | 416 | 376 | 490 | 676 | 610 | 569 | 874 | 924 | 1254 | 1466 | 1619 | 1547 | 720 | 1017 | 1544 | 1.499 | 0.584 | 9.99E-05 |
| AT3G57300.2 | INO80 | protein_coding | INO80 ortholog | 1498 | 1161 | 1286 | 2047 | 1840 | 1732 | 3149 | 2852 | 3292 | 4440 | 4885 | 4710 | 2979.67 | 3098 | 4678 | 1.498 | 0.583 | 1.40E-06 |
| AT1G02140.1 | MEE63 | protein_coding | mago nashi family protei | 1444 | 1265 | 1029 | 2047 | 1488 | 1470 | 3035 | 3108 | 2634 | 4440 | 3950 | 3998 | 3412.11 | 2926 | 4129 | 1.498 | 0.583 | 7.79E-07 |
| AT5G02500.1 | MED37E | protein_coding | heat shock cognate prote | 26316 | 24322 | 14820 | 30552 | 26259 | 27109 | 55318 | 59755 | 37940 | 66270 | 69712 | 73721 | 57386.38 | 51004 | 69901 | 1.497 | 0.582 | 0.00137928 |
| AT5G61030.1 | RBG3 | protein_coding | glycine-rich RNA-binding | 123 | 79 | 65 | 107 | 100 | 85 | 259 | 194 | 166 | 232 | 265 | 231 | 633.68 | 206 | 243 | 1.497 | 0.582 | 0.00016587 |
| AT1G74560.3 | NRP1 | protein_coding | NAP1-related protein 1 | 1430 | 857 | 862 | 1577 | 1255 | 1095 | 3006 | 2105 | 2207 | 3421 | 3332 | 2978 | 2827.87 | 2439 | 3244 | 1.497 | 0.582 | 0.00015891 |
| AT1G25098.1 | | other_rna | other RNA | 1363 | 1286 | 1582 | 2107 | 2076 | 1955 | 2865 | 3159 | 4050 | 4570 | 5511 | 5317 | 6694.47 | 3358 | 5133 | 1.497 | 0.582 | 3.74E-05 |
| AT1G17990.1 | | protein_coding | FMN-linked oxidoreducta | 173 | 213 | 210 | 323 | 286 | 331 | 364 | 523 | 538 | 701 | 759 | 900 | 759.19 | 475 | 787 | 1.497 | 0.582 | 3.58E-05 |
| AT4G24460.1 | CLT2 | protein_coding | CRT (chloroquine-resistar | 210 | 212 | 144 | 301 | 299 | 250 | 441 | 521 | 369 | 653 | 794 | 680 | 917.97 | 444 | 709 | 1.497 | 0.582 | 6.93E-06 |
| AT3G56160.1 | BASS4 | protein_coding | Sodium Bile acid symport | 807 | 758 | 621 | 1066 | 1010 | 778 | 1696 | 1862 | 1590 | 2312 | 2681 | 2116 | 2752.24 | 1716 | 2370 | 1.497 | 0.582 | 4.53E-06 |
| AT4G02450.1 | | protein_coding | HSP20-like chaperones su | 285 | 229 | 174 | 422 | 299 | 181 | 599 | 563 | 445 | 915 | 794 | 492 | 2869.88 | 536 | 734 | 1.496 | 0.581 | 0.0005427 |
| AT5G22650.1 | HDT2 | protein_coding | histone deacetylase 2B | 2627 | 1765 | 1338 | 2864 | 2234 | 1961 | 5522 | 4336 | 3425 | 6212 | 5931 | 5333 | 5898.6 | 4428 | 5825 | 1.496 | 0.581 | 0.00016278 |
| AT4G39710.1 | PNSL4 | protein_coding | FK506-binding protein 16 | 848 | 1000 | 732 | 1294 | 1295 | 1222 | 1783 | 2457 | 1874 | 2807 | 3438 | 3323 | 3317.53 | 2038 | 3189 | 1.495 | 0.581 | 1.51E-05 |
| AT1G03630.1 | PORC | protein_coding | protochlorophyllide oxidc | 4826 | 4517 | 2873 | 4545 | 4726 | 3678 | 10145 | 11097 | 7355 | 9859 | 12546 | 10002 | 11534.92 | 9532 | 10802 | 1.495 | 0.58 | 0.00015037 |
| AT5G59980.2 | | protein_coding | Polymerase/histidinol ph | 176 | 122 | 103 | 148 | 136 | 121 | 370 | 300 | 264 | 321 | 361 | 329 | 284.65 | 311 | 337 | 1.495 | 0.58 | 8.75E-05 |
| AT1G06640.1 | | protein_coding | 2-oxoglutarate (2OG) anc | 352 | 343 | 262 | 835 | 661 | 609 | 740 | 843 | 671 | 1811 | 1755 | 1656 | 1381.79 | 751 | 1741 | 1.495 | 0.58 | 4.32E-06 |
| AT3G56260.2 | | protein_coding | | 33 | 24 | 23 | 56 | 39 | 27 | 69 | 59 | 59 | 121 | 104 | 73 | 93.55 | 62 | 99 | 1.494 | 0.579 | 0.00355908 |
| AT4G03440.1 | | protein_coding | Ankyrin repeat family prc | 57 | 45 | 30 | 73 | 43 | 49 | 120 | 111 | 77 | 158 | 114 | 133 | 117.08 | 103 | 135 | 1.494 | 0.579 | 0.00277345 |
| AT3G03370.2 | | protein_coding | | 34 | 26 | 23 | 76 | 67 | 52 | 71 | 64 | 59 | 165 | 178 | 141 | 108.01 | 65 | 161 | 1.493 | 0.578 | 0.00131721 |
| AT4G33010.1 | GLDP1 | protein_coding | glycine decarboxylase P-p | 22451 | 21277 | 16168 | 28748 | 27993 | 21852 | 47194 | 52274 | 41391 | 62357 | 74315 | 59425 | 64682.6 | 46953 | 65366 | 1.492 | 0.578 | 0.00063755 |
| AT3G48310.1 | CYP71A22 | protein_coding | cytochrome P450, family | 269 | 251 | 237 | 397 | 234 | 236 | 565 | 617 | 607 | 861 | 621 | 642 | 603.53 | 596 | 708 | 1.493 | 0.578 | 2.38E-05 |
| AT3G23710.1 | TIC22L | protein_coding | Tic22-like family protein | 202 | 158 | 157 | 224 | 163 | 154 | 425 | 388 | 402 | 486 | 433 | 419 | 471.26 | 405 | 446 | 1.493 | 0.578 | 1.62E-05 |
| AT4G00490.1 | BAM2 | protein_coding | beta-amylase 2 | 567 | 499 | 433 | 915 | 707 | 647 | 1192 | 1226 | 1109 | 1985 | 1877 | 1759 | 1445.5 | 1176 | 1874 | 1.493 | 0.578 | 1.27E-06 |
| AT5G03390.1 | | protein_coding | Protein of unknown funct | 58 | 41 | 26 | 84 | 53 | 60 | 122 | 101 | 67 | 182 | 141 | 163 | 190.21 | 97 | 162 | 1.492 | 0.577 | 0.00054213 |
| AT2G21860.1 | | protein_coding | violaxanthin de-epoxidas | 1167 | 907 | 658 | 1295 | 1064 | 932 | 2453 | 2228 | 1685 | 2809 | 2825 | 2535 | 2681.13 | 2122 | 2723 | 1.492 | 0.577 | 1.90E-05 |
| AT1G07370.1 | PCNA | protein_coding | proliferating cellular nucl | 367 | 342 | 228 | 465 | 400 | 357 | 771 | 840 | 584 | 1009 | 1062 | 971 | 981.92 | 732 | 1014 | 1.492 | 0.577 | 6.85E-06 |
| AT3G11170.1 | FAD7 | protein_coding | fatty acid desaturase 7 | 4862 | 3708 | 3624 | 6070 | 5101 | 4198 | 10220 | 9110 | 9278 | 13166 | 13542 | 11416 | 11750.53 | 9536 | 12708 | 1.492 | 0.577 | 3.65E-06 |
| AT1G03760.1 | | protein_coding | Prefoldin chaperone subu | 127 | 111 | 111 | 164 | 105 | 134 | 267 | 273 | 284 | 356 | 279 | 364 | 277.29 | 275 | 333 | 1.49 | 0.576 | 0.00019699 |
| AT4G19870.1 | | protein_coding | Galactose oxidase/kelch r | 62 | 53 | 62 | 69 | 75 | 75 | 130 | 130 | 159 | 150 | 199 | 204 | 178.51 | 140 | 184 | 1.49 | 0.575 | 0.00025415 |
| AT1G27050.1 | ATHB54 | protein_coding | homeobox protein 54 | 97 | 78 | 56 | 99 | 63 | 95 | 204 | 167 | 143 | 215 | 167 | 258 | 301.82 | 180 | 213 | 1.489 | 0.575 | 0.00010948 |
| AT1G50840.1 | POLIA | protein_coding | polymerase gamma 2 | 1191 | 874 | 714 | 1280 | 1016 | 750 | 2504 | 2147 | 1828 | 2776 | 2697 | 2040 | 2177.08 | 2160 | 2504 | 1.49 | 0.575 | 4.14E-05 |
| AT1G24793.1 | LPXC4 | protein_coding | UDP-3-O-acyl N-acetylgly | 60 | 76 | 45 | 62 | 89 | 81 | 126 | 187 | 115 | 134 | 236 | 220 | 263.56 | 143 | 197 | 1.488 | 0.574 | 0.00203101 |
| AT5G62130.2 | | protein_coding | Per1-like family protein | 129 | 120 | 70 | 167 | 136 | 72 | 271 | 295 | 179 | 362 | 361 | 196 | 361.81 | 248 | 306 | 1.489 | 0.574 | 0.00143804 |
| AT3G04840.1 | RPS3AA | protein_coding | Ribosomal protein S3Ae | 8625 | 6063 | 4229 | 9030 | 6377 | 5748 | 18130 | 14896 | 10827 | 19587 | 16929 | 15631 | 17064.25 | 14618 | 17382 | 1.489 | 0.574 | 0.00091794 |
| AT3G44430.1 | | protein_coding | | 325 | 337 | 280 | 407 | 327 | 333 | 683 | 828 | 717 | 883 | 868 | 906 | 1077.65 | 743 | 886 | 1.488 | 0.573 | 7.60E-07 |
| AT5G23100.1 | | protein_coding | Protein of unknown funct | 118 | 131 | 138 | 169 | 88 | 61 | 248 | 322 | 353 | 367 | 234 | 166 | 246.79 | 308 | 256 | 1.486 | 0.571 | 0.00918591 |
| AT1G12260.1 | NAC007 | protein_coding | NAC 007 | 44 | 47 | 49 | 72 | 68 | 46 | 92 | 115 | 125 | 156 | 181 | 125 | 118.49 | 111 | 154 | 1.486 | 0.571 | 0.00595546 |
| AT3G15460.1 | | protein_coding | Ribosomal RNA processin | 289 | 178 | 144 | 294 | 224 | 137 | 607 | 437 | 369 | 638 | 595 | 373 | 499.42 | 471 | 535 | 1.485 | 0.571 | 0.00282959 |
| AT1G42970.1 | GAPB | protein_coding | glyceraldehyde-3-phosph | 32463 | 34560 | 26251 | 34714 | 31276 | 36966 | 68240 | 84908 | 67204 | 75298 | 83031 | 100527 | 92923.31 | 73451 | 86285 | 1.486 | 0.571 | 0.00147521 |
| AT3G25130.1 | | protein_coding | | 56 | 58 | 39 | 82 | 69 | 62 | 118 | 142 | 100 | 178 | 183 | 169 | 137.37 | 120 | 177 | 1.486 | 0.571 | 0.00126421 |
| AT3G52900.1 | | protein_coding | Family of unknown functi | 102 | 114 | 105 | 163 | 149 | 99 | 214 | 280 | 269 | 354 | 396 | 269 | 382.32 | 254 | 340 | 1.485 | 0.571 | 7.25E-05 |
| AT1G67440.1 | emb1688 | protein_coding | Minichromosome mainte | 366 | 282 | 305 | 441 | 359 | 330 | 769 | 693 | 781 | 957 | 953 | 897 | 895.9 | 748 | 936 | 1.486 | 0.571 | 5.76E-07 |
| AT1G30680.1 | T5I8.13 | protein_coding | toprim domain-containin | 979 | 592 | 514 | 952 | 618 | 529 | 2058 | 1454 | 1316 | 2065 | 1641 | 1439 | 1623.17 | 1609 | 1715 | 1.484 | 0.57 | 0.00086368 |
| AT5G07900.1 | | protein_coding | Mitochondrial transcripti | 271 | 178 | 149 | 287 | 207 | 185 | 570 | 437 | 381 | 623 | 550 | 503 | 666.26 | 463 | 559 | 1.485 | 0.57 | 1.04E-05 |
| AT5G58020.1 | | protein_coding | | 484 | 447 | 447 | 733 | 509 | 617 | 1017 | 1098 | 1144 | 1590 | 1351 | 1678 | 1300.24 | 1086 | 1540 | 1.485 | 0.57 | 3.80E-06 |
| AT5G24650.1 | | protein_coding | Mitochondrial import inn | 705 | 575 | 464 | 830 | 679 | 690 | 1482 | 1413 | 1188 | 1800 | 1803 | 1876 | 1974.09 | 1361 | 1826 | 1.485 | 0.57 | 1.94E-06 |
| AT3G60980.1 | | protein_coding | Tetratricopeptide repeat | 292 | 310 | 316 | 443 | 367 | 494 | 614 | 762 | 809 | 961 | 974 | 1343 | 866.06 | 728 | 1093 | 1.484 | 0.569 | 7.94E-05 |
| AT5G01500.1 | TAAC | protein_coding | thylakoid ATP/ADP carrie | 226 | 192 | 115 | 337 | 307 | 236 | 475 | 472 | 294 | 731 | 815 | 642 | 647.48 | 414 | 729 | 1.483 | 0.569 | 5.56E-05 |
| AT5G41020.1 | | protein_coding | myb family transcription | 148 | 83 | 85 | 128 | 89 | 97 | 311 | 204 | 218 | 278 | 236 | 264 | 432.67 | 244 | 259 | 1.484 | 0.569 | 2.49E-05 |
| AT3G56510.1 | | protein_coding | RNA-binding (RRM/RBD/ | 271 | 197 | 174 | 271 | 219 | 225 | 570 | 484 | 445 | 588 | 581 | 612 | 564.56 | 500 | 594 | 1.484 | 0.569 | 3.70E-07 |
| AT5G35630.1 | GLN2 | protein_coding | glutamine synthetase 2 | 38442 | 40158 | 37992 | 46645 | 41205 | 47739 | 80808 | 98661 | 97262 | 101177 | 109390 | 129823 | 104185.98 | 92244 | 113463 | 1.483 | 0.568 | 0.00132673 |
| AT5G02870.1 | RPL4D | protein_coding | Ribosomal protein L4/L1 | 6533 | 5122 | 3215 | 7066 | 5027 | 5504 | 13733 | 12584 | 8231 | 15327 | 13346 | 14968 | 15188.52 | 11516 | 14547 | 1.482 | 0.567 | 0.00082171 |
| AT3G47520.1 | MDH | protein_coding | malate dehydrogenase | 2607 | 1530 | 1388 | 2768 | 1908 | 1591 | 5480 | 3759 | 3553 | 6004 | 5065 | 4327 | 5116.13 | 4264 | 5132 | 1.482 | 0.567 | 0.00029827 |
| AT2G41060.1 | UBA2B | protein_coding | RNA-binding (RRM/RBD/ | 328 | 301 | 365 | 397 | 395 | 495 | 689 | 740 | 934 | 861 | 1049 | 1346 | 879.67 | 788 | 1085 | 1.481 | 0.567 | 0.00017224 |
| AT3G66652.1 | FIPS3 | protein_coding | fip1 motif-containing pro | 194 | 166 | 157 | 204 | 179 | 147 | 408 | 408 | 402 | 442 | 475 | 400 | 337.3 | 406 | 439 | 1.481 | 0.567 | 7.53E-05 |
| AT1G52870.2 | | protein_coding | Peroxisomal membrane 2 | 1418 | 1620 | 1495 | 1951 | 1954 | 1633 | 2981 | 3980 | 3827 | 4232 | 5187 | 4441 | 4298.32 | 3596 | 4620 | 1.482 | 0.567 | 4.44E-05 |
| AT2G20340.1 | ELI5 | protein_coding | Pyridoxal phosphate (PLP | 234 | 230 | 186 | 358 | 377 | 233 | 492 | 565 | 476 | 777 | 1001 | 634 | 652.97 | 511 | 804 | 1.481 | 0.566 | 0.00010512 |
| AT2G02020.1 | NPF8.4 | protein_coding | Major facilitator superfar | 196 | 156 | 104 | 250 | 202 | 141 | 412 | 383 | 266 | 542 | 536 | 383 | 563.68 | 354 | 487 | 1.48 | 0.566 | 9.59E-05 |
| AT5G07950.1 | | protein_coding | | 106 | 77 | 86 | 136 | 108 | 122 | 223 | 189 | 220 | 295 | 287 | 332 | 261.68 | 211 | 305 | 1.481 | 0.566 | 7.31E-05 |
| AT4G34340.1 | TAF8 | protein_coding | TBP-associated factor 8 | 94 | 88 | 82 | 111 | 87 | 77 | 198 | 216 | 210 | 241 | 231 | 209 | 188.64 | 208 | 227 | 1.481 | 0.566 | 4.55E-05 |
| AT2G17265.1 | HSK | protein_coding | homoserine kinase | 460 | 497 | 339 | 621 | 549 | 587 | 967 | 1221 | 868 | 1347 | 1457 | 1596 | 1440.59 | 1019 | 1467 | 1.48 | 0.566 | 2.32E-05 |
| AT3G03840.1 | | protein_coding | SAUR-like auxin-responsi | 86 | 116 | 126 | 149 | 147 | 141 | 181 | 285 | 323 | 323 | 390 | 383 | 250.39 | 263 | 365 | 1.48 | 0.565 | 0.00436899 |
| AT5G44490.1 | | protein_coding | FBD, F-box, Skp2-like and | 114 | 64 | 49 | 79 | 52 | 45 | 240 | 157 | 125 | 171 | 138 | 122 | 162.94 | 174 | 144 | 1.478 | 0.564 | 0.00532091 |
| AT1G64970.1 | VTE4 | protein_coding | gamma-tocopherol meth | 171 | 176 | 102 | 237 | 282 | 164 | 359 | 432 | 261 | 514 | 749 | 446 | 472.58 | 351 | 570 | 1.478 | 0.564 | 0.00278026 |
| AT5G20720.1 | CPN21 | protein_coding | chaperonin 20 | 13207 | 8691 | 7047 | 13613 | 9668 | 9307 | 27762 | 21352 | 18041 | 29528 | 25666 | 25310 | 27843.03 | 22385 | 26835 | 1.479 | 0.564 | 0.0006529 |
| AT1G18020.1 | | protein_coding | FMN-linked oxidoreducta | 132 | 173 | 189 | 244 | 204 | 267 | 277 | 425 | 484 | 529 | 542 | 726 | 630.2 | 395 | 599 | 1.479 | 0.564 | 0.00026721 |
| AT1G29320.1 | | protein_coding | Transducin/WD40 repeat | 362 | 228 | 162 | 370 | 256 | 200 | 761 | 560 | 415 | 803 | 680 | 544 | 643.55 | 579 | 676 | 1.477 | 0.563 | 0.00134815 |

| | | | | | | | | | | | | | | | | | | | | | |
|---|---|---|---|---|---|---|---|---|---|---|---|---|---|---|---|---|---|---|---|---|---|
| AT3G14415.2 | GLO2 | protein_coding | Aldolase-type TIM barrel | 11167 | 10399 | 8232 | 17299 | 13499 | 13429 | 23474 | 25548 | 21074 | 37523 | 35837 | 36519 | 32113.52 | 23365 | 36626 | 1.477 | 0.563 | 7.08E-05 |
| AT4G20840.1 | | protein_coding | FAD-binding Berberine fa | 344 | 290 | 238 | 361 | 307 | 244 | 723 | 712 | 609 | 783 | 815 | 664 | 696.2 | 681 | 754 | 1.478 | 0.563 | 1.81E-06 |
| AT1G63220.1 | | protein_coding | Calcium-dependent lipid- | 204 | 141 | 156 | 239 | 168 | 189 | 429 | 346 | 399 | 518 | 446 | 514 | 474.88 | 391 | 493 | 1.477 | 0.563 | 1.16E-06 |
| AT1G70610.1 | ABCB26 | protein_coding | transporter associated wi | 766 | 731 | 641 | 1115 | 982 | 914 | 1610 | 1796 | 1641 | 2419 | 2607 | 2486 | 2030.37 | 1682 | 2504 | 1.478 | 0.563 | 6.94E-07 |
| AT4G21710.1 | NRPB2 | protein_coding | DNA-directed RNA polym | 1733 | 1350 | 1271 | 2325 | 1890 | 1697 | 3643 | 3317 | 3254 | 5043 | 5018 | 4615 | 3059.25 | 3405 | 4892 | 1.477 | 0.563 | 5.13E-07 |
| AT1G22780.1 | RPS18A | protein_coding | Ribosomal protein S13/S1 | 2508 | 1836 | 1343 | 2510 | 1874 | 1832 | 5272 | 4511 | 3438 | 5444 | 4975 | 4982 | 5336.87 | 4407 | 5134 | 1.477 | 0.562 | 0.00010646 |
| AT1G48030.1 | LPD1 | protein_coding | mitochondrial lipoamide | 3081 | 2622 | 1892 | 3634 | 3164 | 2887 | 6476 | 6442 | 4844 | 7882 | 8400 | 7851 | 10080.84 | 5921 | 8044 | 1.477 | 0.562 | 2.94E-05 |
| AT4G02540.1 | | protein_coding | Cysteine/Histidine-rich C1 | 603 | 438 | 481 | 792 | 609 | 578 | 1268 | 1076 | 1231 | 1718 | 1617 | 1572 | 1429.95 | 1192 | 1636 | 1.476 | 0.562 | 7.81E-06 |
| AT4G22710.1 | CYP706A2 | protein_coding | cytochrome P450, family | 635 | 543 | 356 | 805 | 559 | 434 | 1335 | 1334 | 911 | 1746 | 1484 | 1180 | 1303.92 | 1193 | 1470 | 1.475 | 0.561 | 0.00037868 |
| AT4G39600.1 | | protein_coding | Galactose oxidase/kelch r | 22 | 28 | 23 | 39 | 22 | 33 | 46 | 69 | 59 | 85 | 58 | 90 | 73.68 | 58 | 78 | 1.474 | 0.56 | 0.00853193 |
| AT2G35430.1 | | protein_coding | Zinc finger C-x8-C-x5-C-x3 | 46 | 35 | 35 | 39 | 30 | 35 | 97 | 86 | 90 | 85 | 80 | 95 | 93.78 | 91 | 87 | 1.474 | 0.56 | 0.00578924 |
| AT4G21140.1 | | protein_coding | | 83 | 41 | 41 | 63 | 45 | 49 | 174 | 101 | 105 | 137 | 119 | 133 | 207.09 | 127 | 130 | 1.473 | 0.559 | 0.00085228 |
| AT4G10400.1 | | protein_coding | F-box/RNI-like/FBD-like d | 152 | 118 | 93 | 199 | 167 | 134 | 320 | 290 | 238 | 432 | 443 | 364 | 329.73 | 283 | 413 | 1.473 | 0.559 | 0.00017274 |
| AT4G16990.2 | RLM3 | protein_coding | disease resistance proteir | 4687 | 2999 | 3266 | 6578 | 4364 | 4589 | 9852 | 7368 | 8361 | 14268 | 11585 | 12480 | 9869.8 | 8527 | 12778 | 1.473 | 0.559 | 4.13E-05 |
| AT5G59660.1 | | protein_coding | Leucine-rich repeat prote | 45 | 42 | 43 | 53 | 45 | 29 | 95 | 103 | 110 | 115 | 119 | 79 | 102.87 | 103 | 104 | 1.472 | 0.558 | 0.00137408 |
| AT5G53970.1 | | protein_coding | Tyrosine transaminase fa | 413 | 322 | 375 | 608 | 383 | 534 | 868 | 791 | 960 | 1319 | 1017 | 1452 | 898.71 | 873 | 1263 | 1.472 | 0.558 | 5.35E-05 |
| AT5G05470.1 | EIF2 | protein_coding | eukaryotic translation ini | 414 | 283 | 278 | 440 | 361 | 278 | 870 | 695 | 712 | 954 | 958 | 756 | 815.8 | 759 | 889 | 1.473 | 0.558 | 2.53E-05 |
| AT5G23140.1 | CLPP2 | protein_coding | nuclear-encoded CLP prot | 502 | 418 | 336 | 636 | 419 | 497 | 1055 | 1027 | 860 | 1380 | 1112 | 1352 | 1228.32 | 981 | 1281 | 1.472 | 0.558 | 1.65E-05 |
| AT1G70780.1 | | protein_coding | | 658 | 468 | 497 | 657 | 418 | 456 | 1383 | 1150 | 1272 | 1425 | 1110 | 1240 | 1477.81 | 1268 | 1258 | 1.47 | 0.556 | 9.98E-06 |
| AT4G36090.3 | | protein_coding | oxidoreductase, 2OG-Fe(I | 212 | 152 | 173 | 295 | 209 | 216 | 446 | 373 | 443 | 640 | 555 | 587 | 475.82 | 421 | 594 | 1.47 | 0.556 | 6.36E-06 |
| AT3G29075.1 | | protein_coding | glycine-rich protein | 98 | 101 | 71 | 98 | 78 | 114 | 206 | 248 | 182 | 213 | 207 | 310 | 347.49 | 212 | 243 | 1.469 | 0.555 | 0.00048849 |
| AT4G26450.1 | | protein_coding | | 547 | 495 | 542 | 861 | 649 | 742 | 1150 | 1216 | 1388 | 1868 | 1723 | 2018 | 1351.36 | 1251 | 1870 | 1.469 | 0.555 | 5.49E-06 |
| AT4G24620.1 | PGI1 | protein_coding | phosphoglucose isomeras | 2033 | 1671 | 1434 | 2628 | 2237 | 1894 | 4274 | 4105 | 3671 | 5700 | 5939 | 5151 | 5701.6 | 4017 | 5597 | 1.469 | 0.555 | 1.42E-06 |
| AT4G30150.1 | | protein_coding | | 849 | 512 | 493 | 1016 | 748 | 588 | 1785 | 1258 | 1262 | 2204 | 1986 | 1599 | 1183.75 | 1435 | 1930 | 1.468 | 0.554 | 0.00054601 |
| AT3G04760.1 | | protein_coding | Pentatricopeptide repeat | 1988 | 1476 | 1369 | 2406 | 1771 | 1323 | 4179 | 3626 | 3505 | 5219 | 4702 | 3598 | 3622.09 | 3770 | 4506 | 1.469 | 0.554 | 5.09E-05 |
| AT1G30420.1 | ABCC11 | protein_coding | multidrug resistance-asso | 143 | 134 | 132 | 159 | 189 | 151 | 301 | 329 | 338 | 345 | 502 | 411 | 322.68 | 323 | 419 | 1.467 | 0.553 | 0.00014549 |
| AT2G40400.1 | | protein_coding | Protein of unknown funct | 2840 | 2396 | 1641 | 2811 | 2452 | 1897 | 5970 | 5887 | 4201 | 6097 | 6509 | 5159 | 6398.93 | 5353 | 5922 | 1.467 | 0.553 | 7.98E-05 |
| AT4G32605.1 | | protein_coding | Mitochondrial glycoprote | 173 | 142 | 147 | 225 | 210 | 164 | 364 | 349 | 376 | 488 | 558 | 446 | 451.27 | 363 | 497 | 1.467 | 0.553 | 9.08E-07 |
| AT1G70830.1 | MLP28 | protein_coding | MLP-like protein 28 | 1477 | 2235 | 1154 | 1926 | 2378 | 2563 | 3105 | 5491 | 2954 | 4178 | 6313 | 6970 | 6732.74 | 3850 | 5820 | 1.466 | 0.552 | 0.00864864 |
| AT1G10240.1 | FRS11 | protein_coding | FAR1-related sequence 1 | 194 | 185 | 175 | 219 | 214 | 180 | 408 | 455 | 448 | 475 | 568 | 489 | 460.95 | 437 | 511 | 1.467 | 0.552 | 5.74E-06 |
| AT4G21280.2 | PSBQ1 | protein_coding | photosystem II subunit Q | 16872 | 18306 | 13537 | 21813 | 20352 | 24057 | 35466 | 44974 | 34656 | 47314 | 54030 | 65422 | 72626.04 | 38365 | 55589 | 1.465 | 0.551 | 0.00200924 |
| AT1G25210.1 | LPXC1 | protein_coding | UDP-3-O-acyl N-acetylgly | 60 | 74 | 52 | 76 | 79 | 92 | 126 | 182 | 133 | 165 | 210 | 250 | 262.46 | 147 | 208 | 1.465 | 0.551 | 0.00040866 |
| AT2G26080.1 | GLDP2 | protein_coding | glycine decarboxylase P-p | 9093 | 7425 | 6199 | 14111 | 10051 | 10711 | 19114 | 18242 | 15870 | 30608 | 26683 | 29128 | 21461.12 | 17742 | 28806 | 1.465 | 0.551 | 4.85E-05 |
| AT1G71850.1 | | protein_coding | Ubiquitin carboxyl-termir | 156 | 108 | 102 | 166 | 113 | 111 | 328 | 265 | 261 | 360 | 300 | 302 | 305.72 | 285 | 321 | 1.465 | 0.551 | 3.31E-05 |
| AT3G02890.1 | | protein_coding | RING/FYVE/PHD zinc fing | 390 | 321 | 313 | 481 | 410 | 279 | 820 | 789 | 801 | 1043 | 1088 | 759 | 717.98 | 803 | 963 | 1.465 | 0.551 | 2.39E-05 |
| AT5G66580.1 | | protein_coding | | 94 | 123 | 109 | 145 | 144 | 192 | 198 | 302 | 279 | 315 | 382 | 522 | 346.74 | 260 | 406 | 1.464 | 0.55 | 0.00282487 |
| AT5G62440.1 | | protein_coding | Protein of unknown funct | 182 | 128 | 67 | 163 | 135 | 122 | 383 | 314 | 172 | 354 | 358 | 332 | 580.47 | 290 | 348 | 1.465 | 0.55 | 0.00160478 |
| AT3G12270.1 | PRMT3 | protein_coding | protein arginine methyltr | 399 | 240 | 220 | 373 | 297 | 247 | 839 | 590 | 563 | 809 | 788 | 672 | 725.97 | 664 | 756 | 1.464 | 0.55 | 0.00015087 |
| AT4G33140.1 | | protein_coding | Haloacid dehalogenase-li | 143 | 81 | 95 | 170 | 127 | 113 | 301 | 199 | 243 | 369 | 337 | 307 | 285.94 | 248 | 338 | 1.464 | 0.55 | 0.00011026 |
| AT1G65150.1 | | protein_coding | TRAF-like family protein | 95 | 59 | 47 | 118 | 99 | 85 | 200 | 145 | 120 | 256 | 263 | 231 | 207.54 | 155 | 250 | 1.463 | 0.549 | 0.0007756 |
| AT2G01250.1 | RPL7B | protein_coding | Ribosomal protein L30/L7 | 7584 | 5365 | 4174 | 7354 | 5564 | 5232 | 15942 | 13181 | 10686 | 15952 | 14771 | 14228 | 14638.75 | 13270 | 14984 | 1.463 | 0.549 | 0.00029431 |
| AT1G55490.1 | CPN60B1 | protein_coding | chaperonin 60 beta | 14844 | 10372 | 7726 | 13400 | 11231 | 9555 | 31203 | 25482 | 19779 | 29066 | 29816 | 25984 | 28967.4 | 25488 | 28289 | 1.462 | 0.548 | 0.00123914 |
| AT4G03400.1 | DFL2 | protein_coding | Auxin-responsive GH3 far | 1171 | 971 | 782 | 1459 | 1327 | 1005 | 2462 | 2386 | 2002 | 3165 | 3523 | 2733 | 2518.83 | 2283 | 3140 | 1.462 | 0.548 | 0.00011021 |
| AT4G28190.2 | ULT1 | protein_coding | Developmental regulator | 209 | 150 | 164 | 244 | 152 | 227 | 439 | 369 | 420 | 529 | 404 | 617 | 526.23 | 409 | 517 | 1.461 | 0.547 | 0.00024965 |
| AT3G29200.1 | CM1 | protein_coding | chorismate mutase 1 | 508 | 336 | 293 | 632 | 437 | 370 | 1068 | 825 | 750 | 1371 | 1160 | 1006 | 1072.34 | 881 | 1179 | 1.461 | 0.547 | 0.00014418 |
| AT4G18780.1 | CESA8 | protein_coding | cellulose synthase family | 153 | 191 | 180 | 215 | 198 | 174 | 322 | 469 | 461 | 466 | 526 | 473 | 390.86 | 417 | 488 | 1.46 | 0.546 | 0.00015789 |
| AT4G34650.1 | SQS2 | protein_coding | squalene synthase 2 | 172 | 160 | 163 | 213 | 239 | 172 | 362 | 393 | 417 | 462 | 634 | 468 | 426.12 | 391 | 521 | 1.46 | 0.546 | 8.29E-05 |
| AT3G18980.1 | ETP1 | protein_coding | EIN2 targeting protein1 | 426 | 418 | 369 | 556 | 481 | 412 | 895 | 1027 | 945 | 1206 | 1277 | 1120 | 1095.59 | 956 | 1201 | 1.46 | 0.546 | 8.82E-06 |
| AT1G07350.1 | SR45A | protein_coding | RNA-binding (RRM/RBD/ | 272 | 218 | 149 | 264 | 265 | 211 | 572 | 536 | 381 | 573 | 704 | 574 | 797.85 | 496 | 617 | 1.459 | 0.545 | 0.00010783 |
| AT3G54500.3 | | protein_coding | | 2374 | 2152 | 2442 | 3972 | 2983 | 3372 | 4990 | 5287 | 6252 | 8616 | 7919 | 9170 | 6510.78 | 5510 | 8568 | 1.459 | 0.545 | 1.46E-05 |
| AT1G04480.1 | RPL23C | protein_coding | Ribosomal protein L14p/ | 1410 | 1144 | 667 | 1727 | 1213 | 1370 | 2964 | 2811 | 1708 | 3746 | 3220 | 3726 | 3587.66 | 2494 | 3564 | 1.458 | 0.544 | 0.00286747 |
| AT4G18730.1 | RPL11B | protein_coding | ribosomal protein L16B | 2765 | 2198 | 1285 | 2696 | 2047 | 2082 | 5812 | 5400 | 3290 | 5848 | 5434 | 5662 | 5710.24 | 4834 | 5648 | 1.458 | 0.544 | 0.00143174 |
| AT1G67940.1 | ABCI17 | protein_coding | non-intrinsic ABC protein | 73 | 54 | 52 | 108 | 84 | 85 | 153 | 133 | 133 | 234 | 223 | 231 | 211.79 | 140 | 229 | 1.458 | 0.544 | 6.87E-05 |
| AT4G17390.1 | RPL15B | protein_coding | Ribosomal protein L23/L1 | 3501 | 3197 | 1771 | 3800 | 2924 | 3819 | 7359 | 7854 | 4534 | 8243 | 7763 | 10386 | 9924.67 | 6582 | 8797 | 1.457 | 0.543 | 0.00294642 |
| AT1G02820.1 | | protein_coding | Late embryogenesis abun | 62 | 79 | 52 | 61 | 60 | 52 | 130 | 194 | 133 | 132 | 159 | 141 | 197.4 | 152 | 144 | 1.457 | 0.543 | 0.00106608 |
| AT3G15850.1 | ADS3 | protein_coding | fatty acid desaturase 5 | 4809 | 3786 | 2913 | 6876 | 4926 | 3540 | 10109 | 9302 | 7457 | 14915 | 13077 | 9627 | 10549 | 8956 | 12540 | 1.457 | 0.543 | 0.00058837 |
| AT3G23990.1 | CPN60 | protein_coding | heat shock protein 60 | 2504 | 1410 | 1126 | 2601 | 1805 | 1440 | 5264 | 3464 | 2883 | 5642 | 4792 | 3916 | 3881.65 | 3870 | 4783 | 1.456 | 0.542 | 0.00439152 |
| AT3G05060.1 | NOP5-2 | protein_coding | NOP56-like pre RNA proc | 1643 | 1109 | 807 | 1552 | 1283 | 939 | 3454 | 2725 | 2066 | 3366 | 3406 | 2554 | 2687.4 | 2748 | 3109 | 1.456 | 0.542 | 0.00175956 |
| AT5G44050.1 | | protein_coding | MATE efflux family prote | 148 | 119 | 79 | 174 | 107 | 86 | 311 | 292 | 202 | 377 | 284 | 234 | 342 | 268 | 298 | 1.456 | 0.542 | 0.00150699 |
| AT3G27906.1 | | protein_coding | | 75 | 88 | 69 | 120 | 101 | 115 | 158 | 216 | 177 | 260 | 268 | 313 | 277.53 | 184 | 280 | 1.456 | 0.542 | 0.00010032 |
| AT3G22220.1 | | protein_coding | hAT transposon superfam | 452 | 369 | 388 | 449 | 383 | 404 | 950 | 907 | 993 | 974 | 1017 | 1099 | 768.32 | 950 | 1030 | 1.456 | 0.542 | 1.29E-05 |
| AT1G76050.2 | | protein_coding | Pseudouridine synthase f | 384 | 318 | 318 | 480 | 371 | 424 | 807 | 781 | 814 | 1041 | 985 | 1153 | 1118.28 | 801 | 1060 | 1.455 | 0.541 | 1.78E-06 |
| AT2G21385.4 | | protein_coding | | 1537 | 1258 | 1141 | 1744 | 1537 | 1400 | 3231 | 3091 | 2921 | 3783 | 4080 | 3807 | 3767.24 | 3081 | 3890 | 1.454 | 0.54 | 1.70E-06 |
| AT4G20360.1 | TUFA | protein_coding | RAB GTPase homolog E1 | 24732 | 23145 | 19558 | 26768 | 22091 | 25145 | 51988 | 56863 | 50070 | 58062 | 58646 | 68380 | 67188.7 | 52974 | 61696 | 1.453 | 0.539 | 0.00075591 |
| AT3G57350.1 | NUP93B | protein_coding | Nucleoporin interacting c | 406 | 283 | 274 | 491 | 396 | 333 | 853 | 695 | 701 | 1065 | 1051 | 906 | 732.52 | 750 | 1007 | 1.453 | 0.539 | 1.76E-05 |
| AT5G23580.1 | CPK12 | protein_coding | calmodulin-like domain p | 245 | 157 | 165 | 243 | 243 | 148 | 515 | 386 | 422 | 527 | 645 | 402 | 454.55 | 441 | 525 | 1.452 | 0.538 | 0.00033387 |
| AT3G58610.1 | | protein_coding | ketol-acid reductoisomer | 13508 | 9933 | 8496 | 14244 | 10086 | 11240 | 28395 | 24404 | 21750 | 30897 | 26776 | 30567 | 25822.9 | 24850 | 29413 | 1.452 | 0.538 | 0.00024015 |
| AT1G19485.1 | | protein_coding | Transducin/WD40 repeat | 362 | 314 | 393 | 406 | 353 | 404 | 761 | 771 | 1006 | 881 | 937 | 1099 | 856.42 | 846 | 972 | 1.452 | 0.538 | 9.42E-05 |
| AT5G65490.1 | | protein_coding | | 398 | 271 | 273 | 386 | 294 | 333 | 837 | 666 | 699 | 837 | 781 | 906 | 771.72 | 734 | 841 | 1.452 | 0.538 | 6.97E-06 |
| AT3G18130.1 | RACK1C | protein_coding | receptor for activated C k | 1851 | 1135 | 680 | 2007 | 1409 | 1159 | 3891 | 2788 | 1741 | 4353 | 3741 | 3152 | 3515.92 | 2807 | 3749 | 1.451 | 0.537 | 0.00917957 |
| AT3G60245.1 | RPL37AC | protein_coding | Zinc-binding ribosomal pr | 4408 | 3827 | 2253 | 4011 | 3041 | 3750 | 9266 | 9402 | 5768 | 8700 | 8073 | 10198 | 11466.13 | 8145 | 8990 | 1.45 | 0.537 | 0.00463671 |
| AT2G45830.1 | DTA2 | protein_coding | downstream target of AG | 67 | 67 | 81 | 97 | 54 | 100 | 141 | 165 | 207 | 210 | 143 | 272 | 209.3 | 171 | 208 | 1.451 | 0.537 | 0.00426162 |
| AT3G04230.1 | RPS16B | protein_coding | Ribosomal protein S5 dor | 77 | 71 | 41 | 102 | 80 | 103 | 162 | 174 | 105 | 221 | 212 | 280 | 293.26 | 147 | 238 | 1.45 | 0.537 | 0.00259593 |
| AT3G22310.1 | RH9 | protein_coding | putative mitochondrial R | 279 | 208 | 131 | 260 | 245 | 190 | 586 | 511 | 335 | 564 | 650 | 517 | 876.36 | 477 | 577 | 1.451 | 0.537 | 0.00075521 |
| AT4G37920.1 | | protein_coding | | 770 | 506 | 476 | 771 | 554 | 511 | 1619 | 1243 | 1219 | 1672 | 1471 | 1390 | 1304.24 | 1360 | 1511 | 1.451 | 0.537 | 0.00019922 |
| AT1G31814.1 | FRL2 | protein_coding | FRIGIDA like 2 | 125 | 109 | 141 | 166 | 141 | 164 | 263 | 268 | 361 | 360 | 374 | 446 | 261.12 | 297 | 393 | 1.45 | 0.536 | 0.00365216 |
| AT2G33540.1 | CPL3 | protein_coding | C-terminal domain phosp | 892 | 748 | 721 | 1292 | 1079 | 822 | 1875 | 1838 | 1846 | 2802 | 2864 | 2235 | 1748.33 | 1853 | 2634 | 1.45 | 0.536 | 0.00045663 |
| AT4G19710.2 | AKHSDH2 | protein_coding | aspartate kinase-homose | 2513 | 1852 | 1537 | 2510 | 2279 | 1832 | 5283 | 4550 | 3935 | 5444 | 6050 | 4982 | 4573.88 | 4589 | 5492 | 1.45 | 0.536 | 3.54E-05 |
| AT5G13420.1 | | protein_coding | Aldolase-type TIM barrel | 1130 | 534 | 560 | 1594 | 873 | 787 | 2375 | 1312 | 1434 | 3458 | 2318 | 2140 | 2052.4 | 1707 | 2639 | 1.449 | 0.535 | 0.00658195 |
| AT5G17360.1 | | protein_coding | | 42 | 38 | 39 | 54 | 48 | 51 | 88 | 93 | 100 | 117 | 127 | 139 | 96.59 | 94 | 128 | 1.449 | 0.535 | 0.00254692 |
| AT5G58390.1 | PER67 | protein_coding | Peroxidase superfamily p | 74 | 59 | 40 | 83 | 88 | 75 | 156 | 145 | 102 | 180 | 234 | 204 | 218.69 | 134 | 206 | 1.449 | 0.535 | 0.00138639 |
| AT2G38280.1 | AMPD | protein_coding | AMP deaminase, putative | 1244 | 858 | 755 | 1578 | 1043 | 914 | 2615 | 2108 | 1933 | 3423 | 2769 | 2486 | 2306.9 | 2219 | 2893 | 1.449 | 0.535 | 0.00014517 |
| AT4G37560.1 | | protein_coding | Acetamidase/Formamida | 129 | 120 | 135 | 179 | 173 | 135 | 271 | 295 | 346 | 388 | 459 | 367 | 331.1 | 304 | 405 | 1.449 | 0.535 | 9.08E-05 |
| AT5G58620.1 | | protein_coding | zinc finger (CCCH-type) fa | 196 | 122 | 126 | 219 | 121 | 184 | 412 | 300 | 323 | 475 | 321 | 500 | 320.51 | 345 | 432 | 1.448 | 0.534 | 0.00217089 |
| AT3G57150.1 | CBF5 | protein_coding | homologue of NAP57 | 1424 | 1049 | 659 | 1167 | 991 | 967 | 2993 | 2577 | 1687 | 2531 | 2631 | 2630 | 2866.25 | 2419 | 2597 | 1.448 | 0.534 | 0.0016119 |
| AT4G38495.1 | | protein_coding | | 76 | 55 | 74 | 132 | 80 | 103 | 160 | 135 | 189 | 286 | 212 | 280 | 225.5 | 161 | 259 | 1.447 | 0.534 | 0.00129105 |
| AT4G32450.1 | PCMP-H63 | protein_coding | Pentatricopeptide repeat | 156 | 126 | 145 | 184 | 113 | 143 | 328 | 310 | 371 | 399 | 300 | 389 | 283.01 | 336 | 363 | 1.448 | 0.534 | 0.00049435 |
| AT1G74030.1 | ENO1 | protein_coding | enolase 1 | 572 | 406 | 356 | 719 | 467 | 419 | 1202 | 997 | 911 | 1560 | 1240 | 1139 | 1147.87 | 1037 | 1313 | 1.448 | 0.534 | 0.00019662 |
| AT2G43910.2 | HOL1 | protein_coding | HARMLESS TO OZONE LA | 2834 | 2545 | 2051 | 4165 | 3227 | 3452 | 5957 | 6253 | 5251 | 9034 | 8567 | 9388 | 9491.31 | 5820 | 8996 | 1.448 | 0.534 | 8.08E-06 |
| AT5G04890.1 | RTM2 | protein_coding | HSP20-like chaperones su | 40 | 43 | 56 | 48 | 63 | 48 | 84 | 106 | 143 | 104 | 167 | 131 | 167.15 | 111 | 134 | 1.447 | 0.533 | 0.00832657 |
| AT5G02830.1 | | protein_coding | Tetratricopeptide repeat | 1483 | 975 | 814 | 1807 | 1315 | 860 | 3117 | 2395 | 2084 | 3920 | 3491 | 2339 | 2712.05 | 2532 | 3250 | 1.446 | 0.533 | 0.00233553 |
| AT5G66350.1 | SHI | protein_coding | Lateral root primordium ( | 54 | 42 | 29 | 56 | 37 | 45 | 114 | 103 | 74 | 121 | 98 | 122 | 100.42 | 97 | 114 | 1.446 | 0.532 | 0.00480223 |
| AT1G04070.1 | TOM9-1 | protein_coding | translocase of outer mem | 45 | 43 | 27 | 97 | 50 | 77 | 95 | 106 | 69 | 210 | 133 | 209 | 143.44 | 90 | 184 | 1.446 | 0.532 | 0.00439833 |
| AT3G10090.1 | RPS28A | protein_coding | Nucleic acid-binding, OB- | 365 | 220 | 216 | 417 | 294 | 267 | 767 | 540 | 553 | 905 | 781 | 726 | 760.2 | 620 | 804 | 1.446 | 0.532 | 0.00091129 |
| AT1G36390.1 | | protein_coding | Co-chaperone GrpE famil | 970 | 674 | 586 | 1061 | 742 | 637 | 2039 | 1656 | 1500 | 2301 | 1970 | 1732 | 2050.02 | 1732 | 2001 | 1.446 | 0.532 | 0.00011923 |
| AT5G09650.1 | PPA6 | protein_coding | pyrophosphorylase 6 | 6374 | 4621 | 4031 | 7782 | 6227 | 5107 | 13399 | 11353 | 10320 | 16880 | 16531 | 13888 | 13729.88 | 11691 | 15766 | 1.445 | 0.532 | 0.00011277 |
| AT1G65930.1 | CICDH | protein_coding | cytosolic NADP+-depende | 6946 | 5038 | 5306 | 10325 | 5991 | 7094 | 14601 | 12377 | 13584 | 22396 | 15905 | 19292 | 16495.97 | 13521 | 19198 | 1.445 | 0.531 | 0.00020261 |
| AT3G18600.1 | RH51 | protein_coding | P-loop containing nucleo | 916 | 556 | 393 | 830 | 679 | 490 | 1925 | 1366 | 1006 | 1800 | 1803 | 1333 | 1369.18 | 1432 | 1645 | 1.444 | 0.53 | 0.00565701 |
| AT1G51310.1 | | protein_coding | transferases;tRNA (5-met | 325 | 185 | 170 | 322 | 194 | 199 | 683 | 455 | 435 | 698 | 515 | 541 | 758.33 | 524 | 585 | 1.444 | 0.53 | 0.00096198 |
| AT4G15760.1 | MO1 | protein_coding | monooxygenase 1 | 346 | 326 | 341 | 551 | 268 | 381 | 727 | 801 | 873 | 1195 | 711 | 1036 | 878.49 | 800 | 981 | 1.444 | 0.53 | 0.00065671 |
| AT5G03780.1 | TRFL10 | protein_coding | TRF-like 10 | 61 | 59 | 75 | 79 | 71 | 77 | 128 | 145 | 192 | 171 | 188 | 209 | 149.54 | 155 | 189 | 1.443 | 0.529 | 0.00550823 |
| AT5G55640.1 | | protein_coding | | 101 | 113 | 105 | 148 | 120 | 106 | 212 | 278 | 269 | 321 | 319 | 288 | 298.5 | 253 | 309 | 1.443 | 0.529 | 0.00015226 |
| AT1G53460.1 | | protein_coding | | 205 | 123 | 174 | 234 | 191 | 170 | 431 | 302 | 445 | 508 | 507 | 462 | 428.6 | 393 | 492 | 1.443 | 0.529 | 3.29E-05 |
| AT2G43090.1 | | protein_coding | Aconitase/3-isopropylma | 2050 | 1660 | 1280 | 2562 | 1804 | 1982 | 4309 | 4078 | 3277 | 5557 | 4789 | 5390 | 5464.99 | 3888 | 5245 | 1.443 | 0.529 | 2.32E-05 |
| AT1G27385.4 | | protein_coding | | 401 | 313 | 320 | 475 | 331 | 396 | 843 | 769 | 819 | 1030 | 879 | 1077 | 1146.02 | 810 | 995 | 1.443 | 0.529 | 5.46E-06 |
| AT3G25210.1 | | protein_coding | Tetratricopeptide repeat | 85 | 63 | 97 | 84 | 64 | 94 | 179 | 155 | 248 | 182 | 170 | 256 | 197.19 | 194 | 203 | 1.442 | 0.528 | 0.00737053 |
| AT3G13650.1 | DIR7 | protein_coding | Disease resistance-respor | 23 | 26 | 20 | 44 | 47 | 46 | 48 | 64 | 51 | 95 | 125 | 125 | 107.81 | 54 | 115 | 1.442 | 0.528 | 0.00579695 |
| AT3G53370.1 | S1FA1 | protein_coding | S1FA-like DNA-binding pr | 61 | 37 | 30 | 71 | 60 | 48 | 128 | 91 | 77 | 154 | 159 | 131 | 147.03 | 99 | 148 | 1.442 | 0.528 | 0.00219673 |
| AT1G74060.1 | RPL6B | protein_coding | Ribosomal protein L6 fam | 2584 | 1833 | 1571 | 2589 | 2035 | 2045 | 5432 | 4503 | 4022 | 5616 | 5402 | 5561 | 5357.23 | 4652 | 5526 | 1.442 | 0.528 | 3.80E-05 |
| AT1G31200.1 | PP2A9 | protein_coding | phloem protein 2-A9 | 31 | 23 | 22 | 33 | 39 | 26 | 65 | 57 | 56 | 72 | 104 | 71 | 84.39 | 59 | 82 | 1.441 | 0.527 | 0.00804979 |
| AT3G03920.1 | | protein_coding | H/ACA ribonucleoprotein | 173 | 111 | 94 | 171 | 141 | 105 | 364 | 273 | 241 | 371 | 374 | 286 | 1308.54 | 293 | 344 | 1.44 | 0.526 | 0.00121791 |
| AT2G36950.1 | | protein_coding | Heavy metal transport/de | 108 | 122 | 83 | 152 | 103 | 108 | 227 | 300 | 212 | 330 | 273 | 294 | 371.08 | 246 | 299 | 1.44 | 0.526 | 0.00033714 |
| AT1G77940.1 | RPL30B | protein_coding | Ribosomal protein L7Ae/ | 3079 | 2004 | 1768 | 3682 | 2712 | 2281 | 6472 | 4923 | 4526 | 7987 | 7200 | 6203 | 6777.45 | 5307 | 7130 | 1.44 | 0.526 | 0.00027006 |
| AT1G36180.1 | ACC2 | protein_coding | acetyl-CoA carboxylase 2 | 253 | 199 | 221 | 552 | 421 | 389 | 532 | 489 | 566 | 1197 | 1118 | 1058 | 616.71 | 529 | 1124 | 1.44 | 0.526 | 3.55E-06 |
| AT4G25420.1 | GA20OX1 | protein_coding | 2-oxoglutarate (2OG) and | 47 | 35 | 56 | 56 | 52 | 55 | 99 | 86 | 143 | 121 | 138 | 150 | 137.36 | 109 | 136 | 1.439 | 0.525 | 0.00463764 |
| AT4G39330.1 | CAD9 | protein_coding | cinnamyl alcohol dehydro | 5312 | 4567 | 3276 | 6223 | 5573 | 4082 | 11166 | 11220 | 8387 | 13498 | 14795 | 11101 | 11586.16 | 10258 | 13131 | 1.439 | 0.525 | 0.00032087 |
| AT1G61980.1 | | protein_coding | Mitochondrial transcripti | 172 | 131 | 147 | 231 | 158 | 150 | 362 | 322 | 376 | 501 | 419 | 408 | 369.47 | 353 | 443 | 1.439 | 0.525 | 1.32E-05 |
| AT3G17710.1 | | protein_coding | F-box and associated inte | 64 | 59 | 43 | 64 | 56 | 39 | 135 | 145 | 110 | 139 | 149 | 106 | 107.79 | 130 | 131 | 1.437 | 0.524 | 0.00925151 |
| AT4G38225.3 | | protein_coding | | 435 | 526 | 299 | 526 | 468 | 610 | 914 | 1292 | 765 | 1141 | 1242 | 1659 | 2073.57 | 990 | 1347 | 1.438 | 0.524 | 0.00148563 |
| AT4G36690.1 | U2AF65A | protein_coding | U2 snRNP auxilliary facto | 1260 | 1338 | 1219 | 1542 | 1490 | 2001 | 2649 | 3287 | 3121 | 3345 | 3956 | 5442 | 4011.96 | 3019 | 4248 | 1.438 | 0.524 | 0.00062958 |
| AT4G37910.1 | HSP70-9 | protein_coding | mitochondrial heat shock | 3551 | 2306 | 1844 | 3866 | 2740 | 2652 | 7464 | 5665 | 4721 | 8386 | 7274 | 7212 | 5602.32 | 5950 | 7624 | 1.438 | 0.524 | 0.00044138 |
| AT3G02450.1 | | protein_coding | cell division protein ftsH, | 1434 | 1076 | 816 | 1529 | 1213 | 913 | 3014 | 2644 | 2089 | 3317 | 3220 | 2483 | 2801.55 | 2582 | 3007 | 1.438 | 0.524 | 0.00027722 |
| AT5G01650.2 | | protein_coding | Tautomerase/MIF superfa | 447 | 477 | 318 | 487 | 487 | 512 | 940 | 1172 | 814 | 1056 | 1293 | 1392 | 1524.21 | 975 | 1247 | 1.438 | 0.524 | 0.00024578 |
| AT1G35516.1 | | protein_coding | myb-like transcription fac | 241 | 241 | 256 | 334 | 326 | 373 | 507 | 592 | 655 | 724 | 865 | 1014 | 949.46 | 585 | 868 | 1.438 | 0.524 | 0.00015193 |
| AT5G40200.1 | DEGP9 | protein_coding | DegP protease 9 | 841 | 643 | 669 | 1004 | 793 | 762 | 1768 | 1580 | 1713 | 2178 | 2105 | 2072 | 1622.56 | 1687 | 2118 | 1.438 | 0.524 | 3.47E-06 |
| AT2G46572.1 | | other_rna | other RNA | 45 | 39 | 38 | 38 | 44 | 31 | 95 | 96 | 97 | 82 | 117 | 84 | 86.62 | 96 | 94 | 1.437 | 0.523 | 0.00762467 |
| AT5G02600.2 | NAKR1 | protein_coding | Heavy metal transport/de | 114 | 125 | 82 | 144 | 140 | 154 | 240 | 307 | 210 | 312 | 372 | 419 | 415.65 | 252 | 368 | 1.437 | 0.523 | 0.00043049 |
| AT1G72770.1 | HAB1 | protein_coding | HYPERSENSITIVE TO ABA | 271 | 223 | 285 | 461 | 334 | 348 | 570 | 548 | 730 | 1000 | 887 | 946 | 661.68 | 616 | 944 | 1.437 | 0.523 | 1.51E-05 |
| AT3G14760.1 | | protein_coding | | 102 | 158 | 110 | 143 | 186 | 157 | 214 | 388 | 282 | 310 | 494 | 427 | 383.44 | 295 | 410 | 1.436 | 0.522 | 0.00448153 |
| AT5G22440.1 | RPL10AC | protein_coding | Ribosomal protein L1p/L1 | 2219 | 1290 | 1105 | 2130 | 1544 | 1156 | 4665 | 3169 | 2829 | 4620 | 4099 | 3144 | 3591.39 | 3554 | 3954 | 1.436 | 0.522 | 0.0041406 |
| AT2G23350.1 | PAB4 | protein_coding | poly(A) binding protein 4 | 4455 | 3110 | 2204 | 4957 | 3832 | 3168 | 9365 | 7641 | 5642 | 10752 | 10173 | 8615 | 9117.03 | 7549 | 9847 | 1.436 | 0.522 | 0.00110774 |
| AT1G43670.1 | | protein_coding | Inositol monophosphatas | 4205 | 4080 | 4163 | 4934 | 4474 | 4119 | 8839 | 10024 | 10658 | 10702 | 11877 | 11201 | 12896.44 | 9840 | 11260 | 1.436 | 0.522 | 3.40E-05 |
| AT4G00750.1 | | protein_coding | S-adenosyl-L-methionine- | 262 | 357 | 251 | 330 | 338 | 399 | 551 | 877 | 643 | 716 | 897 | 1085 | 864.28 | 690 | 899 | 1.435 | 0.521 | 0.00113029 |
| AT4G35540.1 | | protein_coding | zinc ion binding;transcrip | 51 | 36 | 51 | 67 | 62 | 61 | 107 | 88 | 131 | 145 | 165 | 166 | 127.09 | 109 | 159 | 1.435 | 0.521 | 0.00099655 |
| AT5G60970.1 | TCP5 | protein_coding | TEOSINTE BRANCHED 1, | 123 | 139 | 128 | 177 | 170 | 137 | 259 | 341 | 328 | 384 | 451 | 373 | 312.59 | 309 | 403 | 1.435 | 0.521 | 0.000303 |
| AT3G54110.1 | PUMP1 | protein_coding | plant uncoupling mitocho | 832 | 542 | 476 | 1156 | 759 | 784 | 1749 | 1332 | 1219 | 2507 | 2015 | 2132 | 2049.79 | 1433 | 2218 | 1.435 | 0.521 | 0.00021893 |
| AT2G01755.1 | | protein_coding | | 194 | 182 | 146 | 277 | 224 | 202 | 408 | 447 | 374 | 601 | 595 | 549 | 599.76 | 410 | 582 | 1.435 | 0.521 | 2.05E-06 |
| AT1G56110.1 | NOP56 | protein_coding | homolog of nucleolar pro | 2643 | 1689 | 1271 | 2724 | 2229 | 1766 | 5556 | 4150 | 3254 | 5909 | 5917 | 4803 | 5091.82 | 4320 | 5543 | 1.434 | 0.52 | 0.00265731 |
| AT3G55990.1 | ESK1 | protein_coding | Plant protein of unknowr | 70 | 85 | 70 | 118 | 105 | 105 | 147 | 209 | 179 | 256 | 279 | 286 | 235.29 | 178 | 274 | 1.434 | 0.52 | 0.00031411 |
| AT4G37925.1 | NDHM | protein_coding | subunit NDH-M of NAD(P | 1019 | 825 | 789 | 1911 | 1377 | 1010 | 2142 | 2027 | 2020 | 4145 | 3656 | 2747 | 2624.23 | 2063 | 3516 | 1.434 | 0.52 | 0.00021284 |
| AT5G42300.1 | UBL5 | protein_coding | ubiquitin-like protein 5 | 1186 | 887 | 825 | 1738 | 1223 | 1253 | 2493 | 2179 | 2112 | 3770 | 3247 | 3407 | 2909.52 | 2261 | 3475 | 1.434 | 0.52 | 1.87E-05 |
| AT5G05730.2 | ASA1 | protein_coding | anthranilate synthase alp | 600 | 572 | 485 | 738 | 665 | 675 | 1261 | 1405 | 1242 | 1601 | 1765 | 1836 | 1438.21 | 1303 | 1734 | 1.434 | 0.52 | 1.28E-05 |
| AT1G17560.1 | HLL | protein_coding | Ribosomal protein L14p/ | 120 | 97 | 80 | 181 | 137 | 95 | 252 | 238 | 205 | 393 | 364 | 258 | 302.3 | 232 | 338 | 1.433 | 0.519 | 0.00390239 |
| AT4G10480.1 | | protein_coding | Nascent polypeptide-asso | 3503 | 2582 | 1945 | 3651 | 2747 | 2540 | 7364 | 6343 | 4979 | 7919 | 7293 | 6907 | 7126.41 | 6229 | 7373 | 1.433 | 0.519 | 0.00023932 |
| AT2G25830.1 | | protein_coding | YebC-related | 1151 | 735 | 709 | 1177 | 961 | 775 | 2419 | 1806 | 1815 | 2553 | 2551 | 2108 | 2464.63 | 2013 | 2404 | 1.433 | 0.519 | 0.00015642 |

| | | | | | | | | | | | | | | | | | | | | | |
|---|---|---|---|---|---|---|---|---|---|---|---|---|---|---|---|---|---|---|---|---|---|
| AT1G19670.1 | CLH1 | protein_coding | chlorophyllase 1 | 396 | 412 | 377 | 766 | 700 | 717 | 832 | 1012 | 965 | 1662 | 1858 | 1950 | 1649.4 | 936 | 1823 | 1.433 | 0.519 | 5.78E-05 |
| AT1G02090.1 | CSN7 | protein_coding | Proteasome component ( | 357 | 344 | 293 | 512 | 403 | 444 | 750 | 845 | 750 | 1111 | 1070 | 1207 | 1127.34 | 782 | 1129 | 1.433 | 0.519 | 8.16E-06 |
| AT1G23950.2 | | protein_coding | Protein of unknown funct | 304 | 286 | 224 | 402 | 334 | 348 | 639 | 703 | 573 | 872 | 887 | 946 | 788.56 | 638 | 902 | 1.433 | 0.519 | 3.43E-06 |
| AT1G48840.1 | | protein_coding | Plant protein of unknowr | 269 | 264 | 274 | 372 | 314 | 332 | 565 | 649 | 701 | 807 | 834 | 903 | 659.91 | 638 | 848 | 1.433 | 0.519 | 2.23E-06 |
| AT4G03270.1 | CYCD6-1 | protein_coding | Cyclin D6;1 | 43 | 42 | 34 | 52 | 49 | 27 | 90 | 103 | 87 | 113 | 130 | 73 | 120.44 | 93 | 105 | 1.432 | 0.518 | 0.00750339 |
| AT1G58210.1 | EMB1674 | protein_coding | kinase interacting family | 80 | 91 | 110 | 110 | 87 | 89 | 168 | 224 | 282 | 239 | 231 | 242 | 207.66 | 225 | 237 | 1.432 | 0.518 | 0.00365831 |
| AT1G07880.2 | MPK13 | protein_coding | Protein kinase superfamil | 87 | 67 | 59 | 139 | 124 | 87 | 183 | 165 | 151 | 302 | 329 | 237 | 245.83 | 166 | 289 | 1.432 | 0.518 | 0.00045658 |
| AT4G10920.1 | KELP | protein_coding | transcriptional coactivatc | 674 | 479 | 536 | 793 | 576 | 503 | 1417 | 1177 | 1372 | 1720 | 1529 | 1368 | 1366.85 | 1322 | 1539 | 1.432 | 0.518 | 6.54E-05 |
| AT4G07825.1 | | protein_coding | | 127 | 110 | 112 | 148 | 136 | 161 | 267 | 270 | 287 | 321 | 361 | 438 | 717.74 | 275 | 373 | 1.432 | 0.518 | 2.80E-05 |
| AT2G44160.1 | MTHFR2 | protein_coding | methylenetetrahydrofola | 3173 | 2357 | 2145 | 4195 | 3253 | 2942 | 6670 | 5791 | 5491 | 9099 | 8636 | 8001 | 6673.83 | 5984 | 8579 | 1.432 | 0.518 | 1.43E-05 |
| AT1G56070.1 | LOS1 | protein_coding | Ribosomal protein S5/Elo | 33299 | 24103 | 17410 | 33970 | 24555 | 22610 | 69997 | 59217 | 44571 | 73684 | 65188 | 61487 | 51838.56 | 57928 | 66786 | 1.431 | 0.517 | 0.0040636 |
| AT4G34200.1 | PGDH1 | protein_coding | D-3-phosphoglycerate de | 2248 | 1569 | 995 | 2430 | 1835 | 1989 | 4725 | 3855 | 2547 | 5271 | 4871 | 5409 | 4660.95 | 3709 | 5184 | 1.431 | 0.517 | 0.00258958 |
| AT5G60870.1 | | protein_coding | Regulator of chromosom | 65 | 57 | 50 | 65 | 49 | 59 | 137 | 140 | 128 | 141 | 130 | 160 | 161.52 | 135 | 144 | 1.431 | 0.517 | 0.00074813 |
| AT5G23900.1 | RPL13D | protein_coding | Ribosomal protein L13e f | 1657 | 1274 | 944 | 1923 | 1305 | 1624 | 3483 | 3130 | 2417 | 4171 | 3464 | 4416 | 4063.25 | 3010 | 4017 | 1.431 | 0.517 | 0.00042394 |
| AT1G31970.1 | RH5 | protein_coding | DEA(D/H)-box RNA helica | 1043 | 707 | 581 | 1055 | 899 | 755 | 2192 | 1737 | 1487 | 2288 | 2387 | 2053 | 2190.21 | 1805 | 2243 | 1.431 | 0.517 | 0.00031715 |
| AT1G12880.1 | NUDT12 | protein_coding | nudix hydrolase homolog | 79 | 71 | 79 | 157 | 134 | 101 | 166 | 174 | 202 | 341 | 356 | 275 | 279.08 | 181 | 324 | 1.431 | 0.517 | 0.00016188 |
| AT2G35605.1 | | protein_coding | SWIB/MDM2 domain sup | 200 | 161 | 145 | 276 | 209 | 160 | 420 | 396 | 371 | 599 | 555 | 435 | 484.65 | 396 | 530 | 1.431 | 0.517 | 4.96E-05 |
| AT1G06000.1 | UGT89C1 | protein_coding | UDP-Glycosyltransferase | 196 | 146 | 82 | 245 | 154 | 174 | 412 | 359 | 210 | 531 | 409 | 473 | 583.3 | 327 | 471 | 1.43 | 0.516 | 0.00128636 |
| AT5G51720.1 | NEET | protein_coding | 2 iron, 2 sulfur cluster bir | 503 | 440 | 270 | 629 | 457 | 446 | 1057 | 1081 | 691 | 1364 | 1213 | 1213 | 3460.29 | 943 | 1263 | 1.43 | 0.516 | 0.0008856 |
| AT1G74070.1 | CYP26-2 | protein_coding | Cyclophilin-like peptidyl-p | 665 | 766 | 531 | 837 | 659 | 656 | 1398 | 1882 | 1359 | 1816 | 1749 | 1784 | 2319.83 | 1546 | 1783 | 1.43 | 0.516 | 6.36E-05 |
| AT3G54660.1 | EMB2360 | protein_coding | glutathione reductase | 3672 | 3491 | 3446 | 4822 | 4189 | 3873 | 7719 | 8577 | 8822 | 10459 | 11121 | 10532 | 10908.45 | 8373 | 10704 | 1.43 | 0.516 | 1.81E-05 |
| AT3G18660.2 | GUX1 | protein_coding | plant glycogenin-like star | 44 | 55 | 36 | 62 | 42 | 48 | 92 | 135 | 92 | 134 | 112 | 131 | 103.8 | 106 | 126 | 1.429 | 0.515 | 0.00697821 |
| AT3G21950.1 | | protein_coding | S-adenosyl-L-methionine- | 140 | 117 | 97 | 149 | 211 | 99 | 294 | 287 | 248 | 323 | 560 | 269 | 411.84 | 276 | 384 | 1.429 | 0.515 | 0.00404628 |
| AT3G54400.1 | | protein_coding | Eukaryotic aspartyl prote | 3238 | 2050 | 1702 | 4574 | 2389 | 3434 | 6807 | 5036 | 4357 | 9921 | 6342 | 9339 | 7270.66 | 5400 | 8534 | 1.429 | 0.515 | 0.00325046 |
| AT4G19550.2 | | protein_coding | zinc ion binding;transcrip | 90 | 62 | 87 | 84 | 55 | 54 | 189 | 152 | 223 | 182 | 146 | 147 | 170.26 | 188 | 158 | 1.429 | 0.515 | 0.00225141 |
| AT1G03530.1 | ATNAF1 | protein_coding | nuclear assembly factor 1 | 190 | 153 | 128 | 143 | 167 | 133 | 399 | 376 | 328 | 310 | 443 | 362 | 397.01 | 368 | 372 | 1.429 | 0.515 | 0.00188738 |
| AT3G45030.1 | RPS20A | protein_coding | Ribosomal protein S10p/S | 1692 | 1229 | 838 | 1817 | 1467 | 1323 | 3557 | 3019 | 2145 | 3941 | 3895 | 3598 | 3672.58 | 2907 | 3811 | 1.429 | 0.515 | 0.00129915 |
| AT3G01550.1 | PPT2 | protein_coding | phosphoenolpyruvate (pe | 529 | 460 | 358 | 757 | 732 | 497 | 1112 | 1130 | 917 | 1642 | 1943 | 1352 | 1392.62 | 1053 | 1646 | 1.429 | 0.515 | 0.00024306 |
| AT4G28440.1 | | protein_coding | Nucleic acid-binding, OB- | 330 | 274 | 220 | 413 | 342 | 338 | 694 | 673 | 563 | 896 | 908 | 919 | 1161.38 | 643 | 908 | 1.429 | 0.515 | 4.52E-05 |
| AT5G13280.1 | AK1 | protein_coding | aspartate kinase 1 | 1056 | 839 | 658 | 1282 | 909 | 897 | 2220 | 2061 | 1685 | 2781 | 2413 | 2439 | 2114.35 | 1989 | 2544 | 1.429 | 0.515 | 3.54E-05 |
| AT1G60989.1 | SCRL7 | protein_coding | SCR-like 7 | 58 | 42 | 50 | 116 | 91 | 72 | 122 | 103 | 128 | 252 | 242 | 196 | 174.98 | 118 | 230 | 1.428 | 0.514 | 0.00134804 |
| AT4G29400.1 | | protein_coding | Protein of unknown funct | 255 | 167 | 138 | 198 | 162 | 154 | 536 | 410 | 353 | 429 | 430 | 419 | 417.7 | 433 | 426 | 1.428 | 0.514 | 0.00082215 |
| AT2G26500.2 | | protein_coding | cytochrome b6f complex | 11727 | 10467 | 9142 | 16209 | 13678 | 12705 | 24651 | 25715 | 23404 | 35159 | 36312 | 34550 | 34411.59 | 24590 | 35340 | 1.428 | 0.514 | 0.00023404 |
| AT5G47360.1 | | protein_coding | Tetratricopeptide repeat | 62 | 69 | 69 | 105 | 79 | 67 | 130 | 170 | 177 | 228 | 210 | 182 | 171.99 | 159 | 207 | 1.427 | 0.513 | 0.00242566 |
| AT1G25054.1 | LPXC4 | protein_coding | UDP-3-O-acyl N-acetylgly | 152 | 154 | 202 | 227 | 210 | 251 | 320 | 378 | 517 | 492 | 558 | 683 | 477.61 | 405 | 578 | 1.427 | 0.513 | 0.0010617 |
| AT1G77550.1 | | protein_coding | tubulin-tyrosine ligases;tu | 566 | 374 | 330 | 530 | 410 | 354 | 1190 | 919 | 845 | 1150 | 1088 | 963 | 893.03 | 985 | 1067 | 1.427 | 0.513 | 9.04E-05 |
| AT1G06720.1 | | protein_coding | P-loop containing nucleo: | 939 | 611 | 485 | 829 | 671 | 513 | 1974 | 1501 | 1242 | 1798 | 1781 | 1395 | 1296.64 | 1572 | 1658 | 1.426 | 0.512 | 0.00411118 |
| AT1G23970.2 | | protein_coding | Protein of unknown funct | 147 | 93 | 87 | 119 | 69 | 87 | 309 | 228 | 223 | 258 | 183 | 237 | 234.32 | 253 | 226 | 1.426 | 0.512 | 0.00207453 |
| AT3G26570.1 | PHT2-1 | protein_coding | phosphate transporter 2; | 2977 | 2788 | 1660 | 4143 | 3480 | 2979 | 6258 | 6850 | 4250 | 8987 | 9239 | 8101 | 10315.98 | 5786 | 8776 | 1.426 | 0.512 | 0.0010913 |
| AT3G03130.1 | | protein_coding | | 282 | 232 | 220 | 306 | 320 | 179 | 593 | 570 | 563 | 664 | 850 | 487 | 559.61 | 575 | 667 | 1.426 | 0.512 | 0.00095859 |
| AT4G18740.1 | | protein_coding | Rho termination factor | 612 | 558 | 666 | 919 | 637 | 738 | 1286 | 1371 | 1705 | 1993 | 1691 | 2007 | 1609.56 | 1454 | 1897 | 1.426 | 0.512 | 0.00032877 |
| AT1G17190.1 | GSTU26 | protein_coding | glutathione S-transferase | 188 | 163 | 182 | 263 | 170 | 172 | 395 | 400 | 466 | 570 | 451 | 468 | 487.38 | 420 | 496 | 1.426 | 0.512 | 1.92E-05 |
| AT5G65010.2 | ASN2 | protein_coding | asparagine synthetase 2 | 6620 | 4117 | 4464 | 8271 | 4700 | 4599 | 13916 | 10115 | 11428 | 17941 | 12477 | 12507 | 11374.86 | 11820 | 14308 | 1.425 | 0.511 | 0.00142934 |
| AT2G45300.1 | | protein_coding | RNA 3'-terminal phospha | 1079 | 811 | 697 | 1513 | 1224 | 1027 | 2268 | 1992 | 1784 | 3282 | 3249 | 2793 | 2366.39 | 2015 | 3108 | 1.425 | 0.511 | 3.56E-05 |
| AT4G26680.1 | | protein_coding | Tetratricopeptide repeat | 51 | 43 | 52 | 52 | 53 | 42 | 107 | 106 | 133 | 113 | 141 | 114 | 95.45 | 115 | 123 | 1.424 | 0.51 | 0.00993827 |
| AT5G01930.1 | MAN6 | protein_coding | Glycosyl hydrolase superf | 64 | 66 | 64 | 89 | 96 | 108 | 135 | 162 | 164 | 193 | 255 | 294 | 212.95 | 154 | 247 | 1.424 | 0.51 | 0.00248345 |
| AT1G44800.1 | | protein_coding | nodulin MtN21 /EamA-lik | 192 | 151 | 118 | 365 | 181 | 204 | 404 | 371 | 302 | 792 | 481 | 555 | 489.76 | 359 | 609 | 1.424 | 0.51 | 0.00210668 |
| AT1G20691.1 | | other_rna | | 106 | 88 | 81 | 95 | 94 | 82 | 223 | 216 | 207 | 206 | 250 | 223 | 190.67 | 215 | 226 | 1.424 | 0.51 | 0.00115088 |
| AT1G26761.1 | | protein_coding | Arabinanase/levansucras | 357 | 350 | 262 | 394 | 360 | 249 | 750 | 860 | 671 | 855 | 956 | 677 | 1072.26 | 760 | 829 | 1.424 | 0.51 | 0.00020519 |
| AT3G56990.1 | EDA7 | protein_coding | embryo sac development | 326 | 145 | 136 | 266 | 229 | 167 | 685 | 356 | 348 | 577 | 608 | 454 | 668.84 | 463 | 546 | 1.424 | 0.509 | 0.00871843 |
| AT2G04400.1 | | protein_coding | Aldolase-type TIM barrel | 666 | 456 | 402 | 877 | 627 | 446 | 1400 | 1120 | 1029 | 1902 | 1665 | 1213 | 1330.77 | 1183 | 1593 | 1.423 | 0.509 | 0.00120066 |
| AT3G02650.1 | | protein_coding | Tetratricopeptide repeat | 521 | 327 | 324 | 552 | 386 | 358 | 1095 | 803 | 829 | 1197 | 1025 | 974 | 876.27 | 909 | 1065 | 1.424 | 0.509 | 0.00016135 |
| AT3G08943.1 | | protein_coding | ARM repeat superfamily | 825 | 611 | 520 | 978 | 855 | 716 | 1734 | 1501 | 1331 | 2121 | 2270 | 1947 | 1724.81 | 1522 | 2113 | 1.423 | 0.509 | 7.60E-05 |
| AT5G64460.2 | | protein_coding | Phosphoglycerate mutase | 1102 | 887 | 899 | 1142 | 1015 | 923 | 2316 | 2179 | 2301 | 2477 | 2695 | 2510 | 2738.01 | 2265 | 2561 | 1.423 | 0.509 | 1.84E-05 |
| AT3G18165.1 | MOS4 | protein_coding | modifier of snc1,4 | 408 | 310 | 296 | 559 | 439 | 416 | 858 | 762 | 758 | 1213 | 1165 | 1131 | 1188.55 | 793 | 1170 | 1.423 | 0.509 | 8.42E-06 |
| AT3G19150.1 | KRP6 | protein_coding | KIP-related protein 6 | 54 | 31 | 32 | 70 | 37 | 51 | 114 | 76 | 82 | 152 | 98 | 139 | 119 | 91 | 130 | 1.422 | 0.508 | 0.00915411 |
| AT4G30990.2 | | protein_coding | ARM repeat superfamily | 2445 | 1399 | 1213 | 2377 | 1960 | 1490 | 5140 | 3437 | 3105 | 5156 | 5203 | 4052 | 2845.59 | 3894 | 4804 | 1.422 | 0.508 | 0.00400347 |
| AT5G35220.1 | EGY1 | protein_coding | Peptidase M50 family pro | 1451 | 982 | 751 | 1532 | 1229 | 777 | 3050 | 2413 | 1923 | 3323 | 3263 | 2113 | 3199.76 | 2462 | 2900 | 1.422 | 0.508 | 0.0036184 |
| AT1G66580.1 | RPL10C | protein_coding | senescence associated ge | 3470 | 2213 | 1930 | 3758 | 2311 | 2138 | 7294 | 5437 | 4941 | 8151 | 6135 | 5814 | 6259.22 | 5891 | 6700 | 1.422 | 0.508 | 0.00183649 |
| AT5G57120.1 | | protein_coding | | 290 | 212 | 170 | 293 | 229 | 187 | 610 | 521 | 435 | 636 | 608 | 509 | 1012.4 | 522 | 584 | 1.422 | 0.508 | 0.00025635 |
| AT5G40770.1 | PHB3 | protein_coding | prohibitin 3 | 934 | 686 | 504 | 1142 | 815 | 921 | 1963 | 1685 | 1290 | 2477 | 2164 | 2505 | 2414.56 | 1646 | 2382 | 1.421 | 0.507 | 0.0007005 |
| AT4G34240.1 | ALDH3I1 | protein_coding | aldehyde dehydrogenase | 1234 | 855 | 835 | 1499 | 1184 | 841 | 2594 | 2101 | 2138 | 3251 | 3143 | 2287 | 2801.48 | 2278 | 2894 | 1.42 | 0.506 | 0.00032508 |
| AT5G51460.3 | TPPA | protein_coding | Haloacid dehalogenase-lil | 585 | 308 | 327 | 850 | 442 | 433 | 1230 | 757 | 837 | 1844 | 1173 | 1178 | 1139.86 | 941 | 1398 | 1.419 | 0.505 | 0.00573911 |
| AT5G48610.1 | | protein_coding | | 159 | 107 | 141 | 185 | 161 | 127 | 334 | 263 | 361 | 401 | 427 | 345 | 347.03 | 319 | 391 | 1.419 | 0.505 | 0.0004566 |
| AT5G60790.1 | ABCF1 | protein_coding | ABC transporter family pr | 3325 | 2289 | 1872 | 3707 | 2790 | 2514 | 6989 | 5624 | 4792 | 8041 | 7407 | 6837 | 5083.3 | 5802 | 7428 | 1.419 | 0.505 | 0.00025654 |
| AT5G56210.1 | WIP2 | protein_coding | WPP domain interacting | 162 | 185 | 200 | 214 | 222 | 173 | 341 | 455 | 512 | 464 | 589 | 470 | 564.41 | 436 | 508 | 1.419 | 0.505 | 0.00016865 |
| AT3G04790.1 | RPI3 | protein_coding | Ribose 5-phosphate isom | 4248 | 3841 | 3034 | 5122 | 4388 | 4926 | 8930 | 9437 | 7767 | 11110 | 11649 | 13396 | 13636.26 | 8711 | 12052 | 1.419 | 0.505 | 0.00012463 |
| AT5G64420.1 | | protein_coding | DNA polymerase V family | 1694 | 1189 | 1093 | 1742 | 1449 | 1206 | 3561 | 2921 | 2798 | 3779 | 3847 | 3280 | 2540.44 | 3093 | 3635 | 1.419 | 0.505 | 9.09E-05 |
| AT2G29570.1 | PCNA2 | protein_coding | proliferating cell nuclear | 551 | 454 | 370 | 581 | 532 | 478 | 1158 | 1115 | 947 | 1260 | 1412 | 1300 | 1265.45 | 1073 | 1324 | 1.419 | 0.505 | 6.70E-05 |
| AT5G58340.1 | | protein_coding | myb-like HTH transcriptic | 70 | 98 | 77 | 88 | 92 | 89 | 147 | 241 | 197 | 191 | 244 | 242 | 200.71 | 195 | 226 | 1.419 | 0.504 | 0.00302801 |
| AT3G48500.2 | PDE312 | protein_coding | Nucleic acid-binding, OB- | 1610 | 992 | 841 | 1283 | 1051 | 874 | 3384 | 2437 | 2153 | 2783 | 2790 | 2377 | 2643.83 | 2658 | 2650 | 1.419 | 0.504 | 0.00122448 |
| AT5G42790.1 | PAF1 | protein_coding | proteasome alpha subuni | 900 | 715 | 666 | 1122 | 737 | 917 | 1892 | 1757 | 1705 | 2434 | 1957 | 2494 | 2464.71 | 1785 | 2295 | 1.418 | 0.504 | 3.54E-05 |
| AT2G04039.3 | | protein_coding | | 1270 | 1120 | 1071 | 1699 | 1496 | 1334 | 2670 | 2752 | 2742 | 3685 | 3972 | 3628 | 3737.67 | 2721 | 3762 | 1.418 | 0.504 | 4.36E-06 |
| AT5G62840.1 | | protein_coding | Phosphoglycerate mutase | 234 | 191 | 103 | 267 | 208 | 162 | 492 | 469 | 264 | 579 | 552 | 441 | 717.12 | 408 | 524 | 1.417 | 0.503 | 0.00253305 |
| AT1G32470.1 | GDH3 | protein_coding | Single hybrid motif super | 7903 | 6643 | 6765 | 10780 | 8786 | 8900 | 16613 | 16321 | 17319 | 23383 | 23325 | 24203 | 23229.91 | 16751 | 23637 | 1.417 | 0.503 | 0.00012299 |
| AT5G19760.1 | DTC | protein_coding | Mitochondrial substrate ( | 3988 | 3244 | 2693 | 5060 | 3775 | 4064 | 8383 | 7970 | 6894 | 10976 | 10022 | 11052 | 10125.7 | 7749 | 10683 | 1.417 | 0.503 | 3.56E-05 |
| AT5G01170.1 | | protein_coding | Protein of unknown funct | 46 | 44 | 44 | 35 | 40 | 52 | 97 | 108 | 113 | 76 | 106 | 141 | 131.47 | 106 | 108 | 1.416 | 0.502 | 0.00476516 |
| AT3G57780.1 | | protein_coding | | 252 | 224 | 156 | 345 | 298 | 197 | 530 | 550 | 399 | 748 | 791 | 536 | 596.29 | 493 | 692 | 1.416 | 0.502 | 0.00127417 |
| AT1G15390.1 | PDF1A | protein_coding | peptide deformylase 1A | 485 | 364 | 259 | 486 | 341 | 344 | 1020 | 894 | 663 | 1054 | 905 | 935 | 1084.46 | 859 | 965 | 1.416 | 0.502 | 0.0004986 |
| AT5G19930.1 | | protein_coding | Protein of unknown funct | 53 | 67 | 60 | 88 | 72 | 66 | 111 | 165 | 154 | 191 | 191 | 179 | 211.14 | 143 | 187 | 1.416 | 0.502 | 0.00046386 |
| AT2G43110.1 | | protein_coding | | 355 | 239 | 202 | 316 | 255 | 212 | 746 | 587 | 517 | 685 | 677 | 577 | 621.05 | 617 | 646 | 1.416 | 0.502 | 0.00044769 |
| AT1G12830.1 | | protein_coding | | 137 | 118 | 100 | 143 | 106 | 121 | 288 | 290 | 256 | 310 | 281 | 329 | 326.54 | 278 | 307 | 1.416 | 0.502 | 0.00014586 |
| AT4G29750.1 | | protein_coding | CRS1 / YhbY (CRM) doma | 1067 | 811 | 764 | 1044 | 790 | 769 | 2243 | 1992 | 1956 | 2265 | 2097 | 2091 | 1995.03 | 2064 | 2151 | 1.416 | 0.502 | 8.18E-06 |
| AT1G48450.1 | | protein_coding | Protein of unknown funct | 2003 | 1703 | 1516 | 2373 | 2018 | 2018 | 4210 | 4184 | 3881 | 5147 | 5357 | 5488 | 4953.95 | 4092 | 5331 | 1.417 | 0.502 | 4.56E-06 |
| AT2G43100.1 | IPMI2 | protein_coding | isopropylmalate isomeras | 757 | 716 | 312 | 1307 | 1134 | 946 | 1591 | 1759 | 799 | 2835 | 3011 | 2573 | 3932.97 | 1383 | 2806 | 1.416 | 0.501 | 0.00936921 |
| AT1G12900.1 | GAPA2 | protein_coding | glyceraldehyde 3-phosph | 38483 | 39185 | 27995 | 36760 | 35915 | 29995 | 80894 | 96270 | 71669 | 79736 | 95346 | 81570 | 108043.77 | 82944 | 85551 | 1.415 | 0.501 | 0.00563105 |
| AT2G22000.1 | PEP6 | protein_coding | elicitor peptide 6 precurs | 48 | 45 | 46 | 67 | 75 | 46 | 101 | 111 | 118 | 145 | 199 | 125 | 131.06 | 110 | 156 | 1.415 | 0.501 | 0.00526956 |
| AT3G02150.2 | TCP13 | protein_coding | plastid transcription facto | 219 | 158 | 159 | 278 | 160 | 250 | 460 | 388 | 407 | 603 | 425 | 680 | 384.28 | 418 | 569 | 1.415 | 0.501 | 0.00176608 |
| AT1G08130.1 | LIG1 | protein_coding | DNA ligase 1 | 725 | 651 | 592 | 823 | 794 | 722 | 1524 | 1599 | 1516 | 1785 | 2108 | 1963 | 1604.96 | 1546 | 1952 | 1.415 | 0.501 | 1.53E-05 |
| AT5G17310.2 | UGP2 | protein_coding | UDP-glucose pyrophosph | 969 | 815 | 700 | 1310 | 952 | 916 | 2037 | 2002 | 1792 | 2842 | 2527 | 2491 | 2291.51 | 1944 | 2620 | 1.415 | 0.501 | 1.48E-05 |
| AT5G62430.1 | CDF1 | protein_coding | cycling DOF factor 1 | 337 | 309 | 257 | 434 | 405 | 343 | 708 | 759 | 658 | 941 | 1075 | 933 | 767.63 | 708 | 983 | 1.415 | 0.501 | 7.37E-06 |
| AT1G65310.1 | XTH17 | protein_coding | xyloglucan endotransgluc | 263 | 257 | 318 | 421 | 353 | 386 | 553 | 631 | 814 | 913 | 937 | 1050 | 637.45 | 666 | 967 | 1.414 | 0.5 | 0.00084541 |
| AT1G03400.1 | | protein_coding | 2-oxoglutarate (2OG) anc | 133 | 165 | 167 | 235 | 169 | 237 | 280 | 405 | 428 | 510 | 449 | 645 | 493.59 | 371 | 535 | 1.414 | 0.5 | 0.00076197 |
| AT3G26630.1 | PCMP-A6 | protein_coding | Tetratricopeptide repeat | 350 | 274 | 225 | 360 | 296 | 190 | 736 | 673 | 576 | 781 | 786 | 517 | 739 | 662 | 695 | 1.415 | 0.5 | 0.00072206 |
| AT3G27350.2 | | protein_coding | | 214 | 189 | 272 | 355 | 282 | 300 | 450 | 464 | 696 | 770 | 749 | 816 | 539.12 | 537 | 778 | 1.414 | 0.5 | 0.00041787 |
| AT2G16070.2 | PDV2 | protein_coding | plastid division2 | 283 | 250 | 265 | 320 | 273 | 281 | 595 | 614 | 678 | 694 | 725 | 764 | 706.41 | 629 | 728 | 1.414 | 0.5 | 7.32E-06 |
| AT4G03280.1 | PETC | protein_coding | photosynthetic electron t | 14819 | 16842 | 13216 | 18953 | 16350 | 20945 | 31151 | 41378 | 33834 | 41111 | 43405 | 56959 | 63580.98 | 35454 | 47158 | 1.413 | 0.499 | 0.00496599 |
| AT1G12200.1 | | protein_coding | Flavin-binding monooxyg | 99 | 89 | 56 | 210 | 161 | 112 | 208 | 219 | 143 | 456 | 427 | 305 | 345.42 | 190 | 396 | 1.413 | 0.499 | 0.00173338 |
| AT4G36648.1 | | other_rna | other RNA | 677 | 580 | 758 | 942 | 704 | 987 | 1423 | 1425 | 1941 | 2043 | 1869 | 2684 | 1918.27 | 1596 | 2199 | 1.414 | 0.499 | 0.00141684 |
| AT1G45231.2 | | protein_coding | S-adenosyl-L-methionine- | 151 | 104 | 96 | 177 | 151 | 119 | 317 | 256 | 246 | 384 | 401 | 324 | 310.31 | 273 | 370 | 1.413 | 0.499 | 0.00036452 |
| AT4G24280.1 | HSP70-6 | protein_coding | chloroplast heat shock pr | 10509 | 7473 | 5772 | 9747 | 7635 | 6650 | 22091 | 18360 | 14777 | 21142 | 20269 | 18084 | 18202.91 | 18409 | 19832 | 1.412 | 0.498 | 0.00108047 |
| AT5G08790.1 | NAC081 | protein_coding | NAC (No Apical Meristem | 210 | 153 | 148 | 325 | 167 | 204 | 441 | 376 | 379 | 705 | 443 | 555 | 462.24 | 399 | 568 | 1.412 | 0.498 | 0.00078156 |
| AT4G00895.1 | | protein_coding | ATPase, F1 complex, OSC | 1096 | 960 | 728 | 1601 | 1060 | 1034 | 2304 | 2359 | 1864 | 3473 | 2814 | 2812 | 3341.07 | 2176 | 3033 | 1.412 | 0.498 | 0.00017783 |
| AT5G58330.1 | | protein_coding | lactate/malate dehydrog | 5231 | 4032 | 3490 | 6338 | 4998 | 4409 | 10996 | 9906 | 8935 | 13748 | 13269 | 11990 | 12794.55 | 9946 | 13002 | 1.413 | 0.498 | 8.54E-05 |
| AT2G47990.1 | SWA1 | protein_coding | transducin family protein | 319 | 284 | 168 | 265 | 187 | 246 | 671 | 698 | 430 | 575 | 496 | 669 | 759.31 | 600 | 580 | 1.411 | 0.497 | 0.00424249 |
| AT1G77030.1 | RH29 | protein_coding | hydrolases, acting on acic | 524 | 311 | 281 | 489 | 380 | 291 | 1101 | 764 | 719 | 1061 | 1009 | 791 | 1004.15 | 861 | 954 | 1.411 | 0.497 | 0.00225081 |
| AT3G18030.1 | HAL3A | protein_coding | HAL3-like protein A | 107 | 101 | 113 | 98 | 95 | 122 | 225 | 248 | 289 | 213 | 252 | 332 | 275.91 | 254 | 266 | 1.412 | 0.497 | 0.00137556 |
| AT1G61970.1 | | protein_coding | Mitochondrial transcripti | 158 | 132 | 159 | 208 | 188 | 184 | 332 | 324 | 407 | 451 | 499 | 500 | 367.5 | 354 | 483 | 1.412 | 0.497 | 0.00019347 |
| AT2G22870.1 | EMB2001 | protein_coding | P-loop containing nucleo: | 347 | 191 | 165 | 313 | 203 | 147 | 729 | 469 | 422 | 679 | 539 | 400 | 619.91 | 540 | 539 | 1.41 | 0.496 | 0.00740488 |
| AT4G24700.1 | | protein_coding | | 73 | 53 | 56 | 111 | 81 | 65 | 153 | 130 | 143 | 241 | 215 | 177 | 160.58 | 142 | 211 | 1.41 | 0.496 | 0.00398819 |
| AT3G25230.2 | FKBP62 | protein_coding | rotamase FKBP 1 | 1502 | 1044 | 891 | 1733 | 1380 | 1065 | 3157 | 2565 | 2281 | 3759 | 3664 | 2896 | 2982.74 | 2668 | 3440 | 1.41 | 0.496 | 0.00050392 |
| AT2G41170.1 | | protein_coding | F-box family protein | 210 | 206 | 200 | 343 | 283 | 299 | 441 | 506 | 512 | 744 | 751 | 813 | 560.61 | 486 | 769 | 1.41 | 0.496 | 3.14E-06 |
| AT3G07030.1 | | protein_coding | Alba DNA/RNA-binding p | 55 | 42 | 44 | 45 | 37 | 26 | 116 | 103 | 113 | 98 | 98 | 71 | 267.13 | 111 | 89 | 1.409 | 0.495 | 0.00100869 |
| AT1G16000.1 | | protein_coding | | 232 | 127 | 101 | 218 | 159 | 125 | 488 | 312 | 259 | 473 | 422 | 340 | 414 | 353 | 412 | 1.408 | 0.494 | 0.00709558 |
| AT4G26850.1 | VTC2 | protein_coding | mannose-1-phosphate gu | 2774 | 2362 | 1448 | 4090 | 3002 | 2220 | 5831 | 5803 | 3707 | 8872 | 7970 | 6037 | 6239.98 | 5114 | 7626 | 1.409 | 0.494 | 0.00400646 |
| AT5G03910.1 | ABCB29 | protein_coding | ABC2 homolog 12 | 837 | 723 | 666 | 824 | 778 | 623 | 1759 | 1776 | 1705 | 1787 | 2065 | 1694 | 1850.31 | 1747 | 1849 | 1.409 | 0.494 | 2.30E-05 |
| AT2G01120.2 | ORC4 | protein_coding | origin recognition comple | 182 | 140 | 131 | 213 | 172 | 159 | 383 | 344 | 335 | 462 | 457 | 432 | 287.28 | 354 | 450 | 1.408 | 0.493 | 0.00114676 |
| AT3G55750.1 | RPL35AD | protein_coding | Ribosomal protein L35Ae | 3460 | 2205 | 2026 | 3505 | 2715 | 2111 | 7273 | 5417 | 5187 | 7603 | 7208 | 5741 | 6201.82 | 5959 | 6851 | 1.407 | 0.493 | 0.00104451 |
| AT4G08940.1 | | protein_coding | Ubiquitin carboxyl-termir | 115 | 92 | 85 | 116 | 83 | 79 | 242 | 226 | 218 | 252 | 220 | 215 | 216.55 | 229 | 229 | 1.407 | 0.493 | 0.0007442 |
| AT3G59150.1 | | protein_coding | F-box/RNI-like superfamil | 81 | 74 | 85 | 131 | 98 | 95 | 170 | 182 | 218 | 284 | 260 | 258 | 183.12 | 190 | 267 | 1.407 | 0.493 | 0.00064504 |
| AT1G20920.1 | RH42 | protein_coding | P-loop containing nucleo: | 544 | 461 | 519 | 810 | 737 | 693 | 1144 | 1133 | 1329 | 1757 | 1957 | 1885 | 1643.76 | 1202 | 1866 | 1.407 | 0.493 | 3.37E-05 |
| AT5G65360.1 | HTR2 | protein_coding | Histone superfamily prot | 1299 | 1368 | 973 | 1433 | 1375 | 1511 | 2731 | 3361 | 2491 | 3108 | 3650 | 4109 | 3641.65 | 2861 | 3622 | 1.407 | 0.492 | 0.0004951 |
| AT3G01660.1 | | protein_coding | S-adenosyl-L-methionine- | 242 | 210 | 172 | 301 | 225 | 174 | 509 | 516 | 440 | 653 | 597 | 473 | 525.36 | 488 | 574 | 1.406 | 0.492 | 0.00037099 |
| AT2G24820.1 | TIC55 | protein_coding | translocon at the inner er | 1948 | 1623 | 1280 | 2929 | 2067 | 1836 | 4095 | 3987 | 3277 | 6353 | 5487 | 4993 | 4513.35 | 3786 | 5611 | 1.406 | 0.492 | 0.00012358 |
| AT3G24190.1 | | protein_coding | Protein kinase superfamil | 1624 | 1343 | 1110 | 2390 | 1918 | 1757 | 3414 | 3300 | 2842 | 5184 | 5092 | 4778 | 3683.57 | 3185 | 5018 | 1.406 | 0.492 | 2.43E-05 |
| AT5G58290.1 | RPT3 | protein_coding | regulatory particle triple- | 1573 | 1405 | 1299 | 2278 | 1663 | 1985 | 3307 | 3452 | 3326 | 4941 | 4415 | 5398 | 4403.55 | 3362 | 4918 | 1.406 | 0.492 | 2.05E-05 |
| AT2G30590.1 | WRKY21 | protein_coding | WRKY DNA-binding prote | 395 | 307 | 310 | 508 | 403 | 360 | 830 | 754 | 794 | 1102 | 1070 | 979 | 860.56 | 793 | 1050 | 1.405 | 0.491 | 3.05E-05 |
| AT2G37190.1 | RPL12A | protein_coding | Ribosomal protein L11 fa | 2373 | 1891 | 1278 | 2296 | 1695 | 2010 | 4988 | 4646 | 3272 | 4980 | 4500 | 5466 | 5573.75 | 4302 | 4982 | 1.404 | 0.49 | 0.00106024 |
| AT2G33435.1 | | protein_coding | RNA-binding (RRM/RBD/ | 387 | 320 | 361 | 374 | 297 | 318 | 814 | 786 | 924 | 811 | 788 | 865 | 583.84 | 841 | 821 | 1.404 | 0.49 | 0.00078594 |
| AT5G55740.1 | CRR21 | protein_coding | Tetratricopeptide repeat | 588 | 460 | 389 | 698 | 523 | 380 | 1236 | 1130 | 996 | 1514 | 1388 | 1033 | 1303.82 | 1121 | 1312 | 1.404 | 0.49 | 0.00044725 |
| AT5G17170.1 | ENH1 | protein_coding | rubredoxin family proteir | 2660 | 1813 | 1507 | 3516 | 2582 | 2315 | 5592 | 4454 | 3858 | 7627 | 6855 | 6296 | 6755.92 | 4635 | 6926 | 1.404 | 0.49 | 0.00039628 |
| AT1G50450.1 | | protein_coding | Saccharopine dehydroger | 828 | 717 | 595 | 836 | 776 | 816 | 1741 | 1762 | 1523 | 1813 | 2060 | 2219 | 2568.39 | 1675 | 2031 | 1.404 | 0.49 | 6.90E-05 |
| AT3G08950.1 | HCC1 | protein_coding | electron transport SCO1/ | 189 | 97 | 85 | 178 | 144 | 97 | 397 | 238 | 218 | 386 | 382 | 264 | 312.45 | 284 | 344 | 1.403 | 0.489 | 0.00859617 |
| AT1G04250.1 | IAA17 | protein_coding | AUX/IAA transcriptional r | 622 | 646 | 420 | 1264 | 1013 | 1323 | 1307 | 1587 | 1075 | 2742 | 2689 | 3598 | 2101.85 | 1323 | 3010 | 1.403 | 0.489 | 0.00145023 |
| AT3G04710.1 | TPR10 | protein_coding | ankyrin repeat family pro | 286 | 214 | 163 | 292 | 222 | 166 | 601 | 526 | 417 | 633 | 589 | 451 | 536.97 | 515 | 558 | 1.404 | 0.489 | 0.00066012 |
| AT5G67630.1 | | protein_coding | P-loop containing nucleo: | 452 | 337 | 256 | 489 | 372 | 372 | 950 | 828 | 655 | 1061 | 988 | 1012 | 824.98 | 811 | 1020 | 1.403 | 0.489 | 0.0002886 |
| AT3G55330.1 | PPL1 | protein_coding | PsbP-like protein 1 | 2252 | 2122 | 1913 | 3098 | 2894 | 2308 | 4734 | 5213 | 4897 | 6720 | 7683 | 6276 | 7512.21 | 4948 | 6893 | 1.403 | 0.488 | 5.58E-05 |
| AT1G76860.1 | LSM3B | protein_coding | Small nuclear ribonucleo | 254 | 232 | 206 | 334 | 255 | 318 | 534 | 570 | 527 | 724 | 677 | 865 | 720.57 | 544 | 755 | 1.402 | 0.488 | 3.92E-05 |
| AT4G22240.1 | PAP2 | protein_coding | Plastid-lipid associated pr | 1730 | 1689 | 1399 | 2406 | 1743 | 1976 | 3637 | 4150 | 3582 | 5219 | 4627 | 5374 | 5041.28 | 3790 | 5073 | 1.402 | 0.488 | 2.85E-05 |
| AT5G19440.1 | | protein_coding | NAD(P)-binding Rossman | 564 | 456 | 278 | 821 | 472 | 399 | 1186 | 1120 | 712 | 1781 | 1253 | 1085 | 1353.33 | 1006 | 1373 | 1.402 | 0.487 | 0.00822552 |
| AT3G17830.1 | | protein_coding | Molecular chaperone Hsp | 194 | 109 | 115 | 126 | 132 | 115 | 408 | 268 | 294 | 273 | 350 | 313 | 346.18 | 323 | 312 | 1.401 | 0.487 | 0.00278667 |
| AT1G21280.1 | | protein_coding | | 72 | 74 | 70 | 85 | 78 | 92 | 151 | 182 | 179 | 184 | 207 | 250 | 193.07 | 171 | 214 | 1.401 | 0.487 | 0.00077901 |
| AT4G02390.1 | PARP2 | protein_coding | poly(ADP-ribose) polyme | 143 | 131 | 159 | 212 | 182 | 174 | 301 | 322 | 407 | 460 | 483 | 473 | 342.7 | 343 | 472 | 1.402 | 0.487 | 0.00039634 |
| AT1G31920.1 | PCMP-H11 | protein_coding | Tetratricopeptide repeat | 560 | 538 | 479 | 676 | 623 | 477 | 1177 | 1322 | 1226 | 1466 | 1654 | 1297 | 1029.11 | 1242 | 1472 | 1.401 | 0.487 | 6.12E-05 |
| AT3G04500.1 | | protein_coding | RNA-binding (RRM/RBD/ | 324 | 232 | 261 | 460 | 350 | 383 | 681 | 570 | 668 | 998 | 929 | 1042 | 887.63 | 640 | 990 | 1.402 | 0.487 | 2.06E-05 |

| | | | | | | | | | | | | | | | | | | | | | |
|---|---|---|---|---|---|---|---|---|---|---|---|---|---|---|---|---|---|---|---|---|---|
| AT1G78070.1 | | protein_coding | Transducin/WD40 repeat | 203 | 261 | 253 | 294 | 323 | 270 | 427 | 641 | 648 | 638 | 857 | 734 | 658.34 | 572 | 743 | 1.401 | 0.486 | 0.0038236 |
| AT3G08640.1 | | protein_coding | Protein of unknown funct | 627 | 396 | 318 | 733 | 559 | 487 | 1318 | 973 | 814 | 1590 | 1484 | 1324 | 1546.59 | 1035 | 1466 | 1.401 | 0.486 | 0.00108103 |
| AT4G05631.1 | | protein_coding | | 2647 | 2174 | 2286 | 1623 | 1285 | 1315 | 5564 | 5341 | 5852 | 3520 | 3411 | 3576 | 2332.77 | 5586 | 3502 | 1.401 | 0.486 | 0.00082325 |
| AT5G18200.1 | | protein_coding | UTP:galactose-1-phospha | 83 | 94 | 87 | 161 | 146 | 142 | 174 | 231 | 223 | 349 | 388 | 386 | 288.03 | 209 | 374 | 1.401 | 0.486 | 0.00014929 |
| AT1G70670.1 | PXG4 | protein_coding | Caleosin-related family p | 152 | 148 | 131 | 256 | 203 | 162 | 320 | 364 | 335 | 555 | 539 | 441 | 443.12 | 340 | 512 | 1.4 | 0.486 | 9.28E-05 |
| AT1G47250.1 | PAF2 | protein_coding | 20S proteasome alpha su | 309 | 336 | 310 | 476 | 394 | 431 | 650 | 825 | 794 | 1032 | 1046 | 1172 | 1002.1 | 756 | 1083 | 1.4 | 0.486 | 9.28E-05 |
| AT5G13680.1 | ELP1 | protein_coding | IKI3 family protein | 665 | 521 | 488 | 682 | 632 | 519 | 1398 | 1280 | 1249 | 1479 | 1678 | 1411 | 1185.42 | 1309 | 1523 | 1.4 | 0.486 | 3.13E-05 |
| AT1G63050.1 | LPLAT2 | protein_coding | MBOAT (membrane bour | 295 | 288 | 242 | 346 | 300 | 244 | 620 | 708 | 620 | 751 | 796 | 664 | 679.1 | 649 | 737 | 1.401 | 0.486 | 1.48E-05 |
| AT3G52340.1 | SPP3B | protein_coding | sucrose-6F-phosphate ph | 1369 | 723 | 706 | 1490 | 835 | 774 | 2878 | 1776 | 1807 | 3232 | 2217 | 2105 | 2255.64 | 2154 | 2518 | 1.399 | 0.485 | 0.00773181 |
| AT2G42950.1 | | protein_coding | Magnesium transporter C | 120 | 101 | 76 | 151 | 121 | 74 | 252 | 248 | 195 | 328 | 321 | 201 | 226.82 | 232 | 283 | 1.4 | 0.485 | 0.00381832 |
| AT5G58350.1 | WNK4 | protein_coding | with no lysine (K) kinase | 148 | 116 | 120 | 292 | 143 | 158 | 311 | 285 | 307 | 633 | 380 | 430 | 346.33 | 301 | 481 | 1.4 | 0.485 | 0.00230101 |
| AT2G37990.1 | | protein_coding | ribosome biogenesis regu | 869 | 521 | 499 | 861 | 647 | 531 | 1827 | 1280 | 1277 | 1868 | 1718 | 1444 | 1618.77 | 1461 | 1677 | 1.399 | 0.485 | 0.00129723 |
| AT5G13240.1 | | protein_coding | transcription regulators | 619 | 460 | 467 | 831 | 564 | 607 | 1301 | 1130 | 1196 | 1803 | 1497 | 1651 | 1507.17 | 1209 | 1650 | 1.399 | 0.485 | 0.00022657 |
| AT1G50920.1 | | protein_coding | Nucleolar GTP-binding pr | 3099 | 2194 | 1807 | 3172 | 2507 | 2343 | 6514 | 5390 | 4626 | 6880 | 6656 | 6372 | 5117.47 | 5510 | 6636 | 1.4 | 0.485 | 0.00017065 |
| AT1G29965.1 | RPL18AA | protein_coding | Ribosomal protein L18ae/ | 277 | 170 | 145 | 269 | 204 | 148 | 582 | 418 | 371 | 583 | 542 | 402 | 672.12 | 457 | 509 | 1.398 | 0.484 | 0.00330211 |
| AT3G13940.1 | | protein_coding | DNA binding;DNA-directe | 355 | 220 | 208 | 338 | 277 | 236 | 746 | 540 | 532 | 733 | 735 | 642 | 649.93 | 606 | 703 | 1.399 | 0.484 | 0.0006303 |
| AT1G20580.1 | SMD3B | protein_coding | Small nuclear ribonucleop | 325 | 308 | 253 | 363 | 309 | 374 | 683 | 757 | 648 | 787 | 820 | 1017 | 1150.6 | 696 | 875 | 1.399 | 0.484 | 0.00027367 |
| AT2G40760.1 | STR7 | protein_coding | Rhodanese/Cell cycle con | 377 | 342 | 313 | 403 | 284 | 296 | 792 | 840 | 801 | 874 | 754 | 805 | 908.73 | 811 | 811 | 1.399 | 0.484 | 2.24E-05 |
| AT3G21550.1 | DMP2 | protein_coding | DUF679 domain membra | 124 | 156 | 127 | 254 | 185 | 184 | 261 | 383 | 325 | 551 | 491 | 500 | 372.45 | 323 | 514 | 1.398 | 0.483 | 0.00629002 |
| AT5G49500.1 | SRP-54B | protein_coding | Signal recognition particl | 48 | 43 | 50 | 48 | 48 | 38 | 101 | 106 | 128 | 104 | 127 | 103 | 125.61 | 112 | 111 | 1.398 | 0.483 | 0.00558657 |
| AT2G39460.1 | RPL23AA | protein_coding | ribosomal protein L23AA | 5224 | 3539 | 2826 | 4844 | 3514 | 3199 | 10981 | 8695 | 7235 | 10507 | 9329 | 8699 | 10486.15 | 8970 | 9512 | 1.398 | 0.483 | 0.00161158 |
| AT4G10690.1 | | transposable_elem | transposable element ger | 115 | 72 | 73 | 205 | 141 | 122 | 242 | 177 | 187 | 445 | 374 | 332 | 268.92 | 202 | 384 | 1.397 | 0.483 | 0.00123325 |
| AT2G17370.1 | HMG2 | protein_coding | 3-hydroxy-3-methylglutar | 157 | 147 | 115 | 154 | 142 | 152 | 330 | 361 | 294 | 334 | 377 | 413 | 308.7 | 328 | 375 | 1.398 | 0.483 | 0.00078879 |
| AT5G63830.1 | | protein_coding | HIT-type Zinc finger famil | 83 | 56 | 79 | 104 | 76 | 91 | 174 | 138 | 202 | 226 | 202 | 247 | 180.07 | 171 | 225 | 1.396 | 0.482 | 0.00236085 |
| AT5G65900.1 | RH27 | protein_coding | DEA(D/H)-box RNA helica | 428 | 281 | 281 | 451 | 345 | 262 | 900 | 690 | 719 | 978 | 916 | 712 | 810.35 | 770 | 869 | 1.397 | 0.482 | 0.0009934 |
| AT1G41880.1 | RPL35AB | protein_coding | Ribosomal protein L35Ae | 3297 | 2176 | 1878 | 3295 | 2415 | 2123 | 6931 | 5346 | 4808 | 7147 | 6411 | 5773 | 6457.55 | 5695 | 6444 | 1.396 | 0.481 | 0.00181726 |
| AT4G04620.1 | ATG8B | protein_coding | Ubiquitin-like superfamily | 75 | 99 | 84 | 133 | 93 | 108 | 158 | 243 | 215 | 288 | 247 | 294 | 259.97 | 205 | 276 | 1.396 | 0.481 | 0.00128178 |
| AT3G60965.1 | | transposable_elem | transposable element ger | 211 | 162 | 143 | 216 | 181 | 207 | 444 | 398 | 366 | 469 | 481 | 563 | 390.66 | 403 | 504 | 1.395 | 0.481 | 0.00011291 |
| AT1G05670.1 | | protein_coding | Pentatricopeptide repeat | 197 | 154 | 137 | 153 | 137 | 100 | 414 | 378 | 351 | 332 | 364 | 272 | 306.96 | 381 | 323 | 1.395 | 0.48 | 0.00067663 |
| AT3G21770.1 | PER30 | protein_coding | Peroxidase superfamily p | 66 | 44 | 51 | 102 | 96 | 84 | 139 | 108 | 131 | 221 | 255 | 228 | 167.69 | 126 | 235 | 1.394 | 0.479 | 0.00137204 |
| AT4G16410.1 | | protein_coding | | 1700 | 1415 | 1107 | 2369 | 1887 | 1458 | 3574 | 3476 | 2834 | 5139 | 5010 | 3965 | 4556.99 | 3295 | 4705 | 1.393 | 0.479 | 0.00035044 |
| AT3G02820.1 | | protein_coding | zinc knuckle (CCHC-type) | 126 | 130 | 108 | 131 | 131 | 122 | 265 | 319 | 276 | 284 | 348 | 332 | 285.5 | 287 | 321 | 1.394 | 0.479 | 0.00024182 |
| AT5G26860.1 | LON1 | protein_coding | lon protease 1 | 901 | 549 | 423 | 943 | 640 | 518 | 1894 | 1349 | 1083 | 2045 | 1699 | 1409 | 1405.57 | 1442 | 1718 | 1.393 | 0.478 | 0.00966909 |
| AT5G65710.1 | HSL2 | protein_coding | HAESA-like 2 | 84 | 99 | 104 | 124 | 122 | 100 | 177 | 243 | 266 | 269 | 324 | 272 | 224.66 | 229 | 288 | 1.393 | 0.478 | 0.00555898 |
| AT1G05600.1 | | protein_coding | Tetratricopeptide repeat | 173 | 102 | 91 | 136 | 133 | 99 | 364 | 251 | 233 | 295 | 353 | 269 | 267.23 | 283 | 306 | 1.393 | 0.478 | 0.00249919 |
| AT3G22320.1 | NRPB5A | protein_coding | Eukaryotic rpb5 RNA poly | 982 | 739 | 726 | 1288 | 966 | 837 | 2064 | 1816 | 1859 | 2794 | 2565 | 2276 | 2062.41 | 1913 | 2545 | 1.393 | 0.478 | 0.00013218 |
| AT5G01030.1 | | protein_coding | Protein of unknown funct | 443 | 286 | 347 | 501 | 371 | 330 | 931 | 703 | 888 | 1087 | 985 | 897 | 734.65 | 841 | 990 | 1.392 | 0.478 | 0.00010445 |
| AT4G03120.1 | | protein_coding | C2H2 and C2HC zinc finge | 156 | 128 | 109 | 168 | 136 | 153 | 328 | 314 | 279 | 364 | 361 | 416 | 425.01 | 307 | 380 | 1.393 | 0.478 | 5.99E-05 |
| AT5G52630.1 | RPA3A | protein_coding | Nucleic acid-binding, OB- | 217 | 123 | 94 | 202 | 180 | 124 | 456 | 302 | 241 | 438 | 478 | 337 | 401.82 | 333 | 418 | 1.391 | 0.477 | 0.00764115 |
| AT3G01120.1 | CGS1 | protein_coding | Pyridoxal phosphate (PLP | 4746 | 4241 | 2704 | 6341 | 5309 | 3922 | 9976 | 10419 | 6922 | 13754 | 14094 | 10666 | 12760.01 | 9106 | 12838 | 1.392 | 0.477 | 0.00247654 |
| AT3G04520.1 | THA2 | protein_coding | threonine aldolase 2 | 175 | 200 | 140 | 322 | 198 | 241 | 368 | 491 | 358 | 698 | 526 | 655 | 561.12 | 406 | 626 | 1.392 | 0.477 | 0.0003561 |
| AT4G31700.1 | RPS6A | protein_coding | ribosomal protein S6 | 7951 | 5467 | 3701 | 7320 | 4973 | 4973 | 16714 | 13431 | 9475 | 15878 | 13202 | 13524 | 14468.36 | 13207 | 14201 | 1.391 | 0.476 | 0.00702898 |
| AT3G11070.1 | | protein_coding | Outer membrane OMP85 | 493 | 395 | 258 | 571 | 413 | 458 | 1036 | 970 | 660 | 1239 | 1096 | 1246 | 981.95 | 889 | 1194 | 1.391 | 0.476 | 0.00189077 |
| AT1G15430.1 | | protein_coding | Protein of unknown funct | 73 | 79 | 58 | 112 | 84 | 104 | 153 | 194 | 148 | 243 | 223 | 283 | 212.13 | 165 | 250 | 1.391 | 0.476 | 0.0011609 |
| AT5G41520.1 | RPS10B | protein_coding | RNA binding Plectin/S10 | 1988 | 1598 | 1201 | 2235 | 1704 | 1949 | 4179 | 3926 | 3075 | 4848 | 4524 | 5300 | 6129.95 | 3727 | 4891 | 1.391 | 0.476 | 0.00071455 |
| AT4G29380.1 | | protein_coding | protein kinase family pro | 613 | 535 | 612 | 701 | 639 | 705 | 1289 | 1314 | 1567 | 1521 | 1696 | 1917 | 1191.89 | 1390 | 1711 | 1.39 | 0.476 | 0.00024152 |
| AT5G01270.2 | CPL2 | protein_coding | carboxyl-terminal domain | 612 | 473 | 418 | 671 | 574 | 486 | 1286 | 1162 | 1070 | 1455 | 1524 | 1322 | 1194.74 | 1173 | 1434 | 1.39 | 0.476 | 5.51E-05 |
| AT1G27130.1 | GSTU13 | protein_coding | glutathione S-transferase | 243 | 333 | 190 | 463 | 324 | 406 | 511 | 818 | 486 | 1004 | 860 | 1104 | 1344.51 | 605 | 989 | 1.39 | 0.475 | 0.0039283 |
| AT3G24170.1 | ATGR1 | protein_coding | glutathione-disulfide redu | 1566 | 962 | 935 | 2611 | 1489 | 1493 | 3292 | 2363 | 2394 | 5664 | 3953 | 4060 | 3192.77 | 2683 | 4559 | 1.39 | 0.475 | 0.00211241 |
| AT5G52470.1 | MED36B | protein_coding | fibrillarin 1 | 1572 | 1242 | 955 | 1880 | 1422 | 1387 | 3304 | 3051 | 2445 | 4078 | 3775 | 3772 | 4078.93 | 2933 | 3875 | 1.389 | 0.475 | 0.00051452 |
| AT5G48990.1 | | protein_coding | Galactose oxidase/kelch r | 126 | 105 | 139 | 171 | 143 | 130 | 265 | 258 | 356 | 371 | 380 | 354 | 283.8 | 293 | 368 | 1.389 | 0.475 | 0.00051245 |
| AT4G38150.1 | | protein_coding | Pentatricopeptide repeat | 116 | 95 | 66 | 124 | 106 | 106 | 244 | 233 | 169 | 269 | 281 | 288 | 311.99 | 215 | 279 | 1.389 | 0.474 | 0.00172614 |
| AT5G56660.1 | ILL2 | protein_coding | IAA-leucine resistant (ILR | 139 | 104 | 99 | 225 | 151 | 110 | 292 | 256 | 253 | 488 | 401 | 299 | 323.72 | 267 | 396 | 1.389 | 0.474 | 0.00161922 |
| AT3G54470.1 | PYRE-F | protein_coding | uridine 5'-monophosphat | 1639 | 1053 | 906 | 1709 | 1334 | 1056 | 3445 | 2587 | 2319 | 3707 | 3541 | 2872 | 3045.65 | 2784 | 3373 | 1.389 | 0.474 | 0.00160917 |
| AT1G26770.2 | EXPA10 | protein_coding | expansin A10 | 235 | 270 | 178 | 284 | 269 | 253 | 494 | 663 | 456 | 616 | 714 | 688 | 774.14 | 538 | 673 | 1.389 | 0.474 | 0.00061927 |
| AT3G22890.1 | APS1 | protein_coding | ATP sulfurylase 1 | 3352 | 3277 | 1799 | 4252 | 4085 | 3177 | 7046 | 8051 | 4606 | 9223 | 10845 | 8640 | 9215.63 | 6568 | 9569 | 1.388 | 0.473 | 0.00660501 |
| AT3G62250.1 | RPS27AC | protein_coding | ubiquitin 5 | 4433 | 3968 | 2525 | 4782 | 3474 | 4223 | 9318 | 9749 | 6464 | 10373 | 9223 | 11484 | 11687.07 | 8510 | 10360 | 1.388 | 0.473 | 0.00354109 |
| AT4G39040.1 | | protein_coding | RNA-binding CRS1 / YhbY | 1403 | 1092 | 967 | 1445 | 1149 | 1229 | 2949 | 2683 | 2476 | 3134 | 3050 | 3342 | 3452.13 | 2703 | 3175 | 1.388 | 0.473 | 4.14E-05 |
| AT3G29160.1 | KIN11 | protein_coding | SNF1 kinase homolog 11 | 598 | 473 | 535 | 696 | 509 | 547 | 1257 | 1162 | 1370 | 1510 | 1351 | 1488 | 1216.02 | 1263 | 1450 | 1.387 | 0.472 | 5.23E-05 |
| AT1G70782.1 | CPuORF28 | protein_coding | conserved peptide upstre | 626 | 481 | 500 | 671 | 381 | 478 | 1316 | 1182 | 1280 | 1455 | 1011 | 1300 | 1502.62 | 1259 | 1255 | 1.386 | 0.471 | 0.00045265 |
| AT5G24400.1 | EMB2024 | protein_coding | NagB/RpiA/CoA transfera | 490 | 383 | 401 | 692 | 486 | 503 | 1030 | 941 | 1027 | 1501 | 1290 | 1368 | 1230.81 | 999 | 1386 | 1.386 | 0.471 | 3.58E-05 |
| AT1G12000.1 | PFP-BETA1 | protein_coding | Phosphofructokinase fam | 1781 | 1515 | 1439 | 2431 | 1941 | 1996 | 3744 | 3722 | 3684 | 5273 | 5153 | 5428 | 4020.06 | 3717 | 5285 | 1.386 | 0.471 | 1.37E-05 |
| AT2G33050.1 | AtRLP26 | protein_coding | receptor like protein 26 | 377 | 289 | 207 | 465 | 324 | 255 | 792 | 710 | 530 | 1009 | 860 | 693 | 889.68 | 677 | 854 | 1.385 | 0.47 | 0.00309887 |
| AT5G08620.1 | RH25 | protein_coding | DEA(D/H)-box RNA helica | 514 | 359 | 327 | 494 | 451 | 420 | 1080 | 882 | 837 | 1072 | 1197 | 1142 | 930.82 | 933 | 1137 | 1.385 | 0.47 | 0.00050052 |
| AT1G03150.1 | | protein_coding | Acyl-CoA N-acyltransfera | 279 | 213 | 205 | 272 | 264 | 249 | 586 | 523 | 525 | 590 | 701 | 677 | 652.63 | 545 | 656 | 1.385 | 0.47 | 0.00012002 |
| AT3G53580.1 | DAPF | protein_coding | diaminopimelate epimera | 1213 | 1143 | 954 | 1603 | 1249 | 1370 | 2550 | 2808 | 2442 | 3477 | 3316 | 3726 | 3518.45 | 2600 | 3506 | 1.385 | 0.47 | 7.76E-05 |
| AT5G47435.2 | PURU1 | protein_coding | formyltetrahydrofolate de | 864 | 688 | 723 | 1008 | 759 | 731 | 1816 | 1690 | 1851 | 2186 | 2015 | 1988 | 1927.61 | 1786 | 2063 | 1.385 | 0.47 | 5.13E-05 |
| AT5G22800.1 | EMB86 | protein_coding | Alanyl-tRNA synthetase, | 3039 | 2542 | 2307 | 2755 | 2609 | 2229 | 6388 | 6245 | 5906 | 5976 | 6926 | 6062 | 5985.75 | 6180 | 6321 | 1.386 | 0.47 | 2.96E-05 |
| AT1G09800.1 | | protein_coding | Pseudouridine synthase f | 86 | 73 | 55 | 80 | 74 | 63 | 181 | 179 | 141 | 174 | 196 | 171 | 182.41 | 167 | 180 | 1.384 | 0.469 | 0.00848102 |
| AT4G27090.1 | RPL14B | protein_coding | Ribosomal protein L14 | 6231 | 4833 | 3911 | 6293 | 4834 | 4893 | 13098 | 11874 | 10012 | 13650 | 12833 | 13306 | 12403.8 | 11661 | 13263 | 1.384 | 0.469 | 0.00067257 |
| AT1G79920.1 | HSP70-15 | protein_coding | Heat shock protein 70 (H | 2640 | 2173 | 1520 | 3140 | 2593 | 2155 | 5549 | 5339 | 3891 | 6811 | 6884 | 5860 | 5993.74 | 4926 | 6518 | 1.384 | 0.469 | 0.00052956 |
| AT1G16880.1 | ACR11 | protein_coding | uridylyltransferase-relate | 5579 | 4349 | 3990 | 7813 | 6207 | 5142 | 11727 | 10685 | 10215 | 16947 | 16478 | 13983 | 16608.8 | 10876 | 15803 | 1.384 | 0.469 | 0.00038035 |
| AT3G52220.1 | | protein_coding | | 560 | 520 | 543 | 738 | 559 | 614 | 1177 | 1278 | 1390 | 1601 | 1484 | 1670 | 1449.33 | 1282 | 1585 | 1.384 | 0.469 | 9.42E-05 |
| AT3G13235.1 | | protein_coding | ubiquitin family protein | 670 | 500 | 465 | 798 | 580 | 620 | 1408 | 1228 | 1190 | 1731 | 1540 | 1686 | 1276.42 | 1275 | 1652 | 1.384 | 0.469 | 6.88E-05 |
| AT2G43770.1 | | protein_coding | Transducin/WD40 repeat | 494 | 386 | 380 | 590 | 451 | 457 | 1038 | 948 | 973 | 1280 | 1197 | 1243 | 1170.62 | 986 | 1240 | 1.384 | 0.469 | 2.06E-05 |
| AT1G01790.1 | KEA1 | protein_coding | K+ efflux antiporter 1 | 3797 | 3977 | 3961 | 4786 | 5345 | 4557 | 7982 | 9771 | 10140 | 10381 | 14190 | 12392 | 10804.53 | 9298 | 12321 | 1.383 | 0.468 | 0.00110867 |
| AT3G07090.1 | | protein_coding | PPPDE putative thiol pep | 441 | 330 | 280 | 513 | 437 | 300 | 927 | 811 | 717 | 1113 | 1160 | 816 | 942.72 | 818 | 1030 | 1.383 | 0.468 | 0.00064465 |
| AT1G74250.1 | | protein_coding | DNAJ heat shock N-termi | 93 | 70 | 66 | 117 | 88 | 103 | 195 | 172 | 169 | 254 | 234 | 280 | 507.31 | 179 | 256 | 1.383 | 0.468 | 1.17E-05 |
| AT1G23090.1 | SULTR3 | protein_coding | sulfate transporter 91 | 494 | 316 | 299 | 659 | 380 | 542 | 1038 | 776 | 765 | 1429 | 1009 | 1474 | 1031.8 | 860 | 1304 | 1.383 | 0.467 | 0.00229481 |
| AT1G15310.1 | SRP-54A | protein_coding | signal recognition particl | 50 | 51 | 42 | 82 | 75 | 56 | 105 | 125 | 108 | 178 | 199 | 152 | 160.98 | 113 | 176 | 1.383 | 0.467 | 0.00186133 |
| AT3G46600.1 | SCL30 | protein_coding | GRAS family transcription | 298 | 335 | 372 | 407 | 353 | 423 | 626 | 823 | 952 | 883 | 937 | 1150 | 711.23 | 800 | 990 | 1.382 | 0.467 | 0.00096091 |
| AT5G40880.1 | ZFWD3 | protein_coding | WD-40 repeat family prot | 118 | 99 | 83 | 138 | 116 | 94 | 248 | 243 | 212 | 299 | 308 | 256 | 247.54 | 234 | 288 | 1.382 | 0.467 | 0.00079608 |
| AT5G41070.1 | DRB5 | protein_coding | dsRNA-binding protein 5 | 88 | 81 | 90 | 124 | 123 | 142 | 185 | 199 | 230 | 269 | 327 | 386 | 312.59 | 205 | 327 | 1.382 | 0.467 | 0.0007903 |
| AT1G15710.1 | TYRAAT2 | protein_coding | prephenate dehydrogena | 257 | 204 | 155 | 283 | 210 | 193 | 540 | 501 | 397 | 614 | 558 | 525 | 708.75 | 479 | 566 | 1.382 | 0.467 | 0.00029724 |
| AT3G59765.1 | | other_rna | other RNA | 84 | 88 | 78 | 140 | 130 | 110 | 177 | 216 | 200 | 304 | 345 | 299 | 277.1 | 198 | 316 | 1.382 | 0.467 | 0.00018958 |
| AT3G54280.2 | BTAF1 | protein_coding | DNA binding;ATP binding | 1747 | 1315 | 1274 | 1797 | 1687 | 1330 | 3672 | 3231 | 3262 | 3898 | 4479 | 3617 | 2653.6 | 3388 | 3998 | 1.382 | 0.467 | 0.00016361 |
| AT1G67620.1 | | protein_coding | Lojap-related protein | 141 | 85 | 78 | 134 | 83 | 112 | 296 | 209 | 200 | 291 | 220 | 305 | 285.77 | 235 | 272 | 1.381 | 0.466 | 0.00500349 |
| AT3G48200.1 | | protein_coding | | 3050 | 2334 | 1975 | 3904 | 2844 | 1996 | 6411 | 5734 | 5056 | 8468 | 7550 | 5428 | 5400.17 | 5734 | 7149 | 1.381 | 0.466 | 0.00244162 |
| AT4G25710.1 | | protein_coding | Galactose oxidase/kelch r | 112 | 79 | 68 | 122 | 74 | 94 | 235 | 194 | 174 | 265 | 196 | 256 | 210.23 | 201 | 239 | 1.381 | 0.466 | 0.00184132 |
| AT3G53500.2 | RS2Z32 | protein_coding | RNA-binding (RRM/RBD/ | 614 | 626 | 503 | 759 | 619 | 771 | 1291 | 1538 | 1288 | 1646 | 1643 | 2097 | 2062.01 | 1372 | 1795 | 1.382 | 0.466 | 0.00036718 |
| AT5G52920.1 | PKP2 | protein_coding | plastidic pyruvate kinase | 4822 | 4242 | 4796 | 5434 | 4454 | 4997 | 10136 | 10422 | 12278 | 11787 | 11824 | 13589 | 10216.09 | 10945 | 12400 | 1.381 | 0.466 | 0.00031859 |
| AT3G14890.1 | ZDP | protein_coding | phosphoesterase | 268 | 252 | 204 | 335 | 310 | 252 | 563 | 619 | 522 | 727 | 823 | 685 | 679.22 | 568 | 745 | 1.381 | 0.466 | 1.92E-05 |
| AT1G16300.1 | GAPCP2 | protein_coding | glyceraldehyde-3-phosph | 186 | 103 | 95 | 184 | 130 | 134 | 391 | 253 | 243 | 399 | 345 | 364 | 341.79 | 296 | 369 | 1.38 | 0.465 | 0.00745632 |
| AT3G59670.1 | | protein_coding | | 370 | 207 | 209 | 414 | 356 | 284 | 778 | 509 | 535 | 898 | 945 | 772 | 643.79 | 607 | 872 | 1.38 | 0.465 | 0.00403429 |
| AT2G18330.1 | | protein_coding | AAA-type ATPase family p | 344 | 227 | 176 | 388 | 256 | 284 | 723 | 558 | 451 | 842 | 680 | 772 | 640.65 | 577 | 765 | 1.38 | 0.465 | 0.00267472 |
| AT4G17870.1 | PYR1 | protein_coding | Polyketide cyclase/dehyd | 456 | 465 | 362 | 598 | 347 | 468 | 959 | 1142 | 927 | 1297 | 921 | 1273 | 1416.63 | 1009 | 1164 | 1.38 | 0.465 | 0.00245012 |
| AT5G47210.1 | | protein_coding | Hyaluronan / mRNA bind | 5044 | 4155 | 2928 | 5386 | 4102 | 4708 | 10603 | 10208 | 7496 | 11683 | 10890 | 12803 | 14833.82 | 9436 | 11792 | 1.38 | 0.465 | 0.00177738 |
| AT1G11800.1 | | protein_coding | endonuclease/exonucleas | 131 | 113 | 125 | 190 | 128 | 140 | 275 | 278 | 320 | 412 | 340 | 381 | 334.72 | 291 | 378 | 1.38 | 0.465 | 0.00038524 |
| AT5G66055.1 | AKRP | protein_coding | ankyrin repeat protein | 737 | 697 | 599 | 780 | 719 | 833 | 1549 | 1712 | 1533 | 1692 | 1909 | 2265 | 2256.57 | 1598 | 1955 | 1.38 | 0.465 | 0.00032711 |
| AT2G38770.1 | EMB2765 | protein_coding | P-loop containing nucleo | 1646 | 1195 | 1148 | 1887 | 1439 | 1301 | 3460 | 2936 | 2939 | 4093 | 3820 | 3538 | 2398.45 | 3112 | 3817 | 1.38 | 0.465 | 0.00029699 |
| AT1G76300.1 | SMD3A | protein_coding | snRNP core protein SMD3 | 224 | 162 | 133 | 246 | 197 | 199 | 471 | 398 | 340 | 534 | 523 | 541 | 580.28 | 403 | 533 | 1.381 | 0.465 | 0.00029686 |
| AT1G80380.2 | GLYK | protein_coding | P-loop containing nucleo | 3518 | 3635 | 3323 | 4211 | 3915 | 4129 | 7395 | 8931 | 8507 | 9134 | 10393 | 11229 | 11892.54 | 8278 | 10252 | 1.38 | 0.465 | 0.0002815 |
| AT2G40900.1 | | protein_coding | nodulin MtN21 /EamA-lik | 66 | 57 | 32 | 121 | 94 | 91 | 139 | 140 | 82 | 262 | 250 | 247 | 185.29 | 120 | 253 | 1.38 | 0.464 | 0.00683788 |
| AT1G34030.1 | RPS18A | protein_coding | Ribosomal protein S13/S1 | 2648 | 1665 | 1289 | 2938 | 2004 | 1668 | 5566 | 4091 | 3300 | 6373 | 5320 | 4536 | 4907.82 | 4319 | 5410 | 1.379 | 0.464 | 0.00572363 |
| AT3G48420.1 | | protein_coding | Haloacid dehalogenase-li | 2673 | 2768 | 2505 | 3492 | 3000 | 3039 | 5619 | 6800 | 6413 | 7574 | 7964 | 8264 | 8342.23 | 6277 | 7934 | 1.379 | 0.464 | 0.0015972 |
| AT2G47450.1 | CAO | protein_coding | chloroplast signal recogni | 1498 | 1888 | 1079 | 1827 | 1710 | 2107 | 3149 | 4638 | 2762 | 3963 | 4540 | 5730 | 8474.68 | 3516 | 4744 | 1.379 | 0.464 | 0.00830841 |
| AT4G18140.1 | SSP4b | protein_coding | SCP1-like small phosphat | 153 | 178 | 207 | 269 | 242 | 250 | 322 | 437 | 530 | 583 | 642 | 680 | 431.57 | 430 | 635 | 1.378 | 0.463 | 0.00399837 |
| AT2G42330.1 | STIPL2 | protein_coding | GC-rich sequence DNA-bi | 95 | 101 | 107 | 158 | 143 | 151 | 200 | 248 | 274 | 343 | 380 | 411 | 326.47 | 241 | 378 | 1.379 | 0.463 | 0.0023574 |
| AT4G16530.1 | | protein_coding | Family of unknown functi | 48 | 47 | 42 | 49 | 28 | 49 | 101 | 115 | 108 | 106 | 74 | 133 | 204.7 | 108 | 104 | 1.378 | 0.463 | 0.00221532 |
| AT1G17760.1 | CSTF77 | protein_coding | Tetratricopeptide repeat | 443 | 357 | 324 | 509 | 420 | 370 | 931 | 877 | 829 | 1104 | 1115 | 1006 | 875.14 | 879 | 1075 | 1.379 | 0.463 | 3.28E-05 |
| AT4G34000.1 | ABF3 | protein_coding | abscisic acid responsive e | 261 | 251 | 211 | 422 | 385 | 370 | 549 | 617 | 540 | 915 | 1022 | 1006 | 798.92 | 569 | 981 | 1.378 | 0.463 | 2.07E-05 |
| AT1G06870.1 | TPP2 | protein_coding | Peptidase S24/S26A/S26 | 235 | 219 | 187 | 301 | 253 | 223 | 494 | 538 | 479 | 653 | 672 | 606 | 616.89 | 504 | 644 | 1.377 | 0.462 | 7.10E-06 |
| AT1G53280.1 | DJ1B | protein_coding | Class I glutamine amidotr | 451 | 323 | 308 | 579 | 427 | 562 | 948 | 794 | 788 | 1256 | 1134 | 1528 | 1135.24 | 843 | 1306 | 1.376 | 0.461 | 0.00096819 |
| AT5G50100.1 | | protein_coding | Putative thiol-disulphide | 397 | 332 | 330 | 626 | 452 | 367 | 835 | 816 | 845 | 1358 | 1200 | 998 | 937.84 | 832 | 1185 | 1.377 | 0.461 | 0.00042262 |
| AT2G28230.1 | MED20A | protein_coding | TATA-binding related fact | 146 | 134 | 128 | 176 | 135 | 108 | 307 | 329 | 328 | 382 | 358 | 294 | 321.96 | 321 | 345 | 1.377 | 0.461 | 0.00018551 |
| AT1G48920.1 | NUCL1 | protein_coding | nucleolin like 1 | 6686 | 4641 | 3256 | 5037 | 4195 | 3334 | 14054 | 11402 | 8336 | 10926 | 11137 | 9067 | 13653.99 | 11264 | 10377 | 1.375 | 0.46 | 0.00668745 |
| AT4G22690.1 | CYP706A1 | protein_coding | cytochrome P450, family | 1387 | 1178 | 755 | 1915 | 1231 | 999 | 2916 | 2894 | 1933 | 4154 | 3268 | 2717 | 3792.18 | 2581 | 3380 | 1.375 | 0.46 | 0.00601929 |
| AT1G20220.1 | | protein_coding | Alba DNA/RNA-binding p | 186 | 161 | 158 | 231 | 162 | 143 | 391 | 396 | 404 | 501 | 430 | 389 | 1289 | 397 | 440 | 1.376 | 0.46 | 0.00066999 |
| AT5G62070.1 | IQD23 | protein_coding | IQ-domain 23 | 220 | 197 | 125 | 244 | 209 | 181 | 462 | 484 | 320 | 529 | 555 | 492 | 504.48 | 422 | 525 | 1.376 | 0.46 | 0.0003234 |
| AT1G15270.1 | | protein_coding | Translation machinery as | 1705 | 1306 | 1258 | 2235 | 1605 | 1611 | 3584 | 3209 | 3221 | 4848 | 4261 | 4381 | 4106.92 | 3338 | 4497 | 1.376 | 0.46 | 9.61E-05 |
| AT2G34570.1 | MEE21 | protein_coding | PIN domain-like family pr | 196 | 141 | 105 | 169 | 141 | 108 | 412 | 346 | 269 | 367 | 374 | 294 | 351.81 | 342 | 345 | 1.375 | 0.459 | 0.00136005 |
| AT4G21070.1 | BRCA1 | protein_coding | breast cancer susceptibili | 308 | 269 | 239 | 324 | 309 | 249 | 647 | 661 | 612 | 703 | 820 | 677 | 501.47 | 640 | 733 | 1.374 | 0.459 | 0.00062921 |
| AT2G42740.1 | RPL11A | protein_coding | ribosomal protein large s | 232 | 165 | 145 | 260 | 204 | 180 | 488 | 405 | 371 | 564 | 542 | 489 | 543.26 | 421 | 532 | 1.375 | 0.459 | 0.00050551 |
| AT5G17890.1 | DAR4 | protein_coding | DA1-related protein 4 | 1185 | 869 | 830 | 1139 | 779 | 784 | 2491 | 2135 | 2125 | 2471 | 2068 | 2132 | 1751.16 | 2250 | 2224 | 1.375 | 0.459 | 0.00016294 |
| AT5G14450.1 | | protein_coding | GDSL-like Lipase/Acylhyd | 242 | 211 | 196 | 327 | 243 | 232 | 509 | 518 | 502 | 709 | 645 | 631 | 580.87 | 510 | 662 | 1.374 | 0.459 | 1.73E-05 |
| AT1G08710.2 | SKIP24 | protein_coding | F-box family protein | 89 | 68 | 60 | 81 | 78 | 63 | 187 | 167 | 154 | 176 | 207 | 171 | 193.73 | 169 | 185 | 1.373 | 0.458 | 0.00986733 |
| AT3G61440.1 | CYSC1 | protein_coding | cysteine synthase C1 | 8365 | 4658 | 5324 | 10438 | 6056 | 7397 | 17584 | 11444 | 13630 | 22641 | 16077 | 20116 | 16924.56 | 14219 | 19611 | 1.374 | 0.458 | 0.00552719 |
| AT5G64150.1 | | protein_coding | RNA methyltransferase fa | 116 | 137 | 108 | 142 | 125 | 144 | 244 | 337 | 276 | 308 | 332 | 392 | 372.53 | 286 | 344 | 1.373 | 0.458 | 0.00163087 |
| AT3G12340.1 | FKBP43 | protein_coding | FKBP-like peptidyl-prolyl | 225 | 146 | 143 | 242 | 201 | 150 | 473 | 359 | 366 | 525 | 534 | 408 | 414.71 | 399 | 489 | 1.373 | 0.458 | 0.00132754 |
| AT5G28919.1 | | protein_coding | | 185 | 192 | 149 | 190 | 215 | 237 | 389 | 472 | 381 | 412 | 571 | 645 | 855.13 | 414 | 543 | 1.374 | 0.458 | 0.00118122 |
| AT4G32690.1 | GLB3 | protein_coding | hemoglobin 3 | 115 | 65 | 68 | 171 | 116 | 114 | 242 | 160 | 174 | 371 | 308 | 310 | 287.95 | 192 | 330 | 1.374 | 0.458 | 0.00092552 |
| AT1G50250.1 | FTSH1 | protein_coding | FTSH protease 1 | 2519 | 2846 | 2609 | 3619 | 3236 | 3724 | 5295 | 6992 | 6679 | 7850 | 8591 | 10127 | 8290.57 | 6322 | 8856 | 1.373 | 0.458 | 0.00083984 |
| AT5G04740.1 | ACR12 | protein_coding | ACT domain-containing p | 1163 | 785 | 784 | 1428 | 1038 | 950 | 2445 | 1929 | 2007 | 3097 | 2756 | 2583 | 2579.69 | 2127 | 2812 | 1.373 | 0.458 | 0.00025215 |
| AT5G44750.2 | REV1 | protein_coding | DNA-directed DNA polym | 669 | 667 | 688 | 891 | 820 | 838 | 1406 | 1639 | 1761 | 1933 | 2177 | 2279 | 1814.64 | 1602 | 2130 | 1.374 | 0.458 | 0.00022554 |
| AT2G07680.1 | ATMRP11 | protein_coding | multidrug resistance-asso | 374 | 296 | 269 | 469 | 448 | 350 | 786 | 727 | 689 | 1017 | 1189 | 952 | 924.65 | 734 | 1053 | 1.373 | 0.458 | 0.0001176 |
| AT3G05070.1 | | protein_coding | | 325 | 246 | 221 | 380 | 278 | 269 | 683 | 604 | 566 | 824 | 738 | 732 | 701.64 | 618 | 765 | 1.373 | 0.458 | 2.86E-05 |
| AT3G16460.1 | JAL34 | protein_coding | Mannose-binding lectin s | 5436 | 4086 | 2760 | 6665 | 4227 | 3751 | 11427 | 10039 | 7066 | 14457 | 11222 | 10201 | 12807.28 | 9511 | 11960 | 1.373 | 0.457 | 0.00863028 |
| AT2G44690.1 | ARAC9 | protein_coding | Arabidopsis RAC-like 9 | 40 | 39 | 40 | 74 | 40 | 50 | 84 | 96 | 102 | 161 | 106 | 136 | 121.19 | 94 | 134 | 1.373 | 0.457 | 0.00774847 |
| AT4G00620.1 | FOLD4 | protein_coding | Amino acid dehydrogenas | 370 | 211 | 182 | 406 | 252 | 264 | 778 | 518 | 466 | 881 | 669 | 718 | 711.47 | 587 | 756 | 1.373 | 0.457 | 0.00565974 |
| AT4G39880.1 | | protein_coding | Ribosomal protein L23/L1 | 328 | 276 | 196 | 298 | 246 | 312 | 689 | 678 | 502 | 646 | 653 | 848 | 965.88 | 623 | 716 | 1.372 | 0.457 | 0.00385606 |
| AT1G32580.1 | MORF5 | protein_coding | plastid developmental pr | 510 | 300 | 287 | 534 | 397 | 388 | 1072 | 737 | 735 | 1158 | 1054 | 1055 | 1230.99 | 848 | 1089 | 1.372 | 0.457 | 0.00125878 |
| AT5G12470.1 | | protein_coding | Protein of unknown funct | 1728 | 1301 | 977 | 1748 | 1422 | 1193 | 3632 | 3196 | 2501 | 3792 | 3775 | 3244 | 4437.16 | 3110 | 3604 | 1.372 | 0.457 | 0.0008471 |
| AT1G31180.1 | IMDH1 | protein_coding | isopropylmalate dehydro | 1451 | 978 | 846 | 1556 | 1188 | 1028 | 3050 | 2403 | 2166 | 3375 | 3154 | 2796 | 3179.14 | 2540 | 3108 | 1.373 | 0.457 | 0.00080304 |
| AT2G19730.1 | RPL28A | protein_coding | Ribosomal L28e protein f | 3165 | 2585 | 1668 | 3087 | 2248 | 2296 | 6653 | 6351 | 4270 | 6696 | 5968 | 6244 | 7767.62 | 5758 | 6303 | 1.372 | 0.456 | 0.00308519 |
| AT1G54270.1 | TIF4A-2 | protein_coding | eif4a-2 | 2446 | 1983 | 1439 | 2960 | 2126 | 2152 | 5142 | 4872 | 3684 | 6421 | 5644 | 5852 | 5894.46 | 4566 | 5972 | 1.372 | 0.456 | 0.00043876 |

| | | | | | | | | | | | | | | | | | | | | |
|---|---|---|---|---|---|---|---|---|---|---|---|---|---|---|---|---|---|---|---|---|
| AT1G50010.1 | TUBA4 | protein_coding | tubulin alpha-2 chain | 3746 | 4728 | 3527 | 4338 | 4580 | 4734 | 7874 | 11616 | 9029 | 9410 | 12159 | 12874 | 14311.76 | 9506 | 11481 | 1.371 | 0.455 | 0.00455635 |
| AT1G79550.1 | PGK | protein_coding | phosphoglycerate kinase | 4397 | 3242 | 2369 | 5588 | 3890 | 3455 | 9243 | 7965 | 6065 | 12121 | 10327 | 9396 | 9463.1 | 7758 | 10615 | 1.371 | 0.455 | 0.00242898 |
| AT3G18210.1 | | protein_coding | 2-oxoglutarate (2OG) and | 403 | 346 | 268 | 475 | 433 | 391 | 847 | 850 | 686 | 1030 | 1150 | 1063 | 921.44 | 794 | 1081 | 1.371 | 0.455 | 8.08E-05 |
| AT5G09920.1 | NRPB4 | protein_coding | RNA polymerase II, Rpb4, | 473 | 350 | 392 | 649 | 507 | 469 | 994 | 860 | 1004 | 1408 | 1346 | 1275 | 1122.18 | 953 | 1343 | 1.371 | 0.455 | 6.52E-05 |
| AT1G44920.1 | | protein_coding | | 1042 | 820 | 756 | 1220 | 932 | 979 | 2190 | 2015 | 1935 | 2646 | 2474 | 2662 | 3231.68 | 2047 | 2594 | 1.371 | 0.455 | 5.44E-05 |
| AT1G12800.1 | | protein_coding | Nucleic acid-binding, OB- | 6404 | 5824 | 4769 | 6533 | 5978 | 5182 | 13462 | 14308 | 12209 | 14171 | 15870 | 14092 | 14254.22 | 13326 | 14711 | 1.369 | 0.454 | 0.00030217 |
| AT5G16760.1 | ITPK1 | protein_coding | Inositol 1,3,4-trisphospha | 717 | 585 | 582 | 823 | 647 | 579 | 1507 | 1437 | 1490 | 1785 | 1718 | 1575 | 1573.26 | 1478 | 1693 | 1.37 | 0.454 | 4.75E-05 |
| AT2G21180.1 | | protein_coding | | 41 | 32 | 40 | 61 | 48 | 53 | 86 | 79 | 102 | 132 | 127 | 144 | 128.98 | 89 | 134 | 1.369 | 0.453 | 0.00674134 |
| AT4G22380.1 | | protein_coding | Ribosomal protein L7Ae/L | 279 | 195 | 196 | 324 | 278 | 203 | 586 | 479 | 502 | 703 | 738 | 552 | 554.48 | 522 | 664 | 1.369 | 0.453 | 0.00501667 |
| AT3G17185.1 | TASIR-ARF | other_rna | TAS3/TASIR-ARF (TRANS- | 120 | 96 | 124 | 237 | 146 | 136 | 252 | 236 | 317 | 514 | 388 | 370 | 362.74 | 268 | 424 | 1.369 | 0.453 | 0.00116007 |
| AT4G02400.1 | | protein_coding | U3 ribonucleoprotein (Ut | 1009 | 686 | 612 | 901 | 713 | 613 | 2121 | 1685 | 1567 | 1954 | 1893 | 1667 | 1580.38 | 1791 | 1838 | 1.369 | 0.453 | 0.00084631 |
| AT1G18500.1 | IPMS1 | protein_coding | methylthioalkylmalate sy | 1634 | 1234 | 1350 | 1605 | 1285 | 1410 | 3435 | 3032 | 3456 | 3481 | 3411 | 3834 | 3300.43 | 3308 | 3575 | 1.369 | 0.453 | 0.00013801 |
| AT5G51010.1 | | protein_coding | Rubredoxin-like superfam | 1613 | 1295 | 1101 | 2076 | 1690 | 1723 | 3391 | 3182 | 2819 | 4503 | 4487 | 4686 | 4801.82 | 3131 | 4559 | 1.369 | 0.453 | 5.77E-05 |
| AT1G25175.1 | | other_rna | other RNA | 978 | 1012 | 1175 | 1601 | 1525 | 1834 | 2056 | 2486 | 3008 | 3473 | 4049 | 4987 | 4650.57 | 2517 | 4170 | 1.368 | 0.452 | 0.0031969 |
| AT5G45330.1 | DCP5-L | protein_coding | decapping 5-like | 117 | 89 | 105 | 115 | 128 | 118 | 246 | 219 | 269 | 249 | 340 | 321 | 286.77 | 245 | 303 | 1.368 | 0.452 | 0.0022091 |
| AT5G20360.1 | Phox3 | protein_coding | Octicosapeptide/Phox/Be | 427 | 277 | 289 | 455 | 342 | 282 | 898 | 681 | 740 | 987 | 908 | 767 | 647.34 | 773 | 887 | 1.368 | 0.452 | 0.00098683 |
| AT1G01990.1 | | protein_coding | | 110 | 121 | 96 | 159 | 129 | 149 | 231 | 297 | 246 | 345 | 342 | 405 | 411.37 | 258 | 364 | 1.368 | 0.452 | 0.00024368 |
| AT4G23570.3 | SGT1A | protein_coding | phosphatase-related | 425 | 364 | 306 | 329 | 288 | 267 | 893 | 894 | 783 | 714 | 765 | 726 | 880.22 | 857 | 735 | 1.368 | 0.452 | 5.20E-05 |
| AT2G40960.1 | | protein_coding | Single-stranded nucleic ac | 228 | 250 | 200 | 176 | 199 | 158 | 479 | 614 | 512 | 382 | 528 | 430 | 711.7 | 535 | 447 | 1.367 | 0.451 | 0.001346 |
| AT1G79050.1 | RECA | protein_coding | recA DNA recombination | 337 | 301 | 207 | 458 | 316 | 293 | 708 | 740 | 530 | 993 | 839 | 797 | 1039.81 | 659 | 876 | 1.367 | 0.451 | 0.00065674 |
| AT1G22410.1 | | protein_coding | Class-II DAHP synthetase | 2072 | 1784 | 1392 | 2745 | 1941 | 1849 | 4355 | 4383 | 3564 | 5954 | 5153 | 5028 | 5033.22 | 4101 | 5378 | 1.367 | 0.451 | 0.00020783 |
| AT4G33060.1 | CYP57 | protein_coding | Cyclophilin-like peptidyl-p | 346 | 337 | 319 | 386 | 309 | 342 | 727 | 828 | 817 | 837 | 820 | 930 | 737.12 | 791 | 862 | 1.367 | 0.451 | 8.50E-05 |
| AT2G28000.1 | CPN60A1 | protein_coding | chaperonin-60alpha | 18089 | 13054 | 9261 | 15888 | 12078 | 11117 | 38024 | 32071 | 23709 | 34463 | 32064 | 30232 | 29410.59 | 31268 | 32253 | 1.366 | 0.45 | 0.00690676 |
| AT1G16970.1 | KU70 | protein_coding | KU70 homolog | 310 | 203 | 216 | 348 | 303 | 198 | 652 | 499 | 553 | 755 | 804 | 538 | 578.83 | 568 | 699 | 1.366 | 0.45 | 0.00151894 |
| AT2G24280.1 | | protein_coding | alpha/beta-Hydrolases su | 1115 | 870 | 895 | 1824 | 1256 | 971 | 2344 | 2137 | 2291 | 3956 | 3334 | 2641 | 2549.01 | 2257 | 3310 | 1.366 | 0.45 | 0.00124489 |
| AT3G03640.1 | BGLU25 | protein_coding | beta glucosidase 25 | 338 | 239 | 231 | 285 | 262 | 230 | 711 | 587 | 591 | 618 | 696 | 625 | 559.16 | 630 | 646 | 1.366 | 0.45 | 0.00114306 |
| AT4G26190.1 | | protein_coding | Haloacid dehalogenase-li | 711 | 521 | 551 | 622 | 544 | 469 | 1495 | 1280 | 1411 | 1349 | 1444 | 1275 | 1304.11 | 1395 | 1356 | 1.366 | 0.45 | 0.00032316 |
| AT2G41460.1 | ARP | protein_coding | apurinic endonuclease-re | 235 | 188 | 177 | 277 | 210 | 179 | 494 | 462 | 453 | 601 | 558 | 487 | 460.94 | 470 | 549 | 1.366 | 0.45 | 0.00011126 |
| AT4G10710.1 | SPT16 | protein_coding | global transcription facto | 1391 | 1277 | 1212 | 1823 | 1472 | 1504 | 2924 | 3137 | 3103 | 3954 | 3908 | 4090 | 2695.81 | 3055 | 3984 | 1.366 | 0.45 | 6.41E-05 |
| AT1G71260.1 | WHY2 | protein_coding | WHIRLY 2 | 473 | 258 | 221 | 424 | 393 | 306 | 994 | 634 | 566 | 920 | 1043 | 832 | 945.62 | 731 | 932 | 1.365 | 0.449 | 0.00809327 |
| AT4G25340.1 | FKBP53 | protein_coding | FK506 BINDING PROTEIN | 772 | 473 | 410 | 687 | 518 | 435 | 1623 | 1162 | 1050 | 1490 | 1375 | 1183 | 1308.23 | 1278 | 1349 | 1.365 | 0.449 | 0.00687573 |
| AT1G76040.2 | CPK29 | protein_coding | calcium-dependent prote | 104 | 88 | 102 | 148 | 138 | 137 | 219 | 216 | 261 | 321 | 366 | 373 | 313.16 | 232 | 353 | 1.365 | 0.449 | 0.00623069 |
| AT1G80750.1 | RPL7A | protein_coding | Ribosomal protein L30/L7 | 1120 | 660 | 587 | 998 | 772 | 598 | 2354 | 1621 | 1503 | 2165 | 2049 | 1626 | 1852.79 | 1826 | 1947 | 1.365 | 0.449 | 0.00549948 |
| AT5G07800.1 | | protein_coding | Flavin-binding monooxyg | 73 | 66 | 74 | 108 | 94 | 69 | 153 | 162 | 189 | 234 | 250 | 188 | 186.49 | 168 | 224 | 1.364 | 0.448 | 0.00992772 |
| AT3G23000.1 | CIPK7 | protein_coding | CBL-interacting protein ki | 771 | 756 | 477 | 972 | 767 | 807 | 1621 | 1857 | 1221 | 2108 | 2036 | 2195 | 1978.42 | 1566 | 2113 | 1.364 | 0.448 | 0.00146543 |
| AT5G08290.1 | YLS8 | protein_coding | mRNA splicing factor, thio | 902 | 828 | 730 | 1303 | 1056 | 1166 | 1896 | 2034 | 1869 | 2826 | 2803 | 3171 | 3084.54 | 1933 | 2933 | 1.364 | 0.448 | 7.29E-05 |
| AT3G14390.1 | LYSA1 | protein_coding | Pyridoxal-dependent dec | 1117 | 741 | 551 | 1177 | 847 | 726 | 2348 | 1821 | 1411 | 2553 | 2249 | 1974 | 2175.15 | 1860 | 2259 | 1.363 | 0.447 | 0.00505108 |
| AT2G31670.1 | | protein_coding | Stress responsive alpha-b | 664 | 461 | 415 | 957 | 561 | 560 | 1396 | 1133 | 1062 | 2076 | 1489 | 1523 | 1972.99 | 1197 | 1696 | 1.363 | 0.447 | 0.00192651 |
| AT2G19540.1 | | protein_coding | Transducin family protein | 1032 | 725 | 612 | 1303 | 906 | 896 | 2169 | 1781 | 1567 | 2826 | 2405 | 2437 | 2292.58 | 1839 | 2556 | 1.363 | 0.447 | 0.00111466 |
| AT3G20260.1 | | protein_coding | Protein of unknown funct | 277 | 219 | 225 | 288 | 239 | 177 | 582 | 538 | 576 | 625 | 634 | 481 | 523.59 | 565 | 580 | 1.363 | 0.447 | 0.00047792 |
| AT1G09760.1 | U2A' | protein_coding | U2 small nuclear ribonucl | 701 | 514 | 519 | 876 | 696 | 563 | 1474 | 1263 | 1329 | 1900 | 1848 | 1531 | 1556.77 | 1355 | 1760 | 1.362 | 0.446 | 0.00033195 |
| AT4G15900.1 | PRL1 | protein_coding | pleiotropic regulatory loc | 765 | 542 | 524 | 910 | 694 | 725 | 1608 | 1332 | 1341 | 1974 | 1842 | 1972 | 1917.15 | 1427 | 1929 | 1.362 | 0.446 | 0.00029801 |
| AT4G46800.1 | BOU | protein_coding | Mitochondrial substrate c | 1266 | 1139 | 838 | 1584 | 1287 | 1353 | 2661 | 2798 | 2145 | 3436 | 3417 | 3679 | 3657.45 | 2535 | 3511 | 1.362 | 0.446 | 0.00020915 |
| AT2G16380.1 | SFH7 | protein_coding | Sec14p-like phosphatidyl | 444 | 429 | 395 | 698 | 521 | 498 | 933 | 1054 | 1011 | 1514 | 1383 | 1354 | 1083.1 | 999 | 1417 | 1.362 | 0.446 | 7.14E-05 |
| AT4G26860.2 | | protein_coding | Predicted pyridoxal phos | 726 | 617 | 541 | 876 | 784 | 776 | 1526 | 1516 | 1385 | 1900 | 2081 | 2110 | 2270.74 | 1476 | 2030 | 1.362 | 0.446 | 5.50E-05 |
| AT1G48550.1 | | protein_coding | Vacuolar protein sorting- | 125 | 97 | 119 | 135 | 132 | 130 | 263 | 238 | 305 | 293 | 350 | 354 | 346.55 | 269 | 332 | 1.362 | 0.445 | 0.00110016 |
| AT1G32750.1 | TAF1 | protein_coding | HAC13 protein (HAC13) | 1172 | 1032 | 1078 | 1498 | 1224 | 1043 | 2464 | 2535 | 2760 | 3249 | 3249 | 2836 | 2166.88 | 2586 | 3111 | 1.362 | 0.445 | 9.95E-05 |
| AT4G23940.1 | | protein_coding | FtsH extracellular proteas | 1281 | 1088 | 964 | 1571 | 1397 | 1201 | 2693 | 2673 | 2468 | 3408 | 3709 | 3266 | 2993.15 | 2611 | 3461 | 1.361 | 0.445 | 7.97E-05 |
| AT2G19385.1 | | protein_coding | zinc ion binding | 229 | 146 | 108 | 217 | 174 | 138 | 481 | 359 | 276 | 471 | 462 | 375 | 433.21 | 372 | 436 | 1.361 | 0.444 | 0.00438675 |
| AT5G40490.1 | | protein_coding | RNA-binding (RRM/RBD/ | 103 | 68 | 75 | 104 | 94 | 84 | 217 | 167 | 192 | 226 | 250 | 228 | 485.21 | 192 | 235 | 1.361 | 0.444 | 0.00081313 |
| AT2G46590.2 | DOF2.5 | protein_coding | Dof-type zinc finger DNA- | 201 | 134 | 177 | 362 | 253 | 286 | 423 | 329 | 453 | 785 | 672 | 778 | 572.66 | 402 | 745 | 1.36 | 0.444 | 0.00030539 |
| AT3G25840.1 | | protein_coding | Protein kinase superfamil | 653 | 530 | 562 | 776 | 713 | 619 | 1373 | 1302 | 1439 | 1683 | 1893 | 1683 | 1647.57 | 1371 | 1753 | 1.361 | 0.444 | 0.00012474 |
| AT1G14810.1 | | protein_coding | semialdehyde dehydroge | 1634 | 1161 | 784 | 1644 | 1332 | 1115 | 3435 | 2852 | 2007 | 3566 | 3536 | 3032 | 3711.65 | 2765 | 3378 | 1.359 | 0.443 | 0.00567208 |
| AT2G44640.1 | | protein_coding | | 931 | 677 | 438 | 834 | 665 | 602 | 1957 | 1663 | 1121 | 1809 | 1765 | 1637 | 1937.52 | 1580 | 1737 | 1.359 | 0.443 | 0.00445274 |
| AT5G04940.1 | SUVH1 | protein_coding | SU(VAR)3-9 homolog 1 | 428 | 331 | 348 | 491 | 407 | 382 | 900 | 813 | 891 | 1065 | 1080 | 1039 | 849.84 | 868 | 1061 | 1.359 | 0.443 | 6.85E-05 |
| AT2G35750.1 | | protein_coding | | 15792 | 17262 | 12934 | 15132 | 14045 | 15786 | 33196 | 42410 | 33112 | 32823 | 37286 | 42929 | 18570.63 | 36239 | 37679 | 1.358 | 0.442 | 0.00883467 |
| AT3G20290.1 | RPS8A | protein_coding | Ribosomal protein S8e fa | 8356 | 7121 | 4625 | 8545 | 6431 | 7627 | 17565 | 17495 | 11840 | 18535 | 17073 | 20741 | 20875.98 | 15633 | 18783 | 1.358 | 0.442 | 0.00831305 |
| AT3G15950.1 | NAI2 | protein_coding | DNA topoisomerase-relat | 7853 | 5437 | 4688 | 10012 | 6317 | 5778 | 16508 | 13358 | 12002 | 21717 | 16770 | 15713 | 12590.65 | 13956 | 18067 | 1.359 | 0.442 | 0.00433441 |
| AT2G27470.1 | NF-YB11 | protein_coding | nuclear factor Y, subunit | 9 | 9 | 6 | 9 | 16 | 7 | 19 | 22 | 15 | 20 | 42 | 19 | 191.57 | 19 | 27 | 1.359 | 0.442 | 0.00375878 |
| AT1G17270.1 | | protein_coding | O-fucosyltransferase fam | 106 | 122 | 89 | 123 | 128 | 111 | 223 | 300 | 228 | 267 | 340 | 302 | 249.9 | 250 | 303 | 1.359 | 0.442 | 0.00280397 |
| AT4G02630.1 | | protein_coding | Protein kinase superfamil | 131 | 146 | 109 | 210 | 229 | 127 | 275 | 359 | 279 | 456 | 608 | 345 | 443.32 | 304 | 470 | 1.357 | 0.441 | 0.00887165 |
| AT2G30490.1 | CYP73A5 | protein_coding | cinnamate-4-hydroxylase | 1953 | 1597 | 1183 | 2765 | 2000 | 1883 | 4105 | 3924 | 3029 | 5998 | 5310 | 5121 | 4117.2 | 3686 | 5476 | 1.358 | 0.441 | 0.0007757 |
| AT2G21090.1 | ABCG15 | protein_coding | ABC-2 type transporter fa | 83 | 71 | 50 | 141 | 115 | 103 | 174 | 174 | 128 | 306 | 305 | 280 | 190.79 | 159 | 297 | 1.357 | 0.44 | 0.00665541 |
| AT3G16990.1 | TENA_E | protein_coding | Haem oxygenase-like, mu | 35 | 42 | 36 | 68 | 54 | 42 | 74 | 103 | 92 | 147 | 143 | 114 | 136.48 | 90 | 135 | 1.357 | 0.44 | 0.00627165 |
| AT4G27340.1 | | protein_coding | Met-10+ like family prote | 439 | 256 | 227 | 348 | 268 | 233 | 923 | 629 | 581 | 755 | 711 | 634 | 717.65 | 711 | 700 | 1.357 | 0.44 | 0.00395346 |
| AT5G03740.1 | HDT3 | protein_coding | histone deacetylase 2C | 592 | 560 | 386 | 800 | 656 | 536 | 1244 | 1376 | 988 | 1735 | 1742 | 1458 | 1777.27 | 1203 | 1645 | 1.357 | 0.44 | 0.0009063 |
| AT3G43510.1 | | transposable_eler | transposable element ger | 233 | 151 | 178 | 251 | 178 | 193 | 490 | 371 | 456 | 544 | 473 | 525 | 464.6 | 439 | 514 | 1.357 | 0.44 | 0.00043292 |
| AT3G24460.1 | | protein_coding | Serinc-domain containing | 110 | 99 | 95 | 106 | 106 | 75 | 231 | 243 | 243 | 230 | 281 | 204 | 195.62 | 239 | 238 | 1.356 | 0.439 | 0.00518997 |
| AT2G19480.1 | NAP1 | protein_coding | nucleosome assembly pro | 2624 | 1686 | 1680 | 2680 | 1970 | 1834 | 5516 | 4142 | 4301 | 5813 | 5230 | 4987 | 4877.66 | 4653 | 5343 | 1.355 | 0.439 | 0.00100648 |
| AT3G01790.1 | | protein_coding | Ribosomal protein L13 fa | 309 | 248 | 206 | 343 | 246 | 275 | 650 | 609 | 527 | 744 | 653 | 748 | 831.45 | 595 | 715 | 1.356 | 0.439 | 0.00092995 |
| AT4G27690.1 | VPS26B | protein_coding | vacuolar protein sorting 2 | 205 | 164 | 205 | 210 | 149 | 183 | 431 | 403 | 525 | 456 | 396 | 498 | 421.87 | 453 | 450 | 1.356 | 0.439 | 0.00070023 |
| AT1G10270.1 | GRP23 | protein_coding | glutamine-rich protein 23 | 230 | 188 | 133 | 264 | 212 | 228 | 483 | 462 | 340 | 573 | 563 | 620 | 542.29 | 428 | 585 | 1.355 | 0.439 | 0.00054816 |
| AT2G18440.1 | GUT15 | other_rna | GUT15 (GENE WITH UNST | 848 | 743 | 891 | 1071 | 947 | 916 | 1783 | 1825 | 2281 | 2323 | 2514 | 2491 | 3285.08 | 1963 | 2443 | 1.356 | 0.439 | 0.00033371 |
| AT2G39740.1 | | protein_coding | Nucleotidyltransferase fa | 227 | 185 | 162 | 306 | 210 | 194 | 477 | 455 | 415 | 664 | 558 | 528 | 457.65 | 449 | 583 | 1.356 | 0.439 | 0.00017661 |
| AT5G55220.1 | TIG | protein_coding | trigger factor type chaper | 4867 | 3519 | 3155 | 4606 | 3613 | 3283 | 10231 | 8646 | 8077 | 9991 | 9592 | 8928 | 8767.35 | 8985 | 9504 | 1.354 | 0.438 | 0.00115417 |
| AT1G70700.1 | TIFY7 | protein_coding | TIFY domain/Divergent C | 264 | 272 | 218 | 448 | 331 | 295 | 555 | 668 | 558 | 972 | 879 | 802 | 698.34 | 594 | 884 | 1.354 | 0.438 | 0.00084072 |
| AT1G15410.1 | | protein_coding | aspartate-glutamate race | 340 | 267 | 259 | 558 | 369 | 347 | 715 | 656 | 663 | 1210 | 980 | 944 | 864.61 | 678 | 1045 | 1.355 | 0.438 | 0.00046674 |
| AT5G40930.1 | TOM20-4 | protein_coding | translocase of outer mem | 314 | 217 | 216 | 374 | 246 | 276 | 660 | 533 | 553 | 811 | 653 | 751 | 751.05 | 582 | 738 | 1.354 | 0.438 | 0.00037717 |
| AT4G35630.1 | PSAT1 | protein_coding | phosphoserine aminotrar | 829 | 500 | 497 | 935 | 608 | 522 | 1743 | 1228 | 1272 | 2028 | 1614 | 1420 | 1361.74 | 1414 | 1687 | 1.354 | 0.437 | 0.00542638 |
| AT3G21240.1 | 4CL2 | protein_coding | 4-coumarate:CoA ligase 2 | 76 | 64 | 49 | 94 | 92 | 63 | 160 | 157 | 125 | 204 | 244 | 171 | 217.61 | 147 | 206 | 1.354 | 0.437 | 0.00488748 |
| AT1G72130.1 | NPF5.11 | protein_coding | Major facilitator superfar | 654 | 460 | 359 | 750 | 568 | 423 | 1375 | 1130 | 919 | 1627 | 1508 | 1150 | 1538.95 | 1141 | 1428 | 1.353 | 0.437 | 0.00428238 |
| AT1G47860.1 | | transposable_eler | transposable element ger | 134 | 138 | 153 | 215 | 143 | 183 | 282 | 339 | 392 | 466 | 380 | 498 | 358.93 | 338 | 448 | 1.354 | 0.437 | 0.00398708 |
| AT3G53870.1 | RPS3B | protein_coding | Ribosomal protein S3 fam | 5523 | 4315 | 3072 | 5852 | 4135 | 4128 | 11610 | 10601 | 7865 | 12694 | 10977 | 11226 | 11293.42 | 10025 | 11632 | 1.353 | 0.437 | 0.00344387 |
| AT5G13740.1 | ZIF1 | protein_coding | zinc induced facilitator 1 | 340 | 327 | 420 | 472 | 475 | 388 | 715 | 803 | 1075 | 1024 | 1261 | 1055 | 905.21 | 864 | 1113 | 1.354 | 0.437 | 0.00177942 |
| AT1G70730.3 | PGM2 | protein_coding | Phosphoglucomutase/phe | 2688 | 1848 | 1540 | 2803 | 2089 | 1925 | 5650 | 4540 | 3942 | 6080 | 5546 | 5235 | 5334.87 | 4711 | 5620 | 1.354 | 0.437 | 0.00096827 |
| AT3G26240.1 | | protein_coding | Cysteine/Histidine-rich C1 | 379 | 332 | 382 | 404 | 293 | 327 | 797 | 816 | 978 | 876 | 778 | 889 | 668.53 | 864 | 848 | 1.354 | 0.437 | 0.00071494 |
| AT4G32770.1 | VTE1 | protein_coding | tocopherol cyclase, chlorc | 367 | 362 | 324 | 396 | 360 | 296 | 771 | 889 | 829 | 859 | 956 | 805 | 957.19 | 830 | 873 | 1.353 | 0.437 | 0.00046691 |
| AT5G10360.1 | RPS6B | protein_coding | Ribosomal protein S6e | 6181 | 4466 | 3052 | 6191 | 4437 | 4697 | 12993 | 10972 | 7813 | 13429 | 11779 | 12773 | 12801.03 | 10593 | 12660 | 1.353 | 0.436 | 0.00873785 |
| AT5G27750.1 | | protein_coding | F-box/FBD-like domains c | 84 | 77 | 56 | 100 | 90 | 66 | 177 | 189 | 143 | 217 | 239 | 179 | 179.74 | 170 | 212 | 1.353 | 0.436 | 0.00852907 |
| AT5G08180.1 | | protein_coding | Ribosomal protein L7Ae/L | 1148 | 729 | 604 | 1126 | 899 | 708 | 2413 | 1791 | 1546 | 2442 | 2387 | 1925 | 2108.09 | 1917 | 2251 | 1.353 | 0.436 | 0.00478052 |
| AT1G06550.1 | | protein_coding | ATP-dependent caseinoly | 492 | 321 | 300 | 672 | 524 | 406 | 1034 | 789 | 768 | 1458 | 1391 | 1104 | 1242.27 | 864 | 1318 | 1.353 | 0.436 | 0.00172242 |
| AT1G72970.1 | HTH | protein_coding | Glucose-methanol-cholin | 2096 | 1672 | 1162 | 2644 | 2241 | 1856 | 4406 | 4108 | 2975 | 5735 | 5949 | 5047 | 4574.79 | 3830 | 5577 | 1.353 | 0.436 | 0.00165315 |
| AT1G21060.1 | | protein_coding | Protein of unknown funct | 416 | 326 | 386 | 629 | 509 | 570 | 874 | 801 | 988 | 1364 | 1351 | 1550 | 1061.1 | 888 | 1422 | 1.353 | 0.436 | 0.00048602 |
| AT3G26560.1 | | protein_coding | ATP-dependent RNA helic | 807 | 738 | 569 | 961 | 844 | 692 | 1696 | 1813 | 1457 | 2085 | 2241 | 1882 | 1820.76 | 1655 | 2069 | 1.353 | 0.436 | 0.00047728 |
| AT1G31340.1 | RUB1 | protein_coding | related to ubiquitin 1 | 850 | 615 | 607 | 1112 | 897 | 826 | 1772 | 1511 | 1554 | 2412 | 2381 | 2246 | 1970.97 | 1612 | 2346 | 1.353 | 0.436 | 0.00031288 |
| AT5G08540.1 | | protein_coding | | 937 | 846 | 766 | 1105 | 862 | 1014 | 1970 | 2078 | 1961 | 2397 | 2288 | 2758 | 2632.75 | 2003 | 2481 | 1.353 | 0.436 | 0.00019444 |
| AT3G09100.2 | | protein_coding | mRNA capping enzyme fa | 814 | 705 | 772 | 1078 | 958 | 873 | 1711 | 1732 | 1976 | 2338 | 2543 | 2374 | 1709.52 | 1806 | 2418 | 1.353 | 0.436 | 0.00016232 |
| AT5G54640.1 | TSA1 | protein_coding | tryptophan synthase alph | 794 | 527 | 433 | 1088 | 691 | 568 | 1669 | 1295 | 1109 | 2360 | 1834 | 1545 | 1899.88 | 1358 | 1913 | 1.352 | 0.435 | 0.00627134 |
| AT5G03850.1 | RPS28A | protein_coding | Nucleic acid-binding, OB- | 2276 | 1561 | 1286 | 2540 | 1801 | 1612 | 4784 | 3835 | 3292 | 5510 | 4781 | 4384 | 4746.83 | 3970 | 4892 | 1.352 | 0.435 | 0.00442234 |
| AT4G22730.1 | | protein_coding | Leucine-rich repeat prote | 252 | 219 | 178 | 277 | 257 | 192 | 530 | 538 | 456 | 601 | 682 | 522 | 436.86 | 508 | 602 | 1.352 | 0.435 | 0.00098616 |
| AT1G73710.1 | | protein_coding | Pentatricopeptide repeat | 536 | 421 | 373 | 528 | 417 | 344 | 1127 | 1034 | 955 | 1145 | 1107 | 935 | 812.13 | 1039 | 1062 | 1.352 | 0.435 | 0.00032342 |
| AT3G21790.1 | UGT71B7 | protein_coding | UDP-Glycosyltransferase | 282 | 209 | 142 | 326 | 195 | 245 | 593 | 513 | 364 | 707 | 518 | 666 | 601.21 | 490 | 630 | 1.351 | 0.434 | 0.00553308 |
| AT1G77180.1 | SKIP | protein_coding | chromatin protein family | 1162 | 1148 | 1417 | 1459 | 1241 | 1620 | 2443 | 2820 | 3628 | 3165 | 3295 | 4405 | 3009.01 | 2964 | 3622 | 1.351 | 0.434 | 0.00547451 |
| AT1G55460.1 | | protein_coding | DNA/RNA-binding proteir | 70 | 76 | 46 | 83 | 62 | 56 | 147 | 187 | 118 | 180 | 165 | 152 | 438.44 | 151 | 166 | 1.351 | 0.434 | 0.00118386 |
| AT5G56740.1 | HAG2 | protein_coding | histone acetyltransferase | 351 | 310 | 268 | 448 | 419 | 302 | 738 | 762 | 686 | 972 | 1112 | 821 | 783.62 | 729 | 968 | 1.351 | 0.434 | 0.0004008 |
| AT3G04920.1 | RPS24A | protein_coding | Ribosomal protein S24e f | 3256 | 1990 | 1926 | 3155 | 2345 | 1935 | 6844 | 4889 | 4931 | 6843 | 6225 | 5262 | 5813.28 | 5555 | 6110 | 1.35 | 0.433 | 0.00378185 |
| AT3G17800.2 | | protein_coding | Protein of unknown funct | 929 | 1011 | 936 | 1262 | 972 | 1302 | 1953 | 2484 | 2396 | 2737 | 2580 | 3541 | 2699.11 | 2278 | 2953 | 1.35 | 0.433 | 0.00170226 |
| AT5G63060.1 | | protein_coding | Sec14p-like phosphatidyl | 229 | 179 | 176 | 259 | 222 | 169 | 481 | 440 | 451 | 562 | 589 | 460 | 625.6 | 457 | 537 | 1.35 | 0.433 | 0.00109269 |
| AT2G34750.1 | | protein_coding | RNA polymerase I specifi | 1538 | 1049 | 1027 | 1537 | 1225 | 1223 | 3233 | 2577 | 2629 | 3334 | 3252 | 3326 | 2837.29 | 2813 | 3304 | 1.35 | 0.433 | 0.00031602 |
| AT1G33410.1 | SAR1 | protein_coding | SUPPRESSOR OF AUXIN R | 1116 | 826 | 805 | 1271 | 1127 | 984 | 2346 | 2029 | 2061 | 2757 | 2992 | 2676 | 1948.25 | 2145 | 2808 | 1.35 | 0.433 | 0.00029302 |
| AT5G51120.2 | PABN1 | protein_coding | polyadenylate-binding pr | 316 | 280 | 236 | 392 | 292 | 312 | 664 | 688 | 604 | 850 | 775 | 848 | 945.94 | 652 | 824 | 1.35 | 0.432 | 8.66E-05 |
| AT3G58660.1 | | protein_coding | Ribosomal protein L1p/L1 | 532 | 326 | 272 | 423 | 380 | 310 | 1118 | 801 | 696 | 918 | 1009 | 843 | 1273.76 | 872 | 923 | 1.349 | 0.432 | 0.0068764 |
| AT3G24730.1 | | protein_coding | mRNA splicing factor, thio | 91 | 68 | 67 | 97 | 63 | 60 | 191 | 167 | 172 | 210 | 167 | 163 | 190.9 | 177 | 180 | 1.35 | 0.432 | 0.00553045 |
| AT3G08730.1 | ATPK1 | protein_coding | protein-serine kinase 1 | 1682 | 1034 | 953 | 1451 | 1045 | 898 | 3536 | 2540 | 2440 | 3147 | 2774 | 2442 | 2902.18 | 2839 | 2788 | 1.349 | 0.432 | 0.0046953 |
| AT1G11700.1 | | protein_coding | Protein of unknown funct | 63 | 88 | 39 | 118 | 104 | 104 | 132 | 216 | 100 | 256 | 276 | 283 | 306.63 | 149 | 272 | 1.349 | 0.432 | 0.00467535 |
| AT3G20330.1 | PYRB | protein_coding | PYRIMIDINE B | 658 | 390 | 380 | 773 | 593 | 516 | 1383 | 958 | 973 | 1677 | 1574 | 1403 | 1341.05 | 1105 | 1551 | 1.349 | 0.432 | 0.00236593 |
| AT3G56150.1 | TIF3C1 | protein_coding | eukaryotic translation ini | 5646 | 4154 | 3340 | 5986 | 4607 | 4347 | 11868 | 10206 | 8551 | 12984 | 12231 | 11821 | 9593.7 | 10208 | 12345 | 1.349 | 0.431 | 0.00110309 |
| AT1G29465.1 | | protein_coding | | 123 | 101 | 106 | 159 | 144 | 178 | 259 | 248 | 271 | 345 | 382 | 484 | 274.65 | 259 | 404 | 1.348 | 0.431 | 0.00367099 |
| AT2G29200.1 | APUM1 | protein_coding | pumilio 1 | 650 | 399 | 387 | 601 | 451 | 419 | 1366 | 980 | 991 | 1304 | 1197 | 1139 | 892.14 | 1112 | 1213 | 1.348 | 0.431 | 0.00195548 |
| AT5G14590.1 | T15N1_80 | protein_coding | Isocitrate/isopropylmalat | 859 | 574 | 502 | 975 | 664 | 719 | 1806 | 1410 | 1285 | 2115 | 1763 | 1955 | 1703.01 | 1500 | 1944 | 1.348 | 0.431 | 0.00163693 |
| AT5G51110.1 | | protein_coding | Transcriptional coactivato | 2758 | 2454 | 1835 | 2950 | 2539 | 2525 | 5798 | 6029 | 4698 | 6399 | 6740 | 6867 | 8933.86 | 5508 | 6669 | 1.348 | 0.431 | 0.00072593 |
| AT4G02730.1 | WDR5B | protein_coding | Transducin/WD40 repeat | 64 | 59 | 55 | 72 | 58 | 72 | 135 | 145 | 141 | 156 | 154 | 196 | 146.36 | 140 | 169 | 1.347 | 0.43 | 0.00844051 |
| AT5G65860.1 | | protein_coding | ankyrin repeat family pro | 364 | 240 | 180 | 306 | 252 | 214 | 765 | 590 | 461 | 664 | 669 | 582 | 610.55 | 605 | 638 | 1.347 | 0.43 | 0.00297391 |
| AT3G15590.1 | | protein_coding | Tetratricopeptide repeat | 623 | 410 | 424 | 660 | 472 | 418 | 1310 | 1007 | 1085 | 1432 | 1253 | 1137 | 1025.36 | 1134 | 1274 | 1.348 | 0.43 | 0.00141684 |
| AT3G32940.1 | | protein_coding | RNA-binding KH domain- | 129 | 125 | 141 | 169 | 134 | 166 | 271 | 307 | 361 | 367 | 356 | 451 | 501.62 | 313 | 391 | 1.347 | 0.43 | 0.00034861 |
| AT3G12050.1 | | protein_coding | Aha1 domain-containing | 331 | 287 | 241 | 346 | 238 | 270 | 696 | 705 | 617 | 751 | 632 | 734 | 1069.81 | 673 | 706 | 1.347 | 0.43 | 0.00030035 |
| AT1G63770.5 | | protein_coding | Peptidase M1 family prot | 4784 | 3980 | 3894 | 5033 | 4250 | 3924 | 10056 | 9778 | 9969 | 10917 | 11283 | 10671 | 9892.34 | 9934 | 10957 | 1.347 | 0.43 | 0.00015595 |
| AT4G38370.1 | | protein_coding | Phosphoglycerate mutase | 89 | 114 | 80 | 119 | 106 | 108 | 187 | 280 | 205 | 258 | 281 | 294 | 322.19 | 224 | 278 | 1.346 | 0.429 | 0.00171402 |
| AT3G25480.1 | STR4A | protein_coding | Rhodanese/Cell cycle con | 677 | 512 | 457 | 772 | 602 | 478 | 1423 | 1258 | 1170 | 1675 | 1598 | 1300 | 1718.09 | 1284 | 1524 | 1.347 | 0.429 | 0.00062925 |
| AT1G07615.1 | ATOBGM | protein_coding | GTP-binding protein Obg, | 96 | 69 | 63 | 81 | 50 | 65 | 202 | 170 | 161 | 176 | 133 | 177 | 204.97 | 178 | 162 | 1.345 | 0.428 | 0.00502138 |
| AT1G29900.1 | CARB | protein_coding | carbamoyl phosphate syn | 4732 | 3052 | 2618 | 5270 | 3838 | 3567 | 9947 | 7498 | 6702 | 11431 | 10189 | 9700 | 7294.44 | 8049 | 10440 | 1.345 | 0.428 | 0.00257597 |
| AT2G20690.1 | | protein_coding | lumazine-binding family p | 472 | 364 | 303 | 548 | 346 | 394 | 992 | 894 | 776 | 1189 | 919 | 1071 | 1530.73 | 887 | 1060 | 1.346 | 0.428 | 0.00207926 |
| AT5G67385.1 | | protein_coding | Phototropic-responsive N | 2437 | 2002 | 1744 | 3280 | 2637 | 2074 | 5123 | 4919 | 4465 | 7115 | 7001 | 5640 | 5600.78 | 4836 | 6585 | 1.345 | 0.428 | 0.00083938 |
| AT1G58080.1 | HISN1A | protein_coding | ATP phosphoribosyl trans | 2684 | 2230 | 1988 | 2950 | 2230 | 2272 | 5642 | 5479 | 5089 | 6399 | 5920 | 6179 | 6972.4 | 5403 | 6166 | 1.345 | 0.428 | 0.0001744 |
| AT2G13360.1 | AGT1 | protein_coding | alanine:glyoxylate amino | 16847 | 17962 | 17772 | 22813 | 19214 | 22806 | 35414 | 44129 | 45497 | 49484 | 51009 | 62020 | 52626.57 | 41680 | 54171 | 1.344 | 0.427 | 0.00948941 |
| AT1G09310.1 | | protein_coding | Protein of unknown funct | 13495 | 13180 | 10019 | 20424 | 17303 | 15526 | 28367 | 32381 | 25649 | 44302 | 45935 | 42222 | 46152.3 | 28799 | 44153 | 1.345 | 0.427 | 0.00581608 |
| AT3G52950.1 | CBSCBSPB3 | protein_coding | CBS / octicosapeptide/Ph | 499 | 349 | 300 | 574 | 379 | 329 | 1049 | 857 | 768 | 1245 | 1006 | 895 | 873.59 | 891 | 1049 | 1.344 | 0.427 | 0.00202296 |
| AT3G15351.1 | | protein_coding | | 76 | 75 | 68 | 101 | 97 | 71 | 160 | 184 | 174 | 219 | 258 | 193 | 219.25 | 173 | 223 | 1.344 | 0.426 | 0.00559365 |
| AT5G67030.1 | ZEP | protein_coding | zeaxanthin epoxidase (ZE | 5102 | 5365 | 5174 | 5823 | 5635 | 5001 | 10725 | 13181 | 13246 | 12631 | 14960 | 13600 | 13686.76 | 12384 | 13730 | 1.343 | 0.426 | 0.00145432 |
| AT1G64770.3 | PNSB2 | protein_coding | NDH-dependent cyclic ele | 1855 | 1390 | 1085 | 1705 | 1493 | 1252 | 3899 | 3415 | 2778 | 3698 | 3964 | 3405 | 4325.21 | 3364 | 3689 | 1.343 | 0.426 | 0.00144254 |
| AT5G26880.1 | AGL26 | protein_coding | AGAMOUS-like 26 | 322 | 284 | 266 | 431 | 347 | 427 | 677 | 698 | 681 | 935 | 921 | 1161 | 1042.07 | 685 | 1006 | 1.343 | 0.426 | 0.0004945 |
| AT1G23890.2 | | protein_coding | NHL domain-containing p | 77 | 78 | 83 | 136 | 106 | 107 | 162 | 192 | 212 | 295 | 281 | 291 | 225.24 | 189 | 289 | 1.342 | 0.425 | 0.00929828 |
| AT2G31840.1 | | protein_coding | Thioredoxin superfamily p | 355 | 231 | 189 | 317 | 238 | 200 | 746 | 568 | 484 | 688 | 632 | 544 | 910.18 | 599 | 621 | 1.342 | 0.425 | 0.00419265 |
| AT4G38170.1 | FRS9 | protein_coding | FAR1-related sequence 9 | 200 | 177 | 190 | 185 | 190 | 169 | 420 | 435 | 486 | 401 | 504 | 460 | 400.12 | 447 | 455 | 1.342 | 0.425 | 0.00081364 |
| AT4G17530.1 | RABD2C | protein_coding | RAB GTPase homolog 1C | 746 | 664 | 652 | 1158 | 836 | 930 | 1568 | 1631 | 1669 | 2512 | 2219 | 2529 | 2474.08 | 1623 | 2420 | 1.343 | 0.425 | 0.00026862 |

| | | | | | | | | | | | | | | | | | | | | | |
|---|---|---|---|---|---|---|---|---|---|---|---|---|---|---|---|---|---|---|---|---|---|
| AT1G57660.1 | RPL21F | protein_coding | Translation protein SH3-li | 2312 | 1650 | 1147 | 2126 | 1546 | 1457 | 4860 | 4054 | 2936 | 4611 | 4104 | 3962 | 4869.06 | 3950 | 4226 | 1.342 | 0.424 | 0.00829025 |
| AT3G47070.1 | | protein_coding | | 3051 | 3761 | 2903 | 4157 | 3214 | 3974 | 6413 | 9240 | 7432 | 9017 | 8532 | 10807 | 10064.77 | 7695 | 9452 | 1.341 | 0.424 | 0.00472061 |
| AT2G17360.1 | RPS4A | protein_coding | Ribosomal protein S4 (RP | 4767 | 3303 | 2668 | 4918 | 3413 | 3458 | 10021 | 8115 | 6830 | 10668 | 9061 | 9404 | 8649.71 | 8322 | 9711 | 1.342 | 0.424 | 0.00359628 |
| AT1G62740.1 | HOP2 | protein_coding | stress-inducible protein, p | 1016 | 791 | 694 | 1197 | 915 | 825 | 2136 | 1943 | 1777 | 2596 | 2429 | 2244 | 2009.28 | 1952 | 2423 | 1.341 | 0.423 | 0.00049532 |
| AT1G77480.1 | | protein_coding | Eukaryotic aspartyl prote | 310 | 250 | 280 | 332 | 278 | 247 | 652 | 614 | 717 | 720 | 738 | 672 | 592.72 | 661 | 710 | 1.34 | 0.423 | 0.0001731 |
| AT5G13400.1 | NPF6.1 | protein_coding | Major facilitator superfar | 870 | 689 | 470 | 1052 | 868 | 689 | 1829 | 1693 | 1203 | 2282 | 2304 | 1874 | 1970.67 | 1575 | 2153 | 1.339 | 0.422 | 0.00307942 |
| AT5G06110.1 | | protein_coding | DnaJ domain ;Myb-like D | 1401 | 927 | 804 | 1419 | 1194 | 1021 | 2945 | 2277 | 2058 | 3078 | 3170 | 2777 | 2522.27 | 2427 | 3008 | 1.34 | 0.422 | 0.00184949 |
| AT2G24500.1 | FZF | protein_coding | Zinc finger protein 622 | 729 | 513 | 454 | 706 | 521 | 472 | 1532 | 1260 | 1162 | 1531 | 1383 | 1284 | 1347.02 | 1318 | 1399 | 1.339 | 0.422 | 0.00166296 |
| AT5G44785.2 | OSB3 | protein_coding | organellar single-strande | 995 | 575 | 591 | 922 | 675 | 597 | 2092 | 1413 | 1513 | 2000 | 1792 | 1624 | 1752.24 | 1673 | 1805 | 1.339 | 0.421 | 0.00375944 |
| AT3G12345.1 | | protein_coding | | 2002 | 2270 | 2063 | 2654 | 2192 | 2962 | 4208 | 5577 | 5281 | 5757 | 5819 | 8055 | 6785.05 | 5022 | 6544 | 1.339 | 0.421 | 0.00366439 |
| AT1G30580.1 | | protein_coding | GTP binding | 3607 | 2242 | 2196 | 3605 | 2531 | 2314 | 7582 | 5508 | 5622 | 7820 | 6719 | 6293 | 6131.64 | 6237 | 6944 | 1.339 | 0.421 | 0.00236967 |
| AT4G18100.1 | RPL32A | protein_coding | Ribosomal protein L32e | 6265 | 4385 | 3245 | 5860 | 4419 | 3899 | 13169 | 10773 | 8307 | 12711 | 11731 | 10603 | 10841.37 | 10750 | 11682 | 1.338 | 0.42 | 0.00793244 |
| AT1G12230.2 | | protein_coding | Aldolase superfamily prot | 902 | 977 | 1016 | 1125 | 1034 | 1349 | 1896 | 2400 | 2601 | 2440 | 2745 | 3669 | 3192.45 | 2299 | 2951 | 1.338 | 0.42 | 0.00585316 |
| AT2G20020.1 | CAF1 | protein_coding | RNA-binding CRS1 / YhbY | 996 | 662 | 533 | 935 | 678 | 700 | 2094 | 1626 | 1365 | 2028 | 1800 | 1904 | 1796.67 | 1695 | 1911 | 1.338 | 0.42 | 0.0036809 |
| AT5G36950.1 | DEGP10 | protein_coding | DegP protease 10 | 303 | 232 | 198 | 357 | 315 | 211 | 637 | 570 | 507 | 774 | 836 | 574 | 624.45 | 571 | 728 | 1.338 | 0.42 | 0.00228738 |
| AT3G23325.1 | | protein_coding | Splicing factor 3B subunit | 183 | 125 | 108 | 257 | 176 | 173 | 385 | 307 | 276 | 557 | 467 | 470 | 444.91 | 323 | 498 | 1.338 | 0.42 | 0.00139548 |
| AT4G01000.1 | | protein_coding | Ubiquitin-like superfamily | 589 | 554 | 544 | 732 | 546 | 668 | 1238 | 1361 | 1393 | 1588 | 1450 | 1817 | 1586.75 | 1331 | 1618 | 1.338 | 0.42 | 0.00044855 |
| AT2G45680.1 | TCP9 | protein_coding | TCP family transcription f | 95 | 97 | 63 | 146 | 88 | 110 | 200 | 238 | 161 | 317 | 234 | 299 | 346.11 | 200 | 283 | 1.337 | 0.419 | 0.00518053 |
| AT3G63090.1 | | protein_coding | Ubiquitin carboxyl-termin | 150 | 101 | 94 | 136 | 92 | 112 | 315 | 248 | 241 | 295 | 244 | 305 | 269.97 | 268 | 281 | 1.337 | 0.419 | 0.0033241 |
| AT1G31500.4 | CCR4-4 | protein_coding | DNAse I-like superfamily | 463 | 314 | 271 | 473 | 384 | 301 | 973 | 771 | 694 | 1026 | 1019 | 819 | 900.78 | 813 | 955 | 1.337 | 0.419 | 0.002232 |
| AT1G04530.1 | TPR4 | protein_coding | Tetratricopeptide repeat | 810 | 706 | 599 | 1170 | 909 | 772 | 1703 | 1735 | 1533 | 2538 | 2413 | 2099 | 1862.14 | 1657 | 2350 | 1.337 | 0.419 | 0.00049068 |
| AT1G58025.2 | | protein_coding | DNA-binding bromodoma | 484 | 362 | 430 | 441 | 355 | 383 | 1017 | 889 | 1101 | 957 | 942 | 1042 | 846.56 | 1002 | 980 | 1.337 | 0.419 | 0.00042983 |
| AT5G58040.1 | FIPS5 | protein_coding | homolog of yeast FIP1 [V] | 701 | 657 | 553 | 693 | 631 | 633 | 1474 | 1614 | 1416 | 1503 | 1675 | 1721 | 1403.62 | 1501 | 1633 | 1.337 | 0.419 | 0.00040032 |
| AT1G30880.1 | | protein_coding | | 454 | 329 | 367 | 545 | 437 | 492 | 954 | 808 | 940 | 1182 | 1160 | 1338 | 1038.35 | 901 | 1227 | 1.337 | 0.419 | 0.00033347 |
| AT1G67250.1 | | protein_coding | Proteasome maturation f | 375 | 300 | 269 | 442 | 317 | 374 | 788 | 737 | 689 | 959 | 842 | 1017 | 995.99 | 738 | 939 | 1.337 | 0.419 | 0.00031366 |
| AT3G02570.1 | PMI1 | protein_coding | Mannose-6-phosphate iso | 1323 | 1169 | 959 | 1353 | 1068 | 1095 | 2781 | 2872 | 2455 | 2935 | 2835 | 2978 | 3153.11 | 2703 | 2916 | 1.337 | 0.419 | 0.00021095 |
| AT4G09320.1 | NDPK1 | protein_coding | Nucleoside diphosphate k | 6103 | 4475 | 3334 | 6178 | 4794 | 4321 | 12829 | 10994 | 8535 | 13401 | 12727 | 11751 | 13639.06 | 10786 | 12626 | 1.336 | 0.418 | 0.00641006 |
| AT3G54980.1 | | protein_coding | Pentatricopeptide repeat | 359 | 235 | 257 | 292 | 255 | 261 | 755 | 577 | 658 | 633 | 677 | 710 | 481.63 | 663 | 673 | 1.336 | 0.418 | 0.00269716 |
| AT5G17230.3 | PSY1 | protein_coding | PHYTOENE SYNTHASE | 5732 | 4428 | 3561 | 7590 | 5672 | 5087 | 12049 | 10879 | 9116 | 16463 | 15058 | 13834 | 11948.1 | 10681 | 15118 | 1.336 | 0.418 | 0.00166083 |
| AT1G54170.1 | CID3 | protein_coding | CTC-interacting domain 3 | 717 | 621 | 655 | 750 | 734 | 679 | 1507 | 1526 | 1677 | 1627 | 1949 | 1846 | 1389.27 | 1570 | 1807 | 1.336 | 0.418 | 0.00061921 |
| AT3G11270.1 | RPN8B | protein_coding | Mov34/MPN/PAD-1 fami | 263 | 195 | 202 | 301 | 269 | 241 | 553 | 479 | 517 | 653 | 714 | 655 | 577.86 | 516 | 674 | 1.336 | 0.418 | 0.00021117 |
| AT1G24360.1 | | protein_coding | NAD(P)-binding Rossman | 2118 | 1730 | 1716 | 2475 | 1897 | 2026 | 4452 | 4250 | 4393 | 5369 | 5036 | 5510 | 5244.92 | 4365 | 5305 | 1.336 | 0.418 | 0.0001055 |
| AT3G44630.3 | | protein_coding | Disease resistance protei | 1203 | 903 | 901 | 1378 | 1091 | 723 | 2529 | 2219 | 2307 | 2989 | 2896 | 1966 | 1785.08 | 2352 | 2617 | 1.335 | 0.417 | 0.00710799 |
| AT3G19370.1 | | protein_coding | Plant protein of unknowr | 321 | 230 | 180 | 358 | 320 | 217 | 675 | 565 | 461 | 777 | 850 | 590 | 662.13 | 567 | 739 | 1.335 | 0.417 | 0.00497849 |
| AT3G26710.1 | CCB1 | protein_coding | cofactor assembly of com | 2140 | 1574 | 1266 | 2261 | 1606 | 1371 | 4498 | 3867 | 3241 | 4904 | 4264 | 3728 | 4570.05 | 3869 | 4299 | 1.336 | 0.417 | 0.00311654 |
| AT4G24820.1 | RPN7 | protein_coding | 26S proteasome, regulato | 1738 | 1294 | 1351 | 2152 | 1614 | 1560 | 3653 | 3179 | 3459 | 4668 | 4285 | 4242 | 4284.13 | 3430 | 4398 | 1.335 | 0.417 | 0.0002356 |
| AT1G73960.1 | TAF2 | protein_coding | TBP-associated factor 2 | 893 | 751 | 724 | 999 | 904 | 848 | 1877 | 1845 | 1853 | 2167 | 2400 | 2306 | 1707 | 1858 | 2291 | 1.335 | 0.417 | 0.0001604 |
| AT3G03020.1 | | protein_coding | | 79 | 56 | 68 | 117 | 73 | 80 | 166 | 138 | 174 | 254 | 194 | 218 | 194.89 | 159 | 222 | 1.334 | 0.416 | 0.00736403 |
| AT3G06980.1 | RH50 | protein_coding | DEA(D/H)-box RNA helica | 2370 | 1678 | 1291 | 1933 | 1653 | 1293 | 4982 | 4123 | 3305 | 4193 | 4388 | 3516 | 4906.14 | 4137 | 4032 | 1.334 | 0.416 | 0.0038844 |
| AT3G46490.1 | | protein_coding | 2-oxoglutarate (2OG) and | 309 | 331 | 268 | 401 | 415 | 274 | 650 | 813 | 686 | 870 | 1102 | 745 | 939.36 | 716 | 906 | 1.334 | 0.416 | 0.00333718 |
| AT2G35240.1 | MORF6 | protein_coding | plastid developmental pr | 315 | 212 | 200 | 318 | 208 | 221 | 662 | 521 | 512 | 690 | 552 | 601 | 790.15 | 565 | 614 | 1.334 | 0.416 | 0.00330614 |
| AT4G18120.1 | AML3 | protein_coding | MEI2-like 3 | 621 | 572 | 646 | 847 | 738 | 733 | 1305 | 1405 | 1654 | 1837 | 1959 | 1993 | 1689.04 | 1455 | 1930 | 1.334 | 0.416 | 0.00069888 |
| AT1G51680.1 | 4CL1 | protein_coding | 4-coumarate:CoA ligase 1 | 874 | 1066 | 700 | 1030 | 1032 | 694 | 1837 | 2619 | 1792 | 2234 | 2740 | 1887 | 2980.27 | 2083 | 2287 | 1.333 | 0.415 | 0.00721482 |
| AT3G27890.1 | NQR | protein_coding | NADPH:quinone oxidored | 510 | 287 | 312 | 529 | 375 | 294 | 1072 | 705 | 799 | 1147 | 996 | 800 | 1074.55 | 859 | 981 | 1.333 | 0.415 | 0.00656938 |
| AT2G38170.3 | CAX1 | protein_coding | cation exchanger 1 | 4530 | 2855 | 2635 | 4443 | 2802 | 3794 | 9522 | 7014 | 6746 | 9637 | 7439 | 10318 | 7516.48 | 7761 | 9131 | 1.334 | 0.415 | 0.0060456 |
| AT3G53020.1 | RPL24B | protein_coding | Ribosomal protein L24e f | 3179 | 2163 | 1748 | 3290 | 2439 | 2224 | 6682 | 5314 | 4475 | 7136 | 6475 | 6048 | 7418.18 | 5490 | 6553 | 1.333 | 0.415 | 0.00502889 |
| AT4G39970.1 | | protein_coding | Haloacid dehalogenase-li | 1817 | 1816 | 1647 | 2340 | 2329 | 2226 | 3819 | 4462 | 4216 | 5076 | 6183 | 6053 | 5773.49 | 4166 | 5771 | 1.333 | 0.415 | 0.00062537 |
| AT5G48220.1 | | protein_coding | Aldolase-type TIM barrel | 981 | 756 | 680 | 894 | 839 | 749 | 2062 | 1857 | 1741 | 1939 | 2227 | 2037 | 2418.11 | 1887 | 2068 | 1.333 | 0.415 | 0.00035536 |
| AT4G03070.1 | AOP1 | protein_coding | 2-oxoglutarate (2OG) and | 250 | 217 | 179 | 316 | 295 | 161 | 526 | 533 | 458 | 685 | 783 | 438 | 546.39 | 506 | 635 | 1.333 | 0.414 | 0.00987296 |
| AT5G02960.1 | RPS23B | protein_coding | Ribosomal protein S12/S2 | 7338 | 4851 | 3818 | 6584 | 4802 | 4232 | 15425 | 11918 | 9774 | 14281 | 12748 | 11509 | 13114.26 | 12372 | 12846 | 1.333 | 0.414 | 0.00938874 |
| AT4G18440.1 | | protein_coding | L-Aspartase-like family pr | 1211 | 991 | 674 | 1240 | 1212 | 826 | 2546 | 2435 | 1725 | 2690 | 3218 | 2246 | 2703.61 | 2235 | 2718 | 1.332 | 0.414 | 0.00927797 |
| AT1G21770.1 | | protein_coding | Acyl-CoA N-acyltransfera | 208 | 262 | 227 | 465 | 302 | 467 | 437 | 644 | 581 | 1009 | 802 | 1270 | 919.03 | 554 | 1027 | 1.333 | 0.414 | 0.00866233 |
| AT1G18540.1 | RPL6A | protein_coding | Ribosomal protein L6 fan | 6100 | 4268 | 3279 | 5261 | 3733 | 3804 | 12823 | 10486 | 8394 | 11412 | 9910 | 10345 | 11213.16 | 10568 | 10556 | 1.333 | 0.414 | 0.0070855 |
| AT5G18570.1 | OBGL | protein_coding | GTP1/OBG family protein | 507 | 414 | 312 | 517 | 388 | 339 | 1066 | 1017 | 799 | 1121 | 1030 | 922 | 2013.43 | 961 | 1024 | 1.332 | 0.414 | 0.00139962 |
| AT3G11330.1 | PIRL9 | protein_coding | plant intracellular ras gro | 287 | 271 | 251 | 403 | 281 | 258 | 603 | 666 | 643 | 874 | 746 | 702 | 695.97 | 637 | 774 | 1.332 | 0.414 | 0.00027965 |
| AT1G29357.1 | | other_rna | other RNA | 82 | 70 | 105 | 116 | 104 | 104 | 172 | 172 | 269 | 252 | 276 | 283 | 320.84 | 204 | 270 | 1.331 | 0.413 | 0.00405161 |
| AT5G03830.1 | | protein_coding | CDK inhibitor P21 binding | 136 | 109 | 138 | 137 | 112 | 135 | 286 | 268 | 353 | 297 | 297 | 367 | 331.17 | 302 | 320 | 1.331 | 0.413 | 0.00318526 |
| AT5G59730.1 | ATEXO70H7 | protein_coding | exocyst subunit exo70 fa | 637 | 609 | 491 | 807 | 638 | 688 | 1339 | 1496 | 1257 | 1750 | 1694 | 1871 | 1212.29 | 1364 | 1772 | 1.332 | 0.413 | 0.00150537 |
| AT3G52230.1 | | protein_coding | | 996 | 937 | 965 | 1409 | 1258 | 1432 | 2094 | 2302 | 2470 | 3056 | 3340 | 3894 | 3192.74 | 2289 | 3430 | 1.332 | 0.413 | 0.00069352 |
| AT1G09620.1 | | protein_coding | ATP binding;leucine-tRNA | 4526 | 3344 | 2995 | 4646 | 3737 | 3590 | 9514 | 8216 | 7667 | 10078 | 9921 | 9763 | 7192.31 | 8466 | 9921 | 1.331 | 0.413 | 0.00045641 |
| AT4G37000.1 | RCCR | protein_coding | accelerated cell death 2 ( | 773 | 699 | 671 | 889 | 774 | 647 | 1625 | 1717 | 1718 | 1928 | 2055 | 1759 | 2125.3 | 1687 | 1914 | 1.332 | 0.413 | 0.00024671 |
| AT1G49980.1 | | protein_coding | DNA/RNA polymerases su | 257 | 209 | 199 | 243 | 169 | 194 | 540 | 513 | 509 | 527 | 449 | 528 | 547.29 | 521 | 501 | 1.331 | 0.412 | 0.00151068 |
| AT2G21270.3 | UFD1 | protein_coding | ubiquitin fusion degradat | 698 | 523 | 509 | 882 | 721 | 642 | 1467 | 1285 | 1303 | 1913 | 1914 | 1746 | 1591.43 | 1352 | 1858 | 1.33 | 0.412 | 0.00033367 |
| AT5G09620.1 | | protein_coding | Octicosapeptide/Phox/Be | 657 | 672 | 665 | 816 | 632 | 893 | 1381 | 1651 | 1702 | 1770 | 1678 | 2428 | 1594.64 | 1578 | 1959 | 1.33 | 0.411 | 0.00429269 |
| AT1G79930.1 | HSP70-14 | protein_coding | heat shock protein 91 | 1664 | 1357 | 1012 | 1925 | 1486 | 1409 | 3498 | 3334 | 2591 | 4176 | 3945 | 3832 | 3924.79 | 3141 | 3984 | 1.33 | 0.411 | 0.00152785 |
| AT5G51300.1 | | protein_coding | splicing factor-related | 699 | 666 | 618 | 902 | 768 | 913 | 1469 | 1636 | 1582 | 1957 | 2039 | 2483 | 2003.43 | 1562 | 2160 | 1.33 | 0.411 | 0.00090203 |
| AT3G01160.1 | | protein_coding | | 594 | 389 | 335 | 544 | 421 | 399 | 1249 | 956 | 858 | 1180 | 1118 | 1085 | 927.74 | 1021 | 1128 | 1.329 | 0.41 | 0.00362278 |
| AT4G10620.1 | | protein_coding | P-loop containing nucleo | 271 | 158 | 164 | 223 | 184 | 159 | 570 | 388 | 420 | 484 | 488 | 432 | 458.04 | 459 | 468 | 1.329 | 0.41 | 0.00284277 |
| AT4G38160.3 | pde191 | protein_coding | Mitochondrial transcripti | 1143 | 919 | 793 | 1086 | 927 | 1005 | 2403 | 2258 | 2030 | 2356 | 2461 | 2733 | 2679.46 | 2230 | 2517 | 1.329 | 0.41 | 0.00082235 |
| AT3G15354.1 | SPA3 | protein_coding | SPA1-related 3 | 984 | 654 | 575 | 1191 | 790 | 671 | 2068 | 1607 | 1472 | 2583 | 2097 | 1825 | 1583.6 | 1716 | 2168 | 1.328 | 0.409 | 0.00868048 |
| AT2G47790.1 | | protein_coding | Transducin/WD40 repeat | 271 | 216 | 137 | 314 | 223 | 230 | 570 | 531 | 351 | 681 | 592 | 625 | 654.93 | 484 | 633 | 1.328 | 0.409 | 0.00389477 |
| AT2G37860.3 | LCD1 | protein_coding | Protein of unknown funct | 670 | 486 | 393 | 611 | 586 | 428 | 1408 | 1194 | 1006 | 1325 | 1556 | 1164 | 1555.85 | 1203 | 1348 | 1.327 | 0.409 | 0.00373996 |
| AT5G48240.3 | | protein_coding | | 529 | 364 | 314 | 424 | 346 | 299 | 1112 | 894 | 804 | 920 | 919 | 813 | 835.11 | 937 | 884 | 1.328 | 0.409 | 0.00175315 |
| AT3G51920.1 | CML9 | protein_coding | calmodulin 9 | 703 | 502 | 502 | 855 | 610 | 539 | 1478 | 1233 | 1285 | 1855 | 1619 | 1466 | 1553.87 | 1332 | 1647 | 1.327 | 0.409 | 0.00090627 |
| AT2G42680.1 | MBF1A | protein_coding | multiprotein bridging fac | 679 | 621 | 639 | 919 | 756 | 826 | 1427 | 1526 | 1636 | 1993 | 2007 | 2246 | 2227.96 | 1530 | 2082 | 1.328 | 0.409 | 0.00059971 |
| AT3G05530.1 | RPT5A | protein_coding | regulatory particle triple- | 1305 | 1113 | 1070 | 1573 | 1283 | 1353 | 2743 | 2734 | 2739 | 3412 | 3406 | 3679 | 3653.88 | 2739 | 3499 | 1.328 | 0.409 | 0.00015244 |
| AT2G32230.1 | PRORP1 | protein_coding | proteinaceous RNase P 1 | 1281 | 750 | 707 | 1113 | 822 | 618 | 2693 | 1843 | 1810 | 2414 | 2182 | 1681 | 2939.95 | 2115 | 2092 | 1.327 | 0.408 | 0.00964416 |
| AT1G22840.1 | CYTC-1 | protein_coding | CYTOCHROME C-1 | 1064 | 820 | 582 | 1400 | 942 | 1110 | 2237 | 2015 | 1490 | 3037 | 2501 | 3019 | 3061.03 | 1914 | 2852 | 1.327 | 0.408 | 0.00483606 |
| AT3G55620.1 | EIF6-2 | protein_coding | Translation initiation fact | 798 | 706 | 511 | 926 | 745 | 702 | 1677 | 1735 | 1308 | 2009 | 1978 | 1909 | 2259.46 | 1573 | 1965 | 1.327 | 0.408 | 0.00120531 |
| AT3G59400.1 | GUN4 | protein_coding | enzyme binding;tetrapyrr | 1484 | 1780 | 1136 | 1810 | 1531 | 1680 | 3119 | 4373 | 2908 | 3926 | 4064 | 4569 | 6971.57 | 3467 | 4186 | 1.326 | 0.407 | 0.00587923 |
| AT5G17600.1 | ATL52 | protein_coding | RING/U-box superfamily | 106 | 96 | 87 | 126 | 78 | 83 | 223 | 236 | 223 | 273 | 207 | 226 | 235.48 | 227 | 235 | 1.325 | 0.406 | 0.00620542 |
| AT4G19510.1 | | protein_coding | Disease resistance protei | 896 | 677 | 711 | 910 | 744 | 663 | 1883 | 1663 | 1820 | 1974 | 1975 | 1803 | 1411.96 | 1789 | 1917 | 1.325 | 0.406 | 0.00219056 |
| AT4G28740.1 | | protein_coding | | 530 | 410 | 368 | 641 | 483 | 406 | 1114 | 1007 | 942 | 1390 | 1282 | 1104 | 1212.09 | 1021 | 1259 | 1.325 | 0.406 | 0.00152168 |
| AT1G71720.1 | | protein_coding | Nucleic acid-binding prot | 516 | 480 | 406 | 523 | 408 | 373 | 1085 | 1179 | 1039 | 1134 | 1083 | 1014 | 1591.05 | 1101 | 1077 | 1.325 | 0.406 | 0.0005375 |
| AT1G55890.1 | | protein_coding | Tetratricopeptide repeat | 420 | 338 | 304 | 440 | 358 | 303 | 883 | 830 | 778 | 954 | 950 | 824 | 857.24 | 830 | 909 | 1.325 | 0.406 | 0.00022078 |
| AT3G05590.1 | RPL18B | protein_coding | ribosomal protein L18 | 5821 | 4326 | 3154 | 5818 | 4532 | 4305 | 12236 | 10628 | 8074 | 12620 | 12031 | 11707 | 12887.43 | 10313 | 12119 | 1.324 | 0.405 | 0.00912287 |
| AT5G51740.1 | | protein_coding | Peptidase family M48 far | 130 | 83 | 119 | 182 | 140 | 186 | 273 | 204 | 305 | 395 | 372 | 506 | 321.83 | 261 | 424 | 1.324 | 0.405 | 0.00700824 |
| AT1G02840.1 | SR34 | protein_coding | RNA-binding (RRM/RBD/ | 618 | 686 | 606 | 826 | 704 | 940 | 1299 | 1685 | 1551 | 1792 | 1869 | 2556 | 2212.91 | 1512 | 2072 | 1.324 | 0.405 | 0.00494354 |
| AT2G25720.1 | | protein_coding | | 79 | 78 | 76 | 115 | 89 | 103 | 166 | 192 | 195 | 249 | 236 | 280 | 254.2 | 184 | 255 | 1.324 | 0.405 | 0.00299066 |
| AT2G33620.1 | AHL10 | protein_coding | AT hook motif DNA-bindi | 199 | 193 | 175 | 235 | 250 | 219 | 418 | 474 | 448 | 510 | 664 | 596 | 624.62 | 447 | 590 | 1.324 | 0.405 | 0.00240934 |
| AT2G35390.2 | PRS1 | protein_coding | Phosphoribosyltransferas | 930 | 606 | 563 | 1281 | 758 | 698 | 1955 | 1489 | 1441 | 2779 | 2012 | 1898 | 2163.71 | 1628 | 2230 | 1.323 | 0.404 | 0.00806787 |
| AT5G06980.4 | | protein_coding | | 910 | 489 | 644 | 1012 | 687 | 830 | 1913 | 1201 | 1649 | 2195 | 1824 | 2257 | 1541.04 | 1588 | 2092 | 1.323 | 0.404 | 0.00730715 |
| AT1G26230.1 | CPN60B4 | protein_coding | TCP-1/cpn60 chaperonin | 721 | 578 | 443 | 735 | 696 | 466 | 1516 | 1420 | 1134 | 1594 | 1848 | 1267 | 1544.48 | 1357 | 1570 | 1.323 | 0.404 | 0.00680085 |
| AT1G80630.1 | | protein_coding | RNI-like superfamily prot | 232 | 187 | 206 | 244 | 194 | 175 | 488 | 459 | 527 | 529 | 515 | 476 | 431.46 | 491 | 507 | 1.323 | 0.404 | 0.00186929 |
| AT5G45190.2 | CYCT1-5 | protein_coding | Cyclin family protein | 447 | 415 | 495 | 526 | 430 | 488 | 940 | 1020 | 1267 | 1141 | 1142 | 1327 | 1126.68 | 1076 | 1203 | 1.323 | 0.404 | 0.00146573 |
| AT5G53120.6 | SPMS | protein_coding | spermidine synthase 3 | 182 | 183 | 148 | 244 | 220 | 171 | 383 | 450 | 379 | 529 | 584 | 465 | 468.12 | 404 | 526 | 1.323 | 0.404 | 0.0013302 |
| AT4G37440.1 | | protein_coding | | 769 | 578 | 644 | 850 | 609 | 684 | 1616 | 1420 | 1649 | 1844 | 1617 | 1860 | 1430.81 | 1562 | 1774 | 1.323 | 0.404 | 0.0011729 |
| AT1G25260.1 | | protein_coding | Ribosomal protein L10 fa | 1009 | 680 | 621 | 1207 | 909 | 709 | 2121 | 1671 | 1590 | 2618 | 2413 | 1928 | 1965.63 | 1794 | 2320 | 1.322 | 0.403 | 0.00530773 |
| AT4G11240.1 | TOPP6 | protein_coding | Calcineurin-like metallo-p | 320 | 218 | 273 | 365 | 282 | 393 | 673 | 536 | 699 | 792 | 749 | 1069 | 817.24 | 636 | 870 | 1.322 | 0.403 | 0.003289 |
| AT3G50880.1 | | protein_coding | DNA glycosylase superfar | 174 | 150 | 200 | 279 | 219 | 251 | 366 | 369 | 512 | 605 | 581 | 683 | 538.85 | 416 | 623 | 1.323 | 0.403 | 0.00125347 |
| AT3G63520.1 | CCD1 | protein_coding | carotenoid cleavage diox | 3996 | 3397 | 3434 | 4882 | 3776 | 3839 | 8400 | 8346 | 8791 | 10590 | 10024 | 10440 | 10611.53 | 8512 | 10351 | 1.322 | 0.403 | 0.00047947 |
| AT1G16250.1 | | protein_coding | Galactose oxidase/kelch r | 155 | 158 | 159 | 160 | 142 | 153 | 326 | 388 | 407 | 347 | 377 | 416 | 321.73 | 374 | 380 | 1.322 | 0.402 | 0.00301755 |
| AT5G10070.2 | | protein_coding | RNase L inhibitor protein- | 223 | 194 | 177 | 283 | 184 | 188 | 469 | 477 | 453 | 614 | 488 | 511 | 528.9 | 466 | 538 | 1.321 | 0.402 | 0.00129332 |
| AT4G29510.1 | PRMT11 | protein_coding | arginine methyltransferas | 1005 | 739 | 573 | 893 | 774 | 669 | 2113 | 1816 | 1467 | 1937 | 2055 | 1819 | 1868.3 | 1799 | 1937 | 1.321 | 0.401 | 0.00213333 |
| AT2G25870.1 | | protein_coding | haloacid dehalogenase-lil | 1517 | 1163 | 900 | 1525 | 1243 | 1095 | 3189 | 2857 | 2304 | 3308 | 3300 | 2978 | 3372.87 | 2783 | 3195 | 1.32 | 0.401 | 0.00166504 |
| AT5G65630.1 | GTE7 | protein_coding | global transcription facto | 1256 | 963 | 961 | 1145 | 915 | 990 | 2640 | 2366 | 2460 | 2484 | 2429 | 2692 | 2598.06 | 2489 | 2535 | 1.32 | 0.401 | 0.00064855 |
| AT1G32100.1 | PRR1 | protein_coding | pinoresinol reductase 1 | 180 | 176 | 185 | 192 | 165 | 218 | 378 | 432 | 474 | 416 | 438 | 593 | 455.85 | 428 | 482 | 1.32 | 0.4 | 0.0050089 |
| AT5G02770.1 | MOS11 | protein_coding | | 1347 | 1177 | 1263 | 1646 | 1645 | 1507 | 2831 | 2892 | 3233 | 3570 | 4367 | 4098 | 3863.85 | 2985 | 4012 | 1.32 | 0.4 | 0.00101486 |
| AT5G12130.1 | PDE149 | protein_coding | integral membrane TerC | 921 | 683 | 644 | 937 | 838 | 696 | 1936 | 1678 | 1649 | 2032 | 2225 | 1893 | 2136.05 | 1754 | 2050 | 1.32 | 0.4 | 0.0009932 |
| AT3G13290.1 | VCR | protein_coding | varicose-related | 763 | 676 | 636 | 917 | 800 | 724 | 1604 | 1661 | 1628 | 1989 | 2124 | 1969 | 1456.68 | 1631 | 2027 | 1.32 | 0.4 | 0.00058482 |
| AT3G48190.1 | ATM | protein_coding | ataxia-telangiectasia mut | 1230 | 1008 | 1008 | 1354 | 1256 | 1112 | 2586 | 2476 | 2581 | 2937 | 3334 | 3024 | 1885.38 | 2548 | 3098 | 1.319 | 0.4 | 0.00044301 |
| AT4G30100.1 | | protein_coding | P-loop containing nucleo | 1340 | 1255 | 1194 | 1538 | 1193 | 1205 | 2817 | 3083 | 3057 | 3336 | 3167 | 3277 | 2709.03 | 2986 | 3260 | 1.32 | 0.4 | 0.00038342 |
| AT5G64680.3 | | protein_coding | | 190 | 120 | 127 | 205 | 146 | 157 | 399 | 295 | 325 | 445 | 388 | 427 | 404.65 | 340 | 420 | 1.319 | 0.399 | 0.00096147 |
| AT5G09330.1 | VNI1 | protein_coding | NAC domain containing p | 426 | 366 | 321 | 550 | 374 | 434 | 895 | 899 | 822 | 1193 | 993 | 1180 | 1390.09 | 872 | 1122 | 1.319 | 0.399 | 0.00094139 |
| AT3G52120.1 | | protein_coding | SWAP (Suppressor-of-Wh | 611 | 540 | 472 | 796 | 633 | 551 | 1284 | 1327 | 1208 | 1727 | 1680 | 1498 | 1392.07 | 1273 | 1635 | 1.319 | 0.399 | 0.00084831 |
| AT5G10170.1 | IPS3 | protein_coding | myo-inositol-1-phosphate | 410 | 337 | 325 | 417 | 346 | 329 | 862 | 828 | 832 | 905 | 919 | 895 | 1105.22 | 841 | 906 | 1.319 | 0.399 | 0.00025251 |
| AT1G02500.1 | SAM1 | protein_coding | S-adenosylmethionine sy | 2864 | 3212 | 2672 | 3800 | 3095 | 4379 | 6020 | 7891 | 6840 | 8243 | 8217 | 11908 | 8155.56 | 6917 | 9456 | 1.318 | 0.398 | 0.00904057 |
| AT2G43460.1 | RPL38B | protein_coding | Ribosomal L38e protein f | 2523 | 1910 | 1575 | 2818 | 2014 | 1862 | 5304 | 4693 | 4032 | 6113 | 5347 | 5064 | 5518.88 | 4676 | 5508 | 1.318 | 0.398 | 0.00217528 |
| AT2G27450.2 | CPA | protein_coding | nitrilase-like protein 1 | 696 | 491 | 467 | 832 | 611 | 568 | 1463 | 1206 | 1196 | 1805 | 1622 | 1545 | 1565.16 | 1288 | 1657 | 1.318 | 0.398 | 0.00194569 |
| AT5G03760.1 | CSLA9 | protein_coding | Nucleotide-diphospho-su | 2066 | 1606 | 1495 | 2473 | 2242 | 1816 | 4343 | 3946 | 3827 | 5364 | 5952 | 4939 | 4238.14 | 4039 | 5418 | 1.318 | 0.398 | 0.00106681 |
| AT2G19270.1 | | protein_coding | | 324 | 290 | 280 | 403 | 310 | 321 | 681 | 712 | 717 | 874 | 823 | 873 | 834.63 | 703 | 857 | 1.317 | 0.398 | 0.00015064 |
| AT5G02270.2 | ABCI20 | protein_coding | non-intrinsic ABC protein | 178 | 125 | 93 | 218 | 126 | 120 | 374 | 307 | 238 | 473 | 335 | 326 | 470.08 | 306 | 378 | 1.316 | 0.397 | 0.00867017 |
| AT5G02590.1 | | protein_coding | Tetratricopeptide repeat | 152 | 110 | 89 | 138 | 103 | 106 | 320 | 270 | 228 | 299 | 273 | 288 | 398.14 | 273 | 287 | 1.317 | 0.397 | 0.00329122 |
| AT3G14750.1 | FLXL1 | protein_coding | | 138 | 119 | 121 | 143 | 135 | 117 | 290 | 292 | 310 | 310 | 358 | 318 | 255.56 | 297 | 329 | 1.317 | 0.397 | 0.00297574 |
| AT3G06720.1 | IMPA1 | protein_coding | importin alpha isoform 1 | 1387 | 973 | 845 | 1709 | 1124 | 1127 | 2916 | 2390 | 2163 | 3707 | 2984 | 3065 | 2718.47 | 2490 | 3252 | 1.317 | 0.397 | 0.00288156 |
| AT1G15910.1 | FDM1 | protein_coding | XH/XS domain-containing | 405 | 381 | 371 | 422 | 416 | 438 | 851 | 936 | 950 | 915 | 1104 | 1191 | 824.28 | 912 | 1070 | 1.317 | 0.397 | 0.00140642 |
| AT2G04520.1 | | protein_coding | Nucleic acid-binding, OB- | 1597 | 1206 | 1258 | 1735 | 1373 | 1273 | 3357 | 2963 | 3221 | 3763 | 3645 | 3462 | 3307.59 | 3180 | 3623 | 1.317 | 0.397 | 0.00081724 |
| AT4G28730.1 | GRXC5 | protein_coding | Glutaredoxin family prote | 436 | 320 | 340 | 428 | 357 | 297 | 917 | 786 | 870 | 928 | 948 | 808 | 928.47 | 858 | 895 | 1.316 | 0.397 | 0.00050398 |
| AT5G16050.1 | GRF5 | protein_coding | general regulatory factor | 1132 | 1046 | 989 | 1506 | 1258 | 1165 | 2380 | 2570 | 2532 | 3267 | 3340 | 3168 | 2743.38 | 2494 | 3258 | 1.316 | 0.397 | 0.00043968 |
| AT5G51940.1 | NRPB6A | protein_coding | RNA polymerase Rpb6 | 300 | 249 | 239 | 379 | 284 | 317 | 631 | 612 | 612 | 822 | 754 | 862 | 751.05 | 618 | 813 | 1.317 | 0.397 | 0.00025132 |
| AT1G52300.1 | RPL37B | protein_coding | Zinc-binding ribosomal pr | 3912 | 3354 | 2215 | 3973 | 2755 | 3126 | 8223 | 8240 | 5671 | 8618 | 7314 | 8501 | 8475.03 | 7378 | 8144 | 1.316 | 0.396 | 0.00994698 |
| AT1G31280.1 | AGO2 | protein_coding | Argonaute family protein | 125 | 161 | 126 | 173 | 168 | 143 | 263 | 396 | 323 | 375 | 446 | 389 | 370.58 | 327 | 403 | 1.316 | 0.396 | 0.00575982 |
| AT3G02065.2 | RH41 | protein_coding | P-loop containing nucleo | 220 | 182 | 190 | 243 | 185 | 173 | 462 | 447 | 486 | 527 | 491 | 470 | 512.57 | 465 | 496 | 1.315 | 0.396 | 0.00028808 |
| AT1G11475.1 | NRPB10 | protein_coding | RNA polymerases N / 8 kI | 350 | 231 | 223 | 443 | 345 | 265 | 736 | 568 | 571 | 961 | 916 | 721 | 770.85 | 625 | 866 | 1.315 | 0.395 | 0.00751606 |
| AT5G63890.2 | HISN8 | protein_coding | histidinol dehydrogenase | 1404 | 941 | 890 | 1625 | 1072 | 1062 | 2951 | 2312 | 2278 | 3525 | 2846 | 2888 | 2903.43 | 2514 | 3086 | 1.315 | 0.395 | 0.00323018 |
| AT5G45410.1 | | protein_coding | | 548 | 497 | 380 | 840 | 605 | 500 | 1152 | 1221 | 973 | 1822 | 1606 | 1360 | 1584.48 | 1115 | 1596 | 1.315 | 0.395 | 0.00225479 |
| AT4G22310.1 | MPC4 | protein_coding | Uncharacterised protein f | 1027 | 794 | 869 | 1653 | 1087 | 1068 | 2159 | 1951 | 2225 | 3586 | 2886 | 2904 | 2819.56 | 2112 | 3125 | 1.315 | 0.395 | 0.0012455 |
| AT4G39460.1 | SAMC1 | protein_coding | S-adenosylmethionine ca | 1002 | 872 | 854 | 1150 | 961 | 947 | 2106 | 2142 | 2186 | 2494 | 2551 | 2575 | 2681.12 | 2145 | 2540 | 1.314 | 0.395 | 0.00047867 |
| AT5G58090.1 | | protein_coding | O-Glycosyl hydrolases far | 466 | 538 | 418 | 672 | 596 | 805 | 980 | 1322 | 1070 | 1458 | 1582 | 2189 | 1330.54 | 1124 | 1743 | 1.314 | 0.394 | 0.00989665 |
| AT4G26430.1 | CSN6B | protein_coding | COP9 signalosome subun | 171 | 126 | 108 | 191 | 148 | 101 | 359 | 310 | 276 | 414 | 393 | 275 | 366.83 | 315 | 361 | 1.314 | 0.394 | 0.00729379 |
| AT2G02880.1 | | protein_coding | mucin-related | 138 | 120 | 94 | 132 | 96 | 74 | 290 | 295 | 241 | 286 | 255 | 201 | 247.88 | 275 | 247 | 1.314 | 0.394 | 0.00612592 |
| AT5G04600.1 | | protein_coding | RNA-binding (RRM/RBD/ | 855 | 508 | 573 | 883 | 699 | 608 | 1797 | 1248 | 1467 | 1915 | 1856 | 1653 | 1805.47 | 1504 | 1808 | 1.314 | 0.394 | 0.00489553 |
| AT5G06360.1 | | protein_coding | Ribosomal protein S8e fa | 1166 | 818 | 751 | 1208 | 890 | 822 | 2451 | 2010 | 1923 | 2620 | 2363 | 2235 | 2403.35 | 2128 | 2406 | 1.314 | 0.394 | 0.00215717 |
| AT1G62390.1 | Phox2 | protein_coding | Octicosapeptide/Phox/Be | 3382 | 2228 | 2165 | 3180 | 2459 | 2230 | 7109 | 5474 | 5543 | 6898 | 6528 | 6064 | 5406.39 | 6042 | 6497 | 1.314 | 0.394 | 0.00187503 |
| AT2G19180.1 | | protein_coding | | 127 | 120 | 146 | 169 | 149 | 172 | 267 | 295 | 374 | 367 | 396 | 468 | 480.82 | 312 | 410 | 1.314 | 0.394 | 0.00124666 |

| | | | | | | | | | | | | | | | | | | | | |
|---|---|---|---|---|---|---|---|---|---|---|---|---|---|---|---|---|---|---|---|---|
| AT3G19800.1 | | protein_coding | Protein of unknown funct | 477 | 460 | 400 | 660 | 556 | 461 | 1003 | 1130 | 1024 | 1432 | 1476 | 1254 | 1734.43 | 1052 | 1387 | 1.314 | 0.394 | 0.00099182 |
| AT1G20960.1 | emb1507 | protein_coding | U5 small nuclear ribonucl | 3916 | 3183 | 2872 | 4120 | 3548 | 3316 | 8232 | 7820 | 7353 | 8937 | 9419 | 9018 | 6573.94 | 7802 | 9125 | 1.314 | 0.394 | 0.00066759 |
| AT3G12670.1 | emb2742 | protein_coding | CTP synthase family prote | 1246 | 949 | 918 | 1277 | 1141 | 1088 | 2619 | 2332 | 2350 | 2770 | 3029 | 2959 | 2497.22 | 2434 | 2919 | 1.314 | 0.394 | 0.00061191 |
| AT4G30950.1 | FAD6 | protein_coding | fatty acid desaturase 6 | 8218 | 8036 | 6978 | 9642 | 8706 | 8736 | 17275 | 19743 | 17864 | 20914 | 23112 | 23757 | 22623.13 | 18294 | 22594 | 1.313 | 0.393 | 0.00336246 |
| AT3G02710.1 | | protein_coding | ARM repeat superfamily | 396 | 301 | 280 | 417 | 302 | 361 | 832 | 740 | 717 | 905 | 802 | 982 | 870.92 | 763 | 896 | 1.313 | 0.393 | 0.00094178 |
| AT5G39530.1 | | protein_coding | Protein of unknown funct | 2131 | 2194 | 1938 | 2852 | 2554 | 2277 | 4480 | 5390 | 4961 | 6186 | 6780 | 6192 | 5298.45 | 4944 | 6386 | 1.313 | 0.393 | 0.00082781 |
| AT5G58900.4 | | protein_coding | F-box/RNI-like superfamil | 168 | 111 | 120 | 172 | 121 | 96 | 353 | 273 | 307 | 373 | 321 | 261 | 303.58 | 311 | 318 | 1.312 | 0.392 | 0.0080969 |
| AT4G32720.1 | LA1 | protein_coding | La protein 1 | 937 | 646 | 564 | 807 | 675 | 557 | 1970 | 1587 | 1444 | 1750 | 1792 | 1515 | 1812.21 | 1667 | 1686 | 1.313 | 0.392 | 0.00386057 |
| AT1G71960.1 | ABCG25 | protein_coding | ATP-binding casette fami | 147 | 118 | 88 | 201 | 149 | 130 | 309 | 290 | 225 | 436 | 396 | 354 | 332.12 | 275 | 395 | 1.312 | 0.392 | 0.00203666 |
| AT2G02910.1 | | protein_coding | Protein of unknown funct | 178 | 142 | 164 | 201 | 159 | 133 | 374 | 349 | 420 | 436 | 422 | 362 | 409.13 | 381 | 407 | 1.312 | 0.392 | 0.00126304 |
| AT5G20170.1 | MED17 | protein_coding | RNA polymerase II transc | 304 | 254 | 210 | 304 | 242 | 233 | 639 | 624 | 538 | 659 | 642 | 634 | 569.7 | 600 | 645 | 1.312 | 0.392 | 0.00067046 |
| AT4G20010.1 | OSB2 | protein_coding | plastid transcriptionally a | 514 | 370 | 319 | 563 | 404 | 348 | 1080 | 909 | 817 | 1221 | 1073 | 946 | 1073.04 | 935 | 1080 | 1.311 | 0.391 | 0.0025991 |
| AT5G27670.1 | HTA7 | protein_coding | histone H2A 7 | 699 | 650 | 573 | 772 | 734 | 720 | 1469 | 1597 | 1467 | 1675 | 1949 | 1958 | 1797.6 | 1511 | 1861 | 1.311 | 0.391 | 0.00077107 |
| AT4G29120.1 | | protein_coding | 6-phosphogluconate deh | 314 | 270 | 232 | 384 | 287 | 271 | 660 | 663 | 594 | 833 | 762 | 737 | 955.16 | 639 | 777 | 1.312 | 0.391 | 0.00073012 |
| AT5G05780.1 | RPN8A | protein_coding | RP non-ATPase subunit 8 | 1454 | 1128 | 1094 | 1740 | 1282 | 1442 | 3056 | 2771 | 2801 | 3774 | 3403 | 3921 | 3075.01 | 2876 | 3699 | 1.311 | 0.391 | 0.00060735 |
| AT2G39760.1 | BPM3 | protein_coding | BTB/POZ/MATH-domains | 468 | 391 | 389 | 593 | 482 | 431 | 984 | 961 | 996 | 1286 | 1280 | 1172 | 1009.59 | 980 | 1246 | 1.311 | 0.391 | 0.00042851 |
| AT5G01780.2 | | protein_coding | 2-oxoglutarate-depender | 206 | 189 | 158 | 186 | 163 | 144 | 433 | 464 | 404 | 403 | 433 | 392 | 442.33 | 434 | 409 | 1.311 | 0.391 | 0.00026405 |
| AT4G00200.1 | | protein_coding | splicing factor PWI doma | 778 | 703 | 776 | 837 | 842 | 965 | 1635 | 1727 | 1987 | 1816 | 2235 | 2624 | 2683.16 | 1783 | 2225 | 1.31 | 0.39 | 0.00498412 |
| AT1G73230.1 | | protein_coding | Nascent polypeptide-assc | 2878 | 2027 | 1789 | 3183 | 2427 | 2194 | 6050 | 4980 | 4580 | 6904 | 6443 | 5966 | 6341.08 | 5203 | 6438 | 1.311 | 0.39 | 0.00197785 |
| AT3G15180.2 | | protein_coding | ARM repeat superfamily | 294 | 193 | 194 | 375 | 265 | 206 | 618 | 474 | 497 | 813 | 704 | 560 | 574.56 | 530 | 692 | 1.309 | 0.389 | 0.00544088 |
| AT3G24570.1 | | protein_coding | Peroxisomal membrane 2 | 273 | 283 | 224 | 378 | 286 | 347 | 574 | 695 | 573 | 820 | 759 | 944 | 895.21 | 614 | 841 | 1.309 | 0.389 | 0.00153641 |
| AT1G72040.1 | | protein_coding | P-loop containing nucleo | 874 | 624 | 570 | 889 | 797 | 719 | 1837 | 1533 | 1459 | 1928 | 2116 | 1955 | 1891.08 | 1610 | 2000 | 1.309 | 0.389 | 0.00139754 |
| AT4G23840.1 | | protein_coding | Leucine-rich repeat (LRR) | 129 | 104 | 112 | 150 | 127 | 123 | 271 | 256 | 287 | 325 | 337 | 334 | 276.4 | 271 | 332 | 1.309 | 0.389 | 0.00106179 |
| AT4G32190.1 | | protein_coding | Myosin heavy chain-relat | 832 | 726 | 688 | 850 | 721 | 633 | 1749 | 1784 | 1761 | 1844 | 1914 | 1721 | 1668.54 | 1765 | 1826 | 1.309 | 0.389 | 0.00051008 |
| AT3G52090.2 | NRPB11 | protein_coding | DNA-directed RNA polym | 480 | 396 | 405 | 753 | 585 | 597 | 1009 | 973 | 1037 | 1633 | 1553 | 1624 | 1313.18 | 1006 | 1603 | 1.309 | 0.389 | 0.00050778 |
| AT1G70470.1 | | protein_coding | | 60 | 69 | 64 | 106 | 75 | 74 | 126 | 170 | 164 | 230 | 199 | 201 | 183.24 | 153 | 210 | 1.308 | 0.388 | 0.00681383 |
| AT1G07470.1 | | protein_coding | Transcription factor IIA, a | 284 | 270 | 223 | 353 | 342 | 327 | 597 | 663 | 571 | 766 | 908 | 889 | 787.7 | 610 | 854 | 1.309 | 0.388 | 0.00120663 |
| AT4G17020.3 | | protein_coding | transcription factor-relat | 303 | 225 | 292 | 377 | 317 | 281 | 637 | 553 | 748 | 818 | 842 | 764 | 589.21 | 646 | 808 | 1.309 | 0.388 | 0.00119016 |
| AT2G03820.1 | | protein_coding | nonsense-mediated mRN | 1369 | 1070 | 951 | 1615 | 1192 | 1200 | 2878 | 2629 | 2435 | 3503 | 3164 | 3263 | 2809.89 | 2647 | 3310 | 1.308 | 0.388 | 0.00088485 |
| AT5G26760.2 | | protein_coding | | 559 | 452 | 505 | 597 | 576 | 518 | 1175 | 1110 | 1293 | 1295 | 1529 | 1409 | 1098.27 | 1193 | 1411 | 1.309 | 0.388 | 0.0007486 |
| AT5G10400.1 | HTR2 | protein_coding | Histone superfamily prot | 677 | 701 | 465 | 862 | 739 | 775 | 1423 | 1722 | 1190 | 1870 | 1962 | 2108 | 1856.21 | 1445 | 1980 | 1.308 | 0.387 | 0.00314643 |
| AT4G03430.1 | STA1 | protein_coding | pre-mRNA splicing factor | 794 | 769 | 709 | 986 | 763 | 858 | 1669 | 1889 | 1815 | 2139 | 2026 | 2333 | 1657.65 | 1791 | 2166 | 1.307 | 0.387 | 0.0008452 |
| AT1G22960.1 | | protein_coding | Pentatricopeptide repeat | 187 | 148 | 124 | 144 | 113 | 97 | 393 | 364 | 317 | 312 | 300 | 264 | 255.74 | 358 | 292 | 1.307 | 0.386 | 0.00693486 |
| AT3G58840.1 | PMD1 | protein_coding | Tropomyosin-related | 44 | 24 | 28 | 35 | 28 | 14 | 92 | 59 | 72 | 76 | 74 | 38 | 335.12 | 74 | 63 | 1.307 | 0.386 | 0.00523411 |
| AT4G01100.2 | ADNT1 | protein_coding | adenine nucleotide transp | 1625 | 1204 | 1007 | 1975 | 1355 | 1280 | 3416 | 2958 | 2578 | 4284 | 3597 | 3481 | 3453.7 | 2984 | 3787 | 1.307 | 0.386 | 0.00298001 |
| AT5G05450.1 | RH18 | protein_coding | P-loop containing nucleo | 591 | 422 | 412 | 531 | 374 | 452 | 1242 | 1037 | 1055 | 1152 | 993 | 1229 | 1009.84 | 1111 | 1125 | 1.307 | 0.386 | 0.00183885 |
| AT4G23890.1 | ndhS | protein_coding | | 1461 | 1416 | 1547 | 1932 | 1658 | 1538 | 3071 | 3479 | 3960 | 4191 | 4402 | 4182 | 3996.55 | 3503 | 4258 | 1.307 | 0.386 | 0.00146753 |
| AT4G39150.1 | | protein_coding | DNAJ heat shock N-termi | 1014 | 740 | 682 | 1046 | 846 | 722 | 2132 | 1818 | 1746 | 2269 | 2246 | 1963 | 2139.73 | 1899 | 2159 | 1.307 | 0.386 | 0.0013501 |
| AT1G31190.1 | IMPL1 | protein_coding | myo-inositol monophospl | 631 | 612 | 551 | 751 | 609 | 699 | 1326 | 1504 | 1411 | 1629 | 1617 | 1901 | 1804.84 | 1414 | 1716 | 1.307 | 0.386 | 0.00102361 |
| AT4G20830.1 | | protein_coding | FAD-binding Berberine fa | 369 | 254 | 290 | 486 | 259 | 361 | 776 | 624 | 742 | 1054 | 688 | 982 | 782 | 714 | 908 | 1.306 | 0.385 | 0.00699463 |
| AT4G33865.1 | RPS29A | protein_coding | Ribosomal protein S14p/ | 1639 | 1312 | 999 | 2016 | 1427 | 1445 | 3445 | 3223 | 2558 | 4373 | 3788 | 3930 | 4779.6 | 3075 | 4030 | 1.306 | 0.385 | 0.00610171 |
| AT3G22590.1 | CDC37 | protein_coding | PLANT HOMOLOGOUS TO | 262 | 188 | 256 | 295 | 242 | 256 | 551 | 462 | 655 | 640 | 642 | 696 | 579.17 | 556 | 659 | 1.306 | 0.385 | 0.00321388 |
| AT5G21930.1 | PAA2 | protein_coding | P-type ATPase of Arabido | 1835 | 1826 | 1847 | 1974 | 1706 | 1969 | 3857 | 4486 | 4728 | 4282 | 4529 | 5355 | 4695.59 | 4357 | 4722 | 1.306 | 0.385 | 0.00215529 |
| AT2G03980.1 | | protein_coding | GDSL-like Lipase/Acylhyd | 438 | 316 | 293 | 563 | 388 | 388 | 921 | 776 | 750 | 1221 | 1030 | 1055 | 898.51 | 816 | 1102 | 1.306 | 0.385 | 0.00137025 |
| AT1G70490.1 | ARF2-A | protein_coding | Ras-related small GTP-bir | 1461 | 1508 | 1134 | 1961 | 1760 | 1689 | 3071 | 3705 | 2903 | 4254 | 4672 | 4593 | 5048.83 | 3226 | 4506 | 1.306 | 0.385 | 0.00115503 |
| AT4G30760.1 | | protein_coding | Putative endonuclease or | 481 | 413 | 338 | 641 | 559 | 530 | 1011 | 1015 | 865 | 1390 | 1484 | 1441 | 1410.62 | 964 | 1438 | 1.306 | 0.385 | 0.00106155 |
| AT3G08010.1 | ATAB2 | protein_coding | RNA binding | 1144 | 986 | 908 | 1332 | 1093 | 967 | 2405 | 2422 | 2325 | 2889 | 2902 | 2630 | 3445.13 | 2384 | 2807 | 1.306 | 0.385 | 0.00065027 |
| AT1G54380.1 | | protein_coding | spliceosome protein-relat | 288 | 204 | 224 | 315 | 267 | 229 | 605 | 501 | 573 | 683 | 709 | 623 | 589.83 | 560 | 672 | 1.306 | 0.385 | 0.000419 |
| AT1G74710.2 | ICS1 | protein_coding | ADC synthase superfamily | 243 | 231 | 221 | 238 | 189 | 160 | 511 | 568 | 566 | 516 | 502 | 435 | 411.22 | 548 | 484 | 1.305 | 0.384 | 0.00631983 |
| AT3G07230.1 | | protein_coding | wound-responsive protei | 3922 | 3653 | 4136 | 6391 | 5201 | 4656 | 8244 | 8975 | 10588 | 13863 | 13807 | 12662 | 6976.61 | 9269 | 13444 | 1.305 | 0.384 | 0.00340616 |
| AT5G12140.1 | CYS1 | protein_coding | cystatin-1 | 788 | 822 | 708 | 1217 | 1108 | 1184 | 1656 | 2020 | 1813 | 2640 | 2941 | 3220 | 2941.12 | 1830 | 2934 | 1.305 | 0.384 | 0.00167943 |
| AT2G34730.1 | WAP | protein_coding | myosin heavy chain-relat | 250 | 183 | 232 | 278 | 235 | 229 | 526 | 450 | 594 | 603 | 624 | 623 | 482.92 | 523 | 617 | 1.305 | 0.384 | 0.00150499 |
| AT3G12800.1 | SDRB | protein_coding | short-chain dehydrogena | 901 | 701 | 558 | 1141 | 801 | 745 | 1894 | 1722 | 1429 | 2475 | 2126 | 2026 | 2277.01 | 1682 | 2209 | 1.304 | 0.383 | 0.00247999 |
| AT1G75210.1 | | protein_coding | HAD-superfamily hydrola | 1051 | 879 | 925 | 1183 | 991 | 801 | 2209 | 2160 | 2368 | 2566 | 2631 | 2178 | 2368.33 | 2246 | 2458 | 1.304 | 0.383 | 0.0009104 |
| AT1G61010.1 | CPSF73-I | protein_coding | cleavage and polyadenyla | 644 | 532 | 509 | 644 | 578 | 539 | 1354 | 1307 | 1303 | 1397 | 1534 | 1466 | 1239.75 | 1321 | 1466 | 1.304 | 0.383 | 0.00086163 |
| AT5G58870.1 | FTSH9 | protein_coding | FTSH protease 9 | 1193 | 1016 | 1065 | 1669 | 1388 | 1313 | 2508 | 2496 | 2726 | 3620 | 3685 | 3571 | 2807.56 | 2577 | 3625 | 1.304 | 0.383 | 0.00056025 |
| AT4G28830.1 | | protein_coding | S-adenosyl-L-methionine- | 118 | 97 | 94 | 133 | 107 | 112 | 248 | 238 | 241 | 288 | 284 | 305 | 272.6 | 242 | 292 | 1.304 | 0.382 | 0.00517199 |
| AT1G05860.1 | | protein_coding | | 174 | 111 | 128 | 178 | 139 | 147 | 366 | 273 | 328 | 386 | 369 | 400 | 343.32 | 322 | 385 | 1.303 | 0.382 | 0.00274016 |
| AT4G02840.2 | | protein_coding | Small nuclear ribonucleop | 350 | 298 | 260 | 383 | 329 | 306 | 736 | 732 | 666 | 831 | 873 | 832 | 907.92 | 711 | 845 | 1.303 | 0.382 | 0.00146379 |
| AT3G05500.1 | | protein_coding | Rubber elongation factor | 462 | 429 | 330 | 491 | 405 | 446 | 971 | 1054 | 845 | 1065 | 1075 | 1213 | 1075.57 | 957 | 1118 | 1.302 | 0.381 | 0.00264657 |
| AT1G73130.1 | | protein_coding | | 144 | 100 | 133 | 141 | 123 | 106 | 303 | 246 | 340 | 306 | 327 | 288 | 246.76 | 296 | 307 | 1.302 | 0.38 | 0.00798822 |
| AT5G15090.1 | VDAC3 | protein_coding | voltage dependent anion | 2298 | 1619 | 1343 | 2823 | 1972 | 2029 | 4831 | 3978 | 3438 | 6123 | 5235 | 5518 | 5027.15 | 4082 | 5625 | 1.301 | 0.38 | 0.00432053 |
| AT3G06790.1 | MORF3 | protein_coding | plastid developmental pr | 202 | 184 | 121 | 221 | 193 | 220 | 425 | 452 | 310 | 479 | 512 | 598 | 647.78 | 396 | 530 | 1.301 | 0.38 | 0.00412214 |
| AT1G22950.1 | | protein_coding | 2-oxoglutarate (2OG) anc | 235 | 189 | 178 | 282 | 254 | 193 | 494 | 464 | 456 | 612 | 674 | 525 | 526.55 | 471 | 604 | 1.301 | 0.38 | 0.00122558 |
| AT5G50320.1 | HAG3 | protein_coding | radical SAM domain-cont | 368 | 342 | 295 | 434 | 346 | 404 | 774 | 840 | 755 | 941 | 919 | 1099 | 852.43 | 790 | 986 | 1.301 | 0.38 | 0.00086707 |
| AT1G53090.1 | SPA4 | protein_coding | SPA1-related 4 | 423 | 304 | 355 | 563 | 402 | 361 | 889 | 747 | 909 | 1221 | 1067 | 982 | 710.68 | 848 | 1090 | 1.301 | 0.379 | 0.003925 |
| AT2G20950.4 | | protein_coding | Arabidopsis phospholipas | 101 | 120 | 68 | 129 | 102 | 87 | 212 | 295 | 174 | 280 | 271 | 237 | 624.74 | 227 | 263 | 1.3 | 0.379 | 0.00331691 |
| AT5G53580.1 | PLR1 | protein_coding | NAD(P)-linked oxidoredu | 1330 | 1065 | 869 | 1289 | 1078 | 922 | 2796 | 2617 | 2225 | 2796 | 2862 | 2507 | 2881.6 | 2546 | 2722 | 1.301 | 0.379 | 0.00193595 |
| AT2G05620.1 | PGR5 | protein_coding | proton gradient regulatio | 1920 | 1584 | 1441 | 2517 | 1976 | 1769 | 4036 | 3892 | 3689 | 5460 | 5246 | 4811 | 5220.04 | 3872 | 5172 | 1.3 | 0.379 | 0.00158279 |
| AT1G57770.1 | | protein_coding | FAD/NAD(P)-binding oxid | 1023 | 960 | 862 | 1217 | 1023 | 1109 | 2150 | 2359 | 2207 | 2640 | 2716 | 3016 | 3050.09 | 2239 | 2791 | 1.3 | 0.379 | 0.00089822 |
| AT5G25560.2 | | protein_coding | CHY-type/CTCHY-type/RII | 584 | 531 | 702 | 855 | 841 | 748 | 1228 | 1305 | 1797 | 1855 | 2233 | 2034 | 2099.21 | 1443 | 2041 | 1.299 | 0.378 | 0.00813404 |
| AT2G34970.1 | | protein_coding | Trimeric LpxA-like enzym | 813 | 543 | 487 | 814 | 601 | 529 | 1709 | 1334 | 1247 | 1766 | 1596 | 1439 | 1212.32 | 1430 | 1600 | 1.3 | 0.378 | 0.0054128 |
| AT1G09140.1 | SR30 | protein_coding | SERINE-ARGININE PROTE | 589 | 450 | 458 | 606 | 548 | 569 | 1238 | 1106 | 1173 | 1314 | 1455 | 1547 | 1516.35 | 1172 | 1439 | 1.299 | 0.378 | 0.00188798 |
| AT1G26170.1 | | protein_coding | ARM repeat superfamily | 702 | 537 | 511 | 799 | 634 | 557 | 1476 | 1319 | 1308 | 1733 | 1683 | 1515 | 1351.78 | 1368 | 1644 | 1.3 | 0.378 | 0.00117387 |
| AT3G14100.1 | UBP1C | protein_coding | RNA-binding (RRM/RBD/ | 1328 | 1155 | 1180 | 2063 | 1710 | 1633 | 2792 | 2838 | 3021 | 4475 | 4540 | 4441 | 3162.41 | 2884 | 4485 | 1.3 | 0.378 | 0.00097001 |
| AT1G11910.1 | APA1 | protein_coding | aspartic proteinase A1 | 4538 | 4672 | 4094 | 5642 | 5268 | 5376 | 9539 | 11478 | 10481 | 12238 | 13985 | 14620 | 11452.41 | 10499 | 13614 | 1.298 | 0.377 | 0.00294817 |
| AT3G49560.1 | | protein_coding | Mitochondrial import inn | 1087 | 845 | 782 | 989 | 843 | 858 | 2285 | 2076 | 2002 | 2145 | 2238 | 2333 | 2475.91 | 2121 | 2239 | 1.298 | 0.377 | 0.00171712 |
| AT1G56200.1 | emb1303 | protein_coding | embryo defective 1303 | 488 | 376 | 377 | 418 | 335 | 482 | 1026 | 924 | 965 | 907 | 889 | 1311 | 2161.5 | 972 | 1036 | 1.298 | 0.376 | 0.00622961 |
| AT1G14300.2 | | protein_coding | ARM repeat superfamily | 210 | 130 | 121 | 171 | 141 | 130 | 441 | 319 | 310 | 371 | 374 | 354 | 381.11 | 357 | 366 | 1.297 | 0.376 | 0.00402848 |
| AT4G02920.2 | | protein_coding | | 5156 | 4300 | 3746 | 5204 | 3849 | 3133 | 10838 | 10564 | 9590 | 11288 | 10218 | 8520 | 9906.19 | 10331 | 10009 | 1.298 | 0.376 | 0.00356227 |
| AT5G20130.1 | | protein_coding | | 382 | 314 | 270 | 471 | 381 | 311 | 803 | 771 | 691 | 1022 | 1011 | 846 | 1077.6 | 755 | 960 | 1.297 | 0.376 | 0.00279202 |
| AT5G44320.1 | | protein_coding | Eukaryotic translation ini | 1300 | 988 | 874 | 1458 | 1099 | 1032 | 2733 | 2427 | 2237 | 3163 | 2918 | 2806 | 2418.16 | 2466 | 2962 | 1.298 | 0.376 | 0.00177182 |
| AT5G06290.1 | | protein_coding | SAC3/GANP/Nin1/mts3/e | 534 | 482 | 504 | 627 | 562 | 526 | 1123 | 1184 | 1290 | 1360 | 1492 | 1430 | 1148.86 | 1199 | 1427 | 1.297 | 0.376 | 0.00115793 |
| AT1G06430.1 | FTSH8 | protein_coding | FTSH protease 8 | 2006 | 1743 | 1825 | 2855 | 2074 | 2085 | 4217 | 4282 | 4672 | 6193 | 5506 | 5670 | 4251.49 | 4390 | 5790 | 1.298 | 0.376 | 0.00100665 |
| AT4G35460.1 | NTR1 | protein_coding | NADPH-dependent thiore | 350 | 320 | 344 | 414 | 322 | 406 | 736 | 786 | 881 | 898 | 855 | 1104 | 1035.1 | 801 | 952 | 1.297 | 0.375 | 0.00263767 |
| AT5G09250.1 | KIWI | protein_coding | ssDNA-binding transcript | 159 | 123 | 139 | 171 | 134 | 141 | 334 | 302 | 356 | 371 | 356 | 383 | 383.44 | 331 | 370 | 1.297 | 0.375 | 0.00142606 |
| AT4G10760.1 | MTA | protein_coding | mRNAadenosine methyla | 457 | 348 | 320 | 499 | 403 | 360 | 961 | 855 | 819 | 1082 | 1070 | 979 | 898.81 | 878 | 1044 | 1.297 | 0.375 | 0.00095824 |
| AT2G28130.1 | | protein_coding | | 184 | 155 | 167 | 217 | 135 | 142 | 387 | 381 | 428 | 471 | 358 | 386 | 371.18 | 399 | 405 | 1.296 | 0.374 | 0.0073553 |
| AT5G48545.1 | HINT3 | protein_coding | histidine triad nucleotide | 230 | 276 | 226 | 361 | 270 | 322 | 483 | 678 | 579 | 783 | 717 | 876 | 894.6 | 580 | 792 | 1.296 | 0.374 | 0.00582818 |
| AT1G79490.1 | EMB2217 | protein_coding | Pentatricopeptide repeat | 165 | 136 | 107 | 157 | 137 | 128 | 347 | 334 | 274 | 341 | 364 | 348 | 279.78 | 318 | 351 | 1.295 | 0.373 | 0.00521377 |
| AT5G13710.1 | SMT1 | protein_coding | sterol methyltransferase | 2232 | 1708 | 2014 | 2797 | 1765 | 2230 | 4692 | 4196 | 5156 | 6067 | 4686 | 6064 | 5522.73 | 4681 | 5606 | 1.295 | 0.373 | 0.00379063 |
| AT1G12250.1 | | protein_coding | Pentapeptide repeat-con | 591 | 694 | 576 | 665 | 598 | 658 | 1242 | 1705 | 1475 | 1442 | 1588 | 1789 | 2028.92 | 1474 | 1606 | 1.295 | 0.373 | 0.00371271 |
| AT5G37850.1 | PK | protein_coding | pfkB-like carbohydrate ki | 606 | 488 | 487 | 640 | 495 | 490 | 1274 | 1199 | 1247 | 1388 | 1314 | 1333 | 1344.06 | 1240 | 1345 | 1.295 | 0.373 | 0.0011715 |
| AT5G35210.1 | | protein_coding | metalloendopeptidases;z | 2317 | 1975 | 2113 | 2494 | 2154 | 2069 | 4871 | 4852 | 5409 | 5410 | 5718 | 5627 | 4005.1 | 5044 | 5585 | 1.295 | 0.373 | 0.00089657 |
| AT3G12400.1 | ELC | protein_coding | Ubiquitin-conjugating en | 238 | 211 | 176 | 282 | 243 | 271 | 500 | 518 | 451 | 612 | 645 | 737 | 671.17 | 490 | 665 | 1.295 | 0.373 | 0.00050632 |
| AT5G24020.1 | MIND1 | protein_coding | septum site-determining | 457 | 425 | 295 | 490 | 403 | 455 | 961 | 1044 | 755 | 1063 | 1070 | 1237 | 1148.44 | 920 | 1123 | 1.294 | 0.372 | 0.00556057 |
| AT4G36520.1 | | protein_coding | Chaperone DnaJ-domain | 227 | 218 | 241 | 246 | 260 | 237 | 477 | 536 | 617 | 534 | 690 | 645 | 602.28 | 543 | 623 | 1.294 | 0.372 | 0.00513389 |
| AT4G17520.1 | | protein_coding | Hyaluronan / mRNA bind | 1183 | 1027 | 827 | 1337 | 952 | 1265 | 2487 | 2523 | 2117 | 2900 | 2527 | 3440 | 5267.24 | 2376 | 2956 | 1.294 | 0.372 | 0.00485283 |
| AT4G30960.1 | CIPK6 | protein_coding | SOS3-interacting protein | 2306 | 2417 | 1871 | 2718 | 2314 | 2420 | 4847 | 5938 | 4790 | 5896 | 6143 | 6581 | 6224.01 | 5192 | 6207 | 1.294 | 0.372 | 0.00316516 |
| AT3G54130.1 | | protein_coding | Josephin family protein | 99 | 101 | 92 | 156 | 115 | 118 | 208 | 248 | 236 | 338 | 305 | 321 | 274.37 | 231 | 321 | 1.295 | 0.372 | 0.00278674 |
| AT3G54480.1 | SKIP5 | protein_coding | SKP1/ASK-interacting pro | 253 | 266 | 268 | 358 | 329 | 324 | 532 | 654 | 686 | 777 | 873 | 881 | 835.78 | 624 | 844 | 1.294 | 0.372 | 0.00274765 |
| AT2G25670.1 | | protein_coding | | 1235 | 921 | 826 | 1267 | 1005 | 965 | 2596 | 2263 | 2115 | 2748 | 2668 | 2624 | 2753.65 | 2325 | 2680 | 1.294 | 0.372 | 0.0016703 |
| AT5G13610.1 | | protein_coding | Protein of unknown funct | 259 | 190 | 158 | 306 | 224 | 212 | 544 | 467 | 404 | 664 | 595 | 577 | 580.63 | 472 | 612 | 1.294 | 0.372 | 0.00136908 |
| AT1G75660.1 | XRN3 | protein_coding | 5'-3' exoribonuclease 3 | 1526 | 1234 | 1302 | 1531 | 1305 | 1433 | 3208 | 3032 | 3333 | 3321 | 3464 | 3897 | 2768.61 | 3191 | 3561 | 1.294 | 0.372 | 0.00129922 |
| AT4G11900.1 | | protein_coding | S-locus lectin protein kina | 308 | 286 | 261 | 428 | 284 | 228 | 647 | 703 | 668 | 928 | 754 | 620 | 592.28 | 673 | 767 | 1.293 | 0.371 | 0.00797706 |
| AT3G16520.3 | UGT88A1 | protein_coding | UDP-glucosyl transferase | 1349 | 1459 | 1013 | 1566 | 1242 | 959 | 2836 | 3584 | 2593 | 3397 | 3297 | 2608 | 4685.01 | 3004 | 3101 | 1.294 | 0.371 | 0.00680574 |
| AT5G10490.1 | MSL2 | protein_coding | MSCS-like 2 | 904 | 601 | 620 | 1165 | 833 | 761 | 1900 | 1477 | 1587 | 2527 | 2211 | 2069 | 1799.15 | 1655 | 2269 | 1.293 | 0.371 | 0.00360698 |
| AT4G18700.1 | CIPK12 | protein_coding | CBL-interacting protein ki | 1084 | 829 | 815 | 1386 | 1003 | 901 | 2279 | 2037 | 2086 | 3006 | 2663 | 2450 | 2217.85 | 2134 | 2706 | 1.294 | 0.371 | 0.00292008 |
| AT2G11910.1 | | protein_coding | | 883 | 796 | 663 | 885 | 762 | 778 | 1856 | 1956 | 1697 | 1920 | 2023 | 2116 | 2203.38 | 1836 | 2020 | 1.293 | 0.371 | 0.00277254 |
| AT3G62190.1 | | protein_coding | Chaperone DnaJ-domain | 236 | 243 | 277 | 349 | 268 | 267 | 496 | 597 | 709 | 757 | 711 | 726 | 731.53 | 601 | 731 | 1.294 | 0.371 | 0.00213899 |
| AT5G59460.1 | | protein_coding | scarecrow-like transcripti | 180 | 152 | 141 | 200 | 157 | 160 | 378 | 373 | 361 | 434 | 417 | 435 | 358.75 | 371 | 429 | 1.293 | 0.371 | 0.00476198 |
| AT1G33270.1 | | protein_coding | Acyl transferase/acyl hyd | 292 | 232 | 255 | 432 | 307 | 315 | 614 | 570 | 653 | 937 | 815 | 857 | 690.17 | 612 | 870 | 1.293 | 0.37 | 0.00245064 |
| AT2G43750.1 | OASB | protein_coding | O-acetylserine (thiol) lyas | 3343 | 3043 | 2703 | 3541 | 3118 | 3281 | 7027 | 7476 | 6920 | 7681 | 8278 | 8922 | 7905.09 | 7141 | 8294 | 1.293 | 0.37 | 0.00134476 |
| AT5G05180.1 | | protein_coding | | 432 | 310 | 343 | 516 | 391 | 372 | 908 | 762 | 878 | 1119 | 1038 | 1012 | 921.33 | 849 | 1056 | 1.292 | 0.37 | 0.00130233 |
| AT1G01170.1 | | protein_coding | Protein of unknown funct | 688 | 597 | 641 | 905 | 736 | 736 | 1446 | 1467 | 1641 | 1963 | 1954 | 2002 | 1929.1 | 1518 | 1973 | 1.292 | 0.37 | 0.00124932 |
| AT3G01180.1 | SS2 | protein_coding | starch synthase 2 | 886 | 715 | 697 | 941 | 820 | 731 | 1862 | 1757 | 1784 | 2041 | 2177 | 1988 | 1924.3 | 1801 | 2069 | 1.292 | 0.37 | 0.00070232 |
| AT3G13580.3 | RPL7D | protein_coding | Ribosomal protein L30/L7 | 1371 | 907 | 900 | 1253 | 881 | 923 | 2882 | 2228 | 2304 | 2718 | 2339 | 2510 | 2432.17 | 2471 | 2522 | 1.291 | 0.369 | 0.00660544 |
| AT4G24060.1 | DOF4.6 | protein_coding | Dof-type zinc finger DNA- | 260 | 219 | 249 | 307 | 300 | 313 | 547 | 538 | 637 | 666 | 796 | 851 | 671.14 | 574 | 771 | 1.291 | 0.369 | 0.00232227 |
| AT5G04710.1 | | protein_coding | Zn-dependent exopeptid | 481 | 356 | 329 | 490 | 358 | 328 | 1011 | 875 | 842 | 1063 | 950 | 892 | 1326.9 | 909 | 968 | 1.291 | 0.369 | 0.00211181 |
| AT1G09800.1 | RPS18A | protein_coding | S18 ribosomal protein | 2850 | 2199 | 2005 | 3110 | 2348 | 2290 | 5991 | 5403 | 5133 | 6746 | 6233 | 6228 | 5936.14 | 5509 | 6402 | 1.292 | 0.369 | 0.00160322 |
| AT2G02500.1 | ISPD | protein_coding | Nucleotide-diphospho-su | 642 | 564 | 615 | 802 | 645 | 610 | 1350 | 1386 | 1574 | 1740 | 1712 | 1659 | 1638.06 | 1437 | 1704 | 1.292 | 0.369 | 0.00109632 |
| AT5G02720.1 | | protein_coding | | 404 | 349 | 340 | 456 | 372 | 425 | 849 | 857 | 870 | 989 | 988 | 1156 | 933.81 | 859 | 1044 | 1.291 | 0.369 | 0.00108194 |
| AT5G24690.1 | | protein_coding | Protein of unknown funct | 3613 | 2651 | 2599 | 4032 | 3134 | 3019 | 7595 | 6513 | 6654 | 8746 | 8320 | 8210 | 7262.47 | 6921 | 8425 | 1.291 | 0.368 | 0.00121317 |
| AT1G62600.1 | | protein_coding | Flavin-binding monooxyg | 287 | 290 | 285 | 358 | 269 | 303 | 603 | 712 | 730 | 777 | 714 | 824 | 750.6 | 682 | 772 | 1.291 | 0.368 | 0.00092567 |
| AT1G06950.1 | TIC110 | protein_coding | translocon at the inner er | 6201 | 5024 | 3905 | 5926 | 4825 | 4466 | 13035 | 12343 | 9997 | 12854 | 12809 | 12145 | 10648.35 | 11792 | 12603 | 1.289 | 0.367 | 0.00451377 |
| AT5G22040.1 | | protein_coding | | 151 | 109 | 146 | 162 | 122 | 141 | 317 | 268 | 374 | 351 | 324 | 383 | 396.32 | 320 | 353 | 1.289 | 0.366 | 0.00650402 |
| AT4G26840.1 | SUMO1 | protein_coding | small ubiquitin-like modi | 486 | 563 | 419 | 545 | 478 | 520 | 1022 | 1383 | 1073 | 1182 | 1269 | 1414 | 1765.66 | 1159 | 1288 | 1.289 | 0.366 | 0.00600375 |
| AT5G23820.1 | | protein_coding | MD-2-related lipid recog | 2364 | 1847 | 1522 | 4247 | 3158 | 2797 | 4969 | 4538 | 3896 | 9212 | 8384 | 7606 | 8572.42 | 4468 | 8401 | 1.289 | 0.366 | 0.00551469 |
| AT5G13970.1 | | protein_coding | | 210 | 176 | 196 | 218 | 178 | 203 | 441 | 432 | 502 | 473 | 473 | 552 | 449.86 | 458 | 499 | 1.289 | 0.366 | 0.00160208 |
| AT1G14650.1 | | protein_coding | SWAP (Suppressor-of-Wh | 911 | 787 | 737 | 991 | 825 | 811 | 1915 | 1934 | 1887 | 2150 | 2190 | 2205 | 2014.01 | 1912 | 2182 | 1.289 | 0.366 | 0.00093289 |
| AT3G59970.3 | MTHFR1 | protein_coding | methylenetetrahydrofola | 8221 | 6490 | 5359 | 9034 | 7692 | 6126 | 17281 | 15945 | 13719 | 19596 | 20420 | 16659 | 17176.11 | 15648 | 18892 | 1.288 | 0.365 | 0.00781405 |
| AT2G29890.3 | VLN1 | protein_coding | villin-like 1 | 393 | 364 | 450 | 492 | 463 | 408 | 826 | 894 | 1152 | 1067 | 1229 | 1110 | 894.29 | 957 | 1135 | 1.288 | 0.365 | 0.00453972 |
| AT4G09040.1 | | protein_coding | RNA-binding (RRM/RBD/ | 1729 | 1463 | 1398 | 1766 | 1606 | 1464 | 3634 | 3594 | 3579 | 3831 | 4264 | 3981 | 3992.41 | 3602 | 4025 | 1.288 | 0.365 | 0.00115645 |
| AT2G03070.1 | MED8 | protein_coding | mediator subunit 8 | 353 | 313 | 291 | 405 | 325 | 337 | 742 | 769 | 745 | 878 | 863 | 916 | 808.56 | 752 | 886 | 1.288 | 0.365 | 0.00063889 |
| AT1G55900.1 | TIM50 | protein_coding | Haloacid dehalogenase-li | 375 | 235 | 210 | 376 | 260 | 266 | 788 | 577 | 538 | 816 | 690 | 723 | 731.82 | 634 | 743 | 1.287 | 0.364 | 0.00525045 |
| AT3G61140.1 | CSN1 | protein_coding | 26S proteasome, regulato | 655 | 527 | 414 | 822 | 595 | 566 | 1377 | 1295 | 1060 | 1783 | 1580 | 1539 | 1316.57 | 1244 | 1634 | 1.287 | 0.364 | 0.00448628 |
| AT2G33730.1 | RH21 | protein_coding | P-loop containing nucleo | 725 | 615 | 605 | 745 | 734 | 794 | 1524 | 1511 | 1549 | 1616 | 1949 | 2159 | 1693.52 | 1528 | 1908 | 1.287 | 0.364 | 0.00274256 |
| AT5G03190.2 | CPUORF47 | protein_coding | conserved peptide upstre | 181 | 196 | 152 | 242 | 271 | 198 | 380 | 482 | 389 | 525 | 719 | 538 | 543.25 | 417 | 594 | 1.286 | 0.363 | 0.00862436 |
| AT5G42070.1 | | protein_coding | | 555 | 581 | 435 | 610 | 540 | 563 | 1167 | 1427 | 1114 | 1323 | 1434 | 1531 | 1631.8 | 1236 | 1429 | 1.286 | 0.363 | 0.00480586 |
| AT2G17320.1 | | protein_coding | Uncharacterised conserve | 361 | 337 | 396 | 412 | 375 | 420 | 759 | 828 | 1014 | 894 | 996 | 1142 | 942.36 | 867 | 1011 | 1.286 | 0.363 | 0.00468055 |
| AT3G23100.1 | XRCC4 | protein_coding | homolog of human DNA l | 111 | 105 | 88 | 103 | 99 | 87 | 233 | 258 | 225 | 223 | 263 | 237 | 268.43 | 239 | 241 | 1.286 | 0.363 | 0.00431163 |
| AT1G73490.2 | | protein_coding | RNA-binding (RRM/RBD/ | 714 | 530 | 497 | 814 | 583 | 541 | 1501 | 1302 | 1272 | 1766 | 1548 | 1471 | 1564.58 | 1358 | 1595 | 1.287 | 0.363 | 0.00283612 |
| AT5G01350.1 | | protein_coding | | 679 | 534 | 607 | 934 | 749 | 771 | 1427 | 1312 | 1554 | 2026 | 1988 | 2097 | 1635.06 | 1431 | 2037 | 1.286 | 0.363 | 0.00149444 |
| AT4G34290.1 | | protein_coding | SWIB/MDM2 domain sup | 711 | 605 | 411 | 813 | 681 | 584 | 1495 | 1486 | 1052 | 1763 | 1808 | 1588 | 1953.79 | 1344 | 1720 | 1.285 | 0.362 | 0.00834549 |
| AT4G38020.1 | | protein_coding | tRNA/rRNA methyltransfe | 172 | 147 | 114 | 192 | 137 | 131 | 362 | 361 | 292 | 416 | 364 | 356 | 416.7 | 338 | 379 | 1.285 | 0.362 | 0.00644185 |

| | | | | | | | | | | | | | | | | | | | | | |
|---|---|---|---|---|---|---|---|---|---|---|---|---|---|---|---|---|---|---|---|---|---|
| AT5G61990.1 | | protein_coding | Pentatricopeptide repeat | 226 | 169 | 209 | 233 | 205 | 181 | 475 | 415 | 535 | 505 | 544 | 492 | 414.93 | 475 | 514 | 1.285 | 0.362 | 0.00228493 |
| AT5G51200.1 | NUP205 | protein_coding | Protein of unknown funct | 2842 | 2329 | 2250 | 3496 | 2531 | 2242 | 5974 | 5722 | 5760 | 7583 | 6719 | 6097 | 5023.6 | 5819 | 6800 | 1.286 | 0.362 | 0.00204514 |
| AT5G66720.1 | | protein_coding | Protein phosphatase 2C f | 1146 | 1019 | 1032 | 1326 | 1044 | 1085 | 2409 | 2503 | 2642 | 2876 | 2772 | 2951 | 2952.47 | 2518 | 2866 | 1.285 | 0.362 | 0.0010359 |
| AT2G16500.1 | SPE1 | protein_coding | arginine decarboxylase 1 | 1539 | 1480 | 989 | 1673 | 1186 | 1178 | 3235 | 3636 | 2532 | 3629 | 3149 | 3204 | 3912.18 | 3134 | 3327 | 1.284 | 0.361 | 0.00802331 |
| AT3G09820.1 | ADK1 | protein_coding | adenosine kinase 1 | 1767 | 1438 | 1049 | 2118 | 1660 | 1465 | 3714 | 3533 | 2686 | 4594 | 4407 | 3984 | 7038.55 | 3311 | 4328 | 1.284 | 0.361 | 0.0057496 |
| AT5G13010.1 | EMB3011 | protein_coding | RNA helicase family prote | 1289 | 1103 | 1205 | 1591 | 1293 | 1404 | 2710 | 2710 | 3085 | 3451 | 3433 | 3818 | 2478.52 | 2835 | 3567 | 1.284 | 0.361 | 0.00255689 |
| AT5G42270.1 | FTSH5 | protein_coding | FtsH extracellular proteas | 9231 | 8611 | 8619 | 10231 | 8538 | 9750 | 19404 | 21156 | 22065 | 22192 | 22666 | 26515 | 24473.55 | 20875 | 23791 | 1.283 | 0.36 | 0.00936378 |
| AT5G19220.1 | ADG2 | protein_coding | ADP glucose pyrophosph | 7191 | 6527 | 6085 | 7079 | 6677 | 6031 | 15116 | 16036 | 15578 | 15355 | 17726 | 16401 | 16419.03 | 15577 | 16494 | 1.283 | 0.36 | 0.00527839 |
| AT2G35450.1 | | protein_coding | catalytics;hydrolases | 394 | 392 | 356 | 390 | 372 | 347 | 828 | 963 | 911 | 846 | 988 | 944 | 1162.81 | 901 | 926 | 1.283 | 0.36 | 0.00239031 |
| AT5G47680.1 | TRM10 | protein_coding | | 262 | 245 | 226 | 300 | 247 | 272 | 551 | 602 | 579 | 651 | 656 | 740 | 683.41 | 577 | 682 | 1.283 | 0.36 | 0.00154524 |
| AT1G58270.1 | ZW9 | protein_coding | TRAF-like family protein | 1087 | 1048 | 786 | 1341 | 1508 | 1111 | 2285 | 2575 | 2012 | 2909 | 4003 | 3021 | 3217.91 | 2291 | 3311 | 1.283 | 0.359 | 0.00903695 |
| AT1G45170.1 | OEP24A | protein_coding | | 204 | 174 | 135 | 260 | 176 | 165 | 429 | 427 | 346 | 564 | 467 | 449 | 478.4 | 401 | 493 | 1.282 | 0.358 | 0.00686439 |
| AT4G33495.1 | RPD1 | protein_coding | Ubiquitin carboxyl-termir | 132 | 140 | 126 | 177 | 129 | 123 | 277 | 344 | 323 | 384 | 342 | 334 | 332.69 | 315 | 353 | 1.281 | 0.358 | 0.00482656 |
| AT5G23250.1 | | protein_coding | Succinyl-CoA ligase, alpha | 532 | 358 | 318 | 566 | 416 | 410 | 1118 | 880 | 814 | 1228 | 1104 | 1115 | 1009.14 | 937 | 1149 | 1.282 | 0.358 | 0.00449138 |
| AT4G04770.1 | ABCI8 | protein_coding | ATP binding cassette prot | 3045 | 2417 | 2392 | 3639 | 2575 | 2491 | 6401 | 5938 | 6124 | 7893 | 6836 | 6774 | 6919.17 | 6154 | 7168 | 1.282 | 0.358 | 0.00228018 |
| AT3G48070.2 | | protein_coding | RING/U-box superfamily | 297 | 260 | 230 | 331 | 241 | 241 | 624 | 639 | 589 | 718 | 640 | 655 | 679.86 | 617 | 671 | 1.282 | 0.358 | 0.00072042 |
| AT3G27925.1 | DEGP1 | protein_coding | DegP protease 1 | 1148 | 1376 | 1081 | 1451 | 1273 | 1269 | 2413 | 3381 | 2767 | 3147 | 3380 | 3451 | 4853.48 | 2854 | 3326 | 1.28 | 0.357 | 0.00514973 |
| AT3G20050.1 | CCT1 | protein_coding | T-complex protein 1 alph | 2586 | 1945 | 1638 | 3045 | 2409 | 2207 | 5436 | 4779 | 4193 | 6605 | 6395 | 6002 | 5413.03 | 4803 | 6334 | 1.281 | 0.357 | 0.00329654 |
| AT3G57610.1 | PURA | protein_coding | adenylosuccinate synthas | 2086 | 1743 | 1369 | 2386 | 1778 | 1814 | 4385 | 4282 | 3505 | 5175 | 4720 | 4933 | 4706.02 | 4057 | 4943 | 1.28 | 0.357 | 0.00271981 |
| AT5G57590.1 | BIO3-BIO1 | protein_coding | adenosylmethionine-8-ar | 145 | 122 | 96 | 141 | 141 | 89 | 305 | 300 | 246 | 306 | 374 | 242 | 382.77 | 284 | 307 | 1.28 | 0.356 | 0.00885333 |
| AT1G47420.1 | SDH5 | protein_coding | succinate dehydrogenase | 1533 | 1064 | 1025 | 1898 | 1274 | 1280 | 3222 | 2614 | 2624 | 4117 | 3382 | 3481 | 3573.85 | 2820 | 3660 | 1.28 | 0.356 | 0.00504469 |
| AT5G12010.1 | | protein_coding | | 268 | 198 | 188 | 279 | 178 | 204 | 563 | 486 | 481 | 605 | 473 | 555 | 543.91 | 510 | 544 | 1.28 | 0.356 | 0.00336483 |
| AT1G42550.1 | PMI1 | protein_coding | plastid movement impair | 730 | 643 | 571 | 780 | 556 | 697 | 1535 | 1580 | 1462 | 1692 | 1476 | 1895 | 2189.67 | 1526 | 1688 | 1.28 | 0.356 | 0.00272266 |
| AT3G54190.1 | | protein_coding | Transducin/WD40 repeat | 892 | 755 | 887 | 1018 | 950 | 844 | 1875 | 1855 | 2271 | 2208 | 2522 | 2295 | 1939.98 | 2000 | 2342 | 1.28 | 0.356 | 0.00258107 |
| AT4G27830.1 | BGLU10 | protein_coding | beta glucosidase 10 | 583 | 509 | 484 | 651 | 519 | 374 | 1226 | 1251 | 1239 | 1412 | 1378 | 1017 | 1051.99 | 1239 | 1269 | 1.279 | 0.355 | 0.00942089 |
| AT5G59870.1 | HTA6 | protein_coding | histone H2A 6 | 1272 | 1138 | 841 | 1095 | 1125 | 1028 | 2674 | 2796 | 2153 | 2375 | 2987 | 2796 | 3300.09 | 2541 | 2719 | 1.279 | 0.355 | 0.00621528 |
| AT2G21970.1 | sept-02 | protein_coding | stress enhanced protein 2 | 378 | 328 | 278 | 693 | 467 | 513 | 795 | 806 | 712 | 1503 | 1240 | 1395 | 1136.39 | 771 | 1379 | 1.279 | 0.355 | 0.00369673 |
| AT1G27340.1 | FBX6 | protein_coding | Galactose oxidase/kelch r | 468 | 412 | 317 | 578 | 484 | 437 | 984 | 1012 | 812 | 1254 | 1285 | 1188 | 1122.1 | 936 | 1242 | 1.279 | 0.355 | 0.00254501 |
| AT5G22330.1 | RIN1 | protein_coding | P-loop containing nucleo | 505 | 370 | 354 | 632 | 465 | 430 | 1062 | 909 | 906 | 1371 | 1234 | 1169 | 1019.23 | 959 | 1258 | 1.279 | 0.355 | 0.00246131 |
| AT1G48860.1 | | protein_coding | RNA 3'-terminal phospha | 935 | 756 | 657 | 1087 | 819 | 776 | 1965 | 1857 | 1682 | 2358 | 2174 | 2110 | 2243.82 | 1835 | 2214 | 1.279 | 0.355 | 0.00180696 |
| AT3G04605.1 | | protein_coding | | 564 | 495 | 423 | 575 | 504 | 464 | 1186 | 1216 | 1083 | 1247 | 1338 | 1262 | 1107 | 1162 | 1282 | 1.279 | 0.355 | 0.00170948 |
| AT3G21390.1 | | protein_coding | Mitochondrial substrate c | 227 | 175 | 167 | 311 | 246 | 222 | 477 | 430 | 428 | 675 | 653 | 604 | 614.61 | 445 | 644 | 1.279 | 0.355 | 0.00170806 |
| AT3G01370.1 | ATCFM2 | protein_coding | CRM family member 2 | 1971 | 1868 | 1694 | 2044 | 1706 | 1775 | 4143 | 4589 | 4337 | 4434 | 4529 | 4827 | 4460.02 | 4356 | 4597 | 1.279 | 0.355 | 0.00140163 |
| AT3G13330.1 | PA200 | protein_coding | proteasome activating pr | 1415 | 1119 | 1091 | 1731 | 1392 | 1340 | 2974 | 2749 | 2793 | 3755 | 3695 | 3644 | 2363.1 | 2839 | 3698 | 1.279 | 0.355 | 0.00120257 |
| AT4G27585.1 | | protein_coding | SPFH/Band 7/PHB domai | 374 | 294 | 248 | 459 | 341 | 322 | 786 | 722 | 635 | 996 | 905 | 876 | 863.98 | 714 | 926 | 1.278 | 0.354 | 0.0033806 |
| AT4G00660.2 | RH8 | protein_coding | RNAhelicase-like 8 | 922 | 795 | 713 | 1088 | 903 | 710 | 1938 | 1953 | 1825 | 2360 | 2397 | 1931 | 1847.47 | 1905 | 2229 | 1.278 | 0.354 | 0.00255284 |
| AT3G57870.1 | SCE1 | protein_coding | sumo conjugation enzym | 1502 | 1222 | 1338 | 1945 | 1587 | 1634 | 3157 | 3002 | 3425 | 4219 | 4213 | 4444 | 3999.57 | 3195 | 4292 | 1.278 | 0.354 | 0.0021317 |
| AT5G19990.1 | RPT6A | protein_coding | regulatory particle triple- | 1244 | 1102 | 961 | 1387 | 1110 | 1222 | 2615 | 2707 | 2460 | 3009 | 2947 | 3323 | 2899.57 | 2594 | 3093 | 1.278 | 0.354 | 0.00185797 |
| AT1G17880.1 | BTF3 | protein_coding | basic transcription factor | 3305 | 2318 | 2016 | 3387 | 2580 | 2251 | 6947 | 5695 | 5161 | 7347 | 6849 | 6121 | 6562.64 | 5934 | 6772 | 1.278 | 0.353 | 0.00754347 |
| AT4G36020.1 | CSP1 | protein_coding | cold shock domain protei | 111 | 116 | 86 | 120 | 89 | 121 | 233 | 285 | 220 | 260 | 236 | 329 | 677.29 | 246 | 275 | 1.277 | 0.353 | 0.00599699 |
| AT5G64400.2 | | protein_coding | | 838 | 754 | 571 | 1173 | 845 | 910 | 1762 | 1852 | 1462 | 2544 | 2243 | 2475 | 4155.9 | 1692 | 2421 | 1.277 | 0.353 | 0.00584567 |
| AT2G37410.1 | TIM17-2 | protein_coding | translocase inner membr | 740 | 584 | 505 | 900 | 682 | 650 | 1556 | 1435 | 1293 | 1952 | 1811 | 1768 | 1853.21 | 1428 | 1844 | 1.277 | 0.353 | 0.00337265 |
| AT3G47600.1 | MYB94 | protein_coding | myb domain protein 94 | 178 | 129 | 142 | 165 | 152 | 133 | 374 | 317 | 364 | 358 | 404 | 362 | 359.71 | 352 | 375 | 1.277 | 0.353 | 0.00274126 |
| AT5G61670.1 | | protein_coding | | 825 | 850 | 722 | 1135 | 1034 | 938 | 1734 | 2088 | 1848 | 2462 | 2745 | 2551 | 2403.18 | 1890 | 2586 | 1.277 | 0.353 | 0.0020355 |
| AT5G58130.1 | ROS3 | protein_coding | RNA-binding (RRM/RBD/ | 564 | 478 | 484 | 627 | 503 | 547 | 1186 | 1174 | 1239 | 1360 | 1335 | 1488 | 1026.75 | 1200 | 1394 | 1.277 | 0.353 | 0.00141002 |
| AT1G66510.1 | | protein_coding | AAR2 protein family | 142 | 99 | 90 | 132 | 97 | 89 | 298 | 243 | 230 | 286 | 258 | 242 | 246.56 | 257 | 262 | 1.277 | 0.352 | 0.0091097 |
| AT1G27200.1 | | protein_coding | Domain of unknown func | 214 | 176 | 174 | 253 | 189 | 195 | 450 | 432 | 445 | 549 | 502 | 530 | 368.47 | 442 | 527 | 1.276 | 0.352 | 0.00871259 |
| AT5G63150.1 | | protein_coding | | 422 | 257 | 307 | 420 | 303 | 287 | 887 | 631 | 786 | 911 | 804 | 780 | 750.1 | 768 | 832 | 1.276 | 0.352 | 0.00564966 |
| AT3G56860.3 | UBA2A | protein_coding | UBP1-associated protein | 995 | 1000 | 1004 | 1194 | 1099 | 1244 | 2092 | 2457 | 2570 | 2590 | 2918 | 3383 | 2806.35 | 2373 | 2964 | 1.276 | 0.352 | 0.00562072 |
| AT3G16640.1 | TCTP | protein_coding | translationally controlled | 4294 | 3252 | 3151 | 5427 | 4354 | 3945 | 9026 | 7990 | 8067 | 11772 | 11559 | 10728 | 15321.76 | 8361 | 11353 | 1.276 | 0.352 | 0.00461711 |
| AT1G03300.1 | ATDUF1 | protein_coding | DOMAIN OF UNKNOWN | 141 | 112 | 117 | 154 | 129 | 117 | 296 | 275 | 300 | 334 | 342 | 318 | 292.91 | 290 | 331 | 1.277 | 0.352 | 0.00434286 |
| AT2G03870.2 | LSM7 | protein_coding | Small nuclear ribonucleo | 368 | 303 | 299 | 448 | 349 | 406 | 774 | 744 | 765 | 972 | 927 | 1104 | 968.23 | 761 | 1001 | 1.276 | 0.352 | 0.00183502 |
| AT5G64280.1 | DIT2-2 | protein_coding | dicarboxylate transporter | 131 | 122 | 137 | 204 | 177 | 143 | 275 | 300 | 351 | 442 | 470 | 389 | 719.17 | 309 | 434 | 1.276 | 0.351 | 0.00882491 |
| AT2G41220.1 | GLU2 | protein_coding | glutamate synthase 2 | 958 | 700 | 650 | 1022 | 772 | 699 | 2014 | 1720 | 1664 | 2217 | 2049 | 1901 | 1467.2 | 1799 | 2056 | 1.276 | 0.351 | 0.00625301 |
| AT3G58470.1 | | protein_coding | nucleic acid binding;meth | 204 | 157 | 138 | 215 | 154 | 193 | 429 | 386 | 353 | 466 | 409 | 525 | 493.19 | 389 | 467 | 1.275 | 0.351 | 0.00618703 |
| AT1G06220.2 | MEE5 | protein_coding | Ribosomal protein S5/Elo | 2066 | 1536 | 1395 | 2122 | 1873 | 1527 | 4343 | 3774 | 3571 | 4603 | 4972 | 4153 | 3596.77 | 3896 | 4576 | 1.275 | 0.351 | 0.00383166 |
| AT5G45680.1 | FKBP13 | protein_coding | FK506-binding protein 13 | 1745 | 1696 | 1692 | 2091 | 1938 | 1684 | 3668 | 4167 | 4332 | 4536 | 5145 | 4580 | 5452.18 | 4056 | 4754 | 1.276 | 0.351 | 0.00273021 |
| AT3G44110.1 | ATJ3 | protein_coding | DNAJ homologue 3 | 3039 | 2493 | 2046 | 3512 | 2693 | 2642 | 6388 | 6125 | 5238 | 7618 | 7149 | 7185 | 6282.33 | 5917 | 7317 | 1.276 | 0.351 | 0.00271554 |
| AT1G77840.1 | | protein_coding | Translation initiation fact | 801 | 644 | 561 | 704 | 592 | 571 | 1684 | 1582 | 1436 | 1527 | 1572 | 1553 | 1564.6 | 1567 | 1551 | 1.276 | 0.351 | 0.00221413 |
| AT3G46740.1 | TOC75-3 | protein_coding | translocon at the outer e | 4027 | 3062 | 2507 | 4018 | 3076 | 3005 | 8465 | 7523 | 6418 | 8715 | 8166 | 8172 | 7580.04 | 7469 | 8351 | 1.275 | 0.35 | 0.00515706 |
| AT3G19670.1 | PRP40B | protein_coding | pre-mRNA-processing pro | 471 | 404 | 395 | 440 | 418 | 384 | 990 | 993 | 1011 | 954 | 1110 | 1044 | 966.35 | 998 | 1036 | 1.274 | 0.35 | 0.00212074 |
| AT3G04870.1 | ZDS1 | protein_coding | zeta-carotene desaturase | 2476 | 1903 | 1720 | 2675 | 1873 | 1820 | 5205 | 4675 | 4403 | 5802 | 4972 | 4949 | 4781.46 | 4761 | 5241 | 1.273 | 0.349 | 0.0034628 |
| AT3G24430.1 | HCF101 | protein_coding | ATP binding | 2329 | 2213 | 1929 | 2570 | 2225 | 2309 | 4896 | 5437 | 4938 | 5575 | 5907 | 6279 | 6445.44 | 5090 | 5920 | 1.274 | 0.349 | 0.00178238 |
| AT1G60170.1 | emb1220 | protein_coding | pre-mRNA processing rib | 845 | 726 | 752 | 1082 | 861 | 964 | 1776 | 1784 | 1925 | 2347 | 2286 | 2622 | 1939.39 | 1828 | 2418 | 1.273 | 0.349 | 0.00177635 |
| AT2G26190.1 | | protein_coding | calmodulin-binding famil | 348 | 401 | 335 | 434 | 376 | 404 | 732 | 985 | 858 | 941 | 998 | 1099 | 791.73 | 858 | 1013 | 1.273 | 0.348 | 0.00654069 |
| AT5G56000.1 | HSP90-4 | protein_coding | HEAT SHOCK PROTEIN 81 | 1310 | 1130 | 833 | 1342 | 1140 | 928 | 2754 | 2776 | 2133 | 2911 | 3026 | 2524 | 1982.3 | 2554 | 2820 | 1.272 | 0.347 | 0.00923973 |
| AT1G47500.1 | RBP47C' | protein_coding | RNA-binding protein 47C | 715 | 481 | 431 | 647 | 544 | 464 | 1503 | 1182 | 1103 | 1403 | 1444 | 1262 | 1372.98 | 1263 | 1370 | 1.272 | 0.347 | 0.00900904 |
| AT5G11340.1 | | protein_coding | Acyl-CoA N-acyltransfera | 377 | 237 | 234 | 436 | 281 | 298 | 792 | 582 | 599 | 946 | 746 | 810 | 766.93 | 658 | 834 | 1.272 | 0.347 | 0.00796601 |
| AT2G27770.1 | | protein_coding | Plant protein of unknowr | 353 | 332 | 337 | 414 | 375 | 269 | 742 | 816 | 863 | 898 | 996 | 732 | 693.39 | 807 | 875 | 1.271 | 0.346 | 0.0051533 |
| AT5G08680.1 | | protein_coding | ATP synthase alpha/beta | 2637 | 2052 | 1626 | 3448 | 2529 | 2578 | 5543 | 5041 | 4163 | 7479 | 6714 | 7011 | 6110.33 | 4916 | 7068 | 1.271 | 0.346 | 0.00499611 |
| AT1G03910.2 | CTN | protein_coding | | 637 | 554 | 445 | 622 | 519 | 594 | 1339 | 1361 | 1139 | 1349 | 1378 | 1615 | 1380.57 | 1280 | 1447 | 1.271 | 0.346 | 0.00477604 |
| AT5G46430.1 | RPL32B | protein_coding | Ribosomal protein L32e | 2522 | 1948 | 1621 | 2616 | 2053 | 2048 | 5301 | 4786 | 4150 | 5674 | 5450 | 5569 | 5392.37 | 4746 | 5564 | 1.271 | 0.346 | 0.00455334 |
| AT2G35635.1 | RUB2 | protein_coding | ubiquitin 7 | 1044 | 741 | 707 | 1288 | 973 | 879 | 2195 | 1821 | 1810 | 2794 | 2583 | 2390 | 2376.49 | 1942 | 2589 | 1.271 | 0.346 | 0.00451575 |
| AT1G50900.1 | LTD | protein_coding | Ankyrin repeat family pro | 2056 | 2212 | 1564 | 2409 | 2056 | 2087 | 4322 | 5434 | 4004 | 5225 | 5458 | 5675 | 6671.84 | 4587 | 5453 | 1.27 | 0.345 | 0.00676002 |
| AT3G50380.1 | | protein_coding | Protein of unknown funct | 1907 | 1625 | 1689 | 2195 | 1957 | 1689 | 4009 | 3992 | 4324 | 4761 | 5195 | 4593 | 3069.46 | 4108 | 4850 | 1.271 | 0.345 | 0.00213859 |
| AT3G06200.1 | GK3 | protein_coding | P-loop containing nucleo | 200 | 166 | 151 | 215 | 199 | 195 | 420 | 408 | 387 | 466 | 528 | 530 | 550.12 | 405 | 508 | 1.27 | 0.345 | 0.00080816 |
| AT5G13090.1 | | protein_coding | | 138 | 123 | 122 | 141 | 119 | 155 | 290 | 302 | 312 | 306 | 316 | 422 | 362.08 | 301 | 348 | 1.27 | 0.344 | 0.00930404 |
| AT5G56910.1 | | protein_coding | Proteinase inhibitor I25, | 116 | 131 | 119 | 114 | 107 | 107 | 244 | 322 | 305 | 247 | 284 | 291 | 337.53 | 290 | 274 | 1.27 | 0.344 | 0.00599996 |
| AT3G14400.1 | UBP25 | protein_coding | ubiquitin-specific proteas | 995 | 790 | 757 | 1142 | 878 | 718 | 2092 | 1941 | 1938 | 2477 | 2331 | 1953 | 1980.07 | 1990 | 2254 | 1.269 | 0.344 | 0.00572094 |
| AT1G04950.1 | TAF6 | protein_coding | TATA BOX ASSOCIATED F | 331 | 277 | 301 | 403 | 412 | 366 | 696 | 681 | 771 | 874 | 1094 | 995 | 833.08 | 716 | 988 | 1.269 | 0.344 | 0.00498759 |
| AT5G36790.1 | PGLP1B | protein_coding | Haloacid dehalogenase-li | 4112 | 3250 | 3129 | 4334 | 3736 | 3051 | 8644 | 7985 | 8010 | 9401 | 9918 | 8297 | 9489.94 | 8213 | 9205 | 1.27 | 0.344 | 0.00462992 |
| AT5G36700.1 | PGLP1B | protein_coding | 2-phosphoglycolate phos | 4027 | 3305 | 3055 | 4214 | 3718 | 3195 | 8465 | 8120 | 7821 | 9141 | 9870 | 8689 | 9460 | 8135 | 9233 | 1.269 | 0.344 | 0.00359564 |
| AT1G17130.2 | | protein_coding | Family of unknown functi | 511 | 400 | 383 | 584 | 513 | 472 | 1074 | 983 | 981 | 1267 | 1362 | 1284 | 1204.64 | 1013 | 1304 | 1.27 | 0.344 | 0.00240462 |
| AT1G10580.1 | | protein_coding | Transducin/WD40 repeat | 309 | 301 | 273 | 379 | 306 | 338 | 650 | 740 | 699 | 822 | 812 | 919 | 1038.81 | 696 | 851 | 1.269 | 0.344 | 0.00218453 |
| AT3G43690.1 | | transposable_elen | transposable element ger | 130 | 129 | 117 | 174 | 144 | 140 | 273 | 317 | 300 | 377 | 382 | 381 | 729.57 | 297 | 380 | 1.27 | 0.344 | 0.00193595 |
| AT3G27090.1 | | protein_coding | DCD (Development and C | 2897 | 2134 | 2685 | 3303 | 2487 | 2768 | 6090 | 5243 | 6874 | 7165 | 6602 | 7527 | 5530.39 | 6069 | 7098 | 1.268 | 0.343 | 0.00815971 |
| AT4G29040.1 | RPT2A | protein_coding | regulatory particle AAA-A | 1105 | 878 | 882 | 1217 | 946 | 952 | 2323 | 2157 | 2258 | 2640 | 2511 | 2589 | 3214.49 | 2246 | 2580 | 1.268 | 0.343 | 0.00203094 |
| AT1G19710.1 | | protein_coding | UDP-Glycosyltransferase | 245 | 199 | 235 | 335 | 233 | 225 | 515 | 489 | 602 | 727 | 619 | 612 | 511.61 | 535 | 653 | 1.267 | 0.342 | 0.00804816 |
| AT3G12640.1 | | protein_coding | RNA binding (RRM/RBD/ | 552 | 453 | 497 | 574 | 467 | 460 | 1160 | 1113 | 1272 | 1245 | 1240 | 1251 | 1059.18 | 1182 | 1245 | 1.267 | 0.342 | 0.00291701 |
| AT5G15390.1 | | protein_coding | tRNA/rRNA methyltransf | 199 | 178 | 143 | 203 | 190 | 211 | 418 | 437 | 366 | 440 | 504 | 574 | 665.95 | 407 | 506 | 1.268 | 0.342 | 0.00278118 |
| AT4G34555.1 | RPS25D | protein_coding | Ribosomal protein S25 fa | 347 | 233 | 231 | 353 | 299 | 276 | 729 | 572 | 591 | 766 | 794 | 751 | 744.23 | 631 | 770 | 1.266 | 0.341 | 0.00470426 |
| AT3G03960.1 | CCT8 | protein_coding | TCP-1/cpn60 chaperonin | 1798 | 1420 | 1292 | 1989 | 1710 | 1532 | 3780 | 3489 | 3308 | 4314 | 4540 | 4166 | 3690.95 | 3526 | 4340 | 1.266 | 0.341 | 0.00271534 |
| AT5G26160.1 | | protein_coding | | 427 | 290 | 326 | 389 | 285 | 274 | 898 | 712 | 835 | 844 | 757 | 745 | 563.13 | 815 | 782 | 1.265 | 0.34 | 0.00979362 |
| AT4G30600.1 | | protein_coding | signal recognition particle | 1557 | 1243 | 1383 | 1783 | 1254 | 1514 | 3273 | 3054 | 3541 | 3867 | 3329 | 4117 | 3355.75 | 3289 | 3771 | 1.266 | 0.34 | 0.00789695 |
| AT1G65440.1 | GTB1 | protein_coding | global transcription facto | 1163 | 962 | 897 | 1247 | 1100 | 806 | 2445 | 2363 | 2296 | 2705 | 2920 | 2192 | 2419.58 | 2368 | 2606 | 1.266 | 0.34 | 0.00617422 |
| AT4G37130.1 | NUP58 | protein_coding | hydroxyproline-rich glyco | 175 | 157 | 144 | 205 | 163 | 195 | 368 | 386 | 369 | 445 | 433 | 530 | 530.67 | 374 | 469 | 1.265 | 0.339 | 0.00462688 |
| AT2G41140.1 | CRK1 | protein_coding | CDPK-related kinase 1 | 666 | 521 | 516 | 796 | 628 | 544 | 1400 | 1280 | 1321 | 1727 | 1667 | 1479 | 1299.39 | 1334 | 1624 | 1.265 | 0.339 | 0.00443964 |
| AT1G80070.1 | SUS2 | protein_coding | Pre-mRNA-processing-spl | 7999 | 6767 | 6128 | 9616 | 7912 | 7439 | 16814 | 16625 | 15688 | 20858 | 21005 | 20230 | 12674.44 | 16376 | 20698 | 1.265 | 0.339 | 0.00403131 |
| AT5G37130.1 | | protein_coding | Protein prenylyltransfera | 919 | 733 | 720 | 806 | 777 | 741 | 1932 | 1801 | 1843 | 1748 | 2063 | 2015 | 1678.33 | 1859 | 1942 | 1.265 | 0.339 | 0.00334648 |
| AT1G10570.1 | ULP1C | protein_coding | Cysteine proteinases supe | 336 | 289 | 355 | 407 | 289 | 367 | 706 | 710 | 909 | 883 | 767 | 998 | 764.94 | 775 | 883 | 1.264 | 0.338 | 0.00983836 |
| AT3G10670.1 | ABCI6 | protein_coding | non-intrinsic ABC protein | 1503 | 1180 | 884 | 1451 | 1078 | 1014 | 3159 | 2899 | 2263 | 3147 | 2862 | 2758 | 3265.68 | 2774 | 2922 | 1.264 | 0.338 | 0.00946633 |
| AT3G21220.1 | MKK5 | protein_coding | MAP kinase kinase 5 | 168 | 130 | 117 | 192 | 129 | 150 | 353 | 319 | 300 | 416 | 342 | 408 | 501.61 | 324 | 389 | 1.264 | 0.338 | 0.00567664 |
| AT4G31850.1 | PGR3 | protein_coding | proton gradient regulatio | 1782 | 1357 | 1415 | 1518 | 1323 | 1161 | 3746 | 3334 | 3622 | 3293 | 3512 | 3157 | 2917.22 | 3567 | 3321 | 1.264 | 0.338 | 0.00461324 |
| AT1G16470.1 | PAB1 | protein_coding | proteasome subunit PAB | 969 | 674 | 736 | 1083 | 789 | 778 | 2037 | 1656 | 1884 | 2349 | 2095 | 2116 | 2152.52 | 1859 | 2187 | 1.264 | 0.338 | 0.00362869 |
| AT1G03560.1 | | protein_coding | Pentatricopeptide repeat | 137 | 114 | 120 | 148 | 126 | 90 | 288 | 280 | 307 | 321 | 335 | 245 | 329.6 | 292 | 300 | 1.263 | 0.337 | 0.00685527 |
| AT5G27760.1 | | protein_coding | Hypoxia-responsive famil | 252 | 189 | 198 | 395 | 269 | 244 | 530 | 464 | 507 | 857 | 714 | 664 | 669.49 | 500 | 745 | 1.263 | 0.337 | 0.00498402 |
| AT5G28740.1 | | protein_coding | Tetratricopeptide repeat | 1168 | 854 | 810 | 1232 | 1036 | 971 | 2455 | 2098 | 2074 | 2672 | 2750 | 2641 | 2018.46 | 2209 | 2688 | 1.263 | 0.337 | 0.00364538 |
| AT2G45000.1 | NUP62 | protein_coding | structural constituent of | 143 | 119 | 92 | 178 | 150 | 126 | 301 | 292 | 236 | 386 | 398 | 343 | 592.7 | 276 | 376 | 1.263 | 0.337 | 0.00329715 |
| AT1G44900.1 | MCM2 | protein_coding | minichromosome mainte | 973 | 753 | 639 | 1008 | 871 | 650 | 2045 | 1850 | 1636 | 2186 | 2312 | 1768 | 1562.8 | 1844 | 2089 | 1.262 | 0.336 | 0.00996266 |
| AT1G29990.1 | PFD6 | protein_coding | prefoldin 6 | 404 | 311 | 306 | 464 | 408 | 288 | 849 | 764 | 783 | 1006 | 1083 | 783 | 876.48 | 799 | 957 | 1.262 | 0.336 | 0.00951159 |
| AT3G10390.1 | FLD | protein_coding | Flavin containing amine c | 244 | 230 | 219 | 282 | 215 | 181 | 513 | 565 | 561 | 612 | 571 | 492 | 534.01 | 546 | 558 | 1.262 | 0.336 | 0.00255606 |
| AT5G40910.1 | | protein_coding | Disease resistance protei | 841 | 686 | 832 | 1183 | 941 | 872 | 1768 | 1685 | 2130 | 2566 | 2498 | 2371 | 1725.04 | 1861 | 2478 | 1.262 | 0.335 | 0.00970617 |
| AT1G53120.1 | | protein_coding | RNA-binding S4 domain- | 432 | 315 | 272 | 353 | 244 | 219 | 908 | 774 | 696 | 766 | 648 | 596 | 819.57 | 793 | 670 | 1.262 | 0.335 | 0.00691452 |
| AT5G53070.1 | | protein_coding | Ribosomal protein L9/RN | 248 | 216 | 173 | 241 | 200 | 183 | 521 | 531 | 443 | 523 | 531 | 498 | 534.98 | 498 | 517 | 1.262 | 0.335 | 0.00680456 |
| AT3G06750.1 | | protein_coding | hydroxyproline-rich glyco | 204 | 199 | 147 | 287 | 208 | 223 | 429 | 489 | 376 | 623 | 552 | 606 | 1070.42 | 431 | 594 | 1.262 | 0.335 | 0.00674159 |
| AT2G46250.1 | | protein_coding | myosin heavy chain-relat | 163 | 192 | 170 | 177 | 167 | 172 | 343 | 472 | 435 | 384 | 443 | 468 | 456.61 | 417 | 432 | 1.261 | 0.335 | 0.00672288 |
| AT5G51410.1 | | protein_coding | LUC7 N_terminus domair | 337 | 357 | 333 | 355 | 320 | 368 | 708 | 877 | 853 | 770 | 850 | 1001 | 806.24 | 813 | 874 | 1.26 | 0.334 | 0.00921221 |
| AT3G49000.1 | | protein_coding | RNA polymerase III subur | 348 | 246 | 251 | 416 | 299 | 251 | 732 | 604 | 643 | 902 | 794 | 683 | 651.02 | 660 | 793 | 1.261 | 0.334 | 0.00511752 |
| AT4G33760.1 | | protein_coding | tRNA synthetase class II ( | 849 | 735 | 613 | 879 | 675 | 627 | 1785 | 1806 | 1569 | 1907 | 1792 | 1705 | 1919.9 | 1720 | 1801 | 1.261 | 0.334 | 0.00408317 |
| AT3G03950.3 | ECT1 | protein_coding | evolutionarily conserved | 484 | 379 | 362 | 509 | 397 | 356 | 1017 | 931 | 927 | 1104 | 1054 | 968 | 968.51 | 958 | 1042 | 1.26 | 0.334 | 0.00382314 |
| AT2G17930.1 | | protein_coding | Phosphatidylinositol 3- ar | 3229 | 2902 | 2932 | 4485 | 4025 | 3693 | 6788 | 7130 | 7506 | 9728 | 10685 | 10043 | 5338.64 | 7141 | 10152 | 1.26 | 0.333 | 0.00344462 |
| AT2G04378.2 | | protein_coding | Beta-galactosidase relate | 154 | 127 | 119 | 192 | 202 | 169 | 324 | 312 | 305 | 416 | 536 | 460 | 850.99 | 314 | 471 | 1.259 | 0.333 | 0.00329792 |
| AT3G57990.1 | | protein_coding | | 456 | 433 | 361 | 496 | 441 | 471 | 959 | 1064 | 924 | 1076 | 1171 | 1281 | 1339.65 | 982 | 1176 | 1.259 | 0.332 | 0.00576739 |
| AT2G27680.1 | | protein_coding | NAD(P)-linked oxidoredu | 2641 | 2367 | 2111 | 2638 | 2244 | 2483 | 5552 | 5815 | 5404 | 5722 | 5957 | 6752 | 7316.82 | 5590 | 6144 | 1.259 | 0.332 | 0.00460105 |
| AT4G32610.1 | | protein_coding | copper ion binding | 428 | 442 | 390 | 480 | 400 | 432 | 900 | 1086 | 998 | 1041 | 1062 | 1175 | 1013.51 | 995 | 1093 | 1.259 | 0.332 | 0.00419476 |
| AT5G53060.1 | | protein_coding | RNA-binding KH domain- | 285 | 215 | 220 | 333 | 269 | 258 | 599 | 528 | 563 | 722 | 714 | 702 | 564.99 | 563 | 713 | 1.259 | 0.332 | 0.00160333 |
| AT3G47910.2 | | protein_coding | Ubiquitin carboxyl-termir | 2039 | 1505 | 1394 | 1940 | 1571 | 1408 | 4286 | 3698 | 3569 | 4208 | 4171 | 3829 | 2886.2 | 3851 | 4069 | 1.258 | 0.331 | 0.00719779 |
| AT3G19170.1 | PREP1 | protein_coding | presequence protease 1 | 8654 | 7148 | 6876 | 8056 | 6631 | 5891 | 18191 | 17561 | 17603 | 17474 | 17604 | 16020 | 16150.25 | 17785 | 17033 | 1.258 | 0.331 | 0.00709345 |
| AT2G41680.1 | NTRC | protein_coding | NADPH-dependent thiore | 4280 | 4112 | 3280 | 4350 | 3664 | 3586 | 8997 | 10102 | 8397 | 9436 | 9727 | 9752 | 10661.19 | 9165 | 9638 | 1.258 | 0.331 | 0.00624853 |
| AT1G67730.1 | KCR1 | protein_coding | beta-ketoacyl reductase 1 | 1327 | 1162 | 969 | 1600 | 1307 | 1166 | 2789 | 2855 | 2481 | 3471 | 3470 | 3171 | 3014.52 | 2708 | 3371 | 1.258 | 0.331 | 0.00528656 |
| AT1G73990.1 | SPPA | protein_coding | signal peptide peptidase | 1127 | 878 | 930 | 1089 | 932 | 924 | 2369 | 2157 | 2381 | 2362 | 2474 | 2513 | 2441.16 | 2302 | 2450 | 1.258 | 0.331 | 0.00295413 |
| AT1G08410.1 | LSG1-2 | protein_coding | P-loop containing nucleo | 498 | 369 | 327 | 470 | 419 | 346 | 1047 | 907 | 837 | 1019 | 1112 | 941 | 815.17 | 930 | 1024 | 1.257 | 0.33 | 0.00896953 |
| AT5G63490.1 | CBSCBSPB1 | protein_coding | CBS / octicosapeptide/Ph | 333 | 264 | 266 | 329 | 298 | 237 | 700 | 649 | 681 | 714 | 791 | 645 | 611.55 | 677 | 717 | 1.257 | 0.33 | 0.00744966 |
| AT4G37120.1 | SMP2 | protein_coding | Pre-mRNA splicing Prp18 | 1105 | 777 | 886 | 1235 | 819 | 835 | 2323 | 1909 | 2268 | 2679 | 2174 | 2271 | 2192.45 | 2167 | 2375 | 1.257 | 0.33 | 0.00725163 |
| AT2G47330.1 | RH24 | protein_coding | P-loop containing nucleo | 517 | 465 | 418 | 537 | 442 | 512 | 1087 | 1142 | 1070 | 1165 | 1173 | 1392 | 1074.11 | 1100 | 1243 | 1.257 | 0.33 | 0.00445877 |
| AT5G03340.1 | CDC48E | protein_coding | ATPase, AAA-type, CDC48 | 538 | 470 | 433 | 669 | 508 | 464 | 1131 | 1155 | 1109 | 1451 | 1349 | 1262 | 1106.67 | 1132 | 1354 | 1.257 | 0.33 | 0.00405625 |
| AT1G80040.3 | | protein_coding | | 577 | 557 | 476 | 741 | 570 | 628 | 1213 | 1368 | 1219 | 1607 | 1513 | 1708 | 1742.92 | 1267 | 1609 | 1.257 | 0.33 | 0.00370449 |
| AT4G32880.1 | ATHB-8 | protein_coding | homeobox gene 8 | 365 | 416 | 323 | 525 | 446 | 467 | 767 | 1022 | 827 | 1139 | 1184 | 1270 | 1056.85 | 872 | 1198 | 1.256 | 0.329 | 0.00989543 |
| AT5G49940.1 | NIFU2 | protein_coding | NIFU-like protein 2 | 1111 | 905 | 913 | 1327 | 993 | 981 | 2335 | 2223 | 2337 | 2878 | 2636 | 2668 | 2879.11 | 2298 | 2727 | 1.256 | 0.329 | 0.00305913 |
| AT2G35120.1 | GDH2 | protein_coding | Single hybrid motif super | 239 | 194 | 171 | 321 | 262 | 222 | 502 | 477 | 438 | 696 | 696 | 604 | 624.94 | 472 | 665 | 1.255 | 0.328 | 0.0053575 |
| AT1G11750.2 | CLPP6 | protein_coding | CLP protease proteolytic | 2254 | 1766 | 1865 | 2494 | 1848 | 1985 | 4738 | 4339 | 4775 | 5410 | 4906 | 5398 | 5709.41 | 4617 | 5238 | 1.255 | 0.328 | 0.00360537 |
| AT1G64520.1 | RPN12A | protein_coding | regulatory particle non-A | 1214 | 920 | 951 | 1361 | 1089 | 1072 | 2552 | 2260 | 2435 | 2952 | 2891 | 2915 | 2771.6 | 2416 | 2919 | 1.255 | 0.328 | 0.00339411 |
| AT5G02120.1 | HLIP | protein_coding | one helix protein | 1779 | 1699 | 1496 | 2378 | 1876 | 1871 | 3740 | 4174 | 3830 | 5158 | 4980 | 5088 | 5510.87 | 3915 | 5075 | 1.255 | 0.328 | 0.00270675 |
| AT2G42920.1 | PCMP-E75 | protein_coding | Pentatricopeptide repeat | 245 | 208 | 210 | 260 | 221 | 224 | 515 | 511 | 538 | 564 | 587 | 609 | 613.18 | 521 | 587 | 1.255 | 0.328 | 0.00210123 |
| AT5G59450.1 | SCL11 | protein_coding | GRAS family transcriptior | 300 | 299 | 306 | 318 | 285 | 258 | 631 | 735 | 783 | 690 | 757 | 702 | 558.69 | 716 | 716 | 1.255 | 0.327 | 0.0094449 |

| | | | | | | | | | | | | | | | | | | | | | |
|---|---|---|---|---|---|---|---|---|---|---|---|---|---|---|---|---|---|---|---|---|---|
| AT1G20100.1 | | protein_coding | | 696 | 491 | 576 | 672 | 466 | 465 | 1463 | 1206 | 1475 | 1458 | 1237 | 1265 | 1456.01 | 1381 | 1320 | 1.253 | 0.326 | 0.00935197 |
| AT5G42390.1 | | protein_coding | Insulinase (Peptidase fam | 6610 | 5159 | 4576 | 6804 | 5466 | 4665 | 13895 | 12675 | 11715 | 14759 | 14511 | 12686 | 12090.46 | 12762 | 13985 | 1.253 | 0.326 | 0.00889133 |
| AT3G61740.1 | ATX3 | protein_coding | SET domain protein 14 | 360 | 297 | 329 | 334 | 361 | 353 | 757 | 730 | 842 | 724 | 958 | 960 | 715.93 | 776 | 881 | 1.253 | 0.326 | 0.00791399 |
| AT5G62390.1 | BAG7 | protein_coding | BCL-2-associated athanog | 1410 | 946 | 1100 | 1465 | 1053 | 1132 | 2964 | 2324 | 2816 | 3178 | 2795 | 3078 | 3298.02 | 2701 | 3017 | 1.253 | 0.326 | 0.00774778 |
| AT1G43560.1 | Aty2 | protein_coding | thioredoxin Y2 | 979 | 989 | 749 | 1189 | 1011 | 900 | 2058 | 2430 | 1917 | 2579 | 2684 | 2447 | 3831.16 | 2135 | 2570 | 1.254 | 0.326 | 0.0060856 |
| AT3G04340.1 | emb2458 | protein_coding | FtsH extracellular proteas | 2908 | 2141 | 2066 | 2707 | 2222 | 1902 | 6113 | 5260 | 5289 | 5872 | 5899 | 5172 | 5272.29 | 5554 | 5648 | 1.254 | 0.326 | 0.00485629 |
| AT1G44780.1 | | protein_coding | | 163 | 141 | 136 | 144 | 119 | 95 | 343 | 346 | 348 | 312 | 316 | 258 | 320.84 | 346 | 295 | 1.252 | 0.325 | 0.0074698 |
| AT1G10030.1 | ERG28 | protein_coding | homolog of yeast ergoste | 216 | 164 | 160 | 198 | 187 | 163 | 454 | 403 | 410 | 429 | 496 | 443 | 486.32 | 422 | 456 | 1.253 | 0.325 | 0.00372869 |
| AT1G66330.1 | | protein_coding | senescence-associated fa | 1042 | 896 | 910 | 1567 | 1053 | 1163 | 2190 | 2201 | 2330 | 3399 | 2795 | 3163 | 2400.53 | 2240 | 3119 | 1.252 | 0.324 | 0.00775462 |
| AT2G40600.1 | | protein_coding | appr-1-p processing enzy | 174 | 187 | 149 | 255 | 231 | 203 | 366 | 459 | 381 | 553 | 613 | 552 | 736.17 | 402 | 573 | 1.252 | 0.324 | 0.00272451 |
| AT3G03300.1 | DCL2 | protein_coding | dicer-like 2 | 502 | 410 | 449 | 550 | 484 | 438 | 1055 | 1007 | 1149 | 1193 | 1285 | 1191 | 965.9 | 1070 | 1223 | 1.251 | 0.323 | 0.0084294 |
| AT3G58510.1 | RH11 | protein_coding | DEA(D/H)-box RNA helica | 282 | 260 | 225 | 302 | 309 | 245 | 593 | 639 | 576 | 655 | 820 | 666 | 1439.42 | 603 | 714 | 1.251 | 0.323 | 0.00778022 |
| AT4G15802.1 | HSBP | protein_coding | heat shock factor binding | 888 | 655 | 745 | 1133 | 844 | 895 | 1867 | 1609 | 1907 | 2458 | 2241 | 2434 | 2506.89 | 1794 | 2378 | 1.251 | 0.323 | 0.00491604 |
| AT1G53750.1 | RPT1A | protein_coding | regulatory particle triple- | 2520 | 1944 | 1839 | 3199 | 2320 | 2212 | 5297 | 4776 | 4708 | 6939 | 6159 | 6015 | 5258.57 | 4927 | 6371 | 1.251 | 0.323 | 0.00423519 |
| AT1G04810.1 | RPN2B | protein_coding | 26S proteasome regulato | 938 | 852 | 743 | 1056 | 955 | 969 | 1972 | 2093 | 1902 | 2291 | 2535 | 2635 | 2105.75 | 1989 | 2487 | 1.251 | 0.323 | 0.00346093 |
| AT2G20420.1 | | protein_coding | ATP citrate lyase (ACL) fa | 2664 | 1881 | 1721 | 3202 | 2440 | 2240 | 5600 | 4621 | 4406 | 6945 | 6478 | 6092 | 5246.53 | 4876 | 6505 | 1.25 | 0.322 | 0.00836658 |
| AT5G53850.2 | | protein_coding | haloacid dehalogenase-lil | 1681 | 1241 | 1326 | 1947 | 1578 | 1662 | 3534 | 3049 | 3395 | 4223 | 4189 | 4520 | 3941.75 | 3326 | 4311 | 1.25 | 0.322 | 0.00471872 |
| AT1G50380.1 | | protein_coding | Prolyl oligopeptidase fam | 448 | 334 | 335 | 417 | 338 | 328 | 942 | 821 | 858 | 905 | 897 | 892 | 854.47 | 874 | 898 | 1.25 | 0.322 | 0.00436214 |
| AT5G43130.2 | TAF4B | protein_coding | TBP-associated factor 4 | 728 | 632 | 662 | 786 | 671 | 681 | 1530 | 1553 | 1695 | 1705 | 1781 | 1852 | 1582.83 | 1593 | 1779 | 1.249 | 0.321 | 0.00624029 |
| AT5G49880.1 | MAD1 | protein_coding | mitotic checkpoint family | 452 | 387 | 431 | 493 | 413 | 427 | 950 | 951 | 1103 | 1069 | 1096 | 1161 | 892.91 | 1001 | 1109 | 1.249 | 0.321 | 0.00494445 |
| AT5G07240.1 | IQD24 | protein_coding | IQ-domain 24 | 591 | 564 | 423 | 661 | 566 | 467 | 1242 | 1386 | 1083 | 1434 | 1503 | 1270 | 1364.18 | 1237 | 1402 | 1.248 | 0.32 | 0.00989326 |
| AT4G20910.1 | HEN1 | protein_coding | double-stranded RNA bin | 309 | 252 | 225 | 337 | 308 | 215 | 650 | 619 | 576 | 731 | 818 | 585 | 521.41 | 615 | 711 | 1.248 | 0.32 | 0.00969669 |
| AT5G03080.1 | LPPG | protein_coding | Phosphatidic acid phosph | 204 | 159 | 143 | 204 | 152 | 196 | 429 | 391 | 366 | 442 | 404 | 533 | 514.28 | 395 | 460 | 1.248 | 0.32 | 0.00796095 |
| AT3G53280.1 | CYP71B5 | protein_coding | cytochrome p450 71b5 | 375 | 331 | 270 | 389 | 251 | 276 | 788 | 813 | 691 | 844 | 666 | 751 | 670.64 | 764 | 754 | 1.249 | 0.32 | 0.00660196 |
| AT5G63040.1 | | protein_coding | | 226 | 178 | 136 | 280 | 240 | 183 | 475 | 437 | 348 | 607 | 637 | 498 | 651.65 | 420 | 581 | 1.249 | 0.32 | 0.00652292 |
| AT4G17330.1 | ATG2484-1 | protein_coding | G2484-1 protein | 3246 | 2757 | 2594 | 3883 | 3266 | 2637 | 6823 | 6773 | 6641 | 8423 | 8670 | 7171 | 5808.23 | 6746 | 8088 | 1.248 | 0.32 | 0.00558539 |
| AT3G20780.1 | TOP6B | protein_coding | topoisomerase 6 subunit | 246 | 232 | 238 | 239 | 221 | 173 | 517 | 570 | 609 | 518 | 587 | 470 | 495.34 | 565 | 525 | 1.248 | 0.319 | 0.00846172 |
| AT4G31180.1 | | protein_coding | Class II aminoacyl-tRNA a | 1002 | 911 | 733 | 1106 | 973 | 961 | 2106 | 2238 | 1877 | 2399 | 2583 | 2613 | 2576.32 | 2074 | 2532 | 1.248 | 0.319 | 0.0073509 |
| AT3G53110.1 | RH38 | protein_coding | P-loop containing nucleo | 1237 | 1007 | 1007 | 1457 | 1061 | 1094 | 2600 | 2474 | 2578 | 3160 | 2817 | 2975 | 2613.98 | 2551 | 2984 | 1.248 | 0.319 | 0.004002 |
| AT2G33250.1 | | protein_coding | | 242 | 196 | 200 | 252 | 193 | 200 | 509 | 482 | 512 | 547 | 512 | 544 | 609.81 | 501 | 534 | 1.247 | 0.319 | 0.00187506 |
| AT1G11440.1 | | protein_coding | | 198 | 191 | 152 | 225 | 187 | 151 | 416 | 469 | 389 | 488 | 496 | 411 | 426.96 | 425 | 465 | 1.247 | 0.318 | 0.00877831 |
| AT4G02570.1 | CUL1 | protein_coding | cullin 1 | 2983 | 2262 | 2384 | 3614 | 2867 | 2657 | 6270 | 5557 | 6103 | 7839 | 7611 | 7226 | 6006.13 | 5977 | 7559 | 1.247 | 0.318 | 0.00456909 |
| AT3G54760.1 | | protein_coding | dentin sialophosphoprote | 1336 | 1059 | 1127 | 1291 | 1165 | 987 | 2808 | 2602 | 2885 | 2800 | 3093 | 2684 | 2407.42 | 2765 | 2859 | 1.246 | 0.317 | 0.00615056 |
| AT2G42247.1 | | other_rna | other RNA | 117 | 111 | 93 | 134 | 98 | 98 | 246 | 273 | 238 | 291 | 260 | 267 | 361.95 | 252 | 273 | 1.246 | 0.317 | 0.00535116 |
| AT1G55480.1 | ZKT | protein_coding | protein containing PDZ d | 5016 | 4154 | 4301 | 4766 | 4185 | 3900 | 10544 | 10206 | 11011 | 10338 | 11110 | 10606 | 11337.5 | 10587 | 10685 | 1.245 | 0.316 | 0.00654363 |
| AT3G56120.1 | | protein_coding | S-adenosyl-L-methionine- | 411 | 313 | 302 | 442 | 370 | 293 | 864 | 769 | 773 | 959 | 982 | 797 | 743.59 | 802 | 913 | 1.245 | 0.316 | 0.00481004 |
| AT1G04190.1 | TPR3 | protein_coding | Tetratricopeptide repeat | 472 | 376 | 347 | 529 | 417 | 343 | 992 | 924 | 888 | 1147 | 1107 | 933 | 957.95 | 935 | 1062 | 1.244 | 0.315 | 0.0077897 |
| AT5G18620.2 | CHR17 | protein_coding | chromatin remodeling fa | 1283 | 1085 | 1028 | 1394 | 1236 | 1172 | 2697 | 2666 | 2632 | 3024 | 3281 | 3187 | 2258.3 | 2665 | 3164 | 1.244 | 0.315 | 0.00492256 |
| AT2G47580.1 | U1A | protein_coding | spliceosomal protein U1A | 521 | 424 | 363 | 639 | 482 | 427 | 1095 | 1042 | 929 | 1386 | 1280 | 1161 | 1192.49 | 1022 | 1276 | 1.244 | 0.314 | 0.00858476 |
| AT3G59990.1 | MAP2B | protein_coding | methionine aminopeptid | 1189 | 985 | 991 | 1427 | 1136 | 1203 | 2499 | 2420 | 2537 | 3095 | 3016 | 3271 | 2705.73 | 2485 | 3127 | 1.243 | 0.314 | 0.00439364 |
| AT3G19650.1 | | protein_coding | cyclin-related | 225 | 196 | 156 | 196 | 167 | 170 | 473 | 482 | 399 | 425 | 443 | 462 | 499.5 | 451 | 443 | 1.242 | 0.313 | 0.00705894 |
| AT5G58230.1 | MSI1 | protein_coding | Transducin/WD40 repeat | 598 | 464 | 441 | 623 | 503 | 504 | 1257 | 1140 | 1129 | 1351 | 1335 | 1371 | 1242.04 | 1175 | 1352 | 1.242 | 0.313 | 0.00639081 |
| AT1G32500.1 | ABCI7 | protein_coding | non-intrinsic ABC protein | 2528 | 2021 | 1843 | 3180 | 2307 | 2442 | 5314 | 4965 | 4718 | 6898 | 6125 | 6641 | 5419.86 | 4999 | 6555 | 1.243 | 0.313 | 0.004978 |
| AT4G33510.1 | DHS2 | protein_coding | 3-deoxy-d-arabino-heptu | 2466 | 2461 | 1862 | 2728 | 2352 | 2306 | 5184 | 6046 | 4767 | 5917 | 6244 | 6271 | 6538.38 | 5332 | 6144 | 1.241 | 0.312 | 0.00859971 |
| AT1G33520.1 | MOS2 | protein_coding | D111/G-patch domain-co | 140 | 110 | 118 | 137 | 114 | 124 | 294 | 270 | 302 | 297 | 303 | 337 | 529.95 | 289 | 312 | 1.241 | 0.311 | 0.00375079 |
| AT4G04320.1 | | protein_coding | malonyl-CoA decarboxyla | 398 | 367 | 358 | 447 | 349 | 331 | 837 | 902 | 917 | 970 | 927 | 900 | 874.33 | 885 | 932 | 1.239 | 0.309 | 0.00552387 |
| AT1G02330.1 | | protein_coding | | 326 | 243 | 278 | 280 | 243 | 238 | 685 | 597 | 712 | 607 | 645 | 647 | 687.1 | 665 | 633 | 1.239 | 0.309 | 0.00344191 |
| AT1G01040.2 | DCL1 | protein_coding | dicer-like 1 | 1031 | 893 | 920 | 1179 | 1054 | 1000 | 2167 | 2194 | 2355 | 2557 | 2798 | 2719 | 1990.94 | 2239 | 2691 | 1.238 | 0.308 | 0.00511134 |
| AT5G44000.1 | | protein_coding | Glutathione S-transferase | 335 | 325 | 294 | 384 | 308 | 367 | 704 | 798 | 753 | 833 | 818 | 998 | 1046.46 | 752 | 883 | 1.237 | 0.307 | 0.00837863 |
| AT1G43710.1 | SDC | protein_coding | Pyridoxal phosphate (PLP | 1488 | 1173 | 1120 | 1532 | 1147 | 1073 | 3128 | 2882 | 2867 | 3323 | 3045 | 2918 | 3410.87 | 2959 | 3095 | 1.236 | 0.306 | 0.00819794 |
| AT1G66680.1 | AR401 | protein_coding | S-adenosyl-L-methionine- | 321 | 275 | 293 | 419 | 325 | 368 | 675 | 676 | 750 | 909 | 863 | 1001 | 804.62 | 700 | 924 | 1.236 | 0.305 | 0.00497188 |
| AT5G14250.1 | CSN3 | protein_coding | Proteasome component ( | 557 | 384 | 399 | 595 | 467 | 472 | 1171 | 943 | 1021 | 1291 | 1240 | 1284 | 1124.69 | 1045 | 1272 | 1.234 | 0.304 | 0.00816129 |
| AT4G27000.1 | RBP45C | protein_coding | RNA-binding (RRM/RBD/ | 1442 | 1283 | 1241 | 1670 | 1262 | 1493 | 3031 | 3152 | 3177 | 3622 | 3350 | 4060 | 3194.73 | 3120 | 3677 | 1.233 | 0.303 | 0.00941015 |
| AT4G00231.1 | MEE50 | protein_coding | ARM repeat superfamily | 209 | 177 | 200 | 228 | 179 | 177 | 439 | 435 | 512 | 495 | 475 | 481 | 414.35 | 462 | 484 | 1.233 | 0.302 | 0.00763837 |
| AT4G22350.2 | | protein_coding | Ubiquitin C-terminal hyd | 306 | 241 | 224 | 278 | 233 | 234 | 643 | 592 | 573 | 603 | 619 | 636 | 576.02 | 603 | 619 | 1.233 | 0.302 | 0.00451353 |
| AT4G08960.1 | | protein_coding | phosphotyrosyl phosphat | 206 | 167 | 180 | 265 | 199 | 175 | 433 | 410 | 461 | 575 | 528 | 476 | 551.75 | 435 | 526 | 1.231 | 0.3 | 0.00851832 |
| AT4G31040.1 | | protein_coding | CemA-like proton extrusi | 1329 | 1109 | 1036 | 1281 | 1001 | 906 | 2794 | 2725 | 2652 | 2779 | 2657 | 2464 | 2876.38 | 2724 | 2633 | 1.23 | 0.299 | 0.00946599 |
| AT4G08170.2 | ITPK3 | protein_coding | Inositol 1,3,4-trisphospha | 223 | 185 | 209 | 250 | 249 | 199 | 469 | 455 | 535 | 542 | 661 | 541 | 565.68 | 486 | 581 | 1.229 | 0.298 | 0.00949145 |
| AT1G01350.1 | | protein_coding | Zinc finger (CCCH-type/C | 315 | 245 | 249 | 267 | 215 | 236 | 662 | 602 | 637 | 579 | 571 | 642 | 579 | 634 | 597 | 1.228 | 0.297 | 0.00479526 |
| AT3G56840.1 | | protein_coding | FAD-dependent oxidored | 228 | 179 | 179 | 268 | 198 | 173 | 479 | 440 | 458 | 581 | 526 | 470 | 499.92 | 459 | 526 | 1.227 | 0.295 | 0.00689837 |
| AT5G39080.1 | | protein_coding | HXXXD-type acyl-transfer | 358 | 312 | 254 | 407 | 313 | 318 | 753 | 767 | 650 | 883 | 831 | 865 | 1018.46 | 723 | 860 | 1.225 | 0.293 | 0.0081231 |
| AT1G61890.1 | | protein_coding | MATE efflux family prote | 208 | 206 | 172 | 312 | 242 | 252 | 437 | 506 | 440 | 677 | 642 | 685 | 558.92 | 461 | 668 | 1.225 | 0.293 | 0.00691054 |
| AT1G74170.1 | AtRLP13 | protein_coding | receptor like protein 13 | 377 | 276 | 295 | 364 | 280 | 240 | 792 | 678 | 755 | 790 | 743 | 653 | 658.4 | 742 | 729 | 1.224 | 0.291 | 0.00954387 |
| AT2G40650.1 | PRP38 | protein_coding | PRP38 family protein | 352 | 278 | 255 | 349 | 298 | 284 | 740 | 683 | 653 | 757 | 791 | 772 | 799.66 | 692 | 773 | 1.223 | 0.291 | 0.00723568 |
| AT2G20270.2 | GRXS12 | protein_coding | Thioredoxin superfamily | 878 | 759 | 633 | 1010 | 804 | 774 | 1846 | 1865 | 1621 | 2191 | 2134 | 2105 | 2073.26 | 1777 | 2143 | 1.222 | 0.289 | 0.00935716 |
| AT5G13060.1 | ABAP1 | protein_coding | ARMADILLO BTB protein | 306 | 252 | 217 | 252 | 235 | 213 | 643 | 619 | 556 | 547 | 624 | 579 | 501.86 | 606 | 583 | 1.222 | 0.289 | 0.008646 |
| AT2G30575.1 | GAUT5 | protein_coding | los glycosyltransferase 5 | 362 | 321 | 312 | 392 | 278 | 294 | 761 | 789 | 799 | 850 | 738 | 800 | 723.33 | 783 | 796 | 1.216 | 0.283 | 0.00559436 |
| AT5G20930.1 | TOUSLED | protein_coding | Protein kinase superfamil | 286 | 245 | 241 | 324 | 235 | 251 | 601 | 602 | 617 | 703 | 624 | 683 | 560.94 | 607 | 670 | 1.214 | 0.28 | 0.00883337 |
| AT5G20060.2 | | protein_coding | alpha/beta-Hydrolases su | 765 | 609 | 585 | 869 | 710 | 689 | 1608 | 1496 | 1498 | 1885 | 1885 | 1874 | 2114.43 | 1534 | 1881 | 1.213 | 0.279 | 0.00990255 |
| AT3G10220.1 | TFCB | protein_coding | tubulin folding cofactor B | 258 | 189 | 209 | 255 | 201 | 200 | 542 | 464 | 535 | 553 | 534 | 544 | 579.41 | 514 | 544 | 1.208 | 0.273 | 0.00827214 |
| AT5G43500.1 | ARP9 | protein_coding | actin-related protein 9 | 369 | 312 | 330 | 295 | 246 | 234 | 776 | 767 | 845 | 640 | 653 | 636 | 713.13 | 796 | 643 | 0.83 | -0.269 | 0.00771279 |
| AT3G18990.1 | VRN1 | protein_coding | AP2/B3-like transcription | 401 | 323 | 334 | 288 | 252 | 265 | 843 | 794 | 855 | 625 | 669 | 721 | 768.41 | 831 | 672 | 0.825 | -0.278 | 0.00952451 |
| AT5G55530.1 | | protein_coding | Calcium-dependent lipid- | 457 | 412 | 396 | 468 | 376 | 363 | 961 | 1012 | 1014 | 1015 | 998 | 987 | 868.93 | 996 | 1000 | 0.823 | -0.281 | 0.00945069 |
| AT3G46440.1 | UXS5 | protein_coding | UDP-XYL synthase 5 | 310 | 294 | 296 | 267 | 227 | 224 | 652 | 722 | 758 | 579 | 603 | 609 | 690 | 711 | 597 | 0.822 | -0.283 | 0.00799477 |
| AT4G35950.1 | ARAC6 | protein_coding | RAC-like 6 | 445 | 395 | 343 | 335 | 270 | 234 | 935 | 970 | 878 | 727 | 717 | 636 | 897.14 | 928 | 693 | 0.82 | -0.286 | 0.00873529 |
| AT2G38840.1 | | protein_coding | Guanylate-binding family | 361 | 310 | 321 | 321 | 247 | 262 | 759 | 762 | 822 | 696 | 656 | 712 | 680.85 | 781 | 688 | 0.82 | -0.286 | 0.00843873 |
| AT2G30270.1 | | protein_coding | Protein of unknown funct | 193 | 156 | 143 | 168 | 122 | 118 | 406 | 383 | 366 | 364 | 324 | 321 | 443.24 | 385 | 336 | 0.82 | -0.286 | 0.00614969 |
| AT1G68100.1 | IAR1 | protein_coding | ZIP metal ion transporter | 328 | 304 | 298 | 313 | 228 | 251 | 689 | 747 | 763 | 679 | 605 | 683 | 652.9 | 733 | 656 | 0.819 | -0.287 | 0.00848357 |
| AT1G32530.1 | MIP1 | protein_coding | RING/U-box superfamily | 291 | 286 | 271 | 248 | 173 | 179 | 612 | 703 | 694 | 538 | 459 | 487 | 603.72 | 670 | 495 | 0.818 | -0.289 | 0.00943917 |
| AT5G39830.1 | DEGP8 | protein_coding | Trypsin family protein wi | 1269 | 1218 | 1050 | 1007 | 836 | 757 | 2668 | 2992 | 2688 | 2184 | 2219 | 2059 | 2805.53 | 2783 | 2154 | 0.818 | -0.29 | 0.00995651 |
| AT5G46420.1 | | protein_coding | 16S rRNA processing prot | 2957 | 2461 | 2227 | 1762 | 1513 | 1485 | 6216 | 6046 | 5701 | 3822 | 4017 | 4038 | 4528.22 | 5988 | 3959 | 0.817 | -0.291 | 0.00747128 |
| AT3G58690.1 | | protein_coding | Protein kinase superfamil | 290 | 217 | 198 | 180 | 147 | 143 | 610 | 533 | 507 | 390 | 390 | 389 | 436.57 | 550 | 390 | 0.815 | -0.295 | 0.0074032 |
| AT5G52960.1 | | protein_coding | | 1240 | 955 | 897 | 850 | 609 | 610 | 2607 | 2346 | 2296 | 1844 | 1617 | 1659 | 2264.79 | 2416 | 1707 | 0.814 | -0.297 | 0.00939414 |
| AT2G32000.1 | | protein_coding | DNA topoisomerase, type | 364 | 344 | 331 | 297 | 262 | 258 | 765 | 845 | 847 | 644 | 696 | 702 | 728.8 | 819 | 681 | 0.814 | -0.297 | 0.00773516 |
| AT2G05250.1 | | protein_coding | DNAJ heat shock N-termi | 331 | 271 | 271 | 247 | 204 | 232 | 696 | 666 | 694 | 536 | 542 | 631 | 533.65 | 685 | 570 | 0.814 | -0.297 | 0.00653795 |
| AT2G28390.1 | | protein_coding | SAND family protein | 584 | 517 | 505 | 486 | 437 | 406 | 1228 | 1270 | 1293 | 1054 | 1160 | 1104 | 1094.38 | 1264 | 1106 | 0.813 | -0.299 | 0.00852619 |
| AT2G25480.1 | | protein_coding | TPX2 (targeting protein f | 409 | 334 | 303 | 219 | 207 | 180 | 860 | 821 | 776 | 475 | 550 | 489 | 690.96 | 819 | 505 | 0.813 | -0.299 | 0.00372678 |
| AT4G21770.1 | | protein_coding | Pseudouridine synthase f | 458 | 361 | 333 | 269 | 221 | 225 | 963 | 887 | 853 | 583 | 587 | 612 | 812.62 | 901 | 594 | 0.812 | -0.3 | 0.00459707 |
| AT4G01940.1 | NIFU1 | protein_coding | NFU domain protein 1 | 1139 | 933 | 957 | 906 | 721 | 705 | 2394 | 2292 | 2450 | 1965 | 1914 | 1917 | 2338.41 | 2379 | 1932 | 0.812 | -0.301 | 0.00645921 |
| AT4G21790.1 | TOM1 | protein_coding | tobamovirus multiplicatio | 1027 | 842 | 850 | 884 | 721 | 659 | 2159 | 2069 | 2176 | 1917 | 1914 | 1792 | 2024.32 | 2135 | 1874 | 0.812 | -0.301 | 0.00585533 |
| AT3G63150.1 | MIRO2 | protein_coding | MIRO-related GTP-ase 2 | 499 | 359 | 366 | 304 | 278 | 289 | 1049 | 882 | 937 | 659 | 738 | 786 | 910.69 | 956 | 728 | 0.811 | -0.303 | 0.00997037 |
| AT1G61430.1 | | protein_coding | S-locus lectin protein kin | 172 | 139 | 124 | 103 | 99 | 94 | 362 | 341 | 317 | 223 | 263 | 256 | 309.08 | 340 | 247 | 0.81 | -0.303 | 0.00968087 |
| AT5G65950.1 | | protein_coding | | 1042 | 848 | 896 | 883 | 727 | 746 | 2190 | 2083 | 2294 | 1915 | 1930 | 2029 | 1853.76 | 2189 | 1958 | 0.81 | -0.303 | 0.00792908 |
| AT1G32550.2 | FdC1 | protein_coding | 2Fe-2S ferredoxin-like su | 745 | 708 | 657 | 572 | 538 | 549 | 1566 | 1739 | 1682 | 1241 | 1428 | 1493 | 1771.66 | 1662 | 1387 | 0.81 | -0.304 | 0.00904923 |
| AT2G42590.3 | GRF9 | protein_coding | general regulatory factor | 3786 | 3368 | 3323 | 2978 | 2572 | 2476 | 7958 | 8275 | 8507 | 6460 | 6828 | 6733 | 7930.94 | 8247 | 6674 | 0.81 | -0.304 | 0.00678467 |
| AT2G24650.1 | REM13 | protein_coding | DNA binding;DNA bindin | 534 | 397 | 471 | 348 | 305 | 283 | 1123 | 975 | 1206 | 755 | 810 | 770 | 729.1 | 1101 | 778 | 0.81 | -0.304 | 0.00671075 |
| AT5G10950.1 | | protein_coding | Tudor/PWWP/MBT super | 275 | 208 | 233 | 186 | 148 | 151 | 578 | 511 | 596 | 403 | 393 | 411 | 485.99 | 562 | 402 | 0.809 | -0.305 | 0.00845778 |
| AT5G08710.1 | | protein_coding | Regulator of chromosom | 139 | 117 | 106 | 93 | 76 | 88 | 292 | 287 | 271 | 202 | 202 | 239 | 364.5 | 283 | 214 | 0.809 | -0.306 | 0.00814928 |
| AT5G03520.1 | RABE1D | protein_coding | RAB GTPase homolog 8C | 1174 | 1002 | 1045 | 1014 | 667 | 713 | 2468 | 2462 | 2675 | 2199 | 1771 | 1939 | 2326.93 | 2535 | 1970 | 0.808 | -0.307 | 0.00959982 |
| AT3G51140.1 | | protein_coding | Protein of unknown funct | 1057 | 876 | 750 | 858 | 619 | 687 | 2222 | 2152 | 1920 | 1861 | 1643 | 1868 | 2261.51 | 2098 | 1791 | 0.808 | -0.307 | 0.00717952 |
| AT5G23330.1 | RIBF1 | protein_coding | Nucleotidylyl transferase | 154 | 135 | 120 | 104 | 95 | 93 | 324 | 332 | 307 | 226 | 252 | 253 | 318.02 | 321 | 244 | 0.808 | -0.307 | 0.00704373 |
| AT1G15570.1 | CYCA2-3 | protein_coding | CYCLIN A2;3 | 242 | 242 | 210 | 179 | 149 | 126 | 509 | 595 | 538 | 388 | 396 | 343 | 446.78 | 547 | 376 | 0.808 | -0.308 | 0.00691394 |
| AT4G35510.1 | | protein_coding | | 137 | 129 | 117 | 112 | 86 | 80 | 288 | 317 | 300 | 243 | 228 | 218 | 313.11 | 302 | 230 | 0.808 | -0.308 | 0.00668914 |
| AT5G09740.1 | HAM2 | protein_coding | histone acetyltransferase | 219 | 187 | 219 | 167 | 135 | 129 | 460 | 459 | 561 | 362 | 358 | 351 | 399.49 | 493 | 357 | 0.807 | -0.31 | 0.00766278 |
| AT2G37120.1 | S1FA2 | protein_coding | S1FA-like DNA-binding pr | 206 | 186 | 225 | 168 | 137 | 148 | 433 | 457 | 576 | 364 | 364 | 402 | 425.03 | 489 | 377 | 0.806 | -0.311 | 0.00923673 |
| AT1G75350.1 | RPL31 | protein_coding | Ribosomal protein L31 | 5371 | 4848 | 4869 | 4060 | 3429 | 3561 | 11290 | 11911 | 12465 | 8807 | 9103 | 9684 | 11734.01 | 11889 | 9198 | 0.806 | -0.311 | 0.00858084 |
| AT1G04910.1 | | protein_coding | O-fucosyltransferase fam | 459 | 410 | 446 | 345 | 273 | 314 | 965 | 1007 | 1142 | 748 | 725 | 854 | 849.9 | 1038 | 776 | 0.806 | -0.311 | 0.00670031 |
| AT3G28710.1 | VHA-D1 | protein_coding | ATPase, V0/A0 complex, | 1920 | 1642 | 1787 | 1715 | 1466 | 1466 | 4036 | 4034 | 4575 | 3720 | 3892 | 3987 | 3991.01 | 4215 | 3866 | 0.806 | -0.311 | 0.00564329 |
| AT5G07360.1 | | protein_coding | Amidase family protein | 199 | 152 | 155 | 143 | 92 | 102 | 418 | 373 | 397 | 310 | 244 | 277 | 323.06 | 396 | 277 | 0.806 | -0.312 | 0.00930613 |
| AT5G10540.1 | CYOP | protein_coding | Zincin-like metalloprotea | 1473 | 1335 | 1321 | 1250 | 930 | 1079 | 3096 | 3280 | 3382 | 2711 | 2469 | 2934 | 2865.55 | 3253 | 2705 | 0.806 | -0.312 | 0.00684261 |
| AT3G14150.1 | GLO3 | protein_coding | Aldolase-type TIM barrel | 349 | 300 | 295 | 222 | 230 | 192 | 734 | 737 | 755 | 482 | 611 | 522 | 661.49 | 742 | 538 | 0.806 | -0.312 | 0.0063375 |
| AT5G47490.1 | SEC16B | protein_coding | RGPR-related | 668 | 570 | 559 | 524 | 486 | 445 | 1404 | 1400 | 1431 | 1137 | 1290 | 1210 | 1101.82 | 1412 | 1212 | 0.806 | -0.312 | 0.00632117 |
| AT2G05755.1 | | protein_coding | Nodulin MtN21 /EamA-lil | 112 | 92 | 98 | 99 | 77 | 72 | 235 | 226 | 251 | 215 | 204 | 196 | 298.25 | 237 | 205 | 0.805 | -0.313 | 0.0094396 |
| AT2G33600.1 | | protein_coding | NAD(P)-binding Rossman | 370 | 291 | 226 | 219 | 197 | 159 | 778 | 715 | 579 | 475 | 523 | 432 | 636.01 | 691 | 477 | 0.805 | -0.313 | 0.00918414 |
| AT3G49310.1 | | protein_coding | Major facilitator superfa | 364 | 290 | 270 | 336 | 264 | 266 | 765 | 712 | 691 | 729 | 701 | 723 | 623.91 | 723 | 718 | 0.805 | -0.313 | 0.0067902 |
| AT5G64020.1 | TBL14 | protein_coding | TRICHOME BIREFRINGEN | 271 | 229 | 222 | 208 | 194 | 215 | 570 | 563 | 568 | 451 | 515 | 585 | 506.61 | 567 | 517 | 0.805 | -0.313 | 0.00553554 |
| AT2G43360.1 | BIO2 | protein_coding | Radical SAM superfamily | 1123 | 915 | 984 | 881 | 737 | 693 | 2361 | 2248 | 2519 | 1911 | 1957 | 1885 | 2189.54 | 2376 | 1918 | 0.805 | -0.313 | 0.00448099 |
| AT1G66430.1 | | protein_coding | pfkB-like carbohydrate ki | 1137 | 960 | 915 | 945 | 817 | 812 | 2390 | 2359 | 2342 | 2050 | 2169 | 2208 | 2513.2 | 2364 | 2142 | 0.804 | -0.314 | 0.00516923 |
| AT2G16930.2 | | protein_coding | Ribosomal protein L27 fa | 315 | 243 | 271 | 277 | 211 | 233 | 662 | 597 | 694 | 601 | 560 | 634 | 775.84 | 651 | 598 | 0.805 | -0.314 | 0.00369758 |
| AT4G38360.2 | LAZ1 | protein_coding | Protein of unknown funct | 304 | 253 | 233 | 299 | 207 | 201 | 639 | 622 | 596 | 649 | 550 | 547 | 723.57 | 619 | 582 | 0.805 | -0.314 | 0.00359866 |
| AT4G11630.1 | | protein_coding | Ribosomal protein L19 fa | 291 | 201 | 189 | 219 | 162 | 152 | 612 | 494 | 484 | 475 | 430 | 413 | 538.04 | 530 | 439 | 0.803 | -0.316 | 0.00665789 |
| AT4G27650.1 | PEL1 | protein_coding | Eukaryotic release factor | 389 | 291 | 290 | 280 | 194 | 238 | 818 | 715 | 742 | 607 | 515 | 647 | 632.42 | 758 | 590 | 0.804 | -0.316 | 0.00492716 |
| AT5G02040.1 | PRA1A1 | protein_coding | prenylated RAB acceptor | 175 | 147 | 162 | 158 | 140 | 142 | 368 | 361 | 415 | 343 | 372 | 386 | 390.13 | 381 | 367 | 0.803 | -0.317 | 0.00783354 |
| AT4G32330.1 | | protein_coding | TPX2 (targeting protein f | 1836 | 1536 | 1654 | 1445 | 1314 | 1117 | 3859 | 3774 | 4234 | 3134 | 3488 | 3038 | 3418.78 | 3956 | 3220 | 0.802 | -0.318 | 0.005451 |
| AT2G37940.1 | IPCS2 | protein_coding | Arabidopsis Inositol phos | 667 | 626 | 673 | 621 | 504 | 536 | 1402 | 1538 | 1723 | 1347 | 1338 | 1458 | 1459.19 | 1554 | 1381 | 0.801 | -0.319 | 0.00824989 |
| AT3G11500.1 | | protein_coding | Small nuclear ribonucleo | 380 | 332 | 394 | 355 | 279 | 255 | 799 | 816 | 1009 | 770 | 741 | 693 | 831.59 | 875 | 735 | 0.801 | -0.319 | 0.00770159 |
| AT1G18840.1 | IQD30 | protein_coding | IQ-domain 30 | 630 | 590 | 595 | 590 | 492 | 478 | 1324 | 1450 | 1523 | 1280 | 1306 | 1300 | 1181.42 | 1432 | 1295 | 0.802 | -0.319 | 0.00572395 |
| AT2G42130.4 | PAP13 | protein_coding | Plastid-lipid associated p | 979 | 881 | 882 | 665 | 594 | 565 | 2058 | 2164 | 2258 | 1442 | 1577 | 1536 | 1956.21 | 2160 | 1518 | 0.802 | -0.319 | 0.00421375 |
| AT3G14930.1 | HEME1 | protein_coding | Uroporphyrinogen decar | 4933 | 4342 | 3428 | 2941 | 2618 | 2272 | 10370 | 10667 | 8776 | 6379 | 6950 | 6179 | 9773.12 | 9938 | 6503 | 0.801 | -0.32 | 0.00985257 |
| AT2G05210.1 | POT1A | protein_coding | Nucleic acid-binding, OB- | 380 | 367 | 358 | 296 | 297 | 283 | 799 | 902 | 917 | 642 | 788 | 770 | 792.44 | 873 | 733 | 0.801 | -0.32 | 0.00493765 |
| AT4G29010.1 | AIM1 | protein_coding | Enoyl-CoA hydratase/iso | 2582 | 2115 | 2074 | 1989 | 1598 | 1558 | 5428 | 5196 | 5310 | 4314 | 4242 | 4237 | 4213.29 | 5311 | 4264 | 0.801 | -0.32 | 0.00337316 |
| AT4G18975.1 | | protein_coding | Pentatricopeptide repeat | 421 | 328 | 316 | 274 | 239 | 220 | 885 | 806 | 809 | 594 | 634 | 598 | 745.33 | 833 | 609 | 0.801 | -0.32 | 0.00267422 |
| AT1G19650.1 | SFH4 | protein_coding | Sec14p-like phosphatidyl | 434 | 417 | 379 | 315 | 277 | 223 | 912 | 1024 | 970 | 683 | 735 | 606 | 803.17 | 969 | 675 | 0.801 | -0.321 | 0.00899425 |
| AT1G38065.1 | | protein_coding | O-fucosyltransferase fam | 169 | 156 | 149 | 138 | 120 | 95 | 355 | 383 | 381 | 299 | 319 | 258 | 365.06 | 373 | 292 | 0.8 | -0.321 | 0.0040946 |
| AT3G21160.1 | MNS2 | protein_coding | alpha-mannosidase 2 | 406 | 284 | 277 | 284 | 220 | 229 | 853 | 698 | 709 | 616 | 584 | 623 | 633.19 | 753 | 608 | 0.8 | -0.322 | 0.00891231 |
| AT2G45010.1 | | protein_coding | PLAC8 family protein | 205 | 168 | 165 | 140 | 110 | 149 | 431 | 413 | 422 | 304 | 292 | 405 | 429.17 | 422 | 334 | 0.8 | -0.322 | 0.0078543 |
| AT5G55040.1 | | protein_coding | DNA-binding bromodom | 768 | 675 | 649 | 546 | 461 | 450 | 1614 | 1658 | 1661 | 1184 | 1224 | 1224 | 1289.88 | 1644 | 1211 | 0.8 | -0.322 | 0.00623877 |
| AT1G35190.1 | | protein_coding | 2-oxoglutarate (2OG) an | 304 | 256 | 282 | 271 | 191 | 207 | 639 | 629 | 722 | 588 | 507 | 563 | 573.25 | 663 | 553 | 0.8 | -0.322 | 0.00470262 |
| AT3G62330.1 | | protein_coding | Zinc knuckle (CCHC-type) | 379 | 302 | 322 | 251 | 197 | 216 | 797 | 742 | 824 | 544 | 523 | 587 | 567.42 | 788 | 551 | 0.8 | -0.322 | 0.0033097 |
| AT3G01920.1 | | protein_coding | DHBP synthase RibB-like | 239 | 216 | 200 | 150 | 152 | 174 | 502 | 531 | 512 | 325 | 404 | 473 | 673.09 | 515 | 401 | 0.799 | -0.323 | 0.00648014 |
| AT1G53670.1 | MSRB1 | protein_coding | methionine sulfoxide red | 1722 | 1682 | 1868 | 1167 | 984 | 1039 | 3620 | 4132 | 4782 | 2531 | 2612 | 2825 | 3372.1 | 4178 | 2656 | 0.799 | -0.324 | 0.00886221 |
| AT2G47690.1 | | protein_coding | NADH-ubiquinone oxidor | 280 | 217 | 230 | 280 | 177 | 168 | 589 | 533 | 589 | 607 | 470 | 457 | 693.2 | 570 | 511 | 0.799 | -0.324 | 0.00788077 |
| AT5G57360.2 | ADO1 | protein_coding | Galactose oxidase/kelch | 564 | 569 | 502 | 465 | 390 | 473 | 1186 | 1398 | 1285 | 1009 | 1035 | 1286 | 1111.71 | 1290 | 1110 | 0.798 | -0.325 | 0.0087498 |
| AT3G60130.1 | BGLU16 | protein_coding | beta glucosidase 16 | 549 | 544 | 547 | 404 | 318 | 374 | 1154 | 1337 | 1400 | 876 | 844 | 1017 | 1261.7 | 1297 | 912 | 0.798 | -0.325 | 0.00861412 |
| AT3G27470.1 | | protein_coding | Protein of unknown funct | 253 | 226 | 212 | 233 | 185 | 151 | 532 | 555 | 543 | 505 | 491 | 411 | 458.6 | 543 | 469 | 0.799 | -0.325 | 0.00681411 |

| | | | | | | | | | | | | | | | | | | | | | |
|---|---|---|---|---|---|---|---|---|---|---|---|---|---|---|---|---|---|---|---|---|---|
| AT1G07080.1 | | protein_coding | Thioredoxin superfamily | 418 | 411 | 335 | 409 | 333 | 317 | 879 | 1010 | 858 | 887 | 884 | 862 | 1371.15 | 916 | 878 | 0.798 | -0.325 | 0.00614112 |
| AT4G05530.1 | SDRA | protein_coding | indole-3-butyric acid resp | 325 | 279 | 261 | 239 | 198 | 197 | 683 | 685 | 668 | 518 | 526 | 536 | 649.25 | 679 | 527 | 0.798 | -0.326 | 0.0012081 |
| AT1G28100.4 | | protein_coding | | 222 | 223 | 191 | 116 | 128 | 92 | 467 | 548 | 489 | 252 | 340 | 250 | 468.65 | 501 | 281 | 0.797 | -0.327 | 0.00831697 |
| AT5G38380.1 | | protein_coding | | 200 | 149 | 174 | 140 | 134 | 126 | 420 | 366 | 445 | 304 | 356 | 343 | 413.03 | 410 | 334 | 0.797 | -0.327 | 0.00556222 |
| AT3G18820.1 | RABG3F | protein_coding | RAB GTPase homolog G3 | 1589 | 1107 | 1246 | 1246 | 821 | 945 | 3340 | 2720 | 3190 | 2703 | 2180 | 2570 | 2962.76 | 3083 | 2484 | 0.797 | -0.328 | 0.0093662 |
| AT1G68830.1 | STN7 | protein_coding | STT7 homolog STN7 | 2300 | 2319 | 2317 | 2034 | 1807 | 1797 | 4835 | 5697 | 5932 | 4412 | 4797 | 4887 | 4997.43 | 5488 | 4699 | 0.796 | -0.328 | 0.00593455 |
| AT2G30470.1 | VAL1 | protein_coding | high-level expression of s | 313 | 323 | 310 | 254 | 191 | 207 | 658 | 794 | 794 | 551 | 507 | 563 | 655.31 | 749 | 540 | 0.797 | -0.328 | 0.00539539 |
| AT3G15090.1 | | protein_coding | GroES-like zinc-binding al | 256 | 183 | 225 | 174 | 139 | 142 | 538 | 450 | 576 | 377 | 369 | 386 | 436.47 | 521 | 377 | 0.797 | -0.328 | 0.00425571 |
| AT4G35910.1 | CTU2 | protein_coding | Adenine nucleotide alpha | 212 | 159 | 154 | 133 | 106 | 111 | 446 | 391 | 394 | 288 | 281 | 302 | 337.83 | 410 | 290 | 0.796 | -0.329 | 0.00968337 |
| AT1G53165.3 | ATMAP4K | protein_coding | Protein kinase superfamil | 511 | 358 | 477 | 346 | 288 | 308 | 1074 | 880 | 1221 | 751 | 765 | 838 | 708.49 | 1058 | 785 | 0.796 | -0.329 | 0.00905994 |
| AT4G02790.1 | DGP3 | protein_coding | GTP-binding family prote | 1304 | 1016 | 913 | 924 | 723 | 606 | 2741 | 2496 | 2337 | 2004 | 1919 | 1648 | 2293.92 | 2525 | 1857 | 0.796 | -0.329 | 0.00609462 |
| AT1G55130.1 | TMN6 | protein_coding | Endomembrane protein 7 | 401 | 330 | 288 | 295 | 265 | 190 | 843 | 811 | 737 | 640 | 704 | 517 | 660.1 | 797 | 620 | 0.796 | -0.329 | 0.00579011 |
| AT5G53310.1 | | protein_coding | myosin heavy chain-relat | 475 | 374 | 449 | 378 | 322 | 355 | 998 | 919 | 1149 | 820 | 855 | 965 | 897.96 | 1022 | 880 | 0.796 | -0.329 | 0.0050153 |
| AT3G45230.1 | | protein_coding | hydroxyproline-rich glyco | 132 | 134 | 95 | 131 | 81 | 89 | 277 | 329 | 243 | 284 | 215 | 242 | 478.23 | 283 | 247 | 0.796 | -0.33 | 0.00726113 |
| AT1G68820.1 | | protein_coding | Transmembrane Fragile-X | 230 | 214 | 217 | 202 | 175 | 162 | 483 | 526 | 556 | 438 | 465 | 441 | 487.13 | 522 | 448 | 0.796 | -0.33 | 0.00614741 |
| AT1G51160.1 | | protein_coding | SNARE-like superfamily p | 406 | 353 | 368 | 357 | 287 | 299 | 853 | 867 | 942 | 774 | 762 | 813 | 943.17 | 887 | 783 | 0.796 | -0.33 | 0.00307572 |
| AT4G33670.1 | LGALDH | protein_coding | NAD(P)-linked oxidoredu | 663 | 565 | 557 | 380 | 358 | 413 | 1394 | 1388 | 1426 | 824 | 950 | 1123 | 1440.26 | 1403 | 966 | 0.795 | -0.331 | 0.00728034 |
| AT3G12290.1 | FOLD2 | protein_coding | Amino acid dehydrogena | 1309 | 1150 | 1022 | 1000 | 787 | 931 | 2752 | 2825 | 2616 | 2169 | 2089 | 2532 | 3111.27 | 2731 | 2263 | 0.795 | -0.331 | 0.0038793 |
| AT1G14270.1 | | protein_coding | CAAX amino terminal pro | 888 | 789 | 666 | 598 | 562 | 495 | 1867 | 1938 | 1705 | 1297 | 1492 | 1346 | 2026.93 | 1837 | 1378 | 0.795 | -0.331 | 0.00324049 |
| AT3G47860.1 | CHL | protein_coding | chloroplastic lipocalin | 1273 | 963 | 1138 | 913 | 853 | 852 | 2676 | 2366 | 2913 | 1980 | 2265 | 2317 | 2457.36 | 2652 | 2187 | 0.794 | -0.332 | 0.00920377 |
| AT3G24870.1 | EAF1B | protein_coding | Helicase/SANT-associatec | 788 | 772 | 842 | 671 | 615 | 598 | 1656 | 1897 | 2156 | 1455 | 1633 | 1626 | 1243.58 | 1903 | 1571 | 0.794 | -0.332 | 0.007931 |
| AT4G09760.2 | F17A8.110 | protein_coding | Protein kinase superfamil | 578 | 604 | 567 | 363 | 350 | 271 | 1215 | 1484 | 1452 | 787 | 929 | 737 | 1187.51 | 1384 | 818 | 0.794 | -0.332 | 0.00764626 |
| AT4G06598.1 | | protein_coding | | 207 | 157 | 173 | 133 | 92 | 106 | 435 | 386 | 443 | 288 | 244 | 288 | 348.38 | 421 | 273 | 0.794 | -0.332 | 0.00417498 |
| AT5G11580.1 | | protein_coding | Regulator of chromosom | 1006 | 888 | 792 | 627 | 493 | 552 | 2115 | 2182 | 2028 | 1360 | 1309 | 1501 | 2237.13 | 2108 | 1390 | 0.794 | -0.332 | 0.00264203 |
| AT1G27480.1 | LCAT1 | protein_coding | alpha/beta-Hydrolases su | 526 | 582 | 433 | 572 | 442 | 390 | 1106 | 1430 | 1109 | 1241 | 1173 | 1061 | 1325.06 | 1215 | 1158 | 0.794 | -0.333 | 0.00952651 |
| AT5G07470.1 | MSRA3 | protein_coding | peptidemethionine sulfox | 336 | 338 | 296 | 255 | 235 | 272 | 706 | 830 | 758 | 553 | 624 | 740 | 963.34 | 765 | 639 | 0.794 | -0.333 | 0.00864969 |
| AT3G53760.1 | GCP4 | protein_coding | GAMMA-TUBULIN COMP | 247 | 191 | 173 | 185 | 187 | 201 | 519 | 469 | 443 | 401 | 496 | 547 | 412.68 | 477 | 481 | 0.794 | -0.333 | 0.00760223 |
| AT2G20440.1 | | protein_coding | Ypt/Rab-GAP domain of g | 159 | 121 | 133 | 127 | 86 | 88 | 334 | 297 | 340 | 275 | 228 | 239 | 264.08 | 324 | 247 | 0.794 | -0.334 | 0.00974652 |
| AT2G02410.1 | | protein_coding | | 238 | 204 | 200 | 146 | 113 | 132 | 500 | 501 | 512 | 317 | 300 | 359 | 399.5 | 504 | 325 | 0.793 | -0.334 | 0.00365466 |
| AT1G58250.2 | SAB | protein_coding | Golgi-body localisation pr | 2183 | 1849 | 1826 | 1681 | 1443 | 1292 | 4589 | 4543 | 4675 | 3646 | 3831 | 3514 | 2763.16 | 4602 | 3664 | 0.793 | -0.334 | 0.0027429 |
| AT1G28150.1 | | protein_coding | | 395 | 367 | 405 | 329 | 305 | 343 | 830 | 902 | 1037 | 714 | 810 | 933 | 1030.85 | 923 | 819 | 0.793 | -0.335 | 0.00743796 |
| AT2G32080.1 | PURA1 | protein_coding | purin-rich alpha 1 | 847 | 715 | 714 | 875 | 632 | 509 | 1780 | 1757 | 1828 | 1898 | 1678 | 1384 | 2025.85 | 1788 | 1653 | 0.793 | -0.335 | 0.00699281 |
| AT4G19110.2 | | protein_coding | Protein kinase superfamil | 1361 | 1160 | 1182 | 1178 | 980 | 938 | 2861 | 2850 | 3026 | 2555 | 2602 | 2551 | 2895.85 | 2912 | 2569 | 0.793 | -0.335 | 0.00330809 |
| AT2G30460.1 | | protein_coding | Nucleotide/sugar transpo | 277 | 207 | 188 | 229 | 164 | 139 | 582 | 509 | 481 | 497 | 435 | 378 | 552.84 | 524 | 437 | 0.793 | -0.335 | 0.00318724 |
| AT1G31850.1 | | protein_coding | S-adenosyl-L-methionine- | 882 | 682 | 750 | 653 | 477 | 559 | 1854 | 1676 | 1920 | 1416 | 1266 | 1520 | 1636.01 | 1817 | 1401 | 0.792 | -0.336 | 0.00462069 |
| AT4G29790.1 | | protein_coding | | 610 | 536 | 583 | 468 | 427 | 397 | 1282 | 1317 | 1493 | 1015 | 1134 | 1080 | 915.42 | 1364 | 1076 | 0.792 | -0.336 | 0.00390394 |
| AT3G02860.2 | | protein_coding | zinc ion binding | 241 | 194 | 186 | 151 | 127 | 135 | 507 | 477 | 476 | 328 | 337 | 367 | 386.05 | 487 | 344 | 0.792 | -0.336 | 0.00307284 |
| AT4G02460.1 | PMS1 | protein_coding | DNA mismatch repair pro | 218 | 166 | 168 | 147 | 140 | 115 | 458 | 408 | 430 | 319 | 372 | 313 | 319.14 | 432 | 335 | 0.791 | -0.337 | 0.00694729 |
| AT3G09360.1 | | protein_coding | Cyclin/Brf1-like TBP-bindi | 209 | 164 | 141 | 113 | 99 | 107 | 439 | 403 | 361 | 245 | 263 | 291 | 321.65 | 401 | 266 | 0.792 | -0.337 | 0.00554878 |
| AT1G15470.1 | | protein_coding | Transducin/WD40 repeat | 235 | 203 | 180 | 168 | 169 | 140 | 494 | 499 | 461 | 364 | 449 | 381 | 473.39 | 485 | 398 | 0.792 | -0.337 | 0.00538376 |
| AT4G25360.1 | TBL18 | protein_coding | TRICHOME BIREFRINGEN | 336 | 292 | 305 | 292 | 258 | 288 | 706 | 717 | 781 | 633 | 685 | 783 | 616.96 | 735 | 700 | 0.792 | -0.337 | 0.00429233 |
| AT5G36170.1 | PRFB1 | protein_coding | high chlorophyll fluoresce | 955 | 836 | 888 | 650 | 571 | 569 | 2007 | 2054 | 2273 | 1410 | 1516 | 1547 | 2118.03 | 2111 | 1491 | 0.792 | -0.337 | 0.00285313 |
| AT4G36890.1 | IRX14 | protein_coding | Nucleotide-diphospho-su | 326 | 239 | 259 | 250 | 197 | 198 | 685 | 587 | 663 | 542 | 523 | 538 | 556.39 | 645 | 534 | 0.792 | -0.337 | 0.00140558 |
| AT2G21380.1 | | protein_coding | Kinesin motor family prot | 1900 | 2014 | 1739 | 1448 | 1410 | 1110 | 3994 | 4948 | 4452 | 3141 | 3743 | 3019 | 3362.88 | 4465 | 3301 | 0.79 | -0.339 | 0.0083486 |
| AT5G28040.1 | | protein_coding | DNA-binding storekeeper | 218 | 222 | 199 | 194 | 173 | 210 | 458 | 545 | 509 | 421 | 459 | 571 | 499.68 | 504 | 484 | 0.79 | -0.339 | 0.00798027 |
| AT4G01026.1 | PYL7 | protein_coding | PYR1-like 7 | 168 | 162 | 153 | 202 | 190 | 149 | 353 | 398 | 392 | 438 | 504 | 405 | 426.16 | 381 | 449 | 0.79 | -0.339 | 0.00752407 |
| AT4G34120.1 | CBSX2 | protein_coding | Cystathionine beta-synth | 763 | 699 | 548 | 563 | 485 | 395 | 1604 | 1717 | 1403 | 1221 | 1288 | 1074 | 2666.55 | 1575 | 1194 | 0.79 | -0.339 | 0.00509359 |
| AT3G55830.1 | EPC1 | protein_coding | Nucleotide-diphospho-su | 453 | 363 | 406 | 391 | 296 | 334 | 952 | 892 | 1039 | 848 | 786 | 908 | 792.22 | 961 | 847 | 0.79 | -0.339 | 0.00322121 |
| AT2G28930.1 | APK1B | protein_coding | protein kinase 1B | 459 | 363 | 352 | 315 | 234 | 233 | 965 | 892 | 901 | 683 | 621 | 634 | 840.37 | 919 | 646 | 0.791 | -0.339 | 0.00268464 |
| AT3G48330.1 | PIMT1 | protein_coding | protein-l-isoaspartate me | 176 | 136 | 149 | 151 | 117 | 118 | 370 | 334 | 381 | 328 | 311 | 321 | 394.17 | 362 | 320 | 0.791 | -0.339 | 0.0022094 |
| AT2G18910.1 | | protein_coding | hydroxyproline-rich glyco | 361 | 270 | 291 | 311 | 187 | 193 | 759 | 663 | 745 | 675 | 496 | 525 | 774.89 | 722 | 565 | 0.79 | -0.34 | 0.00879711 |
| AT4G22290.1 | | protein_coding | Ubiquitin-specific protea | 334 | 380 | 309 | 257 | 254 | 272 | 702 | 934 | 791 | 557 | 674 | 740 | 814.65 | 809 | 657 | 0.79 | -0.34 | 0.0080491 |
| AT3G27390.1 | | protein_coding | | 821 | 617 | 585 | 508 | 360 | 371 | 1726 | 1516 | 1498 | 1102 | 956 | 1009 | 1218.66 | 1580 | 1022 | 0.79 | -0.34 | 0.00366904 |
| AT5G50180.1 | | protein_coding | Protein kinase superfamil | 483 | 443 | 507 | 346 | 336 | 326 | 1015 | 1088 | 1298 | 751 | 892 | 887 | 1063.59 | 1134 | 843 | 0.79 | -0.341 | 0.00936 |
| AT2G17033.2 | | protein_coding | pentatricopeptide (PPR) r | 961 | 667 | 646 | 609 | 423 | 446 | 2020 | 1639 | 1654 | 1321 | 1123 | 1213 | 1422.42 | 1771 | 1219 | 0.79 | -0.341 | 0.00846224 |
| AT2G18730.1 | DGK3 | protein_coding | diacylglycerol kinase 3 | 553 | 396 | 396 | 481 | 305 | 322 | 1162 | 973 | 1014 | 1043 | 810 | 876 | 1166.57 | 1050 | 910 | 0.79 | -0.341 | 0.0061763 |
| AT4G39860.1 | | protein_coding | | 279 | 291 | 231 | 202 | 205 | 189 | 586 | 715 | 591 | 438 | 544 | 514 | 588.81 | 631 | 499 | 0.789 | -0.341 | 0.00519311 |
| AT3G25470.1 | | protein_coding | bacterial hemolysin-relat | 1400 | 1007 | 1048 | 924 | 695 | 655 | 2943 | 2474 | 2683 | 2004 | 1845 | 1781 | 2465.97 | 2700 | 1877 | 0.789 | -0.341 | 0.00356782 |
| AT2G03480.1 | QUL2 | protein_coding | QUASIMODO2 LIKE 2 | 312 | 282 | 257 | 231 | 215 | 206 | 656 | 693 | 658 | 501 | 571 | 560 | 601 | 669 | 544 | 0.789 | -0.341 | 0.00145957 |
| AT3G51470.1 | | protein_coding | Protein phosphatase 2C f | 151 | 139 | 104 | 121 | 89 | 95 | 317 | 341 | 266 | 262 | 236 | 258 | 334.63 | 308 | 252 | 0.789 | -0.342 | 0.00710076 |
| AT2G03620.1 | MRS2-5 | protein_coding | magnesium transporter 3 | 153 | 135 | 155 | 148 | 126 | 112 | 322 | 332 | 397 | 321 | 335 | 305 | 292.21 | 350 | 320 | 0.789 | -0.343 | 0.00916307 |
| AT1G27680.1 | APL2 | protein_coding | ADPGLC-PPase large subu | 289 | 266 | 203 | 226 | 223 | 145 | 607 | 654 | 520 | 490 | 592 | 394 | 617.42 | 594 | 492 | 0.788 | -0.343 | 0.00742493 |
| AT2G02220.1 | PSKR1 | protein_coding | phytosulfokin receptor 1 | 383 | 311 | 357 | 290 | 222 | 287 | 805 | 764 | 914 | 629 | 589 | 780 | 564.16 | 828 | 666 | 0.788 | -0.343 | 0.00660074 |
| AT4G38420.1 | sks9 | protein_coding | SKU5 similar 9 | 613 | 590 | 427 | 549 | 447 | 421 | 1289 | 1450 | 1093 | 1191 | 1187 | 1145 | 1500.41 | 1277 | 1174 | 0.789 | -0.343 | 0.00634159 |
| AT5G35430.1 | | protein_coding | Tetratricopeptide repeat | 829 | 644 | 598 | 666 | 519 | 488 | 1743 | 1582 | 1531 | 1445 | 1378 | 1327 | 1207.12 | 1619 | 1383 | 0.789 | -0.343 | 0.0025355 |
| AT1G32160.1 | | protein_coding | Protein of unknown funct | 1195 | 1168 | 1191 | 987 | 925 | 795 | 2512 | 2870 | 3049 | 2141 | 2456 | 2162 | 2588.22 | 2810 | 2253 | 0.788 | -0.344 | 0.00570867 |
| AT2G38630.1 | | protein_coding | Transducin/WD40 repeat | 238 | 237 | 234 | 189 | 167 | 149 | 500 | 582 | 599 | 410 | 443 | 405 | 465.24 | 560 | 419 | 0.788 | -0.344 | 0.00475129 |
| AT1G50360.1 | VIII-A | protein_coding | P-loop containing nucleo | 382 | 350 | 399 | 314 | 266 | 273 | 803 | 860 | 1021 | 681 | 706 | 742 | 682.12 | 895 | 710 | 0.788 | -0.344 | 0.00352059 |
| AT5G16910.1 | CSLD2 | protein_coding | cellulose-synthase like D2 | 407 | 387 | 343 | 310 | 232 | 250 | 856 | 951 | 878 | 672 | 616 | 680 | 674.58 | 895 | 656 | 0.788 | -0.344 | 0.00327215 |
| AT2G25950.1 | | protein_coding | Protein of unknown funct | 382 | 270 | 296 | 182 | 203 | 138 | 803 | 663 | 758 | 395 | 539 | 375 | 645.39 | 741 | 436 | 0.787 | -0.345 | 0.00767081 |
| AT3G13060.2 | ECT5 | protein_coding | evolutionarily conserved | 527 | 525 | 472 | 423 | 407 | 408 | 1108 | 1290 | 1208 | 918 | 1080 | 1110 | 1134.57 | 1202 | 1036 | 0.787 | -0.345 | 0.00349235 |
| AT4G36440.1 | | protein_coding | | 406 | 266 | 342 | 270 | 197 | 228 | 853 | 654 | 876 | 586 | 523 | 620 | 682.51 | 794 | 576 | 0.787 | -0.345 | 0.00337002 |
| AT2G39480.1 | ABCB6 | protein_coding | P-glycoprotein 6 | 1009 | 936 | 1021 | 740 | 717 | 719 | 2121 | 2300 | 2614 | 1605 | 1903 | 1955 | 1556.74 | 2345 | 1821 | 0.787 | -0.346 | 0.00814447 |
| AT3G02600.1 | LPP3 | protein_coding | lipid phosphate phosphat | 539 | 486 | 398 | 410 | 319 | 371 | 1133 | 1194 | 1019 | 889 | 847 | 1009 | 1181.79 | 1115 | 915 | 0.787 | -0.346 | 0.00343518 |
| AT5G35930.1 | | protein_coding | AMP-dependent syntheta | 316 | 269 | 261 | 221 | 171 | 188 | 664 | 661 | 668 | 479 | 454 | 511 | 501.3 | 664 | 481 | 0.787 | -0.346 | 0.00266687 |
| AT3G53900.2 | UPP | protein_coding | uracil phosphoribosyltran | 645 | 564 | 558 | 509 | 424 | 428 | 1356 | 1386 | 1429 | 1104 | 1126 | 1164 | 1665.98 | 1390 | 1131 | 0.787 | -0.346 | 0.00180958 |
| AT3G24880.1 | EAF1A | protein_coding | Helicase/SANT-associatec | 745 | 676 | 641 | 582 | 503 | 498 | 1566 | 1661 | 1641 | 1262 | 1335 | 1354 | 1003.94 | 1623 | 1317 | 0.787 | -0.346 | 0.00168255 |
| AT5G05800.1 | LIMYB | protein_coding | | 334 | 265 | 270 | 253 | 192 | 196 | 702 | 651 | 691 | 549 | 510 | 533 | 581.83 | 681 | 531 | 0.787 | -0.346 | 0.00106461 |
| AT5G16800.3 | | protein_coding | Acyl-CoA N-acyltransfera | 149 | 159 | 169 | 143 | 144 | 137 | 313 | 391 | 433 | 310 | 382 | 373 | 373.03 | 379 | 355 | 0.786 | -0.347 | 0.00847831 |
| AT1G13000.1 | | protein_coding | Protein of unknown funct | 318 | 254 | 330 | 254 | 169 | 225 | 668 | 624 | 845 | 551 | 449 | 612 | 574.02 | 712 | 537 | 0.786 | -0.347 | 0.00832108 |
| AT1G26130.2 | ALA12 | protein_coding | ATPase E1-E2 type family | 456 | 479 | 430 | 311 | 308 | 239 | 959 | 1177 | 1101 | 675 | 818 | 650 | 688.34 | 1079 | 714 | 0.786 | -0.347 | 0.00609456 |
| AT5G12200.1 | PYD2 | protein_coding | pyrimidine 2 | 414 | 280 | 341 | 331 | 241 | 265 | 870 | 688 | 873 | 718 | 640 | 721 | 725.2 | 810 | 693 | 0.786 | -0.347 | 0.00244889 |
| AT5G05570.1 | | protein_coding | transducin family protein | 493 | 441 | 443 | 398 | 306 | 337 | 1036 | 1083 | 1134 | 863 | 812 | 916 | 820.4 | 1084 | 864 | 0.786 | -0.347 | 0.00151591 |
| AT5G43430.1 | ETFB | protein_coding | electron transfer flavopro | 223 | 176 | 242 | 197 | 167 | 179 | 469 | 432 | 620 | 427 | 443 | 487 | 473.31 | 507 | 452 | 0.786 | -0.348 | 0.00593741 |
| AT2G38610.1 | | protein_coding | RNA-binding KH domain- | 336 | 288 | 310 | 301 | 220 | 261 | 706 | 708 | 794 | 653 | 584 | 710 | 760.7 | 736 | 649 | 0.786 | -0.348 | 0.00183641 |
| AT3G58790.1 | GAUT15 | protein_coding | galacturonosyltransferase | 247 | 251 | 229 | 209 | 220 | 195 | 519 | 617 | 586 | 453 | 584 | 530 | 480.88 | 574 | 522 | 0.785 | -0.349 | 0.0085483 |
| AT5G01950.1 | | protein_coding | Leucine-rich repeat prote | 312 | 317 | 259 | 243 | 244 | 210 | 656 | 779 | 663 | 527 | 648 | 571 | 772.81 | 699 | 582 | 0.785 | -0.349 | 0.00264232 |
| AT3G53920.1 | SIGC | protein_coding | RNApolymerase sigma-su | 601 | 552 | 549 | 461 | 359 | 356 | 1263 | 1356 | 1405 | 1000 | 953 | 968 | 1279.77 | 1341 | 974 | 0.785 | -0.349 | 0.00248974 |
| AT3G55140.1 | | protein_coding | Pectin lyase-like superfan | 311 | 262 | 281 | 232 | 200 | 217 | 654 | 644 | 719 | 503 | 531 | 590 | 603.97 | 672 | 541 | 0.785 | -0.349 | 0.0014532 |
| AT5G57210.1 | | protein_coding | Ypt/Rab-GAP domain of g | 332 | 317 | 378 | 287 | 287 | 271 | 698 | 779 | 968 | 623 | 762 | 737 | 648.38 | 815 | 707 | 0.785 | -0.35 | 0.00783181 |
| AT5G62200.1 | | protein_coding | Embryo-specific protein 3 | 141 | 149 | 134 | 131 | 84 | 109 | 296 | 366 | 343 | 284 | 223 | 296 | 345.7 | 335 | 268 | 0.784 | -0.35 | 0.00583089 |
| AT2G45670.1 | LPEAT2 | protein_coding | calcineurin B subunit-rela | 1263 | 1016 | 1101 | 1002 | 833 | 762 | 2655 | 2496 | 2819 | 2173 | 2211 | 2072 | 2468.9 | 2657 | 2152 | 0.784 | -0.35 | 0.00227716 |
| AT2G17580.1 | | protein_coding | Polynucleotide adenylyltr | 360 | 267 | 242 | 276 | 204 | 176 | 757 | 656 | 620 | 599 | 542 | 479 | 510.46 | 678 | 540 | 0.784 | -0.351 | 0.0052732 |
| AT4G18520.1 | PCMP-A2 | protein_coding | Pentatricopeptide repeat | 356 | 271 | 274 | 166 | 187 | 168 | 748 | 666 | 701 | 360 | 496 | 457 | 556.76 | 705 | 438 | 0.784 | -0.351 | 0.00385028 |
| AT3G48760.1 | PAT05 | protein_coding | DHHC-type zinc finger far | 583 | 532 | 525 | 470 | 366 | 407 | 1226 | 1307 | 1344 | 1019 | 972 | 1107 | 1065.93 | 1292 | 1033 | 0.784 | -0.351 | 0.00177821 |
| AT3G09210.1 | PTAC13 | protein_coding | plastid transcriptionally a | 1303 | 1127 | 1078 | 850 | 725 | 739 | 2739 | 2769 | 2760 | 1844 | 1925 | 2010 | 2822.02 | 2756 | 1926 | 0.784 | -0.351 | 0.00142643 |
| AT1G52260.1 | PDIL1-5 | protein_coding | PDI-like 1-5 | 173 | 164 | 147 | 135 | 106 | 117 | 364 | 403 | 376 | 293 | 281 | 318 | 302.03 | 381 | 297 | 0.783 | -0.352 | 0.0037321 |
| AT2G45980.1 | | protein_coding | | 577 | 523 | 586 | 418 | 351 | 442 | 1213 | 1285 | 1500 | 907 | 932 | 1202 | 1133.78 | 1333 | 1014 | 0.783 | -0.353 | 0.00745397 |
| AT5G45560.1 | EDR2L | protein_coding | Pleckstrin homology (PH) | 179 | 168 | 170 | 136 | 119 | 103 | 376 | 413 | 435 | 295 | 316 | 280 | 328.67 | 408 | 297 | 0.783 | -0.353 | 0.00326357 |
| AT1G35220.1 | | protein_coding | | 432 | 303 | 299 | 335 | 280 | 245 | 908 | 744 | 765 | 727 | 743 | 666 | 782.77 | 806 | 712 | 0.782 | -0.354 | 0.00322867 |
| AT5G02790.1 | GSTL3 | protein_coding | Glutathione S-transferase | 2494 | 1913 | 2042 | 2372 | 1661 | 1521 | 5243 | 4700 | 5228 | 5145 | 4410 | 4136 | 4958.76 | 5057 | 4564 | 0.782 | -0.354 | 0.00280928 |
| AT3G28970.1 | AAR3 | protein_coding | Domain of unknown func | 353 | 268 | 285 | 237 | 193 | 200 | 742 | 658 | 730 | 514 | 512 | 544 | 538.4 | 710 | 523 | 0.783 | -0.354 | 0.00279524 |
| AT2G34090.2 | MEE18 | protein_coding | maternal effect embryo a | 347 | 273 | 277 | 185 | 161 | 155 | 729 | 671 | 709 | 401 | 427 | 422 | 652.53 | 703 | 417 | 0.782 | -0.354 | 0.00233395 |
| AT4G30996.1 | | protein_coding | Protein of unknown funct | 377 | 312 | 326 | 339 | 244 | 279 | 792 | 767 | 835 | 735 | 648 | 759 | 850.03 | 798 | 714 | 0.783 | -0.354 | 0.00106612 |
| AT3G45190.1 | | protein_coding | SIT4 phosphatase-associa | 1062 | 994 | 912 | 841 | 803 | 674 | 2232 | 2442 | 2335 | 1824 | 2132 | 1833 | 1606.95 | 2336 | 1930 | 0.782 | -0.355 | 0.00240775 |
| AT5G37780.3 | CAM4 | protein_coding | calmodulin 1 | 1104 | 951 | 1011 | 911 | 666 | 685 | 2321 | 2336 | 2588 | 1976 | 1768 | 1863 | 2512.53 | 2415 | 1869 | 0.782 | -0.355 | 0.00211376 |
| AT2G43070.1 | SPPL3 | protein_coding | SIGNAL PEPTIDE PEPTIDA | 700 | 520 | 534 | 539 | 428 | 389 | 1471 | 1278 | 1367 | 1169 | 1136 | 1058 | 1253.79 | 1372 | 1121 | 0.782 | -0.355 | 0.00206816 |
| AT1G14000.1 | VIK | protein_coding | VH1-interacting kinase | 992 | 844 | 883 | 763 | 594 | 610 | 2085 | 2074 | 2261 | 1655 | 1577 | 1659 | 1754.81 | 2140 | 1630 | 0.782 | -0.355 | 0.00163115 |
| AT1G76730.1 | COG0212 | protein_coding | NagB/RpiA/CoA transfera | 731 | 682 | 617 | 592 | 506 | 525 | 1537 | 1676 | 1580 | 1284 | 1343 | 1428 | 1667.97 | 1598 | 1352 | 0.782 | -0.355 | 0.00152365 |
| AT1G50890.1 | SP2L | protein_coding | ARM repeat superfamily | 149 | 149 | 134 | 147 | 111 | 132 | 313 | 366 | 343 | 319 | 295 | 359 | 336.89 | 341 | 324 | 0.781 | -0.356 | 0.00786133 |
| AT4G37760.1 | SQE3 | protein_coding | squalene epoxidase 3 | 975 | 879 | 837 | 913 | 758 | 792 | 2050 | 2160 | 2143 | 1980 | 2012 | 2154 | 1836.52 | 2118 | 2049 | 0.781 | -0.356 | 0.00304234 |
| AT5G51710.1 | KEA5 | protein_coding | K+ efflux antiporter 5 | 265 | 242 | 217 | 216 | 148 | 177 | 557 | 595 | 556 | 469 | 393 | 481 | 464.42 | 569 | 448 | 0.781 | -0.356 | 0.00280481 |
| AT4G18370.1 | DEGP5 | protein_coding | DEGP protease 5 | 1098 | 956 | 853 | 810 | 692 | 635 | 2308 | 2349 | 2184 | 1757 | 1837 | 1727 | 2359.18 | 2280 | 1774 | 0.781 | -0.356 | 0.00229129 |
| AT2G40316.1 | | protein_coding | | 253 | 200 | 189 | 218 | 196 | 153 | 532 | 491 | 484 | 473 | 520 | 416 | 552.48 | 502 | 470 | 0.781 | -0.356 | 0.00199673 |
| AT3G63170.1 | FAP1 | protein_coding | Chalcone-flavanone isom | 410 | 332 | 348 | 317 | 238 | 299 | 862 | 816 | 891 | 688 | 632 | 813 | 974.31 | 856 | 711 | 0.782 | -0.356 | 0.00198543 |
| AT1G20810.1 | FKBP18 | protein_coding | FKBP-like peptidyl-prolyl | 929 | 922 | 843 | 606 | 580 | 530 | 1953 | 2265 | 2158 | 1314 | 1540 | 1441 | 2134.47 | 2125 | 1432 | 0.782 | -0.356 | 0.0018264 |
| AT3G06410.1 | | protein_coding | Zinc finger C-x8-C-x5-C-x3 | 233 | 241 | 223 | 233 | 195 | 218 | 490 | 592 | 571 | 505 | 518 | 593 | 615.51 | 551 | 539 | 0.782 | -0.356 | 0.00164205 |
| AT4G19670.1 | | protein_coding | RING/U-box superfamily | 253 | 242 | 268 | 276 | 210 | 249 | 532 | 595 | 686 | 599 | 558 | 677 | 536.28 | 604 | 611 | 0.781 | -0.357 | 0.00575695 |
| AT2G19790.1 | | protein_coding | SNARE-like superfamily p | 492 | 427 | 474 | 392 | 305 | 346 | 1034 | 1049 | 1213 | 850 | 810 | 941 | 1148.04 | 1099 | 867 | 0.781 | -0.357 | 0.00338692 |
| AT5G35080.1 | OS9 | protein_coding | | 281 | 247 | 218 | 195 | 167 | 182 | 591 | 607 | 558 | 423 | 443 | 495 | 495.62 | 585 | 454 | 0.781 | -0.357 | 0.00226001 |
| AT3G49390.1 | CID10 | protein_coding | CTC-interacting domain 1 | 431 | 329 | 357 | 349 | 245 | 289 | 906 | 808 | 914 | 757 | 650 | 786 | 811.3 | 876 | 731 | 0.781 | -0.357 | 0.00199189 |
| AT5G56340.1 | ATCRT1 | protein_coding | RING/U-box superfamily | 248 | 225 | 243 | 251 | 159 | 180 | 521 | 553 | 622 | 544 | 422 | 489 | 508.95 | 565 | 485 | 0.78 | -0.358 | 0.00574775 |
| AT3G11230.2 | | protein_coding | Yippee family putative zir | 147 | 122 | 148 | 161 | 104 | 130 | 309 | 300 | 379 | 349 | 276 | 354 | 351.44 | 329 | 326 | 0.78 | -0.359 | 0.00524782 |
| AT2G29390.1 | SMO2-1 | protein_coding | sterol 4-alpha-methyl-oxi | 325 | 289 | 267 | 258 | 258 | 187 | 683 | 710 | 684 | 560 | 685 | 509 | 719.27 | 692 | 585 | 0.78 | -0.359 | 0.00355894 |
| AT1G08660.1 | MGP2 | protein_coding | MALE GAMETOPHYTE DE | 505 | 418 | 491 | 369 | 337 | 352 | 1062 | 1027 | 1257 | 800 | 895 | 957 | 1095.29 | 1115 | 884 | 0.78 | -0.359 | 0.00284201 |
| AT4G15020.1 | | protein_coding | hAT transposon superfan | 482 | 425 | 388 | 486 | 519 | 382 | 1013 | 1044 | 993 | 1054 | 1378 | 1039 | 920.95 | 1017 | 1157 | 0.779 | -0.36 | 0.00416667 |
| AT5G07842.1 | CPuORF16 | protein_coding | conserved peptide upstre | 54 | 56 | 61 | 55 | 47 | 50 | 114 | 138 | 156 | 119 | 125 | 136 | 233.42 | 136 | 127 | 0.779 | -0.361 | 0.00786569 |
| AT1G50170.1 | ATSIRB | protein_coding | sirohydrochlorin ferroche | 383 | 275 | 217 | 237 | 170 | 147 | 805 | 676 | 556 | 514 | 451 | 400 | 663.71 | 679 | 455 | 0.779 | -0.361 | 0.00700953 |
| AT5G16520.1 | | protein_coding | | 209 | 181 | 174 | 163 | 136 | 100 | 439 | 445 | 445 | 354 | 361 | 272 | 385.88 | 443 | 329 | 0.778 | -0.361 | 0.00291839 |
| AT3G07140.1 | | protein_coding | GPI transamidase compo | 603 | 418 | 435 | 360 | 331 | 284 | 1268 | 1027 | 1114 | 781 | 879 | 772 | 856.63 | 1136 | 811 | 0.779 | -0.361 | 0.00236811 |
| AT2G30070.1 | POT1 | protein_coding | potassium transporter 1 | 458 | 274 | 308 | 271 | 198 | 191 | 963 | 673 | 788 | 588 | 526 | 519 | 608.51 | 808 | 544 | 0.778 | -0.362 | 0.00513557 |
| AT5G26850.1 | | protein_coding | Uncharacterized protein | 1210 | 1265 | 1245 | 1128 | 1081 | 1084 | 2544 | 3108 | 3187 | 2447 | 2870 | 2948 | 2338.14 | 2946 | 2755 | 0.778 | -0.362 | 0.00462208 |
| AT1G13210.1 | ALA11 | protein_coding | autoinhibited Ca2+/ATPa | 519 | 452 | 430 | 401 | 302 | 310 | 1091 | 1110 | 1101 | 870 | 802 | 843 | 869.76 | 1101 | 838 | 0.778 | -0.362 | 0.00120873 |
| AT3G53780.2 | ATRBL4 | protein_coding | RHOMBOID-like protein 4 | 153 | 128 | 130 | 127 | 110 | 133 | 322 | 314 | 333 | 275 | 292 | 362 | 294.57 | 323 | 310 | 0.778 | -0.363 | 0.00541928 |
| AT3G19460.2 | RTNLB11 | protein_coding | Reticulon family protein | 215 | 204 | 246 | 240 | 199 | 189 | 452 | 501 | 630 | 521 | 528 | 514 | 574.32 | 528 | 521 | 0.778 | -0.363 | 0.00305024 |
| AT2G46060.1 | | protein_coding | transmembrane protein-r | 390 | 365 | 327 | 344 | 273 | 236 | 820 | 897 | 837 | 746 | 725 | 642 | 694.1 | 851 | 704 | 0.778 | -0.363 | 0.00077345 |
| AT3G18090.1 | NRPD2B | protein_coding | nuclear RNA polymerase | 237 | 208 | 171 | 148 | 147 | 92 | 498 | 511 | 438 | 321 | 390 | 250 | 364.66 | 482 | 320 | 0.776 | -0.365 | 0.00818369 |
| AT2G20740.1 | TOM2AH3 | protein_coding | Tetraspanin family protei | 271 | 319 | 309 | 248 | 227 | 247 | 570 | 784 | 791 | 538 | 603 | 672 | 778.25 | 715 | 604 | 0.777 | -0.365 | 0.00576382 |
| AT4G11820.2 | HMGS | protein_coding | hydroxymethylglutaryl-Co | 1134 | 1053 | 1222 | 836 | 707 | 728 | 2384 | 2587 | 3128 | 1813 | 1877 | 1980 | 2418.56 | 2700 | 1890 | 0.776 | -0.365 | 0.00330625 |
| AT5G17640.1 | | protein_coding | Protein of unknown funct | 327 | 266 | 278 | 275 | 179 | 222 | 687 | 654 | 712 | 597 | 475 | 604 | 728.32 | 684 | 559 | 0.776 | -0.365 | 0.00242884 |
| AT1G06890.1 | | protein_coding | nodulin MtN21 /EamA-lik | 695 | 605 | 664 | 547 | 467 | 417 | 1461 | 1486 | 1700 | 1186 | 1240 | 1134 | 1435.32 | 1549 | 1187 | 0.776 | -0.365 | 0.00169032 |
| AT3G53470.2 | | protein_coding | | 348 | 403 | 323 | 343 | 244 | 287 | 732 | 990 | 827 | 744 | 648 | 780 | 1030.3 | 850 | 724 | 0.776 | -0.366 | 0.00764893 |
| AT3G12980.1 | HAC5 | protein_coding | histone acetyltransferase | 895 | 859 | 834 | 652 | 618 | 606 | 1881 | 2110 | 2135 | 1414 | 1641 | 1648 | 1402.38 | 2042 | 1568 | 0.776 | -0.366 | 0.00476807 |
| AT2G38480.1 | | protein_coding | Uncharacterised protein | 123 | 91 | 90 | 99 | 87 | 86 | 259 | 224 | 230 | 215 | 231 | 234 | 279.09 | 238 | 227 | 0.776 | -0.366 | 0.00309138 |
| AT5G16660.1 | | protein_coding | | 1183 | 925 | 1069 | 897 | 649 | 732 | 2487 | 2273 | 2737 | 1946 | 1723 | 1991 | 2235.66 | 2499 | 1887 | 0.776 | -0.366 | 0.00295909 |

| | | | | | | | | | | | | | | | | | | | | | |
|---|---|---|---|---|---|---|---|---|---|---|---|---|---|---|---|---|---|---|---|---|---|
| AT5G54730.1 | ATG18F | protein_coding | homolog of yeast autoph | 278 | 238 | 268 | 220 | 158 | 190 | 584 | 585 | 686 | 477 | 419 | 517 | 438.52 | 618 | 471 | 0.776 | -0.366 | 0.00260188 |
| AT3G56720.3 | | protein_coding | | 673 | 630 | 644 | 450 | 400 | 407 | 1415 | 1548 | 1649 | 976 | 1062 | 1107 | 1245.5 | 1537 | 1048 | 0.776 | -0.366 | 0.00252628 |
| AT5G62640.3 | ELF5 | protein_coding | proline-rich family protei | 337 | 310 | 333 | 239 | 218 | 222 | 708 | 762 | 853 | 518 | 579 | 604 | 940.3 | 774 | 567 | 0.776 | -0.366 | 0.00127401 |
| AT2G18465.1 | | protein_coding | Chaperone DnaJ-domain | 143 | 96 | 119 | 119 | 103 | 85 | 301 | 236 | 305 | 258 | 273 | 231 | 250.89 | 281 | 254 | 0.775 | -0.367 | 0.00932574 |
| AT3G45050.2 | | protein_coding | | 554 | 393 | 364 | 399 | 293 | 246 | 1165 | 966 | 932 | 865 | 778 | 669 | 1039.2 | 1021 | 771 | 0.776 | -0.367 | 0.00426467 |
| AT4G04470.1 | PMP22 | protein_coding | Peroxisomal membrane 2 | 246 | 167 | 183 | 178 | 149 | 145 | 517 | 410 | 468 | 386 | 396 | 394 | 450.37 | 465 | 392 | 0.775 | -0.367 | 0.00227827 |
| AT3G16270.1 | | protein_coding | ENTH/VHS family protein | 945 | 780 | 828 | 839 | 682 | 685 | 1986 | 1916 | 2120 | 1820 | 1811 | 1863 | 1536.72 | 2007 | 1831 | 0.775 | -0.367 | 0.0018216 |
| AT1G29810.1 | | protein_coding | Transcriptional coactivato | 106 | 88 | 81 | 60 | 50 | 58 | 223 | 216 | 207 | 130 | 133 | 158 | 254.66 | 215 | 140 | 0.775 | -0.368 | 0.00841619 |
| AT4G33300.1 | ADR1-L1 | protein_coding | ADR1-like 1 | 621 | 678 | 561 | 512 | 427 | 515 | 1305 | 1666 | 1436 | 1111 | 1134 | 1401 | 1345.95 | 1469 | 1215 | 0.775 | -0.368 | 0.00498018 |
| AT5G51600.1 | MAP65-3 | protein_coding | Microtubule associated p | 393 | 380 | 355 | 282 | 294 | 219 | 826 | 934 | 909 | 612 | 781 | 596 | 639.53 | 890 | 663 | 0.775 | -0.368 | 0.0039484 |
| AT5G22875.1 | | protein_coding | | 127 | 119 | 124 | 121 | 98 | 91 | 267 | 292 | 317 | 262 | 260 | 247 | 324.48 | 292 | 256 | 0.775 | -0.368 | 0.00274571 |
| AT4G12610.2 | RAP74 | protein_coding | transcription activators;D | 321 | 258 | 262 | 187 | 191 | 180 | 675 | 634 | 671 | 406 | 507 | 489 | 631.79 | 660 | 467 | 0.775 | -0.368 | 0.00178787 |
| AT3G04350.1 | | protein_coding | Plant protein of unknowr | 329 | 206 | 236 | 251 | 190 | 186 | 692 | 506 | 604 | 544 | 504 | 506 | 483.68 | 601 | 518 | 0.774 | -0.369 | 0.0053554 |
| AT3G60810.1 | | protein_coding | | 360 | 275 | 306 | 220 | 219 | 183 | 757 | 676 | 783 | 477 | 581 | 498 | 691.29 | 739 | 519 | 0.774 | -0.369 | 0.00187821 |
| AT1G54710.1 | ATG18H | protein_coding | homolog of yeast autoph | 533 | 488 | 485 | 485 | 451 | 444 | 1120 | 1199 | 1242 | 1052 | 1197 | 1207 | 880.62 | 1187 | 1152 | 0.774 | -0.369 | 0.00181436 |
| AT1G68000.1 | PIS1 | protein_coding | phosphatidylinositol synt | 391 | 342 | 379 | 348 | 285 | 276 | 822 | 840 | 970 | 755 | 757 | 751 | 860.9 | 877 | 754 | 0.774 | -0.369 | 0.00149648 |
| AT1G01650.1 | SPPL4 | protein_coding | SIGNAL PEPTIDE PEPTIDA | 280 | 213 | 279 | 212 | 198 | 236 | 589 | 523 | 714 | 460 | 526 | 642 | 572.08 | 609 | 543 | 0.774 | -0.37 | 0.00527582 |
| AT2G39805.2 | | protein_coding | Integral membrane Yip1 | 195 | 156 | 188 | 160 | 138 | 138 | 410 | 383 | 481 | 347 | 366 | 375 | 372.63 | 425 | 363 | 0.774 | -0.37 | 0.00332641 |
| AT5G53010.1 | | protein_coding | calcium-transporting ATP | 233 | 179 | 183 | 143 | 139 | 112 | 490 | 440 | 468 | 310 | 369 | 305 | 379.18 | 466 | 328 | 0.774 | -0.37 | 0.00226188 |
| AT3G03810.1 | EDA30 | protein_coding | O-fucosyltransferase fam | 325 | 311 | 275 | 255 | 219 | 194 | 683 | 764 | 704 | 553 | 581 | 528 | 556.23 | 717 | 554 | 0.774 | -0.37 | 0.00095914 |
| AT1G45230.1 | | protein_coding | Protein of unknown funct | 796 | 598 | 503 | 521 | 398 | 349 | 1673 | 1469 | 1288 | 1130 | 1057 | 949 | 1314.43 | 1477 | 1045 | 0.773 | -0.371 | 0.00304321 |
| AT3G18060.1 | | protein_coding | transducin family protein | 2231 | 1940 | 1718 | 1762 | 1424 | 1280 | 4690 | 4766 | 4398 | 3822 | 3780 | 3481 | 3622.21 | 4618 | 3694 | 0.773 | -0.371 | 0.00180625 |
| AT2G23390.1 | | protein_coding | | 1203 | 1032 | 941 | 846 | 730 | 589 | 2529 | 2535 | 2409 | 1835 | 1938 | 1602 | 2271.79 | 2491 | 1792 | 0.773 | -0.371 | 0.00151174 |
| AT1G30890.1 | | protein_coding | Integral membrane HRF1 | 296 | 250 | 222 | 233 | 181 | 168 | 622 | 614 | 568 | 505 | 481 | 457 | 536.25 | 601 | 481 | 0.773 | -0.371 | 0.00038384 |
| AT3G61150.1 | HDG1 | protein_coding | homeodomain GLABROU | 475 | 456 | 371 | 324 | 262 | 360 | 998 | 1120 | 950 | 703 | 696 | 979 | 812.83 | 1023 | 793 | 0.773 | -0.372 | 0.00491451 |
| AT4G30060.1 | | protein_coding | Core-2/I-branching beta- | 233 | 181 | 171 | 204 | 148 | 113 | 490 | 445 | 438 | 442 | 393 | 307 | 368.09 | 458 | 381 | 0.773 | -0.372 | 0.0044863 |
| AT3G09740.1 | SYP71 | protein_coding | syntaxin of plants 71 | 1280 | 1074 | 1184 | 959 | 773 | 908 | 2691 | 2639 | 3031 | 2080 | 2052 | 2469 | 2435.53 | 2787 | 2200 | 0.772 | -0.372 | 0.00267063 |
| AT5G10560.1 | BXL6 | protein_coding | Glycosyl hydrolase family | 1471 | 1367 | 1302 | 819 | 789 | 806 | 3092 | 3358 | 3333 | 1776 | 2095 | 2192 | 2318.5 | 3261 | 2021 | 0.773 | -0.372 | 0.00192013 |
| AT2G47320.1 | CYP21-3 | protein_coding | Cyclophilin-like peptidyl- | 697 | 586 | 644 | 604 | 454 | 421 | 1465 | 1440 | 1649 | 1310 | 1205 | 1145 | 1405.44 | 1518 | 1220 | 0.773 | -0.372 | 0.00160194 |
| AT2G26570.1 | WEB1 | protein_coding | Plant protein of unknowr | 909 | 890 | 813 | 738 | 617 | 588 | 1911 | 2187 | 2081 | 1601 | 1638 | 1599 | 1541.07 | 2060 | 1613 | 0.773 | -0.372 | 0.00126073 |
| AT1G16858.1 | CPuORF55 | protein_coding | conserved peptide upstre | 333 | 335 | 341 | 329 | 257 | 262 | 700 | 823 | 873 | 714 | 682 | 712 | 709.21 | 799 | 703 | 0.773 | -0.372 | 0.00123985 |
| AT1G69850.1 | NPF4.6 | protein_coding | nitrate transporter 1:2 | 716 | 476 | 491 | 505 | 362 | 321 | 1505 | 1169 | 1257 | 1095 | 961 | 873 | 1031.54 | 1310 | 976 | 0.772 | -0.373 | 0.00551661 |
| AT4G16760.1 | ACX1 | protein_coding | acyl-CoA oxidase 1 | 1308 | 922 | 899 | 1092 | 705 | 945 | 2750 | 2265 | 2301 | 2369 | 1872 | 2570 | 2242.13 | 2439 | 2270 | 0.772 | -0.373 | 0.0054985 |
| AT5G55230.2 | MAP65-1 | protein_coding | microtubule-associated p | 2099 | 1924 | 1837 | 1443 | 1346 | 1242 | 4412 | 4727 | 4703 | 3130 | 3573 | 3378 | 3785.98 | 4614 | 3360 | 0.772 | -0.373 | 0.00098296 |
| AT4G17140.3 | | protein_coding | pleckstrin homology (PH) | 4162 | 3529 | 4075 | 3625 | 2756 | 2775 | 8749 | 8670 | 10432 | 7863 | 7317 | 7546 | 5426.8 | 9284 | 7575 | 0.772 | -0.374 | 0.00276453 |
| AT1G73390.1 | | protein_coding | Endosomal targeting BRC | 417 | 346 | 314 | 336 | 281 | 215 | 877 | 850 | 804 | 729 | 746 | 585 | 668.33 | 844 | 687 | 0.772 | -0.374 | 0.00271641 |
| AT1G50440.1 | | protein_coding | RING/FYVE/PHD zinc fing | 319 | 301 | 279 | 227 | 224 | 235 | 671 | 740 | 714 | 492 | 595 | 639 | 649.64 | 708 | 575 | 0.771 | -0.374 | 0.00266291 |
| AT2G25310.1 | | protein_coding | Protein of unknown funct | 384 | 336 | 381 | 352 | 289 | 295 | 807 | 825 | 975 | 764 | 767 | 802 | 859.13 | 869 | 778 | 0.772 | -0.374 | 0.00093505 |
| AT1G64510.1 | RPS6 | protein_coding | Translation elongation fa | 4952 | 5434 | 4559 | 4035 | 3453 | 4111 | 10409 | 13350 | 11671 | 8752 | 9167 | 11180 | 13212.42 | 11810 | 9700 | 0.771 | -0.375 | 0.00638772 |
| AT5G55990.1 | CBL2 | protein_coding | calcineurin B-like protein | 338 | 350 | 364 | 245 | 277 | 211 | 711 | 860 | 932 | 531 | 735 | 574 | 884.5 | 834 | 613 | 0.771 | -0.375 | 0.0057332 |
| AT4G34140.1 | | protein_coding | D111/G-patch domain-co | 289 | 250 | 264 | 188 | 189 | 137 | 607 | 614 | 676 | 408 | 502 | 373 | 533.5 | 632 | 428 | 0.771 | -0.375 | 0.00181884 |
| AT5G01220.1 | SQD2 | protein_coding | sulfoquinovosyldiacylglyc | 1049 | 674 | 618 | 818 | 581 | 476 | 2205 | 1656 | 1582 | 1774 | 1542 | 1294 | 1929.53 | 1814 | 1537 | 0.771 | -0.376 | 0.0084781 |
| AT2G04270.5 | RNE | protein_coding | RNase E/G-like | 1142 | 878 | 808 | 613 | 554 | 388 | 2401 | 2157 | 2069 | 1330 | 1471 | 1055 | 1646.15 | 2209 | 1285 | 0.77 | -0.376 | 0.00349811 |
| AT3G53000.1 | PP2A15 | protein_coding | phloem protein 2-A15 | 235 | 200 | 224 | 201 | 173 | 166 | 494 | 491 | 573 | 436 | 459 | 451 | 486.05 | 519 | 449 | 0.771 | -0.376 | 0.00211977 |
| AT1G22620.1 | SAC1 | protein_coding | Phosphoinositide phosph | 630 | 601 | 582 | 467 | 380 | 391 | 1324 | 1477 | 1490 | 1013 | 1009 | 1063 | 1102.11 | 1430 | 1028 | 0.771 | -0.376 | 0.00101713 |
| AT1G27630.1 | CYCT1-3 | protein_coding | cyclin T 1;3 | 297 | 361 | 395 | 239 | 198 | 222 | 624 | 887 | 1011 | 518 | 526 | 604 | 866.22 | 841 | 549 | 0.77 | -0.377 | 0.00888367 |
| AT3G30380.1 | | protein_coding | alpha/beta-Hydrolases su | 174 | 181 | 196 | 131 | 125 | 133 | 366 | 445 | 502 | 284 | 332 | 362 | 357.2 | 438 | 326 | 0.77 | -0.377 | 0.00657005 |
| AT1G05720.1 | | protein_coding | selenoprotein family prot | 224 | 193 | 232 | 193 | 160 | 167 | 471 | 474 | 594 | 419 | 425 | 454 | 464.47 | 513 | 433 | 0.77 | -0.377 | 0.00221884 |
| AT1G31410.1 | | protein_coding | putrescine-binding peripl | 441 | 485 | 398 | 357 | 318 | 336 | 927 | 1192 | 1019 | 774 | 844 | 914 | 1010.76 | 1046 | 844 | 0.77 | -0.377 | 0.0019341 |
| AT1G35340.1 | | protein_coding | ATP-dependent protease | 847 | 677 | 781 | 575 | 456 | 446 | 1780 | 1663 | 1999 | 1247 | 1211 | 1213 | 1829.14 | 1814 | 1224 | 0.77 | -0.377 | 0.00123695 |
| AT1G61030.1 | | protein_coding | WAPL (Wings apart-like p | 319 | 264 | 258 | 189 | 195 | 183 | 671 | 649 | 660 | 410 | 518 | 498 | 439.63 | 660 | 475 | 0.77 | -0.377 | 0.00103454 |
| AT2G24395.1 | | protein_coding | chaperone protein dnaJ-r | 223 | 210 | 193 | 151 | 116 | 112 | 469 | 516 | 494 | 328 | 308 | 305 | 488.43 | 493 | 314 | 0.77 | -0.377 | 0.00037817 |
| AT5G13270.1 | PCMP-H90 | protein_coding | Pentatricopeptide repeat | 219 | 196 | 160 | 209 | 170 | 163 | 460 | 482 | 410 | 453 | 451 | 443 | 367.79 | 451 | 449 | 0.769 | -0.378 | 0.00346418 |
| AT2G44660.1 | | protein_coding | ALG6, ALG8 glycosyltrans | 169 | 136 | 116 | 136 | 99 | 89 | 355 | 334 | 297 | 295 | 263 | 242 | 327.58 | 329 | 267 | 0.77 | -0.378 | 0.00253362 |
| AT3G24140.1 | FAMA | protein_coding | basic helix-loop-helix (bH | 246 | 233 | 250 | 210 | 176 | 193 | 517 | 572 | 640 | 456 | 467 | 525 | 491.59 | 576 | 483 | 0.769 | -0.379 | 0.00589192 |
| AT3G60190.1 | DRP1E | protein_coding | DYNAMIN-like 1E | 952 | 993 | 1025 | 723 | 693 | 733 | 2001 | 2440 | 2624 | 1568 | 1840 | 1993 | 1870.93 | 2355 | 1800 | 0.769 | -0.379 | 0.00407377 |
| AT4G04840.1 | MSRB6 | protein_coding | methionine sulfoxide red | 1205 | 1236 | 884 | 1556 | 1124 | 1021 | 2533 | 3037 | 2263 | 3375 | 2984 | 2777 | 3565.08 | 2611 | 3045 | 0.769 | -0.379 | 0.00402384 |
| AT2G01260.1 | | protein_coding | Protein of unknown funct | 376 | 361 | 314 | 351 | 261 | 296 | 790 | 887 | 804 | 761 | 693 | 805 | 833.63 | 827 | 753 | 0.769 | -0.379 | 0.00080554 |
| AT1G07420.1 | SMO2-2 | protein_coding | sterol 4-alpha-methyl-oxi | 421 | 313 | 415 | 333 | 251 | 235 | 885 | 769 | 1062 | 722 | 666 | 639 | 1152.39 | 905 | 676 | 0.769 | -0.38 | 0.00358048 |
| AT1G76700.1 | ATJ10 | protein_coding | DNAJ heat shock N-termi | 279 | 196 | 226 | 219 | 163 | 196 | 586 | 482 | 579 | 475 | 433 | 533 | 434.35 | 549 | 480 | 0.769 | -0.38 | 0.00326727 |
| AT3G58460.2 | RBL15 | protein_coding | RHOMBOID-like protein 1 | 361 | 341 | 320 | 320 | 280 | 295 | 759 | 838 | 819 | 694 | 743 | 802 | 881.95 | 805 | 746 | 0.768 | -0.38 | 0.00109527 |
| AT3G25660.1 | GATA | protein_coding | Amidase family protein | 2248 | 1948 | 1646 | 1389 | 1308 | 1234 | 4725 | 4786 | 4214 | 3013 | 3472 | 3356 | 5333.23 | 4575 | 3280 | 0.768 | -0.38 | 0.00077343 |
| AT4G19530.1 | | protein_coding | disease resistance proteir | 4843 | 3822 | 3263 | 2832 | 2126 | 1616 | 10180 | 9390 | 8353 | 6143 | 5644 | 4395 | 5706.19 | 9308 | 5394 | 0.768 | -0.381 | 0.00401574 |
| AT1G14230.1 | APY4 | protein_coding | GDA1/CD39 nucleoside p | 233 | 200 | 231 | 216 | 200 | 234 | 490 | 491 | 591 | 469 | 531 | 636 | 492.24 | 524 | 545 | 0.767 | -0.382 | 0.00368414 |
| AT5G03900.2 | | protein_coding | Iron-sulphur cluster biosy | 1128 | 917 | 1016 | 799 | 709 | 704 | 2371 | 2253 | 2601 | 1733 | 1882 | 1914 | 2149.32 | 2408 | 1843 | 0.767 | -0.382 | 0.00116794 |
| AT3G15920.1 | | protein_coding | Phox (PX) domain-contai | 434 | 348 | 383 | 304 | 267 | 294 | 912 | 855 | 981 | 659 | 709 | 800 | 706.86 | 916 | 723 | 0.768 | -0.382 | 0.00092583 |
| AT5G16210.1 | | protein_coding | HEAT repeat-containing p | 899 | 727 | 729 | 726 | 569 | 584 | 1890 | 1786 | 1866 | 1575 | 1511 | 1588 | 1418.96 | 1847 | 1558 | 0.767 | -0.382 | 0.00086525 |
| AT4G16180.2 | | protein_coding | | 716 | 631 | 655 | 585 | 461 | 472 | 1505 | 1550 | 1677 | 1269 | 1224 | 1284 | 1234.59 | 1577 | 1259 | 0.767 | -0.383 | 0.00080847 |
| AT5G02240.1 | | protein_coding | NAD(P)-binding Rossman | 2256 | 2042 | 1836 | 1815 | 1556 | 1711 | 4742 | 5017 | 4700 | 3937 | 4131 | 4653 | 5440.29 | 4820 | 4240 | 0.767 | -0.383 | 0.00058566 |
| AT4G22580.1 | | protein_coding | Exostosin family protein | 135 | 131 | 115 | 129 | 74 | 95 | 284 | 322 | 294 | 280 | 196 | 258 | 292.84 | 300 | 245 | 0.766 | -0.384 | 0.00666974 |
| AT3G15070.2 | | protein_coding | Phosphoglycerate mutase | 131 | 130 | 116 | 102 | 106 | 85 | 275 | 319 | 297 | 221 | 281 | 231 | 258.26 | 297 | 244 | 0.766 | -0.384 | 0.0058234 |
| AT3G61700.2 | | protein_coding | Plant protein 1589 of unl | 623 | 394 | 417 | 453 | 374 | 401 | 1310 | 968 | 1068 | 983 | 993 | 1090 | 1248.04 | 1115 | 1022 | 0.766 | -0.384 | 0.00452224 |
| AT2G17540.2 | | protein_coding | | 244 | 264 | 249 | 145 | 169 | 162 | 513 | 649 | 637 | 315 | 449 | 441 | 534.34 | 600 | 402 | 0.766 | -0.384 | 0.00434913 |
| AT3G15880.1 | | protein_coding | | 264 | 209 | 223 | 158 | 114 | 145 | 555 | 513 | 571 | 343 | 303 | 394 | 427.18 | 546 | 347 | 0.766 | -0.384 | 0.00118926 |
| AT1G78280.1 | | protein_coding | transferases, transferring | 472 | 405 | 455 | 466 | 329 | 337 | 992 | 995 | 1165 | 1011 | 873 | 916 | 798.25 | 1051 | 933 | 0.766 | -0.384 | 0.00063446 |
| AT1G76260.1 | DWA2 | protein_coding | DWD (DDB1-binding WD- | 158 | 142 | 143 | 90 | 86 | 90 | 332 | 349 | 366 | 195 | 228 | 245 | 311.63 | 349 | 223 | 0.766 | -0.385 | 0.00196499 |
| AT5G06830.1 | | protein_coding | | 501 | 421 | 504 | 355 | 334 | 298 | 1053 | 1034 | 1290 | 770 | 887 | 810 | 913.38 | 1126 | 822 | 0.766 | -0.385 | 0.00118195 |
| AT3G55530.1 | SDIR1 | protein_coding | RING/U-box superfamily | 112 | 111 | 111 | 95 | 98 | 90 | 235 | 273 | 284 | 206 | 260 | 245 | 274.03 | 264 | 237 | 0.765 | -0.386 | 0.00543403 |
| AT4G35740.1 | RECQL3 | protein_coding | DEAD/DEAH box RNA hel | 184 | 151 | 165 | 128 | 103 | 125 | 387 | 371 | 422 | 278 | 273 | 340 | 307.48 | 393 | 297 | 0.765 | -0.386 | 0.00218506 |
| AT2G03340.1 | WRKY3 | protein_coding | WRKY DNA-binding prote | 656 | 649 | 600 | 463 | 395 | 359 | 1379 | 1594 | 1536 | 1004 | 1049 | 976 | 1232.64 | 1503 | 1010 | 0.765 | -0.386 | 0.00139785 |
| AT1G45050.1 | UBC15 | protein_coding | Ubiquitin-conjugating en | 130 | 108 | 118 | 117 | 82 | 92 | 273 | 265 | 302 | 254 | 218 | 250 | 255.55 | 280 | 241 | 0.765 | -0.387 | 0.00529761 |
| AT1G18485.1 | PCMP-H8 | protein_coding | Pentatricopeptide repeat | 266 | 228 | 200 | 161 | 142 | 131 | 559 | 560 | 512 | 349 | 377 | 356 | 441.38 | 544 | 361 | 0.765 | -0.387 | 0.00094909 |
| AT1G09270.1 | IMPA4 | protein_coding | importin alpha isoform 4 | 853 | 754 | 736 | 690 | 579 | 648 | 1793 | 1852 | 1884 | 1497 | 1537 | 1762 | 1714.67 | 1843 | 1599 | 0.765 | -0.387 | 0.00075002 |
| AT5G62100.1 | BAG2 | protein_coding | BCL-2-associated athanog | 208 | 165 | 124 | 177 | 146 | 98 | 437 | 405 | 317 | 384 | 388 | 267 | 371.74 | 386 | 346 | 0.764 | -0.388 | 0.0070114 |
| AT2G43560.1 | FKBP16-3 | protein_coding | FKBP-like peptidyl-prolyl | 2424 | 2749 | 2379 | 1837 | 1736 | 1789 | 5095 | 6754 | 6090 | 3985 | 4609 | 4865 | 6438.63 | 5980 | 4486 | 0.764 | -0.388 | 0.0025638 |
| AT5G39040.1 | ABCB27 | protein_coding | transporter associated wi | 533 | 475 | 451 | 394 | 328 | 269 | 1120 | 1167 | 1155 | 855 | 871 | 732 | 926.87 | 1147 | 819 | 0.764 | -0.388 | 0.00056138 |
| AT4G03080.1 | BSL1 | protein_coding | BRI1 suppressor 1 (BSU1) | 1201 | 1117 | 1000 | 1003 | 875 | 862 | 2525 | 2744 | 2560 | 2176 | 2323 | 2344 | 2338.24 | 2610 | 2281 | 0.764 | -0.388 | 0.00045962 |
| AT1G31020.1 | ATO2 | protein_coding | thioredoxin O2 | 103 | 64 | 89 | 91 | 75 | 62 | 217 | 157 | 228 | 197 | 199 | 169 | 216.28 | 201 | 188 | 0.764 | -0.389 | 0.0094583 |
| AT3G21820.1 | ATXR2 | protein_coding | histone-lysine N-methyltr | 292 | 211 | 180 | 220 | 167 | 134 | 614 | 518 | 461 | 477 | 443 | 364 | 478.92 | 531 | 428 | 0.764 | -0.389 | 0.00378983 |
| AT5G47650.1 | NUDT2 | protein_coding | nudix hydrolase homolog | 168 | 167 | 167 | 178 | 119 | 139 | 353 | 410 | 428 | 386 | 316 | 378 | 398.98 | 397 | 360 | 0.764 | -0.389 | 0.00210024 |
| AT4G31080.2 | | protein_coding | Protein of unknown funct | 581 | 455 | 542 | 397 | 336 | 432 | 1221 | 1118 | 1388 | 861 | 892 | 1175 | 1060.29 | 1242 | 976 | 0.763 | -0.39 | 0.00533436 |
| AT1G53800.2 | | protein_coding | | 1341 | 1089 | 864 | 900 | 707 | 545 | 2819 | 2675 | 2212 | 1952 | 1877 | 1482 | 2316.61 | 2569 | 1770 | 0.763 | -0.39 | 0.00308274 |
| AT5G51340.1 | | protein_coding | Tetratricopeptide repeat | 303 | 259 | 224 | 256 | 217 | 215 | 637 | 636 | 573 | 555 | 576 | 585 | 690.15 | 615 | 572 | 0.763 | -0.39 | 0.00015048 |
| AT3G19850.1 | | protein_coding | Phototropic-responsive N | 765 | 541 | 531 | 504 | 313 | 359 | 1608 | 1329 | 1359 | 1093 | 831 | 976 | 939.64 | 1432 | 967 | 0.763 | -0.391 | 0.00247701 |
| AT2G22530.1 | | protein_coding | Alkaline-phosphatase-like | 186 | 189 | 178 | 198 | 150 | 134 | 391 | 464 | 456 | 429 | 398 | 364 | 376.99 | 437 | 397 | 0.763 | -0.391 | 0.00215876 |
| AT5G53000.1 | TAP46 | protein_coding | 2A phosphatase associate | 506 | 450 | 470 | 383 | 320 | 340 | 1064 | 1106 | 1203 | 831 | 850 | 925 | 987.97 | 1124 | 869 | 0.763 | -0.391 | 0.0008096 |
| AT5G16540.1 | ZFN3 | protein_coding | zinc finger nuclease 3 | 434 | 368 | 393 | 394 | 351 | 236 | 912 | 904 | 1006 | 855 | 932 | 642 | 1148.97 | 941 | 810 | 0.762 | -0.392 | 0.00362452 |
| AT3G56010.1 | | protein_coding | | 1309 | 1105 | 997 | 1023 | 772 | 693 | 2752 | 2715 | 2552 | 2219 | 2049 | 1885 | 2441.04 | 2673 | 2051 | 0.762 | -0.392 | 0.00065984 |
| AT4G19400.1 | | protein_coding | Profilin family protein | 100 | 79 | 83 | 58 | 44 | 44 | 210 | 194 | 212 | 126 | 117 | 120 | 177.18 | 205 | 121 | 0.761 | -0.393 | 0.00758736 |
| AT5G48840.1 | PANC | protein_coding | homolog of bacterial PAN | 241 | 211 | 158 | 166 | 127 | 81 | 507 | 518 | 404 | 360 | 337 | 220 | 447.4 | 476 | 306 | 0.761 | -0.393 | 0.0046471 |
| AT1G67660.1 | | protein_coding | Restriction endonuclease | 389 | 361 | 319 | 245 | 235 | 185 | 818 | 887 | 817 | 531 | 624 | 503 | 840.48 | 841 | 553 | 0.762 | -0.393 | 0.00043797 |
| AT2G46070.1 | MPK12 | protein_coding | mitogen-activated protei | 151 | 167 | 167 | 182 | 154 | 156 | 317 | 410 | 428 | 395 | 409 | 424 | 465.38 | 385 | 409 | 0.761 | -0.394 | 0.00576106 |
| AT2G14285.1 | | protein_coding | Small nuclear ribonucleo | 178 | 140 | 104 | 126 | 94 | 81 | 374 | 344 | 266 | 273 | 250 | 220 | 372.06 | 328 | 248 | 0.761 | -0.394 | 0.00444226 |
| AT1G78995.1 | | protein_coding | | 389 | 430 | 400 | 268 | 230 | 173 | 818 | 1056 | 1024 | 581 | 611 | 470 | 866.59 | 966 | 554 | 0.761 | -0.394 | 0.00286918 |
| AT5G13390.1 | NEF1 | protein_coding | no exine formation 1 | 722 | 558 | 510 | 567 | 480 | 402 | 1518 | 1371 | 1306 | 1230 | 1274 | 1093 | 1285.72 | 1398 | 1199 | 0.761 | -0.394 | 0.00126873 |
| AT5G05560.1 | APC1 | protein_coding | E3 ubiquitin ligase, putat | 442 | 357 | 298 | 377 | 329 | 273 | 929 | 877 | 763 | 818 | 873 | 742 | 669.24 | 856 | 811 | 0.761 | -0.394 | 0.00068912 |
| AT3G48820.1 | | protein_coding | Glycosyltransferase famil | 307 | 267 | 270 | 245 | 186 | 198 | 645 | 656 | 691 | 531 | 494 | 538 | 519.48 | 664 | 521 | 0.76 | -0.395 | 0.00045903 |
| AT5G60940.1 | CSTF50 | protein_coding | Transducin/WD40 repeat | 350 | 268 | 254 | 249 | 203 | 186 | 736 | 658 | 650 | 540 | 539 | 506 | 616.8 | 681 | 528 | 0.76 | -0.395 | 0.00014882 |
| AT4G24730.1 | NRT3.2 | protein_coding | Calcineurin-like metallo-p | 142 | 127 | 118 | 150 | 104 | 101 | 298 | 312 | 302 | 325 | 276 | 275 | 311.26 | 304 | 292 | 0.76 | -0.396 | 0.0073476 |
| AT5G18590.1 | | protein_coding | Galactose oxidase/kelch r | 981 | 879 | 934 | 751 | 650 | 642 | 2062 | 2160 | 2391 | 1629 | 1726 | 1746 | 1801.87 | 2204 | 1700 | 0.76 | -0.396 | 0.0005753 |
| AT4G23270.1 | CRK19 | protein_coding | cysteine-rich RLK (RECEPT | 853 | 834 | 755 | 557 | 401 | 347 | 1793 | 2049 | 1933 | 1208 | 1065 | 944 | 1677.19 | 1925 | 1072 | 0.76 | -0.397 | 0.00175228 |
| AT4G26965.1 | | protein_coding | NADH:ubiquinone oxidor | 112 | 86 | 94 | 85 | 45 | 50 | 235 | 211 | 241 | 184 | 119 | 136 | 201.78 | 229 | 146 | 0.759 | -0.398 | 0.00754996 |
| AT1G32790.2 | CID11 | protein_coding | CTC-interacting domain 1 | 783 | 819 | 783 | 702 | 555 | 706 | 1646 | 2012 | 2005 | 1523 | 1473 | 1920 | 2019.21 | 1888 | 1639 | 0.759 | -0.398 | 0.00212073 |
| AT5G02290.1 | NAK | protein_coding | Protein kinase superfamil | 395 | 324 | 344 | 275 | 186 | 251 | 830 | 796 | 881 | 597 | 494 | 683 | 636.13 | 836 | 591 | 0.759 | -0.398 | 0.00096443 |
| AT4G30580.1 | LPAT1 | protein_coding | Phospholipid/glycerol acy | 1176 | 902 | 929 | 752 | 638 | 610 | 2472 | 2216 | 2378 | 1631 | 1694 | 1659 | 1951.31 | 2355 | 1661 | 0.759 | -0.398 | 0.00044688 |
| AT1G49750.1 | | protein_coding | Leucine-rich repeat (LRR) | 720 | 568 | 445 | 649 | 570 | 388 | 1513 | 1395 | 1139 | 1408 | 1513 | 1055 | 2212.15 | 1349 | 1325 | 0.759 | -0.399 | 0.00552411 |
| AT5G02380.1 | MT2B | protein_coding | metallothionein 2B | 8162 | 8374 | 7071 | 9466 | 7229 | 7867 | 17157 | 20573 | 18102 | 20533 | 19191 | 21394 | 25743.17 | 18611 | 20373 | 0.758 | -0.399 | 0.00448765 |
| AT2G23770.1 | LYK4 | protein_coding | protein kinase family pro | 138 | 117 | 112 | 109 | 101 | 81 | 290 | 287 | 287 | 236 | 268 | 220 | 247.38 | 288 | 241 | 0.759 | -0.399 | 0.00321752 |
| AT5G46020.1 | | protein_coding | | 1564 | 1580 | 1566 | 1135 | 978 | 1175 | 3288 | 3882 | 4009 | 2462 | 2596 | 3195 | 3317.13 | 3726 | 2751 | 0.758 | -0.399 | 0.00181505 |
| AT1G66980.1 | GDPDL2 | protein_coding | suppressor of npr1-1 con | 2257 | 1960 | 1912 | 1203 | 1285 | 1039 | 4744 | 4815 | 4895 | 2609 | 3411 | 2825 | 3310.97 | 4818 | 2948 | 0.758 | -0.399 | 0.00120976 |
| AT2G38980.1 | | pseudogene | | 178 | 169 | 189 | 110 | 114 | 88 | 374 | 415 | 484 | 239 | 303 | 239 | 367.24 | 424 | 260 | 0.758 | -0.4 | 0.00991584 |
| AT1G55730.1 | CAX5 | protein_coding | cation exchanger 5 | 144 | 120 | 141 | 174 | 124 | 93 | 303 | 295 | 361 | 377 | 329 | 253 | 299.56 | 320 | 320 | 0.758 | -0.4 | 0.00552939 |
| AT2G38260.1 | | pseudogene | | 297 | 248 | 301 | 192 | 223 | 201 | 624 | 609 | 771 | 416 | 592 | 547 | 490.55 | 668 | 518 | 0.758 | -0.4 | 0.00328114 |
| AT4G35560.2 | | protein_coding | Transducin/WD40 repeat | 232 | 210 | 229 | 156 | 167 | 154 | 488 | 516 | 586 | 338 | 443 | 419 | 377.49 | 530 | 400 | 0.758 | -0.4 | 0.00298023 |
| AT5G25980.2 | TGG2 | protein_coding | glucoside glucohydrolase | 1158 | 1311 | 1077 | 857 | 782 | 777 | 2434 | 3221 | 2757 | 1859 | 2076 | 2113 | 2854.9 | 2804 | 2016 | 0.758 | -0.4 | 0.00247711 |
| AT1G31360.1 | MED34 | protein_coding | RECQ helicase L2 | 245 | 169 | 189 | 128 | 97 | 126 | 515 | 415 | 484 | 278 | 258 | 343 | 335.24 | 471 | 293 | 0.757 | -0.401 | 0.00798393 |
| AT5G39760.1 | ZHD10 | protein_coding | homeobox protein 23 | 74 | 79 | 47 | 43 | 33 | 31 | 156 | 194 | 120 | 93 | 88 | 84 | 374.12 | 157 | 88 | 0.757 | -0.401 | 0.00186071 |
| AT3G57170.1 | | protein_coding | N-acetylglucosaminyl tra | 400 | 312 | 339 | 242 | 261 | 211 | 841 | 767 | 868 | 525 | 693 | 574 | 762.04 | 825 | 597 | 0.757 | -0.401 | 0.00130539 |
| AT2G18290.1 | APC10 | protein_coding | anaphase promoting com | 575 | 459 | 407 | 381 | 309 | 273 | 1209 | 1128 | 1042 | 826 | 820 | 742 | 1028.23 | 1126 | 796 | 0.757 | -0.401 | 0.00047374 |
| AT2G43200.1 | | protein_coding | S-adenosyl-L-methionine- | 191 | 199 | 124 | 98 | 108 | 65 | 401 | 489 | 317 | 213 | 287 | 177 | 435.34 | 402 | 226 | 0.757 | -0.402 | 0.00801329 |
| AT5G07840.1 | | protein_coding | Ankyrin repeat family pro | 52 | 60 | 52 | 42 | 30 | 42 | 109 | 147 | 133 | 91 | 80 | 114 | 210.4 | 130 | 95 | 0.757 | -0.402 | 0.00582107 |
| AT3G50920.1 | LPPE1 | protein_coding | Phosphatidic acid phosph | 152 | 121 | 103 | 159 | 89 | 94 | 320 | 297 | 264 | 345 | 236 | 256 | 335.63 | 294 | 279 | 0.757 | -0.402 | 0.00330331 |
| AT1G55880.1 | | protein_coding | Pyridoxal-5'-phosphate-d | 155 | 109 | 117 | 100 | 82 | 82 | 326 | 268 | 300 | 217 | 218 | 223 | 238.37 | 298 | 219 | 0.757 | -0.402 | 0.00314284 |
| AT3G07940.1 | AGD11 | protein_coding | Calcium-dependent ARF-t | 231 | 224 | 217 | 160 | 178 | 121 | 486 | 550 | 556 | 347 | 473 | 329 | 504.56 | 531 | 383 | 0.757 | -0.402 | 0.00237256 |
| AT5G15050.1 | | protein_coding | Core-2/I-branching beta- | 299 | 298 | 265 | 238 | 219 | 185 | 629 | 732 | 678 | 516 | 581 | 503 | 696.14 | 680 | 533 | 0.756 | -0.403 | 0.00032357 |
| AT5G23920.1 | | protein_coding | | 606 | 501 | 379 | 533 | 349 | 309 | 1274 | 1231 | 970 | 1156 | 927 | 840 | 1293.37 | 1158 | 974 | 0.756 | -0.404 | 0.00397444 |
| AT1G10200.1 | WLIM1 | protein_coding | GATA type zinc finger tra | 1387 | 1380 | 1305 | 1120 | 1024 | 930 | 2916 | 3390 | 3341 | 2429 | 2718 | 2529 | 3299.98 | 3216 | 2559 | 0.756 | -0.404 | 0.00107037 |
| AT5G01460.1 | | protein_coding | LMBR1-like membrane pr | 549 | 524 | 557 | 478 | 400 | 441 | 1154 | 1287 | 1426 | 1037 | 1062 | 1199 | 1362.64 | 1289 | 1099 | 0.756 | -0.404 | 0.00092326 |
| AT1G73760.1 | | protein_coding | RING/U-box superfamily | 449 | 460 | 408 | 320 | 280 | 255 | 944 | 1130 | 1045 | 694 | 743 | 693 | 961.14 | 1040 | 710 | 0.756 | -0.404 | 0.00054369 |
| AT1G10050.1 | | protein_coding | glycosyl hydrolase family | 104 | 100 | 88 | 74 | 58 | 49 | 219 | 246 | 225 | 161 | 154 | 133 | 162.66 | 230 | 149 | 0.755 | -0.405 | 0.0098003 |
| AT5G22320.1 | | protein_coding | Leucine-rich repeat (LRR) | 332 | 224 | 251 | 237 | 144 | 124 | 698 | 550 | 643 | 514 | 382 | 337 | 462.03 | 630 | 411 | 0.755 | -0.405 | 0.00600593 |
| AT1G28250.1 | | protein_coding | | 194 | 165 | 168 | 189 | 153 | 129 | 408 | 405 | 430 | 410 | 406 | 351 | 394.07 | 414 | 389 | 0.755 | -0.405 | 0.00194242 |
| AT1G02300.1 | | protein_coding | Cysteine proteinases supe | 372 | 343 | 323 | 273 | 285 | 220 | 782 | 843 | 827 | 592 | 757 | 598 | 790.38 | 817 | 649 | 0.755 | -0.405 | 0.00094843 |
| AT3G50500.2 | SRK2D | protein_coding | SNF1-related protein kina | 1674 | 1756 | 1926 | 1135 | 1117 | 1244 | 3519 | 4314 | 4931 | 2462 | 2965 | 3383 | 3799.57 | 4255 | 2937 | 0.755 | -0.406 | 0.00545014 |
| AT5G02030.1 | BLH9 | protein_coding | POX (plant homeobox) fa | 50 | 53 | 39 | 42 | 44 | 43 | 105 | 130 | 100 | 91 | 117 | 117 | 174.3 | 112 | 108 | 0.754 | -0.407 | 0.00324382 |

| | | | | | | | | | | | | | | | | | | | | | |
|---|---|---|---|---|---|---|---|---|---|---|---|---|---|---|---|---|---|---|---|---|---|
| AT2G46090.1 | LCKB2 | protein_coding | Diacylglycerol kinase fam | 366 | 216 | 242 | 230 | 180 | 175 | 769 | 531 | 620 | 499 | 478 | 476 | 617.53 | 640 | 484 | 0.754 | -0.407 | 0.0019866 |
| AT1G02145.3 | ALG12 | protein_coding | homolog of asparagine-li | 376 | 271 | 274 | 202 | 168 | 160 | 790 | 666 | 701 | 438 | 446 | 435 | 595.08 | 719 | 440 | 0.754 | -0.407 | 0.00090695 |
| AT1G60630.1 | | protein_coding | Leucine-rich repeat prote | 89 | 114 | 106 | 90 | 80 | 70 | 187 | 280 | 271 | 195 | 212 | 190 | 222.77 | 246 | 199 | 0.754 | -0.408 | 0.00859913 |
| AT2G44420.1 | | protein_coding | protein N-terminal aspara | 119 | 138 | 112 | 103 | 80 | 76 | 250 | 339 | 287 | 223 | 212 | 207 | 241.67 | 292 | 214 | 0.754 | -0.408 | 0.00691087 |
| AT2G40380.1 | PRA1B2 | protein_coding | prenylated RAB acceptor | 79 | 65 | 45 | 57 | 46 | 51 | 166 | 160 | 115 | 124 | 122 | 139 | 412.04 | 147 | 128 | 0.754 | -0.408 | 0.00094365 |
| AT3G52840.1 | BGAL2 | protein_coding | beta-galactosidase 2 | 3014 | 2888 | 2860 | 2205 | 2043 | 1964 | 6336 | 7095 | 7322 | 4783 | 5424 | 5341 | 5666.29 | 6918 | 5183 | 0.754 | -0.408 | 0.00053133 |
| AT2G38670.1 | PECT1 | protein_coding | phosphorylethanolamine | 838 | 828 | 770 | 571 | 501 | 528 | 1762 | 2034 | 1971 | 1239 | 1330 | 1436 | 1944.29 | 1922 | 1335 | 0.753 | -0.408 | 0.00043286 |
| AT3G52730.1 | | protein_coding | ubiquinol-cytochrome C r | 1068 | 772 | 696 | 795 | 581 | 631 | 2245 | 1897 | 1782 | 1724 | 1542 | 1716 | 2121.37 | 1975 | 1661 | 0.753 | -0.409 | 0.00112521 |
| AT3G55420.1 | | protein_coding | | 107 | 95 | 99 | 98 | 84 | 99 | 225 | 233 | 253 | 213 | 223 | 269 | 243.31 | 237 | 235 | 0.753 | -0.41 | 0.00236473 |
| AT2G34720.1 | NFYA4 | protein_coding | nuclear factor Y, subunit | 268 | 245 | 222 | 215 | 185 | 203 | 563 | 602 | 568 | 466 | 491 | 552 | 522.13 | 578 | 503 | 0.753 | -0.41 | 0.00052767 |
| AT1G80940.1 | | protein_coding | | 483 | 468 | 420 | 403 | 303 | 330 | 1015 | 1150 | 1075 | 874 | 804 | 897 | 984.1 | 1080 | 858 | 0.753 | -0.41 | 0.00035187 |
| AT5G63880.2 | VPS20.1 | protein_coding | SNF7 family protein | 213 | 207 | 261 | 173 | 123 | 152 | 448 | 509 | 668 | 375 | 327 | 413 | 440.59 | 542 | 372 | 0.752 | -0.411 | 0.00848517 |
| AT5G27380.1 | GSH2 | protein_coding | glutathione synthetase 2 | 2294 | 1726 | 1821 | 1506 | 1042 | 1016 | 4822 | 4240 | 4662 | 3267 | 2766 | 2763 | 3953.32 | 4575 | 2932 | 0.752 | -0.411 | 0.00053849 |
| AT4G33530.1 | POT13 | protein_coding | K+ uptake permease 5 | 1686 | 1460 | 1299 | 1274 | 1084 | 913 | 3544 | 3587 | 3326 | 2763 | 2878 | 2483 | 2866.42 | 3486 | 2708 | 0.752 | -0.411 | 0.00041194 |
| AT5G61910.1 | BON1 | protein_coding | DCD (Development and C | 353 | 345 | 280 | 236 | 224 | 209 | 742 | 848 | 717 | 512 | 595 | 568 | 537.11 | 769 | 558 | 0.752 | -0.412 | 0.00070192 |
| AT5G11720.1 | | protein_coding | Glycosyl hydrolases famil | 826 | 826 | 699 | 458 | 408 | 414 | 1736 | 2029 | 1789 | 993 | 1083 | 1126 | 1392.37 | 1851 | 1067 | 0.752 | -0.412 | 0.00058586 |
| AT2G39930.1 | ISA1 | protein_coding | isoamylase 1 | 1532 | 1287 | 1278 | 983 | 833 | 731 | 3220 | 3162 | 3272 | 2132 | 2211 | 1988 | 2667.6 | 3218 | 2110 | 0.751 | -0.412 | 0.00029743 |
| AT4G11660.1 | HSFB2B | protein_coding | winged-helix DNA-bindin | 79 | 61 | 59 | 56 | 46 | 34 | 166 | 150 | 151 | 121 | 122 | 92 | 147.92 | 156 | 112 | 0.751 | -0.413 | 0.00921756 |
| AT1G15020.2 | QSOX1 | protein_coding | quiescin-sulfhydryl oxidas | 194 | 210 | 212 | 150 | 135 | 174 | 408 | 516 | 543 | 325 | 358 | 473 | 394.84 | 489 | 385 | 0.751 | -0.413 | 0.00363954 |
| AT4G32530.2 | VHA-c''1 | protein_coding | ATPase, F0/V0 complex, s | 625 | 594 | 572 | 671 | 490 | 484 | 1314 | 1459 | 1464 | 1455 | 1301 | 1316 | 1685.09 | 1412 | 1357 | 0.751 | -0.413 | 0.00031705 |
| AT5G09320.1 | VPS9B | protein_coding | Vacuolar sorting protein 9 | 60 | 55 | 58 | 52 | 61 | 55 | 126 | 135 | 148 | 113 | 162 | 150 | 155.94 | 136 | 142 | 0.751 | -0.414 | 0.00786991 |
| AT2G29050.1 | ATRBL1 | protein_coding | RHOMBOID-like 1 | 115 | 81 | 87 | 75 | 86 | 67 | 242 | 199 | 223 | 163 | 228 | 182 | 264.6 | 221 | 191 | 0.751 | -0.414 | 0.00645807 |
| AT3G52760.1 | | protein_coding | Integral membrane Yip1 f | 146 | 165 | 131 | 134 | 111 | 102 | 307 | 405 | 335 | 291 | 295 | 277 | 391.9 | 349 | 288 | 0.751 | -0.414 | 0.00063767 |
| AT4G39420.2 | | protein_coding | | 2633 | 2214 | 2079 | 2075 | 1790 | 1522 | 5535 | 5439 | 5322 | 4501 | 4752 | 4139 | 3251.87 | 5432 | 4464 | 0.75 | -0.414 | 0.00057149 |
| AT3G11397.1 | PRA1A3 | protein_coding | prenylated RAB acceptor | 268 | 221 | 211 | 176 | 156 | 172 | 563 | 543 | 540 | 382 | 414 | 468 | 530.47 | 549 | 421 | 0.751 | -0.414 | 0.0001566 |
| AT3G12570.1 | FYD | protein_coding | FYD | 450 | 458 | 481 | 430 | 318 | 418 | 946 | 1125 | 1231 | 933 | 844 | 1137 | 988.06 | 1101 | 971 | 0.75 | -0.415 | 0.00266818 |
| AT4G06676.1 | | protein_coding | | 88 | 83 | 78 | 53 | 47 | 44 | 185 | 204 | 200 | 115 | 125 | 120 | 158.87 | 196 | 120 | 0.75 | -0.415 | 0.00264843 |
| AT4G33180.1 | | protein_coding | alpha/beta-Hydrolases su | 135 | 159 | 151 | 143 | 99 | 121 | 284 | 391 | 387 | 310 | 263 | 329 | 362.06 | 354 | 301 | 0.75 | -0.415 | 0.00226727 |
| AT1G11200.1 | | protein_coding | Protein of unknown funct | 226 | 185 | 223 | 164 | 136 | 103 | 475 | 455 | 571 | 356 | 361 | 280 | 415.78 | 500 | 332 | 0.75 | -0.415 | 0.00074944 |
| AT2G15230.1 | LIP1 | protein_coding | lipase 1 | 162 | 136 | 172 | 146 | 115 | 143 | 341 | 334 | 440 | 317 | 305 | 389 | 318.34 | 372 | 337 | 0.75 | -0.416 | 0.00504615 |
| AT2G25280.1 | | protein_coding | | 403 | 320 | 412 | 339 | 257 | 289 | 847 | 786 | 1055 | 735 | 682 | 786 | 757.11 | 896 | 734 | 0.749 | -0.416 | 0.00083051 |
| AT2G47220.1 | DUF5 | protein_coding | DOMAIN OF UNKNOWN | 202 | 167 | 139 | 104 | 105 | 90 | 425 | 410 | 356 | 226 | 279 | 245 | 315.99 | 397 | 250 | 0.749 | -0.417 | 0.00287818 |
| AT5G41180.1 | | protein_coding | leucine-rich repeat transr | 155 | 149 | 131 | 141 | 116 | 93 | 326 | 366 | 335 | 306 | 308 | 253 | 266.43 | 342 | 289 | 0.748 | -0.418 | 0.00257208 |
| AT5G41100.1 | | protein_coding | | 203 | 191 | 167 | 149 | 123 | 131 | 427 | 469 | 428 | 323 | 327 | 356 | 357.73 | 441 | 335 | 0.748 | -0.418 | 0.00248528 |
| AT2G19600.1 | KEA4 | protein_coding | K+ efflux antiporter 4 | 303 | 279 | 243 | 230 | 203 | 209 | 637 | 685 | 622 | 499 | 539 | 568 | 603.94 | 648 | 535 | 0.749 | -0.418 | 0.00046058 |
| AT4G18050.1 | ABCB9 | protein_coding | P-glycoprotein 9 | 154 | 121 | 135 | 128 | 125 | 115 | 324 | 297 | 346 | 278 | 332 | 313 | 246.5 | 322 | 308 | 0.748 | -0.419 | 0.00206021 |
| AT1G09420.2 | G6PD4 | protein_coding | glucose-6-phosphate deh | 254 | 191 | 241 | 165 | 132 | 120 | 534 | 469 | 617 | 358 | 350 | 326 | 367.68 | 540 | 345 | 0.748 | -0.419 | 0.0012062 |
| AT2G20790.1 | AP5M | protein_coding | clathrin adaptor complex | 505 | 438 | 417 | 335 | 323 | 295 | 1062 | 1076 | 1068 | 727 | 857 | 802 | 898.93 | 1069 | 795 | 0.748 | -0.419 | 0.00022891 |
| AT2G46560.1 | | protein_coding | transducin family protein | 1961 | 1719 | 1557 | 1554 | 1239 | 1191 | 4122 | 4223 | 3986 | 3371 | 3289 | 3239 | 2706.7 | 4110 | 3300 | 0.748 | -0.419 | 0.00018338 |
| AT1G14990.2 | | protein_coding | | 131 | 86 | 94 | 127 | 81 | 80 | 275 | 211 | 241 | 275 | 215 | 218 | 281.76 | 242 | 236 | 0.747 | -0.42 | 0.00103019 |
| AT3G57030.1 | SSL10 | protein_coding | Calcium-dependent phos | 405 | 420 | 375 | 291 | 225 | 280 | 851 | 1032 | 960 | 631 | 597 | 761 | 712.66 | 948 | 663 | 0.748 | -0.42 | 0.00074907 |
| AT4G30920.1 | | protein_coding | Cytosol aminopeptidase f | 1641 | 1328 | 1289 | 1010 | 802 | 1008 | 3450 | 3263 | 3300 | 2191 | 2129 | 2741 | 2799.99 | 3338 | 2354 | 0.747 | -0.42 | 0.0006745 |
| AT2G05230.1 | | protein_coding | DNAJ heat shock N-termi | 342 | 269 | 309 | 275 | 224 | 213 | 719 | 661 | 791 | 597 | 595 | 579 | 541.46 | 724 | 590 | 0.747 | -0.42 | 0.00012507 |
| AT1G12090.1 | ELP | protein_coding | extensin-like protein | 5924 | 6095 | 4200 | 4953 | 3968 | 3778 | 12453 | 14974 | 10752 | 10744 | 10534 | 10274 | 31260.47 | 12726 | 10517 | 0.747 | -0.421 | 0.00659983 |
| AT1G76120.1 | | protein_coding | Pseudouridine synthase f | 155 | 121 | 114 | 105 | 93 | 62 | 326 | 297 | 292 | 228 | 247 | 169 | 251.4 | 305 | 215 | 0.747 | -0.421 | 0.00400597 |
| AT4G09560.1 | RMR4 | protein_coding | Protease-associated (PA) | 121 | 121 | 103 | 95 | 88 | 91 | 254 | 297 | 264 | 206 | 234 | 247 | 266.9 | 272 | 229 | 0.747 | -0.421 | 0.00205303 |
| AT3G23340.1 | ckl10 | protein_coding | casein kinase I-like 10 | 299 | 289 | 275 | 206 | 223 | 151 | 629 | 710 | 704 | 447 | 592 | 411 | 614.61 | 681 | 483 | 0.747 | -0.421 | 0.00126995 |
| AT3G45980.1 | H2B | protein_coding | Histone superfamily prot | 1613 | 1918 | 1652 | 1234 | 1185 | 1256 | 3391 | 4712 | 4229 | 2677 | 3146 | 3416 | 3699.32 | 4111 | 3080 | 0.747 | -0.422 | 0.00171403 |
| AT1G07950.1 | MED22B | protein_coding | Surfeit locus protein 5 sul | 127 | 94 | 100 | 104 | 95 | 77 | 267 | 231 | 256 | 226 | 252 | 209 | 257.78 | 251 | 229 | 0.747 | -0.422 | 0.00143329 |
| AT2G45950.2 | ASK20 | protein_coding | SKP1-like 20 | 407 | 277 | 265 | 238 | 180 | 169 | 856 | 681 | 678 | 516 | 478 | 460 | 566.84 | 738 | 485 | 0.746 | -0.422 | 0.00042248 |
| AT5G02410.1 | | protein_coding | DIE2/ALG10 family | 183 | 128 | 128 | 116 | 94 | 68 | 385 | 314 | 328 | 252 | 250 | 185 | 311.99 | 342 | 229 | 0.746 | -0.423 | 0.00158512 |
| AT5G10350.1 | PABN3 | protein_coding | RNA-binding (RRM/RBD/ | 371 | 308 | 259 | 267 | 179 | 235 | 780 | 757 | 663 | 579 | 475 | 639 | 891.3 | 733 | 564 | 0.746 | -0.423 | 0.00076953 |
| AT2G04410.1 | | protein_coding | RPM1-interacting protein | 614 | 523 | 518 | 481 | 405 | 403 | 1291 | 1285 | 1326 | 1043 | 1075 | 1096 | 1287.26 | 1301 | 1071 | 0.746 | -0.423 | 0.00021566 |
| AT4G19410.1 | PAE7 | protein_coding | Pectinacetylesterase fam | 2260 | 1832 | 1735 | 1896 | 969 | 1444 | 4751 | 4501 | 4442 | 4113 | 2572 | 3927 | 4991.47 | 4565 | 3537 | 0.746 | -0.424 | 0.00411178 |
| AT2G28890.1 | PLL4 | protein_coding | poltergeist like 4 | 417 | 314 | 292 | 327 | 184 | 204 | 877 | 771 | 748 | 709 | 488 | 555 | 576.46 | 799 | 584 | 0.745 | -0.424 | 0.00104038 |
| AT5G37480.1 | | protein_coding | | 157 | 125 | 127 | 95 | 100 | 71 | 330 | 307 | 325 | 206 | 265 | 193 | 296.12 | 321 | 221 | 0.745 | -0.424 | 0.00100851 |
| AT3G18500.3 | CCR4-3 | protein_coding | DNAse I-like superfamily | 415 | 489 | 420 | 332 | 289 | 253 | 872 | 1201 | 1075 | 720 | 767 | 688 | 1216.73 | 1049 | 725 | 0.745 | -0.424 | 0.00091171 |
| AT5G11640.1 | | protein_coding | Thioredoxin superfamily | 241 | 156 | 217 | 141 | 113 | 104 | 507 | 383 | 556 | 306 | 300 | 283 | 434.48 | 482 | 296 | 0.745 | -0.424 | 0.00074822 |
| AT3G15480.1 | | protein_coding | Protein of unknown funct | 680 | 614 | 625 | 611 | 496 | 543 | 1429 | 1508 | 1600 | 1325 | 1317 | 1477 | 1594.96 | 1512 | 1373 | 0.745 | -0.425 | 0.00081966 |
| AT4G27130.1 | | protein_coding | Translation initiation fact | 442 | 488 | 523 | 352 | 302 | 438 | 929 | 1199 | 1339 | 764 | 802 | 1191 | 1470.57 | 1156 | 919 | 0.744 | -0.426 | 0.00917421 |
| AT2G46890.1 | | protein_coding | Protein of unknown funct | 98 | 67 | 60 | 85 | 95 | 58 | 206 | 165 | 154 | 184 | 252 | 158 | 244.15 | 175 | 198 | 0.744 | -0.426 | 0.00789796 |
| AT5G66880.1 | SRK2I | protein_coding | sucrose nonfermenting 1 | 433 | 538 | 474 | 361 | 375 | 327 | 910 | 1322 | 1213 | 783 | 996 | 889 | 940.52 | 1148 | 889 | 0.744 | -0.426 | 0.00362018 |
| AT2G18850.1 | | protein_coding | SET domain-containing p | 168 | 159 | 170 | 101 | 106 | 98 | 353 | 391 | 435 | 219 | 281 | 267 | 299.49 | 393 | 256 | 0.744 | -0.427 | 0.00456568 |
| AT4G14040.1 | SBP2 | protein_coding | selenium-binding protein | 3755 | 2991 | 2358 | 6443 | 5077 | 4241 | 7893 | 7348 | 6037 | 13975 | 13478 | 11533 | 12215.55 | 7093 | 12995 | 0.744 | -0.427 | 0.00165545 |
| AT3G59500.1 | | protein_coding | Integral membrane HRF1 | 392 | 297 | 310 | 316 | 249 | 229 | 824 | 730 | 794 | 685 | 661 | 623 | 732.02 | 783 | 656 | 0.744 | -0.427 | 8.01E-05 |
| AT1G06830.1 | GRXS11 | protein_coding | Glutaredoxin family prot | 135 | 160 | 102 | 151 | 179 | 157 | 284 | 393 | 261 | 328 | 475 | 427 | 394.05 | 313 | 410 | 0.743 | -0.428 | 0.00700751 |
| AT1G29630.2 | EXO1 | protein_coding | 5'-3' exonuclease family | 201 | 148 | 155 | 152 | 117 | 127 | 423 | 364 | 397 | 330 | 311 | 345 | 339.11 | 395 | 329 | 0.743 | -0.428 | 0.00194318 |
| AT2G25730.2 | | protein_coding | | 1554 | 1606 | 1605 | 1082 | 1118 | 992 | 3267 | 3946 | 4109 | 2347 | 2968 | 2698 | 2519.18 | 3774 | 2671 | 0.743 | -0.428 | 0.00125052 |
| AT1G58350.1 | ZW18 | protein_coding | Putative serine esterase | 573 | 496 | 623 | 511 | 375 | 421 | 1204 | 1219 | 1595 | 1108 | 996 | 1145 | 1033.72 | 1339 | 1083 | 0.743 | -0.428 | 0.00103982 |
| AT2G45630.2 | | protein_coding | D-isomer specific 2-hydro | 79 | 77 | 70 | 55 | 50 | 56 | 166 | 189 | 179 | 119 | 133 | 152 | 201.35 | 178 | 135 | 0.743 | -0.429 | 0.00323158 |
| AT3G52110.1 | | protein_coding | | 201 | 203 | 215 | 157 | 139 | 110 | 423 | 499 | 550 | 341 | 369 | 299 | 389.15 | 491 | 336 | 0.742 | -0.43 | 0.00283052 |
| AT3G19490.1 | ATNHD1 | protein_coding | sodium:hydrogen antipor | 876 | 700 | 572 | 663 | 545 | 421 | 1841 | 1720 | 1464 | 1438 | 1447 | 1145 | 1823.29 | 1675 | 1343 | 0.742 | -0.43 | 0.00074024 |
| AT1G73470.3 | | protein_coding | | 439 | 448 | 444 | 324 | 336 | 303 | 923 | 1101 | 1137 | 703 | 892 | 824 | 1141.01 | 1054 | 806 | 0.742 | -0.43 | 0.000509 |
| AT5G40710.1 | | protein_coding | zinc finger (C2H2 type) fa | 140 | 107 | 142 | 105 | 88 | 83 | 294 | 263 | 364 | 228 | 234 | 226 | 253.23 | 307 | 229 | 0.742 | -0.431 | 0.00338906 |
| AT4G18060.1 | | protein_coding | SH3 domain-containing p | 561 | 436 | 502 | 455 | 342 | 424 | 1179 | 1071 | 1285 | 987 | 908 | 1153 | 1149.39 | 1178 | 1016 | 0.742 | -0.431 | 0.0005374 |
| AT1G78590.1 | NADK3 | protein_coding | NAD(H) kinase 3 | 195 | 192 | 189 | 128 | 136 | 132 | 410 | 472 | 484 | 278 | 361 | 359 | 401.94 | 455 | 333 | 0.742 | -0.431 | 0.00052819 |
| AT4G37670.2 | NAGS2 | protein_coding | N-acetyl-l-glutamate synt | 129 | 111 | 100 | 94 | 81 | 72 | 271 | 273 | 256 | 204 | 215 | 196 | 281.03 | 267 | 205 | 0.742 | -0.431 | 0.00032492 |
| AT3G11780.2 | | protein_coding | MD-2-related lipid recogr | 832 | 560 | 720 | 648 | 425 | 501 | 1749 | 1376 | 1843 | 1406 | 1128 | 1362 | 1720.59 | 1656 | 1299 | 0.741 | -0.432 | 0.00121126 |
| AT5G41340.1 | UBC4 | protein_coding | ubiquitin conjugating enz | 379 | 340 | 411 | 301 | 285 | 269 | 797 | 835 | 1052 | 653 | 757 | 732 | 821.47 | 895 | 714 | 0.741 | -0.432 | 0.00068266 |
| AT5G08000.1 | PDCB2 | protein_coding | glucan endo-1,3-beta-glu | 122 | 168 | 130 | 122 | 140 | 97 | 256 | 413 | 333 | 265 | 372 | 264 | 479.22 | 334 | 300 | 0.74 | -0.434 | 0.0081623 |
| AT5G04910.1 | | protein_coding | | 86 | 78 | 92 | 67 | 61 | 63 | 181 | 192 | 236 | 145 | 162 | 171 | 217.79 | 203 | 159 | 0.74 | -0.434 | 0.00154848 |
| AT2G19260.1 | | protein_coding | RING/FYVE/PHD zinc fing | 212 | 175 | 192 | 142 | 117 | 91 | 446 | 430 | 492 | 308 | 311 | 247 | 307.08 | 456 | 289 | 0.74 | -0.434 | 0.0013255 |
| AT3G09162.1 | | protein_coding | | 108 | 107 | 107 | 78 | 81 | 61 | 227 | 263 | 274 | 169 | 215 | 166 | 246.48 | 255 | 183 | 0.74 | -0.435 | 0.00788361 |
| AT5G58920.1 | | protein_coding | | 171 | 117 | 130 | 153 | 100 | 99 | 359 | 287 | 333 | 332 | 265 | 269 | 367.55 | 326 | 289 | 0.74 | -0.435 | 0.00112203 |
| AT2G22300.1 | CMTA3 | protein_coding | signal responsive 1 | 820 | 703 | 844 | 555 | 520 | 524 | 1724 | 1727 | 2161 | 1204 | 1380 | 1425 | 1289.38 | 1871 | 1336 | 0.74 | -0.435 | 0.00072152 |
| AT3G44820.1 | | protein_coding | Phototropic-responsive N | 260 | 238 | 242 | 215 | 157 | 156 | 547 | 585 | 620 | 466 | 417 | 424 | 424.97 | 584 | 436 | 0.74 | -0.435 | 0.00057028 |
| AT5G20140.2 | | protein_coding | SOUL heme-binding famil | 937 | 856 | 804 | 638 | 520 | 506 | 1970 | 2103 | 2058 | 1384 | 1380 | 1376 | 1953.35 | 2044 | 1380 | 0.74 | -0.435 | 8.28E-05 |
| AT4G21326.1 | ATSBT3.12 | protein_coding | subtilase 3.12 | 62 | 88 | 86 | 100 | 89 | 106 | 130 | 216 | 220 | 217 | 236 | 288 | 260.53 | 189 | 247 | 0.739 | -0.436 | 0.00808502 |
| AT4G00165.1 | | protein_coding | Bifunctional inhibitor/lipi | 462 | 414 | 262 | 473 | 315 | 273 | 971 | 1017 | 671 | 1026 | 836 | 742 | 1380.58 | 886 | 868 | 0.739 | -0.436 | 0.00508762 |
| AT3G52960.1 | PRXIIE | protein_coding | Thioredoxin superfamily | 2659 | 2721 | 1946 | 2116 | 1727 | 2087 | 5589 | 6685 | 4982 | 4590 | 4585 | 5675 | 7622 | 5752 | 4950 | 0.739 | -0.436 | 0.0019594 |
| AT2G32520.1 | | protein_coding | alpha/beta-Hydrolases su | 410 | 307 | 295 | 389 | 260 | 294 | 862 | 754 | 755 | 844 | 690 | 800 | 858 | 790 | 778 | 0.739 | -0.436 | 0.00016011 |
| AT4G35220.1 | | protein_coding | Cyclase family protein | 186 | 143 | 139 | 121 | 116 | 111 | 391 | 351 | 356 | 262 | 308 | 302 | 434.36 | 366 | 291 | 0.739 | -0.437 | 0.00047125 |
| AT3G58570.1 | RH52 | protein_coding | P-loop containing nucleo | 362 | 358 | 359 | 272 | 246 | 262 | 761 | 880 | 919 | 590 | 653 | 712 | 989.66 | 853 | 652 | 0.739 | -0.437 | 0.000245 |
| AT4G09310.1 | | protein_coding | SPla/RYanodine receptor | 197 | 145 | 160 | 143 | 127 | 100 | 414 | 356 | 410 | 310 | 337 | 272 | 376.15 | 393 | 306 | 0.738 | -0.438 | 0.00573457 |
| AT1G14290.1 | SBH2 | protein_coding | sphingoid base hydroxyla | 640 | 643 | 754 | 486 | 433 | 463 | 1345 | 1580 | 1930 | 1054 | 1150 | 1259 | 1375.28 | 1618 | 1154 | 0.738 | -0.438 | 0.00193904 |
| AT1G27070.1 | | protein_coding | 5'-AMP-activated protein | 219 | 202 | 193 | 207 | 179 | 134 | 460 | 496 | 494 | 449 | 475 | 364 | 398.47 | 483 | 429 | 0.738 | -0.438 | 0.00133332 |
| AT4G38320.1 | HHP5 | protein_coding | heptahelical protein 5 | 210 | 180 | 163 | 204 | 130 | 115 | 441 | 442 | 417 | 442 | 345 | 313 | 425.48 | 433 | 367 | 0.738 | -0.438 | 0.00056149 |
| AT5G56250.2 | HAP8 | protein_coding | hapless 8 | 541 | 391 | 415 | 408 | 265 | 291 | 1137 | 961 | 1062 | 885 | 704 | 791 | 744.14 | 1053 | 793 | 0.738 | -0.438 | 0.0003372 |
| AT3G51610.1 | | protein_coding | | 232 | 222 | 209 | 207 | 152 | 170 | 488 | 545 | 535 | 449 | 404 | 462 | 541.61 | 523 | 438 | 0.738 | -0.438 | 7.45E-05 |
| AT4G17680.1 | | protein_coding | SBP (S-ribonuclease bindi | 99 | 84 | 70 | 65 | 47 | 54 | 208 | 206 | 179 | 141 | 125 | 147 | 252.96 | 198 | 138 | 0.738 | -0.439 | 0.00105254 |
| AT1G77000.1 | SKP2B | protein_coding | RNI-like superfamily prot | 89 | 74 | 65 | 68 | 75 | 52 | 187 | 182 | 166 | 147 | 199 | 141 | 179.56 | 178 | 162 | 0.737 | -0.44 | 0.00690677 |
| AT5G49555.1 | | protein_coding | FAD/NAD(P)-binding oxid | 430 | 384 | 366 | 351 | 299 | 262 | 904 | 943 | 937 | 761 | 794 | 712 | 927.44 | 928 | 756 | 0.737 | -0.44 | 3.80E-05 |
| AT4G16110.1 | ARR2 | protein_coding | response regulator 2 | 150 | 160 | 162 | 134 | 125 | 160 | 315 | 393 | 415 | 291 | 332 | 435 | 307.94 | 374 | 353 | 0.737 | -0.441 | 0.00556129 |
| AT1G34300.1 | | protein_coding | lectin protein kinase fami | 182 | 196 | 210 | 168 | 140 | 180 | 383 | 482 | 538 | 364 | 372 | 489 | 398.83 | 468 | 408 | 0.736 | -0.441 | 0.00228631 |
| AT5G18525.1 | | protein_coding | protein serine/threonine | 545 | 549 | 574 | 369 | 407 | 425 | 1146 | 1349 | 1469 | 800 | 1080 | 1156 | 957.01 | 1321 | 1012 | 0.736 | -0.441 | 0.00177562 |
| AT1G64440.1 | UGE4 | protein_coding | NAD(P)-binding Rossman | 221 | 133 | 136 | 188 | 131 | 150 | 465 | 327 | 348 | 408 | 348 | 408 | 450.13 | 380 | 388 | 0.737 | -0.441 | 0.00051182 |
| AT3G02540.1 | RAD23C | protein_coding | Rad23 UV excision repair | 869 | 1054 | 1072 | 853 | 848 | 1055 | 1827 | 2589 | 2744 | 1850 | 2251 | 2869 | 2684.08 | 2387 | 2323 | 0.736 | -0.442 | 0.00744969 |
| AT1G34570.1 | | protein_coding | Essential protein Yae1, N | 282 | 191 | 139 | 203 | 157 | 110 | 593 | 469 | 356 | 440 | 417 | 299 | 498.03 | 473 | 385 | 0.736 | -0.442 | 0.00463845 |
| AT4G01550.1 | NAC69 | protein_coding | NAC domain containing p | 230 | 209 | 216 | 157 | 135 | 122 | 483 | 513 | 553 | 341 | 358 | 332 | 351.46 | 516 | 344 | 0.736 | -0.442 | 0.00151368 |
| AT2G26810.1 | | protein_coding | Putative methyltransfera | 133 | 123 | 116 | 100 | 77 | 81 | 280 | 302 | 297 | 217 | 204 | 220 | 257.06 | 293 | 214 | 0.736 | -0.442 | 0.00058221 |
| AT1G12620.1 | | protein_coding | Pentatricopeptide repeat | 175 | 143 | 142 | 124 | 117 | 96 | 368 | 351 | 364 | 269 | 311 | 261 | 275.58 | 361 | 280 | 0.736 | -0.442 | 0.00035853 |
| AT1G77580.2 | FPP1 | protein_coding | Plant protein of unknowr | 502 | 450 | 412 | 317 | 342 | 262 | 1055 | 1106 | 1055 | 688 | 908 | 712 | 738.58 | 1072 | 769 | 0.736 | -0.442 | 0.00033013 |
| AT1G17510.1 | | protein_coding | | 65 | 66 | 54 | 66 | 50 | 63 | 137 | 162 | 138 | 143 | 133 | 171 | 186.13 | 146 | 149 | 0.735 | -0.443 | 0.00139876 |
| AT1G51550.1 | | protein_coding | Kelch repeat-containing F | 147 | 137 | 139 | 87 | 99 | 100 | 309 | 337 | 356 | 189 | 263 | 272 | 249.19 | 334 | 241 | 0.735 | -0.444 | 0.00269609 |
| AT4G33950.1 | SRK2E | protein_coding | Protein kinase superfamil | 252 | 230 | 245 | 229 | 162 | 145 | 530 | 565 | 627 | 497 | 430 | 394 | 525.67 | 574 | 440 | 0.735 | -0.444 | 0.00012866 |
| AT5G22580.1 | | protein_coding | Stress responsive A/B Bar | 216 | 250 | 237 | 140 | 154 | 168 | 454 | 614 | 607 | 304 | 409 | 457 | 668.24 | 558 | 390 | 0.734 | -0.445 | 0.00279708 |
| AT1G13390.1 | | protein_coding | | 83 | 94 | 68 | 59 | 44 | 51 | 174 | 231 | 174 | 128 | 117 | 139 | 236.4 | 193 | 128 | 0.735 | -0.445 | 0.00078503 |
| AT2G20130.1 | LCV1 | protein_coding | like COV 1 | 281 | 222 | 256 | 202 | 153 | 197 | 591 | 545 | 655 | 438 | 406 | 536 | 474.66 | 597 | 460 | 0.735 | -0.445 | 0.0006885 |
| AT4G36860.1 | DAR1 | protein_coding | LIM domain-containing p | 500 | 389 | 413 | 359 | 254 | 265 | 1051 | 956 | 1057 | 779 | 674 | 721 | 750.03 | 1021 | 725 | 0.735 | -0.445 | 4.63E-05 |
| AT1G74600.1 | PCMP-E69 | protein_coding | pentatricopeptide (PPR) r | 132 | 90 | 94 | 91 | 81 | 75 | 277 | 221 | 241 | 197 | 215 | 204 | 192.96 | 246 | 205 | 0.734 | -0.446 | 0.00686701 |
| AT3G01330.1 | E2FF | protein_coding | DP-E2F-like protein 3 | 95 | 70 | 75 | 80 | 82 | 51 | 200 | 172 | 192 | 174 | 218 | 139 | 174.97 | 188 | 177 | 0.734 | -0.446 | 0.00548933 |
| AT5G19070.1 | | protein_coding | SNARE associated Golgi p | 159 | 134 | 121 | 108 | 74 | 123 | 334 | 329 | 310 | 234 | 196 | 334 | 306.88 | 324 | 255 | 0.734 | -0.446 | 0.0022598 |
| AT2G27340.2 | | protein_coding | N-acetylglucosaminylpho | 112 | 106 | 104 | 108 | 70 | 72 | 235 | 260 | 266 | 234 | 186 | 196 | 256.97 | 254 | 205 | 0.734 | -0.446 | 0.00212231 |
| AT4G19985.1 | | protein_coding | Acyl-CoA N-acyltransfera | 455 | 329 | 384 | 351 | 311 | 200 | 956 | 808 | 983 | 761 | 826 | 544 | 1100.24 | 916 | 710 | 0.734 | -0.446 | 0.00123925 |
| AT5G61580.1 | PFK4 | protein_coding | phosphofructokinase 4 | 468 | 336 | 389 | 291 | 191 | 217 | 984 | 825 | 996 | 631 | 507 | 590 | 799.97 | 935 | 576 | 0.734 | -0.446 | 0.00021489 |
| AT5G05830.1 | | protein_coding | RING/FYVE/PHD zinc fing | 171 | 149 | 179 | 145 | 130 | 125 | 359 | 366 | 458 | 315 | 345 | 340 | 406.36 | 394 | 333 | 0.734 | -0.446 | 8.93E-05 |
| AT3G46000.1 | ADF2 | protein_coding | actin depolymerizing fact | 986 | 976 | 894 | 846 | 754 | 730 | 2073 | 2398 | 2289 | 1835 | 2002 | 1985 | 2249.55 | 2253 | 1941 | 0.734 | -0.447 | 0.00011473 |
| AT3G26380.1 | | protein_coding | Melibiase family protein | 363 | 275 | 292 | 245 | 199 | 213 | 763 | 676 | 748 | 531 | 528 | 579 | 577.43 | 729 | 546 | 0.733 | -0.447 | 2.36E-05 |
| AT5G10745.1 | | protein_coding | | 172 | 157 | 185 | 142 | 134 | 139 | 362 | 386 | 474 | 308 | 356 | 378 | 380.44 | 407 | 347 | 0.733 | -0.448 | 0.0010293 |
| AT1G51940.1 | LYK3 | protein_coding | protein kinase family pro | 1410 | 1270 | 1511 | 951 | 799 | 841 | 2964 | 3120 | 3868 | 2063 | 2121 | 2287 | 2756.82 | 3317 | 2157 | 0.733 | -0.448 | 0.00072783 |
| AT5G55810.2 | NMNAT | protein_coding | nicotinate/nicotinamide | 67 | 60 | 74 | 54 | 40 | 43 | 141 | 147 | 189 | 117 | 106 | 117 | 128.82 | 159 | 113 | 0.732 | -0.449 | 0.00640787 |
| AT4G28590.1 | | protein_coding | | 155 | 110 | 90 | 129 | 110 | 109 | 326 | 270 | 230 | 280 | 292 | 296 | 333.82 | 275 | 289 | 0.732 | -0.449 | 0.00127254 |
| AT1G74940.1 | | protein_coding | Protein of unknown funct | 226 | 326 | 303 | 227 | 266 | 206 | 475 | 801 | 776 | 492 | 706 | 560 | 622.34 | 684 | 586 | 0.732 | -0.45 | 0.00977817 |
| AT5G59770.1 | | protein_coding | Protein-tyrosine phospha | 320 | 287 | 250 | 268 | 224 | 155 | 673 | 705 | 640 | 581 | 595 | 422 | 694.08 | 673 | 533 | 0.732 | -0.45 | 0.0005635 |
| AT4G23400.1 | PIP1-5 | protein_coding | plasma membrane intrins | 1950 | 2441 | 1958 | 1544 | 1259 | 1494 | 4099 | 5997 | 5013 | 3349 | 3342 | 4063 | 5472.52 | 5036 | 3585 | 0.732 | -0.451 | 0.00157412 |
| AT3G55760.1 | | protein_coding | | 144 | 181 | 157 | 80 | 85 | 86 | 303 | 445 | 402 | 174 | 226 | 234 | 632.62 | 383 | 211 | 0.732 | -0.451 | 0.00104636 |
| AT1G06240.1 | | protein_coding | Protein of unknown funct | 129 | 117 | 91 | 112 | 80 | 83 | 271 | 287 | 233 | 243 | 212 | 226 | 363.31 | 264 | 227 | 0.732 | -0.451 | 0.00073299 |
| AT4G33470.1 | HDA14 | protein_coding | histone deacetylase 14 | 592 | 606 | 552 | 337 | 356 | 340 | 1244 | 1489 | 1413 | 731 | 945 | 925 | 1310.37 | 1382 | 867 | 0.732 | -0.451 | 0.0003625 |
| AT3G24515.1 | UBC37 | protein_coding | ubiquitin-conjugating enz | 139 | 91 | 136 | 74 | 63 | 66 | 292 | 224 | 348 | 161 | 167 | 179 | 235.59 | 288 | 169 | 0.731 | -0.452 | 0.00623052 |
| AT3G17350.1 | | protein_coding | | 98 | 118 | 80 | 54 | 66 | 70 | 206 | 290 | 205 | 117 | 175 | 190 | 259.89 | 234 | 161 | 0.731 | -0.452 | 0.00403786 |
| AT5G16450.1 | | protein_coding | Ribonuclease E inhibitor | 88 | 79 | 78 | 75 | 51 | 59 | 185 | 194 | 200 | 163 | 135 | 160 | 194.21 | 193 | 153 | 0.731 | -0.452 | 0.00168737 |
| AT1G43130.1 | LCV2 | protein_coding | like COV 2 | 289 | 244 | 282 | 215 | 183 | 209 | 607 | 599 | 722 | 466 | 486 | 568 | 577.48 | 643 | 507 | 0.731 | -0.452 | 0.00040668 |
| AT2G26200.1 | | protein_coding | S-adenosyl-L-methionine- | 460 | 400 | 465 | 316 | 229 | 225 | 967 | 983 | 1190 | 685 | 608 | 612 | 843 | 1047 | 635 | 0.731 | -0.452 | 0.00024219 |
| AT4G03150.1 | | protein_coding | | 410 | 363 | 263 | 328 | 264 | 190 | 862 | 892 | 673 | 711 | 701 | 517 | 741.39 | 809 | 643 | 0.731 | -0.453 | 0.0006154 |
| AT5G42240.1 | SCPL42 | protein_coding | serine carboxypeptidase- | 742 | 717 | 544 | 449 | 408 | 394 | 1560 | 1762 | 1393 | 974 | 1083 | 1071 | 1400.94 | 1572 | 1043 | 0.731 | -0.453 | 0.00019126 |

| | | | | | | | | | | | | | | | | | | | | | |
|---|---|---|---|---|---|---|---|---|---|---|---|---|---|---|---|---|---|---|---|---|---|
| AT5G01015.1 | | protein_coding | | 57 | 75 | 47 | 57 | 49 | 48 | 120 | 184 | 120 | 124 | 130 | 131 | 397.11 | 141 | 128 | 0.73 | -0.454 | 0.0041686 |
| AT3G45630.1 | | protein_coding | RNA binding (RRM/RBD/I | 1042 | 822 | 782 | 738 | 593 | 508 | 2190 | 2020 | 2002 | 1601 | 1574 | 1381 | 1530.35 | 2071 | 1519 | 0.73 | -0.454 | 0.00018502 |
| AT1G20260.1 | VHA-B3 | protein_coding | ATPase, V1 complex, subu | 4394 | 3588 | 3546 | 3197 | 2495 | 2304 | 9237 | 8815 | 9078 | 6935 | 6624 | 6266 | 7947.74 | 9043 | 6608 | 0.73 | -0.454 | 6.00E-05 |
| AT5G24060.2 | | protein_coding | Pentatricopeptide repeat | 533 | 433 | 392 | 275 | 265 | 247 | 1120 | 1064 | 1004 | 597 | 704 | 672 | 1106.16 | 1063 | 658 | 0.73 | -0.454 | 4.75E-05 |
| AT1G71710.1 | | protein_coding | DNAse I-like superfamily | 705 | 687 | 599 | 555 | 407 | 380 | 1482 | 1688 | 1533 | 1204 | 1080 | 1033 | 1290.44 | 1568 | 1106 | 0.73 | -0.455 | 0.00031201 |
| AT3G05520.2 | | protein_coding | Subunits of heterodimeri | 378 | 329 | 373 | 277 | 265 | 252 | 795 | 808 | 955 | 601 | 704 | 685 | 817.86 | 853 | 663 | 0.73 | -0.455 | 0.00016712 |
| AT3G56580.2 | | protein_coding | RING/U-box superfamily | 240 | 238 | 226 | 192 | 182 | 201 | 504 | 585 | 579 | 416 | 483 | 547 | 602.34 | 556 | 482 | 0.73 | -0.455 | 9.17E-05 |
| AT1G49180.1 | | protein_coding | protein kinase family pro | 138 | 149 | 187 | 97 | 109 | 98 | 290 | 366 | 479 | 210 | 289 | 267 | 304.77 | 378 | 255 | 0.729 | -0.456 | 0.00917067 |
| AT4G39100.1 | SHL1 | protein_coding | PHD finger family protein | 399 | 375 | 398 | 334 | 292 | 329 | 839 | 921 | 1019 | 724 | 775 | 895 | 876 | 926 | 798 | 0.729 | -0.456 | 0.00025636 |
| AT1G29750.2 | RKF1 | protein_coding | receptor-like kinase in flo | 484 | 413 | 401 | 316 | 283 | 259 | 1017 | 1015 | 1027 | 685 | 751 | 704 | 700.64 | 1020 | 713 | 0.729 | -0.456 | 8.79E-06 |
| AT4G28010.1 | | protein_coding | Tetratricopeptide repeat | 112 | 81 | 97 | 74 | 62 | 55 | 235 | 199 | 248 | 161 | 165 | 150 | 164.88 | 227 | 159 | 0.728 | -0.457 | 0.00412192 |
| AT1G49300.1 | RABG3E | protein_coding | RAB GTPase homolog G3 | 581 | 520 | 605 | 385 | 321 | 419 | 1221 | 1278 | 1549 | 835 | 852 | 1139 | 1423.43 | 1349 | 942 | 0.728 | -0.457 | 0.0008584 |
| AT3G27990.1 | | other_rna | other RNA | 82 | 79 | 68 | 55 | 41 | 49 | 172 | 194 | 174 | 119 | 109 | 133 | 244.1 | 180 | 120 | 0.729 | -0.457 | 0.00054811 |
| AT3G58160.1 | XI-J | protein_coding | P-loop containing nucleo | 210 | 157 | 164 | 132 | 113 | 90 | 441 | 386 | 420 | 286 | 300 | 245 | 323.05 | 416 | 277 | 0.728 | -0.457 | 0.00042141 |
| AT2G29170.1 | | protein_coding | NAD(P)-binding Rossman | 93 | 100 | 67 | 26 | 25 | 27 | 195 | 246 | 172 | 56 | 66 | 73 | 251.95 | 204 | 65 | 0.728 | -0.458 | 0.00574137 |
| AT3G05800.1 | BHLH150 | protein_coding | AtBS1(activation-tagged | 134 | 223 | 162 | 158 | 115 | 150 | 282 | 548 | 415 | 343 | 305 | 408 | 745.41 | 415 | 352 | 0.728 | -0.458 | 0.00517932 |
| AT1G50500.2 | VPS53 | protein_coding | Membrane trafficking VP | 697 | 656 | 682 | 522 | 416 | 494 | 1465 | 1612 | 1746 | 1132 | 1104 | 1343 | 1312.54 | 1608 | 1193 | 0.728 | -0.458 | 0.0004358 |
| AT2G42700.2 | MIP3 | protein_coding | | 353 | 270 | 280 | 247 | 212 | 202 | 742 | 663 | 717 | 536 | 563 | 549 | 550.9 | 707 | 549 | 0.728 | -0.458 | 0.00010101 |
| AT1G06645.1 | | protein_coding | 2-oxoglutarate (2OG) and | 183 | 198 | 161 | 146 | 111 | 120 | 385 | 486 | 412 | 317 | 295 | 326 | 413.22 | 428 | 313 | 0.727 | -0.459 | 0.00050252 |
| AT5G03220.1 | MED7A | protein_coding | Mediator complex, subur | 190 | 150 | 169 | 137 | 85 | 96 | 399 | 369 | 433 | 297 | 226 | 261 | 281.29 | 400 | 261 | 0.727 | -0.46 | 0.002341 |
| AT5G63140.1 | PAP29 | protein_coding | purple acid phosphatase | 297 | 215 | 161 | 151 | 100 | 128 | 624 | 528 | 412 | 328 | 265 | 348 | 431.03 | 521 | 314 | 0.727 | -0.46 | 0.00148949 |
| AT1G70900.1 | | protein_coding | | 291 | 224 | 184 | 278 | 185 | 158 | 612 | 550 | 471 | 603 | 491 | 430 | 544.53 | 544 | 508 | 0.727 | -0.46 | 0.00090541 |
| AT2G22080.1 | | protein_coding | | 150 | 159 | 188 | 102 | 92 | 109 | 315 | 391 | 481 | 221 | 244 | 296 | 689.36 | 396 | 254 | 0.727 | -0.46 | 0.00082182 |
| AT5G09770.1 | | protein_coding | Ribosomal protein L17 fa | 250 | 167 | 158 | 178 | 142 | 160 | 526 | 410 | 404 | 386 | 377 | 435 | 519.5 | 447 | 399 | 0.726 | -0.461 | 0.00026265 |
| AT2G47830.1 | MTPC1 | protein_coding | Cation efflux family prote | 206 | 166 | 186 | 169 | 142 | 129 | 433 | 408 | 476 | 367 | 377 | 351 | 395.01 | 439 | 365 | 0.726 | -0.462 | 0.00043002 |
| AT5G44520.1 | RPI4 | protein_coding | NagB/RpiA/CoA transfera | 852 | 701 | 573 | 562 | 499 | 371 | 1791 | 1722 | 1467 | 1219 | 1325 | 1009 | 1417.92 | 1660 | 1184 | 0.726 | -0.462 | 0.00034042 |
| AT1G11670.1 | | protein_coding | MATE efflux family prote | 297 | 276 | 207 | 245 | 180 | 145 | 624 | 678 | 530 | 531 | 478 | 394 | 625.8 | 611 | 468 | 0.726 | -0.462 | 0.00022114 |
| AT5G65760.1 | | protein_coding | Serine carboxypeptidase | 477 | 381 | 405 | 358 | 246 | 308 | 1003 | 936 | 1037 | 777 | 653 | 838 | 815.41 | 992 | 756 | 0.726 | -0.462 | 8.45E-05 |
| AT3G26090.1 | RGS1 | protein_coding | G-protein coupled recept | 185 | 159 | 156 | 164 | 100 | 125 | 389 | 391 | 399 | 356 | 265 | 340 | 321.73 | 393 | 320 | 0.725 | -0.463 | 0.0031397 |
| AT5G47820.1 | FRA1 | protein_coding | P-loop containing nucleo | 1416 | 1298 | 1286 | 1041 | 947 | 937 | 2977 | 3189 | 3292 | 2258 | 2514 | 2548 | 2206.24 | 3153 | 2440 | 0.725 | -0.463 | 3.89E-05 |
| AT1G08210.1 | | protein_coding | Eukaryotic aspartyl prote | 185 | 183 | 199 | 128 | 137 | 179 | 389 | 450 | 509 | 278 | 364 | 487 | 412.95 | 449 | 376 | 0.725 | -0.464 | 0.00279668 |
| AT3G30300.1 | | protein_coding | O-fucosyltransferase fam | 747 | 728 | 711 | 606 | 582 | 439 | 1570 | 1789 | 1820 | 1314 | 1545 | 1194 | 1368.54 | 1726 | 1351 | 0.725 | -0.464 | 0.00033426 |
| AT1G60860.1 | AGD2 | protein_coding | ARF-GAP domain 2 | 269 | 287 | 302 | 207 | 197 | 156 | 565 | 705 | 773 | 449 | 523 | 424 | 517.24 | 681 | 465 | 0.725 | -0.464 | 0.00030803 |
| AT5G42420.1 | | protein_coding | Nucleotide-sugar transpo | 521 | 484 | 494 | 340 | 312 | 295 | 1095 | 1189 | 1265 | 737 | 828 | 802 | 1103.18 | 1183 | 789 | 0.725 | -0.464 | 0.00012025 |
| AT3G03140.1 | | protein_coding | Tudor/PWWP/MBT super | 245 | 233 | 233 | 174 | 117 | 144 | 515 | 572 | 596 | 377 | 311 | 392 | 361.5 | 561 | 360 | 0.725 | -0.465 | 0.00044112 |
| AT5G13460.1 | IQD11 | protein_coding | IQ-domain 11 | 348 | 322 | 388 | 292 | 258 | 300 | 732 | 791 | 993 | 633 | 685 | 816 | 627.7 | 839 | 711 | 0.724 | -0.465 | 0.00037113 |
| AT1G78010.1 | | protein_coding | tRNA modification GTPas | 345 | 259 | 210 | 212 | 180 | 140 | 725 | 636 | 538 | 460 | 478 | 381 | 622.86 | 633 | 440 | 0.725 | -0.465 | 0.00028614 |
| AT2G21120.1 | | protein_coding | Protein of unknown funct | 518 | 393 | 448 | 393 | 279 | 304 | 1089 | 966 | 1147 | 852 | 741 | 827 | 892.48 | 1067 | 807 | 0.724 | -0.465 | 4.54E-05 |
| AT1G54570.1 | | protein_coding | Esterase/lipase/thioester | 367 | 294 | 282 | 220 | 193 | 167 | 771 | 722 | 722 | 477 | 512 | 454 | 565.63 | 738 | 481 | 0.724 | -0.465 | 9.36E-06 |
| AT2G46980.2 | | protein_coding | | 275 | 217 | 187 | 188 | 159 | 140 | 578 | 533 | 479 | 408 | 422 | 381 | 366.8 | 530 | 404 | 0.724 | -0.466 | 0.00020903 |
| AT1G15830.1 | | protein_coding | | 1 | 5 | 2 | 1 | 0 | 0 | 2 | 12 | 5 | 2 | 0 | 0 | 174.24 | 6 | 1 | 0.724 | -0.467 | 0.00506311 |
| AT1G14130.1 | | protein_coding | 2-oxoglutarate (2OG) and | 97 | 72 | 56 | 80 | 57 | 54 | 204 | 177 | 143 | 174 | 151 | 147 | 151.02 | 175 | 157 | 0.723 | -0.467 | 0.00473661 |
| AT4G14430.1 | ECI2 | protein_coding | indole-3-butyric acid resp | 268 | 235 | 168 | 319 | 214 | 314 | 563 | 577 | 430 | 692 | 568 | 854 | 567.1 | 523 | 705 | 0.723 | -0.467 | 0.00360064 |
| AT1G63020.1 | NRPD1 | protein_coding | nuclear RNA polymerase | 252 | 190 | 163 | 217 | 182 | 125 | 530 | 467 | 417 | 471 | 483 | 340 | 321.03 | 471 | 431 | 0.724 | -0.467 | 0.00336124 |
| AT4G02410.1 | LECRK43 | protein_coding | Concanavalin A-like lectir | 165 | 191 | 162 | 145 | 97 | 112 | 347 | 469 | 415 | 315 | 258 | 305 | 286.57 | 410 | 293 | 0.723 | -0.468 | 0.00391054 |
| AT4G39140.1 | | protein_coding | RING/U-box superfamily | 157 | 172 | 151 | 149 | 120 | 149 | 330 | 423 | 387 | 323 | 319 | 405 | 356.66 | 380 | 349 | 0.723 | -0.468 | 0.00096868 |
| AT2G43850.1 | | protein_coding | Integrin-linked protein ki | 169 | 151 | 146 | 95 | 88 | 90 | 355 | 371 | 374 | 206 | 234 | 245 | 278.98 | 367 | 228 | 0.722 | -0.47 | 0.00160381 |
| AT1G77490.1 | APXT | protein_coding | thylakoidal ascorbate per | 2239 | 2535 | 2291 | 1482 | 1488 | 1480 | 4707 | 6228 | 5865 | 3215 | 3950 | 4025 | 6160.78 | 5600 | 3730 | 0.722 | -0.47 | 0.00033298 |
| AT2G38000.1 | | protein_coding | chaperone protein dnaJ-r | 219 | 208 | 209 | 140 | 139 | 133 | 460 | 511 | 535 | 304 | 369 | 362 | 395.31 | 502 | 345 | 0.722 | -0.47 | 0.00012189 |
| AT3G50960.1 | PLP3A | protein_coding | phosducin-like protein 3 | 98 | 63 | 73 | 38 | 44 | 35 | 206 | 155 | 187 | 82 | 117 | 95 | 145.78 | 183 | 98 | 0.722 | -0.471 | 0.0094195 |
| AT3G16320.1 | CDC27A | protein_coding | Tetratricopeptide repeat | 85 | 59 | 50 | 56 | 39 | 41 | 179 | 145 | 128 | 121 | 104 | 111 | 119.51 | 151 | 112 | 0.721 | -0.472 | 0.00595313 |
| AT1G62430.1 | CDS1 | protein_coding | CDP-diacylglycerol syntha | 891 | 961 | 763 | 622 | 639 | 422 | 1873 | 2361 | 1953 | 1349 | 1696 | 1148 | 1724.58 | 2062 | 1398 | 0.721 | -0.472 | 0.00123395 |
| AT3G14560.1 | | protein_coding | | 114 | 102 | 116 | 123 | 102 | 100 | 240 | 251 | 297 | 267 | 271 | 272 | 201.4 | 263 | 270 | 0.721 | -0.473 | 0.0099961 |
| AT4G10330.1 | | protein_coding | glycine-rich protein | 73 | 69 | 56 | 63 | 37 | 48 | 153 | 170 | 143 | 137 | 98 | 131 | 169.26 | 155 | 122 | 0.72 | -0.473 | 0.00098302 |
| AT4G25680.1 | | protein_coding | PPPDE putative thiol pep | 348 | 268 | 267 | 259 | 206 | 250 | 732 | 658 | 684 | 562 | 547 | 680 | 680.7 | 691 | 596 | 0.72 | -0.473 | 1.56E-05 |
| AT2G27200.1 | LSG1-1 | protein_coding | P-loop containing nucleo | 188 | 191 | 196 | 108 | 93 | 145 | 395 | 469 | 502 | 234 | 247 | 394 | 293.57 | 455 | 292 | 0.72 | -0.474 | 0.00793139 |
| AT1G50640.1 | ERF3 | protein_coding | ethylene responsive elem | 261 | 359 | 287 | 240 | 169 | 266 | 549 | 882 | 735 | 521 | 449 | 723 | 681.78 | 722 | 564 | 0.72 | -0.474 | 0.00501314 |
| AT2G15695.1 | | protein_coding | Protein of unknown funct | 350 | 310 | 349 | 237 | 204 | 311 | 736 | 762 | 893 | 514 | 542 | 846 | 664.59 | 797 | 634 | 0.72 | -0.474 | 0.00282979 |
| AT1G63480.1 | AHL12 | protein_coding | AT hook motif DNA-bindi | 127 | 127 | 97 | 99 | 91 | 80 | 267 | 312 | 248 | 215 | 242 | 218 | 340.76 | 276 | 225 | 0.72 | -0.474 | 4.81E-05 |
| AT4G37660.1 | | protein_coding | Ribosomal protein L12/ A | 148 | 131 | 83 | 98 | 62 | 95 | 311 | 322 | 212 | 213 | 165 | 258 | 308.43 | 282 | 212 | 0.719 | -0.475 | 0.00708595 |
| AT2G30330.1 | BLOS1 | protein_coding | GCN5L1 family protein | 70 | 80 | 47 | 75 | 47 | 54 | 147 | 197 | 120 | 163 | 125 | 147 | 177.94 | 155 | 145 | 0.719 | -0.475 | 0.00578594 |
| AT5G63850.1 | AAP4 | protein_coding | amino acid permease 4 | 419 | 343 | 354 | 302 | 349 | 407 | 881 | 843 | 906 | 655 | 927 | 1107 | 887.36 | 877 | 896 | 0.719 | -0.475 | 0.00503558 |
| AT3G05160.1 | | protein_coding | Major facilitator superfar | 358 | 390 | 373 | 255 | 251 | 246 | 753 | 958 | 955 | 553 | 666 | 669 | 1118.38 | 889 | 629 | 0.719 | -0.475 | 0.00013961 |
| AT1G65260.1 | VIPP1 | protein_coding | plastid transcriptionally a | 2749 | 2505 | 2672 | 2120 | 1859 | 1812 | 5779 | 6154 | 6840 | 4598 | 4935 | 4928 | 7062.38 | 6258 | 4820 | 0.719 | -0.475 | 5.43E-05 |
| AT1G13910.1 | | protein_coding | Leucine-rich repeat (LRR) | 90 | 86 | 85 | 49 | 62 | 40 | 189 | 211 | 218 | 106 | 165 | 109 | 164.43 | 206 | 127 | 0.719 | -0.476 | 0.00556415 |
| AT2G01170.1 | BAT1 | protein_coding | bidirectional amino acid t | 258 | 356 | 245 | 144 | 178 | 182 | 542 | 875 | 627 | 312 | 473 | 495 | 825.45 | 681 | 427 | 0.719 | -0.476 | 0.00488261 |
| AT5G52120.1 | PP2A14 | protein_coding | phloem protein 2-A14 | 76 | 84 | 63 | 41 | 36 | 35 | 160 | 206 | 161 | 89 | 96 | 95 | 141.32 | 176 | 93 | 0.719 | -0.476 | 0.00283864 |
| AT1G21730.1 | | protein_coding | P-loop containing nucleo | 569 | 544 | 609 | 346 | 420 | 360 | 1196 | 1337 | 1559 | 751 | 1115 | 979 | 987.99 | 1364 | 948 | 0.719 | -0.476 | 0.00117379 |
| AT5G13720.1 | | protein_coding | Uncharacterised protein f | 350 | 343 | 292 | 224 | 218 | 283 | 736 | 843 | 748 | 486 | 579 | 770 | 1053.26 | 776 | 612 | 0.719 | -0.476 | 0.00076881 |
| AT5G20180.1 | | protein_coding | Ribosomal protein L36 | 377 | 234 | 224 | 233 | 208 | 196 | 792 | 575 | 573 | 505 | 552 | 533 | 591.33 | 647 | 530 | 0.719 | -0.476 | 0.00031469 |
| AT4G39900.1 | | protein_coding | | 360 | 281 | 290 | 297 | 223 | 205 | 757 | 690 | 742 | 644 | 592 | 557 | 932.68 | 730 | 598 | 0.719 | -0.476 | 5.40E-05 |
| AT1G34340.1 | | protein_coding | alpha/beta-Hydrolases su | 322 | 277 | 308 | 207 | 166 | 175 | 677 | 681 | 788 | 449 | 441 | 476 | 801.63 | 715 | 455 | 0.719 | -0.476 | 1.99E-05 |
| AT2G41340.1 | NRPD5B | protein_coding | RNA polymerase II fifth la | 61 | 46 | 48 | 36 | 23 | 22 | 128 | 113 | 123 | 78 | 61 | 60 | 108.5 | 121 | 66 | 0.719 | -0.477 | 0.00440089 |
| AT1G17350.2 | | protein_coding | NADH:ubiquinone oxidor | 160 | 122 | 131 | 97 | 71 | 57 | 336 | 300 | 335 | 210 | 188 | 155 | 264.5 | 324 | 184 | 0.718 | -0.478 | 0.00032134 |
| AT3G05210.1 | ERCC1 | protein_coding | nucleotide repair protein, | 127 | 151 | 123 | 132 | 115 | 107 | 267 | 371 | 315 | 286 | 305 | 291 | 356.77 | 318 | 294 | 0.718 | -0.478 | 0.00010125 |
| AT4G13150.1 | | protein_coding | | 388 | 287 | 333 | 460 | 349 | 373 | 816 | 705 | 853 | 998 | 927 | 1014 | 1225.43 | 791 | 980 | 0.718 | -0.479 | 3.46E-05 |
| AT1G60160.1 | POT12 | protein_coding | Potassium transporter fa | 709 | 584 | 603 | 418 | 377 | 398 | 1490 | 1435 | 1544 | 907 | 1001 | 1082 | 1103.92 | 1490 | 997 | 0.717 | -0.479 | 2.05E-05 |
| AT1G73850.1 | | protein_coding | Protein of unknown funct | 75 | 73 | 89 | 83 | 65 | 78 | 158 | 179 | 228 | 180 | 173 | 212 | 169.21 | 188 | 188 | 0.717 | -0.48 | 0.00328105 |
| AT2G17840.1 | ERD7 | protein_coding | Senescence/dehydration- | 321 | 311 | 430 | 240 | 234 | 270 | 675 | 764 | 1101 | 521 | 621 | 734 | 889.73 | 847 | 625 | 0.717 | -0.48 | 0.0031779 |
| AT3G14960.1 | B3GALT13 | protein_coding | Galactosyltransferase fan | 150 | 150 | 130 | 92 | 90 | 96 | 315 | 369 | 333 | 200 | 239 | 261 | 297.78 | 339 | 233 | 0.717 | -0.48 | 0.00027446 |
| AT5G07590.1 | | protein_coding | Transducin/WD40 repeat | 169 | 157 | 154 | 118 | 130 | 137 | 355 | 386 | 394 | 256 | 345 | 373 | 395.69 | 378 | 325 | 0.717 | -0.48 | 0.00019831 |
| AT1G79820.1 | SGB1 | protein_coding | Major facilitator superfar | 98 | 124 | 91 | 85 | 89 | 83 | 206 | 305 | 233 | 184 | 236 | 226 | 248.25 | 248 | 215 | 0.717 | -0.481 | 0.00861461 |
| AT5G19940.1 | PAP8 | protein_coding | Plastid-lipid associated pr | 6799 | 7328 | 6785 | 4377 | 3951 | 4595 | 14292 | 18004 | 17370 | 9494 | 10489 | 12496 | 15903.17 | 16555 | 10826 | 0.717 | -0.481 | 0.00068817 |
| AT2G38130.1 | MAK3 | protein_coding | Acyl-CoA N-acyltransfera | 234 | 201 | 253 | 184 | 128 | 161 | 492 | 494 | 648 | 399 | 340 | 438 | 439.15 | 545 | 392 | 0.716 | -0.481 | 0.00053211 |
| AT4G18230.1 | | protein_coding | | 216 | 190 | 179 | 179 | 146 | 125 | 454 | 467 | 458 | 388 | 388 | 340 | 416.78 | 460 | 372 | 0.716 | -0.481 | 9.92E-05 |
| AT5G04270.1 | PAT15 | protein_coding | DHHC-type zinc finger far | 80 | 71 | 98 | 55 | 38 | 56 | 168 | 174 | 251 | 119 | 101 | 152 | 166.46 | 198 | 124 | 0.716 | -0.482 | 0.00856504 |
| AT4G18280.1 | | protein_coding | glycine-rich cell wall prot | 73 | 75 | 49 | 48 | 38 | 61 | 153 | 184 | 125 | 104 | 101 | 166 | 170.8 | 154 | 124 | 0.716 | -0.482 | 0.00495941 |
| AT5G41240.1 | GSTT2 | protein_coding | glutathione S-transferase | 190 | 121 | 208 | 98 | 94 | 101 | 399 | 297 | 532 | 213 | 250 | 275 | 302.31 | 409 | 246 | 0.716 | -0.482 | 0.00449845 |
| AT4G29820.1 | CFIS1 | protein_coding | homolog of CFIM-25 | 82 | 68 | 67 | 62 | 50 | 54 | 172 | 167 | 172 | 134 | 133 | 147 | 147.65 | 170 | 138 | 0.716 | -0.482 | 0.00105035 |
| AT5G57887.1 | | protein_coding | | 76 | 72 | 97 | 83 | 54 | 66 | 160 | 177 | 248 | 180 | 143 | 179 | 275 | 195 | 167 | 0.716 | -0.482 | 0.00100695 |
| AT5G14720.1 | | protein_coding | Protein kinase superfamil | 1740 | 1642 | 1933 | 1220 | 1166 | 1258 | 3658 | 4034 | 4949 | 2646 | 3095 | 3421 | 3058.84 | 4214 | 3054 | 0.716 | -0.482 | 0.00042322 |
| AT1G18060.1 | | protein_coding | | 1604 | 1674 | 1521 | 1193 | 1163 | 1047 | 3372 | 4113 | 3894 | 2588 | 3087 | 2847 | 4110.71 | 3793 | 2841 | 0.716 | -0.482 | 0.00010096 |
| AT1G02110.1 | | protein_coding | Protein of unknown funct | 120 | 142 | 104 | 79 | 88 | 106 | 252 | 349 | 266 | 171 | 234 | 288 | 411.51 | 289 | 231 | 0.715 | -0.483 | 0.00119657 |
| AT3G63180.1 | ATTKL | protein_coding | TIC-like | 344 | 333 | 311 | 233 | 218 | 242 | 723 | 818 | 796 | 505 | 579 | 658 | 691.33 | 779 | 581 | 0.716 | -0.483 | 2.66E-05 |
| AT3G29400.1 | ATEXO70E1 | protein_coding | exocyst subunit exo70 fa | 405 | 382 | 370 | 299 | 235 | 213 | 851 | 939 | 947 | 649 | 624 | 579 | 610.25 | 912 | 617 | 0.715 | -0.483 | 1.92E-05 |
| AT3G61080.1 | | protein_coding | Protein kinase superfamil | 535 | 583 | 544 | 421 | 401 | 381 | 1125 | 1432 | 1393 | 913 | 1065 | 1036 | 1211.94 | 1317 | 1005 | 0.715 | -0.484 | 0.0001189 |
| AT3G14940.1 | PPC3 | protein_coding | phosphoenolpyruvate car | 264 | 227 | 167 | 178 | 209 | 163 | 555 | 558 | 428 | 386 | 555 | 443 | 404.37 | 514 | 461 | 0.715 | -0.485 | 0.00409375 |
| AT5G54890.1 | | protein_coding | RNA-binding CRS1 / YhbY | 83 | 81 | 86 | 67 | 72 | 58 | 174 | 199 | 220 | 145 | 191 | 158 | 182.35 | 198 | 165 | 0.715 | -0.485 | 0.00231255 |
| AT1G14340.1 | | protein_coding | RNA-binding (RRM/RBD/ | 116 | 96 | 90 | 77 | 70 | 54 | 244 | 236 | 230 | 167 | 186 | 147 | 206.34 | 237 | 167 | 0.714 | -0.485 | 0.00088905 |
| AT1G28710.1 | | protein_coding | Nucleotide-diphospho-su | 160 | 150 | 104 | 118 | 111 | 126 | 336 | 369 | 266 | 256 | 295 | 343 | 416.42 | 324 | 298 | 0.715 | -0.485 | 0.00060786 |
| AT1G20630.1 | CAT1 | protein_coding | catalase 1 | 1265 | 1189 | 1231 | 774 | 647 | 775 | 2659 | 2921 | 3151 | 1679 | 1718 | 2108 | 2642.22 | 2910 | 1835 | 0.714 | -0.485 | 0.00014008 |
| AT5G11040.1 | TRS120 | protein_coding | TRS120 | 876 | 709 | 591 | 539 | 508 | 431 | 1841 | 1742 | 1513 | 1169 | 1349 | 1172 | 1330.03 | 1699 | 1230 | 0.714 | -0.485 | 8.74E-05 |
| AT5G02430.1 | | protein_coding | Transducin/WD40 repeat | 65 | 80 | 66 | 60 | 43 | 49 | 137 | 197 | 169 | 130 | 114 | 133 | 127.46 | 168 | 126 | 0.714 | -0.486 | 0.00609367 |
| AT3G15040.1 | | protein_coding | Protein of unknown funct | 94 | 135 | 119 | 73 | 62 | 97 | 198 | 332 | 305 | 158 | 165 | 264 | 341.41 | 278 | 196 | 0.714 | -0.486 | 0.00438657 |
| AT3G14450.1 | CID9 | protein_coding | CTC-interacting domain 9 | 145 | 124 | 100 | 86 | 84 | 70 | 305 | 305 | 256 | 187 | 223 | 190 | 260.06 | 289 | 200 | 0.714 | -0.486 | 0.00033524 |
| AT1G52220.1 | CURT1C | protein_coding | | 5938 | 5406 | 5158 | 3874 | 3380 | 2981 | 12482 | 13282 | 13205 | 8403 | 8973 | 8107 | 12789.93 | 12990 | 8494 | 0.714 | -0.486 | 5.06E-05 |
| AT1G08620.1 | JMJ16 | protein_coding | Transcription factor jumo | 730 | 622 | 664 | 538 | 499 | 461 | 1535 | 1528 | 1700 | 1167 | 1325 | 1254 | 1036.81 | 1588 | 1249 | 0.714 | -0.486 | 3.68E-05 |
| AT1G52340.1 | ABA2 | protein_coding | NAD(P)-binding Rossman | 198 | 189 | 155 | 131 | 133 | 116 | 416 | 464 | 397 | 284 | 353 | 315 | 393.54 | 426 | 317 | 0.714 | -0.486 | 3.04E-05 |
| AT5G62930.1 | | protein_coding | SGNH hydrolase-type est | 206 | 188 | 164 | 136 | 104 | 119 | 433 | 462 | 420 | 295 | 276 | 324 | 473.26 | 438 | 298 | 0.714 | -0.486 | 1.04E-05 |
| AT5G48470.1 | | protein_coding | | 549 | 343 | 272 | 367 | 268 | 205 | 1154 | 843 | 696 | 796 | 711 | 557 | 790.47 | 898 | 688 | 0.714 | -0.487 | 0.00281912 |
| AT2G46240.1 | BAG6 | protein_coding | BCL-2-associated athano | 251 | 284 | 290 | 162 | 160 | 152 | 528 | 698 | 742 | 351 | 425 | 413 | 447.67 | 656 | 396 | 0.713 | -0.487 | 0.00205782 |
| AT1G19270.1 | DA1 | protein_coding | DA1 | 173 | 181 | 192 | 110 | 121 | 100 | 364 | 445 | 492 | 239 | 321 | 272 | 310.59 | 434 | 277 | 0.714 | -0.487 | 0.00119129 |
| AT1G68080.1 | | protein_coding | 2-oxoglutarate (2OG) and | 105 | 88 | 92 | 82 | 63 | 63 | 221 | 216 | 236 | 178 | 167 | 171 | 220 | 224 | 172 | 0.714 | -0.487 | 0.00054847 |
| AT3G53250.1 | | protein_coding | SAUR-like auxin-responsi | 55 | 59 | 50 | 32 | 24 | 25 | 116 | 145 | 128 | 69 | 64 | 68 | 106.46 | 130 | 67 | 0.713 | -0.488 | 0.00661953 |
| AT4G34950.1 | | protein_coding | Major facilitator superfar | 1246 | 1020 | 680 | 532 | 345 | 344 | 2619 | 2506 | 1741 | 1154 | 916 | 935 | 2037.02 | 2289 | 1002 | 0.713 | -0.488 | 0.00142033 |
| AT1G64850.1 | | protein_coding | Calcium-binding EF hand | 510 | 406 | 389 | 448 | 327 | 267 | 1072 | 997 | 996 | 972 | 868 | 726 | 1033.03 | 1022 | 855 | 0.713 | -0.488 | 8.54E-05 |
| AT5G41620.1 | | protein_coding | | 34 | 48 | 44 | 33 | 30 | 24 | 71 | 118 | 113 | 72 | 80 | 65 | 111.57 | 101 | 72 | 0.712 | -0.489 | 0.00597443 |
| AT4G11330.1 | MPK5 | protein_coding | MAP kinase 5 | 309 | 340 | 390 | 259 | 236 | 265 | 650 | 835 | 998 | 562 | 627 | 721 | 638.61 | 828 | 637 | 0.713 | -0.489 | 0.00177313 |
| AT2G19340.1 | | protein_coding | Oligosaccharyltransferase | 85 | 83 | 77 | 60 | 49 | 55 | 179 | 204 | 197 | 130 | 130 | 150 | 195.67 | 193 | 137 | 0.712 | -0.489 | 0.00056798 |
| AT3G47590.1 | | protein_coding | alpha/beta-Hydrolases su | 315 | 190 | 216 | 150 | 115 | 117 | 662 | 467 | 553 | 325 | 305 | 318 | 369.78 | 561 | 316 | 0.713 | -0.489 | 0.00025164 |
| AT5G06100.2 | MYB33 | protein_coding | myb domain protein 33 | 226 | 182 | 190 | 174 | 132 | 112 | 475 | 447 | 486 | 377 | 350 | 305 | 392.15 | 469 | 344 | 0.713 | -0.489 | 0.00011889 |
| AT3G06300.1 | P4H2 | protein_coding | P4H isoform 2 | 186 | 148 | 105 | 146 | 102 | 105 | 391 | 364 | 269 | 317 | 271 | 286 | 325.7 | 341 | 291 | 0.712 | -0.49 | 0.00067963 |
| AT3G07080.1 | | protein_coding | EamA-like transporter far | 208 | 213 | 225 | 177 | 142 | 113 | 437 | 523 | 576 | 384 | 377 | 307 | 397.64 | 512 | 356 | 0.712 | -0.49 | 0.00041156 |
| AT5G41220.1 | GSTT3 | protein_coding | glutathione S-transferase | 229 | 180 | 233 | 153 | 136 | 151 | 481 | 442 | 596 | 332 | 361 | 411 | 393.76 | 506 | 368 | 0.712 | -0.49 | 0.00025763 |
| AT2G21240.1 | BPC4 | protein_coding | basic pentacysteine 4 | 398 | 377 | 463 | 287 | 256 | 315 | 837 | 926 | 1185 | 623 | 680 | 857 | 856.61 | 983 | 720 | 0.711 | -0.491 | 0.00079464 |
| AT2G28720.1 | | protein_coding | Histone superfamily prot | 837 | 730 | 886 | 415 | 445 | 443 | 1759 | 1793 | 2268 | 900 | 1181 | 1205 | 1665.91 | 1940 | 1095 | 0.711 | -0.492 | 0.00069958 |
| AT2G36410.1 | | protein_coding | Family of unknown functi | 780 | 856 | 942 | 617 | 539 | 605 | 1640 | 2103 | 2412 | 1338 | 1431 | 1645 | 1849.73 | 2052 | 1471 | 0.711 | -0.492 | 0.00036958 |
| AT2G01870.1 | | protein_coding | | 383 | 302 | 336 | 275 | 218 | 155 | 805 | 742 | 860 | 597 | 579 | 422 | 644.45 | 802 | 533 | 0.711 | -0.492 | 0.00025536 |
| AT1G50320.1 | ATHX | protein_coding | thioredoxin X | 1449 | 1653 | 1290 | 1195 | 1112 | 1255 | 3046 | 4061 | 3302 | 2592 | 2952 | 3413 | 4718.65 | 3470 | 2986 | 0.711 | -0.492 | 0.00022579 |
| AT1G34010.1 | | protein_coding | | 176 | 150 | 162 | 125 | 123 | 100 | 370 | 369 | 415 | 271 | 327 | 272 | 363.97 | 385 | 290 | 0.711 | -0.492 | 7.90E-05 |
| AT5G55710.1 | TIC20-V | protein_coding | | 665 | 533 | 471 | 480 | 361 | 338 | 1398 | 1309 | 1206 | 1041 | 958 | 919 | 1794.05 | 1304 | 973 | 0.711 | -0.492 | 1.69E-05 |
| AT2G42640.1 | | protein_coding | Mitogen activated protei | 90 | 65 | 81 | 58 | 49 | 48 | 189 | 160 | 207 | 126 | 130 | 131 | 134.35 | 185 | 129 | 0.711 | -0.493 | 0.00986327 |
| AT2G25300.1 | B3GALT9 | protein_coding | Galactosyltransferase fan | 118 | 97 | 116 | 80 | 79 | 70 | 248 | 238 | 297 | 174 | 210 | 190 | 195.28 | 261 | 191 | 0.71 | -0.493 | 0.00106133 |
| AT5G60710.1 | | protein_coding | Zinc finger (C3HC4-type R | 797 | 764 | 625 | 596 | 433 | 464 | 1675 | 1877 | 1600 | 1293 | 1150 | 1262 | 1576.93 | 1717 | 1235 | 0.711 | -0.493 | 4.19E-05 |
| AT5G46340.1 | | protein_coding | O-acetyltransferase famil | 320 | 237 | 272 | 247 | 225 | 207 | 673 | 582 | 696 | 536 | 597 | 563 | 586.65 | 650 | 565 | 0.711 | -0.493 | 2.32E-05 |
| AT5G54750.2 | | protein_coding | Transport protein particle | 352 | 276 | 275 | 264 | 198 | 203 | 740 | 678 | 704 | 573 | 526 | 552 | 718.09 | 707 | 550 | 0.71 | -0.493 | 2.09E-06 |
| AT4G36515.1 | | protein_coding | | 52 | 48 | 35 | 39 | 31 | 32 | 109 | 118 | 90 | 85 | 82 | 87 | 115.67 | 106 | 85 | 0.71 | -0.494 | 0.00306688 |
| AT1G08390.1 | | protein_coding | | 51 | 70 | 48 | 37 | 30 | 43 | 107 | 172 | 123 | 80 | 80 | 117 | 168.01 | 134 | 92 | 0.709 | -0.495 | 0.00441937 |
| AT2G30230.1 | | protein_coding | | 664 | 510 | 567 | 810 | 608 | 424 | 1396 | 1253 | 1452 | 1757 | 1614 | 1153 | 802.61 | 1367 | 1508 | 0.709 | -0.495 | 0.00389115 |
| AT5G26340.1 | STP13 | protein_coding | Major facilitator superfar | 234 | 192 | 142 | 121 | 82 | 110 | 492 | 472 | 364 | 262 | 218 | 299 | 334.77 | 443 | 260 | 0.709 | -0.495 | 0.00070375 |
| AT1G63800.1 | UBC5 | protein_coding | ubiquitin-conjugating enz | 418 | 456 | 667 | 307 | 272 | 349 | 879 | 1120 | 1708 | 666 | 722 | 949 | 969.1 | 1236 | 779 | 0.709 | -0.496 | 0.00825886 |
| AT5G07810.1 | | protein_coding | SNF2 domain-containing | 140 | 144 | 167 | 129 | 145 | 101 | 294 | 354 | 428 | 280 | 385 | 275 | 258.55 | 359 | 313 | 0.709 | -0.496 | 0.00490384 |
| AT5G62680.1 | NPF2.11 | protein_coding | Major facilitator superfar | 734 | 675 | 566 | 684 | 771 | 609 | 1543 | 1658 | 1449 | 1484 | 2047 | 1656 | 1256.94 | 1550 | 1729 | 0.709 | -0.496 | 0.00022157 |
| AT1G17280.1 | UBC34 | protein_coding | ubiquitin-conjugating enz | 430 | 405 | 471 | 340 | 293 | 284 | 904 | 995 | 1206 | 737 | 778 | 772 | 905.56 | 1035 | 762 | 0.709 | -0.496 | 0.00012536 |
| AT2G17140.1 | | protein_coding | Pentatricopeptide repeat | 77 | 54 | 57 | 58 | 66 | 37 | 162 | 133 | 146 | 126 | 175 | 101 | 133.07 | 147 | 134 | 0.708 | -0.497 | 0.00625865 |

| | | | | | | | | | | | | | | | | | | | | | |
|---|---|---|---|---|---|---|---|---|---|---|---|---|---|---|---|---|---|---|---|---|---|
| AT5G50550.1 | | protein_coding | Transducin/WD40 repeat | 142 | 90 | 120 | 80 | 51 | 46 | 298 | 221 | 307 | 174 | 135 | 125 | 229.01 | 275 | 145 | 0.709 | -0.497 | 0.00113037 |
| AT4G25780.1 | | protein_coding | CAP (Cysteine-rich secret | 154 | 150 | 114 | 109 | 88 | 91 | 324 | 369 | 292 | 236 | 234 | 247 | 347.9 | 328 | 239 | 0.708 | -0.497 | 0.00032126 |
| AT1G07970.1 | | protein_coding | | 189 | 172 | 158 | 174 | 135 | 113 | 397 | 423 | 404 | 377 | 358 | 307 | 398.97 | 408 | 347 | 0.708 | -0.497 | 2.78E-05 |
| AT1G20160.1 | CRSP | protein_coding | Subtilisin-like serine endo | 80 | 79 | 60 | 146 | 138 | 140 | 168 | 194 | 154 | 317 | 366 | 381 | 174.91 | 172 | 355 | 0.708 | -0.498 | 0.00818542 |
| AT1G01230.1 | | protein_coding | ORMDL family protein | 310 | 282 | 342 | 260 | 229 | 286 | 652 | 693 | 876 | 564 | 608 | 778 | 687.41 | 740 | 650 | 0.708 | -0.498 | 0.00030399 |
| AT1G01820.1 | PEX11C | protein_coding | peroxin 11c | 537 | 479 | 492 | 386 | 257 | 282 | 1129 | 1177 | 1260 | 837 | 682 | 767 | 1127.48 | 1189 | 762 | 0.708 | -0.498 | 3.42E-05 |
| AT2G36960.3 | TKI1 | protein_coding | TSL-kinase interacting pro | 273 | 250 | 228 | 179 | 155 | 180 | 574 | 614 | 584 | 388 | 411 | 489 | 429.64 | 591 | 429 | 0.708 | -0.499 | 0.00012358 |
| AT2G27960.1 | CKS1 | protein_coding | cyclin-dependent kinase-s | 257 | 264 | 248 | 190 | 182 | 209 | 540 | 649 | 635 | 412 | 483 | 568 | 722.47 | 608 | 488 | 0.708 | -0.499 | 3.67E-05 |
| AT5G36120.1 | CCB3 | protein_coding | cofactor assembly, compl | 705 | 601 | 545 | 522 | 417 | 322 | 1482 | 1477 | 1395 | 1132 | 1107 | 876 | 1672.82 | 1451 | 1038 | 0.708 | -0.499 | 3.40E-05 |
| AT4G35750.1 | | protein_coding | SEC14 cytosolic factor far | 897 | 748 | 941 | 717 | 543 | 923 | 1886 | 1838 | 2409 | 1555 | 1442 | 2510 | 1878.56 | 2044 | 1836 | 0.707 | -0.5 | 0.00614519 |
| AT2G47020.1 | | protein_coding | Peptide chain release fact | 109 | 90 | 101 | 60 | 52 | 60 | 229 | 221 | 259 | 130 | 138 | 163 | 173.23 | 236 | 144 | 0.707 | -0.5 | 0.00314783 |
| AT1G05562.1 | | other_rna | | 123 | 166 | 113 | 121 | 87 | 103 | 259 | 408 | 289 | 262 | 231 | 280 | 335.39 | 319 | 258 | 0.706 | -0.501 | 0.00226097 |
| AT4G19390.1 | | protein_coding | Uncharacterised protein f | 95 | 108 | 119 | 79 | 58 | 94 | 200 | 265 | 305 | 171 | 154 | 256 | 276.67 | 257 | 194 | 0.707 | -0.501 | 0.00214884 |
| AT3G20510.1 | FAX6 | protein_coding | Transmembrane proteins | 150 | 120 | 103 | 165 | 109 | 112 | 315 | 295 | 264 | 358 | 289 | 305 | 350.35 | 291 | 317 | 0.707 | -0.501 | 0.00076234 |
| AT4G32710.1 | PERK14 | protein_coding | Protein kinase superfamil | 195 | 147 | 126 | 137 | 103 | 91 | 410 | 361 | 323 | 297 | 273 | 247 | 386.83 | 365 | 272 | 0.707 | -0.501 | 2.73E-05 |
| AT5G44510.1 | TAO1 | protein_coding | target of AVRB operation | 569 | 517 | 501 | 345 | 313 | 234 | 1196 | 1270 | 1283 | 748 | 831 | 636 | 957.17 | 1250 | 738 | 0.706 | -0.502 | 8.37E-05 |
| AT5G23750.1 | | protein_coding | Remorin family protein | 118 | 130 | 106 | 112 | 82 | 78 | 248 | 319 | 271 | 243 | 218 | 212 | 320.98 | 279 | 224 | 0.706 | -0.503 | 0.00235347 |
| AT2G16365.1 | | protein_coding | F-box family protein | 776 | 675 | 960 | 553 | 405 | 502 | 1631 | 1658 | 2458 | 1200 | 1075 | 1365 | 1360.45 | 1916 | 1213 | 0.706 | -0.503 | 0.00154245 |
| AT5G44010.1 | | protein_coding | | 106 | 90 | 99 | 79 | 94 | 66 | 223 | 221 | 253 | 171 | 250 | 179 | 226.56 | 232 | 200 | 0.706 | -0.503 | 0.00144158 |
| AT2G25920.1 | | protein_coding | | 99 | 132 | 107 | 80 | 66 | 76 | 208 | 324 | 274 | 174 | 175 | 207 | 279.95 | 269 | 185 | 0.706 | -0.503 | 0.00065273 |
| AT4G03415.1 | | protein_coding | Protein phosphatase 2C f | 583 | 567 | 573 | 430 | 420 | 481 | 1226 | 1393 | 1467 | 933 | 1115 | 1308 | 1214.43 | 1362 | 1119 | 0.706 | -0.503 | 0.00018897 |
| AT1G13270.1 | MAP1B | protein_coding | methionine aminopeptida | 939 | 757 | 746 | 616 | 492 | 438 | 1974 | 1860 | 1910 | 1336 | 1306 | 1191 | 1681.43 | 1915 | 1278 | 0.706 | -0.503 | 9.36E-06 |
| AT1G66410.2 | CAM4 | protein_coding | calmodulin 4 | 1222 | 1150 | 1068 | 956 | 797 | 769 | 2569 | 2825 | 2734 | 2074 | 2116 | 2091 | 2681.25 | 2709 | 2094 | 0.706 | -0.503 | 4.80E-06 |
| AT4G30850.1 | HHP2 | protein_coding | heptahelical transmembr | 66 | 70 | 42 | 67 | 56 | 44 | 139 | 172 | 108 | 145 | 149 | 120 | 211.12 | 140 | 138 | 0.705 | -0.504 | 0.00246299 |
| AT1G78460.1 | | protein_coding | SOUL heme-binding famil | 404 | 395 | 360 | 495 | 255 | 406 | 849 | 970 | 922 | 1074 | 677 | 1104 | 1259.47 | 914 | 952 | 0.705 | -0.504 | 0.00085269 |
| AT5G35560.1 | | protein_coding | DENN (AEX-3) domain-co | 782 | 781 | 750 | 603 | 557 | 480 | 1644 | 1919 | 1920 | 1308 | 1479 | 1305 | 1459.36 | 1828 | 1364 | 0.705 | -0.504 | 6.41E-05 |
| AT5G10620.1 | | protein_coding | methyltransferases | 76 | 88 | 66 | 49 | 47 | 56 | 160 | 216 | 169 | 106 | 125 | 152 | 201.38 | 182 | 128 | 0.705 | -0.505 | 0.00072261 |
| AT4G03205.2 | CPX2 | protein_coding | Coproporphyrinogen III o: | 295 | 310 | 296 | 226 | 202 | 227 | 620 | 762 | 758 | 490 | 536 | 617 | 633.18 | 713 | 548 | 0.705 | -0.505 | 1.56E-05 |
| AT4G16566.1 | HINT4 | protein_coding | histidine triad nucleotide | 151 | 111 | 98 | 130 | 63 | 68 | 317 | 273 | 251 | 282 | 167 | 185 | 282.34 | 280 | 211 | 0.704 | -0.506 | 0.00150525 |
| AT1G08350.2 | TMN5 | protein_coding | Endomembrane protein 7 | 203 | 140 | 140 | 101 | 95 | 74 | 427 | 344 | 358 | 219 | 252 | 201 | 283.98 | 376 | 224 | 0.704 | -0.506 | 0.00054442 |
| AT1G18335.1 | | protein_coding | Acyl-CoA N-acyltransfera: | 137 | 103 | 122 | 84 | 60 | 76 | 288 | 253 | 312 | 182 | 159 | 207 | 223.39 | 284 | 183 | 0.704 | -0.506 | 0.00037898 |
| AT3G21260.3 | GLTP3 | protein_coding | Glycolipid transfer protei | 90 | 89 | 86 | 86 | 68 | 61 | 189 | 219 | 220 | 187 | 181 | 166 | 200.9 | 209 | 178 | 0.704 | -0.506 | 0.00037205 |
| AT1G18750.1 | AGL65 | protein_coding | AGAMOUS-like 65 | 74 | 45 | 72 | 42 | 43 | 36 | 156 | 111 | 184 | 91 | 114 | 98 | 111.58 | 150 | 101 | 0.703 | -0.507 | 0.00737999 |
| AT5G19140.1 | ATAILP1 | protein_coding | Aluminium induced prote | 3412 | 4556 | 4410 | 2756 | 3247 | 2988 | 7172 | 11193 | 11290 | 5978 | 8620 | 8126 | 10859.23 | 9885 | 7575 | 0.704 | -0.507 | 0.00359764 |
| AT4G22840.1 | BASS6 | protein_coding | Sodium Bile acid symport | 64 | 41 | 42 | 29 | 23 | 21 | 135 | 101 | 108 | 63 | 61 | 57 | 114.82 | 115 | 60 | 0.704 | -0.507 | 0.00309581 |
| AT2G43190.1 | | protein_coding | ribonuclease P family pro | 96 | 77 | 79 | 89 | 56 | 66 | 202 | 189 | 202 | 193 | 149 | 179 | 167.15 | 198 | 174 | 0.704 | -0.507 | 0.00136087 |
| AT5G02370.1 | | protein_coding | ATP binding microtubule | 187 | 237 | 227 | 168 | 147 | 105 | 393 | 582 | 581 | 364 | 390 | 286 | 356.71 | 519 | 347 | 0.703 | -0.508 | 0.0071606 |
| AT4G09020.1 | ISA3 | protein_coding | isoamylase 3 | 832 | 899 | 787 | 599 | 663 | 519 | 1749 | 2209 | 2015 | 1299 | 1760 | 1411 | 1967.46 | 1991 | 1490 | 0.703 | -0.508 | 0.00040713 |
| AT1G30590.1 | | protein_coding | RNA polymerase I specific | 766 | 601 | 646 | 491 | 461 | 376 | 1610 | 1477 | 1654 | 1065 | 1224 | 1023 | 1376.33 | 1580 | 1104 | 0.703 | -0.508 | 1.20E-05 |
| AT5G16720.1 | MYOB3 | protein_coding | Protein of unknown funct | 167 | 178 | 219 | 173 | 127 | 139 | 351 | 437 | 561 | 375 | 337 | 378 | 349.46 | 450 | 363 | 0.703 | -0.509 | 0.00079062 |
| AT2G23310.1 | RER1C | protein_coding | Rer1 family protein | 342 | 260 | 291 | 201 | 197 | 194 | 719 | 639 | 745 | 436 | 523 | 528 | 725.05 | 701 | 496 | 0.703 | -0.509 | 8.65E-05 |
| AT3G23170.1 | | protein_coding | | 91 | 130 | 61 | 86 | 71 | 90 | 191 | 319 | 156 | 187 | 188 | 245 | 355.17 | 222 | 207 | 0.702 | -0.51 | 0.00317839 |
| AT2G16710.3 | | protein_coding | Iron-sulphur cluster biosy | 134 | 160 | 159 | 120 | 105 | 94 | 282 | 393 | 407 | 260 | 279 | 256 | 339.45 | 361 | 265 | 0.702 | -0.51 | 0.00032224 |
| AT2G40070.1 | | protein_coding | | 321 | 310 | 339 | 246 | 219 | 261 | 675 | 762 | 868 | 534 | 581 | 710 | 817.04 | 768 | 608 | 0.702 | -0.51 | 5.61E-05 |
| AT4G21215.2 | | protein_coding | | 234 | 215 | 217 | 168 | 205 | 185 | 492 | 528 | 556 | 364 | 544 | 503 | 565.87 | 525 | 470 | 0.702 | -0.511 | 0.0002107 |
| AT3G45100.1 | SETH2 | protein_coding | UDP-Glycosyltransferase | 213 | 184 | 186 | 120 | 112 | 126 | 448 | 452 | 476 | 260 | 297 | 343 | 366.94 | 459 | 300 | 0.702 | -0.511 | 2.97E-05 |
| AT2G24040.1 | | protein_coding | Low temperature and sal | 75 | 53 | 52 | 63 | 48 | 55 | 158 | 130 | 133 | 137 | 127 | 150 | 148.25 | 140 | 138 | 0.701 | -0.512 | 0.00070497 |
| AT5G08050.1 | | protein_coding | Protein of unknown funct | 3058 | 3121 | 2328 | 2263 | 1971 | 1818 | 6428 | 7668 | 5960 | 4909 | 5233 | 4944 | 7213.56 | 6685 | 5029 | 0.701 | -0.512 | 4.33E-05 |
| AT3G23200.1 | | protein_coding | Uncharacterised protein f | 44 | 27 | 34 | 32 | 28 | 27 | 92 | 66 | 87 | 69 | 74 | 73 | 81.38 | 82 | 72 | 0.701 | -0.513 | 0.00731885 |
| AT2G05160.1 | | protein_coding | CCCH-type zinc fingerfam | 156 | 174 | 174 | 85 | 99 | 67 | 328 | 427 | 445 | 184 | 263 | 182 | 258.15 | 400 | 210 | 0.701 | -0.513 | 0.00473989 |
| AT2G04890.1 | SCL21 | protein_coding | SCARECROW-like 21 | 131 | 125 | 111 | 108 | 57 | 79 | 275 | 307 | 284 | 234 | 151 | 215 | 218.8 | 289 | 200 | 0.701 | -0.513 | 0.00224489 |
| AT3G04970.1 | PAT17 | protein_coding | DHHC-type zinc finger far | 80 | 77 | 83 | 60 | 54 | 57 | 168 | 189 | 212 | 130 | 143 | 155 | 174.43 | 190 | 143 | 0.701 | -0.513 | 0.00214944 |
| AT3G59840.1 | | protein_coding | | 157 | 162 | 86 | 142 | 111 | 135 | 330 | 398 | 220 | 308 | 295 | 367 | 457.18 | 316 | 323 | 0.701 | -0.513 | 0.00178777 |
| AT2G28305.1 | LOG1 | protein_coding | Putative lysine decarboxy | 281 | 203 | 199 | 216 | 158 | 124 | 591 | 499 | 509 | 469 | 419 | 337 | 499.45 | 533 | 408 | 0.701 | -0.513 | 0.00036436 |
| AT1G72175.1 | | protein_coding | RING/U-box protein with | 90 | 72 | 82 | 59 | 49 | 54 | 189 | 177 | 210 | 128 | 130 | 147 | 197.98 | 192 | 135 | 0.701 | -0.513 | 0.00010146 |
| AT3G23280.1 | XBAT35 | protein_coding | XB3 ortholog 5 in Arabido | 842 | 984 | 1024 | 559 | 599 | 764 | 1770 | 2418 | 2622 | 1213 | 1590 | 2078 | 2039.98 | 2270 | 1627 | 0.7 | -0.514 | 0.00368037 |
| AT1G18150.2 | MPK8 | protein_coding | Protein kinase superfamil | 742 | 790 | 675 | 502 | 550 | 380 | 1560 | 1941 | 1728 | 1089 | 1460 | 1033 | 1742.52 | 1743 | 1194 | 0.7 | -0.514 | 0.00040159 |
| AT5G65250.1 | | protein_coding | | 292 | 262 | 250 | 209 | 144 | 181 | 614 | 644 | 640 | 453 | 382 | 492 | 617.35 | 633 | 442 | 0.701 | -0.514 | 6.79E-06 |
| AT3G04445.1 | | pseudogene | | 69 | 66 | 69 | 57 | 31 | 47 | 145 | 162 | 177 | 124 | 82 | 128 | 112.7 | 161 | 111 | 0.7 | -0.515 | 0.00507154 |
| AT2G17290.1 | CPK6 | protein_coding | Calcium-dependent prote | 357 | 210 | 266 | 194 | 103 | 110 | 750 | 516 | 681 | 421 | 273 | 299 | 383.69 | 649 | 331 | 0.7 | -0.515 | 0.00257042 |
| AT4G18270.1 | ATTRANS | protein_coding | translocase 11 | 101 | 139 | 99 | 85 | 99 | 53 | 212 | 341 | 253 | 184 | 263 | 144 | 346.82 | 269 | 197 | 0.7 | -0.515 | 0.00204786 |
| AT1G72890.2 | | protein_coding | Disease resistance protei | 165 | 143 | 113 | 128 | 124 | 75 | 347 | 351 | 289 | 278 | 329 | 204 | 257.39 | 329 | 270 | 0.699 | -0.516 | 0.00579853 |
| AT4G10140.1 | | protein_coding | | 110 | 115 | 142 | 104 | 77 | 97 | 231 | 283 | 364 | 226 | 204 | 264 | 278.55 | 293 | 231 | 0.699 | -0.517 | 0.00163983 |
| AT1G10090.1 | | protein_coding | Early-responsive to dehyc | 710 | 692 | 688 | 498 | 495 | 416 | 1492 | 1700 | 1761 | 1080 | 1314 | 1131 | 1211.29 | 1651 | 1175 | 0.699 | -0.517 | 4.16E-05 |
| AT5G46880.1 | HDG5 | protein_coding | homeobox-7 | 180 | 181 | 150 | 114 | 103 | 109 | 378 | 445 | 384 | 247 | 273 | 296 | 300.96 | 402 | 272 | 0.698 | -0.518 | 0.00018488 |
| AT4G01690.1 | PPOX1 | protein_coding | Flavin containing amine c | 3100 | 3118 | 2487 | 1876 | 1539 | 1768 | 6516 | 7660 | 6367 | 4069 | 4086 | 4808 | 6570.21 | 6848 | 4321 | 0.698 | -0.519 | 1.69E-05 |
| AT1G52040.1 | MBP1 | protein_coding | myrosinase-binding prote | 104 | 85 | 86 | 28 | 33 | 29 | 219 | 209 | 220 | 61 | 88 | 79 | 188.93 | 216 | 76 | 0.697 | -0.521 | 0.00090823 |
| AT1G22070.1 | TGA3 | protein_coding | TGA1A-related gene 3 | 512 | 373 | 294 | 304 | 205 | 181 | 1076 | 916 | 753 | 659 | 544 | 492 | 756.56 | 915 | 565 | 0.697 | -0.521 | 0.00025322 |
| AT4G14440.1 | ECI3 | protein_coding | 3-hydroxyacyl-CoA dehyd | 478 | 487 | 338 | 666 | 525 | 622 | 1005 | 1196 | 865 | 1445 | 1394 | 1691 | 2076.83 | 1022 | 1510 | 0.697 | -0.521 | 0.00018541 |
| AT3G22300.1 | RPS10 | protein_coding | ribosomal protein S10 | 279 | 230 | 140 | 218 | 126 | 134 | 586 | 565 | 358 | 473 | 335 | 364 | 543.13 | 503 | 391 | 0.696 | -0.522 | 0.00081492 |
| AT4G13930.1 | SHM4 | protein_coding | serine hydroxymethyltrar | 4907 | 3904 | 2651 | 5795 | 4072 | 3854 | 10315 | 9591 | 6787 | 12570 | 10810 | 10481 | 11191.15 | 8898 | 11287 | 0.696 | -0.522 | 0.00070107 |
| AT1G09450.1 | | protein_coding | Protein kinase superfamil | 103 | 97 | 98 | 80 | 79 | 64 | 217 | 238 | 251 | 174 | 210 | 174 | 181.13 | 235 | 186 | 0.697 | -0.522 | 0.0006912 |
| AT2G17150.1 | NLP1 | protein_coding | Plant regulator RWP-RK f | 253 | 188 | 222 | 147 | 139 | 151 | 532 | 462 | 568 | 319 | 369 | 411 | 353.79 | 521 | 366 | 0.696 | -0.522 | 6.33E-05 |
| AT3G62930.1 | GRXS6 | protein_coding | Thioredoxin superfamily p | 45 | 47 | 42 | 52 | 40 | 52 | 95 | 115 | 108 | 113 | 106 | 141 | 110.31 | 106 | 120 | 0.696 | -0.523 | 0.0029861 |
| AT3G16350.1 | | protein_coding | Homeodomain-like super | 137 | 112 | 85 | 97 | 72 | 75 | 288 | 275 | 218 | 210 | 191 | 204 | 228.7 | 260 | 202 | 0.696 | -0.523 | 0.00079971 |
| AT2G37035.1 | | protein_coding | | 157 | 159 | 143 | 137 | 100 | 101 | 330 | 391 | 366 | 297 | 265 | 275 | 326.6 | 362 | 279 | 0.696 | -0.523 | 0.00012083 |
| AT1G04290.1 | | protein_coding | Thioesterase superfamily | 238 | 167 | 158 | 190 | 146 | 120 | 500 | 410 | 404 | 412 | 388 | 326 | 467.12 | 438 | 375 | 0.696 | -0.523 | 2.15E-05 |
| AT3G07640.1 | | protein_coding | | 273 | 257 | 208 | 194 | 167 | 162 | 574 | 631 | 532 | 421 | 443 | 441 | 564.98 | 579 | 435 | 0.696 | -0.523 | 1.67E-05 |
| AT4G23790.1 | TBL24 | protein_coding | TRICHOME BIREFRINGEN | 260 | 212 | 194 | 162 | 144 | 127 | 547 | 521 | 497 | 351 | 382 | 345 | 417.82 | 522 | 359 | 0.696 | -0.523 | 1.40E-05 |
| AT4G12340.1 | | protein_coding | copper ion binding | 466 | 300 | 274 | 306 | 190 | 230 | 980 | 737 | 701 | 664 | 504 | 625 | 686.78 | 806 | 598 | 0.695 | -0.524 | 0.00010638 |
| AT3G09160.1 | | protein_coding | RNA-binding (RRM/RBD/ | 69 | 92 | 89 | 73 | 57 | 66 | 145 | 226 | 228 | 158 | 151 | 179 | 278.35 | 200 | 163 | 0.695 | -0.525 | 0.0004966 |
| AT5G37500.1 | GORK | protein_coding | gated outwardly-rectifyin | 107 | 101 | 129 | 41 | 56 | 63 | 225 | 248 | 330 | 89 | 149 | 171 | 189.74 | 268 | 136 | 0.695 | -0.526 | 0.00474339 |
| AT5G50650.1 | | protein_coding | Transducin/WD40 repeat | 128 | 126 | 116 | 70 | 57 | 68 | 269 | 310 | 297 | 152 | 151 | 185 | 239.07 | 292 | 163 | 0.695 | -0.526 | 0.00110433 |
| AT4G18890.1 | BEH3 | protein_coding | BES1/BZR1 homolog 3 | 86 | 107 | 89 | 105 | 87 | 66 | 181 | 263 | 228 | 228 | 231 | 179 | 201.58 | 224 | 213 | 0.695 | -0.526 | 0.00088168 |
| AT3G05570.1 | | protein_coding | | 95 | 84 | 82 | 57 | 50 | 42 | 200 | 206 | 210 | 124 | 133 | 114 | 183.78 | 205 | 124 | 0.694 | -0.526 | 0.00052284 |
| AT5G03500.1 | MED7B | protein_coding | Mediator complex, subur | 177 | 140 | 151 | 124 | 88 | 79 | 372 | 344 | 387 | 269 | 234 | 215 | 284.39 | 368 | 239 | 0.694 | -0.526 | 8.83E-05 |
| AT3G20150.1 | | protein_coding | Kinesin motor family prot | 478 | 397 | 407 | 341 | 339 | 224 | 1005 | 975 | 1042 | 740 | 900 | 609 | 686.93 | 1007 | 750 | 0.695 | -0.526 | 6.59E-05 |
| AT5G04170.1 | CML50 | protein_coding | Calcium-binding EF-hand | 386 | 498 | 428 | 313 | 291 | 396 | 811 | 1223 | 1096 | 679 | 773 | 1077 | 1079.5 | 1043 | 843 | 0.694 | -0.527 | 0.00140346 |
| AT5G47790.1 | | protein_coding | SMAD/FHA domain-conta | 200 | 160 | 144 | 122 | 106 | 125 | 420 | 393 | 369 | 265 | 281 | 340 | 342.75 | 394 | 295 | 0.694 | -0.527 | 0.00012949 |
| AT1G29240.1 | | protein_coding | Protein of unknown funct | 57 | 60 | 68 | 55 | 66 | 47 | 120 | 147 | 174 | 119 | 175 | 128 | 111.66 | 147 | 141 | 0.694 | -0.528 | 0.00780919 |
| AT2G39690.1 | | protein_coding | Protein of unknown funct | 83 | 93 | 85 | 59 | 78 | 59 | 174 | 228 | 218 | 128 | 207 | 160 | 176.32 | 207 | 165 | 0.693 | -0.528 | 0.00707574 |
| AT3G12587.1 | OST4A | protein_coding | Oligosaccaryltransferase | 126 | 112 | 108 | 96 | 75 | 100 | 265 | 275 | 276 | 208 | 199 | 272 | 262.76 | 272 | 226 | 0.693 | -0.528 | 0.00086772 |
| AT5G43970.1 | TOM9-2 | protein_coding | translocase of outer mem | 442 | 417 | 272 | 316 | 228 | 277 | 929 | 1024 | 696 | 685 | 605 | 753 | 881.37 | 883 | 681 | 0.693 | -0.528 | 0.00010597 |
| AT2G36840.1 | ACR10 | protein_coding | ACT-like superfamily prot | 85 | 86 | 85 | 50 | 57 | 54 | 179 | 211 | 218 | 108 | 151 | 147 | 149.81 | 203 | 135 | 0.693 | -0.529 | 0.00234105 |
| AT1G30330.2 | ARF6 | protein_coding | auxin response factor 6 | 664 | 770 | 787 | 476 | 495 | 606 | 1396 | 1892 | 2015 | 1032 | 1314 | 1648 | 1888.5 | 1768 | 1331 | 0.693 | -0.529 | 0.0022091 |
| AT5G49610.1 | | protein_coding | F-box family protein | 118 | 79 | 84 | 76 | 68 | 46 | 248 | 194 | 215 | 165 | 181 | 125 | 167.32 | 219 | 157 | 0.693 | -0.529 | 0.00058015 |
| AT1G69450.1 | | protein_coding | Early-responsive to dehyc | 213 | 142 | 156 | 155 | 114 | 101 | 448 | 349 | 399 | 336 | 303 | 275 | 343.19 | 399 | 305 | 0.693 | -0.529 | 0.00015808 |
| AT1G30755.1 | | protein_coding | Protein of unknown funct | 69 | 63 | 52 | 62 | 29 | 49 | 145 | 155 | 133 | 134 | 77 | 133 | 126.94 | 144 | 115 | 0.693 | -0.53 | 0.00252926 |
| AT1G73150.1 | GTE3 | protein_coding | global transcription facto | 156 | 134 | 110 | 95 | 83 | 81 | 328 | 329 | 282 | 206 | 220 | 220 | 197.86 | 313 | 215 | 0.693 | -0.53 | 0.00080122 |
| AT5G65640.1 | BHLH93 | protein_coding | beta HLH protein 93 | 194 | 105 | 124 | 172 | 112 | 111 | 408 | 258 | 317 | 373 | 297 | 302 | 327.71 | 328 | 324 | 0.693 | -0.53 | 0.00038813 |
| AT4G36180.1 | | protein_coding | Leucine-rich receptor-like | 650 | 575 | 474 | 450 | 500 | 327 | 1366 | 1413 | 1213 | 976 | 1327 | 889 | 1162.75 | 1331 | 1064 | 0.693 | -0.53 | 0.00037872 |
| AT5G05960.1 | | protein_coding | Bifunctional inhibitor/lipi | 55 | 42 | 35 | 74 | 70 | 41 | 116 | 103 | 90 | 161 | 186 | 111 | 156.21 | 103 | 153 | 0.692 | -0.531 | 0.00764921 |
| AT3G28370.1 | | protein_coding | | 195 | 171 | 204 | 95 | 114 | 87 | 410 | 420 | 522 | 206 | 303 | 237 | 315.75 | 451 | 249 | 0.692 | -0.531 | 0.00037823 |
| AT1G47310.1 | | protein_coding | | 287 | 243 | 300 | 200 | 150 | 201 | 603 | 597 | 768 | 434 | 398 | 547 | 491.69 | 656 | 460 | 0.692 | -0.531 | 0.00016186 |
| AT1G43620.1 | UGT80B1 | protein_coding | UDP-Glycosyltransferase | 458 | 488 | 423 | 320 | 305 | 257 | 963 | 1199 | 1083 | 694 | 810 | 699 | 990.49 | 1082 | 734 | 0.692 | -0.531 | 3.18E-05 |
| AT2G45880.1 | BAM7 | protein_coding | beta-amylase 7 | 214 | 208 | 206 | 173 | 144 | 156 | 450 | 511 | 527 | 375 | 382 | 424 | 376.87 | 496 | 394 | 0.692 | -0.531 | 7.59E-06 |
| AT1G19690.1 | | protein_coding | NAD(P)-binding Rossman | 149 | 127 | 140 | 111 | 94 | 94 | 313 | 312 | 358 | 241 | 250 | 256 | 316 | 328 | 249 | 0.692 | -0.532 | 2.63E-05 |
| AT1G67180.1 | | protein_coding | zinc finger (C3HC4-type R | 108 | 70 | 65 | 63 | 54 | 40 | 227 | 172 | 166 | 137 | 143 | 109 | 158.2 | 188 | 130 | 0.691 | -0.534 | 0.00644476 |
| AT3G27220.1 | | protein_coding | Galactose oxidase/kelch r | 42 | 50 | 62 | 48 | 31 | 42 | 88 | 123 | 159 | 104 | 82 | 114 | 114.64 | 123 | 100 | 0.691 | -0.534 | 0.00532694 |
| AT3G48980.1 | | protein_coding | Arabidopsis thaliana prot | 53 | 70 | 53 | 52 | 49 | 57 | 111 | 172 | 136 | 113 | 130 | 155 | 154.76 | 140 | 133 | 0.691 | -0.534 | 0.00367561 |
| AT1G49880.1 | ERV1 | protein_coding | Erv1/Alr family protein | 76 | 58 | 55 | 42 | 39 | 55 | 160 | 142 | 141 | 91 | 104 | 150 | 142.41 | 148 | 115 | 0.691 | -0.534 | 0.00361065 |
| AT1G34418.1 | | other_rna | other RNA | 97 | 121 | 105 | 83 | 68 | 80 | 204 | 297 | 269 | 180 | 181 | 218 | 319.87 | 257 | 193 | 0.691 | -0.534 | 0.00062395 |
| AT4G09250.1 | | protein_coding | SPla/RYanodine receptor | 129 | 121 | 99 | 107 | 97 | 96 | 271 | 297 | 253 | 232 | 258 | 261 | 241.58 | 274 | 250 | 0.69 | -0.534 | 5.01E-05 |
| AT3G44100.1 | | protein_coding | MD-2-related lipid recogr | 357 | 299 | 247 | 257 | 235 | 203 | 750 | 735 | 632 | 557 | 624 | 552 | 746.16 | 706 | 578 | 0.69 | -0.535 | 2.76E-06 |
| AT4G12432.1 | CPuORF26 | protein_coding | conserved peptide upstre | 36 | 43 | 49 | 32 | 24 | 28 | 76 | 106 | 125 | 69 | 64 | 76 | 81.46 | 102 | 70 | 0.69 | -0.536 | 0.00995036 |
| AT1G60430.1 | ARPC3 | protein_coding | actin-related protein C3 | 131 | 117 | 157 | 103 | 114 | 121 | 275 | 287 | 402 | 223 | 303 | 329 | 289.11 | 321 | 285 | 0.69 | -0.536 | 0.00182065 |
| AT1G54650.2 | | protein_coding | Methyltransferase family | 99 | 76 | 77 | 71 | 57 | 59 | 208 | 187 | 197 | 154 | 151 | 160 | 180.02 | 197 | 155 | 0.69 | -0.536 | 0.00110762 |
| AT2G32010.1 | CVL1 | protein_coding | CVP2 like 1 | 364 | 467 | 482 | 367 | 351 | 327 | 765 | 1147 | 1234 | 796 | 932 | 889 | 820.85 | 1049 | 872 | 0.69 | -0.536 | 0.00054546 |
| AT5G35740.1 | | protein_coding | Carbohydrate-binding X8 | 145 | 128 | 103 | 75 | 67 | 92 | 305 | 314 | 264 | 163 | 178 | 250 | 320.95 | 294 | 197 | 0.69 | -0.536 | 0.00020129 |
| AT3G13550.1 | COP10 | protein_coding | Ubiquitin-conjugating en: | 191 | 122 | 144 | 105 | 83 | 84 | 401 | 300 | 369 | 228 | 220 | 228 | 305.66 | 357 | 225 | 0.689 | -0.536 | 8.49E-05 |
| AT4G08950.1 | EXO | protein_coding | Phosphate-responsive 1 f | 588 | 348 | 387 | 576 | 288 | 509 | 1236 | 855 | 991 | 1249 | 765 | 1384 | 1144.74 | 1027 | 1133 | 0.689 | -0.537 | 0.00529751 |
| AT1G80460.1 | GLPK | protein_coding | Actin-like ATPase superfa | 750 | 786 | 728 | 534 | 428 | 690 | 1577 | 1931 | 1864 | 1158 | 1136 | 1876 | 1466.86 | 1791 | 1390 | 0.689 | -0.537 | 0.00107772 |
| AT3G29270.1 | | protein_coding | RING/U-box superfamily | | 181 | 186 | 218 | 158 | 164 | 171 | 380 | 457 | 558 | 343 | 435 | 465 | 402.89 | 465 | 414 | 0.689 | -0.538 | 0.0003839 |
| AT1G05570.1 | CALS1 | protein_coding | callose synthase 1 | 549 | 515 | 456 | 377 | 288 | 232 | 1154 | 1265 | 1167 | 818 | 765 | 631 | 753.29 | 1195 | 738 | 0.689 | -0.538 | 0.00011424 |
| AT5G40500.1 | | protein_coding | | 383 | 378 | 376 | 277 | 237 | 231 | 805 | 929 | 963 | 601 | 629 | 628 | 899.38 | 899 | 619 | 0.689 | -0.538 | 8.85E-06 |
| AT4G21210.1 | RP1 | protein_coding | PPDK regulatory protein | 2289 | 2129 | 1628 | 1583 | 1282 | 1212 | 4812 | 5231 | 4168 | 3434 | 3403 | 3296 | 5089.33 | 4737 | 3378 | 0.689 | -0.538 | 4.26E-06 |
| AT5G44550.1 | | protein_coding | Uncharacterised protein f | 58 | 71 | 58 | 71 | 59 | 53 | 122 | 174 | 148 | 154 | 157 | 144 | 222.5 | 148 | 152 | 0.688 | -0.54 | 0.00071712 |
| AT3G17020.1 | | protein_coding | Adenine nucleotide alpha | 690 | 709 | 839 | 542 | 492 | 637 | 1450 | 1742 | 2148 | 1176 | 1306 | 1732 | 2029.59 | 1780 | 1405 | 0.688 | -0.54 | 0.00043352 |
| AT3G20270.1 | | protein_coding | lipid-binding serum glyco | 265 | 213 | 197 | 175 | 122 | 96 | 557 | 523 | 504 | 380 | 324 | 261 | 385.17 | 528 | 322 | 0.688 | -0.54 | 3.33E-05 |
| AT4G19660.1 | NPR4 | protein_coding | NPR1-like protein 4 | 255 | 208 | 215 | 226 | 179 | 144 | 536 | 511 | 550 | 490 | 475 | 392 | 418.49 | 532 | 452 | 0.688 | -0.54 | 1.59E-05 |
| AT4G09200.1 | | protein_coding | SPla/RYanodine receptor | 169 | 136 | 156 | 140 | 127 | 107 | 355 | 334 | 399 | 304 | 337 | 291 | 370.38 | 363 | 311 | 0.688 | -0.54 | 1.33E-05 |
| AT4G04040.1 | PFP-BETA2 | protein_coding | Phosphofructokinase fam | 1937 | 1438 | 1476 | 1315 | 929 | 939 | 4072 | 3533 | 3779 | 2852 | 2466 | 2554 | 3189.03 | 3795 | 2624 | 0.688 | -0.54 | 3.39E-06 |
| AT3G19930.1 | STP4 | protein_coding | sugar transporter 4 | 747 | 482 | 705 | 488 | 306 | 509 | 1570 | 1184 | 1805 | 1059 | 812 | 1384 | 1138.34 | 1520 | 1085 | 0.687 | -0.541 | 0.00168739 |
| AT1G55980.1 | | protein_coding | FAD/NAD(P)-binding oxid | 141 | 138 | 121 | 92 | 69 | 62 | 296 | 339 | 310 | 200 | 183 | 169 | 220.41 | 315 | 184 | 0.687 | -0.542 | 0.00011404 |
| AT4G32270.1 | | protein_coding | Ubiquitin-like superfamily | 132 | 128 | 135 | 88 | 90 | 95 | 277 | 314 | 346 | 191 | 239 | 258 | 306.66 | 312 | 229 | 0.687 | -0.542 | 4.06E-05 |
| AT2G44980.2 | | protein_coding | SNF2 domain-containing | 188 | 129 | 147 | 102 | 63 | 83 | 395 | 317 | 376 | 221 | 167 | 226 | 237.89 | 363 | 205 | 0.686 | -0.543 | 0.00070127 |
| AT1G47720.1 | OSB1 | protein_coding | Primosome PriB/single-st | 79 | 53 | 66 | 50 | 43 | 35 | 166 | 130 | 169 | 108 | 114 | 95 | 130.81 | 155 | 106 | 0.686 | -0.543 | 0.00045599 |
| AT1G56720.1 | | protein_coding | Protein kinase superfamil | 329 | 408 | 385 | 288 | 257 | 273 | 692 | 1002 | 986 | 625 | 682 | 742 | 813.84 | 893 | 683 | 0.686 | -0.543 | 9.55E-05 |
| AT2G39890.1 | PROT1 | protein_coding | proline transporter 1 | 271 | 274 | 260 | 240 | 242 | 160 | 570 | 673 | 666 | 521 | 642 | 435 | 633.52 | 636 | 533 | 0.686 | -0.543 | 4.33E-05 |
| AT4G26520.1 | | protein_coding | Aldolase superfamily prot | 336 | 502 | 297 | 229 | 228 | 299 | 706 | 1233 | 760 | 497 | 605 | 813 | 1212.46 | 900 | 638 | 0.686 | -0.544 | 0.00344371 |
| AT2G20725.1 | | protein_coding | CAAX amino terminal pro | 479 | 446 | 402 | 306 | 268 | 254 | 1007 | 1096 | 1029 | 664 | 711 | 691 | 1056.92 | 1044 | 689 | 0.686 | -0.544 | 1.07E-06 |
| AT4G04925.1 | | protein_coding | | 32 | 36 | 21 | 24 | 26 | 43 | 67 | 88 | 54 | 52 | 69 | 117 | 168.18 | 70 | 79 | 0.685 | -0.545 | 0.00372885 |
| AT1G71697.1 | CK1 | protein_coding | choline kinase 1 | 116 | 121 | 118 | 104 | 105 | 100 | 244 | 297 | 302 | 226 | 279 | 272 | 232.35 | 281 | 259 | 0.685 | -0.545 | 0.00039215 |
| AT3G46870.1 | | protein_coding | Pentatricopeptide repeat | 116 | 92 | 71 | 116 | 94 | 60 | 244 | 226 | 182 | 252 | 250 | 163 | 249.93 | 217 | 222 | 0.686 | -0.545 | 0.00037681 |
| AT3G43210.1 | NACK2 | protein_coding | ATP binding microtubule | 187 | 174 | 189 | 138 | 154 | 117 | 393 | 427 | 484 | 299 | 409 | 318 | 302.24 | 435 | 342 | 0.685 | -0.545 | 0.00034289 |

| | | | | | | | | | | | | | | | | | | | | | |
|---|---|---|---|---|---|---|---|---|---|---|---|---|---|---|---|---|---|---|---|---|---|
| AT4G04370.1 | PCMP-E99 | protein_coding | Tetratricopeptide repeat | 200 | 158 | 178 | 109 | 97 | 80 | 420 | 388 | 456 | 236 | 258 | 218 | 273.55 | 421 | 237 | 0.686 | -0.545 | 4.71E-05 |
| AT5G65210.2 | TGA1 | protein_coding | bZIP transcription factor f | 149 | 111 | 86 | 137 | 111 | 72 | 313 | 273 | 220 | 297 | 295 | 196 | 289.41 | 269 | 263 | 0.684 | -0.547 | 0.00281036 |
| AT2G34170.1 | | protein_coding | Protein of unknown funct | 340 | 480 | 473 | 317 | 348 | 313 | 715 | 1179 | 1211 | 688 | 924 | 851 | 833.94 | 1035 | 821 | 0.684 | -0.547 | 0.00192901 |
| AT2G45850.1 | AHL9 | protein_coding | AT hook motif DNA-bindi | 341 | 403 | 372 | 286 | 266 | 323 | 717 | 990 | 952 | 620 | 706 | 878 | 820.64 | 886 | 735 | 0.684 | -0.547 | 0.00012292 |
| AT1G56700.1 | | protein_coding | Peptidase C15, pyroglutai | 495 | 426 | 500 | 401 | 336 | 263 | 1041 | 1047 | 1280 | 870 | 892 | 715 | 1059.81 | 1123 | 826 | 0.684 | -0.547 | 2.54E-05 |
| AT3G26050.1 | | protein_coding | TPX2 (targeting protein fo | 95 | 88 | 90 | 73 | 99 | 49 | 200 | 216 | 230 | 158 | 263 | 133 | 161.1 | 215 | 185 | 0.684 | -0.548 | 0.00899979 |
| AT5G55050.1 | | protein_coding | GDSL-like Lipase/Acylhydi | 96 | 58 | 49 | 108 | 66 | 45 | 202 | 142 | 125 | 234 | 175 | 122 | 188.53 | 156 | 177 | 0.684 | -0.548 | 0.00506375 |
| AT3G26490.1 | | protein_coding | Phototropic-responsive N | 197 | 251 | 151 | 124 | 91 | 101 | 414 | 617 | 387 | 269 | 242 | 275 | 399.36 | 473 | 262 | 0.684 | -0.548 | 0.00067333 |
| AT2G19400.1 | | protein_coding | AGC (cAMP-dependent, c | 116 | 118 | 89 | 69 | 74 | 54 | 244 | 290 | 228 | 150 | 196 | 147 | 255.24 | 254 | 164 | 0.684 | -0.548 | 0.00043644 |
| AT1G68220.1 | | protein_coding | Protein of unknown funct | 189 | 175 | 156 | 88 | 109 | 115 | 397 | 430 | 399 | 191 | 289 | 313 | 391.01 | 409 | 264 | 0.684 | -0.548 | 0.00036829 |
| AT2G42220.1 | STR9 | protein_coding | Rhodanese/Cell cycle con | 7361 | 7797 | 6579 | 4789 | 4479 | 4477 | 15473 | 19156 | 16843 | 10388 | 11891 | 12175 | 17268.06 | 17157 | 11485 | 0.684 | -0.548 | 4.78E-05 |
| AT1G54050.1 | HSP17.4B | protein_coding | HSP20-like chaperones su | 135 | 91 | 72 | 88 | 67 | 43 | 284 | 224 | 184 | 191 | 178 | 117 | 208.45 | 231 | 162 | 0.684 | -0.549 | 0.00174665 |
| AT5G09270.1 | | protein_coding | | 101 | 68 | 76 | 65 | 68 | 50 | 212 | 167 | 195 | 141 | 181 | 136 | 186.13 | 191 | 153 | 0.684 | -0.549 | 0.00081639 |
| AT5G01230.1 | | protein_coding | S-adenosyl-L-methionine- | 190 | 139 | 144 | 149 | 115 | 86 | 399 | 341 | 369 | 323 | 305 | 234 | 317.58 | 370 | 287 | 0.683 | -0.549 | 3.47E-05 |
| AT5G40240.2 | | protein_coding | nodulin MtN21 /EamA-lik | 108 | 79 | 79 | 73 | 48 | 41 | 227 | 194 | 202 | 158 | 127 | 111 | 205.14 | 208 | 132 | 0.683 | -0.55 | 0.00181428 |
| AT3G15900.1 | | protein_coding | | 269 | 287 | 268 | 192 | 201 | 185 | 565 | 705 | 686 | 416 | 534 | 503 | 612.15 | 652 | 484 | 0.683 | -0.55 | 2.15E-05 |
| AT5G23410.1 | | other_rna | | 74 | 58 | 88 | 41 | 50 | 58 | 156 | 142 | 225 | 89 | 133 | 158 | 159.63 | 174 | 127 | 0.683 | -0.551 | 0.00723988 |
| AT3G59570.1 | | protein_coding | Ypt/Rab-GAP domain of g | 96 | 75 | 82 | 55 | 60 | 48 | 202 | 184 | 210 | 119 | 159 | 131 | 147.74 | 199 | 136 | 0.683 | -0.551 | 0.00397149 |
| AT4G02405.1 | | protein_coding | S-adenosyl-L-methionine- | 203 | 144 | 141 | 129 | 83 | 103 | 427 | 354 | 361 | 280 | 220 | 280 | 361.06 | 381 | 260 | 0.683 | -0.551 | 2.00E-05 |
| AT1G11740.1 | | protein_coding | ankyrin repeat family pro | 103 | 132 | 131 | 74 | 41 | 76 | 217 | 324 | 335 | 161 | 109 | 207 | 434.17 | 292 | 159 | 0.682 | -0.552 | 0.00244641 |
| AT3G61980.1 | | protein_coding | serine protease inhibitor, | 43 | 50 | 59 | 41 | 44 | 44 | 90 | 123 | 151 | 89 | 117 | 120 | 132.52 | 121 | 109 | 0.682 | -0.552 | 0.00190518 |
| AT1G68590.1 | | protein_coding | Ribosomal protein PSRP-3 | 4916 | 3878 | 3613 | 3112 | 2647 | 2383 | 10334 | 9528 | 9250 | 6750 | 7027 | 6480 | 9844.41 | 9704 | 6752 | 0.682 | -0.552 | 2.62E-06 |
| AT3G51000.1 | | protein_coding | alpha/beta-Hydrolases su | 175 | 134 | 145 | 86 | 54 | 79 | 368 | 329 | 371 | 187 | 143 | 215 | 233.26 | 356 | 182 | 0.682 | -0.553 | 0.00079228 |
| AT5G64600.1 | | protein_coding | O-fucosyltransferase fam | 174 | 134 | 149 | 108 | 104 | 96 | 366 | 329 | 381 | 234 | 276 | 261 | 283.62 | 359 | 257 | 0.682 | -0.553 | 3.24E-05 |
| AT4G02550.3 | | protein_coding | | 135 | 105 | 114 | 76 | 69 | 73 | 284 | 258 | 292 | 165 | 183 | 199 | 235.99 | 278 | 182 | 0.682 | -0.553 | 2.71E-05 |
| AT4G29940.1 | PRH | protein_coding | pathogenesis related hon | 241 | 154 | 196 | 121 | 109 | 127 | 507 | 378 | 502 | 262 | 289 | 345 | 350.44 | 462 | 299 | 0.681 | -0.555 | 6.68E-05 |
| AT3G43590.1 | | protein_coding | zinc knuckle (CCHC-type) | 699 | 515 | 483 | 374 | 298 | 307 | 1469 | 1265 | 1237 | 811 | 791 | 835 | 961.68 | 1324 | 812 | 0.681 | -0.555 | 4.93E-06 |
| AT2G05185.1 | | protein_coding | | 46 | 37 | 53 | 32 | 41 | 37 | 97 | 91 | 136 | 69 | 109 | 101 | 105.12 | 108 | 93 | 0.68 | -0.556 | 0.00543434 |
| AT5G62720.1 | | protein_coding | Integral membrane HPP f | 781 | 951 | 801 | 480 | 474 | 713 | 1642 | 2336 | 2051 | 1041 | 1258 | 1939 | 3629.68 | 2010 | 1413 | 0.68 | -0.556 | 0.0017036 |
| AT1G67800.2 | | protein_coding | Copine (Calcium-depende | 97 | 90 | 74 | 78 | 59 | 50 | 204 | 221 | 189 | 169 | 157 | 136 | 189.77 | 205 | 154 | 0.68 | -0.556 | 0.00027201 |
| AT2G02100.1 | PDF2.2 | protein_coding | low-molecular-weight cys | 852 | 1036 | 910 | 780 | 690 | 965 | 1791 | 2545 | 2330 | 1692 | 1832 | 2624 | 3138.46 | 2222 | 2049 | 0.679 | -0.557 | 0.00090821 |
| AT2G17525.1 | | protein_coding | Pentatricopeptide repeat | 145 | 110 | 114 | 91 | 94 | 76 | 305 | 270 | 292 | 197 | 250 | 207 | 226.48 | 289 | 218 | 0.68 | -0.557 | 0.00033562 |
| AT2G34585.1 | | protein_coding | | 248 | 225 | 210 | 215 | 154 | 192 | 521 | 553 | 538 | 466 | 409 | 522 | 635.48 | 537 | 466 | 0.68 | -0.557 | 1.19E-06 |
| AT5G67290.1 | | protein_coding | FAD-dependent oxidored | 204 | 188 | 174 | 152 | 106 | 122 | 429 | 462 | 445 | 330 | 281 | 332 | 447.04 | 445 | 314 | 0.68 | -0.557 | 4.08E-07 |
| AT4G12070.1 | | protein_coding | | 132 | 136 | 114 | 79 | 62 | 79 | 277 | 334 | 292 | 171 | 165 | 215 | 218.45 | 301 | 184 | 0.679 | -0.558 | 0.00134672 |
| AT5G06180.1 | | protein_coding | Protein of unknown funct | 84 | 93 | 76 | 57 | 37 | 48 | 177 | 228 | 195 | 124 | 98 | 131 | 165.68 | 200 | 118 | 0.679 | -0.558 | 0.00067574 |
| AT1G73790.1 | GIP2 | protein_coding | Protein of unknown funct | 62 | 51 | 53 | 51 | 27 | 37 | 130 | 125 | 136 | 111 | 72 | 101 | 123.18 | 130 | 95 | 0.679 | -0.558 | 0.00055308 |
| AT5G10700.1 | | protein_coding | Peptidyl-tRNA hydrolase | 69 | 78 | 67 | 53 | 48 | 52 | 145 | 192 | 172 | 115 | 127 | 141 | 194.24 | 170 | 128 | 0.679 | -0.558 | 0.00034358 |
| AT5G07250.1 | ATRBL3 | protein_coding | RHOMBOID-like protein 3 | 160 | 174 | 169 | 118 | 96 | 116 | 336 | 427 | 433 | 256 | 255 | 315 | 364.95 | 399 | 275 | 0.679 | -0.558 | 5.86E-05 |
| AT3G02580.1 | STE1 | protein_coding | sterol 1 | 510 | 370 | 354 | 371 | 249 | 242 | 1072 | 909 | 906 | 805 | 661 | 658 | 938.75 | 962 | 708 | 0.679 | -0.559 | 3.06E-06 |
| AT2G28900.1 | OEP161 | protein_coding | outer plastid envelope pr | 1103 | 1310 | 1202 | 726 | 628 | 1012 | 2319 | 3218 | 3077 | 1575 | 1667 | 2752 | 3876.2 | 2871 | 1998 | 0.678 | -0.56 | 0.00223025 |
| AT5G61300.1 | | protein_coding | | 193 | 122 | 129 | 144 | 90 | 90 | 406 | 300 | 330 | 312 | 239 | 245 | 265.27 | 345 | 265 | 0.678 | -0.56 | 0.00075705 |
| AT1G78600.2 | BBX22 | protein_coding | light-regulated zinc finger | 1065 | 1303 | 1116 | 721 | 742 | 704 | 2239 | 3201 | 2857 | 1564 | 1970 | 1914 | 2709.68 | 2766 | 1816 | 0.679 | -0.56 | 0.00014368 |
| AT3G09900.1 | RABE1E | protein_coding | RAB GTPase homolog E1E | 331 | 317 | 357 | 242 | 249 | 208 | 696 | 779 | 914 | 525 | 661 | 566 | 633.9 | 796 | 584 | 0.678 | -0.56 | 2.01E-05 |
| AT2G16780.1 | MSI2 | protein_coding | Transducin family protein | 274 | 281 | 220 | 257 | 221 | 238 | 576 | 690 | 563 | 557 | 587 | 647 | 680.5 | 610 | 597 | 0.678 | -0.561 | 1.13E-06 |
| AT2G02760.1 | UBC2 | protein_coding | ubiquiting-conjugating er | 826 | 819 | 903 | 619 | 525 | 556 | 1736 | 2012 | 2312 | 1343 | 1394 | 1512 | 2258.12 | 2020 | 1416 | 0.677 | -0.562 | 1.54E-05 |
| AT4G11850.1 | PLDGAMMA1 | protein_coding | phospholipase D gamma | 318 | 228 | 261 | 295 | 189 | 186 | 668 | 560 | 668 | 640 | 502 | 506 | 546.27 | 632 | 549 | 0.677 | -0.562 | 5.16E-06 |
| AT5G44230.1 | PCMP-H17 | protein_coding | Pentatricopeptide repeat | 88 | 54 | 59 | 78 | 56 | 39 | 185 | 133 | 151 | 169 | 149 | 106 | 123.3 | 156 | 141 | 0.677 | -0.563 | 0.00768722 |
| AT1G53920.1 | GLIP5 | protein_coding | GDSL-motif lipase 5 | 107 | 59 | 80 | 65 | 43 | 33 | 225 | 145 | 205 | 141 | 114 | 90 | 149.82 | 192 | 115 | 0.677 | -0.563 | 0.00257748 |
| AT1G73570.1 | HRD3B | protein_coding | HCP-like superfamily prot | 70 | 80 | 48 | 50 | 41 | 53 | 147 | 197 | 123 | 108 | 109 | 144 | 166.52 | 156 | 120 | 0.677 | -0.563 | 0.00057438 |
| AT1G15140.1 | | protein_coding | FAD/NAD(P)-binding oxid | 560 | 530 | 307 | 428 | 316 | 326 | 1177 | 1302 | 786 | 928 | 839 | 887 | 2848.1 | 1088 | 885 | 0.677 | -0.563 | 0.00014872 |
| AT5G59950.5 | ALY1 | protein_coding | RNA-binding (RRM/RBD/ | 461 | 458 | 376 | 366 | 287 | 254 | 969 | 1125 | 963 | 794 | 762 | 691 | 1237.25 | 1019 | 749 | 0.677 | -0.563 | 3.94E-06 |
| AT3G26540.1 | PCMP-A5 | protein_coding | Tetratricopeptide repeat | 80 | 59 | 45 | 50 | 50 | 37 | 168 | 145 | 115 | 108 | 133 | 101 | 148.82 | 143 | 114 | 0.676 | -0.564 | 0.0009293 |
| AT1G10660.2 | | protein_coding | | 381 | 239 | 334 | 235 | 168 | 195 | 801 | 587 | 855 | 510 | 446 | 530 | 614.61 | 748 | 495 | 0.676 | -0.565 | 4.74E-05 |
| AT3G02180.1 | SP1L3 | protein_coding | SPIRAL1-like3 | 776 | 721 | 693 | 626 | 522 | 579 | 1631 | 1771 | 1774 | 1358 | 1386 | 1575 | 1797.71 | 1725 | 1440 | 0.676 | -0.565 | 6.17E-07 |
| AT3G45420.1 | LECRK14 | protein_coding | Concanavalin A-like lectin | 40 | 42 | 45 | 41 | 32 | 32 | 84 | 103 | 115 | 89 | 85 | 87 | 89.81 | 101 | 87 | 0.675 | -0.566 | 0.00308485 |
| AT4G34215.1 | | protein_coding | Domain of unknown func | 189 | 220 | 171 | 144 | 128 | 185 | 397 | 540 | 438 | 312 | 340 | 503 | 508.51 | 458 | 385 | 0.675 | -0.566 | 0.00020099 |
| AT2G37450.2 | | protein_coding | nodulin MtN21 /EamA-lik | 587 | 543 | 456 | 303 | 300 | 240 | 1234 | 1334 | 1167 | 657 | 796 | 653 | 1031.75 | 1245 | 702 | 0.676 | -0.566 | 1.54E-05 |
| AT1G24440.1 | | protein_coding | RING/U-box superfamily | 158 | 156 | 214 | 109 | 115 | 135 | 332 | 383 | 548 | 236 | 305 | 367 | 309.48 | 421 | 303 | 0.675 | -0.567 | 0.00210051 |
| AT5G59000.1 | | protein_coding | RING/FYVE/PHD zinc fing | 82 | 92 | 80 | 46 | 68 | 61 | 172 | 226 | 205 | 100 | 181 | 166 | 198.83 | 201 | 149 | 0.675 | -0.567 | 0.00089729 |
| AT1G16916.1 | | protein_coding | | 91 | 70 | 61 | 61 | 54 | 61 | 191 | 172 | 156 | 132 | 143 | 166 | 168.8 | 173 | 147 | 0.675 | -0.567 | 0.00030178 |
| AT2G26240.1 | FAX7 | protein_coding | Transmembrane proteins | 135 | 116 | 97 | 129 | 92 | 138 | 284 | 285 | 248 | 280 | 244 | 375 | 416.21 | 272 | 300 | 0.675 | -0.567 | 1.00E-04 |
| AT4G32120.1 | B3GALT10 | protein_coding | Galactosyltransferase fan | 214 | 192 | 205 | 161 | 147 | 143 | 450 | 472 | 525 | 349 | 390 | 389 | 452.94 | 482 | 376 | 0.675 | -0.567 | 5.58E-07 |
| AT5G17580.1 | | protein_coding | Phototropic-responsive N | 103 | 92 | 70 | 52 | 44 | 39 | 217 | 226 | 179 | 113 | 117 | 106 | 161.24 | 207 | 112 | 0.674 | -0.569 | 0.00059979 |
| AT1G11960.1 | | protein_coding | ERD (early-responsive to | 187 | 164 | 196 | 86 | 109 | 111 | 393 | 403 | 502 | 187 | 289 | 302 | 294.84 | 433 | 259 | 0.674 | -0.569 | 0.00028709 |
| AT3G46550.1 | FLA4 | protein_coding | Fasciclin-like arabinogala | 124 | 132 | 79 | 87 | 86 | 62 | 261 | 324 | 202 | 189 | 228 | 169 | 688.75 | 262 | 195 | 0.674 | -0.569 | 0.00023807 |
| AT3G09470.1 | | protein_coding | Major facilitator superfar | 294 | 267 | 277 | 219 | 174 | 180 | 618 | 656 | 709 | 475 | 462 | 489 | 606.04 | 661 | 475 | 0.674 | -0.57 | 1.59E-07 |
| AT5G57880.1 | MPS1 | protein_coding | multipolar spindle 1 | 138 | 98 | 168 | 83 | 63 | 90 | 290 | 241 | 430 | 180 | 167 | 245 | 237.71 | 320 | 197 | 0.673 | -0.571 | 0.00502877 |
| AT1G70518.1 | | other_rna | | 30 | 31 | 34 | 39 | 24 | 27 | 63 | 76 | 87 | 85 | 64 | 73 | 82.96 | 75 | 74 | 0.673 | -0.571 | 0.00350919 |
| AT3G21510.1 | AHP1 | protein_coding | histidine-containing phos | 64 | 40 | 58 | 54 | 36 | 29 | 135 | 98 | 148 | 117 | 96 | 79 | 127.7 | 127 | 97 | 0.673 | -0.572 | 0.00198707 |
| AT1G26665.2 | MED10B | protein_coding | Mediator complex, subur | 146 | 132 | 200 | 100 | 74 | 103 | 307 | 324 | 512 | 217 | 196 | 280 | 318.01 | 381 | 231 | 0.673 | -0.572 | 0.0014959 |
| AT3G53270.1 | | protein_coding | Small nuclear RNA activa | 149 | 151 | 175 | 134 | 115 | 93 | 313 | 371 | 448 | 291 | 305 | 253 | 343.72 | 377 | 283 | 0.673 | -0.572 | 5.22E-05 |
| AT5G66070.2 | ATL27 | protein_coding | RING/U-box superfamily | 52 | 39 | 43 | 30 | 26 | 30 | 109 | 96 | 110 | 65 | 69 | 82 | 129.5 | 105 | 72 | 0.672 | -0.573 | 0.00020436 |
| AT4G14320.2 | RPL36AB | protein_coding | Zinc-binding ribosomal pr | 1949 | 1641 | 1141 | 2004 | 1505 | 1646 | 4097 | 4032 | 2921 | 4347 | 3995 | 4476 | 5538.07 | 3683 | 4273 | 0.672 | -0.573 | 0.00010308 |
| AT3G51390.1 | PAT10 | protein_coding | DHHC-type zinc finger far | 204 | 193 | 228 | 162 | 135 | 139 | 429 | 474 | 584 | 351 | 358 | 378 | 482.14 | 496 | 362 | 0.672 | -0.573 | 9.38E-06 |
| AT3G03890.1 | | protein_coding | FMN binding | 506 | 435 | 424 | 366 | 249 | 228 | 1064 | 1069 | 1085 | 794 | 661 | 620 | 1127.9 | 1073 | 692 | 0.672 | -0.573 | 1.44E-06 |
| AT4G23430.2 | TIC32 | protein_coding | NAD(P)-binding Rossman | 491 | 424 | 382 | 366 | 277 | 283 | 1032 | 1042 | 978 | 794 | 735 | 770 | 976.52 | 1017 | 766 | 0.672 | -0.573 | 3.08E-07 |
| AT1G56120.1 | | protein_coding | Leucine-rich repeat transı | 54 | 39 | 31 | 40 | 22 | 26 | 114 | 96 | 79 | 87 | 58 | 71 | 89.83 | 96 | 72 | 0.672 | -0.574 | 0.00301348 |
| AT4G01410.1 | | protein_coding | Late embryogenesis abun | 166 | 102 | 104 | 85 | 65 | 92 | 349 | 251 | 266 | 184 | 173 | 250 | 276.95 | 289 | 202 | 0.672 | -0.574 | 0.00095696 |
| AT2G47160.2 | BOR1 | protein_coding | HCO3- transporter family | 509 | 312 | 385 | 312 | 269 | 318 | 1070 | 767 | 986 | 677 | 714 | 865 | 866.6 | 941 | 752 | 0.672 | -0.574 | 3.67E-05 |
| AT3G05165.1 | | protein_coding | Major facilitator superfar | 602 | 394 | 308 | 375 | 250 | 148 | 1265 | 968 | 788 | 813 | 664 | 402 | 1191.65 | 1007 | 626 | 0.671 | -0.575 | 0.00287027 |
| AT4G17040.1 | CLPR4 | protein_coding | CLP protease R subunit 4 | 3208 | 2590 | 2699 | 2172 | 1780 | 1755 | 6743 | 6363 | 6910 | 4711 | 4725 | 4773 | 6045.54 | 6672 | 4736 | 0.671 | -0.575 | 1.20E-07 |
| AT2G37380.1 | MAKR3 | protein_coding | | 70 | 82 | 68 | 53 | 52 | 36 | 147 | 201 | 174 | 115 | 138 | 98 | 168.57 | 174 | 117 | 0.671 | -0.576 | 0.00141883 |
| AT4G23920.1 | UGE2 | protein_coding | UDP-D-glucose/UDP-D-ga | 118 | 168 | 123 | 99 | 95 | 112 | 248 | 413 | 315 | 215 | 252 | 305 | 334.89 | 325 | 257 | 0.671 | -0.576 | 0.00025932 |
| AT1G63830.2 | | protein_coding | PLAC8 family protein | 287 | 293 | 245 | 199 | 219 | 222 | 603 | 720 | 627 | 432 | 581 | 604 | 898.27 | 650 | 539 | 0.671 | -0.576 | 1.24E-05 |
| AT1G11180.2 | SCAMP5 | protein_coding | Secretory carrier membra | 184 | 153 | 172 | 112 | 110 | 92 | 387 | 376 | 440 | 243 | 292 | 250 | 364.93 | 401 | 262 | 0.671 | -0.576 | 1.09E-05 |
| AT1G70430.1 | | protein_coding | Protein kinase superfamil | 83 | 69 | 65 | 38 | 38 | 32 | 174 | 170 | 166 | 82 | 101 | 87 | 107.35 | 170 | 90 | 0.67 | -0.577 | 0.00658857 |
| AT4G14960.2 | TUBA6 | protein_coding | Tubulin/FtsZ family prote | 7257 | 8082 | 6350 | 8451 | 8138 | 7765 | 15255 | 19856 | 16256 | 18331 | 21605 | 21116 | 28332.06 | 17122 | 20351 | 0.67 | -0.577 | 0.00010999 |
| AT5G11070.1 | | protein_coding | | 252 | 385 | 319 | 261 | 234 | 337 | 530 | 946 | 817 | 566 | 621 | 916 | 816.4 | 764 | 701 | 0.67 | -0.578 | 0.00787471 |
| AT3G14130.1 | GLO4 | protein_coding | Aldolase-type TIM barrel | 93 | 72 | 120 | 85 | 83 | 80 | 195 | 177 | 307 | 184 | 220 | 218 | 198.49 | 226 | 207 | 0.67 | -0.578 | 0.00192831 |
| AT3G28950.1 | | protein_coding | AIG2-like (avirulence indu | 95 | 107 | 116 | 67 | 59 | 62 | 200 | 263 | 297 | 145 | 157 | 169 | 208.75 | 253 | 157 | 0.67 | -0.578 | 0.00116198 |
| AT2G40020.3 | | protein_coding | Nucleolar histone methyl | 113 | 142 | 97 | 70 | 88 | 51 | 238 | 349 | 248 | 152 | 234 | 139 | 295.25 | 278 | 175 | 0.67 | -0.578 | 0.00099655 |
| AT1G68790.1 | CRWN3 | protein_coding | little nuclei3 | 759 | 656 | 680 | 457 | 381 | 320 | 1595 | 1612 | 1741 | 991 | 1011 | 870 | 1229.37 | 1649 | 957 | 0.67 | -0.578 | 6.48E-07 |
| AT3G17070.1 | PER29 | protein_coding | Peroxidase family protein | 83 | 108 | 79 | 89 | 101 | 123 | 174 | 265 | 202 | 193 | 268 | 334 | 319.07 | 214 | 265 | 0.67 | -0.579 | 0.00230339 |
| AT1G26810.1 | B3GALT15 | protein_coding | galactosyltransferase1 | 124 | 108 | 99 | 57 | 61 | 48 | 261 | 265 | 253 | 124 | 162 | 131 | 182.67 | 260 | 139 | 0.669 | -0.58 | 0.0002345 |
| AT3G06170.1 | | protein_coding | Serinc-domain containing | 301 | 332 | 317 | 226 | 208 | 290 | 633 | 816 | 812 | 490 | 552 | 789 | 710.07 | 754 | 610 | 0.669 | -0.58 | 0.0001387 |
| AT5G64500.1 | | protein_coding | Major facilitator superfar | 596 | 583 | 608 | 427 | 368 | 357 | 1253 | 1432 | 1557 | 926 | 977 | 971 | 1237.12 | 1414 | 958 | 0.669 | -0.58 | 4.36E-06 |
| AT3G15110.1 | | protein_coding | | 545 | 530 | 429 | 353 | 289 | 335 | 1146 | 1302 | 1098 | 766 | 767 | 911 | 1427.02 | 1182 | 815 | 0.669 | -0.58 | 2.23E-06 |
| AT2G30910.1 | ARPC1A | protein_coding | actin-related protein C1A | 227 | 165 | 179 | 164 | 132 | 120 | 477 | 405 | 458 | 356 | 350 | 326 | 370.93 | 447 | 344 | 0.669 | -0.58 | 4.26E-07 |
| AT5G24930.1 | ATCOL4 | protein_coding | CONSTANS-like 4 | 442 | 662 | 521 | 321 | 288 | 495 | 929 | 1626 | 1334 | 696 | 765 | 1346 | 3588.77 | 1296 | 936 | 0.669 | -0.581 | 0.00445162 |
| AT4G28652.1 | | other_rna | | 74 | 62 | 91 | 43 | 43 | 60 | 156 | 152 | 233 | 93 | 114 | 163 | 143.07 | 180 | 123 | 0.669 | -0.581 | 0.00315972 |
| AT3G01690.1 | | protein_coding | alpha/beta-Hydrolases su | 729 | 1029 | 832 | 547 | 569 | 806 | 1532 | 2528 | 2130 | 1186 | 1511 | 2192 | 2523.26 | 2063 | 1630 | 0.668 | -0.582 | 0.00215756 |
| AT4G15680.1 | GRXS4 | protein_coding | Thioredoxin superfamily | 95 | 72 | 64 | 118 | 84 | 122 | 200 | 177 | 164 | 256 | 223 | 332 | 184.66 | 180 | 270 | 0.668 | -0.582 | 0.0017484 |
| AT3G48100.1 | ARR5 | protein_coding | response regulator 5 | 189 | 229 | 194 | 122 | 160 | 133 | 397 | 563 | 497 | 265 | 425 | 362 | 409.36 | 486 | 351 | 0.668 | -0.582 | 0.00036534 |
| AT1G15250.1 | RPL37A | protein_coding | Zinc-binding ribosomal pr | 239 | 173 | 114 | 129 | 107 | 114 | 502 | 425 | 292 | 280 | 284 | 310 | 444.39 | 406 | 291 | 0.668 | -0.582 | 0.0001468 |
| AT5G18550.1 | | protein_coding | Zinc finger C-x8-C-x5-C-x3 | 149 | 160 | 127 | 109 | 123 | 131 | 313 | 393 | 325 | 236 | 327 | 356 | 498.32 | 344 | 306 | 0.668 | -0.582 | 5.82E-06 |
| AT1G52710.1 | | protein_coding | Rubredoxin-like superfam | 36 | 55 | 52 | 30 | 23 | 25 | 76 | 135 | 133 | 65 | 61 | 68 | 92.73 | 115 | 65 | 0.667 | -0.583 | 0.00617784 |
| AT1G75960.1 | AAE8 | protein_coding | AMP-dependent syntheta | 187 | 214 | 124 | 138 | 74 | 99 | 393 | 526 | 317 | 299 | 196 | 269 | 484.26 | 412 | 255 | 0.667 | -0.583 | 0.00029433 |
| AT4G14010.1 | RALFL32 | protein_coding | ralf-like 32 | 50 | 39 | 56 | 93 | 65 | 65 | 105 | 96 | 143 | 202 | 173 | 177 | 218.42 | 115 | 184 | 0.668 | -0.583 | 0.00022724 |
| AT2G26510.1 | NAT3 | protein_coding | Xanthine/uracil permease | 445 | 414 | 414 | 265 | 260 | 241 | 935 | 1017 | 1060 | 575 | 690 | 655 | 902.39 | 1004 | 640 | 0.668 | -0.583 | 3.12E-07 |
| AT4G12130.1 | | protein_coding | Glycine cleavage T-protei | 91 | 68 | 54 | 70 | 27 | 53 | 191 | 167 | 138 | 152 | 72 | 144 | 159.75 | 165 | 123 | 0.667 | -0.584 | 0.00189554 |
| AT1G58470.1 | RBP1 | protein_coding | RNA-binding protein 1 | 79 | 60 | 61 | 39 | 39 | 26 | 166 | 147 | 156 | 85 | 104 | 71 | 106.42 | 156 | 87 | 0.667 | -0.584 | 0.00159592 |
| AT1G08920.2 | SUGTL2 | protein_coding | ERD (early response to de | 164 | 209 | 174 | 130 | 139 | 90 | 345 | 513 | 445 | 282 | 369 | 245 | 334.54 | 434 | 299 | 0.667 | -0.584 | 0.00045141 |
| AT1G59580.1 | MPK2 | protein_coding | mitogen-activated protei | 288 | 242 | 315 | 214 | 162 | 204 | 605 | 595 | 806 | 464 | 430 | 555 | 517.26 | 669 | 483 | 0.667 | -0.584 | 1.76E-05 |
| AT3G28100.1 | | protein_coding | nodulin MtN21 /EamA-lik | 70 | 43 | 26 | 45 | 41 | 27 | 147 | 106 | 67 | 98 | 109 | 73 | 110.7 | 107 | 93 | 0.667 | -0.585 | 0.00932017 |
| AT5G08100.1 | | protein_coding | N-terminal nucleophile ar | 165 | 172 | 160 | 125 | 93 | 126 | 347 | 423 | 410 | 271 | 247 | 343 | 526.83 | 393 | 287 | 0.666 | -0.585 | 2.27E-06 |
| AT1G22830.1 | PCMP-E24 | protein_coding | Tetratricopeptide repeat | 55 | 45 | 54 | 48 | 50 | 31 | 116 | 111 | 138 | 104 | 133 | 84 | 108.51 | 122 | 107 | 0.666 | -0.586 | 0.00180577 |
| AT5G24170.1 | | protein_coding | Got1/Sft2-like vescicle tra | 95 | 96 | 77 | 80 | 52 | 54 | 200 | 236 | 197 | 174 | 138 | 147 | 192.84 | 211 | 153 | 0.666 | -0.586 | 0.0002451 |
| AT1G03600.1 | PSB27-1 | protein_coding | photosystem II family pro | 3617 | 4747 | 3888 | 2826 | 2699 | 3732 | 7603 | 11663 | 9954 | 6130 | 7165 | 10149 | 13627.42 | 9740 | 7815 | 0.666 | -0.587 | 0.00119428 |
| AT4G30140.1 | CDEF1 | protein_coding | GDSL-like Lipase/Acylhydi | 139 | 119 | 97 | 113 | 76 | 72 | 292 | 292 | 248 | 245 | 202 | 196 | 280.18 | 277 | 214 | 0.666 | -0.587 | 0.000181 |
| AT4G21585.1 | ENDO4 | protein_coding | endonuclease 4 | 80 | 62 | 86 | 57 | 57 | 38 | 168 | 152 | 220 | 124 | 151 | 103 | 135.52 | 180 | 126 | 0.665 | -0.588 | 0.00087708 |
| AT2G45890.1 | ROPGEF4 | protein_coding | RHO guanyl-nucleotide e | 20 | 24 | 24 | 27 | 19 | 22 | 42 | 59 | 61 | 59 | 50 | 60 | 54.35 | 54 | 56 | 0.665 | -0.589 | 0.00682502 |
| AT1G03440.1 | | protein_coding | Leucine-rich repeat (LRR) | 220 | 355 | 246 | 224 | 209 | 247 | 462 | 872 | 630 | 486 | 555 | 672 | 702.53 | 655 | 571 | 0.665 | -0.589 | 0.00098497 |
| AT2G21090.1 | PCMP-E48 | protein_coding | Pentatricopeptide repeat | 151 | 104 | 91 | 83 | 94 | 101 | 317 | 256 | 233 | 180 | 250 | 275 | 240.24 | 269 | 235 | 0.665 | -0.589 | 0.00054617 |
| AT5G45780.1 | | protein_coding | Leucine-rich repeat prote | 153 | 110 | 100 | 133 | 93 | 72 | 322 | 270 | 256 | 288 | 247 | 196 | 273.31 | 283 | 244 | 0.665 | -0.589 | 4.20E-05 |
| AT3G52150.1 | | protein_coding | RNA-binding (RRM/RBD/ | 4811 | 4177 | 3885 | 3181 | 2705 | 2540 | 10113 | 10262 | 9946 | 6900 | 7181 | 6907 | 9552.89 | 10107 | 6996 | 0.664 | -0.59 | 1.82E-07 |
| AT5G28830.1 | | protein_coding | calcium-binding EF hand | 116 | 84 | 105 | 90 | 48 | 59 | 244 | 206 | 269 | 195 | 127 | 160 | 161.66 | 240 | 161 | 0.664 | -0.591 | 0.0005099 |
| AT4G25270.1 | PCMP-E53 | protein_coding | Tetratricopeptide repeat | 124 | 105 | 120 | 43 | 39 | 58 | 261 | 258 | 307 | 93 | 104 | 158 | 185.47 | 275 | 118 | 0.664 | -0.591 | 0.0003092 |
| AT1G22350.1 | | pseudogene | | 38 | 24 | 11 | 17 | 17 | 9 | 80 | 59 | 28 | 37 | 45 | 24 | 84.82 | 56 | 35 | 0.664 | -0.592 | 0.00889403 |
| AT2G21880.1 | RABG2 | protein_coding | RAB GTPase homolog 7A | 95 | 90 | 69 | 60 | 55 | 51 | 200 | 221 | 177 | 130 | 146 | 139 | 194.87 | 199 | 138 | 0.664 | -0.592 | 5.26E-05 |
| AT2G46505.1 | SDH4 | protein_coding | succinate dehydrogenase | 649 | 491 | 428 | 562 | 394 | 354 | 1364 | 1206 | 1096 | 1219 | 1046 | 963 | 1304.9 | 1222 | 1076 | 0.663 | -0.592 | 7.10E-06 |
| AT1G08900.1 | SUGTL3 | protein_coding | Major facilitator superfar | 181 | 186 | 217 | 84 | 95 | 111 | 380 | 457 | 556 | 182 | 252 | 302 | 368.86 | 464 | 245 | 0.662 | -0.594 | 0.00041482 |
| AT1G17870.1 | EGY3 | protein_coding | ethylene-dependent grav | 90 | 101 | 71 | 73 | 71 | 50 | 189 | 248 | 182 | 158 | 188 | 136 | 199.76 | 206 | 161 | 0.663 | -0.594 | 0.00027411 |
| AT1G66900.1 | | protein_coding | alpha/beta-Hydrolases su | 240 | 255 | 221 | 171 | 177 | 155 | 504 | 626 | 566 | 371 | 470 | 422 | 477.99 | 565 | 421 | 0.663 | -0.594 | 7.51E-06 |
| AT3G11090.1 | LBD21 | protein_coding | LOB domain-containing p | 211 | 212 | 164 | 182 | 150 | 118 | 444 | 521 | 420 | 395 | 398 | 321 | 502.07 | 462 | 371 | 0.662 | -0.595 | 8.88E-05 |
| AT3G47630.2 | | protein_coding | | 135 | 118 | 110 | 91 | 68 | 68 | 284 | 290 | 282 | 197 | 181 | 185 | 244.39 | 285 | 188 | 0.662 | -0.595 | 2.39E-06 |
| AT4G14110.1 | CSN8 | protein_coding | COP9 signalosome, subur | 294 | 233 | 220 | 324 | 293 | 280 | 618 | 572 | 563 | 703 | 778 | 761 | 809.31 | 584 | 747 | 0.662 | -0.595 | 4.16E-08 |
| AT1G64890.1 | | protein_coding | Major facilitator superfar | 94 | 110 | 99 | 41 | 44 | 77 | 198 | 270 | 253 | 89 | 117 | 209 | 205.91 | 240 | 138 | 0.661 | -0.596 | 0.00465048 |
| AT1G12440.1 | SAP1 | protein_coding | A20/AN1-like zinc finger | 728 | 741 | 841 | 563 | 467 | 557 | 1530 | 1821 | 2153 | 1221 | 1240 | 1515 | 1716.36 | 1835 | 1325 | 0.662 | -0.596 | 9.74E-06 |
| AT5G53900.2 | | protein_coding | Serine/threonine-protein | 233 | 229 | 200 | 166 | 145 | 158 | 490 | 563 | 512 | 360 | 385 | 430 | 472.01 | 522 | 392 | 0.662 | -0.596 | 1.87E-06 |
| AT5G27410.1 | | protein_coding | D-aminoacid aminotransf | 246 | 177 | 149 | 132 | 116 | 102 | 517 | 435 | 381 | 286 | 308 | 277 | 383 | 444 | 290 | 0.661 | -0.597 | 4.06E-06 |
| AT3G46780.1 | PTAC16 | protein_coding | plastid transcriptionally a | 10166 | 15514 | 10467 | 5982 | 6358 | 8377 | 21370 | 38115 | 26796 | 12976 | 16879 | 22781 | 33331.91 | 28760 | 17545 | 0.661 | -0.598 | 0.00541145 |
| AT3G14430.1 | | protein_coding | | 57 | 88 | 85 | 84 | 79 | 84 | 120 | 216 | 218 | 182 | 210 | 228 | 191.7 | 185 | 207 | 0.661 | -0.598 | 0.00189971 |
| AT5G45480.1 | | protein_coding | Protein of unknown funct | 806 | 617 | 479 | 520 | 331 | 289 | 1694 | 1516 | 1226 | 1128 | 879 | 786 | 920.39 | 1479 | 931 | 0.661 | -0.598 | 5.69E-05 |
| AT1G48320.1 | DHNAT1 | protein_coding | Thioesterase superfamily | 67 | 33 | 40 | 51 | 24 | 29 | 141 | 81 | 102 | 111 | 64 | 79 | 113 | 108 | 85 | 0.66 | -0.599 | 0.00527395 |

| | | | | | | | | | | | | | | | | | | | | | |
|---|---|---|---|---|---|---|---|---|---|---|---|---|---|---|---|---|---|---|---|---|---|
| AT1G14170.3 | | protein_coding | RNA-binding KH domain- | 149 | 148 | 202 | 132 | 116 | 135 | 313 | 364 | 517 | 286 | 308 | 367 | 315.74 | 398 | 320 | 0.66 | -0.599 | 0.00042677 |
| AT1G04110.1 | SBT1.2 | protein_coding | Subtilase family protein | 222 | 220 | 263 | 200 | 150 | 190 | 467 | 540 | 673 | 434 | 398 | 517 | 424.67 | 560 | 450 | 0.66 | -0.599 | 0.00014709 |
| AT1G48360.2 | | protein_coding | zinc ion binding;nucleic a | 223 | 189 | 208 | 150 | 102 | 107 | 469 | 464 | 532 | 325 | 271 | 291 | 352.11 | 488 | 296 | 0.66 | -0.599 | 4.36E-06 |
| AT1G30845.1 | | protein_coding | | 40 | 28 | 35 | 32 | 32 | 27 | 84 | 69 | 90 | 69 | 85 | 73 | 78.18 | 81 | 76 | 0.66 | -0.6 | 0.00825603 |
| AT2G25930.1 | ELF3 | protein_coding | hydroxyproline-rich glyco | 778 | 654 | 800 | 417 | 369 | 436 | 1635 | 1607 | 2048 | 905 | 980 | 1186 | 1397.94 | 1763 | 1024 | 0.66 | -0.6 | 1.30E-05 |
| AT2G24350.1 | | protein_coding | RNA binding (RRM/RBD/ | 80 | 61 | 64 | 55 | 49 | 32 | 168 | 150 | 164 | 119 | 130 | 87 | 122.23 | 161 | 112 | 0.66 | -0.601 | 0.00210143 |
| AT2G02770.1 | | protein_coding | 4'-phosphopantetheinyl t | 110 | 74 | 96 | 107 | 64 | 82 | 231 | 182 | 246 | 232 | 170 | 223 | 229.34 | 220 | 208 | 0.659 | -0.601 | 0.00023385 |
| AT4G10100.1 | CNX7 | protein_coding | co-factor for nitrate, redu | 163 | 131 | 112 | 106 | 94 | 110 | 343 | 322 | 287 | 230 | 250 | 299 | 321.05 | 317 | 260 | 0.659 | -0.601 | 1.00E-05 |
| AT3G47640.1 | BHLH47 | protein_coding | basic helix-loop-helix (bH | 69 | 89 | 101 | 56 | 51 | 60 | 145 | 219 | 259 | 121 | 135 | 163 | 152.58 | 208 | 140 | 0.659 | -0.602 | 0.00709175 |
| AT5G65420.3 | CYCD4-1 | protein_coding | CYCLIN D4;1 | 99 | 71 | 61 | 48 | 37 | 42 | 208 | 174 | 156 | 104 | 98 | 114 | 130.85 | 179 | 105 | 0.659 | -0.602 | 0.00156101 |
| AT4G18530.1 | | protein_coding | Protein of unknown funct | 144 | 144 | 165 | 98 | 106 | 79 | 303 | 354 | 422 | 213 | 281 | 215 | 304.11 | 360 | 236 | 0.659 | -0.602 | 4.55E-05 |
| AT2G21140.1 | PRP2 | protein_coding | proline-rich protein 2 | 469 | 566 | 412 | 375 | 349 | 352 | 986 | 1391 | 1055 | 813 | 927 | 957 | 1717.81 | 1144 | 899 | 0.659 | -0.602 | 2.20E-05 |
| AT4G12790.1 | | protein_coding | P-loop containing nucleo | 565 | 439 | 463 | 703 | 660 | 628 | 1188 | 1079 | 1185 | 1525 | 1752 | 1708 | 1791.21 | 1151 | 1662 | 0.659 | -0.602 | 8.37E-07 |
| AT5G66960.1 | | protein_coding | Prolyl oligopeptidase fam | 61 | 42 | 38 | 44 | 27 | 20 | 128 | 103 | 97 | 95 | 72 | 54 | 86.26 | 109 | 74 | 0.659 | -0.603 | 0.00823573 |
| AT5G63800.1 | BGAL6 | protein_coding | Glycosyl hydrolase family | 749 | 1149 | 802 | 766 | 801 | 623 | 1574 | 2823 | 2053 | 1662 | 2126 | 1694 | 1907.14 | 2150 | 1827 | 0.659 | -0.603 | 0.00052288 |
| AT5G66360.2 | | protein_coding | Ribosomal RNA adenine d | 100 | 113 | 109 | 103 | 79 | 79 | 210 | 278 | 279 | 223 | 210 | 215 | 243.07 | 256 | 216 | 0.658 | -0.603 | 1.62E-05 |
| AT5G67060.1 | HEC1 | protein_coding | basic helix-loop-helix (bH | 14 | 10 | 11 | 13 | 9 | 7 | 29 | 25 | 28 | 28 | 24 | 19 | 88.14 | 27 | 24 | 0.658 | -0.604 | 0.00062321 |
| AT3G60180.2 | UMK1 | protein_coding | P-loop containing nucleo | 72 | 54 | 56 | 42 | 24 | 29 | 151 | 133 | 143 | 91 | 64 | 79 | 112.28 | 142 | 78 | 0.658 | -0.604 | 0.00032552 |
| AT2G23755.1 | | protein_coding | | 82 | 83 | 73 | 64 | 56 | 31 | 172 | 204 | 187 | 139 | 149 | 84 | 172.03 | 188 | 124 | 0.658 | -0.605 | 0.00063876 |
| AT4G13195.1 | CLE44 | protein_coding | CLAVATA3/ESR-RELATED | 38 | 36 | 19 | 57 | 39 | 37 | 80 | 88 | 49 | 124 | 104 | 101 | 140.76 | 72 | 110 | 0.658 | -0.605 | 0.00037165 |
| AT3G07200.1 | | protein_coding | RING/U-box superfamily | 158 | 141 | 103 | 102 | 80 | 79 | 332 | 346 | 264 | 221 | 212 | 215 | 283.62 | 314 | 216 | 0.657 | -0.605 | 1.12E-05 |
| AT4G03410.2 | | protein_coding | Peroxisomal membrane 2 | 264 | 220 | 266 | 211 | 146 | 162 | 555 | 540 | 681 | 458 | 388 | 441 | 530.53 | 592 | 429 | 0.658 | -0.605 | 1.92E-06 |
| AT1G56140.1 | | protein_coding | Leucine-rich repeat trans | 277 | 213 | 204 | 154 | 117 | 122 | 582 | 523 | 522 | 334 | 311 | 332 | 385.27 | 542 | 326 | 0.657 | -0.605 | 1.56E-06 |
| AT3G22060.1 | CRRSP38 | protein_coding | Receptor-like protein kin | 161 | 119 | 125 | 134 | 51 | 69 | 338 | 292 | 320 | 291 | 135 | 188 | 245.92 | 317 | 205 | 0.657 | -0.606 | 0.00127445 |
| AT3G19960.2 | VIII-1 | protein_coding | myosin 1 | 1128 | 1036 | 997 | 763 | 713 | 624 | 2371 | 2545 | 2552 | 1655 | 1893 | 1697 | 2040.68 | 2489 | 1748 | 0.657 | -0.606 | 9.91E-08 |
| AT1G15885.1 | | protein_coding | | 36 | 34 | 42 | 34 | 23 | 26 | 76 | 84 | 108 | 74 | 61 | 71 | 79.76 | 89 | 69 | 0.656 | -0.608 | 0.0077786 |
| AT1G50270.1 | PCMP-E42 | protein_coding | Pentatricopeptide repeat | 80 | 81 | 61 | 51 | 45 | 45 | 168 | 199 | 156 | 111 | 119 | 122 | 128.8 | 174 | 117 | 0.656 | -0.608 | 0.00036243 |
| AT3G21740.1 | APO4 | protein_coding | Arabidopsis thaliana prot | 76 | 72 | 83 | 71 | 53 | 54 | 160 | 177 | 212 | 154 | 141 | 147 | 173.9 | 183 | 147 | 0.656 | -0.608 | 4.30E-05 |
| AT4G13330.1 | | protein_coding | S-adenosyl-L-methionine- | 50 | 27 | 33 | 54 | 28 | 22 | 105 | 66 | 84 | 117 | 74 | 60 | 105.65 | 85 | 84 | 0.656 | -0.609 | 0.0027408 |
| AT4G31910.1 | | protein_coding | HXXXD-type acyl-transfer | 68 | 81 | 58 | 68 | 74 | 47 | 143 | 199 | 148 | 147 | 196 | 128 | 154.45 | 163 | 157 | 0.656 | -0.609 | 0.00148443 |
| AT1G69210.1 | | protein_coding | Uncharacterised protein f | 74 | 53 | 39 | 67 | 41 | 41 | 156 | 130 | 100 | 145 | 109 | 111 | 141.47 | 129 | 122 | 0.656 | -0.609 | 0.00081123 |
| AT4G26550.1 | | protein_coding | Got1/Sft2-like vescicle tra | 103 | 110 | 91 | 87 | 66 | 74 | 217 | 270 | 233 | 189 | 175 | 201 | 226.5 | 240 | 188 | 0.656 | -0.609 | 1.80E-05 |
| AT4G05330.1 | AGD13 | protein_coding | ARF-GAP domain 13 | 290 | 224 | 259 | 148 | 131 | 104 | 610 | 550 | 663 | 321 | 348 | 283 | 439.36 | 608 | 317 | 0.656 | -0.609 | 3.69E-06 |
| AT2G06010.1 | ORG4 | protein_coding | OBP3-responsive gene 4 | 179 | 136 | 167 | 126 | 97 | 98 | 376 | 334 | 428 | 273 | 258 | 267 | 327.53 | 379 | 266 | 0.656 | -0.609 | 1.24E-06 |
| AT2G16790.1 | | protein_coding | P-loop containing nucleo | 45 | 43 | 34 | 17 | 26 | 21 | 95 | 106 | 87 | 37 | 69 | 57 | 69.51 | 96 | 54 | 0.655 | -0.61 | 0.00499365 |
| AT1G56260.1 | | protein_coding | | 99 | 60 | 56 | 71 | 50 | 44 | 208 | 147 | 143 | 154 | 133 | 120 | 135.94 | 166 | 136 | 0.655 | -0.61 | 0.00060818 |
| AT5G04360.1 | PU1 | protein_coding | limit dextrinase | 900 | 1092 | 961 | 562 | 568 | 556 | 1892 | 2683 | 2460 | 1219 | 1508 | 1512 | 1828.81 | 2345 | 1413 | 0.655 | -0.61 | 1.57E-05 |
| AT2G44190.1 | EDE1 | protein_coding | Family of unknown functi | 64 | 57 | 53 | 45 | 50 | 43 | 135 | 140 | 136 | 98 | 133 | 117 | 113.75 | 137 | 116 | 0.655 | -0.611 | 0.00167509 |
| AT4G15560.1 | DXS | protein_coding | Deoxyxylulose-5-phospha | 10238 | 10742 | 9169 | 11637 | 11143 | 10108 | 21521 | 26391 | 23473 | 25242 | 29582 | 27488 | 28231.08 | 23795 | 27437 | 0.655 | -0.611 | 1.96E-05 |
| AT4G19120.1 | ERD3 | protein_coding | S-adenosyl-L-methionine- | 1271 | 1226 | 1212 | 926 | 842 | 937 | 2672 | 3012 | 3103 | 2009 | 2235 | 2548 | 2473.42 | 2929 | 2264 | 0.655 | -0.611 | 1.27E-05 |
| AT5G49290.1 | ATRLP56 | protein_coding | receptor like protein 56 | 50 | 46 | 39 | 29 | 23 | 24 | 105 | 113 | 100 | 63 | 61 | 65 | 67.89 | 106 | 63 | 0.654 | -0.612 | 0.00278722 |
| AT1G02410.1 | COX11 | protein_coding | cytochrome c oxidase ass | 287 | 247 | 304 | 205 | 181 | 150 | 603 | 607 | 778 | 445 | 481 | 408 | 544.03 | 663 | 445 | 0.654 | -0.612 | 5.49E-07 |
| AT4G01935.1 | | protein_coding | | 220 | 222 | 214 | 139 | 125 | 92 | 462 | 545 | 548 | 302 | 332 | 250 | 446.48 | 518 | 295 | 0.654 | -0.613 | 2.03E-06 |
| AT1G28050.1 | COL15 | protein_coding | B-box type zinc finger pro | 93 | 105 | 116 | 52 | 47 | 82 | 195 | 258 | 297 | 113 | 125 | 223 | 282.72 | 250 | 154 | 0.653 | -0.614 | 0.00130232 |
| AT5G65400.1 | | protein_coding | alpha/beta-Hydrolases su | 115 | 90 | 92 | 64 | 50 | 53 | 242 | 221 | 236 | 139 | 133 | 144 | 205.16 | 233 | 139 | 0.653 | -0.614 | 4.27E-05 |
| AT5G23575.1 | | protein_coding | Transmembrane CLPTM1 | 1186 | 1170 | 1360 | 624 | 643 | 595 | 2493 | 2874 | 3482 | 1354 | 1707 | 1618 | 2464.39 | 2950 | 1560 | 0.653 | -0.614 | 2.22E-05 |
| AT5G66790.1 | WAKL21 | protein_coding | Protein kinase superfamil | 27 | 41 | 30 | 26 | 21 | 27 | 57 | 101 | 77 | 56 | 56 | 73 | 68.23 | 78 | 62 | 0.653 | -0.615 | 0.00494196 |
| AT1G79460.1 | GA2 | protein_coding | Terpenoid cyclases/Prote | 151 | 162 | 200 | 93 | 93 | 63 | 317 | 398 | 512 | 202 | 247 | 171 | 330.24 | 409 | 207 | 0.653 | -0.615 | 0.00032012 |
| AT1G26660.2 | | protein_coding | Prefoldin chaperone subu | 75 | 52 | 52 | 53 | 31 | 41 | 158 | 128 | 133 | 115 | 82 | 111 | 143.27 | 140 | 103 | 0.653 | -0.615 | 0.00012623 |
| AT4G33640.1 | | protein_coding | | 269 | 362 | 301 | 253 | 200 | 289 | 565 | 889 | 771 | 549 | 531 | 786 | 886.99 | 742 | 622 | 0.652 | -0.616 | 0.00020189 |
| AT1G48175.1 | emb2191 | protein_coding | Cytidine/deoxycytidylate | 124 | 102 | 103 | 85 | 64 | 78 | 261 | 251 | 264 | 184 | 170 | 212 | 234.38 | 259 | 189 | 0.653 | -0.616 | 2.08E-05 |
| AT3G22070.1 | | protein_coding | proline-rich family protei | 106 | 94 | 92 | 65 | 39 | 19 | 223 | 231 | 236 | 141 | 104 | 52 | 132.91 | 230 | 99 | 0.652 | -0.618 | 0.00797068 |
| AT1G14360.1 | UTR3 | protein_coding | UDP-galactose transporte | 491 | 245 | 219 | 289 | 175 | 132 | 1032 | 602 | 561 | 627 | 465 | 359 | 575.65 | 732 | 484 | 0.651 | -0.618 | 0.00273055 |
| AT5G54930.1 | | protein_coding | AT hook motif-containing | 75 | 95 | 76 | 54 | 44 | 43 | 158 | 233 | 195 | 117 | 117 | 117 | 178.61 | 195 | 117 | 0.652 | -0.618 | 4.73E-05 |
| AT1G75190.1 | | protein_coding | | 52 | 65 | 55 | 20 | 42 | 47 | 109 | 160 | 141 | 43 | 112 | 128 | 218.23 | 137 | 94 | 0.651 | -0.619 | 0.0009481 |
| AT2G07723.1 | | pseudogene | | 2599 | 2752 | 2596 | 3009 | 2789 | 3151 | 5463 | 6761 | 6646 | 6527 | 7404 | 8569 | 3469.13 | 6290 | 7500 | 0.65 | -0.621 | 0.00230767 |
| AT1G12920.1 | ERF1-2 | protein_coding | eukaryotic release factor | 1823 | 1122 | 1073 | 856 | 646 | 582 | 3832 | 2757 | 2747 | 1857 | 1715 | 1583 | 2230.77 | 3112 | 1718 | 0.65 | -0.621 | 8.77E-06 |
| AT5G26570.1 | GWD3 | protein_coding | catalytics;carbohydrate k | 2548 | 2588 | 2189 | 1522 | 1333 | 1213 | 5356 | 6358 | 5604 | 3301 | 3539 | 3299 | 5010.08 | 5773 | 3380 | 0.65 | -0.621 | 2.42E-07 |
| AT4G29110.1 | | protein_coding | | 49 | 43 | 38 | 40 | 38 | 33 | 103 | 106 | 97 | 87 | 101 | 90 | 107.67 | 102 | 93 | 0.65 | -0.622 | 0.00316194 |
| AT2G42485.1 | | other_rna | other RNA | 90 | 80 | 76 | 60 | 44 | 45 | 189 | 197 | 195 | 130 | 117 | 122 | 176.44 | 194 | 123 | 0.65 | -0.622 | 7.19E-05 |
| AT5G47540.1 | | protein_coding | Mo25 family protein | 542 | 361 | 414 | 356 | 262 | 302 | 1139 | 887 | 1060 | 772 | 696 | 821 | 928.04 | 1029 | 763 | 0.649 | -0.623 | 2.06E-07 |
| AT4G01995.1 | | protein_coding | | 348 | 322 | 318 | 187 | 190 | 131 | 732 | 791 | 814 | 406 | 504 | 356 | 705.63 | 779 | 422 | 0.649 | -0.625 | 2.11E-07 |
| AT3G47530.1 | PCMP-H76 | protein_coding | Pentatricopeptide repeat | 80 | 51 | 49 | 60 | 43 | 42 | 168 | 125 | 125 | 130 | 114 | 114 | 126.59 | 139 | 119 | 0.648 | -0.626 | 0.00030944 |
| AT5G60460.1 | | protein_coding | Preprotein translocase Se | 48 | 36 | 40 | 47 | 21 | 40 | 101 | 88 | 102 | 102 | 56 | 109 | 106.39 | 97 | 89 | 0.647 | -0.627 | 0.00229181 |
| AT5G56270.1 | WRKY2 | protein_coding | WRKY DNA-binding prote | 128 | 117 | 128 | 68 | 63 | 64 | 269 | 287 | 328 | 147 | 167 | 174 | 214.8 | 295 | 163 | 0.647 | -0.627 | 2.01E-05 |
| AT4G29000.1 | TCX5 | protein_coding | Tesmin/TSO1-like CXC do | 210 | 164 | 167 | 161 | 131 | 128 | 441 | 403 | 428 | 349 | 348 | 348 | 298.19 | 424 | 348 | 0.647 | -0.627 | 7.32E-06 |
| AT2G01918.1 | PQL3 | protein_coding | PsbQ-like 3 | 173 | 184 | 166 | 140 | 94 | 83 | 364 | 452 | 425 | 304 | 250 | 226 | 348.14 | 414 | 260 | 0.648 | -0.627 | 5.81E-06 |
| AT2G36390.1 | SBE2.1 | protein_coding | starch branching enzyme | 1680 | 1531 | 1504 | 1252 | 1131 | 1066 | 3531 | 3761 | 3850 | 2716 | 3003 | 2899 | 3667.6 | 3714 | 2873 | 0.648 | -0.627 | 3.12E-08 |
| AT5G65890.1 | ACR1 | protein_coding | ACT domain repeat 1 | 83 | 103 | 81 | 53 | 79 | 53 | 174 | 253 | 207 | 115 | 210 | 144 | 200.61 | 211 | 156 | 0.647 | -0.628 | 0.00060509 |
| AT1G66050.1 | ORTH5 | protein_coding | Zinc finger (C3HC4-type F | 52 | 46 | 47 | 56 | 52 | 50 | 109 | 113 | 120 | 121 | 138 | 136 | 115.1 | 114 | 132 | 0.647 | -0.628 | 0.00039955 |
| AT3G25740.1 | MAP1C | protein_coding | methionine aminopeptida | 192 | 174 | 203 | 135 | 114 | 100 | 404 | 427 | 520 | 293 | 303 | 272 | 398.45 | 450 | 289 | 0.647 | -0.628 | 5.42E-07 |
| AT2G44850.1 | | protein_coding | | 50 | 33 | 65 | 44 | 36 | 46 | 105 | 81 | 166 | 95 | 96 | 125 | 103.19 | 117 | 105 | 0.646 | -0.63 | 0.00439422 |
| AT4G30700.1 | DYW9 | protein_coding | Pentatricopeptide repeat | 84 | 48 | 43 | 36 | 43 | 31 | 177 | 118 | 110 | 78 | 114 | 84 | 107.1 | 135 | 92 | 0.646 | -0.63 | 0.00332748 |
| AT1G69700.1 | HVA22C | protein_coding | HVA22 homologue C | 409 | 296 | 281 | 252 | 212 | 151 | 860 | 727 | 719 | 547 | 563 | 411 | 679.43 | 769 | 507 | 0.646 | -0.63 | 3.85E-07 |
| AT4G38280.1 | | protein_coding | | 233 | 189 | 213 | 128 | 126 | 132 | 490 | 464 | 545 | 278 | 335 | 359 | 482.88 | 500 | 324 | 0.646 | -0.63 | 7.26E-08 |
| AT2G22890.1 | FAD4L2 | protein_coding | Kua-ubiquitin conjugating | 26 | 39 | 43 | 20 | 27 | 20 | 55 | 96 | 110 | 43 | 72 | 54 | 77.82 | 87 | 56 | 0.646 | -0.631 | 0.00917238 |
| AT3G52905.1 | | protein_coding | Polynucleotidyl transfera | 104 | 69 | 75 | 65 | 49 | 36 | 219 | 170 | 192 | 141 | 130 | 98 | 172.16 | 194 | 123 | 0.646 | -0.631 | 0.00023835 |
| AT1G23360.1 | MENG | protein_coding | S-adenosyl-L-methionine- | 427 | 363 | 383 | 273 | 214 | 195 | 898 | 892 | 981 | 592 | 568 | 530 | 848.14 | 924 | 563 | 0.646 | -0.631 | 8.23E-09 |
| AT3G45600.1 | TET3 | protein_coding | tetraspanin3 | 440 | 591 | 555 | 306 | 431 | 448 | 925 | 1452 | 1421 | 664 | 1144 | 1218 | 1165.51 | 1266 | 1009 | 0.645 | -0.632 | 0.00172616 |
| AT4G23010.3 | UTR2 | protein_coding | UDP-galactose transporte | 389 | 233 | 234 | 249 | 130 | 122 | 818 | 572 | 599 | 540 | 345 | 332 | 525.11 | 663 | 406 | 0.645 | -0.632 | 0.00011625 |
| AT1G17455.1 | EFL4 | protein_coding | ELF4-like 4 | 153 | 112 | 155 | 81 | 70 | 70 | 322 | 275 | 397 | 176 | 186 | 190 | 235.49 | 331 | 184 | 0.645 | -0.633 | 0.00050003 |
| AT4G13430.1 | IIL1 | protein_coding | isopropyl malate isomera | 5562 | 4213 | 3254 | 6665 | 5680 | 4497 | 11692 | 10351 | 8330 | 14457 | 15079 | 12229 | 14927.01 | 10124 | 13922 | 0.645 | -0.633 | 1.38E-05 |
| AT1G04620.1 | HCAR | protein_coding | coenzyme F420 hydrogen | 458 | 506 | 500 | 307 | 307 | 319 | 963 | 1243 | 1280 | 666 | 815 | 868 | 950.64 | 1162 | 783 | 0.645 | -0.633 | 4.70E-06 |
| AT5G56630.1 | PFK7 | protein_coding | phosphofructokinase 7 | 573 | 460 | 454 | 416 | 336 | 267 | 1204 | 1130 | 1162 | 902 | 892 | 726 | 973.66 | 1165 | 840 | 0.645 | -0.633 | 6.87E-08 |
| AT5G25570.3 | | protein_coding | | 47 | 43 | 66 | 29 | 31 | 23 | 99 | 106 | 169 | 63 | 82 | 63 | 91.09 | 125 | 69 | 0.644 | -0.634 | 0.00606477 |
| AT5G14550.1 | | protein_coding | Core-2/I-branching beta- | 192 | 159 | 240 | 114 | 78 | 95 | 404 | 391 | 614 | 247 | 207 | 258 | 363.37 | 470 | 237 | 0.644 | -0.634 | 0.00024862 |
| AT2G30930.1 | | protein_coding | | 462 | 612 | 466 | 358 | 249 | 340 | 971 | 1504 | 1193 | 777 | 661 | 925 | 1474.74 | 1223 | 788 | 0.644 | -0.634 | 5.21E-05 |
| AT1G56230.1 | | protein_coding | Protein of unknown funct | 161 | 216 | 164 | 126 | 115 | 117 | 338 | 531 | 420 | 273 | 305 | 318 | 353.87 | 430 | 299 | 0.644 | -0.634 | 3.34E-05 |
| AT1G35430.1 | | protein_coding | | 21 | 22 | 18 | 13 | 11 | 23 | 44 | 54 | 46 | 28 | 29 | 63 | 68.85 | 48 | 40 | 0.644 | -0.635 | 0.00648705 |
| AT5G25210.1 | | protein_coding | | 112 | 105 | 147 | 74 | 76 | 103 | 235 | 258 | 376 | 161 | 202 | 280 | 215.12 | 290 | 214 | 0.643 | -0.636 | 0.00456594 |
| AT5G38850.1 | | protein_coding | Disease resistance protei | 308 | 303 | 368 | 212 | 156 | 155 | 647 | 744 | 942 | 460 | 414 | 422 | 403.54 | 778 | 432 | 0.643 | -0.636 | 0.00011566 |
| AT1G19310.1 | | protein_coding | RING/U-box superfamily | 97 | 119 | 78 | 74 | 79 | 107 | 204 | 292 | 200 | 161 | 210 | 291 | 294.23 | 232 | 221 | 0.643 | -0.637 | 0.00071904 |
| AT3G03180.1 | | protein_coding | Got1/Sft2-like vescicle tra | 41 | 47 | 64 | 38 | 31 | 44 | 86 | 115 | 164 | 82 | 82 | 120 | 111.64 | 122 | 95 | 0.643 | -0.638 | 0.00177929 |
| AT5G10800.1 | | protein_coding | RNA recognition motif (R | 116 | 150 | 116 | 51 | 69 | 68 | 244 | 369 | 297 | 111 | 183 | 185 | 222.57 | 303 | 160 | 0.642 | -0.639 | 0.00037416 |
| AT5G50330.1 | | protein_coding | Protein kinase superfamil | 85 | 94 | 101 | 71 | 41 | 66 | 179 | 231 | 259 | 154 | 109 | 179 | 165.18 | 223 | 147 | 0.642 | -0.64 | 0.00083453 |
| AT1G01470.1 | LEA14 | protein_coding | Late embryogenesis abun | 253 | 288 | 285 | 191 | 208 | 199 | 532 | 708 | 730 | 414 | 552 | 541 | 758.09 | 657 | 502 | 0.642 | -0.64 | 1.69E-06 |
| AT5G39730.1 | | protein_coding | AIG2-like (avirulence indu | 536 | 353 | 435 | 304 | 260 | 246 | 1127 | 867 | 1114 | 659 | 690 | 669 | 776.68 | 1036 | 673 | 0.642 | -0.64 | 6.25E-07 |
| AT3G43440.1 | TIFY3A | protein_coding | jasmonate-zim-domain pr | 96 | 137 | 111 | 68 | 77 | 60 | 202 | 337 | 284 | 147 | 204 | 163 | 220.61 | 274 | 171 | 0.641 | -0.641 | 0.00014547 |
| AT1G13360.1 | | protein_coding | | 427 | 669 | 361 | 244 | 273 | 280 | 898 | 1644 | 924 | 529 | 725 | 761 | 1025.27 | 1155 | 672 | 0.641 | -0.642 | 0.00079957 |
| AT2G25737.1 | | protein_coding | Sulfite exporter TauE/Saf | 107 | 92 | 107 | 88 | 78 | 71 | 225 | 226 | 274 | 191 | 207 | 193 | 245.23 | 242 | 197 | 0.641 | -0.642 | 1.14E-06 |
| AT1G49350.1 | | protein_coding | pfkB-like carbohydrate ki | 111 | 86 | 72 | 50 | 43 | 60 | 233 | 211 | 184 | 108 | 114 | 163 | 168.09 | 209 | 128 | 0.64 | -0.643 | 0.00018026 |
| AT3G44020.1 | | protein_coding | thylakoid lumenal P17.1 p | 304 | 285 | 250 | 253 | 185 | 144 | 639 | 700 | 640 | 549 | 491 | 392 | 623.42 | 660 | 477 | 0.64 | -0.643 | 7.81E-08 |
| AT4G15180.1 | ATXR3 | protein_coding | SET domain protein 2 | 1778 | 1588 | 1516 | 1876 | 1738 | 1475 | 3737 | 3901 | 3881 | 4069 | 4614 | 4011 | 3034.4 | 3840 | 4231 | 0.64 | -0.643 | 1.32E-08 |
| AT5G02080.1 | | protein_coding | DNA / pantothenate meta | 39 | 40 | 38 | 27 | 24 | 20 | 82 | 98 | 97 | 59 | 64 | 54 | 74.5 | 92 | 59 | 0.64 | -0.644 | 0.00239013 |
| AT5G59600.1 | PCMP-E1 | protein_coding | Tetratricopeptide repeat | 55 | 46 | 41 | 50 | 21 | 26 | 116 | 113 | 105 | 108 | 56 | 71 | 94.8 | 111 | 78 | 0.64 | -0.644 | 0.00146341 |
| AT3G46640.1 | LUX | protein_coding | Homeodomain-like super | 230 | 267 | 319 | 152 | 93 | 189 | 483 | 656 | 817 | 330 | 247 | 514 | 650.06 | 652 | 364 | 0.64 | -0.644 | 0.00094502 |
| AT1G06460.1 | ACD32.1 | protein_coding | alpha-crystallin domain 3 | 1583 | 1837 | 1899 | 918 | 983 | 835 | 3328 | 4513 | 4862 | 1991 | 2610 | 2271 | 5154.9 | 4234 | 2291 | 0.64 | -0.644 | 2.44E-05 |
| AT5G57790.1 | | protein_coding | | 50 | 44 | 51 | 23 | 37 | 33 | 105 | 108 | 131 | 50 | 98 | 90 | 108.8 | 115 | 79 | 0.64 | -0.645 | 0.00179452 |
| AT3G18080.1 | BGLU44 | protein_coding | B-S glucosidase 44 | 4660 | 4616 | 4549 | 2383 | 2795 | 2509 | 9796 | 11341 | 11646 | 5169 | 7420 | 6823 | 8807.55 | 10928 | 6471 | 0.639 | -0.645 | 1.63E-05 |
| AT1G61370.1 | | protein_coding | S-locus lectin protein kina | 85 | 41 | 48 | 70 | 36 | 36 | 179 | 101 | 123 | 152 | 96 | 98 | 113.07 | 134 | 115 | 0.639 | -0.646 | 0.00543958 |
| AT3G50700.1 | IDD2 | protein_coding | indeterminate(ID)-domai | 418 | 602 | 655 | 264 | 330 | 370 | 879 | 1479 | 1677 | 573 | 876 | 1006 | 979.94 | 1345 | 818 | 0.639 | -0.646 | 0.00193703 |
| AT1G62730.1 | | protein_coding | Terpenoid synthases supe | 74 | 42 | 47 | 58 | 40 | 31 | 156 | 103 | 120 | 126 | 106 | 84 | 138.28 | 126 | 105 | 0.639 | -0.646 | 0.00018191 |
| AT4G01330.2 | | protein_coding | Protein kinase superfamil | 585 | 732 | 808 | 529 | 401 | 575 | 1230 | 1798 | 2069 | 1147 | 1065 | 1564 | 1509.14 | 1699 | 1259 | 0.639 | -0.647 | 0.00016497 |
| AT5G25840.1 | | protein_coding | Protein of unknown funct | 166 | 302 | 246 | 184 | 194 | 250 | 349 | 742 | 630 | 399 | 515 | 680 | 650.34 | 574 | 531 | 0.638 | -0.648 | 0.00193073 |
| AT4G14830.1 | HSP1 | protein_coding | | 26 | 26 | 18 | 20 | 22 | 31 | 55 | 64 | 46 | 43 | 58 | 84 | 106.53 | 55 | 62 | 0.638 | -0.648 | 0.00108674 |
| AT3G03600.1 | RPS2 | protein_coding | ribosomal protein S2 | 329 | 242 | 194 | 213 | 120 | 161 | 692 | 595 | 497 | 462 | 319 | 438 | 547.86 | 595 | 406 | 0.638 | -0.648 | 4.74E-06 |
| AT5G55860.1 | | protein_coding | Plant protein of unknowr | 265 | 193 | 232 | 176 | 149 | 148 | 557 | 474 | 594 | 382 | 396 | 402 | 355.11 | 542 | 393 | 0.638 | -0.649 | 9.33E-07 |
| AT2G34860.1 | EDA3 | protein_coding | DnaJ/Hsp40 cysteine-rich | 1992 | 1910 | 1627 | 1195 | 1104 | 1088 | 4187 | 4693 | 4165 | 2592 | 2931 | 2959 | 4223.2 | 4348 | 2827 | 0.637 | -0.65 | 2.16E-08 |
| AT4G09520.1 | | protein_coding | Cofactor-independent ph | 231 | 163 | 173 | 142 | 113 | 79 | 486 | 400 | 443 | 308 | 300 | 215 | 336.18 | 443 | 274 | 0.637 | -0.651 | 2.40E-06 |
| AT3G47160.2 | | protein_coding | RING/U-box superfamily | 375 | 410 | 649 | 200 | 234 | 324 | 788 | 1007 | 1661 | 434 | 621 | 881 | 786.01 | 1152 | 645 | 0.636 | -0.652 | 0.00560148 |
| AT4G15450.1 | | protein_coding | Senescence/dehydration- | 52 | 88 | 110 | 97 | 99 | 117 | 109 | 216 | 282 | 210 | 263 | 318 | 285.51 | 202 | 264 | 0.636 | -0.652 | 0.002253 |
| AT2G01390.1 | | protein_coding | Tetratricopeptide repeat | 58 | 52 | 35 | 34 | 38 | 29 | 122 | 128 | 90 | 74 | 101 | 79 | 108.31 | 113 | 85 | 0.636 | -0.652 | 0.00017778 |
| AT4G11410.1 | | protein_coding | NAD(P)-binding Rossman | 228 | 186 | 161 | 134 | 76 | 82 | 479 | 457 | 412 | 291 | 202 | 223 | 356.87 | 449 | 239 | 0.637 | -0.652 | 3.97E-07 |
| AT4G23240.1 | CRK16 | protein_coding | cysteine-rich RLK (RECEPT | 47 | 38 | 41 | 33 | 23 | 11 | 99 | 93 | 105 | 72 | 61 | 30 | 104.47 | 99 | 54 | 0.636 | -0.653 | 0.00050945 |
| AT4G16710.1 | | protein_coding | glycosyltransferase family | 155 | 153 | 218 | 120 | 114 | 125 | 326 | 376 | 558 | 260 | 303 | 340 | 347.63 | 420 | 301 | 0.636 | -0.653 | 0.00013767 |
| AT5G30495.2 | | protein_coding | Fcf2 pre-rRNA processing | 192 | 151 | 146 | 89 | 88 | 70 | 404 | 371 | 374 | 193 | 234 | 190 | 308 | 383 | 206 | 0.635 | -0.654 | 8.16E-06 |
| AT1G59840.1 | CCB4 | protein_coding | cofactor assembly of com | 443 | 353 | 316 | 247 | 195 | 176 | 931 | 867 | 809 | 536 | 518 | 479 | 814.59 | 869 | 511 | 0.636 | -0.654 | 3.90E-08 |
| AT1G30010.1 | | protein_coding | Intron maturase, type II fa | 117 | 89 | 77 | 83 | 67 | 37 | 246 | 219 | 197 | 180 | 178 | 101 | 207.65 | 221 | 153 | 0.635 | -0.655 | 0.00013749 |
| AT4G13170.1 | RPL13AC | protein_coding | Ribosomal protein L13 fa | 837 | 646 | 464 | 833 | 656 | 635 | 1759 | 1587 | 1188 | 1807 | 1742 | 1727 | 2374.9 | 1511 | 1759 | 0.635 | -0.655 | 1.98E-05 |
| AT3G04930.1 | | protein_coding | DNA-binding storekeeper | 294 | 266 | 218 | 161 | 131 | 177 | 618 | 654 | 558 | 349 | 348 | 481 | 512.82 | 610 | 393 | 0.635 | -0.655 | 4.95E-07 |
| AT1G07740.1 | | protein_coding | Tetratricopeptide repeat | 45 | 39 | 30 | 34 | 28 | 19 | 95 | 96 | 77 | 74 | 74 | 52 | 102.35 | 89 | 67 | 0.635 | -0.656 | 0.00025741 |
| AT2G21195.2 | | protein_coding | | 131 | 116 | 114 | 100 | 90 | 92 | 275 | 285 | 292 | 217 | 239 | 250 | 321.42 | 284 | 235 | 0.635 | -0.656 | 1.45E-08 |
| AT5G59400.2 | | protein_coding | | 201 | 199 | 189 | 162 | 128 | 118 | 423 | 489 | 484 | 351 | 340 | 321 | 470.96 | 465 | 337 | 0.634 | -0.657 | 8.28E-09 |
| AT4G15700.1 | GRXS3 | protein_coding | Thioredoxin superfamily | 64 | 62 | 78 | 88 | 52 | 114 | 135 | 152 | 200 | 191 | 138 | 310 | 172.89 | 162 | 213 | 0.634 | -0.658 | 0.00855108 |
| AT2G36120.1 | DOT1 | protein_coding | Glycine-rich protein famil | 63 | 60 | 40 | 65 | 56 | 49 | 132 | 147 | 102 | 141 | 149 | 133 | 739.11 | 127 | 141 | 0.634 | -0.658 | 1.69E-06 |
| AT5G55840.1 | | protein_coding | Pentatricopeptide repeat | 235 | 186 | 167 | 152 | 110 | 97 | 494 | 457 | 428 | 330 | 292 | 264 | 287.42 | 460 | 295 | 0.634 | -0.658 | 7.81E-07 |
| AT2G31440.1 | | protein_coding | | 149 | 132 | 135 | 86 | 93 | 102 | 313 | 324 | 346 | 187 | 247 | 277 | 359.26 | 328 | 237 | 0.634 | -0.658 | 1.36E-07 |
| AT2G18170.1 | MPK7 | protein_coding | MAP kinase 7 | 212 | 241 | 284 | 127 | 137 | 160 | 446 | 592 | 727 | 275 | 364 | 435 | 418.65 | 588 | 358 | 0.633 | -0.659 | 0.00021598 |
| AT4G37380.1 | ELI1 | protein_coding | Tetratricopeptide repeat | 157 | 129 | 126 | 114 | 101 | 113 | 330 | 317 | 323 | 247 | 268 | 307 | 284.02 | 323 | 274 | 0.633 | -0.659 | 1.68E-05 |
| AT5G57815.1 | COX6B-2 | protein_coding | Cytochrome c oxidase, su | 229 | 213 | 246 | 194 | 169 | 135 | 481 | 523 | 630 | 421 | 449 | 367 | 560.51 | 545 | 412 | 0.633 | -0.659 | 4.16E-08 |
| AT1G13040.1 | | protein_coding | Pentatricopeptide repeat | 48 | 31 | 43 | 39 | 23 | 21 | 101 | 76 | 110 | 85 | 61 | 57 | 81.61 | 96 | 68 | 0.633 | -0.66 | 0.00210497 |
| AT4G14746.1 | | protein_coding | | 84 | 65 | 46 | 101 | 59 | 37 | 177 | 160 | 118 | 219 | 157 | 101 | 211.84 | 152 | 159 | 0.633 | -0.66 | 0.00101585 |
| AT1G30810.1 | JMJ18 | protein_coding | Transcription factor jumo | 678 | 935 | 837 | 593 | 258 | 223 | 1425 | 2297 | 2143 | 1286 | 685 | 606 | 954.23 | 1955 | 859 | 0.632 | -0.661 | 0.00364855 |
| AT4G12990.1 | | protein_coding | | 37 | 52 | 36 | 47 | 35 | 27 | 78 | 128 | 92 | 102 | 93 | 73 | 135.32 | 99 | 89 | 0.632 | -0.661 | 6.28E-05 |
| AT1G13740.1 | AFP2 | protein_coding | ABI five binding protein 2 | 155 | 162 | 133 | 81 | 85 | 76 | 326 | 398 | 340 | 176 | 226 | 207 | 299.18 | 355 | 203 | 0.632 | -0.661 | 1.71E-07 |

| | | | | | | | | | | | | | | | | | | | | | |
|---|---|---|---|---|---|---|---|---|---|---|---|---|---|---|---|---|---|---|---|---|---|
| AT3G50210.1 | | protein_coding | 2-oxoglutarate (2OG) and | 308 | 227 | 228 | 167 | 134 | 119 | 647 | 558 | 584 | 362 | 356 | 324 | 446.98 | 596 | 347 | 0.632 | -0.661 | 5.30E-09 |
| AT5G53090.1 | | protein_coding | NAD(P)-binding Rossman | 44 | 45 | 39 | 28 | 30 | 23 | 92 | 111 | 100 | 61 | 80 | 63 | 94.6 | 101 | 68 | 0.632 | -0.662 | 0.0018214 |
| AT1G21840.1 | UREF | protein_coding | urease accessory protein | 75 | 64 | 73 | 65 | 54 | 72 | 158 | 157 | 187 | 141 | 143 | 196 | 169.63 | 167 | 160 | 0.632 | -0.663 | 1.50E-05 |
| AT4G22770.1 | AHL2 | protein_coding | AT hook motif DNA-bindi | 20 | 26 | 16 | 14 | 20 | 19 | 42 | 64 | 41 | 30 | 53 | 52 | 61.4 | 49 | 45 | 0.631 | -0.665 | 0.00526381 |
| AT5G49150.1 | GEX2 | protein_coding | gamete expressed 2 | 58 | 51 | 46 | 51 | 47 | 31 | 122 | 125 | 118 | 111 | 125 | 84 | 97.99 | 122 | 107 | 0.631 | -0.665 | 0.00171741 |
| AT5G05090.1 | | protein_coding | Homeodomain-like super | 41 | 71 | 72 | 35 | 48 | 49 | 86 | 174 | 184 | 76 | 127 | 133 | 225.55 | 148 | 112 | 0.631 | -0.665 | 0.0011184 |
| AT2G27190.1 | PAP12 | protein_coding | purple acid phosphatase | 942 | 604 | 606 | 658 | 442 | 374 | 1980 | 1484 | 1551 | 1427 | 1173 | 1017 | 1355.92 | 1672 | 1206 | 0.631 | -0.665 | 4.99E-06 |
| AT1G72070.1 | | protein_coding | Chaperone DnaJ-domain | 47 | 24 | 28 | 34 | 20 | 33 | 99 | 59 | 72 | 74 | 53 | 90 | 63.43 | 77 | 72 | 0.63 | -0.666 | 0.00721521 |
| AT1G56570.1 | PCMP-E64 | protein_coding | Tetratricopeptide repeat | 62 | 54 | 43 | 49 | 33 | 33 | 130 | 133 | 110 | 106 | 88 | 90 | 109.65 | 124 | 95 | 0.63 | -0.666 | 0.00010252 |
| AT1G11150.1 | | transposable_elem | transposable element ger | 78 | 73 | 73 | 40 | 42 | 49 | 164 | 179 | 187 | 87 | 112 | 133 | 158.84 | 177 | 111 | 0.63 | -0.666 | 7.12E-05 |
| AT5G63780.1 | SHA1 | protein_coding | RING/FYVE/PHD zinc fing | 955 | 1071 | 905 | 697 | 646 | 467 | 2007 | 2631 | 2317 | 1512 | 1715 | 1270 | 2506.48 | 2318 | 1499 | 0.63 | -0.666 | 2.52E-06 |
| AT5G18430.1 | | protein_coding | GDSL-like Lipase/Acylhyd | 109 | 153 | 153 | 135 | 119 | 153 | 229 | 376 | 392 | 293 | 316 | 416 | 332.55 | 332 | 342 | 0.63 | -0.667 | 0.00181728 |
| AT5G21020.2 | | protein_coding | | 436 | 620 | 716 | 428 | 548 | 645 | 917 | 1523 | 1833 | 928 | 1455 | 1754 | 1642.9 | 1424 | 1379 | 0.629 | -0.668 | 0.00369425 |
| AT5G27320.1 | GID1C | protein_coding | alpha/beta-Hydrolases su | 236 | 238 | 225 | 147 | 136 | 181 | 496 | 585 | 576 | 319 | 361 | 492 | 392.25 | 552 | 391 | 0.629 | -0.668 | 4.41E-05 |
| AT1G30620.1 | MUR4 | protein_coding | NAD(P)-binding Rossman | 96 | 104 | 73 | 76 | 69 | 50 | 202 | 256 | 187 | 165 | 183 | 136 | 205.13 | 215 | 161 | 0.63 | -0.668 | 4.01E-05 |
| AT1G17940.1 | | protein_coding | Endosomal targeting BRC | 230 | 214 | 231 | 150 | 126 | 147 | 483 | 526 | 591 | 325 | 335 | 400 | 502.67 | 533 | 353 | 0.629 | -0.668 | 2.47E-08 |
| AT5G02440.1 | | protein_coding | | 43 | 44 | 51 | 38 | 42 | 39 | 90 | 108 | 131 | 82 | 112 | 106 | 98.32 | 110 | 100 | 0.629 | -0.669 | 0.0008576 |
| AT3G63160.1 | | protein_coding | | 2293 | 3027 | 2286 | 2569 | 2280 | 2332 | 4820 | 7437 | 5852 | 5572 | 6053 | 6342 | 11038.42 | 6036 | 5989 | 0.629 | -0.669 | 6.75E-06 |
| AT4G28030.1 | | protein_coding | Acyl-CoA N-acyltransfera | 545 | 604 | 521 | 381 | 333 | 362 | 1146 | 1484 | 1334 | 826 | 884 | 984 | 1801.25 | 1321 | 898 | 0.629 | -0.669 | 5.96E-08 |
| AT3G49420.1 | | protein_coding | Got1/Sft2-like vescicle tra | 83 | 64 | 70 | 54 | 36 | 38 | 174 | 157 | 179 | 117 | 96 | 103 | 139.74 | 170 | 105 | 0.628 | -0.671 | 1.51E-05 |
| AT3G32980.1 | PER32 | protein_coding | Peroxidase superfamily p | 245 | 137 | 144 | 214 | 138 | 106 | 515 | 337 | 369 | 464 | 366 | 288 | 487.99 | 407 | 373 | 0.628 | -0.672 | 0.00097337 |
| AT1G10865.1 | | protein_coding | | 136 | 95 | 119 | 106 | 56 | 57 | 286 | 233 | 305 | 230 | 149 | 155 | 197.38 | 275 | 178 | 0.627 | -0.672 | 0.00020104 |
| AT4G30710.1 | QWRF8 | protein_coding | Family of unknown functi | 303 | 340 | 274 | 218 | 217 | 216 | 637 | 835 | 701 | 473 | 576 | 587 | 584.15 | 724 | 545 | 0.628 | -0.672 | 2.84E-07 |
| AT4G01670.1 | | protein_coding | | 53 | 42 | 36 | 20 | 13 | 29 | 111 | 103 | 92 | 43 | 35 | 79 | 87.54 | 102 | 52 | 0.627 | -0.673 | 0.00249191 |
| AT5G04220.2 | SYT3 | protein_coding | Calcium-dependent lipid- | 211 | 204 | 179 | 124 | 119 | 107 | 444 | 501 | 458 | 269 | 316 | 291 | 378.17 | 468 | 292 | 0.627 | -0.673 | 1.86E-07 |
| AT2G24190.2 | SDR2B | protein_coding | NAD(P)-binding Rossman | 95 | 73 | 64 | 53 | 40 | 29 | 200 | 179 | 164 | 115 | 106 | 79 | 142.14 | 181 | 100 | 0.627 | -0.674 | 0.00036368 |
| AT5G62560.1 | PUB41 | protein_coding | RING/U-box superfamily | 78 | 83 | 79 | 52 | 44 | 72 | 164 | 204 | 202 | 113 | 117 | 196 | 238.9 | 190 | 142 | 0.627 | -0.674 | 2.68E-05 |
| AT1G31600.1 | | protein_coding | RNA-binding (RRM/RBD/ | 301 | 243 | 267 | 227 | 196 | 207 | 633 | 597 | 684 | 492 | 520 | 563 | 684.32 | 638 | 525 | 0.626 | -0.675 | 8.52E-10 |
| AT4G11360.1 | RHA1B | protein_coding | RING-H2 finger A1B | 39 | 70 | 66 | 47 | 46 | 73 | 82 | 172 | 169 | 102 | 122 | 199 | 210.9 | 141 | 141 | 0.626 | -0.677 | 0.00512008 |
| AT1G68030.1 | | protein_coding | RING/FYVE/PHD zinc fing | 50 | 41 | 62 | 35 | 17 | 23 | 105 | 101 | 159 | 76 | 45 | 63 | 76.54 | 122 | 61 | 0.625 | -0.678 | 0.0025636 |
| AT3G16670.1 | | protein_coding | Pollen Ole e 1 allergen ar | 362 | 503 | 392 | 468 | 380 | 470 | 761 | 1236 | 1004 | 1015 | 1009 | 1278 | 2210.17 | 1000 | 1101 | 0.625 | -0.679 | 1.99E-05 |
| AT3G62890.1 | PCMP-H82 | protein_coding | Pentatricopeptide repeat | 33 | 25 | 19 | 25 | 23 | 14 | 69 | 61 | 49 | 54 | 61 | 38 | 77.27 | 60 | 51 | 0.624 | -0.68 | 0.00122736 |
| AT5G48335.1 | | protein_coding | | 344 | 283 | 270 | 203 | 155 | 148 | 723 | 695 | 691 | 440 | 411 | 402 | 671.43 | 703 | 418 | 0.624 | -0.68 | 2.40E-10 |
| AT3G04485.1 | | other_rna | other RNA | 44 | 65 | 40 | 32 | 29 | 30 | 92 | 160 | 102 | 69 | 77 | 82 | 101.1 | 118 | 76 | 0.624 | -0.681 | 0.00219071 |
| AT4G12080.1 | AHL1 | protein_coding | AT-hook motif nuclear-lo | 28 | 28 | 16 | 15 | 11 | 19 | 59 | 69 | 41 | 33 | 29 | 52 | 55.96 | 56 | 38 | 0.624 | -0.681 | 0.00216551 |
| AT3G03702.1 | | other_rna | other RNA | 50 | 51 | 61 | 40 | 38 | 22 | 105 | 125 | 156 | 87 | 101 | 60 | 105.73 | 129 | 83 | 0.624 | -0.681 | 0.00087489 |
| AT1G21650.3 | SECA2 | protein_coding | Preprotein translocase Se | 484 | 300 | 276 | 455 | 338 | 341 | 1017 | 737 | 707 | 987 | 897 | 927 | 861.49 | 820 | 937 | 0.624 | -0.681 | 6.35E-07 |
| AT5G07830.1 | AtGUS2 | protein_coding | glucuronidase 2 | 358 | 206 | 217 | 281 | 135 | 164 | 753 | 506 | 556 | 610 | 358 | 446 | 488.99 | 605 | 471 | 0.623 | -0.682 | 5.50E-05 |
| AT4G14740.1 | | protein_coding | Plant protein of unknowr | 261 | 310 | 252 | 324 | 313 | 251 | 549 | 762 | 645 | 703 | 831 | 683 | 1127.81 | 652 | 739 | 0.623 | -0.683 | 1.65E-07 |
| AT5G46100.1 | | protein_coding | Pentatricopeptide repeat | 71 | 50 | 44 | 30 | 39 | 21 | 149 | 123 | 113 | 65 | 104 | 57 | 93.11 | 128 | 75 | 0.623 | -0.684 | 0.00379814 |
| AT4G12870.1 | | protein_coding | Gamma interferon respor | 104 | 63 | 45 | 99 | 45 | 35 | 219 | 155 | 115 | 215 | 119 | 95 | 156.6 | 163 | 143 | 0.622 | -0.685 | 0.00420433 |
| AT5G49900.1 | | protein_coding | Beta-glucosidase, GBA2 t | 92 | 82 | 79 | 69 | 76 | 52 | 193 | 201 | 202 | 150 | 202 | 141 | 169.28 | 199 | 164 | 0.622 | -0.685 | 3.43E-05 |
| AT2G27810.1 | NAT12 | protein_coding | nucleobase-ascorbate tra | 792 | 765 | 588 | 457 | 432 | 335 | 1665 | 1879 | 1505 | 991 | 1147 | 911 | 1477.77 | 1683 | 1016 | 0.622 | -0.685 | 2.01E-07 |
| AT4G31560.1 | HCF153 | protein_coding | high chlorophyll fluoresce | 353 | 385 | 277 | 222 | 189 | 227 | 742 | 946 | 709 | 482 | 502 | 617 | 942.48 | 799 | 534 | 0.621 | -0.686 | 2.63E-07 |
| AT1G61500.1 | | protein_coding | S-locus lectin protein kina | 105 | 89 | 111 | 90 | 73 | 77 | 221 | 219 | 284 | 195 | 194 | 209 | 273.94 | 241 | 199 | 0.622 | -0.686 | 8.13E-08 |
| AT1G11170.1 | | protein_coding | Protein of unknown funct | 54 | 41 | 49 | 25 | 31 | 22 | 114 | 101 | 125 | 54 | 82 | 60 | 84.85 | 113 | 65 | 0.621 | -0.687 | 0.00238853 |
| AT1G42480.1 | | protein_coding | | 403 | 274 | 233 | 240 | 160 | 135 | 847 | 673 | 596 | 521 | 425 | 367 | 576.94 | 705 | 438 | 0.621 | -0.687 | 1.88E-06 |
| AT3G51400.1 | | protein_coding | Arabidopsis protein of un | 52 | 73 | 68 | 38 | 31 | 58 | 109 | 179 | 174 | 82 | 82 | 158 | 173.78 | 154 | 107 | 0.621 | -0.688 | 0.00177007 |
| AT4G07950.1 | | protein_coding | DNA-directed RNA polym | 61 | 66 | 50 | 25 | 16 | 27 | 128 | 162 | 128 | 54 | 42 | 73 | 135.97 | 139 | 56 | 0.621 | -0.688 | 6.20E-05 |
| AT5G01175.1 | | other_rna | other RNA | 24 | 20 | 13 | 19 | 28 | 12 | 50 | 49 | 33 | 41 | 74 | 33 | 65.24 | 44 | 49 | 0.62 | -0.689 | 0.00227903 |
| AT5G12930.1 | | protein_coding | | 61 | 54 | 57 | 49 | 44 | 37 | 128 | 133 | 146 | 106 | 117 | 101 | 117.76 | 136 | 108 | 0.62 | -0.689 | 0.00048798 |
| AT5G14080.1 | | protein_coding | Tetratricopeptide repeat | 134 | 96 | 157 | 89 | 66 | 64 | 282 | 236 | 402 | 193 | 175 | 174 | 228.71 | 307 | 181 | 0.62 | -0.689 | 0.00011402 |
| AT4G37608.1 | | protein_coding | | 45 | 23 | 28 | 38 | 14 | 17 | 95 | 57 | 72 | 82 | 37 | 46 | 63.95 | 75 | 55 | 0.62 | -0.69 | 0.00678985 |
| AT3G49710.1 | PCMP-H79 | protein_coding | Pentatricopeptide repeat | 43 | 37 | 31 | 34 | 38 | 34 | 90 | 91 | 79 | 74 | 101 | 92 | 116.09 | 87 | 89 | 0.62 | -0.69 | 0.00013723 |
| AT2G22570.1 | NIC1 | protein_coding | nicotinamidase 1 | 65 | 45 | 61 | 42 | 47 | 40 | 137 | 111 | 156 | 91 | 125 | 109 | 136.6 | 135 | 108 | 0.62 | -0.69 | 0.00010353 |
| AT2G37580.1 | ATL33 | protein_coding | RING/U-box superfamily | 2 | 3 | 1 | 2 | 0 | 1 | 4 | 7 | 3 | 4 | 0 | 3 | 40.84 | 5 | 2 | 0.619 | -0.692 | 0.00855637 |
| AT1G06002.1 | | other_rna | | 18 | 26 | 11 | 11 | 12 | 7 | 38 | 64 | 28 | 24 | 32 | 19 | 63.47 | 43 | 25 | 0.619 | -0.693 | 0.0029321 |
| AT1G26940.1 | CYP23 | protein_coding | Cyclophilin-like peptidyl-p | 160 | 111 | 120 | 66 | 73 | 58 | 336 | 273 | 307 | 143 | 194 | 158 | 231.93 | 305 | 165 | 0.618 | -0.693 | 9.96E-06 |
| AT1G22770.1 | GI | protein_coding | gigantea protein (GI) | 2400 | 2621 | 2776 | 1374 | 1397 | 1471 | 5045 | 6439 | 7107 | 2980 | 3709 | 4000 | 4307.91 | 6197 | 3563 | 0.619 | -0.693 | 3.91E-06 |
| AT1G52030.2 | F-ATMBP | protein_coding | myrosinase-binding prote | 10 | 13 | 10 | 25 | 25 | 15 | 21 | 32 | 26 | 54 | 66 | 41 | 55.72 | 26 | 54 | 0.618 | -0.694 | 0.00500375 |
| AT5G65570.1 | PCMP-H47 | protein_coding | Tetratricopeptide repeat | 44 | 38 | 39 | 28 | 34 | 24 | 92 | 93 | 100 | 61 | 90 | 65 | 76.65 | 95 | 72 | 0.618 | -0.694 | 0.00420042 |
| AT5G62630.1 | HIPL2 | protein_coding | hipl2 protein precursor | 473 | 668 | 487 | 245 | 311 | 202 | 994 | 1641 | 1247 | 531 | 826 | 549 | 881.13 | 1294 | 635 | 0.618 | -0.694 | 0.00013677 |
| AT1G24240.1 | | protein_coding | Ribosomal protein L19 fa | 131 | 129 | 133 | 88 | 72 | 58 | 275 | 317 | 340 | 191 | 191 | 158 | 291.06 | 311 | 180 | 0.618 | -0.694 | 2.11E-08 |
| AT2G39900.1 | WLIN2A | protein_coding | GATA type zinc finger tra | 393 | 398 | 379 | 237 | 229 | 194 | 826 | 978 | 970 | 514 | 608 | 528 | 1090.88 | 925 | 550 | 0.618 | -0.694 | 1.03E-08 |
| AT4G14750.1 | IQD19 | protein_coding | IQ-domain 19 | 207 | 192 | 187 | 283 | 227 | 188 | 435 | 472 | 479 | 614 | 603 | 511 | 501.16 | 462 | 576 | 0.618 | -0.695 | 3.00E-08 |
| AT4G37330.1 | CYP81D4 | protein_coding | cytochrome P450, family | 265 | 261 | 260 | 188 | 185 | 196 | 557 | 641 | 666 | 408 | 491 | 533 | 574.73 | 621 | 477 | 0.618 | -0.695 | 6.15E-09 |
| AT5G63810.1 | BGAL10 | protein_coding | beta-galactosidase 10 | 841 | 1092 | 1620 | 525 | 544 | 953 | 1768 | 2683 | 4147 | 1139 | 1444 | 2592 | 2093.66 | 2866 | 1725 | 0.617 | -0.697 | 0.00934685 |
| AT5G04620.2 | ATBIOF | protein_coding | biotin F | 160 | 177 | 177 | 129 | 106 | 110 | 336 | 435 | 453 | 280 | 281 | 299 | 413.02 | 408 | 287 | 0.617 | -0.697 | 3.96E-08 |
| AT1G51645.1 | | other_rna | other RNA | 50 | 43 | 49 | 28 | 21 | 28 | 105 | 106 | 125 | 61 | 56 | 76 | 83.45 | 112 | 64 | 0.617 | -0.698 | 0.00226689 |
| AT4G32430.1 | PCMP-E40 | protein_coding | Pentatricopeptide repeat | 67 | 32 | 44 | 41 | 37 | 28 | 141 | 79 | 113 | 89 | 98 | 76 | 89.09 | 111 | 88 | 0.616 | -0.698 | 0.00189335 |
| AT1G16960.1 | | protein_coding | Ubiquitin domain-contair | 95 | 115 | 120 | 76 | 99 | 105 | 200 | 283 | 307 | 165 | 263 | 286 | 325.42 | 263 | 238 | 0.616 | -0.699 | 0.00017691 |
| AT3G23840.1 | CER26L | protein_coding | HXXXD-type acyl-transfer | 117 | 168 | 112 | 105 | 140 | 142 | 246 | 413 | 287 | 228 | 372 | 386 | 503.43 | 315 | 329 | 0.616 | -0.699 | 9.98E-05 |
| AT4G24050.1 | | protein_coding | NAD(P)-binding Rossman | 58 | 111 | 106 | 67 | 51 | 74 | 122 | 273 | 271 | 145 | 135 | 201 | 161.86 | 222 | 160 | 0.615 | -0.7 | 0.0040903 |
| AT2G22120.2 | | protein_coding | RING/FYVE/PHD zinc fing | 120 | 124 | 114 | 73 | 71 | 103 | 252 | 305 | 292 | 158 | 188 | 280 | 252.34 | 283 | 209 | 0.615 | -0.7 | 4.92E-05 |
| AT2G18890.1 | | protein_coding | Protein kinase superfamil | 227 | 291 | 201 | 131 | 151 | 91 | 477 | 715 | 515 | 284 | 401 | 247 | 473.04 | 569 | 311 | 0.616 | -0.7 | 1.09E-05 |
| AT5G23400.1 | | protein_coding | Leucine-rich repeat (LRR) | 239 | 226 | 242 | 201 | 160 | 163 | 502 | 555 | 620 | 436 | 425 | 443 | 465.21 | 559 | 435 | 0.616 | -0.7 | 4.58E-08 |
| AT2G14045.1 | | protein_coding | | 397 | 317 | 356 | 306 | 243 | 196 | 835 | 779 | 911 | 664 | 645 | 533 | 712.13 | 842 | 614 | 0.616 | -0.7 | 1.70E-09 |
| AT4G13494.1 | MIR863A | mirna | MIR863a; miRNA | 62 | 78 | 56 | 70 | 56 | 42 | 130 | 192 | 143 | 152 | 149 | 114 | 114.3 | 155 | 138 | 0.615 | -0.702 | 0.00635625 |
| AT5G28913.1 | SADHU4-1 | transposable_elem | transposable element ger | 30 | 56 | 43 | 22 | 33 | 37 | 63 | 138 | 110 | 48 | 88 | 101 | 132.83 | 104 | 79 | 0.615 | -0.702 | 0.00118158 |
| AT1G33350.1 | PCMP-E57 | protein_coding | Pentatricopeptide repeat | 36 | 18 | 31 | 24 | 17 | 20 | 76 | 44 | 79 | 52 | 45 | 54 | 69.84 | 66 | 50 | 0.615 | -0.702 | 0.00078038 |
| AT5G44450.1 | | protein_coding | methyltransferases | 330 | 238 | 271 | 195 | 155 | 193 | 694 | 585 | 694 | 423 | 411 | 525 | 550.72 | 658 | 453 | 0.615 | -0.702 | 6.43E-08 |
| AT5G06690.2 | WCRKC1 | protein_coding | WCRKC thioredoxin 1 | 655 | 833 | 1285 | 255 | 323 | 464 | 1377 | 2047 | 3290 | 553 | 857 | 1262 | 1786.92 | 2238 | 891 | 0.614 | -0.703 | 0.00661251 |
| AT5G12970.1 | | protein_coding | Calcium-dependent lipid- | 16 | 16 | 11 | 25 | 15 | 16 | 34 | 39 | 28 | 54 | 40 | 44 | 39.19 | 34 | 46 | 0.614 | -0.705 | 0.00779558 |
| AT5G40700.2 | | protein_coding | | 82 | 66 | 73 | 49 | 53 | 55 | 172 | 162 | 187 | 106 | 141 | 150 | 146.68 | 174 | 132 | 0.613 | -0.705 | 0.00012267 |
| AT3G12530.1 | GINS2 | protein_coding | PSF2 | 71 | 60 | 55 | 58 | 57 | 38 | 149 | 147 | 141 | 126 | 151 | 103 | 151.6 | 146 | 127 | 0.613 | -0.706 | 2.37E-05 |
| AT1G52080.1 | AR791 | protein_coding | actin binding protein fam | 84 | 66 | 83 | 75 | 48 | 41 | 177 | 162 | 212 | 163 | 127 | 111 | 119.85 | 184 | 134 | 0.612 | -0.707 | 0.00030281 |
| AT1G53035.1 | | protein_coding | | 79 | 81 | 127 | 63 | 42 | 79 | 166 | 199 | 325 | 137 | 112 | 215 | 207.38 | 230 | 155 | 0.612 | -0.708 | 0.00203391 |
| AT1G20230.1 | PCMP-H21 | protein_coding | Pentatricopeptide repeat | 59 | 47 | 62 | 54 | 41 | 27 | 124 | 115 | 159 | 117 | 109 | 73 | 111.41 | 133 | 100 | 0.612 | -0.708 | 0.00015598 |
| AT3G28350.1 | | pseudogene | | 32 | 51 | 47 | 27 | 24 | 24 | 67 | 125 | 120 | 59 | 64 | 65 | 65.25 | 104 | 63 | 0.612 | -0.709 | 0.00683102 |
| AT4G30560.1 | CNGC9 | protein_coding | cyclic nucleotide gated ch | 168 | 118 | 124 | 87 | 88 | 70 | 353 | 290 | 317 | 189 | 234 | 190 | 244.65 | 320 | 204 | 0.612 | -0.709 | 5.30E-07 |
| AT5G02010.1 | ROPGEF7 | protein_coding | RHO guanyl-nucleotide ex | 233 | 187 | 160 | 150 | 123 | 108 | 490 | 459 | 410 | 325 | 327 | 294 | 334.94 | 453 | 315 | 0.611 | -0.711 | 1.95E-08 |
| AT4G26255.1 | | other_rna | other RNA | 11 | 6 | 3 | 13 | 8 | 8 | 23 | 15 | 8 | 28 | 21 | 22 | 35.66 | 15 | 24 | 0.611 | -0.712 | 0.00364096 |
| AT1G18390.2 | | protein_coding | Protein kinase superfamil | 115 | 65 | 82 | 59 | 23 | 55 | 242 | 160 | 210 | 128 | 61 | 150 | 141.23 | 204 | 113 | 0.611 | -0.712 | 0.00079474 |
| AT4G07960.1 | CSLC12 | protein_coding | Cellulose-synthase-like C1 | 75 | 44 | 64 | 26 | 22 | 11 | 158 | 108 | 164 | 56 | 58 | 30 | 93.46 | 143 | 48 | 0.609 | -0.715 | 0.002578 |
| AT3G49290.1 | ABIL2 | protein_coding | ABL interactor-like protei | 278 | 232 | 201 | 149 | 134 | 115 | 584 | 570 | 515 | 323 | 356 | 313 | 533.36 | 556 | 331 | 0.609 | -0.715 | 3.06E-09 |
| AT5G18540.1 | | protein_coding | | 73 | 68 | 88 | 39 | 36 | 45 | 153 | 167 | 225 | 85 | 96 | 122 | 139.45 | 182 | 101 | 0.608 | -0.717 | 7.75E-05 |
| AT4G14550.1 | IAA14 | protein_coding | indole-3-acetic acid induc | 154 | 265 | 221 | 238 | 357 | 335 | 324 | 651 | 566 | 516 | 948 | 911 | 720.16 | 514 | 792 | 0.608 | -0.718 | 0.00069381 |
| AT1G02860.1 | BAH1 | protein_coding | SPX (SYG1/Pho81/XPR1) | 131 | 88 | 90 | 66 | 56 | 60 | 275 | 216 | 230 | 143 | 149 | 163 | 186.61 | 240 | 152 | 0.608 | -0.719 | 8.00E-06 |
| AT2G35795.1 | TIM14-1 | protein_coding | Chaperone DnaJ-domain | 169 | 160 | 157 | 115 | 78 | 108 | 355 | 393 | 402 | 249 | 207 | 294 | 413.04 | 383 | 250 | 0.608 | -0.719 | 2.94E-10 |
| AT1G14660.1 | NHX8 | protein_coding | Na+/H+ exchanger 8 | 112 | 94 | 116 | 56 | 57 | 57 | 235 | 231 | 297 | 121 | 151 | 155 | 191.59 | 254 | 142 | 0.607 | -0.72 | 3.59E-06 |
| AT5G05410.1 | DREB2A | protein_coding | DRE-binding protein 2A | 27 | 21 | 41 | 26 | 17 | 25 | 57 | 52 | 105 | 56 | 45 | 68 | 62.79 | 71 | 56 | 0.607 | -0.721 | 0.00794517 |
| AT1G10160.1 | | transposable_elem | transposable element ger | 94 | 102 | 97 | 68 | 54 | 97 | 198 | 251 | 248 | 147 | 143 | 264 | 247.46 | 232 | 185 | 0.607 | -0.721 | 0.00026572 |
| AT4G20300.2 | | protein_coding | Protein of unknown funct | 348 | 326 | 382 | 374 | 265 | 330 | 732 | 801 | 978 | 811 | 704 | 897 | 869.95 | 837 | 804 | 0.606 | -0.722 | 3.16E-08 |
| AT5G05330.1 | | protein_coding | HMG-box (high mobility | 46 | 42 | 37 | 35 | 28 | 24 | 97 | 103 | 95 | 76 | 74 | 65 | 81.81 | 98 | 72 | 0.606 | -0.723 | 0.0001989 |
| AT5G41690.1 | | protein_coding | RNA-binding (RRM/RBD/ | 60 | 64 | 50 | 33 | 28 | 17 | 126 | 157 | 128 | 72 | 74 | 46 | 122.9 | 137 | 64 | 0.606 | -0.723 | 1.91E-05 |
| AT3G10410.1 | SCPL49 | protein_coding | SERINE CARBOXYPEPTIDA | 682 | 650 | 688 | 633 | 502 | 479 | 1434 | 1597 | 1761 | 1373 | 1333 | 1303 | 1734.57 | 1597 | 1336 | 0.606 | -0.723 | 2.76E-08 |
| AT2G01540.1 | | protein_coding | Calcium-dependent lipid- | 68 | 62 | 61 | 43 | 26 | 25 | 143 | 152 | 156 | 93 | 69 | 68 | 128.65 | 150 | 77 | 0.605 | -0.724 | 2.91E-06 |
| AT5G24970.2 | | protein_coding | Protein kinase superfamil | 190 | 120 | 138 | 135 | 100 | 104 | 399 | 295 | 353 | 293 | 265 | 283 | 320.85 | 349 | 280 | 0.605 | -0.725 | 7.77E-08 |
| AT3G07525.2 | ATG10 | protein_coding | autophagocytosis-associa | 64 | 68 | 67 | 59 | 42 | 54 | 135 | 167 | 172 | 128 | 112 | 147 | 145.21 | 158 | 129 | 0.604 | -0.726 | 4.90E-06 |
| AT5G42250.1 | | protein_coding | Zinc-binding alcohol dehy | 364 | 463 | 426 | 165 | 135 | 144 | 765 | 1138 | 1091 | 358 | 358 | 392 | 1124.77 | 998 | 369 | 0.605 | -0.726 | 6.86E-07 |
| AT1G62422.1 | | protein_coding | | 54 | 54 | 26 | 18 | 10 | 23 | 114 | 133 | 67 | 39 | 27 | 63 | 67.41 | 105 | 43 | 0.604 | -0.727 | 0.00526419 |
| AT4G21650.1 | | protein_coding | Subtilase family protein | 124 | 147 | 125 | 103 | 164 | 116 | 261 | 361 | 320 | 223 | 435 | 315 | 339.72 | 314 | 324 | 0.604 | -0.727 | 0.00049971 |
| AT1G49230.1 | ATL78 | protein_coding | RING/U-box superfamily | 59 | 84 | 51 | 42 | 67 | 48 | 124 | 206 | 131 | 91 | 178 | 131 | 182.64 | 154 | 133 | 0.604 | -0.727 | 0.00028142 |
| AT4G31050.1 | LIP2P | protein_coding | Biotin/lipoate A/B protei | 225 | 229 | 241 | 157 | 128 | 118 | 473 | 563 | 617 | 341 | 340 | 321 | 422.27 | 551 | 334 | 0.604 | -0.727 | 7.39E-08 |
| AT1G30910.1 | | protein_coding | Molybdenum cofactor sul | 398 | 283 | 245 | 198 | 165 | 177 | 837 | 695 | 627 | 429 | 438 | 481 | 693.32 | 720 | 449 | 0.604 | -0.727 | 8.78E-09 |
| AT5G25475.4 | | protein_coding | AP2/B3-like transcription | 45 | 47 | 61 | 44 | 35 | 50 | 95 | 115 | 156 | 95 | 93 | 136 | 117.27 | 122 | 108 | 0.604 | -0.728 | 0.00092682 |
| AT3G05880.1 | RCI2A | protein_coding | Low temperature and sal | 1081 | 1634 | 2471 | 742 | 878 | 1185 | 2272 | 4014 | 6326 | 1609 | 2331 | 3223 | 4912.45 | 4204 | 2388 | 0.603 | -0.729 | 0.00650508 |
| AT4G14680.1 | APS3 | protein_coding | Pseudouridine synthase/a | 478 | 385 | 234 | 491 | 449 | 244 | 1005 | 946 | 599 | 1065 | 1192 | 664 | 1346.72 | 850 | 974 | 0.603 | -0.73 | 0.00012754 |
| AT3G54366.1 | | other_rna | | 108 | 140 | 124 | 98 | 77 | 135 | 227 | 344 | 317 | 213 | 204 | 367 | 355.36 | 296 | 261 | 0.603 | -0.73 | 4.45E-05 |
| AT5G62990.1 | emb1692 | protein_coding | Ubiquitin carboxyl-termir | 139 | 61 | 62 | 102 | 58 | 34 | 292 | 150 | 159 | 221 | 154 | 92 | 193.28 | 200 | 156 | 0.603 | -0.731 | 0.00296455 |
| AT1G53470.1 | MSL4 | protein_coding | mechanosensitive channe | 45 | 43 | 54 | 29 | 31 | 23 | 95 | 106 | 138 | 63 | 82 | 63 | 79.15 | 113 | 69 | 0.603 | -0.731 | 0.00199209 |
| AT4G03820.2 | | protein_coding | Protein of unknown funct | 72 | 66 | 31 | 38 | 36 | 20 | 151 | 162 | 79 | 82 | 96 | 54 | 148.66 | 131 | 77 | 0.602 | -0.731 | 0.00015957 |
| AT1G31580.1 | ECS1 | protein_coding | ECS1 | 674 | 783 | 489 | 782 | 701 | 473 | 1417 | 1924 | 1252 | 1696 | 1861 | 1286 | 2540.76 | 1531 | 1614 | 0.603 | -0.731 | 6.79E-05 |
| AT4G13780.1 | | protein_coding | methionine--tRNA ligase, | 1903 | 1380 | 1471 | 1888 | 1493 | 1434 | 4000 | 3390 | 3766 | 4095 | 3964 | 3900 | 3810.14 | 3719 | 3986 | 0.603 | -0.731 | 9.14E-11 |
| AT4G17100.2 | | protein_coding | | 850 | 1047 | 940 | 490 | 484 | 519 | 1787 | 2572 | 2406 | 1063 | 1285 | 1411 | 2156.78 | 2255 | 1253 | 0.602 | -0.732 | 7.58E-07 |
| AT4G12910.1 | SCPL20 | protein_coding | serine carboxypeptidase- | 177 | 255 | 196 | 207 | 286 | 183 | 372 | 626 | 502 | 449 | 759 | 498 | 665.71 | 500 | 569 | 0.602 | -0.733 | 0.00019237 |
| AT5G46850.2 | | protein_coding | | 102 | 77 | 83 | 57 | 55 | 39 | 214 | 189 | 212 | 124 | 146 | 106 | 158.77 | 205 | 125 | 0.601 | -0.733 | 2.63E-06 |
| AT1G53890.3 | | protein_coding | Protein of unknown funct | 25 | 28 | 24 | 44 | 43 | 27 | 53 | 69 | 61 | 95 | 114 | 73 | 94.86 | 61 | 94 | 0.601 | -0.734 | 0.00057118 |
| AT1G80880.1 | | protein_coding | Tetratricopeptide repeat | 34 | 35 | 35 | 25 | 30 | 25 | 71 | 86 | 90 | 54 | 80 | 68 | 87.2 | 82 | 67 | 0.601 | -0.734 | 0.00044915 |
| AT2G33051.1 | | other_rna | other RNA | 54 | 43 | 49 | 30 | 26 | 27 | 114 | 106 | 125 | 65 | 69 | 73 | 87.7 | 115 | 69 | 0.601 | -0.734 | 7.91E-05 |
| AT2G28480.1 | | protein_coding | RNA-binding CRS1 / YhbY | 186 | 109 | 181 | 81 | 60 | 74 | 391 | 268 | 463 | 176 | 159 | 201 | 270.24 | 374 | 179 | 0.601 | -0.734 | 3.26E-05 |
| AT4G21065.1 | PCMP-H28 | protein_coding | Tetratricopeptide repeat | 54 | 29 | 35 | 27 | 17 | 19 | 114 | 71 | 90 | 59 | 45 | 52 | 68.42 | 92 | 52 | 0.601 | -0.735 | 0.00470331 |
| AT3G03000.1 | CML18 | protein_coding | EF hand calcium-binding | 37 | 46 | 34 | 22 | 24 | 36 | 78 | 113 | 87 | 48 | 64 | 98 | 123.37 | 93 | 70 | 0.6 | -0.736 | 0.00039744 |
| AT3G46180.1 | UTR5 | protein_coding | UDP-galactose transporte | 150 | 123 | 110 | 86 | 81 | 66 | 315 | 302 | 282 | 187 | 215 | 179 | 239.01 | 300 | 194 | 0.601 | -0.736 | 4.04E-08 |
| AT1G52342.1 | | protein_coding | | 62 | 90 | 52 | 46 | 62 | 39 | 130 | 221 | 133 | 100 | 165 | 106 | 163.04 | 161 | 124 | 0.6 | -0.737 | 0.00020845 |
| AT1G10600.1 | AMSH2 | protein_coding | associated molecule with | 202 | 179 | 208 | 120 | 93 | 130 | 425 | 440 | 532 | 260 | 247 | 354 | 378.25 | 466 | 287 | 0.6 | -0.737 | 2.06E-06 |
| AT3G44713.1 | | protein_coding | | 29 | 28 | 26 | 35 | 23 | 22 | 61 | 69 | 67 | 76 | 61 | 60 | 52.77 | 66 | 66 | 0.6 | -0.738 | 0.00336059 |
| AT2G40080.1 | ELF4 | protein_coding | Protein of unknown funct | 0 | 3 | 5 | 2 | 3 | 0 | 0 | 7 | 13 | 4 | 8 | 0 | 134.96 | 7 | 4 | 0.6 | -0.738 | 5.53E-05 |
| AT5G58600.1 | PMR5 | protein_coding | Plant protein of unknowr | 277 | 230 | 261 | 96 | 111 | 156 | 582 | 565 | 668 | 208 | 295 | 424 | 564.01 | 605 | 309 | 0.599 | -0.739 | 3.14E-06 |
| AT5G14910.1 | | protein_coding | Heavy metal transport/de | 3219 | 2928 | 2745 | 1951 | 1714 | 1713 | 6767 | 7194 | 7027 | 4232 | 4550 | 4658 | 6485.05 | 6996 | 4480 | 0.599 | -0.739 | 3.17E-11 |
| AT2G04860.1 | PCMP-E74 | protein_coding | Tetratricopeptide repeat | 48 | 43 | 31 | 37 | 23 | 21 | 101 | 106 | 79 | 80 | 61 | 57 | 71.97 | 95 | 66 | 0.598 | -0.741 | 0.00043509 |
| AT2G29760.1 | PCMP-H33 | protein_coding | Tetratricopeptide repeat | 245 | 155 | 111 | 153 | 126 | 76 | 515 | 381 | 284 | 332 | 335 | 207 | 335.72 | 393 | 291 | 0.598 | -0.741 | 9.88E-05 |
| AT5G42730.1 | | pseudogene | | 71 | 53 | 75 | 47 | 50 | 60 | 149 | 130 | 192 | 102 | 133 | 163 | 149.52 | 157 | 133 | 0.598 | -0.743 | 4.93E-05 |
| AT5G02100.1 | ORP3A | protein_coding | Oxysterol-binding family | 131 | 102 | 125 | 96 | 97 | 91 | 275 | 251 | 320 | 208 | 258 | 247 | 244.97 | 282 | 238 | 0.597 | -0.743 | 1.37E-06 |
| AT1G55970.1 | HAC04 | protein_coding | histone acetyltransferase | 254 | 213 | 202 | 143 | 135 | 113 | 534 | 523 | 517 | 310 | 358 | 307 | 379.32 | 525 | 325 | 0.597 | -0.743 | 1.67E-10 |

| | | | | | | | | | | | | | | | | | | | | | |
|---|---|---|---|---|---|---|---|---|---|---|---|---|---|---|---|---|---|---|---|---|---|
| AT2G36490.1 | ROS1 | protein_coding | demeter-like 1 | 328 | 280 | 323 | 181 | 206 | 165 | 689 | 688 | 827 | 393 | 547 | 449 | 452.05 | 735 | 463 | 0.597 | -0.744 | 1.95E-07 |
| AT3G45160.1 | | protein_coding | Putative membrane lipop | 77 | 47 | 41 | 46 | 64 | 19 | 162 | 115 | 105 | 100 | 170 | 52 | 183.51 | 127 | 107 | 0.597 | -0.745 | 0.00718214 |
| AT2G24220.1 | PUP5 | protein_coding | purine permease 5 | 46 | 56 | 50 | 36 | 22 | 32 | 97 | 138 | 128 | 78 | 58 | 87 | 100.83 | 121 | 74 | 0.597 | -0.745 | 0.00011197 |
| AT4G13050.1 | FATA2 | protein_coding | Acyl-ACP thioesterase | 691 | 534 | 483 | 631 | 462 | 425 | 1453 | 1312 | 1237 | 1369 | 1227 | 1156 | 1522.06 | 1334 | 1251 | 0.597 | -0.745 | 6.19E-10 |
| AT1G61795.1 | RIC9 | protein_coding | PAK-box/P21-Rho-binding | 144 | 170 | 119 | 88 | 64 | 28 | 303 | 418 | 305 | 191 | 170 | 76 | 256.17 | 342 | 146 | 0.596 | -0.746 | 0.00039344 |
| AT3G52860.1 | MED28 | protein_coding | | 70 | 89 | 53 | 59 | 48 | 68 | 147 | 219 | 136 | 128 | 127 | 185 | 191.17 | 167 | 147 | 0.596 | -0.746 | 5.29E-05 |
| AT5G50315.1 | | transposable_elen | transposable element ger | 44 | 20 | 30 | 30 | 24 | 17 | 92 | 49 | 77 | 65 | 64 | 46 | 123.29 | 73 | 58 | 0.596 | -0.747 | 2.74E-05 |
| AT5G48960.1 | | protein_coding | HAD-superfamily hydrola | 502 | 442 | 405 | 483 | 431 | 364 | 1055 | 1086 | 1037 | 1048 | 1144 | 990 | 1081.75 | 1059 | 1061 | 0.596 | -0.748 | 6.71E-10 |
| AT5G41120.1 | | protein_coding | Esterase/lipase/thioester | 64 | 61 | 85 | 57 | 46 | 31 | 135 | 150 | 218 | 124 | 122 | 84 | 135.99 | 168 | 110 | 0.595 | -0.749 | 5.18E-05 |
| AT5G45370.2 | | protein_coding | nodulin MtN21 /EamA-lik | 56 | 54 | 64 | 40 | 29 | 27 | 118 | 133 | 164 | 87 | 77 | 73 | 118.18 | 138 | 79 | 0.595 | -0.749 | 1.63E-05 |
| AT1G74520.1 | HVA22A | protein_coding | HVA22 homologue A | 434 | 379 | 391 | 308 | 277 | 217 | 912 | 931 | 1001 | 668 | 735 | 590 | 864.72 | 948 | 664 | 0.595 | -0.749 | 3.94E-10 |
| AT3G15358.1 | | protein_coding | | 49 | 45 | 64 | 35 | 28 | 30 | 103 | 111 | 164 | 76 | 74 | 82 | 78.56 | 126 | 77 | 0.594 | -0.751 | 0.00092371 |
| AT5G51545.1 | LPA2 | protein_coding | low psii accumulation2 | 610 | 562 | 540 | 334 | 284 | 347 | 1282 | 1381 | 1382 | 724 | 754 | 944 | 1193.03 | 1348 | 807 | 0.594 | -0.751 | 8.12E-10 |
| AT4G12980.1 | | protein_coding | Auxin-responsive family p | 679 | 1308 | 784 | 831 | 1230 | 832 | 1427 | 3214 | 2007 | 1803 | 3265 | 2263 | 2438.79 | 2216 | 2444 | 0.594 | -0.752 | 0.00107447 |
| AT1G19370.1 | | protein_coding | | 516 | 501 | 569 | 249 | 217 | 255 | 1085 | 1231 | 1457 | 540 | 576 | 693 | 830.56 | 1258 | 603 | 0.594 | -0.752 | 2.81E-07 |
| AT3G13160.1 | | protein_coding | Tetratricopeptide repeat | 199 | 149 | 140 | 175 | 147 | 124 | 418 | 366 | 358 | 380 | 390 | 337 | 435.93 | 381 | 369 | 0.594 | -0.752 | 9.03E-11 |
| AT1G53885.1 | | protein_coding | Protein of unknown funct | 9 | 17 | 27 | 20 | 13 | 16 | 19 | 42 | 69 | 43 | 35 | 44 | 63.17 | 43 | 41 | 0.593 | -0.753 | 0.00587343 |
| AT5G48370.1 | | protein_coding | Thioesterase/thiol ester c | 87 | 91 | 88 | 63 | 59 | 46 | 183 | 224 | 225 | 137 | 157 | 125 | 242.09 | 211 | 140 | 0.593 | -0.753 | 1.11E-07 |
| AT4G31060.1 | ERF015 | protein_coding | Integrase-type DNA-bindi | 92 | 103 | 86 | 53 | 38 | 51 | 193 | 253 | 220 | 115 | 101 | 139 | 266.45 | 222 | 118 | 0.594 | -0.753 | 9.91E-09 |
| AT1G12220.1 | RPS5 | protein_coding | Disease resistance protei | 117 | 76 | 85 | 74 | 33 | 37 | 246 | 187 | 218 | 161 | 88 | 101 | 137.03 | 217 | 117 | 0.593 | -0.754 | 0.00166827 |
| AT2G17975.1 | | protein_coding | zinc finger (Ran-binding) | 86 | 67 | 72 | 57 | 53 | 57 | 181 | 165 | 184 | 124 | 141 | 155 | 180.2 | 177 | 140 | 0.593 | -0.754 | 1.68E-06 |
| AT3G58130.1 | | protein_coding | N-acetylglucosaminylpho | 137 | 82 | 114 | 73 | 76 | 53 | 288 | 201 | 292 | 158 | 202 | 144 | 222.19 | 260 | 168 | 0.593 | -0.754 | 1.42E-06 |
| AT5G64000.1 | SAL2 | protein_coding | Inositol monophosphatas | 12 | 12 | 14 | 12 | 12 | 11 | 25 | 29 | 36 | 26 | 32 | 30 | 32.63 | 30 | 29 | 0.592 | -0.755 | 0.00869907 |
| AT5G62350.1 | | protein_coding | Plant invertase/pectin me | 1233 | 2049 | 1151 | 735 | 879 | 930 | 2592 | 5034 | 2947 | 1594 | 2334 | 2529 | 3944.44 | 3524 | 2152 | 0.592 | -0.757 | 0.00021694 |
| AT1G58280.2 | | protein_coding | Phosphoglycerate mutase | 183 | 100 | 104 | 96 | 70 | 58 | 385 | 246 | 266 | 208 | 186 | 158 | 240.55 | 299 | 184 | 0.591 | -0.759 | 9.16E-05 |
| AT1G48270.1 | GCR1 | protein_coding | G-protein-coupled recept | 94 | 88 | 75 | 34 | 46 | 34 | 198 | 216 | 192 | 74 | 122 | 92 | 147.14 | 202 | 96 | 0.591 | -0.759 | 9.18E-07 |
| AT2G28840.1 | XBAT31 | protein_coding | XB3 ortholog 1 in Arabido | 1368 | 1592 | 1857 | 798 | 755 | 1244 | 2876 | 3911 | 4754 | 1731 | 2004 | 3383 | 3439.22 | 3847 | 2373 | 0.591 | -0.76 | 0.00023896 |
| AT5G64860.1 | DPE1 | protein_coding | disproportionating enzym | 868 | 969 | 932 | 636 | 555 | 534 | 1825 | 2381 | 2386 | 1380 | 1473 | 1452 | 2043.62 | 2197 | 1435 | 0.591 | -0.76 | 1.95E-09 |
| AT1G71990.1 | FUT13 | protein_coding | fucosyltransferase 13 | 64 | 70 | 58 | 32 | 32 | 46 | 135 | 172 | 148 | 69 | 85 | 125 | 109.53 | 152 | 93 | 0.59 | -0.761 | 0.00033161 |
| AT3G09430.1 | | protein_coding | | 22 | 15 | 17 | 17 | 17 | 30 | 46 | 37 | 44 | 37 | 45 | 82 | 59.68 | 42 | 55 | 0.59 | -0.762 | 0.00277947 |
| AT1G77170.1 | PCMP-E21 | protein_coding | Tetratricopeptide repeat | 45 | 51 | 25 | 36 | 27 | 33 | 95 | 125 | 64 | 78 | 72 | 90 | 113.28 | 95 | 80 | 0.59 | -0.762 | 0.00023801 |
| AT3G27210.1 | Y-2 | protein_coding | | 27 | 30 | 39 | 19 | 15 | 27 | 57 | 74 | 100 | 41 | 40 | 73 | 66.89 | 77 | 51 | 0.589 | -0.763 | 0.00131108 |
| AT1G01240.1 | | protein_coding | | 142 | 253 | 323 | 137 | 161 | 231 | 298 | 622 | 827 | 297 | 427 | 628 | 463.57 | 582 | 451 | 0.589 | -0.764 | 0.00521697 |
| AT5G61400.1 | | protein_coding | Pentatricopeptide repeat | 51 | 55 | 46 | 30 | 26 | 35 | 107 | 135 | 118 | 65 | 69 | 95 | 83.45 | 120 | 76 | 0.588 | -0.765 | 0.00095085 |
| AT1G20460.1 | | protein_coding | | 68 | 42 | 56 | 43 | 30 | 33 | 143 | 103 | 143 | 93 | 80 | 90 | 106.69 | 130 | 88 | 0.588 | -0.765 | 0.00017988 |
| AT5G07730.1 | | protein_coding | | 50 | 44 | 30 | 28 | 27 | 37 | 105 | 108 | 77 | 61 | 72 | 101 | 99.73 | 97 | 78 | 0.588 | -0.766 | 0.0001588 |
| AT3G53610.1 | RABE1A | protein_coding | RAB GTPase homolog 8 | 439 | 333 | 426 | 282 | 226 | 269 | 923 | 818 | 1091 | 612 | 600 | 732 | 835.84 | 944 | 648 | 0.588 | -0.766 | 6.77E-10 |
| AT4G16750.1 | ERF039 | protein_coding | Integrase-type DNA-bindi | 20 | 20 | 23 | 22 | 16 | 21 | 42 | 49 | 59 | 48 | 42 | 57 | 50.1 | 50 | 49 | 0.588 | -0.767 | 0.00756146 |
| AT4G14970.1 | | protein_coding | | 219 | 192 | 132 | 199 | 174 | 161 | 460 | 472 | 338 | 432 | 462 | 438 | 422.27 | 423 | 444 | 0.587 | -0.767 | 1.39E-08 |
| AT5G65050.3 | AGL31 | protein_coding | AGAMOUS-like 31 | 396 | 367 | 420 | 259 | 223 | 252 | 832 | 902 | 1075 | 562 | 592 | 685 | 861.62 | 936 | 613 | 0.588 | -0.767 | 3.36E-09 |
| AT5G22794.2 | | protein_coding | | 46 | 56 | 61 | 48 | 58 | 37 | 97 | 138 | 156 | 104 | 154 | 101 | 78.83 | 130 | 120 | 0.587 | -0.768 | 0.00821254 |
| AT1G61240.1 | | protein_coding | Protein of unknown funct | 323 | 225 | 195 | 199 | 147 | 140 | 679 | 553 | 499 | 432 | 390 | 381 | 462.13 | 577 | 401 | 0.587 | -0.768 | 6.50E-09 |
| AT1G20693.1 | HMGB2 | protein_coding | high mobility group B2 | 2097 | 2285 | 2773 | 1153 | 1155 | 1314 | 4408 | 5614 | 7099 | 2501 | 3066 | 3573 | 5350.88 | 5707 | 3047 | 0.587 | -0.769 | 3.03E-06 |
| AT5G24920.1 | GDU5 | protein_coding | glutamine dumper 5 | 27 | 35 | 32 | 11 | 23 | 25 | 57 | 86 | 82 | 24 | 61 | 68 | 71.05 | 75 | 51 | 0.586 | -0.77 | 0.00344868 |
| AT4G01575.1 | | protein_coding | serine protease inhibitor, | 18 | 17 | 21 | 10 | 18 | 15 | 38 | 42 | 54 | 22 | 48 | 41 | 45.8 | 45 | 37 | 0.587 | -0.77 | 0.00252833 |
| AT5G28910.2 | | protein_coding | | 61 | 115 | 66 | 37 | 41 | 34 | 128 | 283 | 169 | 80 | 109 | 92 | 133.81 | 193 | 94 | 0.586 | -0.77 | 0.0015939 |
| AT1G46554.1 | | other_rna | other RNA | 42 | 40 | 55 | 15 | 20 | 32 | 88 | 98 | 141 | 33 | 53 | 87 | 94.91 | 109 | 58 | 0.586 | -0.77 | 0.00110904 |
| AT1G09680.1 | | protein_coding | Pentatricopeptide repeat | 42 | 28 | 22 | 34 | 19 | 15 | 88 | 69 | 56 | 74 | 50 | 41 | 63.65 | 71 | 55 | 0.586 | -0.771 | 0.00435395 |
| AT4G26270.1 | PFK3 | protein_coding | phosphofructokinase 3 | 39 | 29 | 41 | 18 | 24 | 16 | 82 | 71 | 105 | 39 | 64 | 44 | 67.79 | 86 | 49 | 0.586 | -0.771 | 0.00065836 |
| AT5G04880.1 | | pseudogene | | 80 | 63 | 78 | 29 | 33 | 26 | 168 | 155 | 200 | 63 | 88 | 71 | 137.51 | 174 | 74 | 0.586 | -0.771 | 8.91E-07 |
| AT4G13200.1 | | protein_coding | | 848 | 666 | 728 | 849 | 642 | 629 | 1783 | 1636 | 1864 | 1842 | 1704 | 1711 | 2280.06 | 1761 | 1752 | 0.585 | -0.773 | 3.26E-12 |
| AT1G03730.1 | | protein_coding | | 28 | 36 | 42 | 32 | 16 | 25 | 59 | 88 | 108 | 69 | 42 | 68 | 97.7 | 85 | 60 | 0.585 | -0.774 | 0.00016911 |
| AT5G27820.1 | | protein_coding | Ribosomal L18p/L5e fami | 182 | 148 | 128 | 131 | 100 | 83 | 383 | 364 | 328 | 284 | 265 | 226 | 370.14 | 358 | 258 | 0.585 | -0.774 | 7.06E-09 |
| AT5G39970.1 | | protein_coding | catalytics | 32 | 17 | 22 | 25 | 21 | 14 | 67 | 42 | 56 | 54 | 56 | 38 | 52.65 | 55 | 49 | 0.584 | -0.775 | 0.00826382 |
| AT5G28920.1 | | protein_coding | | 28 | 21 | 27 | 15 | 8 | 11 | 59 | 52 | 69 | 33 | 21 | 30 | 41 | 60 | 28 | 0.584 | -0.775 | 0.00199265 |
| AT5G42780.1 | ZHD13 | protein_coding | homeobox protein 27 | 16 | 24 | 18 | 14 | 10 | 10 | 34 | 59 | 46 | 30 | 27 | 27 | 52.98 | 46 | 28 | 0.584 | -0.775 | 0.00061645 |
| AT3G47800.1 | | protein_coding | Galactose mutarotase-lik | 409 | 475 | 591 | 229 | 160 | 319 | 860 | 1167 | 1513 | 497 | 425 | 868 | 1166.61 | 1180 | 597 | 0.584 | -0.775 | 0.00024197 |
| AT5G05890.1 | UGT76C5 | protein_coding | UDP-Glycosyltransferase | 233 | 318 | 334 | 164 | 201 | 224 | 490 | 781 | 855 | 356 | 534 | 609 | 563.92 | 709 | 500 | 0.584 | -0.775 | 6.50E-05 |
| AT2G47300.2 | | protein_coding | ribonuclease Ps | 79 | 81 | 54 | 68 | 65 | 44 | 166 | 199 | 138 | 147 | 173 | 120 | 158.77 | 168 | 147 | 0.584 | -0.775 | 3.02E-06 |
| AT2G19170.1 | SBT2.5 | protein_coding | subtilisin-like serine prote | 399 | 364 | 322 | 223 | 263 | 221 | 839 | 894 | 824 | 484 | 698 | 601 | 677.8 | 852 | 594 | 0.584 | -0.775 | 1.72E-09 |
| AT3G02980.1 | MCC1 | protein_coding | MEIOTIC CONTROL OF CR | 31 | 14 | 28 | 20 | 13 | 15 | 65 | 34 | 72 | 43 | 35 | 41 | 52.14 | 57 | 40 | 0.584 | -0.776 | 0.00577714 |
| AT4G26060.1 | | protein_coding | Ribosomal protein L18ae | 37 | 19 | 33 | 21 | 25 | 17 | 78 | 47 | 84 | 46 | 66 | 46 | 57.59 | 70 | 53 | 0.584 | -0.776 | 0.00134704 |
| AT5G10320.1 | | protein_coding | | 106 | 112 | 115 | 73 | 70 | 56 | 223 | 275 | 294 | 158 | 186 | 152 | 204.63 | 264 | 165 | 0.584 | -0.776 | 9.10E-07 |
| AT4G12740.1 | MYH | protein_coding | HhH-GPD base excision D | 22 | 28 | 15 | 35 | 34 | 13 | 46 | 69 | 38 | 76 | 90 | 35 | 63.56 | 51 | 67 | 0.583 | -0.777 | 0.00432872 |
| AT1G67600.1 | | protein_coding | Acid phosphatase/vanadi | 31 | 29 | 30 | 27 | 18 | 19 | 65 | 71 | 77 | 59 | 48 | 52 | 53 | 71 | 53 | 0.584 | -0.777 | 0.00430226 |
| AT1G43580.1 | | protein_coding | Sphingomyelin synthetas | 373 | 324 | 295 | 196 | 161 | 150 | 784 | 796 | 755 | 425 | 427 | 408 | 644.81 | 778 | 420 | 0.584 | -0.777 | 2.48E-13 |
| AT1G52347.1 | | other_rna | other RNA | 40 | 58 | 36 | 28 | 36 | 32 | 84 | 142 | 92 | 61 | 96 | 87 | 107.85 | 106 | 81 | 0.583 | -0.779 | 0.00013161 |
| AT5G56260.1 | | protein_coding | Ribonuclease E inhibitor I | 337 | 282 | 219 | 161 | 107 | 142 | 708 | 693 | 561 | 349 | 284 | 386 | 601.11 | 654 | 340 | 0.583 | -0.779 | 1.32E-09 |
| AT2G07776.2 | | protein_coding | | 416 | 341 | 301 | 496 | 391 | 421 | 874 | 838 | 771 | 1076 | 1038 | 1145 | 486.19 | 828 | 1086 | 0.582 | -0.78 | 0.00357486 |
| AT4G12760.1 | | protein_coding | | 168 | 142 | 167 | 186 | 122 | 159 | 353 | 349 | 428 | 403 | 324 | 432 | 439.48 | 377 | 386 | 0.583 | -0.78 | 3.84E-08 |
| AT2G20480.1 | | protein_coding | | 23 | 17 | 16 | 14 | 11 | 9 | 48 | 42 | 41 | 30 | 29 | 24 | 40.13 | 44 | 28 | 0.582 | -0.782 | 0.00302196 |
| AT4G33620.1 | ULP2A | protein_coding | Cysteine proteinases supe | 139 | 133 | 134 | 97 | 68 | 63 | 292 | 327 | 343 | 210 | 181 | 171 | 197.69 | 321 | 187 | 0.582 | -0.782 | 3.91E-05 |
| AT2G34400.1 | PCMP-E23 | protein_coding | Pentatricopeptide repeat | 27 | 17 | 20 | 31 | 22 | 9 | 57 | 42 | 51 | 67 | 58 | 24 | 64.76 | 50 | 50 | 0.581 | -0.783 | 0.00492611 |
| AT4G19645.1 | | protein_coding | TRAM, LAG1 and CLN8 (T | 44 | 44 | 69 | 26 | 30 | 26 | 92 | 108 | 177 | 56 | 80 | 71 | 86.8 | 126 | 69 | 0.581 | -0.783 | 0.0021271 |
| AT4G14210.1 | PDS | protein_coding | phytoene desaturase 3 | 2659 | 2373 | 2328 | 2491 | 2341 | 2153 | 5589 | 5830 | 5960 | 5403 | 6215 | 5855 | 7019.33 | 5793 | 5824 | 0.581 | -0.783 | 1.73E-11 |
| AT4G11690.1 | | protein_coding | Pentatricopeptide repeat | 34 | 26 | 28 | 17 | 22 | 8 | 71 | 64 | 72 | 37 | 58 | 22 | 48.19 | 69 | 39 | 0.581 | -0.784 | 0.00451677 |
| AT1G24148.1 | | protein_coding | | 69 | 62 | 46 | 57 | 27 | 23 | 145 | 152 | 118 | 124 | 72 | 63 | 130.19 | 138 | 86 | 0.581 | -0.784 | 0.00035808 |
| AT3G04620.1 | | protein_coding | Alba DNA/RNA-binding p | 25 | 22 | 13 | 30 | 28 | 23 | 53 | 54 | 33 | 65 | 74 | 63 | 74.62 | 47 | 67 | 0.581 | -0.784 | 0.00034025 |
| AT5G61560.1 | PUB51 | protein_coding | U-box domain-containing | 97 | 75 | 89 | 63 | 39 | 53 | 204 | 184 | 228 | 137 | 104 | 144 | 124.33 | 205 | 128 | 0.581 | -0.784 | 0.00012838 |
| AT1G55670.1 | PSAG | protein_coding | photosystem I subunit G | 12968 | 17545 | 13909 | 8424 | 8773 | 9530 | 27260 | 43105 | 35608 | 18272 | 23290 | 25916 | 67415.53 | 35324 | 22493 | 0.581 | -0.784 | 5.98E-05 |
| AT1G69572.1 | | other_rna | other RNA | 38 | 30 | 48 | 24 | 18 | 22 | 80 | 74 | 123 | 52 | 48 | 60 | 84.45 | 92 | 53 | 0.58 | -0.785 | 0.00013481 |
| AT5G44580.1 | | protein_coding | | 992 | 511 | 523 | 1011 | 517 | 500 | 2085 | 1255 | 1339 | 2193 | 1373 | 1360 | 1441.88 | 1560 | 1642 | 0.58 | -0.785 | 8.19E-05 |
| AT2G35310.1 | | protein_coding | Transcriptional factor B3 | 33 | 26 | 27 | 23 | 22 | 16 | 69 | 64 | 69 | 50 | 58 | 44 | 51.69 | 67 | 51 | 0.58 | -0.786 | 0.00117911 |
| AT1G04790.1 | | protein_coding | RING/U-box superfamily | 413 | 336 | 322 | 203 | 181 | 175 | 868 | 825 | 824 | 440 | 481 | 476 | 599.7 | 839 | 466 | 0.58 | -0.786 | 9.49E-14 |
| AT1G18265.1 | | protein_coding | Protein of unknown funct | 93 | 129 | 129 | 45 | 48 | 47 | 195 | 317 | 330 | 98 | 127 | 128 | 191.05 | 281 | 118 | 0.58 | -0.787 | 0.00254967 |
| AT1G74670.1 | GASA6 | protein_coding | Gibberellin-regulated fam | 2684 | 4839 | 4566 | 1863 | 3394 | 3100 | 5642 | 11889 | 11689 | 4041 | 9010 | 8430 | 8545.61 | 9740 | 7160 | 0.579 | -0.788 | 0.00342155 |
| AT4G15070.1 | | protein_coding | Cysteine/Histidine-rich C1 | 65 | 49 | 55 | 53 | 31 | 49 | 137 | 120 | 141 | 115 | 82 | 133 | 125.6 | 133 | 110 | 0.579 | -0.788 | 4.61E-06 |
| AT1G67860.1 | | protein_coding | | 24 | 39 | 53 | 45 | 67 | 62 | 50 | 96 | 136 | 98 | 178 | 169 | 196.81 | 94 | 148 | 0.579 | -0.789 | 0.00074909 |
| AT4G40065.1 | | other_rna | other RNA | 123 | 111 | 90 | 58 | 51 | 27 | 259 | 273 | 230 | 126 | 135 | 73 | 249.23 | 254 | 111 | 0.578 | -0.79 | 4.83E-05 |
| AT3G10300.3 | CML49 | protein_coding | Calcium-binding EF-hand | 118 | 128 | 132 | 89 | 61 | 63 | 248 | 314 | 338 | 193 | 162 | 171 | 358.73 | 300 | 175 | 0.578 | -0.79 | 3.60E-08 |
| AT5G61010.1 | ATEXO70E2 | protein_coding | exocyst subunit exo70 fa | 263 | 218 | 184 | 231 | 149 | 77 | 553 | 536 | 471 | 501 | 396 | 209 | 434.85 | 520 | 369 | 0.578 | -0.791 | 0.00020087 |
| AT2G31010.1 | | protein_coding | Protein kinase superfamil | 606 | 352 | 511 | 288 | 173 | 263 | 1274 | 865 | 1308 | 625 | 459 | 715 | 836.42 | 1149 | 600 | 0.578 | -0.791 | 6.53E-07 |
| AT3G06435.1 | | protein_coding | Expressed protein | 30 | 28 | 61 | 67 | 61 | 59 | 63 | 69 | 156 | 145 | 162 | 160 | 83.93 | 96 | 156 | 0.578 | -0.792 | 0.007814 |
| AT3G42640.1 | AHA8 | protein_coding | H(+)-ATPase 8 | 155 | 192 | 204 | 81 | 84 | 106 | 326 | 472 | 522 | 176 | 223 | 288 | 297.95 | 440 | 229 | 0.578 | -0.792 | 8.19E-06 |
| AT5G47240.1 | NUDT8 | protein_coding | nudix hydrolase homolog | 293 | 415 | 438 | 142 | 170 | 93 | 616 | 1020 | 1121 | 308 | 451 | 253 | 726.42 | 919 | 337 | 0.577 | -0.793 | 0.0002087 |
| AT1G13670.1 | | protein_coding | | 67 | 66 | 71 | 45 | 36 | 94 | 141 | 162 | 182 | 98 | 96 | 256 | 145.38 | 162 | 150 | 0.577 | -0.794 | 0.00393475 |
| AT1G33720.1 | CYP76C6 | protein_coding | cytochrome P450, family | 37 | 31 | 36 | 20 | 19 | 16 | 78 | 76 | 92 | 43 | 50 | 44 | 84.94 | 82 | 46 | 0.577 | -0.794 | 5.02E-06 |
| AT1G56000.1 | | protein_coding | FAD/NAD(P)-binding oxid | 241 | 228 | 211 | 171 | 124 | 140 | 507 | 560 | 540 | 371 | 329 | 381 | 451.02 | 536 | 360 | 0.576 | -0.795 | 1.81E-10 |
| AT4G13160.1 | | protein_coding | Protein of unknown funct | 110 | 102 | 111 | 139 | 115 | 124 | 231 | 251 | 284 | 302 | 305 | 337 | 297.37 | 255 | 315 | 0.576 | -0.796 | 1.97E-08 |
| AT5G51100.1 | FSD2 | protein_coding | Fe superoxide dismutase | 552 | 441 | 403 | 426 | 344 | 306 | 1160 | 1083 | 1032 | 924 | 913 | 832 | 1154.5 | 1092 | 890 | 0.576 | -0.796 | 8.16E-10 |
| AT5G54640.1 | RAT5 | protein_coding | Histone superfamily prot | 215 | 238 | 160 | 137 | 132 | 140 | 452 | 585 | 410 | 297 | 350 | 381 | 554.55 | 482 | 343 | 0.576 | -0.796 | 3.66E-10 |
| AT1G65560.1 | | protein_coding | Zinc-binding dehydrogen | 72 | 50 | 38 | 71 | 49 | 41 | 151 | 123 | 97 | 154 | 130 | 111 | 140.86 | 124 | 132 | 0.575 | -0.797 | 3.02E-06 |
| AT1G32090.1 | | protein_coding | early-responsive to dehyd | 667 | 566 | 569 | 312 | 278 | 280 | 1402 | 1391 | 1457 | 677 | 738 | 761 | 1086.15 | 1417 | 725 | 0.575 | -0.797 | 8.13E-12 |
| AT1G12790.1 | | protein_coding | | 65 | 82 | 97 | 31 | 49 | 52 | 137 | 201 | 248 | 67 | 130 | 141 | 128.5 | 195 | 113 | 0.575 | -0.799 | 0.00174352 |
| AT1G62800.2 | ASP4 | protein_coding | aspartate aminotransfera | 115 | 74 | 49 | 76 | 63 | 31 | 242 | 182 | 125 | 165 | 167 | 84 | 196.94 | 183 | 139 | 0.574 | -0.8 | 0.0002812 |
| AT2G34010.1 | | protein_coding | | 100 | 102 | 59 | 65 | 30 | 25 | 210 | 251 | 151 | 141 | 80 | 68 | 199.06 | 204 | 96 | 0.574 | -0.8 | 9.25E-05 |
| AT2G03310.1 | | protein_coding | | 301 | 300 | 283 | 150 | 160 | 132 | 633 | 737 | 724 | 325 | 425 | 359 | 535.79 | 698 | 370 | 0.574 | -0.8 | 1.61E-11 |
| AT4G15000.1 | RPL27C | protein_coding | Ribosomal L27e protein f | 3851 | 3367 | 2209 | 3724 | 2654 | 2957 | 8095 | 8272 | 5655 | 8078 | 7046 | 8041 | 10467.15 | 7341 | 7722 | 0.574 | -0.801 | 2.30E-07 |
| AT5G41960.1 | | protein_coding | | 138 | 99 | 137 | 85 | 57 | 55 | 290 | 243 | 351 | 184 | 151 | 150 | 273.61 | 295 | 162 | 0.574 | -0.801 | 7.27E-08 |
| AT2G35260.1 | | protein_coding | | 1796 | 2016 | 1536 | 868 | 887 | 810 | 3775 | 4953 | 3932 | 1883 | 2355 | 2203 | 4516.22 | 4220 | 2147 | 0.574 | -0.801 | 1.91E-09 |
| AT4G35760.1 | LTO1 | protein_coding | NAD(P)H dehydrogenase | 959 | 846 | 730 | 616 | 549 | 476 | 2016 | 2078 | 1869 | 1336 | 1457 | 1294 | 1687.11 | 1988 | 1362 | 0.574 | -0.801 | 8.05E-13 |
| AT1G75060.1 | | protein_coding | | 34 | 43 | 28 | 15 | 21 | 20 | 71 | 106 | 72 | 33 | 56 | 54 | 58.67 | 83 | 48 | 0.573 | -0.802 | 0.00482775 |
| AT4G14490.1 | | protein_coding | SMAD/FHA domain-conta | 16 | 16 | 19 | 23 | 22 | 23 | 34 | 39 | 49 | 50 | 58 | 63 | 102.69 | 41 | 57 | 0.574 | -0.802 | 5.87E-06 |
| AT2G17730.2 | NIP2 | protein_coding | NEP-interacting protein 2 | 104 | 79 | 88 | 47 | 43 | 68 | 219 | 194 | 225 | 102 | 114 | 185 | 245.72 | 213 | 134 | 0.574 | -0.802 | 3.74E-07 |
| AT4G13070.1 | | protein_coding | RNA-binding CRS1 / YhbY | 128 | 81 | 106 | 112 | 90 | 82 | 269 | 199 | 271 | 243 | 239 | 223 | 254.86 | 246 | 235 | 0.573 | -0.803 | 2.16E-07 |
| AT2G37680.1 | | protein_coding | | 113 | 107 | 116 | 60 | 52 | 40 | 238 | 263 | 297 | 130 | 138 | 109 | 234.02 | 266 | 126 | 0.573 | -0.803 | 3.05E-09 |
| AT4G12880.1 | ENODL19 | protein_coding | early nodulin-like protein | 873 | 686 | 633 | 966 | 853 | 649 | 1835 | 1685 | 1621 | 2095 | 2265 | 1765 | 2718.5 | 1714 | 2042 | 0.573 | -0.803 | 3.97E-11 |
| AT3G45590.1 | SEN1 | protein_coding | splicing endonuclease 1 | 31 | 21 | 32 | 31 | 22 | 19 | 65 | 52 | 82 | 67 | 58 | 52 | 68.01 | 66 | 59 | 0.573 | -0.804 | 0.00022263 |
| AT3G09550.1 | | protein_coding | Ankyrin repeat family pro | 29 | 42 | 28 | 6 | 12 | 9 | 61 | 103 | 72 | 13 | 32 | 24 | 66.52 | 79 | 23 | 0.572 | -0.805 | 0.00131272 |
| AT5G55125.1 | | protein_coding | Ribosomal protein L31 | 172 | 119 | 114 | 137 | 104 | 82 | 362 | 292 | 292 | 297 | 276 | 223 | 370.23 | 315 | 265 | 0.572 | -0.805 | 3.13E-08 |
| AT2G39920.1 | | protein_coding | HAD superfamily, subfam | 187 | 276 | 318 | 71 | 101 | 96 | 393 | 678 | 814 | 154 | 268 | 261 | 442.49 | 628 | 228 | 0.572 | -0.806 | 0.00054794 |
| AT4G29170.1 | MND1 | protein_coding | Mnd1 family protein | 71 | 32 | 60 | 43 | 55 | 44 | 149 | 79 | 154 | 93 | 146 | 120 | 135.35 | 127 | 120 | 0.572 | -0.806 | 0.00015786 |
| AT5G64640.1 | PME64 | protein_coding | Plant invertase/pectin me | 110 | 97 | 74 | 51 | 27 | 41 | 231 | 238 | 189 | 111 | 72 | 111 | 227.26 | 219 | 98 | 0.572 | -0.807 | 1.60E-08 |
| AT4G36945.1 | | protein_coding | PLC-like phosphodiestera | 195 | 177 | 163 | 119 | 99 | 78 | 410 | 435 | 417 | 258 | 263 | 212 | 358.43 | 421 | 244 | 0.571 | -0.807 | 9.11E-12 |
| AT2G47920.1 | NET3C | protein_coding | Kinase interacting (KIP1-l | 41 | 40 | 42 | 24 | 35 | 24 | 86 | 98 | 108 | 52 | 93 | 65 | 105.54 | 97 | 70 | 0.571 | -0.808 | 8.88E-05 |
| AT3G53210.1 | | protein_coding | nodulin MtN21 /EamA-lik | 57 | 55 | 34 | 27 | 29 | 16 | 120 | 135 | 87 | 59 | 77 | 44 | 77.98 | 114 | 60 | 0.571 | -0.809 | 0.00141563 |
| AT1G04990.2 | | protein_coding | Zinc finger C-x8-C-x5-C-x3 | 173 | 138 | 150 | 142 | 98 | 127 | 364 | 339 | 384 | 308 | 260 | 345 | 369.9 | 362 | 304 | 0.571 | -0.809 | 2.92E-10 |
| AT3G26840.1 | | protein_coding | Esterase/lipase/thioester | 108 | 70 | 63 | 83 | 42 | 33 | 227 | 172 | 161 | 180 | 112 | 90 | 132.77 | 187 | 127 | 0.57 | -0.811 | 0.00031596 |
| AT1G07135.1 | | protein_coding | glycine-rich protein | 11 | 7 | 13 | 15 | 2 | 5 | 23 | 17 | 33 | 33 | 5 | 14 | 137.9 | 24 | 17 | 0.569 | -0.813 | 9.94E-05 |
| AT1G78910.1 | | protein_coding | Pseudouridine synthase f | 51 | 60 | 48 | 39 | 38 | 39 | 107 | 147 | 123 | 85 | 101 | 106 | 102.45 | 126 | 97 | 0.569 | -0.813 | 6.16E-05 |
| AT2G33180.1 | | protein_coding | | 752 | 545 | 378 | 360 | 261 | 210 | 1581 | 1339 | 968 | 781 | 693 | 571 | 1220.52 | 1296 | 682 | 0.569 | -0.813 | 2.66E-07 |
| AT5G67640.1 | | protein_coding | | 53 | 38 | 45 | 34 | 22 | 27 | 111 | 93 | 115 | 74 | 58 | 73 | 94.81 | 106 | 68 | 0.568 | -0.815 | 3.77E-06 |
| AT1G55110.1 | IDD7 | protein_coding | indeterminate(ID)-domai | 170 | 191 | 215 | 127 | 148 | 146 | 357 | 469 | 550 | 275 | 393 | 397 | 365.75 | 459 | 355 | 0.569 | -0.815 | 1.44E-06 |
| AT5G11150.1 | VAMP713 | protein_coding | vesicle-associated memb | 230 | 263 | 365 | 159 | 139 | 151 | 483 | 646 | 934 | 345 | 369 | 411 | 582.59 | 688 | 375 | 0.567 | -0.818 | 1.65E-06 |
| AT5G10380.1 | ATL55 | protein_coding | RING/U-box superfamily | 33 | 18 | 23 | 18 | 13 | 10 | 69 | 44 | 59 | 39 | 35 | 27 | 47.01 | 57 | 34 | 0.567 | -0.819 | 0.00182494 |
| AT4G14300.1 | RNP1 | protein_coding | RNA-binding (RRM/RBD/ | 741 | 686 | 570 | 636 | 612 | 600 | 1558 | 1685 | 1459 | 1380 | 1625 | 1632 | 2038.71 | 1567 | 1546 | 0.567 | -0.819 | 1.72E-12 |
| AT1G16825.1 | RTNLB23 | protein_coding | Reticulon family protein | 44 | 32 | 33 | 34 | 21 | 28 | 92 | 79 | 84 | 74 | 56 | 76 | 97.2 | 85 | 69 | 0.566 | -0.82 | 3.49E-06 |
| AT4G33990.1 | EMB2758 | protein_coding | Tetratricopeptide repeat | 77 | 40 | 59 | 37 | 36 | 24 | 162 | 98 | 151 | 80 | 96 | 65 | 102.58 | 137 | 80 | 0.566 | -0.821 | 0.00010804 |
| AT2G47880.1 | GRXC13 | protein_coding | Glutaredoxin family prote | 24 | 23 | 9 | 15 | 16 | 8 | 50 | 57 | 23 | 33 | 42 | 22 | 111.4 | 43 | 32 | 0.566 | -0.821 | 4.78E-05 |
| AT2G16990.1 | | protein_coding | Major facilitator superfar | 32 | 61 | 48 | 28 | 34 | 21 | 67 | 150 | 123 | 61 | 90 | 57 | 107.69 | 113 | 69 | 0.566 | -0.822 | 0.00226026 |
| AT5G16420.1 | | protein_coding | Pentatricopeptide repeat | 43 | 33 | 23 | 33 | 21 | 20 | 90 | 81 | 59 | 72 | 56 | 54 | 78.64 | 77 | 61 | 0.564 | -0.825 | 0.00025805 |
| AT5G63290.1 | | protein_coding | Radical SAM superfamily | 100 | 58 | 56 | 84 | 40 | 34 | 210 | 142 | 143 | 182 | 106 | 92 | 153.33 | 165 | 127 | 0.564 | -0.825 | 0.00018897 |
| AT4G16807.1 | | protein_coding | | 96 | 59 | 70 | 51 | 44 | 35 | 202 | 145 | 179 | 111 | 117 | 95 | 111.22 | 175 | 108 | 0.564 | -0.826 | 0.00039658 |
| AT4G13220.1 | | protein_coding | | 499 | 312 | 277 | 445 | 309 | 247 | 1049 | 767 | 709 | 965 | 820 | 672 | 1295.29 | 842 | 819 | 0.564 | -0.826 | 7.76E-08 |
| AT2G18735.1 | | other_rna | other RNA | 35 | 37 | 37 | 34 | 28 | 28 | 74 | 91 | 95 | 74 | 74 | 76 | 81.68 | 87 | 75 | 0.563 | -0.828 | 0.0003167 |

| | | | | | | | | | | | | | | | | | | | |
|---|---|---|---|---|---|---|---|---|---|---|---|---|---|---|---|---|---|---|---|
| AT5G54520.1 | | protein_coding | Transducin/WD40 repeat | 83 | 62 | 71 | 22 | 23 | 30 | 174 | 152 | 182 | 48 | 61 | 82 | 109.49 | 169 | 64 | 0.563 | -0.828 | 9.79E-05 |
| AT5G56310.1 | PCMP-E13 | protein_coding | Pentatricopeptide repeat | 45 | 35 | 34 | 23 | 18 | 13 | 95 | 86 | 87 | 50 | 48 | 35 | 70.21 | 89 | 44 | 0.563 | -0.828 | 4.90E-05 |
| AT1G12730.1 | | protein_coding | GPI transamidase subunit | 97 | 137 | 141 | 62 | 57 | 48 | 204 | 337 | 361 | 134 | 151 | 131 | 229.52 | 301 | 139 | 0.563 | -0.828 | 5.74E-06 |
| AT5G01200.1 | | protein_coding | Duplicated homeodomain | 11 | 23 | 16 | 23 | 18 | 11 | 23 | 57 | 41 | 50 | 48 | 30 | 47.99 | 40 | 43 | 0.562 | -0.831 | 0.00455919 |
| AT4G13850.1 | RBG2 | protein_coding | glycine-rich RNA-binding | 335 | 274 | 200 | 318 | 221 | 160 | 704 | 673 | 512 | 690 | 587 | 435 | 3436.04 | 630 | 571 | 0.562 | -0.831 | 7.29E-08 |
| AT2G38680.1 | | protein_coding | 5'-nucleotidases;magnesi | 124 | 107 | 98 | 57 | 42 | 46 | 261 | 263 | 251 | 124 | 112 | 125 | 221.84 | 258 | 120 | 0.562 | -0.831 | 3.08E-10 |
| AT1G65230.1 | | protein_coding | Uncharacterized conserve | 1063 | 1351 | 1252 | 768 | 740 | 724 | 2235 | 3319 | 3205 | 1666 | 1965 | 1969 | 2928.61 | 2920 | 1867 | 0.562 | -0.832 | 1.27E-08 |
| AT4G21860.1 | MSRB2 | protein_coding | methionine sulfoxide red | 2033 | 1647 | 1500 | 1124 | 906 | 1065 | 4274 | 4046 | 3840 | 2438 | 2405 | 2896 | 4058.79 | 4053 | 2580 | 0.561 | -0.833 | 3.42E-12 |
| AT1G07730.2 | DIR25 | protein_coding | Disease resistance-respor | 3 | 11 | 4 | 7 | 4 | 7 | 6 | 27 | 10 | 15 | 11 | 19 | 23.97 | 14 | 15 | 0.561 | -0.835 | 0.00755538 |
| AT1G79440.1 | ALDH5F1 | protein_coding | aldehyde dehydrogenase | 801 | 1118 | 1343 | 592 | 482 | 664 | 1684 | 2747 | 3438 | 1284 | 1280 | 1806 | 2325.52 | 2623 | 1457 | 0.56 | -0.835 | 5.16E-05 |
| AT1G18680.1 | | protein_coding | HNH endonuclease doma | 58 | 45 | 52 | 37 | 21 | 37 | 122 | 111 | 133 | 80 | 56 | 101 | 120.53 | 122 | 79 | 0.561 | -0.835 | 3.35E-05 |
| AT1G67910.1 | | protein_coding | | 176 | 143 | 152 | 133 | 83 | 250 | 370 | 351 | 389 | 288 | 220 | 680 | 396 | 370 | 396 | 0.559 | -0.838 | 0.00171321 |
| AT3G55580.1 | | protein_coding | Regulator of chromosome | 31 | 50 | 43 | 21 | 26 | 21 | 65 | 123 | 110 | 46 | 69 | 57 | 87.55 | 99 | 57 | 0.559 | -0.838 | 0.00019656 |
| AT2G35030.1 | PCMP-E15 | protein_coding | Pentatricopeptide repeat | 54 | 37 | 44 | 42 | 43 | 29 | 114 | 91 | 113 | 91 | 114 | 79 | 96.99 | 106 | 95 | 0.559 | -0.839 | 0.00010756 |
| AT2G02750.1 | PCMP-E22 | protein_coding | Pentatricopeptide repeat | 54 | 44 | 63 | 28 | 36 | 32 | 114 | 108 | 161 | 61 | 96 | 87 | 85.41 | 128 | 81 | 0.559 | -0.84 | 0.00018401 |
| AT4G37170.1 | PCMP-H5 | protein_coding | Pentatricopeptide repeat | 42 | 42 | 32 | 38 | 35 | 18 | 88 | 103 | 82 | 82 | 93 | 49 | 84.9 | 91 | 75 | 0.558 | -0.84 | 0.00010834 |
| AT4G12730.1 | FLA2 | protein_coding | FASCICLIN-like arabinogal | 1087 | 1276 | 680 | 1388 | 945 | 1632 | 2285 | 3135 | 1741 | 3011 | 2509 | 4438 | 9405.31 | 2387 | 3319 | 0.558 | -0.841 | 4.86E-05 |
| AT4G15420.1 | | protein_coding | Ubiquitin fusion degradat | 265 | 254 | 273 | 284 | 250 | 312 | 557 | 624 | 699 | 616 | 664 | 848 | 633.94 | 627 | 709 | 0.558 | -0.841 | 2.89E-11 |
| AT4G12620.1 | ORC1B | protein_coding | origin of replication comp | 282 | 194 | 202 | 271 | 169 | 153 | 593 | 477 | 517 | 588 | 449 | 416 | 459.56 | 529 | 484 | 0.558 | -0.842 | 2.92E-10 |
| AT3G50870.1 | GATA18 | protein_coding | GATA type zinc finger tra | 12 | 24 | 18 | 12 | 14 | 20 | 25 | 59 | 46 | 26 | 37 | 54 | 43.55 | 43 | 39 | 0.557 | -0.843 | 0.00852454 |
| AT5G57080.1 | | protein_coding | | 11 | 8 | 11 | 5 | 6 | 9 | 23 | 20 | 28 | 11 | 16 | 24 | 35.92 | 24 | 17 | 0.558 | -0.843 | 0.00140626 |
| AT1G72420.1 | | protein_coding | NADH:ubiquinone oxidor | 98 | 67 | 84 | 90 | 48 | 44 | 206 | 165 | 215 | 195 | 127 | 120 | 190.87 | 195 | 147 | 0.557 | -0.844 | 4.76E-06 |
| AT1G07390.3 | AtRLP1 | protein_coding | receptor like protein 1 | 136 | 44 | 55 | 113 | 45 | 54 | 286 | 108 | 141 | 245 | 119 | 147 | 119.45 | 178 | 170 | 0.557 | -0.845 | 0.0089286 |
| AT5G42440.1 | | protein_coding | Protein kinase superfamil | 13 | 16 | 16 | 13 | 14 | 17 | 27 | 39 | 41 | 28 | 37 | 46 | 45.47 | 36 | 37 | 0.557 | -0.845 | 0.00193296 |
| AT1G22180.2 | | protein_coding | Sec14p-like phosphatidyli | 68 | 66 | 71 | 40 | 38 | 42 | 143 | 162 | 182 | 87 | 101 | 114 | 115.65 | 162 | 101 | 0.556 | -0.846 | 2.42E-05 |
| AT4G26670.1 | TIM22-2 | protein_coding | Mitochondrial import inn | 435 | 479 | 433 | 186 | 220 | 191 | 914 | 1177 | 1109 | 403 | 584 | 519 | 1422.88 | 1067 | 502 | 0.556 | -0.847 | 3.28E-09 |
| AT2G37320.1 | PCMP-E50 | protein_coding | Tetratricopeptide repeat | 25 | 19 | 22 | 23 | 17 | 18 | 53 | 47 | 56 | 50 | 45 | 49 | 53.73 | 52 | 48 | 0.556 | -0.848 | 0.00061648 |
| AT5G13760.1 | | protein_coding | Plasma-membrane cholin | 237 | 216 | 203 | 168 | 143 | 157 | 498 | 531 | 520 | 364 | 380 | 427 | 470.21 | 516 | 390 | 0.556 | -0.848 | 1.67E-13 |
| AT4G29540.2 | LPXA | protein_coding | bacterial transferase hexa | 45 | 48 | 31 | 37 | 26 | 21 | 95 | 118 | 79 | 80 | 69 | 57 | 99.03 | 97 | 69 | 0.555 | -0.849 | 5.99E-06 |
| AT1G62620.1 | | protein_coding | Flavin-binding monooxyg | 17 | 8 | 14 | 8 | 7 | 8 | 36 | 20 | 36 | 17 | 19 | 22 | 33 | 31 | 19 | 0.554 | -0.851 | 0.002632 |
| AT3G49740.1 | PCMP-E84 | protein_coding | Tetratricopeptide repeat | 67 | 37 | 41 | 43 | 29 | 35 | 141 | 91 | 105 | 93 | 77 | 95 | 97.47 | 112 | 88 | 0.554 | -0.851 | 0.00037045 |
| AT3G50420.1 | PCMP-E85 | protein_coding | Pentatricopeptide repeat | 45 | 33 | 28 | 30 | 26 | 24 | 95 | 81 | 72 | 65 | 69 | 65 | 66.19 | 83 | 66 | 0.554 | -0.852 | 2.99E-05 |
| AT5G60190.1 | NEDP1 | protein_coding | Cysteine proteinases supe | 56 | 52 | 62 | 57 | 43 | 39 | 118 | 128 | 159 | 124 | 114 | 106 | 138.45 | 135 | 115 | 0.554 | -0.852 | 1.01E-05 |
| AT2G34200.1 | | protein_coding | RING/FYVE/PHD zinc fing | 82 | 80 | 93 | 25 | 28 | 37 | 172 | 197 | 238 | 54 | 74 | 101 | 138.66 | 202 | 76 | 0.554 | -0.853 | 9.57E-06 |
| AT1G70420.1 | | protein_coding | Protein of unknown funct | 157 | 374 | 399 | 85 | 119 | 121 | 330 | 919 | 1021 | 184 | 316 | 329 | 409.29 | 757 | 276 | 0.552 | -0.856 | 0.00425454 |
| AT1G76590.1 | | protein_coding | PLATZ transcription facto | 32 | 46 | 39 | 14 | 20 | 24 | 67 | 113 | 100 | 30 | 53 | 65 | 78.05 | 93 | 49 | 0.553 | -0.856 | 0.00142659 |
| AT5G37590.1 | | protein_coding | Tetratricopeptide repeat | 189 | 106 | 101 | 98 | 81 | 64 | 397 | 260 | 259 | 213 | 215 | 174 | 267.21 | 305 | 201 | 0.553 | -0.856 | 1.08E-06 |
| AT4G37480.1 | | protein_coding | Chaperone DnaJ-domain | 201 | 185 | 204 | 95 | 82 | 96 | 423 | 455 | 522 | 206 | 218 | 261 | 301.69 | 467 | 228 | 0.552 | -0.856 | 1.48E-10 |
| AT4G15090.1 | FAR1 | protein_coding | FRS (FAR1 Related Seque | 419 | 410 | 435 | 362 | 354 | 356 | 881 | 1007 | 1114 | 785 | 940 | 968 | 947.5 | 1001 | 898 | 0.553 | -0.856 | 4.36E-12 |
| AT4G11980.1 | NUDT14 | protein_coding | nudix hydrolase homolog | 211 | 180 | 126 | 85 | 83 | 76 | 444 | 442 | 323 | 184 | 220 | 207 | 343.71 | 403 | 204 | 0.552 | -0.857 | 1.62E-10 |
| AT2G16530.1 | | protein_coding | 3-oxo-5-alpha-steroid 4-c | 52 | 52 | 53 | 33 | 27 | 23 | 109 | 128 | 136 | 72 | 72 | 63 | 95.42 | 124 | 69 | 0.552 | -0.858 | 2.72E-05 |
| AT3G09060.1 | | protein_coding | Pentatricopeptide repeat | 23 | 20 | 18 | 19 | 16 | 17 | 48 | 49 | 46 | 41 | 42 | 46 | 45.7 | 48 | 43 | 0.551 | -0.859 | 0.0011856 |
| AT1G61360.1 | | protein_coding | S-locus lectin protein kina | 96 | 63 | 49 | 110 | 47 | 51 | 202 | 155 | 125 | 239 | 125 | 139 | 168.93 | 161 | 168 | 0.551 | -0.859 | 5.61E-05 |
| AT1G62260.1 | PCMP-E10 | protein_coding | mitochondrial editing fac | 69 | 38 | 50 | 42 | 27 | 35 | 145 | 93 | 128 | 91 | 72 | 95 | 91.92 | 122 | 86 | 0.551 | -0.859 | 4.20E-05 |
| AT1G25360.1 | PCMP-H74 | protein_coding | Pentatricopeptide repeat | 30 | 30 | 20 | 28 | 21 | 8 | 63 | 74 | 51 | 61 | 56 | 22 | 64.01 | 63 | 46 | 0.551 | -0.86 | 0.00028564 |
| AT1G02350.1 | | protein_coding | protoporphyrinogen oxid | 27 | 45 | 39 | 25 | 23 | 36 | 57 | 111 | 100 | 54 | 61 | 98 | 102.69 | 89 | 71 | 0.551 | -0.86 | 7.84E-05 |
| AT4G17030.1 | EXLB1 | protein_coding | expansin-like B1 | 51 | 43 | 32 | 38 | 36 | 22 | 107 | 106 | 82 | 82 | 96 | 60 | 114.14 | 98 | 79 | 0.551 | -0.86 | 1.77E-06 |
| AT3G27340.1 | | protein_coding | | 232 | 173 | 203 | 151 | 138 | 152 | 488 | 425 | 520 | 328 | 366 | 413 | 451.73 | 478 | 369 | 0.551 | -0.861 | 3.21E-11 |
| AT2G21770.1 | CESA9 | protein_coding | cellulose synthase A9 | 43 | 27 | 29 | 19 | 15 | 11 | 90 | 66 | 74 | 41 | 40 | 30 | 49.18 | 77 | 37 | 0.55 | -0.862 | 0.00056705 |
| AT3G09390.1 | MT2A | protein_coding | metallothionein 2A | 1127 | 1001 | 1174 | 1024 | 555 | 955 | 2369 | 2459 | 3006 | 2221 | 1473 | 2597 | 3154.06 | 2611 | 2097 | 0.55 | -0.862 | 1.32E-06 |
| AT1G71060.1 | | protein_coding | Tetratricopeptide repeat | 94 | 60 | 76 | 54 | 42 | 43 | 198 | 147 | 195 | 117 | 112 | 117 | 134.98 | 180 | 115 | 0.55 | -0.863 | 1.31E-05 |
| AT2G04790.2 | | protein_coding | | 171 | 129 | 126 | 130 | 69 | 55 | 359 | 317 | 323 | 282 | 183 | 150 | 344.28 | 333 | 205 | 0.55 | -0.863 | 2.40E-07 |
| AT4G14560.1 | IAA1 | protein_coding | indole-3-acetic acid induc | 203 | 206 | 301 | 183 | 223 | 339 | 427 | 506 | 771 | 397 | 592 | 922 | 571.93 | 568 | 637 | 0.549 | -0.864 | 0.00017465 |
| AT2G30000.1 | | protein_coding | PHF5-like protein | 58 | 47 | 54 | 65 | 47 | 33 | 122 | 115 | 138 | 141 | 125 | 90 | 170.95 | 125 | 119 | 0.549 | -0.864 | 3.94E-08 |
| AT4G14350.1 | | protein_coding | AGC (cAMP-dependent, c | 629 | 462 | 545 | 585 | 461 | 443 | 1322 | 1135 | 1395 | 1269 | 1224 | 1205 | 1355.58 | 1284 | 1233 | 0.549 | -0.864 | 5.44E-13 |
| AT3G57360.1 | | protein_coding | | 38 | 21 | 19 | 23 | 18 | 10 | 80 | 52 | 49 | 50 | 48 | 27 | 59.66 | 60 | 42 | 0.549 | -0.865 | 0.00141103 |
| AT5G02190.1 | PCS1 | protein_coding | Eukaryotic aspartyl prote | 53 | 59 | 33 | 38 | 32 | 26 | 111 | 145 | 84 | 82 | 85 | 71 | 197.43 | 113 | 79 | 0.548 | -0.866 | 2.02E-06 |
| AT5G66500.1 | PCMP-E38 | protein_coding | Tetratricopeptide repeat | 28 | 20 | 20 | 20 | 21 | 16 | 59 | 49 | 51 | 43 | 56 | 44 | 54.08 | 53 | 48 | 0.548 | -0.867 | 0.00014173 |
| AT5G65740.2 | | protein_coding | zinc ion binding | 184 | 126 | 135 | 87 | 78 | 85 | 387 | 310 | 346 | 189 | 207 | 231 | 283.57 | 348 | 209 | 0.548 | -0.867 | 1.02E-10 |
| AT2G37130.1 | PER21 | protein_coding | Peroxidase superfamily p | 194 | 206 | 141 | 183 | 125 | 133 | 408 | 506 | 361 | 397 | 332 | 362 | 509.66 | 425 | 364 | 0.548 | -0.868 | 0.00162582 |
| AT2G43940.1 | HOL3 | protein_coding | S-adenosyl-L-methionine- | 82 | 58 | 66 | 59 | 31 | 41 | 172 | 142 | 169 | 128 | 82 | 111 | 121.03 | 161 | 107 | 0.548 | -0.869 | 1.57E-05 |
| AT3G27997.1 | | pseudogene | | 23 | 16 | 21 | 14 | 10 | 10 | 48 | 39 | 54 | 30 | 27 | 27 | 49.99 | 47 | 28 | 0.547 | -0.87 | 0.00247395 |
| AT2G35035.1 | URED | protein_coding | urease accessory protein | 27 | 17 | 20 | 17 | 10 | 10 | 57 | 42 | 51 | 37 | 27 | 27 | 46.27 | 50 | 30 | 0.547 | -0.87 | 0.0012258 |
| AT5G18850.1 | | protein_coding | | 66 | 95 | 79 | 54 | 46 | 58 | 139 | 233 | 202 | 117 | 122 | 158 | 223.3 | 191 | 132 | 0.547 | -0.871 | 4.10E-06 |
| AT5G61940.1 | | protein_coding | Ubiquitin carboxyl-termir | 69 | 58 | 54 | 56 | 35 | 28 | 145 | 142 | 138 | 121 | 93 | 76 | 111 | 142 | 97 | 0.546 | -0.874 | 1.41E-05 |
| AT4G15130.1 | CCT2 | protein_coding | phosphorylcholine cytidyl | 308 | 199 | 218 | 215 | 188 | 195 | 647 | 489 | 558 | 466 | 499 | 530 | 595.01 | 565 | 498 | 0.546 | -0.874 | 3.74E-11 |
| AT2G45250.2 | | protein_coding | Integral membrane prote | 216 | 191 | 178 | 109 | 112 | 122 | 454 | 469 | 456 | 236 | 297 | 332 | 397.23 | 460 | 288 | 0.546 | -0.874 | 2.55E-13 |
| AT1G67970.1 | HSFA8 | protein_coding | heat shock transcription f | 52 | 77 | 78 | 38 | 18 | 31 | 109 | 189 | 200 | 82 | 48 | 84 | 127.34 | 166 | 71 | 0.545 | -0.875 | 0.00021657 |
| AT4G14290.1 | | protein_coding | alpha/beta-Hydrolases su | 85 | 87 | 53 | 91 | 88 | 74 | 179 | 214 | 136 | 197 | 234 | 201 | 197.12 | 176 | 211 | 0.545 | -0.875 | 8.42E-08 |
| AT1G52640.1 | | protein_coding | Pentatricopeptide repeat | 16 | 15 | 13 | 18 | 16 | 11 | 34 | 37 | 33 | 39 | 42 | 30 | 35.7 | 35 | 37 | 0.545 | -0.876 | 0.00162726 |
| AT1G09190.1 | PCMP-E70 | protein_coding | Tetratricopeptide repeat | 37 | 16 | 24 | 25 | 31 | 17 | 78 | 39 | 61 | 54 | 82 | 46 | 88.12 | 59 | 61 | 0.545 | -0.876 | 1.46E-05 |
| AT2G23560.1 | MES7 | protein_coding | methyl esterase 7 | 44 | 36 | 25 | 30 | 4 | 13 | 92 | 88 | 64 | 65 | 11 | 35 | 74.7 | 81 | 37 | 0.545 | -0.877 | 0.00609221 |
| AT1G05750.1 | PDE247 | protein_coding | Tetratricopeptide repeat | 70 | 65 | 70 | 52 | 49 | 41 | 147 | 160 | 179 | 113 | 130 | 111 | 160.55 | 162 | 118 | 0.544 | -0.877 | 1.07E-09 |
| AT2G40085.1 | | protein_coding | | 8 | 16 | 14 | 5 | 6 | 6 | 17 | 39 | 36 | 11 | 16 | 16 | 36.2 | 31 | 14 | 0.544 | -0.879 | 0.00067585 |
| AT4G14020.1 | | protein_coding | Rapid alkalinization facto | 165 | 144 | 123 | 220 | 160 | 165 | 347 | 354 | 315 | 477 | 425 | 449 | 528.87 | 339 | 450 | 0.544 | -0.879 | 2.80E-12 |
| AT5G65205.1 | | protein_coding | NAD(P)-binding Rossman | 63 | 67 | 52 | 30 | 40 | 33 | 132 | 165 | 133 | 65 | 106 | 90 | 130.67 | 143 | 87 | 0.543 | -0.88 | 1.85E-07 |
| AT4G14455.1 | BET12 | protein_coding | Target SNARE coiled-coil | 188 | 154 | 181 | 195 | 137 | 150 | 395 | 378 | 463 | 423 | 364 | 408 | 487.9 | 412 | 398 | 0.543 | -0.88 | 7.07E-14 |
| AT2G15630.1 | | protein_coding | Pentatricopeptide repeat | 37 | 22 | 24 | 34 | 28 | 13 | 78 | 54 | 61 | 74 | 74 | 35 | 62.65 | 64 | 61 | 0.543 | -0.881 | 0.00026288 |
| AT3G53170.1 | | protein_coding | Tetratricopeptide repeat | 125 | 89 | 85 | 53 | 44 | 37 | 263 | 219 | 218 | 115 | 117 | 101 | 174.06 | 233 | 111 | 0.543 | -0.881 | 3.18E-08 |
| AT4G14990.1 | | protein_coding | Topoisomerase II-associat | 407 | 357 | 314 | 354 | 328 | 347 | 856 | 877 | 804 | 768 | 871 | 944 | 1034.39 | 846 | 861 | 0.543 | -0.881 | 1.74E-14 |
| AT3G59250.1 | | protein_coding | F-box/RNI-like superfamil | 109 | 95 | 77 | 32 | 42 | 32 | 229 | 233 | 197 | 69 | 112 | 87 | 185.6 | 220 | 89 | 0.542 | -0.882 | 4.38E-06 |
| AT1G05805.1 | BHLH128 | protein_coding | basic helix-loop-helix (bH | 325 | 470 | 403 | 217 | 214 | 344 | 683 | 1155 | 1032 | 471 | 568 | 935 | 1078.3 | 957 | 658 | 0.542 | -0.883 | 1.28E-05 |
| AT1G53903.1 | | protein_coding | Protein of unknown funct | 9 | 17 | 28 | 13 | 19 | 23 | 19 | 42 | 72 | 28 | 50 | 63 | 61.11 | 44 | 47 | 0.542 | -0.884 | 0.00296137 |
| AT4G17440.1 | | protein_coding | Protein of unknown funct | 14 | 20 | 10 | 8 | 11 | 14 | 29 | 49 | 26 | 17 | 29 | 38 | 51.95 | 35 | 28 | 0.542 | -0.884 | 0.00019778 |
| AT5G26040.2 | HDA2 | protein_coding | histone deacetylase 2 | 122 | 134 | 172 | 46 | 64 | 62 | 256 | 329 | 440 | 100 | 170 | 169 | 254.6 | 342 | 146 | 0.542 | -0.884 | 2.02E-06 |
| AT1G70650.1 | | protein_coding | Ran BP2/NZF zinc finger-l | 35 | 19 | 33 | 17 | 19 | 15 | 74 | 47 | 84 | 37 | 50 | 41 | 57.19 | 68 | 43 | 0.541 | -0.885 | 0.00152671 |
| AT2G29070.2 | | protein_coding | Ubiquitin fusion degradat | 66 | 52 | 72 | 66 | 41 | 36 | 139 | 128 | 184 | 143 | 109 | 98 | 133.84 | 150 | 117 | 0.542 | -0.885 | 2.78E-06 |
| AT4G13730.1 | | protein_coding | Ypt/Rab-GAP domain of g | 248 | 236 | 213 | 260 | 231 | 180 | 521 | 580 | 545 | 564 | 613 | 489 | 815.71 | 549 | 555 | 0.541 | -0.886 | 2.58E-15 |
| AT5G15340.1 | PCMP-H91 | protein_coding | Pentatricopeptide repeat | 20 | 17 | 9 | 10 | 10 | 9 | 42 | 42 | 23 | 22 | 27 | 24 | 33.68 | 36 | 24 | 0.541 | -0.887 | 0.00221052 |
| AT3G55390.1 | | protein_coding | Uncharacterised protein f | 11 | 8 | 9 | 15 | 7 | 7 | 23 | 20 | 23 | 33 | 19 | 19 | 69.99 | 22 | 24 | 0.541 | -0.887 | 4.31E-05 |
| AT5G49170.1 | | protein_coding | | 50 | 71 | 65 | 48 | 49 | 66 | 105 | 174 | 166 | 104 | 130 | 179 | 124.59 | 148 | 138 | 0.541 | -0.887 | 3.51E-05 |
| AT4G15410.1 | PUX5 | protein_coding | serine/threonine protein | 146 | 167 | 121 | 171 | 176 | 167 | 307 | 410 | 310 | 371 | 467 | 454 | 642.68 | 342 | 431 | 0.541 | -0.887 | 2.72E-13 |
| AT5G65575.1 | | other_rna | other RNA | 58 | 64 | 59 | 35 | 28 | 31 | 122 | 157 | 151 | 76 | 74 | 84 | 128.92 | 143 | 78 | 0.54 | -0.888 | 3.17E-07 |
| AT4G13590.1 | | protein_coding | Uncharacterized protein f | 1397 | 1069 | 1070 | 1265 | 1054 | 1060 | 2937 | 2626 | 2739 | 2744 | 2798 | 2883 | 3781.98 | 2767 | 2808 | 0.54 | -0.889 | 1.53E-15 |
| AT4G14510.1 | ATCFM3B | protein_coding | CRM family member 3B | 319 | 213 | 181 | 206 | 192 | 140 | 671 | 523 | 463 | 447 | 510 | 381 | 1028.84 | 552 | 446 | 0.539 | -0.89 | 4.06E-11 |
| AT4G29560.1 | | protein_coding | | 38 | 33 | 30 | 27 | 23 | 19 | 80 | 81 | 77 | 59 | 61 | 52 | 73.35 | 79 | 57 | 0.539 | -0.891 | 4.18E-05 |
| AT4G15110.1 | CYP97B3 | protein_coding | cytochrome P450, family | 1848 | 1552 | 1374 | 1597 | 1404 | 1336 | 3885 | 3813 | 3518 | 3464 | 3727 | 3633 | 5097.58 | 3739 | 3608 | 0.539 | -0.891 | 3.32E-16 |
| AT1G68930.1 | PCMP-H22 | protein_coding | pentatricopeptide (PPR) r | 41 | 30 | 25 | 36 | 27 | 14 | 86 | 74 | 64 | 78 | 72 | 38 | 62.83 | 75 | 63 | 0.539 | -0.892 | 0.00012954 |
| AT4G14590.1 | emb2739 | protein_coding | embryo defective 2739 | 72 | 45 | 45 | 62 | 39 | 39 | 151 | 111 | 115 | 134 | 104 | 106 | 101.1 | 126 | 115 | 0.539 | -0.892 | 5.03E-06 |
| AT5G37570.1 | PCMP-E37 | protein_coding | Pentatricopeptide repeat | 42 | 48 | 34 | 29 | 28 | 18 | 88 | 118 | 87 | 63 | 74 | 49 | 78.07 | 98 | 62 | 0.539 | -0.893 | 0.00025381 |
| AT5G18260.1 | | protein_coding | RING/U-box superfamily | 95 | 84 | 77 | 70 | 54 | 47 | 200 | 206 | 197 | 152 | 143 | 128 | 158.44 | 201 | 141 | 0.538 | -0.893 | 4.54E-09 |
| AT1G56690.1 | PCMP-H69 | protein_coding | Pentatricopeptide repeat | 34 | 10 | 17 | 22 | 12 | 18 | 71 | 25 | 44 | 48 | 32 | 49 | 61.7 | 47 | 43 | 0.538 | -0.894 | 0.00057432 |
| AT3G02330.1 | PCMP-E90 | protein_coding | Pentatricopeptide repeat | 80 | 43 | 45 | 36 | 29 | 27 | 168 | 106 | 115 | 78 | 77 | 73 | 92.23 | 130 | 76 | 0.538 | -0.894 | 0.00029988 |
| AT3G22900.1 | NRPD7 | protein_coding | RNA polymerase Rpb7-lik | 40 | 27 | 39 | 35 | 31 | 23 | 84 | 66 | 100 | 76 | 82 | 63 | 79.2 | 83 | 74 | 0.537 | -0.896 | 4.75E-05 |
| AT4G12700.1 | | protein_coding | | 214 | 144 | 188 | 205 | 174 | 158 | 450 | 354 | 481 | 445 | 462 | 430 | 472.44 | 428 | 446 | 0.537 | -0.896 | 5.10E-14 |
| AT1G74088.2 | | protein_coding | | 30 | 23 | 25 | 13 | 15 | 22 | 63 | 57 | 64 | 28 | 40 | 60 | 45.2 | 61 | 43 | 0.537 | -0.897 | 0.00159259 |
| AT4G14420.1 | | protein_coding | HR-like lesion-inducing pr | 913 | 571 | 576 | 917 | 669 | 599 | 1919 | 1403 | 1475 | 1989 | 1776 | 1629 | 2286.33 | 1599 | 1798 | 0.537 | -0.897 | 1.26E-11 |
| AT4G14950.1 | KMS1 | protein_coding | SNARE associated Golgi p | 323 | 262 | 248 | 352 | 288 | 257 | 679 | 644 | 635 | 764 | 765 | 699 | 802.21 | 653 | 743 | 0.537 | -0.897 | 8.95E-15 |
| AT1G74660.1 | MIF1 | protein_coding | mini zinc finger 1 | 15 | 18 | 21 | 15 | 14 | 13 | 32 | 44 | 54 | 33 | 37 | 35 | 41.8 | 43 | 35 | 0.537 | -0.898 | 0.00035725 |
| AT1G27020.1 | | protein_coding | | 1051 | 275 | 327 | 564 | 232 | 292 | 2209 | 676 | 837 | 1223 | 616 | 794 | 1035.87 | 1241 | 878 | 0.536 | -0.899 | 0.00379942 |
| AT1G05820.1 | SPPL5 | protein_coding | SIGNAL PEPTIDE PEPTIDA | 62 | 63 | 38 | 38 | 36 | 19 | 130 | 155 | 97 | 82 | 96 | 52 | 107.14 | 127 | 77 | 0.536 | -0.899 | 5.35E-06 |
| AT4G33985.1 | | protein_coding | Protein of unknown funct | 12 | 13 | 14 | 6 | 6 | 12 | 25 | 32 | 36 | 13 | 16 | 33 | 41.38 | 31 | 21 | 0.536 | -0.9 | 0.00070986 |
| AT2G02360.1 | PP2B10 | protein_coding | phloem protein 2-B10 | 107 | 120 | 122 | 55 | 53 | 50 | 225 | 295 | 312 | 119 | 141 | 136 | 245.52 | 277 | 132 | 0.536 | -0.9 | 4.87E-10 |
| AT2G30420.1 | ETC2 | protein_coding | Homeodomain-like super | 29 | 34 | 34 | 15 | 23 | 20 | 61 | 84 | 87 | 33 | 61 | 54 | 58.88 | 77 | 49 | 0.536 | -0.901 | 0.00088184 |
| AT3G49170.1 | EMB2261 | protein_coding | Tetratricopeptide repeat | 173 | 99 | 87 | 82 | 69 | 51 | 364 | 243 | 223 | 178 | 183 | 139 | 256.13 | 277 | 167 | 0.535 | -0.902 | 1.63E-07 |
| AT5G14565.1 | MIR398C | mirna | MIR398C; miRNA | 212 | 170 | 140 | 136 | 105 | 119 | 446 | 418 | 358 | 295 | 279 | 324 | 463.08 | 407 | 299 | 0.535 | -0.902 | 3.00E-11 |
| AT2G13600.1 | PCMP-E76 | protein_coding | Pentatricopeptide repeat | 84 | 45 | 55 | 47 | 52 | 33 | 177 | 111 | 141 | 102 | 138 | 90 | 125.18 | 143 | 110 | 0.535 | -0.903 | 8.58E-06 |
| AT2G12400.1 | | protein_coding | | 261 | 251 | 289 | 141 | 131 | 129 | 549 | 617 | 740 | 306 | 348 | 351 | 472.05 | 635 | 335 | 0.534 | -0.904 | 2.16E-10 |
| AT1G64170.1 | CHX16 | protein_coding | cation/H+ exchanger 16 | 130 | 81 | 43 | 82 | 40 | 33 | 273 | 199 | 110 | 178 | 106 | 90 | 175.57 | 194 | 125 | 0.534 | -0.906 | 0.00035197 |
| AT4G14605.1 | | protein_coding | Mitochondrial transcripti | 363 | 305 | 304 | 325 | 309 | 225 | 763 | 749 | 778 | 705 | 820 | 612 | 918.16 | 763 | 712 | 0.534 | -0.906 | 4.10E-14 |
| AT2G21130.1 | CYP19-2 | protein_coding | Cyclophilin-like peptidyl-p | 85 | 175 | 122 | 43 | 63 | 72 | 179 | 430 | 312 | 93 | 167 | 196 | 432.06 | 307 | 152 | 0.533 | -0.907 | 6.85E-05 |
| AT3G62000.2 | | protein_coding | S-adenosyl-L-methionine- | 200 | 174 | 142 | 97 | 95 | 90 | 420 | 427 | 364 | 210 | 252 | 245 | 335.15 | 404 | 236 | 0.533 | -0.907 | 3.83E-12 |
| AT4G35480.1 | ATL45 | protein_coding | RING-H2 finger A3B | 7 | 17 | 8 | 3 | 9 | 7 | 15 | 42 | 20 | 7 | 24 | 19 | 114.2 | 26 | 17 | 0.533 | -0.908 | 4.10E-05 |
| AT2G36730.1 | PCMP-E44 | protein_coding | Pentatricopeptide repeat | 34 | 20 | 28 | 22 | 15 | 10 | 71 | 49 | 72 | 48 | 40 | 27 | 91.39 | 64 | 38 | 0.533 | -0.908 | 8.77E-06 |
| AT3G22235.1 | | protein_coding | | 5 | 4 | 1 | 13 | 2 | 7 | 11 | 10 | 3 | 28 | 5 | 19 | 36.91 | 8 | 17 | 0.533 | -0.909 | 0.00035465 |
| AT2G11150.1 | | transposable_elen | transposable element ger | 19 | 16 | 19 | 15 | 6 | 9 | 40 | 39 | 49 | 33 | 16 | 24 | 45.41 | 43 | 24 | 0.533 | -0.909 | 0.00027189 |
| AT5G03350.1 | | protein_coding | Legume lectin family prot | 36 | 34 | 27 | 54 | 51 | 43 | 76 | 84 | 69 | 117 | 135 | 117 | 139.81 | 76 | 123 | 0.533 | -0.909 | 3.14E-09 |
| AT5G28460.1 | | protein_coding | Pentatricopeptide repeat | 188 | 115 | 129 | 126 | 84 | 78 | 395 | 283 | 330 | 273 | 223 | 212 | 270.71 | 336 | 236 | 0.533 | -0.909 | 8.48E-10 |
| AT4G13840.1 | CER26 | protein_coding | HXXXD-type acyl-transfer | 351 | 426 | 274 | 383 | 273 | 291 | 738 | 1047 | 701 | 831 | 725 | 791 | 2436.39 | 829 | 782 | 0.533 | -0.909 | 6.77E-11 |
| AT1G18773.1 | | protein_coding | | 42 | 41 | 44 | 25 | 25 | 23 | 88 | 101 | 113 | 54 | 66 | 63 | 100.69 | 101 | 61 | 0.532 | -0.91 | 2.42E-07 |
| AT1G51610.1 | MTPC4 | protein_coding | Cation efflux family prote | 885 | 870 | 779 | 396 | 380 | 354 | 1860 | 2137 | 1994 | 859 | 1009 | 963 | 1816.16 | 1997 | 944 | 0.532 | -0.91 | 3.35E-13 |
| AT2G21660.1 | RBG7 | protein_coding | cold, circadian rhythm, ar | 694 | 1220 | 1003 | 268 | 299 | 509 | 1459 | 2997 | 2568 | 581 | 794 | 1384 | 14772.98 | 2341 | 920 | 0.532 | -0.911 | 0.00033878 |
| AT1G28190.1 | | protein_coding | | 77 | 63 | 71 | 32 | 50 | 37 | 162 | 155 | 182 | 69 | 133 | 101 | 111.06 | 166 | 101 | 0.531 | -0.912 | 0.00035273 |
| AT4G13260.1 | YUC2 | protein_coding | Flavin-binding monooxyg | 36 | 69 | 92 | 48 | 54 | 100 | 76 | 170 | 236 | 104 | 143 | 272 | 162.57 | 161 | 173 | 0.531 | -0.913 | 0.0034724 |
| AT4G15570.1 | MAA3 | protein_coding | P-loop containing nucleos | 322 | 185 | 207 | 230 | 179 | 178 | 677 | 455 | 530 | 499 | 475 | 484 | 548.93 | 554 | 486 | 0.531 | -0.914 | 7.71E-11 |
| AT4G13640.2 | UNE16 | protein_coding | Homeodomain-like super | 227 | 280 | 258 | 220 | 252 | 329 | 477 | 688 | 660 | 477 | 669 | 895 | 874.19 | 608 | 680 | 0.53 | -0.915 | 1.38E-07 |
| AT5G39410.1 | | protein_coding | Saccharopine dehydroger | 324 | 253 | 274 | 152 | 101 | 139 | 681 | 622 | 701 | 330 | 268 | 378 | 572.99 | 668 | 325 | 0.529 | -0.92 | 8.69E-12 |
| AT3G28850.1 | | protein_coding | Glutaredoxin family prote | 22 | 26 | 22 | 10 | 13 | 15 | 46 | 64 | 56 | 22 | 35 | 41 | 42.81 | 55 | 33 | 0.528 | -0.921 | 0.00087102 |
| AT3G16280.1 | ERF036 | protein_coding | Integrase-type DNA-bindi | 23 | 15 | 19 | 17 | 24 | 14 | 48 | 37 | 49 | 37 | 64 | 38 | 43.02 | 45 | 46 | 0.528 | -0.921 | 0.00045574 |
| AT1G14480.1 | | protein_coding | Ankyrin repeat family pro | 27 | 18 | 21 | 28 | 20 | 10 | 57 | 44 | 54 | 61 | 53 | 27 | 69.95 | 52 | 47 | 0.528 | -0.921 | 0.00023585 |
| AT4G33490.2 | | protein_coding | Eukaryotic aspartyl prote | 236 | 219 | 302 | 168 | 179 | 211 | 496 | 538 | 773 | 364 | 475 | 574 | 490.93 | 602 | 471 | 0.528 | -0.921 | 1.53E-06 |
| AT3G05870.1 | APC11 | protein_coding | anaphase-promoting cor | 47 | 49 | 48 | 43 | 24 | 26 | 99 | 120 | 123 | 93 | 64 | 71 | 115.29 | 114 | 76 | 0.528 | -0.921 | 1.95E-08 |
| AT2G43350.1 | GPX3 | protein_coding | glutathione peroxidase 3 | 443 | 238 | 307 | 240 | 164 | 132 | 931 | 585 | 786 | 521 | 435 | 359 | 534.43 | 767 | 438 | 0.528 | -0.922 | 4.00E-09 |
| AT4G14910.2 | HISN5B | protein_coding | HISTIDINE BIOSYNTHESIS | 423 | 288 | 307 | 370 | 265 | 288 | 889 | 708 | 786 | 803 | 704 | 783 | 1490.1 | 794 | 763 | 0.528 | -0.922 | 9.75E-15 |
| AT1G71420.1 | PCMP-H70 | protein_coding | Tetratricopeptide repeat | 30 | 30 | 30 | 23 | 29 | 17 | 63 | 74 | 77 | 50 | 77 | 46 | 50.66 | 71 | 58 | 0.527 | -0.923 | 0.00089025 |

| | | | | | | | | | | | | | | | | | | | | | |
|---|---|---|---|---|---|---|---|---|---|---|---|---|---|---|---|---|---|---|---|---|---|
| AT4G00200.1 | AHL7 | protein_coding | AT hook motif DNA-bindi | 14 | 26 | 22 | 20 | 20 | 17 | 29 | 64 | 56 | 43 | 53 | 46 | 56.3 | 50 | 47 | 0.527 | -0.923 | 0.00032411 |
| AT4G10000.1 | | protein_coding | Thioredoxin family protei | 272 | 290 | 272 | 130 | 134 | 135 | 572 | 712 | 696 | 282 | 356 | 367 | 551.75 | 660 | 335 | 0.527 | -0.924 | 6.07E-13 |
| AT1G14040.1 | PHO1 | protein_coding | EXS (ERD1/XPR1/SYG1) fa | 75 | 51 | 52 | 72 | 72 | 45 | 158 | 125 | 133 | 156 | 191 | 122 | 147.86 | 139 | 156 | 0.527 | -0.925 | 2.03E-05 |
| AT5G02560.2 | HTA12 | protein_coding | histone H2A 12 | 57 | 67 | 59 | 24 | 28 | 21 | 120 | 165 | 151 | 52 | 74 | 57 | 120.14 | 145 | 61 | 0.527 | -0.925 | 1.75E-05 |
| AT4G38010.1 | PCMP-E45 | protein_coding | Pentatricopeptide repeat | 27 | 17 | 29 | 20 | 15 | 14 | 57 | 42 | 74 | 43 | 40 | 38 | 54.83 | 58 | 40 | 0.526 | -0.926 | 4.09E-05 |
| AT4G12720.4 | NUDT7 | protein_coding | MutT/nudix family protei | 287 | 245 | 245 | 269 | 175 | 198 | 603 | 602 | 627 | 583 | 465 | 538 | 730.08 | 611 | 529 | 0.526 | -0.926 | 1.23E-15 |
| AT4G14310.2 | | protein_coding | Transducin/WD40 repeat | 496 | 430 | 416 | 447 | 476 | 376 | 1043 | 1056 | 1065 | 970 | 1264 | 1023 | 993.46 | 1055 | 1086 | 0.525 | -0.928 | 6.78E-14 |
| AT1G01225.1 | | protein_coding | NC domain-containing pr | 57 | 55 | 51 | 20 | 26 | 16 | 120 | 135 | 131 | 43 | 69 | 44 | 89.09 | 129 | 52 | 0.525 | -0.929 | 8.80E-07 |
| AT4G18930.1 | | protein_coding | RNA ligase/cyclic nucleot | 260 | 222 | 193 | 143 | 87 | 119 | 547 | 545 | 494 | 310 | 231 | 324 | 629.69 | 529 | 288 | 0.525 | -0.929 | 4.22E-16 |
| AT5G63690.1 | | protein_coding | Nucleic acid-binding, OB- | 6 | 4 | 6 | 5 | 6 | 11 | 13 | 10 | 15 | 11 | 16 | 30 | 29.96 | 13 | 19 | 0.525 | -0.93 | 0.00714976 |
| AT5G40405.1 | PCMP-H14 | protein_coding | Tetratricopeptide repeat | 46 | 48 | 23 | 37 | 26 | 26 | 97 | 118 | 59 | 80 | 69 | 71 | 83.83 | 91 | 73 | 0.525 | -0.93 | 8.21E-05 |
| AT4G14200.1 | | protein_coding | Pentatricopeptide repeat | 290 | 289 | 250 | 351 | 351 | 235 | 610 | 710 | 640 | 761 | 932 | 639 | 650.81 | 653 | 777 | 0.525 | -0.931 | 7.18E-13 |
| AT3G28640.1 | PCMP-E79 | protein_coding | Tetratricopeptide repeat | 35 | 31 | 30 | 23 | 25 | 16 | 74 | 76 | 77 | 50 | 66 | 44 | 66.21 | 76 | 53 | 0.524 | -0.932 | 7.06E-05 |
| AT5G42690.2 | | protein_coding | Protein of unknown funct | 36 | 14 | 28 | 15 | 15 | 15 | 76 | 34 | 72 | 33 | 40 | 41 | 43.5 | 61 | 38 | 0.524 | -0.933 | 0.00821464 |
| AT2G04110.1 | | pseudogene | | 98 | 116 | 112 | 55 | 61 | 66 | 206 | 285 | 287 | 119 | 162 | 179 | 231.2 | 259 | 153 | 0.523 | -0.934 | 3.86E-07 |
| AT4G13010.1 | | protein_coding | Oxidoreductase, zinc-binc | 1075 | 1067 | 963 | 1138 | 841 | 1091 | 2260 | 2621 | 2465 | 2468 | 2233 | 2967 | 3507.04 | 2449 | 2556 | 0.523 | -0.934 | 1.53E-13 |
| AT2G01890.1 | PAP8 | protein_coding | purple acid phosphatase | 345 | 589 | 313 | 207 | 346 | 186 | 725 | 1447 | 801 | 449 | 919 | 506 | 709.14 | 991 | 625 | 0.523 | -0.935 | 0.00016714 |
| AT3G55450.2 | PBL1 | protein_coding | PBS1-like 1 | 259 | 271 | 262 | 147 | 121 | 178 | 544 | 666 | 671 | 319 | 321 | 484 | 485.72 | 627 | 375 | 0.523 | -0.935 | 8.79E-11 |
| AT5G15170.1 | TDP1 | protein_coding | tyrosyl-DNA phosphodies | 187 | 180 | 180 | 103 | 100 | 96 | 393 | 442 | 461 | 223 | 265 | 261 | 288.66 | 432 | 250 | 0.523 | -0.935 | 1.04E-12 |
| AT3G09780.1 | CCR1 | protein_coding | CRINKLY4 related 1 | 73 | 77 | 84 | 38 | 44 | 62 | 153 | 189 | 215 | 82 | 117 | 169 | 149.96 | 186 | 123 | 0.523 | -0.936 | 1.44E-05 |
| AT2G37310.1 | PCMP-E49 | protein_coding | Pentatricopeptide repeat | 74 | 49 | 61 | 57 | 44 | 53 | 156 | 120 | 156 | 124 | 117 | 144 | 149.33 | 144 | 128 | 0.523 | -0.936 | 1.21E-08 |
| AT5G50110.1 | | protein_coding | S-adenosyl-L-methionine- | 130 | 93 | 96 | 84 | 70 | 79 | 273 | 228 | 246 | 182 | 186 | 215 | 287.92 | 249 | 194 | 0.522 | -0.937 | 4.78E-15 |
| AT4G04360.1 | | protein_coding | Protein of unknown funct | 50 | 33 | 51 | 25 | 26 | 30 | 105 | 81 | 131 | 54 | 69 | 82 | 98.57 | 106 | 68 | 0.522 | -0.938 | 3.53E-07 |
| AT1G35260.1 | MLP165 | protein_coding | MLP-like protein 165 | 14 | 10 | 3 | 13 | 11 | 11 | 29 | 25 | 8 | 28 | 29 | 30 | 37.01 | 21 | 29 | 0.522 | -0.939 | 0.00903733 |
| AT3G51340.1 | | protein_coding | Eukaryotic aspartyl prote | 19 | 16 | 9 | 22 | 13 | 13 | 40 | 39 | 23 | 48 | 35 | 35 | 45.18 | 34 | 39 | 0.521 | -0.941 | 0.0008207 |
| AT2G40110.1 | | protein_coding | Yippee family putative zir | 89 | 71 | 88 | 53 | 58 | 64 | 187 | 174 | 225 | 115 | 154 | 174 | 170.8 | 195 | 148 | 0.521 | -0.941 | 4.03E-07 |
| AT5G27930.1 | PPC6-7 | protein_coding | Protein phosphatase 2C f | 150 | 155 | 130 | 88 | 96 | 61 | 315 | 381 | 333 | 191 | 255 | 166 | 321.74 | 343 | 204 | 0.521 | -0.941 | 2.82E-11 |
| AT5G35410.1 | CIPK24 | protein_coding | Protein kinase superfamil | 129 | 91 | 100 | 51 | 45 | 54 | 271 | 224 | 256 | 111 | 119 | 147 | 165.99 | 250 | 126 | 0.52 | -0.942 | 4.31E-09 |
| AT1G49140.1 | | protein_coding | Complex I subunit NDUFS | 405 | 427 | 394 | 348 | 244 | 344 | 851 | 1049 | 1009 | 755 | 648 | 935 | 1178.58 | 970 | 779 | 0.521 | -0.942 | 9.74E-12 |
| AT1G13970.1 | | protein_coding | Protein of unknown funct | 52 | 33 | 29 | 38 | 31 | 21 | 109 | 81 | 74 | 82 | 82 | 57 | 64.41 | 88 | 74 | 0.52 | -0.944 | 0.00016969 |
| AT2G06925.1 | PLA2-ALPHA | protein_coding | Phospholipase A2 family | 451 | 446 | 473 | 219 | 175 | 177 | 948 | 1096 | 1211 | 475 | 465 | 481 | 1128.33 | 1085 | 474 | 0.52 | -0.944 | 4.94E-12 |
| AT4G21380.1 | SD18 | protein_coding | receptor kinase 3 | 60 | 48 | 42 | 50 | 34 | 26 | 126 | 118 | 108 | 108 | 90 | 71 | 88.11 | 117 | 90 | 0.519 | -0.946 | 3.61E-05 |
| AT4G14520.1 | NRPB7L | protein_coding | DNA-directed RNA polym | 104 | 80 | 91 | 79 | 66 | 85 | 219 | 197 | 233 | 171 | 175 | 231 | 255.35 | 216 | 192 | 0.519 | -0.947 | 3.45E-10 |
| AT4G13630.1 | | protein_coding | Protein of unknown funct | 349 | 270 | 322 | 271 | 218 | 248 | 734 | 663 | 824 | 588 | 579 | 674 | 726.28 | 740 | 614 | 0.518 | -0.948 | 6.87E-16 |
| AT4G14160.1 | | protein_coding | Sec23/Sec24 protein tran | 1721 | 1338 | 1288 | 1630 | 1297 | 1346 | 3618 | 3287 | 3297 | 3536 | 3443 | 3660 | 3849.16 | 3401 | 3546 | 0.518 | -0.948 | 3.16E-16 |
| AT4G13040.3 | | protein_coding | Integrase-type DNA-bindi | 195 | 114 | 180 | 156 | 105 | 141 | 410 | 280 | 461 | 338 | 279 | 383 | 363.53 | 384 | 333 | 0.518 | -0.949 | 1.43E-08 |
| AT1G28140.1 | | protein_coding | | 928 | 870 | 839 | 419 | 398 | 454 | 1951 | 2137 | 2148 | 909 | 1057 | 1235 | 1762.21 | 2079 | 1067 | 0.518 | -0.949 | 8.85E-14 |
| AT3G18020.1 | | protein_coding | Pentatricopeptide repeat | 70 | 49 | 39 | 48 | 42 | 27 | 147 | 120 | 100 | 104 | 112 | 73 | 95.43 | 122 | 96 | 0.518 | -0.95 | 4.80E-06 |
| AT2G38070.1 | | protein_coding | Protein of unknown funct | 50 | 49 | 40 | 17 | 32 | 37 | 105 | 120 | 102 | 37 | 85 | 101 | 81.21 | 109 | 74 | 0.517 | -0.951 | 0.00899819 |
| AT3G14580.1 | | protein_coding | Pentatricopeptide repeat | 59 | 35 | 25 | 40 | 23 | 21 | 124 | 86 | 64 | 87 | 61 | 57 | 80.06 | 91 | 68 | 0.517 | -0.951 | 4.75E-05 |
| AT5G38730.1 | | protein_coding | Tetratricopeptide repeat | 54 | 39 | 39 | 31 | 37 | 23 | 114 | 96 | 100 | 67 | 98 | 63 | 78.65 | 103 | 76 | 0.517 | -0.952 | 2.49E-05 |
| AT4G14340.1 | CKI1 | protein_coding | casein kinase I | 360 | 314 | 393 | 336 | 308 | 329 | 757 | 771 | 1006 | 729 | 818 | 895 | 865.2 | 845 | 814 | 0.517 | -0.952 | 2.30E-13 |
| AT4G13980.1 | HSFA5 | protein_coding | winged-helix DNA-binding | 193 | 155 | 219 | 159 | 165 | 171 | 406 | 381 | 561 | 345 | 438 | 465 | 403.52 | 449 | 416 | 0.517 | -0.953 | 7.12E-11 |
| AT4G13495.1 | | other_rna | other RNA | 3858 | 3920 | 3452 | 2840 | 3177 | 2424 | 8110 | 9631 | 8837 | 6160 | 8434 | 6592 | 12285.36 | 8859 | 7062 | 0.517 | -0.953 | 1.96E-11 |
| AT2G27920.1 | SCPL51 | protein_coding | serine carboxypeptidase- | 244 | 200 | 171 | 105 | 134 | 89 | 513 | 491 | 438 | 228 | 356 | 242 | 478.18 | 481 | 275 | 0.516 | -0.954 | 3.30E-08 |
| AT5G51570.1 | HIR4 | protein_coding | SPFH/Band 7/PHB domai | 232 | 229 | 217 | 125 | 101 | 132 | 488 | 563 | 556 | 271 | 268 | 359 | 477.26 | 536 | 299 | 0.516 | -0.954 | 2.30E-12 |
| AT4G15470.1 | | protein_coding | Bax inhibitor-1 family pro | 1056 | 1016 | 990 | 1306 | 1108 | 1293 | 2220 | 2496 | 2534 | 2833 | 2941 | 3516 | 3738.2 | 2417 | 3097 | 0.515 | -0.956 | 1.11E-14 |
| AT5G59570.1 | BOA | protein_coding | Homeodomain-like super | 11 | 14 | 19 | 4 | 1 | 17 | 23 | 34 | 49 | 9 | 3 | 46 | 49.71 | 35 | 19 | 0.515 | -0.957 | 0.00227742 |
| AT1G53600.1 | PCMP-E63 | protein_coding | Tetratricopeptide repeat | 44 | 27 | 25 | 26 | 20 | 27 | 92 | 66 | 64 | 56 | 53 | 73 | 60.06 | 74 | 61 | 0.514 | -0.961 | 0.00051727 |
| AT1G34750.1 | | protein_coding | Protein phosphatase 2C f | 48 | 44 | 29 | 27 | 12 | 18 | 101 | 108 | 74 | 59 | 32 | 49 | 72.22 | 94 | 47 | 0.513 | -0.963 | 1.57E-05 |
| AT4G14385.1 | | protein_coding | | 145 | 130 | 148 | 148 | 115 | 138 | 305 | 319 | 379 | 321 | 305 | 375 | 462.09 | 334 | 334 | 0.513 | -0.964 | 2.19E-14 |
| AT4G14410.1 | BHLH104 | protein_coding | basic helix-loop-helix (bH | 343 | 292 | 277 | 335 | 242 | 287 | 721 | 717 | 709 | 727 | 642 | 780 | 1040.78 | 716 | 716 | 0.512 | -0.964 | 4.24E-17 |
| AT4G13770.1 | CYP83A1 | protein_coding | cytochrome P450, family | 3318 | 2907 | 1225 | 3071 | 2814 | 1900 | 6975 | 7142 | 3136 | 6661 | 7471 | 5167 | 8472.21 | 5751 | 6433 | 0.512 | -0.965 | 1.60E-05 |
| AT3G29120.1 | | transposable_elen | transposable element ger | 32 | 33 | 41 | 21 | 23 | 21 | 67 | 81 | 105 | 46 | 61 | 57 | 88.81 | 84 | 55 | 0.512 | -0.965 | 5.46E-07 |
| AT4G14650.1 | | protein_coding | | 39 | 34 | 34 | 44 | 38 | 38 | 82 | 84 | 87 | 95 | 101 | 103 | 99.86 | 84 | 100 | 0.512 | -0.965 | 3.96E-07 |
| AT4G14790.1 | SUV3 | protein_coding | ATP-dependent RNA helic | 212 | 171 | 198 | 192 | 183 | 140 | 446 | 420 | 507 | 416 | 486 | 381 | 532.5 | 458 | 428 | 0.512 | -0.966 | 9.82E-17 |
| AT1G30280.1 | | protein_coding | Chaperone DnaJ-domain | 119 | 186 | 216 | 98 | 100 | 86 | 250 | 457 | 553 | 213 | 265 | 234 | 391.4 | 420 | 237 | 0.512 | -0.967 | 6.15E-07 |
| AT4G13970.1 | | protein_coding | zinc ion binding | 361 | 289 | 322 | 335 | 279 | 274 | 759 | 710 | 824 | 727 | 741 | 745 | 730.4 | 764 | 738 | 0.512 | -0.967 | 4.50E-18 |
| AT4G23660.3 | PPT1 | protein_coding | polyprenyltransferase 1 | 129 | 100 | 122 | 66 | 49 | 38 | 271 | 246 | 312 | 143 | 130 | 103 | 191.98 | 276 | 125 | 0.511 | -0.968 | 1.58E-10 |
| AT5G26000.1 | TGG1 | protein_coding | thioglucoside glucohydro | 18854 | 26327 | 25617 | 14715 | 17541 | 16304 | 39632 | 64681 | 65581 | 31918 | 46567 | 44338 | 51955.79 | 56631 | 40941 | 0.51 | -0.97 | 3.79E-06 |
| AT4G14880.1 | OASA1 | protein_coding | O-acetylserine (thiol) lyas | 5665 | 4635 | 4315 | 5948 | 4492 | 4366 | 11908 | 11387 | 11047 | 12902 | 11925 | 11873 | 16869.53 | 11447 | 12233 | 0.51 | -0.971 | 6.92E-15 |
| AT4G10380.1 | NIP5-1 | protein_coding | NOD26-like intrinsic prot | 23 | 17 | 17 | 34 | 9 | 7 | 48 | 42 | 44 | 74 | 24 | 19 | 60.13 | 45 | 39 | 0.51 | -0.972 | 0.00102379 |
| AT5G14920.1 | GASA14 | protein_coding | Gibberellin-regulated fan | 239 | 234 | 242 | 187 | 119 | 315 | 502 | 575 | 620 | 406 | 316 | 857 | 2693.99 | 566 | 526 | 0.51 | -0.972 | 4.79E-05 |
| AT4G12710.1 | | protein_coding | ARM repeat superfamily | 87 | 84 | 73 | 80 | 64 | 91 | 183 | 206 | 187 | 174 | 170 | 247 | 228.02 | 192 | 197 | 0.51 | -0.972 | 5.87E-11 |
| AT4G40045.1 | | protein_coding | | 180 | 187 | 177 | 90 | 111 | 89 | 378 | 459 | 453 | 195 | 295 | 242 | 467.75 | 430 | 244 | 0.51 | -0.972 | 6.73E-13 |
| AT4G15415.2 | B'GAMMA | protein_coding | Protein phosphatase 2A r | 654 | 589 | 612 | 650 | 505 | 545 | 1375 | 1447 | 1567 | 1410 | 1341 | 1482 | 1483.07 | 1463 | 1411 | 0.509 | -0.973 | 4.48E-16 |
| AT5G53920.1 | | protein_coding | ribosomal protein L11 me | 67 | 47 | 47 | 57 | 30 | 30 | 141 | 115 | 120 | 124 | 80 | 82 | 139.03 | 125 | 95 | 0.509 | -0.975 | 5.53E-09 |
| AT4G13500.1 | | protein_coding | | 1366 | 1188 | 1156 | 1497 | 1137 | 953 | 2871 | 2919 | 2959 | 3247 | 3018 | 2592 | 4003.94 | 2916 | 2952 | 0.508 | -0.976 | 1.50E-15 |
| AT3G62810.1 | | protein_coding | complex 1 family protein | 121 | 93 | 69 | 66 | 41 | 56 | 254 | 228 | 177 | 143 | 109 | 152 | 287.53 | 220 | 135 | 0.507 | -0.979 | 1.45E-08 |
| AT4G14360.1 | | protein_coding | S-adenosyl-L-methionine- | 2044 | 1457 | 1479 | 1785 | 1366 | 1140 | 4297 | 3580 | 3786 | 3872 | 3626 | 3100 | 3739.17 | 3888 | 3533 | 0.507 | -0.979 | 4.45E-15 |
| AT3G26782.1 | PCMP-H34 | protein_coding | Tetratricopeptide repeat | 58 | 42 | 41 | 31 | 37 | 25 | 122 | 103 | 105 | 67 | 98 | 68 | 111.58 | 110 | 78 | 0.507 | -0.981 | 6.77E-08 |
| AT3G45020.1 | | protein_coding | Ribosomal L18p/L5e fami | 38 | 33 | 23 | 21 | 16 | 27 | 80 | 81 | 59 | 46 | 42 | 73 | 68.35 | 73 | 54 | 0.506 | -0.982 | 2.53E-05 |
| AT4G13990.1 | | protein_coding | Exostosin family protein | 91 | 65 | 62 | 66 | 74 | 46 | 191 | 160 | 159 | 143 | 196 | 125 | 175.12 | 170 | 155 | 0.506 | -0.982 | 1.69E-07 |
| AT4G12650.1 | TMN12 | protein_coding | Endomembrane protein 7 | 3242 | 2335 | 2404 | 3035 | 2339 | 1996 | 6815 | 5737 | 6154 | 6583 | 6210 | 5428 | 6114.75 | 6235 | 6074 | 0.506 | -0.982 | 1.54E-15 |
| AT3G09040.1 | PCMP-E88 | protein_coding | Pentatricopeptide repeat | 68 | 46 | 56 | 37 | 31 | 20 | 143 | 113 | 143 | 80 | 82 | 54 | 83.14 | 133 | 72 | 0.506 | -0.984 | 0.00013444 |
| AT5G20630.1 | GER3 | protein_coding | germin 3 | 1430 | 2449 | 1774 | 1477 | 1594 | 2069 | 3006 | 6017 | 4542 | 3204 | 4232 | 5627 | 9116.8 | 4522 | 4354 | 0.506 | -0.984 | 1.11E-05 |
| AT1G22570.1 | NPF5.15 | protein_coding | Major facilitator superfar | 448 | 387 | 316 | 174 | 92 | 137 | 942 | 951 | 809 | 377 | 244 | 373 | 682.78 | 901 | 331 | 0.505 | -0.985 | 5.86E-14 |
| AT1G12710.1 | P2A12 | protein_coding | phloem protein 2-A12 | 161 | 216 | 239 | 86 | 72 | 112 | 338 | 531 | 612 | 187 | 191 | 305 | 399.61 | 494 | 228 | 0.505 | -0.986 | 7.32E-07 |
| AT4G23330.1 | | protein_coding | | 75 | 86 | 61 | 49 | 47 | 41 | 158 | 211 | 156 | 106 | 125 | 111 | 200.03 | 175 | 114 | 0.504 | -0.988 | 8.61E-11 |
| AT4G14800.2 | PBD2 | protein_coding | 20S proteasome beta sub | 595 | 388 | 435 | 646 | 401 | 445 | 1251 | 953 | 1114 | 1401 | 1065 | 1210 | 1530.24 | 1106 | 1225 | 0.504 | -0.988 | 5.55E-14 |
| AT2G41210.1 | PIP5K5 | protein_coding | phosphatidylinositol- 4-p | 94 | 57 | 58 | 43 | 23 | 26 | 198 | 140 | 148 | 93 | 61 | 71 | 112.05 | 162 | 75 | 0.504 | -0.989 | 2.11E-05 |
| AT5G27280.1 | | protein_coding | Zim17-type zinc finger pr | 351 | 407 | 455 | 184 | 233 | 250 | 738 | 1000 | 1165 | 399 | 619 | 680 | 1047.65 | 968 | 566 | 0.503 | -0.99 | 1.14E-08 |
| AT3G58520.1 | | protein_coding | Ubiquitin carboxyl-termir | 59 | 36 | 25 | 32 | 35 | 25 | 124 | 88 | 64 | 69 | 93 | 68 | 124.68 | 92 | 77 | 0.503 | -0.991 | 4.10E-07 |
| AT5G48440.1 | | protein_coding | FAD-dependent oxidored | 134 | 114 | 125 | 65 | 56 | 51 | 282 | 280 | 320 | 141 | 149 | 139 | 240.87 | 294 | 143 | 0.503 | -0.991 | 3.37E-12 |
| AT4G14070.1 | AAE15 | protein_coding | acyl-activating enzyme 15 | 1406 | 1129 | 1059 | 1219 | 943 | 872 | 2956 | 2774 | 2711 | 2644 | 2503 | 2371 | 3359.64 | 2814 | 2506 | 0.503 | -0.991 | 1.19E-18 |
| AT1G10990.2 | | protein_coding | | 36 | 26 | 29 | 32 | 16 | 25 | 76 | 64 | 74 | 69 | 42 | 68 | 102.23 | 71 | 60 | 0.503 | -0.992 | 9.39E-08 |
| AT4G14103.2 | | protein_coding | F-box/RNI-like superfamil | 29 | 31 | 30 | 22 | 21 | 22 | 61 | 76 | 77 | 48 | 56 | 60 | 65.36 | 71 | 55 | 0.502 | -0.994 | 6.49E-06 |
| AT5G61370.1 | | protein_coding | Pentatricopeptide repeat | 67 | 50 | 49 | 54 | 42 | 27 | 141 | 123 | 125 | 117 | 112 | 73 | 127.42 | 130 | 101 | 0.502 | -0.995 | 3.03E-08 |
| AT2G29500.1 | HSP17.6B | protein_coding | HSP20-like chaperones su | 38 | 41 | 22 | 32 | 16 | 17 | 80 | 101 | 56 | 69 | 42 | 46 | 75.55 | 79 | 52 | 0.501 | -0.996 | 0.00011008 |
| AT3G13880.1 | PCMP-E89 | protein_coding | Tetratricopeptide repeat | 87 | 57 | 64 | 42 | 51 | 30 | 183 | 140 | 164 | 91 | 135 | 82 | 129.4 | 162 | 103 | 0.501 | -0.997 | 4.48E-07 |
| AT1G15220.1 | CCMH | protein_coding | cytochrome c biogenesis | 89 | 75 | 72 | 34 | 28 | 41 | 187 | 184 | 184 | 74 | 74 | 111 | 141.15 | 185 | 86 | 0.501 | -0.997 | 2.02E-07 |
| AT1G09910.1 | | protein_coding | Rhamnogalacturonate lya | 186 | 155 | 216 | 103 | 63 | 104 | 391 | 381 | 553 | 223 | 167 | 283 | 306.82 | 442 | 224 | 0.501 | -0.998 | 1.28E-07 |
| AT4G01130.1 | | protein_coding | GDSL-like Lipase/Acylhyd | 256 | 274 | 304 | 97 | 97 | 109 | 538 | 673 | 778 | 210 | 258 | 296 | 522.57 | 663 | 255 | 0.501 | -0.998 | 8.13E-11 |
| AT4G15510.1 | PPD1 | protein_coding | Photosystem II reaction c | 1048 | 906 | 714 | 838 | 753 | 612 | 2203 | 2226 | 1828 | 1818 | 1999 | 1664 | 2896.41 | 2086 | 1827 | 0.501 | -0.998 | 1.08E-15 |
| AT1G03850.1 | GRXS13 | protein_coding | Glutaredoxin family prote | 20 | 19 | 35 | 18 | 11 | 13 | 42 | 47 | 90 | 39 | 29 | 35 | 42.96 | 60 | 34 | 0.5 | -1 | 0.00235505 |
| AT4G26810.1 | | protein_coding | SWIB/MDM2 domain sup | 64 | 37 | 35 | 36 | 29 | 19 | 135 | 91 | 90 | 78 | 77 | 52 | 113.94 | 105 | 69 | 0.5 | -1 | 8.52E-08 |
| AT1G54217.2 | | protein_coding | Ribosomal protein L18ae | 27 | 34 | 30 | 23 | 24 | 20 | 57 | 84 | 77 | 50 | 64 | 54 | 94.54 | 73 | 56 | 0.5 | -1.001 | 1.65E-07 |
| AT1G11530.1 | CXXS1 | protein_coding | C-terminal cysteine resid | 231 | 240 | 414 | 124 | 134 | 204 | 486 | 590 | 1060 | 269 | 356 | 555 | 529.08 | 712 | 393 | 0.499 | -1.003 | 4.80E-05 |
| AT5G39350.1 | PCMP-E16 | protein_coding | Tetratricopeptide repeat | 46 | 26 | 23 | 27 | 28 | 12 | 97 | 64 | 59 | 59 | 74 | 33 | 109.57 | 73 | 55 | 0.498 | -1.006 | 4.49E-07 |
| AT1G53000.1 | KDSB | protein_coding | Nucleotide-diphospho-su | 415 | 305 | 357 | 242 | 162 | 192 | 872 | 749 | 914 | 525 | 430 | 522 | 717.79 | 845 | 492 | 0.498 | -1.006 | 9.76E-18 |
| AT5G46220.1 | | protein_coding | Protein of unknown funct | 78 | 66 | 26 | 36 | 38 | 12 | 164 | 162 | 67 | 78 | 101 | 33 | 130.75 | 131 | 71 | 0.497 | -1.008 | 0.00046268 |
| AT1G76720.1 | | protein_coding | eukaryotic translation ini | 81 | 67 | 87 | 43 | 50 | 41 | 170 | 165 | 223 | 93 | 133 | 111 | 146.94 | 186 | 112 | 0.497 | -1.009 | 7.83E-08 |
| AT4G14465.1 | AHL20 | protein_coding | AT-hook motif nuclear-lo | 14 | 20 | 17 | 38 | 20 | 25 | 29 | 49 | 44 | 82 | 53 | 68 | 82.05 | 41 | 68 | 0.497 | -1.01 | 0.00046941 |
| AT4G14713.1 | TIFY4A | protein_coding | TIFY domain/Divergent C | 59 | 78 | 68 | 47 | 51 | 77 | 124 | 192 | 174 | 102 | 135 | 209 | 220.51 | 163 | 149 | 0.496 | -1.011 | 6.50E-08 |
| AT2G34240.1 | | protein_coding | Protein with domains of | 19 | 17 | 14 | 16 | 18 | 13 | 40 | 42 | 36 | 35 | 48 | 35 | 37.27 | 39 | 39 | 0.496 | -1.012 | 0.00058194 |
| AT3G15130.1 | PCMP-H86 | protein_coding | Tetratricopeptide repeat | 39 | 25 | 23 | 18 | 20 | 20 | 82 | 61 | 59 | 39 | 53 | 54 | 62.74 | 67 | 49 | 0.496 | -1.012 | 7.72E-05 |
| AT4G13100.2 | | protein_coding | RING/U-box superfamily | 101 | 116 | 93 | 111 | 111 | 115 | 212 | 285 | 238 | 241 | 295 | 313 | 525.1 | 245 | 283 | 0.495 | -1.013 | 1.35E-18 |
| AT5G24470.1 | APRR5 | protein_coding | pseudo-response regulato | 616 | 1018 | 1387 | 210 | 291 | 699 | 1295 | 2501 | 3551 | 456 | 773 | 1901 | 1996.31 | 2449 | 1043 | 0.495 | -1.014 | 0.00643387 |
| AT4G13615.1 | | protein_coding | Uncharacterised protein f | 484 | 740 | 404 | 625 | 443 | 734 | 1017 | 1818 | 1034 | 1356 | 1176 | 1996 | 2872.18 | 1290 | 1509 | 0.494 | -1.017 | 6.07E-07 |
| AT4G14900.1 | FRL4B | protein_coding | FRIGIDA-like protein | 479 | 472 | 515 | 538 | 490 | 692 | 1007 | 1160 | 1318 | 1167 | 1301 | 1882 | 1595.48 | 1162 | 1450 | 0.494 | -1.017 | 1.22E-10 |
| AT5G04780.1 | PCMP-H16 | protein_coding | Pentatricopeptide repeat | 56 | 55 | 54 | 40 | 31 | 34 | 118 | 135 | 138 | 87 | 82 | 92 | 101.04 | 130 | 87 | 0.494 | -1.018 | 8.20E-07 |
| AT4G14240.1 | CBSDUF1 | protein_coding | CBS domain-containing p | 689 | 477 | 467 | 632 | 455 | 404 | 1448 | 1172 | 1196 | 1371 | 1208 | 1099 | 1491.38 | 1272 | 1226 | 0.494 | -1.018 | 2.96E-14 |
| AT4G08210.1 | PCMP-E100 | protein_coding | Pentatricopeptide repeat | 36 | 21 | 21 | 25 | 25 | 11 | 76 | 52 | 54 | 54 | 66 | 30 | 45.39 | 61 | 50 | 0.494 | -1.019 | 0.00052384 |
| AT3G62550.1 | | protein_coding | Adenine nucleotide alpha | 520 | 1126 | 1728 | 210 | 363 | 622 | 1093 | 2766 | 4424 | 456 | 964 | 1691 | 2088.94 | 2761 | 1037 | 0.493 | -1.02 | 0.00746505 |
| AT3G17668.1 | ENA | protein_coding | DnaJ/Hsp40 cysteine-rich | 69 | 62 | 59 | 45 | 31 | 41 | 145 | 152 | 151 | 98 | 82 | 111 | 129.77 | 149 | 97 | 0.493 | -1.022 | 1.74E-09 |
| AT5G62360.1 | | protein_coding | Plant invertase/pectin me | 286 | 352 | 272 | 107 | 117 | 85 | 601 | 865 | 696 | 232 | 311 | 231 | 852.04 | 721 | 258 | 0.492 | -1.023 | 6.34E-10 |
| AT1G50732.1 | | protein_coding | | 479 | 369 | 346 | 305 | 274 | 163 | 1007 | 907 | 886 | 662 | 727 | 443 | 731.27 | 933 | 611 | 0.492 | -1.023 | 1.81E-12 |
| AT3G23330.1 | PCMP-H32 | protein_coding | Tetratricopeptide repeat | 73 | 45 | 44 | 45 | 29 | 32 | 153 | 111 | 113 | 98 | 77 | 87 | 100.76 | 126 | 87 | 0.492 | -1.024 | 2.74E-06 |
| AT1G76185.1 | | protein_coding | | 70 | 59 | 54 | 38 | 40 | 32 | 147 | 145 | 138 | 82 | 106 | 87 | 111.23 | 143 | 92 | 0.492 | -1.024 | 5.86E-09 |
| AT3G51750.1 | | protein_coding | | 89 | 65 | 56 | 39 | 38 | 32 | 187 | 160 | 143 | 85 | 101 | 87 | 112.59 | 163 | 91 | 0.492 | -1.024 | 4.29E-09 |
| AT4G14100.1 | | protein_coding | transferases, transferring | 229 | 180 | 154 | 216 | 157 | 168 | 481 | 442 | 394 | 469 | 417 | 457 | 617.48 | 439 | 448 | 0.491 | -1.025 | 1.25E-18 |
| AT5G50450.1 | | protein_coding | HCP-like superfamily prot | 76 | 129 | 149 | 27 | 31 | 83 | 160 | 317 | 381 | 59 | 82 | 226 | 297.43 | 286 | 122 | 0.491 | -1.026 | 0.00119825 |
| AT1G60610.1 | | protein_coding | SBP (S-ribonuclease bindi | 63 | 63 | 60 | 44 | 24 | 37 | 132 | 155 | 154 | 95 | 64 | 101 | 114.41 | 147 | 87 | 0.49 | -1.029 | 5.37E-09 |
| AT4G13370.1 | | protein_coding | Plant protein of unknowr | 157 | 147 | 135 | 176 | 145 | 103 | 330 | 361 | 346 | 382 | 385 | 280 | 372.63 | 346 | 349 | 0.49 | -1.029 | 1.37E-12 |
| AT1G12210.1 | RFL1 | protein_coding | RPS5-like 1 | 44 | 44 | 36 | 29 | 35 | 11 | 92 | 108 | 92 | 63 | 93 | 30 | 74.04 | 97 | 62 | 0.49 | -1.03 | 0.00019144 |
| AT4G12670.1 | | protein_coding | Homeodomain-like super | 75 | 67 | 49 | 59 | 48 | 45 | 158 | 165 | 125 | 128 | 127 | 122 | 233.66 | 149 | 126 | 0.489 | -1.032 | 1.98E-12 |
| AT3G42860.1 | | protein_coding | zinc knuckle (CCHC-type) | 46 | 29 | 33 | 41 | 24 | 27 | 97 | 71 | 84 | 89 | 64 | 73 | 79.91 | 84 | 75 | 0.488 | -1.034 | 3.56E-07 |
| AT4G13250.1 | NYC1 | protein_coding | NAD(P)-binding Rossman | 3037 | 2327 | 2619 | 2661 | 1870 | 1997 | 6384 | 5717 | 6705 | 5772 | 4964 | 5431 | 7685.57 | 6269 | 5389 | 0.488 | -1.036 | 4.02E-17 |
| AT4G14030.1 | SBP1 | protein_coding | selenium-binding protein | 2010 | 1538 | 1661 | 1708 | 1380 | 1453 | 4225 | 3779 | 4252 | 3705 | 3664 | 3951 | 4432.92 | 4085 | 3773 | 0.488 | -1.036 | 4.32E-19 |
| AT3G61170.1 | | protein_coding | Tetratricopeptide repeat | 23 | 14 | 15 | 15 | 20 | 15 | 48 | 34 | 38 | 33 | 53 | 41 | 44.24 | 40 | 42 | 0.487 | -1.038 | 8.25E-05 |
| AT1G23020.2 | FRO3 | protein_coding | ferric reduction oxidase 3 | 620 | 376 | 491 | 339 | 181 | 240 | 1303 | 924 | 1257 | 735 | 481 | 653 | 919.49 | 1161 | 623 | 0.487 | -1.039 | 2.76E-11 |
| AT3G53240.1 | AtRLP45 | protein_coding | receptor like protein 45 | 210 | 229 | 164 | 102 | 88 | 89 | 441 | 563 | 420 | 221 | 234 | 242 | 367.78 | 475 | 232 | 0.487 | -1.039 | 1.15E-14 |
| AT1G57540.1 | | protein_coding | | 52 | 48 | 47 | 28 | 31 | 35 | 109 | 118 | 120 | 61 | 82 | 95 | 118.66 | 116 | 79 | 0.486 | -1.04 | 1.41E-09 |
| AT2G47890.1 | COL13 | protein_coding | B-box type zinc finger pro | 68 | 106 | 93 | 31 | 26 | 36 | 143 | 260 | 238 | 67 | 69 | 98 | 288.19 | 214 | 78 | 0.486 | -1.041 | 9.52E-11 |
| AT5G59080.1 | | protein_coding | | 236 | 328 | 351 | 196 | 248 | 202 | 496 | 806 | 899 | 425 | 658 | 549 | 574.74 | 734 | 544 | 0.486 | -1.042 | 1.09E-07 |
| AT4G13530.1 | | protein_coding | | 351 | 374 | 389 | 398 | 305 | 344 | 738 | 919 | 996 | 863 | 810 | 935 | 1086.25 | 884 | 869 | 0.486 | -1.042 | 5.12E-16 |
| AT4G13270.1 | | protein_coding | Late embryogenesis abun | 209 | 176 | 183 | 195 | 156 | 180 | 439 | 432 | 468 | 423 | 414 | 489 | 655.95 | 446 | 442 | 0.486 | -1.042 | 1.80E-21 |
| AT4G14940.1 | ATAO1 | protein_coding | amine oxidase 1 | 44 | 44 | 36 | 62 | 33 | 34 | 92 | 108 | 92 | 134 | 88 | 92 | 131.24 | 97 | 105 | 0.485 | -1.043 | 7.54E-10 |
| AT2G21710.1 | EMB2219 | protein_coding | Mitochondrial transcripti | 141 | 117 | 93 | 76 | 73 | 47 | 296 | 287 | 238 | 165 | 194 | 128 | 243.48 | 274 | 162 | 0.485 | -1.044 | 4.74E-12 |
| AT4G14840.1 | | protein_coding | | 76 | 64 | 67 | 87 | 62 | 63 | 160 | 157 | 172 | 189 | 165 | 171 | 195.05 | 163 | 175 | 0.485 | -1.044 | 1.47E-12 |
| AT3G22231.1 | PCC1 | protein_coding | pathogen and circadian c | 72 | 51 | 45 | 54 | 57 | 50 | 151 | 125 | 115 | 117 | 151 | 136 | 77.13 | 130 | 135 | 0.484 | -1.048 | 0.00295013 |
| AT4G13340.1 | LRX3 | protein_coding | Leucine-rich repeat (LRR) | 691 | 523 | 418 | 870 | 505 | 479 | 1453 | 1285 | 1070 | 1887 | 1341 | 1303 | 5410.64 | 1269 | 1510 | 0.483 | -1.049 | 7.02E-10 |
| AT3G25970.1 | PCMP-E46 | protein_coding | Pentatricopeptide repeat | 28 | 21 | 20 | 15 | 19 | 13 | 59 | 52 | 51 | 33 | 50 | 35 | 52.66 | 54 | 39 | 0.483 | -1.05 | 4.27E-05 |
| AT1G62770.1 | | protein_coding | Plant invertase/pectin me | 36 | 22 | 29 | 25 | 25 | 57 | 76 | 54 | 74 | 54 | 66 | 155 | 61.35 | 68 | 92 | 0.482 | -1.053 | 0.00151619 |
| AT5G28370.1 | | protein_coding | Pentatricopeptide repeat | 142 | 94 | 99 | 97 | 76 | 71 | 298 | 231 | 253 | 210 | 202 | 193 | 213.59 | 261 | 202 | 0.482 | -1.053 | 3.18E-11 |

| | | | | | | | | | | | | | | | | | | | | | |
|---|---|---|---|---|---|---|---|---|---|---|---|---|---|---|---|---|---|---|---|---|---|
| AT2G27610.1 | PCMP-H60 | protein_coding | Tetratricopeptide repeat | 47 | 26 | 25 | 35 | 23 | 26 | 99 | 64 | 64 | 76 | 61 | 71 | 69.02 | 76 | 69 | 0.481 | -1.055 | 2.34E-05 |
| AT5G57110.1 | ACA8 | protein_coding | autoinhibited Ca2+ -ATPa | 730 | 886 | 987 | 450 | 450 | 399 | 1535 | 2177 | 2527 | 976 | 1195 | 1085 | 1617.05 | 2080 | 1085 | 0.481 | -1.056 | 7.59E-11 |
| AT1G10760.1 | GWD1 | protein_coding | Pyruvate phosphate dikin | 8924 | 9089 | 8198 | 3802 | 3722 | 3185 | 18759 | 22330 | 20987 | 8247 | 9881 | 8661 | 14589.25 | 20692 | 8930 | 0.481 | -1.057 | 8.04E-15 |
| AT4G14480.1 | BLUS1 | protein_coding | Protein kinase superfamil | 232 | 312 | 300 | 216 | 243 | 228 | 488 | 767 | 768 | 469 | 645 | 620 | 632.72 | 674 | 578 | 0.48 | -1.058 | 1.75E-10 |
| AT2G43920.1 | HOL2 | protein_coding | S-adenosyl-L-methionine- | 100 | 111 | 83 | 90 | 64 | 83 | 210 | 273 | 212 | 195 | 170 | 226 | 337.34 | 232 | 197 | 0.48 | -1.059 | 3.02E-13 |
| AT2G37720.1 | TBL15 | protein_coding | TRICHOME BIREFRINGEN | 39 | 39 | 28 | 10 | 14 | 9 | 82 | 96 | 72 | 22 | 37 | 24 | 48.41 | 83 | 28 | 0.48 | -1.06 | 3.34E-05 |
| AT3G61520.1 | | protein_coding | Pentatricopeptide repeat | 184 | 113 | 135 | 95 | 97 | 86 | 387 | 278 | 346 | 206 | 258 | 234 | 262.62 | 337 | 233 | 0.48 | -1.06 | 1.90E-13 |
| AT3G03580.1 | PCMP-H23 | protein_coding | Tetratricopeptide repeat | 88 | 54 | 63 | 51 | 49 | 26 | 185 | 133 | 161 | 111 | 130 | 71 | 121.83 | 160 | 104 | 0.479 | -1.062 | 1.54E-07 |
| AT4G14760.1 | NET1B | protein_coding | kinase interacting (KIP1-li | 1790 | 1414 | 1321 | 1418 | 1262 | 873 | 3763 | 3474 | 3382 | 3076 | 3350 | 2374 | 2603.27 | 3540 | 2933 | 0.479 | -1.062 | 8.42E-15 |
| AT1G52200.1 | PCR8 | protein_coding | PLAC8 family protein | 129 | 47 | 71 | 81 | 20 | 48 | 271 | 115 | 182 | 176 | 53 | 131 | 141.46 | 189 | 120 | 0.478 | -1.065 | 0.00091112 |
| AT4G12770.1 | | protein_coding | Chaperone DnaJ-domain | 570 | 568 | 525 | 590 | 490 | 518 | 1198 | 1395 | 1344 | 1280 | 1301 | 1409 | 1512.88 | 1312 | 1330 | 0.477 | -1.067 | 1.09E-19 |
| AT3G53220.1 | | protein_coding | Thioredoxin superfamily | 31 | 24 | 28 | 38 | 34 | 19 | 65 | 59 | 72 | 82 | 90 | 52 | 83.07 | 65 | 75 | 0.477 | -1.068 | 7.33E-08 |
| AT1G71696.2 | SOL1 | protein_coding | carboxypeptidase D, puta | 258 | 186 | 185 | 124 | 96 | 69 | 542 | 457 | 474 | 269 | 255 | 188 | 352.83 | 491 | 237 | 0.477 | -1.069 | 1.80E-15 |
| AT2G20260.1 | PSAE2 | protein_coding | photosystem I subunit E- | 3053 | 4980 | 3602 | 1807 | 1887 | 2743 | 6418 | 12235 | 9221 | 3920 | 5010 | 7459 | 11757.96 | 9291 | 5463 | 0.476 | -1.07 | 1.06E-06 |
| AT3G25440.1 | | protein_coding | RNA-binding CRS1 / YhbY | 61 | 58 | 60 | 21 | 23 | 28 | 128 | 142 | 154 | 46 | 61 | 76 | 98.46 | 141 | 61 | 0.476 | -1.07 | 5.98E-08 |
| AT5G52110.1 | HCF208 | protein_coding | Protein of unknown funct | 241 | 210 | 226 | 118 | 86 | 89 | 507 | 516 | 579 | 256 | 228 | 242 | 378.04 | 534 | 242 | 0.476 | -1.071 | 2.90E-15 |
| AT4G14190.1 | | protein_coding | Pentatricopeptide repeat | 199 | 154 | 166 | 177 | 143 | 116 | 418 | 378 | 425 | 384 | 380 | 315 | 452.93 | 407 | 360 | 0.476 | -1.072 | 2.03E-18 |
| AT3G59350.1 | PTI13 | protein_coding | Protein kinase superfamil | 480 | 502 | 825 | 260 | 173 | 413 | 1009 | 1233 | 2112 | 564 | 459 | 1123 | 1146.39 | 1451 | 715 | 0.475 | -1.073 | 6.02E-05 |
| AT1G45163.1 | | protein_coding | | 63 | 75 | 80 | 39 | 32 | 27 | 132 | 184 | 205 | 85 | 85 | 73 | 183.28 | 174 | 81 | 0.474 | -1.077 | 4.10E-11 |
| AT3G15140.1 | | protein_coding | Polynucleotidyl transfera | 97 | 73 | 59 | 74 | 63 | 51 | 204 | 179 | 151 | 161 | 167 | 139 | 236.87 | 178 | 156 | 0.474 | -1.077 | 4.36E-12 |
| AT1G17630.1 | PCMP-E72 | protein_coding | Pentatricopeptide repeat | 51 | 37 | 37 | 34 | 22 | 20 | 107 | 91 | 95 | 74 | 58 | 54 | 82.29 | 98 | 62 | 0.473 | -1.079 | 2.64E-05 |
| AT1G65032.1 | | protein_coding | | 44 | 32 | 23 | 23 | 19 | 23 | 92 | 79 | 59 | 50 | 50 | 63 | 88.14 | 77 | 54 | 0.473 | -1.081 | 2.03E-07 |
| AT1G69350.1 | PCMP-E66 | protein_coding | Tetratricopeptide repeat | 38 | 28 | 23 | 23 | 14 | 10 | 80 | 69 | 59 | 50 | 37 | 27 | 46.77 | 69 | 38 | 0.472 | -1.085 | 0.00089472 |
| AT4G13710.1 | | protein_coding | Pectin lyase-like superfan | 49 | 46 | 40 | 51 | 54 | 49 | 103 | 113 | 102 | 111 | 143 | 133 | 133.75 | 106 | 129 | 0.471 | -1.086 | 4.10E-10 |
| AT5G10530.1 | LECRK91 | protein_coding | Concanavalin A-like lectir | 33 | 32 | 33 | 29 | 42 | 17 | 69 | 79 | 84 | 63 | 112 | 46 | 119.34 | 77 | 74 | 0.471 | -1.087 | 2.42E-07 |
| AT4G14710.5 | ARD3 | protein_coding | RmlC-like cupins superfar | 733 | 552 | 549 | 757 | 479 | 488 | 1541 | 1356 | 1405 | 1642 | 1272 | 1327 | 1932.94 | 1434 | 1414 | 0.47 | -1.089 | 1.78E-18 |
| AT1G49490.1 | PEX2 | protein_coding | Leucine-rich repeat (LRR) | 2 | 1 | 3 | 5 | 2 | 2 | 4 | 2 | 8 | 11 | 5 | 5 | 23.19 | 5 | 7 | 0.47 | -1.09 | 0.00225705 |
| AT2G39850.1 | | protein_coding | Subtilisin-like serine endc | 136 | 235 | 170 | 117 | 166 | 124 | 286 | 577 | 435 | 254 | 441 | 337 | 468.31 | 433 | 344 | 0.469 | -1.092 | 2.20E-07 |
| AT2G16750.1 | | protein_coding | Protein kinase protein wi | 142 | 92 | 122 | 128 | 63 | 52 | 298 | 226 | 312 | 278 | 167 | 141 | 220.46 | 279 | 195 | 0.469 | -1.093 | 1.35E-08 |
| AT5G47510.1 | SFH11 | protein_coding | Sec14p-like phosphatidyli | 48 | 25 | 30 | 37 | 35 | 30 | 101 | 61 | 77 | 80 | 93 | 82 | 97.74 | 80 | 85 | 0.468 | -1.095 | 5.18E-09 |
| AT1G56145.2 | | protein_coding | Leucine-rich repeat transı | 149 | 118 | 122 | 104 | 76 | 69 | 313 | 290 | 312 | 226 | 202 | 188 | 250.62 | 305 | 205 | 0.468 | -1.095 | 8.81E-12 |
| AT1G01700.1 | ROPGEF2 | protein_coding | RHO guanyl-nucleotide e | 25 | 13 | 25 | 16 | 20 | 11 | 53 | 32 | 64 | 35 | 53 | 30 | 41.14 | 50 | 39 | 0.467 | -1.098 | 0.00010915 |
| AT3G08820.1 | PCMP-H84 | protein_coding | Pentatricopeptide repeat | 21 | 20 | 10 | 31 | 14 | 10 | 44 | 49 | 26 | 67 | 37 | 27 | 61.76 | 40 | 44 | 0.467 | -1.098 | 6.10E-05 |
| AT4G12750.1 | | protein_coding | Homeodomain-like transc | 463 | 347 | 339 | 263 | 241 | 199 | 973 | 853 | 868 | 570 | 640 | 541 | 799.6 | 898 | 584 | 0.467 | -1.098 | 1.99E-22 |
| AT4G23000.1 | | protein_coding | Calcineurin-like metallo-p | 128 | 89 | 87 | 50 | 38 | 25 | 269 | 219 | 223 | 108 | 101 | 68 | 168.41 | 237 | 92 | 0.467 | -1.099 | 8.28E-10 |
| AT5G36930.2 | | protein_coding | Disease resistance protei | 377 | 246 | 229 | 195 | 140 | 113 | 792 | 604 | 586 | 423 | 372 | 307 | 398.14 | 661 | 367 | 0.467 | -1.099 | 5.55E-14 |
| AT2G35820.1 | | protein_coding | ureidoglycolate hydrolase | 263 | 190 | 248 | 237 | 175 | 176 | 553 | 467 | 635 | 514 | 465 | 479 | 375.89 | 552 | 486 | 0.464 | -1.106 | 2.22E-15 |
| AT5G57630.1 | CIPK21 | protein_coding | CBL-interacting protein ki | 574 | 662 | 951 | 293 | 235 | 352 | 1207 | 1626 | 2435 | 636 | 624 | 957 | 1199.31 | 1756 | 739 | 0.464 | -1.107 | 1.96E-07 |
| AT5G19020.1 | PCMP-E42 | protein_coding | mitochondrial editing fac | 55 | 38 | 41 | 38 | 44 | 39 | 116 | 93 | 105 | 82 | 117 | 106 | 106.14 | 105 | 102 | 0.463 | -1.11 | 1.34E-07 |
| AT4G12680.1 | | protein_coding | | 106 | 85 | 107 | 87 | 71 | 65 | 223 | 209 | 274 | 189 | 188 | 177 | 287.4 | 235 | 185 | 0.463 | -1.11 | 2.91E-17 |
| AT3G12770.1 | PCMP-H43 | protein_coding | mitochondrial editing fac | 48 | 29 | 32 | 28 | 26 | 15 | 101 | 71 | 82 | 61 | 69 | 41 | 87.77 | 85 | 57 | 0.463 | -1.111 | 4.78E-09 |
| AT4G14500.1 | | protein_coding | Polyketide cyclase/dehyd | 738 | 817 | 888 | 675 | 686 | 816 | 1551 | 2007 | 2273 | 1464 | 1821 | 2219 | 2368.07 | 1944 | 1835 | 0.463 | -1.111 | 4.24E-12 |
| AT4G13660.1 | PRR2 | protein_coding | pinoresinol reductase 2 | 33 | 9 | 16 | 19 | 7 | 15 | 69 | 22 | 41 | 41 | 19 | 41 | 44.45 | 44 | 34 | 0.463 | -1.112 | 0.00187438 |
| AT4G23950.2 | | protein_coding | Galactose-binding proteir | 99 | 76 | 81 | 57 | 48 | 36 | 208 | 187 | 207 | 124 | 127 | 98 | 157.81 | 201 | 116 | 0.462 | -1.113 | 1.83E-11 |
| AT4G14342.1 | | protein_coding | Splicing factor 3B subunit | 606 | 432 | 468 | 624 | 459 | 336 | 1274 | 1061 | 1198 | 1354 | 1219 | 914 | 1706.8 | 1178 | 1162 | 0.462 | -1.113 | 3.79E-15 |
| AT5G57230.1 | | protein_coding | Thioredoxin superfamily | 58 | 59 | 41 | 53 | 37 | 28 | 122 | 145 | 105 | 115 | 98 | 76 | 149.91 | 124 | 96 | 0.462 | -1.114 | 3.56E-12 |
| AT4G16835.1 | DYW10 | protein_coding | Tetratricopeptide repeat | 51 | 38 | 30 | 23 | 23 | 20 | 107 | 93 | 77 | 50 | 61 | 54 | 81.88 | 92 | 55 | 0.462 | -1.115 | 7.68E-09 |
| AT5G59700.1 | | protein_coding | Protein kinase superfamil | 56 | 66 | 44 | 27 | 32 | 21 | 118 | 162 | 113 | 59 | 85 | 57 | 79.9 | 131 | 67 | 0.461 | -1.117 | 0.0001489 |
| AT4G13550.1 | | protein_coding | triglyceride lipases;triglyc | 591 | 499 | 530 | 545 | 437 | 394 | 1242 | 1226 | 1357 | 1182 | 1160 | 1071 | 1281.59 | 1275 | 1138 | 0.461 | -1.118 | 1.32E-20 |
| AT1G33330.1 | | protein_coding | Class I peptide chain relea | 146 | 144 | 100 | 65 | 56 | 73 | 307 | 354 | 256 | 141 | 149 | 199 | 369.63 | 306 | 163 | 0.46 | -1.119 | 1.75E-14 |
| AT4G14965.1 | MAPR4 | protein_coding | membrane-associated pr | 254 | 201 | 204 | 231 | 190 | 193 | 534 | 494 | 522 | 501 | 504 | 525 | 723.65 | 517 | 510 | 0.459 | -1.123 | 1.23E-25 |
| AT4G14145.1 | | protein_coding | | 149 | 120 | 150 | 130 | 83 | 130 | 313 | 295 | 384 | 282 | 220 | 354 | 361.99 | 331 | 285 | 0.459 | -1.124 | 9.14E-14 |
| AT4G13830.2 | ATJ20 | protein_coding | DNAJ-like 20 | 158 | 211 | 300 | 224 | 194 | 279 | 332 | 518 | 768 | 486 | 515 | 759 | 1297.75 | 539 | 587 | 0.459 | -1.125 | 2.33E-07 |
| AT4G12800.1 | PSAL | protein_coding | photosystem I subunit l | 16339 | 28036 | 18222 | 15907 | 15306 | 25059 | 34346 | 68879 | 46649 | 34504 | 40634 | 68146 | 145423.08 | 49958 | 47761 | 0.458 | -1.126 | 2.89E-05 |
| AT4G15545.1 | | protein_coding | | 4414 | 3864 | 3960 | 4699 | 3963 | 4200 | 9279 | 9493 | 10138 | 10193 | 10521 | 11422 | 14371.98 | 9637 | 10712 | 0.458 | -1.127 | 3.04E-19 |
| AT4G13870.2 | WEX | protein_coding | Werner syndrome-like ex | 147 | 93 | 111 | 69 | 55 | 70 | 309 | 228 | 284 | 150 | 146 | 190 | 226.07 | 274 | 162 | 0.457 | -1.128 | 1.88E-13 |
| AT4G14770.1 | TCX2 | protein_coding | TESMIN/TSO1-like CXC 2 | 379 | 337 | 325 | 336 | 332 | 243 | 797 | 828 | 832 | 729 | 881 | 661 | 875.44 | 819 | 757 | 0.457 | -1.129 | 2.20E-21 |
| AT1G80570.2 | FBL14 | protein_coding | RNI-like superfamily prot | 25 | 18 | 26 | 13 | 11 | 19 | 53 | 44 | 67 | 28 | 29 | 52 | 46.56 | 55 | 36 | 0.457 | -1.13 | 1.31E-05 |
| AT4G14920.1 | | protein_coding | Acyl-CoA N-acyltransfera | 528 | 475 | 465 | 466 | 426 | 403 | 1110 | 1167 | 1190 | 1011 | 1131 | 1096 | 1260.65 | 1156 | 1079 | 0.456 | -1.131 | 7.37E-23 |
| AT1G74700.1 | NUZ | protein_coding | tRNAse Z1 | 65 | 48 | 47 | 43 | 29 | 31 | 137 | 118 | 120 | 93 | 77 | 84 | 110.74 | 125 | 85 | 0.456 | -1.132 | 5.93E-10 |
| AT5G25500.1 | | protein_coding | | 67 | 44 | 48 | 59 | 42 | 33 | 141 | 108 | 123 | 128 | 112 | 90 | 104.53 | 124 | 110 | 0.455 | -1.136 | 1.54E-06 |
| AT4G15260.1 | | protein_coding | UDP-Glycosyltransferase | 13 | 22 | 23 | 22 | 13 | 37 | 27 | 54 | 59 | 48 | 35 | 101 | 74.62 | 47 | 61 | 0.452 | -1.146 | 0.00039567 |
| AT1G31430.1 | PCMP-E55 | protein_coding | Pentatricopeptide repeat | 37 | 29 | 37 | 18 | 10 | 12 | 78 | 71 | 95 | 39 | 27 | 33 | 54.11 | 81 | 33 | 0.451 | -1.148 | 3.75E-05 |
| AT5G10570.1 | BHLH61 | protein_coding | basic helix-loop-helix (bH | 21 | 42 | 29 | 19 | 13 | 9 | 44 | 103 | 74 | 41 | 35 | 24 | 55.88 | 74 | 33 | 0.451 | -1.148 | 3.48E-05 |
| AT4G14720.1 | TIFY4B | protein_coding | TIFY domain/Divergent C | 161 | 215 | 236 | 150 | 158 | 200 | 338 | 528 | 604 | 325 | 419 | 544 | 544.1 | 490 | 429 | 0.451 | -1.148 | 2.35E-10 |
| AT4G14120.1 | | protein_coding | | 56 | 61 | 47 | 52 | 48 | 25 | 118 | 150 | 120 | 113 | 127 | 68 | 134.1 | 129 | 103 | 0.45 | -1.151 | 1.92E-09 |
| AT5G43290.1 | WRKY49 | protein_coding | WRKY DNA-binding prote | 42 | 9 | 14 | 15 | 4 | 9 | 88 | 22 | 36 | 33 | 11 | 24 | 33.91 | 49 | 23 | 0.45 | -1.152 | 0.0088391 |
| AT1G03100.1 | | protein_coding | Pentatricopeptide repeat | 114 | 54 | 67 | 49 | 44 | 26 | 240 | 133 | 172 | 106 | 117 | 71 | 130.76 | 182 | 98 | 0.45 | -1.152 | 2.34E-06 |
| AT5G61950.1 | | protein_coding | Ubiquitin carboxyl-termir | 137 | 115 | 114 | 68 | 91 | 43 | 288 | 283 | 292 | 147 | 242 | 117 | 218.98 | 288 | 169 | 0.45 | -1.153 | 1.35E-10 |
| AT4G14000.1 | | protein_coding | Putative methyltransfera | 235 | 232 | 200 | 205 | 151 | 170 | 494 | 570 | 512 | 445 | 401 | 462 | 656.76 | 525 | 436 | 0.45 | -1.154 | 5.67E-27 |
| AT2G38465.1 | | protein_coding | | 31 | 35 | 45 | 12 | 14 | 22 | 65 | 86 | 115 | 26 | 37 | 60 | 100.47 | 89 | 41 | 0.449 | -1.156 | 1.60E-08 |
| AT3G52390.2 | | protein_coding | TatD related DNase | 90 | 62 | 52 | 55 | 39 | 45 | 189 | 152 | 133 | 119 | 104 | 122 | 183.34 | 158 | 115 | 0.449 | -1.156 | 1.28E-11 |
| AT4G13345.1 | MEE55 | protein_coding | Serinc-domain containing | 236 | 172 | 191 | 260 | 228 | 181 | 496 | 423 | 489 | 564 | 605 | 492 | 575 | 469 | 554 | 0.448 | -1.157 | 5.58E-20 |
| AT4G12690.1 | | protein_coding | Plant protein of unknowr | 266 | 290 | 289 | 305 | 231 | 285 | 559 | 712 | 740 | 662 | 613 | 775 | 531.4 | 670 | 683 | 0.448 | -1.158 | 2.01E-12 |
| AT4G04410.1 | | transposable_elen | transposable element ger | 164 | 179 | 180 | 168 | 222 | 166 | 345 | 440 | 461 | 364 | 589 | 451 | 308.32 | 415 | 468 | 0.447 | -1.161 | 8.20E-07 |
| AT1G08070.1 | PCMP-H12 | protein_coding | Tetratricopeptide repeat | 44 | 25 | 33 | 25 | 21 | 19 | 92 | 61 | 84 | 54 | 56 | 52 | 62.6 | 79 | 54 | 0.446 | -1.164 | 7.25E-07 |
| AT5G08490.1 | PCMP-E32 | protein_coding | Tetratricopeptide repeat | 27 | 23 | 21 | 19 | 19 | 9 | 57 | 57 | 54 | 41 | 50 | 24 | 45.66 | 56 | 38 | 0.445 | -1.167 | 8.26E-06 |
| AT4G13540.1 | | protein_coding | | 6 | 3 | 2 | 5 | 9 | 9 | 13 | 7 | 5 | 11 | 24 | 24 | 62.76 | 8 | 20 | 0.445 | -1.169 | 1.34E-07 |
| AT3G09540.1 | | protein_coding | Pectin lyase-like superfan | 141 | 172 | 116 | 56 | 57 | 55 | 296 | 423 | 297 | 121 | 151 | 150 | 322.19 | 339 | 141 | 0.444 | -1.171 | 1.05E-16 |
| AT5G46680.1 | | protein_coding | Pentatricopeptide repeat | 36 | 29 | 34 | 27 | 19 | 13 | 76 | 71 | 87 | 59 | 50 | 35 | 65.78 | 78 | 48 | 0.443 | -1.174 | 1.18E-06 |
| AT2G36500.1 | CBSCBSPB2 | protein_coding | CBS / octicosapeptide/Ph | 49 | 42 | 28 | 17 | 12 | 14 | 103 | 103 | 72 | 37 | 32 | 38 | 57.11 | 93 | 36 | 0.443 | -1.175 | 1.08E-06 |
| AT5G03720.1 | HSFA3 | protein_coding | heat shock transcription f | 31 | 44 | 55 | 14 | 19 | 26 | 65 | 108 | 141 | 30 | 50 | 71 | 83.68 | 105 | 50 | 0.442 | -1.177 | 2.27E-06 |
| AT4G36930.1 | SPT | protein_coding | basic helix-loop-helix (bH | 102 | 59 | 53 | 42 | 39 | 45 | 214 | 145 | 136 | 91 | 104 | 122 | 136.46 | 165 | 106 | 0.442 | -1.178 | 5.84E-10 |
| AT4G16146.1 | | protein_coding | cAMP-regulated phospho | 144 | 176 | 207 | 42 | 66 | 51 | 303 | 432 | 530 | 91 | 175 | 139 | 357.31 | 422 | 135 | 0.442 | -1.179 | 3.64E-08 |
| AT4G35130.1 | PCMP-H27 | protein_coding | Tetratricopeptide repeat | 46 | 31 | 31 | 43 | 22 | 29 | 97 | 76 | 79 | 93 | 58 | 79 | 79.58 | 84 | 77 | 0.442 | -1.179 | 1.31E-08 |
| AT4G15240.1 | | protein_coding | Protein of unknown funct | 122 | 138 | 136 | 128 | 106 | 151 | 256 | 339 | 348 | 278 | 281 | 411 | 378.26 | 314 | 323 | 0.442 | -1.179 | 8.27E-15 |
| AT4G13350.1 | NIG | protein_coding | NSP (nuclear shuttle prot | 1114 | 888 | 907 | 1085 | 718 | 843 | 2342 | 2182 | 2322 | 2353 | 1906 | 2292 | 2988.57 | 2282 | 2184 | 0.442 | -1.179 | 1.32E-22 |
| AT4G13360.1 | | protein_coding | ATP-dependent caseinoly | 636 | 433 | 485 | 589 | 349 | 370 | 1337 | 1064 | 1242 | 1278 | 927 | 1006 | 1595.75 | 1214 | 1070 | 0.441 | -1.181 | 3.31E-18 |
| AT1G02900.1 | RALF1 | protein_coding | rapid alkalinization factor | 10 | 13 | 9 | 19 | 10 | 8 | 21 | 32 | 23 | 41 | 27 | 22 | 32.88 | 25 | 30 | 0.44 | -1.183 | 0.00056205 |
| AT5G67245.1 | | protein_coding | | 17 | 16 | 13 | 5 | 2 | 6 | 36 | 39 | 33 | 11 | 5 | 16 | 31.88 | 36 | 11 | 0.44 | -1.183 | 8.94E-05 |
| AT1G14470.1 | PCMP-A4 | protein_coding | Pentatricopeptide repeat | 43 | 24 | 36 | 28 | 20 | 29 | 90 | 59 | 92 | 61 | 53 | 79 | 60.11 | 80 | 64 | 0.44 | -1.184 | 1.57E-05 |
| AT5G52850.1 | PCMP-H31 | protein_coding | Pentatricopeptide repeat | 81 | 72 | 55 | 64 | 72 | 49 | 170 | 177 | 141 | 139 | 191 | 133 | 142.72 | 163 | 154 | 0.439 | -1.187 | 2.58E-08 |
| AT4G35050.1 | MSI3 | protein_coding | Transducin family protein | 200 | 169 | 145 | 142 | 148 | 135 | 420 | 415 | 371 | 308 | 393 | 367 | 439.38 | 402 | 356 | 0.438 | -1.19 | 1.54E-19 |
| AT3G02010.1 | PCMP-H36 | protein_coding | Pentatricopeptide repeat | 46 | 34 | 22 | 29 | 23 | 26 | 97 | 84 | 56 | 63 | 61 | 71 | 70.38 | 79 | 65 | 0.437 | -1.193 | 4.27E-06 |
| AT1G79120.1 | | protein_coding | Ubiquitin carboxyl-termir | 33 | 16 | 24 | 31 | 15 | 12 | 69 | 39 | 61 | 67 | 40 | 33 | 45.41 | 56 | 47 | 0.436 | -1.198 | 5.95E-05 |
| AT4G04330.1 | | protein_coding | Chaperonin-like RbcX pro | 516 | 852 | 954 | 151 | 235 | 228 | 1085 | 2093 | 2442 | 328 | 624 | 620 | 1684.04 | 1873 | 524 | 0.435 | -1.201 | 1.31E-06 |
| AT4G21220.1 | LPXD2 | protein_coding | Trimeric LpxA-like enzym | 34 | 17 | 17 | 26 | 21 | 16 | 71 | 42 | 44 | 56 | 56 | 44 | 66.13 | 52 | 52 | 0.434 | -1.203 | 1.15E-07 |
| AT1G32190.1 | | protein_coding | alpha/beta-Hydrolases su | 60 | 71 | 81 | 22 | 33 | 52 | 126 | 174 | 207 | 48 | 88 | 141 | 169.96 | 169 | 92 | 0.434 | -1.204 | 8.00E-07 |
| AT4G13720.1 | | protein_coding | Inosine triphosphate pyrc | 436 | 318 | 300 | 335 | 320 | 262 | 917 | 781 | 768 | 727 | 850 | 712 | 1167.41 | 822 | 763 | 0.434 | -1.204 | 1.13E-20 |
| AT4G14930.1 | | protein_coding | Survival protein SurE-like | 346 | 258 | 261 | 256 | 210 | 223 | 727 | 634 | 668 | 555 | 558 | 606 | 764.82 | 676 | 573 | 0.434 | -1.204 | 9.07E-28 |
| AT1G14205.1 | | protein_coding | Ribosomal L18p/L5e fami | 52 | 42 | 39 | 19 | 22 | 17 | 109 | 103 | 100 | 41 | 58 | 46 | 97.03 | 104 | 48 | 0.434 | -1.205 | 7.86E-12 |
| AT4G14096.1 | | protein_coding | F-box/RNI-like superfamil | 137 | 116 | 104 | 126 | 100 | 98 | 288 | 285 | 266 | 273 | 265 | 267 | 310.53 | 280 | 268 | 0.433 | -1.206 | 4.06E-19 |
| AT4G14622.1 | CPuORF60 | protein_coding | conserved peptide upstre | 106 | 138 | 168 | 114 | 145 | 108 | 223 | 339 | 430 | 247 | 385 | 294 | 298.21 | 331 | 309 | 0.433 | -1.207 | 7.68E-10 |
| AT4G13510.1 | AMT1-1 | protein_coding | ammonium transporter 1 | 262 | 218 | 119 | 276 | 134 | 156 | 551 | 536 | 305 | 599 | 356 | 424 | 814.28 | 464 | 460 | 0.433 | -1.207 | 1.30E-10 |
| AT4G14180.1 | AtPRD1 | protein_coding | putative recombination ir | 26 | 21 | 16 | 16 | 20 | 17 | 55 | 52 | 41 | 35 | 53 | 46 | 45.83 | 49 | 45 | 0.433 | -1.209 | 2.99E-05 |
| AT1G13750.1 | PAP1 | protein_coding | Purple acid phosphatases | 152 | 69 | 59 | 54 | 33 | 30 | 320 | 170 | 151 | 117 | 88 | 82 | 138.37 | 214 | 96 | 0.433 | -1.209 | 1.39E-05 |
| AT3G61060.2 | PP2A13 | protein_coding | phloem protein 2-A13 | 191 | 334 | 568 | 113 | 184 | 283 | 401 | 821 | 1454 | 245 | 488 | 770 | 551.32 | 892 | 501 | 0.432 | -1.211 | 0.00068007 |
| AT4G14615.1 | | protein_coding | | 307 | 199 | 224 | 244 | 186 | 171 | 645 | 489 | 573 | 529 | 494 | 465 | 688.18 | 569 | 496 | 0.432 | -1.211 | 2.41E-25 |
| AT4G13600.1 | | protein_coding | Carbohydrate-binding X8 | 3 | 2 | 3 | 4 | 4 | 4 | 6 | 5 | 8 | 9 | 11 | 11 | 28.79 | 6 | 10 | 0.432 | -1.212 | 0.00048954 |
| AT5G09950.1 | PCMP-H35 | protein_coding | Tetratricopeptide repeat | 20 | 20 | 12 | 16 | 19 | 9 | 42 | 49 | 31 | 35 | 50 | 24 | 43.98 | 41 | 36 | 0.432 | -1.212 | 2.08E-05 |
| AT2G37950.1 | | protein_coding | RING/FYVE/PHD zinc fing | 8 | 22 | 7 | 8 | 11 | 5 | 17 | 54 | 18 | 17 | 29 | 14 | 68.05 | 30 | 20 | 0.432 | -1.212 | 9.12E-07 |
| AT3G19330.1 | | protein_coding | Protein of unknown funct | 15 | 20 | 10 | 11 | 9 | 11 | 32 | 49 | 26 | 24 | 24 | 30 | 68.69 | 36 | 26 | 0.431 | -1.213 | 5.88E-09 |
| AT3G52605.1 | | other_rna | other RNA | 5 | 14 | 4 | 6 | 2 | 3 | 11 | 34 | 10 | 13 | 5 | 8 | 23.54 | 18 | 9 | 0.431 | -1.215 | 0.00403153 |
| AT4G14890.1 | FdC2 | protein_coding | 2Fe-2S ferredoxin-like sup | 606 | 551 | 387 | 534 | 415 | 490 | 1274 | 1354 | 991 | 1158 | 1102 | 1333 | 3988.55 | 1206 | 1198 | 0.431 | -1.215 | 4.22E-20 |
| AT1G63240.1 | | protein_coding | | 301 | 324 | 328 | 122 | 108 | 114 | 633 | 796 | 840 | 265 | 287 | 310 | 523.04 | 756 | 287 | 0.431 | -1.215 | 2.22E-22 |
| AT4G15010.1 | | protein_coding | Mitochondrial substrate | 175 | 139 | 181 | 161 | 113 | 127 | 368 | 341 | 463 | 349 | 300 | 345 | 512.69 | 391 | 331 | 0.431 | -1.216 | 1.69E-23 |
| AT1G56300.1 | | protein_coding | Chaperone DnaJ-domain | 511 | 675 | 684 | 100 | 199 | 157 | 1074 | 1658 | 1751 | 217 | 528 | 427 | 1331.45 | 1494 | 391 | 0.43 | -1.217 | 7.29E-08 |
| AT3G19250.1 | | protein_coding | Protein of unknown funct | 13 | 11 | 9 | 13 | 4 | 6 | 27 | 27 | 23 | 28 | 11 | 16 | 38.79 | 26 | 18 | 0.43 | -1.218 | 3.95E-06 |
| AT1G66540.1 | | protein_coding | Cytochrome P450 superfa | 44 | 27 | 21 | 21 | 20 | 14 | 92 | 66 | 54 | 46 | 53 | 38 | 88.24 | 71 | 46 | 0.43 | -1.218 | 2.29E-10 |
| AT1G68260.1 | | protein_coding | Thioesterase superfamily | 189 | 164 | 173 | 90 | 75 | 98 | 397 | 403 | 443 | 195 | 199 | 267 | 438.55 | 414 | 220 | 0.43 | -1.219 | 3.39E-22 |
| AT4G19191.1 | PCMP-E1 | protein_coding | Tetratricopeptide repeat | 28 | 23 | 20 | 11 | 10 | 14 | 59 | 57 | 51 | 24 | 27 | 38 | 40.4 | 56 | 30 | 0.429 | -1.22 | 3.05E-06 |
| AT4G16880.1 | | protein_coding | Leucine-rich repeat (LRR) | 686 | 689 | 648 | 307 | 278 | 255 | 1442 | 1693 | 1659 | 666 | 738 | 693 | 1118.16 | 1598 | 699 | 0.429 | -1.221 | 3.76E-17 |
| AT4G22212.1 | | protein_coding | Arabidopsis defensin-like | 17 | 17 | 19 | 15 | 18 | 11 | 36 | 42 | 49 | 33 | 48 | 30 | 52.99 | 42 | 37 | 0.428 | -1.223 | 3.62E-06 |
| AT2G01510.1 | PCMP-H37 | protein_coding | Tetratricopeptide repeat | 35 | 30 | 22 | 20 | 19 | 13 | 74 | 74 | 56 | 43 | 50 | 35 | 57.54 | 68 | 43 | 0.428 | -1.223 | 2.06E-06 |
| AT5G27110.1 | PCMP-E14 | protein_coding | Tetratricopeptide repeat | 47 | 23 | 29 | 46 | 34 | 36 | 99 | 57 | 74 | 100 | 90 | 98 | 96.47 | 77 | 96 | 0.428 | -1.223 | 1.08E-08 |
| AT4G15520.1 | | protein_coding | tRNA/rRNA methyltransf | 129 | 115 | 80 | 101 | 77 | 83 | 271 | 283 | 205 | 219 | 204 | 226 | 324.41 | 253 | 216 | 0.428 | -1.225 | 8.51E-18 |
| AT4G13410.1 | CSLA15 | protein_coding | Nucleotide-diphospho-su | 113 | 52 | 21 | 107 | 35 | 24 | 238 | 128 | 54 | 232 | 93 | 65 | 197.81 | 140 | 130 | 0.427 | -1.226 | 0.00133321 |
| AT4G14905.1 | | protein_coding | Galactose oxidase/kelch r | 98 | 103 | 118 | 139 | 122 | 111 | 206 | 253 | 302 | 302 | 324 | 302 | 346.15 | 254 | 309 | 0.427 | -1.226 | 5.92E-17 |
| AT2G33680.1 | PCMP-E19 | protein_coding | Tetratricopeptide repeat | 53 | 36 | 27 | 25 | 29 | 21 | 111 | 88 | 69 | 54 | 77 | 57 | 81.61 | 89 | 63 | 0.427 | -1.227 | 2.08E-07 |
| AT5G61800.1 | PCMP-E8 | protein_coding | Pentatricopeptide repeat | 12 | 14 | 12 | 8 | 7 | 6 | 25 | 34 | 31 | 17 | 19 | 16 | 32 | 30 | 17 | 0.427 | -1.228 | 0.00010804 |
| AT5G59090.1 | ATSBT4.12 | protein_coding | subtilase 4.12 | 94 | 41 | 78 | 70 | 39 | 61 | 198 | 101 | 200 | 152 | 104 | 166 | 132.83 | 166 | 141 | 0.427 | -1.228 | 1.65E-07 |
| AT3G28660.1 | PCMP-E80 | protein_coding | Tetratricopeptide repeat | 44 | 31 | 25 | 30 | 34 | 17 | 92 | 76 | 64 | 65 | 90 | 46 | 76.01 | 77 | 67 | 0.427 | -1.228 | 4.70E-08 |
| AT4G13520.1 | SMAP1 | protein_coding | small acidic protein 1 | 521 | 519 | 476 | 545 | 414 | 444 | 1095 | 1275 | 1219 | 1182 | 1099 | 1207 | 1576.62 | 1196 | 1163 | 0.427 | -1.229 | 1.93E-25 |
| AT1G65120.2 | | protein_coding | Ubiquitin carboxyl-termir | 39 | 18 | 22 | 16 | 18 | 16 | 82 | 44 | 56 | 35 | 48 | 44 | 50.74 | 61 | 42 | 0.426 | -1.23 | 6.40E-05 |
| AT4G13400.1 | | protein_coding | 2-oxoglutarate (2OG) anc | 81 | 52 | 57 | 51 | 37 | 65 | 170 | 128 | 146 | 111 | 98 | 177 | 154.35 | 148 | 129 | 0.426 | -1.23 | 1.84E-10 |
| AT5G38520.2 | | protein_coding | alpha/beta-Hydrolases su | 2141 | 2261 | 1918 | 942 | 924 | 985 | 4501 | 5555 | 4910 | 2043 | 2453 | 2679 | 4589.77 | 4989 | 2392 | 0.426 | -1.23 | 4.82E-21 |
| AT5G06620.1 | ATXR4 | protein_coding | SET domain protein 38 | 23 | 19 | 16 | 20 | 17 | 19 | 48 | 47 | 41 | 43 | 45 | 52 | 82.76 | 45 | 47 | 0.426 | -1.232 | 1.85E-09 |
| AT1G03650.1 | | protein_coding | Acyl-CoA N-acyltransfera | 31 | 19 | 17 | 19 | 6 | 12 | 65 | 47 | 44 | 41 | 16 | 33 | 52.53 | 52 | 30 | 0.425 | -1.234 | 2.38E-07 |
| AT5G40160.1 | EMB506 | protein_coding | Ankyrin repeat family prc | 809 | 583 | 619 | 409 | 309 | 327 | 1701 | 1432 | 1585 | 887 | 820 | 889 | 1351.49 | 1573 | 865 | 0.424 | -1.236 | 9.69E-22 |
| AT1G71490.1 | PCMP-E67 | protein_coding | Tetratricopeptide repeat | 49 | 26 | 34 | 28 | 19 | 16 | 103 | 64 | 87 | 61 | 50 | 44 | 67.11 | 85 | 52 | 0.424 | -1.237 | 2.54E-07 |
| AT4G14150.1 | KIN12A | protein_coding | phragmoplast-associated | 179 | 183 | 172 | 159 | 151 | 153 | 376 | 450 | 440 | 345 | 401 | 416 | 348.66 | 422 | 387 | 0.424 | -1.239 | 3.19E-21 |
| AT1G65000.1 | | protein_coding | | 50 | 43 | 48 | 23 | 29 | 21 | 105 | 106 | 123 | 50 | 77 | 57 | 74.34 | 111 | 61 | 0.423 | -1.241 | 9.94E-09 |
| AT3G49142.1 | PCMP-H77 | protein_coding | Tetratricopeptide repeat | 42 | 31 | 22 | 21 | 28 | 24 | 88 | 76 | 56 | 46 | 74 | 65 | 58.6 | 73 | 62 | 0.423 | -1.242 | 3.23E-05 |
| AT5G63820.1 | | protein_coding | Protein of unknown funct | 17 | 19 | 54 | 8 | 12 | 23 | 36 | 47 | 138 | 17 | 32 | 63 | 54 | 74 | 37 | 0.422 | -1.244 | 0.00505325 |
| AT4G14620.1 | | protein_coding | Protein of unknown funct | 111 | 166 | 168 | 123 | 104 | 142 | 233 | 408 | 430 | 267 | 276 | 386 | 310.95 | 357 | 310 | 0.422 | -1.244 | 4.95E-10 |

| | | | | | | | | | | | | | | | | | | | | | |
|---|---|---|---|---|---|---|---|---|---|---|---|---|---|---|---|---|---|---|---|---|---|
| AT4G13493.1 | MIR850A | mirna | MIR850a; miRNA | 258 | 227 | 241 | 164 | 186 | 123 | 542 | 558 | 617 | 356 | 494 | 334 | 503.6 | 572 | 395 | 0.422 | -1.244 | 1.48E-18 |
| AT4G15475.1 | FBL4 | protein_coding | F-box/RNI-like superfamil | 231 | 237 | 233 | 188 | 171 | 189 | 486 | 582 | 596 | 408 | 454 | 514 | 654.2 | 555 | 459 | 0.422 | -1.245 | 2.31E-25 |
| AT1G73260.1 | ATKTI1 | protein_coding | kunitz trypsin inhibitor 1 | 18 | 9 | 9 | 31 | 9 | 17 | 38 | 22 | 23 | 67 | 24 | 46 | 39.88 | 28 | 46 | 0.421 | -1.247 | 0.00173599 |
| AT2G21630.1 | | protein_coding | Sec23/Sec24 protein tran | 85 | 80 | 93 | 36 | 28 | 44 | 179 | 197 | 238 | 78 | 74 | 120 | 187.76 | 205 | 91 | 0.419 | -1.253 | 1.33E-15 |
| AT4G15540.1 | | protein_coding | EamA-like transporter far | 531 | 342 | 348 | 524 | 361 | 293 | 1116 | 840 | 891 | 1137 | 958 | 797 | 1551.81 | 949 | 964 | 0.419 | -1.254 | 5.50E-18 |
| AT5G65070.3 | MAF4 | protein_coding | K-box region and MADS-b | 43 | 49 | 43 | 13 | 12 | 12 | 90 | 120 | 110 | 28 | 32 | 33 | 54.12 | 107 | 31 | 0.419 | -1.256 | 0.00018193 |
| AT4G14220.1 | RHF1A | protein_coding | RING-H2 group F1A | 74 | 70 | 76 | 97 | 52 | 53 | 156 | 172 | 195 | 210 | 138 | 144 | 408.93 | 174 | 164 | 0.417 | -1.263 | 1.12E-17 |
| AT4G13670.1 | PTAC5 | protein_coding | plastid transcriptionally a | 839 | 740 | 596 | 596 | 520 | 530 | 1764 | 1818 | 1526 | 1293 | 1380 | 1441 | 2710.46 | 1703 | 1371 | 0.417 | -1.263 | 2.21E-26 |
| AT4G13992.1 | | protein_coding | Cysteine/Histidine-rich C1 | 16 | 19 | 11 | 13 | 13 | 5 | 34 | 47 | 28 | 28 | 35 | 14 | 35.65 | 36 | 26 | 0.416 | -1.264 | 3.08E-05 |
| AT4G02750.1 | PCMP-H24 | protein_coding | Tetratricopeptide repeat | 53 | 35 | 33 | 35 | 25 | 15 | 111 | 86 | 84 | 76 | 66 | 41 | 82.13 | 94 | 61 | 0.416 | -1.266 | 2.31E-09 |
| AT2G30766.1 | | protein_coding | | 30 | 73 | 63 | 32 | 31 | 30 | 63 | 179 | 161 | 69 | 82 | 82 | 82.84 | 134 | 78 | 0.413 | -1.274 | 0.00591292 |
| AT1G49720.2 | ABF1 | protein_coding | abscisic acid responsive e | 277 | 435 | 519 | 92 | 138 | 156 | 582 | 1069 | 1329 | 200 | 366 | 424 | 688.9 | 993 | 330 | 0.414 | -1.274 | 2.06E-07 |
| AT1G76930.1 | ATEXT4 | protein_coding | extensin 4 | 141 | 31 | 26 | 163 | 52 | 30 | 296 | 76 | 67 | 354 | 138 | 82 | 997.5 | 146 | 191 | 0.413 | -1.275 | 0.00315521 |
| AT2G07675.1 | | protein_coding | Ribosomal protein S12/S2 | 700 | 669 | 598 | 749 | 686 | 593 | 1471 | 1644 | 1531 | 1625 | 1821 | 1613 | 811.97 | 1549 | 1686 | 0.413 | -1.276 | 0.00142334 |
| AT4G15080.1 | PAT19 | protein_coding | DHHC-type zinc finger far | 596 | 521 | 493 | 548 | 439 | 436 | 1253 | 1280 | 1262 | 1189 | 1165 | 1186 | 1494.71 | 1265 | 1180 | 0.413 | -1.276 | 1.08E-29 |
| AT1G61460.1 | | protein_coding | S-locus protein kinase, pu | 37 | 38 | 23 | 26 | 18 | 16 | 78 | 93 | 59 | 56 | 48 | 44 | 67.28 | 77 | 49 | 0.412 | -1.278 | 3.29E-08 |
| AT5G58670.1 | PLC1 | protein_coding | phospholipase C1 | 98 | 119 | 74 | 60 | 63 | 47 | 206 | 292 | 189 | 130 | 167 | 128 | 212.16 | 229 | 142 | 0.412 | -1.278 | 1.41E-13 |
| AT2G40720.1 | PCMP-E26 | protein_coding | Tetratricopeptide repeat | 52 | 52 | 46 | 33 | 27 | 38 | 109 | 128 | 118 | 72 | 72 | 103 | 91.7 | 118 | 82 | 0.412 | -1.279 | 1.26E-08 |
| AT4G05590.2 | MPC3 | protein_coding | | 35 | 18 | 15 | 16 | 10 | 14 | 74 | 44 | 38 | 35 | 27 | 38 | 44.42 | 52 | 33 | 0.411 | -1.282 | 3.82E-06 |
| AT3G53360.1 | PCMP-E86 | protein_coding | Tetratricopeptide repeat | 52 | 32 | 33 | 37 | 29 | 19 | 109 | 79 | 84 | 80 | 77 | 52 | 87.27 | 91 | 70 | 0.411 | -1.282 | 5.97E-09 |
| AT1G31690.1 | | protein_coding | Copper amine oxidase far | 131 | 82 | 44 | 26 | 17 | 6 | 275 | 201 | 113 | 56 | 45 | 16 | 156.31 | 196 | 39 | 0.411 | -1.283 | 2.99E-05 |
| AT5G64100.1 | PER69 | protein_coding | Peroxidase superfamily p | 93 | 42 | 18 | 45 | 19 | 27 | 195 | 103 | 46 | 98 | 50 | 73 | 113.18 | 115 | 74 | 0.411 | -1.284 | 0.00258174 |
| AT5G66631.1 | | protein_coding | Tetratricopeptide repeat | 42 | 21 | 21 | 18 | 14 | 17 | 88 | 52 | 54 | 39 | 37 | 46 | 56.57 | 65 | 41 | 0.411 | -1.284 | 1.03E-07 |
| AT4G39952.1 | PCMP-E98 | protein_coding | Pentatricopeptide repeat | 50 | 37 | 24 | 31 | 25 | 21 | 105 | 91 | 61 | 67 | 66 | 57 | 78.16 | 86 | 63 | 0.41 | -1.286 | 3.23E-08 |
| AT1G13410.1 | | protein_coding | Tetratricopeptide repeat | 46 | 22 | 32 | 28 | 19 | 18 | 97 | 54 | 82 | 61 | 50 | 49 | 67 | 78 | 53 | 0.409 | -1.289 | 1.59E-07 |
| AT3G08000.1 | | protein_coding | RNA-binding (RRM/RBD/ | 45 | 29 | 40 | 26 | 20 | 18 | 95 | 71 | 102 | 56 | 53 | 49 | 100.52 | 89 | 53 | 0.409 | -1.289 | 4.81E-14 |
| AT4G14700.1 | ORC1A | protein_coding | origin recognition comple | 35 | 39 | 42 | 39 | 24 | 24 | 74 | 96 | 108 | 85 | 64 | 65 | 74.71 | 93 | 71 | 0.408 | -1.292 | 3.35E-10 |
| AT3G23880.1 | | protein_coding | F-box and associated inte | 118 | 124 | 143 | 69 | 69 | 85 | 248 | 305 | 366 | 150 | 183 | 231 | 209.97 | 306 | 188 | 0.408 | -1.292 | 2.46E-13 |
| AT4G12780.1 | | protein_coding | Chaperone DnaJ-domain | 728 | 629 | 610 | 602 | 516 | 552 | 1530 | 1545 | 1562 | 1306 | 1370 | 1501 | 1747.07 | 1546 | 1392 | 0.408 | -1.292 | 5.84E-30 |
| AT5G20030.1 | | protein_coding | Plant Tudor-like RNA-bin | 69 | 53 | 66 | 21 | 20 | 26 | 145 | 130 | 169 | 46 | 53 | 71 | 88.77 | 148 | 57 | 0.406 | -1.3 | 2.59E-09 |
| AT2G43700.1 | LECRK54 | protein_coding | Concanavalin A-like lectir | 45 | 27 | 27 | 18 | 9 | 7 | 95 | 66 | 69 | 39 | 24 | 19 | 62.01 | 77 | 27 | 0.406 | -1.301 | 5.11E-08 |
| AT2G42530.1 | COR15B | protein_coding | cold regulated 15b | 9 | 42 | 50 | 8 | 24 | 12 | 19 | 103 | 128 | 17 | 64 | 33 | 89.59 | 83 | 38 | 0.405 | -1.303 | 0.00530681 |
| AT5G55450.1 | | protein_coding | Bifunctional inhibitor/lipi | 22 | 10 | 7 | 15 | 8 | 6 | 46 | 25 | 18 | 33 | 21 | 16 | 35.58 | 30 | 23 | 0.405 | -1.304 | 0.00021317 |
| AT4G14870.1 | SECE1 | protein_coding | secE/sec61-gamma prote | 478 | 551 | 464 | 348 | 284 | 401 | 1005 | 1354 | 1188 | 755 | 754 | 1090 | 2067.57 | 1182 | 866 | 0.405 | -1.306 | 2.39E-18 |
| AT5G05380.1 | PRA1B3 | protein_coding | prenylated RAB acceptor | 3 | 13 | 8 | 2 | 3 | 8 | 6 | 32 | 20 | 4 | 8 | 22 | 45.49 | 19 | 11 | 0.403 | -1.31 | 0.00046067 |
| AT2G23040.1 | | protein_coding | | 14 | 10 | 7 | 4 | 5 | 3 | 29 | 25 | 18 | 9 | 13 | 8 | 25.93 | 24 | 10 | 0.401 | -1.317 | 1.55E-05 |
| AT3G01580.1 | PCMP-E87 | protein_coding | Tetratricopeptide repeat | 26 | 21 | 18 | 13 | 11 | 11 | 55 | 52 | 46 | 28 | 29 | 30 | 40.95 | 51 | 29 | 0.401 | -1.317 | 1.70E-07 |
| AT5G48250.1 | COL10 | protein_coding | B-box type zinc finger pro | 265 | 288 | 357 | 146 | 69 | 118 | 557 | 708 | 914 | 317 | 183 | 321 | 611.71 | 726 | 274 | 0.401 | -1.318 | 3.79E-13 |
| AT2G18670.1 | ATL56 | protein_coding | RING/U-box superfamily | 77 | 100 | 107 | 33 | 24 | 30 | 162 | 246 | 274 | 72 | 64 | 82 | 178.7 | 227 | 73 | 0.401 | -1.319 | 1.01E-10 |
| AT4G39530.1 | PCMP-E52 | protein_coding | Tetratricopeptide repeat | 47 | 27 | 34 | 24 | 42 | 22 | 99 | 66 | 87 | 52 | 112 | 60 | 75.94 | 84 | 75 | 0.401 | -1.32 | 5.03E-08 |
| AT4G15030.2 | | protein_coding | | 72 | 69 | 78 | 43 | 36 | 32 | 151 | 170 | 200 | 93 | 96 | 87 | 329.85 | 174 | 92 | 0.4 | -1.321 | 4.30E-22 |
| AT4G14330.1 | | protein_coding | P-loop containing nucleo | 355 | 282 | 293 | 237 | 296 | 186 | 746 | 693 | 750 | 514 | 786 | 506 | 623 | 730 | 602 | 0.4 | -1.322 | 9.64E-19 |
| AT4G14270.1 | CID2 | protein_coding | | 260 | 406 | 469 | 159 | 241 | 372 | 547 | 997 | 1201 | 345 | 640 | 1012 | 1209.35 | 915 | 666 | 0.4 | -1.323 | 3.00E-06 |
| AT3G62270.1 | BOR2 | protein_coding | HCO3- transporter family | 121 | 77 | 58 | 99 | 97 | 42 | 254 | 189 | 148 | 215 | 258 | 114 | 190.89 | 197 | 196 | 0.4 | -1.323 | 2.69E-07 |
| AT5G56870.1 | BGAL4 | protein_coding | beta-galactosidase 4 | 389 | 565 | 1059 | 217 | 153 | 388 | 818 | 1388 | 2711 | 471 | 406 | 1055 | 766.82 | 1639 | 644 | 0.399 | -1.324 | 0.00020519 |
| AT2G19450.1 | DGAT1 | protein_coding | membrane bound O-acyl | 171 | 165 | 199 | 68 | 36 | 81 | 359 | 405 | 509 | 147 | 96 | 220 | 379.77 | 424 | 154 | 0.399 | -1.325 | 1.67E-14 |
| AT2G23985.2 | | protein_coding | | 106 | 91 | 77 | 50 | 29 | 34 | 223 | 224 | 197 | 108 | 77 | 92 | 155.03 | 215 | 92 | 0.399 | -1.326 | 7.79E-14 |
| AT3G60330.1 | AHA7 | protein_coding | H(+)-ATPase 7 | 39 | 11 | 18 | 49 | 18 | 16 | 82 | 27 | 46 | 106 | 48 | 44 | 57.76 | 52 | 66 | 0.398 | -1.331 | 6.26E-05 |
| AT3G58590.1 | | protein_coding | Pentatricopeptide repeat | 57 | 32 | 24 | 36 | 28 | 35 | 120 | 79 | 61 | 78 | 74 | 95 | 87.11 | 87 | 82 | 0.397 | -1.332 | 1.64E-09 |
| AT5G61380.1 | APRR1 | protein_coding | CCT motif -containing res | 510 | 624 | 920 | 337 | 256 | 303 | 1072 | 1533 | 2355 | 731 | 680 | 824 | 1146.58 | 1653 | 745 | 0.397 | -1.332 | 8.10E-11 |
| AT1G29090.1 | | protein_coding | Cysteine proteinases supe | 62 | 21 | 18 | 48 | 18 | 22 | 130 | 52 | 46 | 104 | 48 | 60 | 70.69 | 76 | 71 | 0.396 | -1.336 | 0.00186109 |
| AT4G13210.2 | | protein_coding | Pectin lyase-like superfan | 16 | 11 | 18 | 16 | 16 | 13 | 34 | 27 | 46 | 35 | 42 | 35 | 44.99 | 36 | 37 | 0.395 | -1.341 | 1.35E-06 |
| AT3G19508.1 | | protein_coding | | 45 | 39 | 35 | 52 | 33 | 30 | 95 | 96 | 90 | 113 | 88 | 82 | 116.4 | 94 | 94 | 0.395 | -1.342 | 8.35E-15 |
| AT1G21930.1 | | protein_coding | | 177 | 142 | 173 | 68 | 81 | 73 | 372 | 349 | 443 | 147 | 215 | 199 | 316.41 | 388 | 187 | 0.393 | -1.347 | 2.68E-17 |
| AT1G33170.1 | | protein_coding | S-adenosyl-L-methionine- | 580 | 573 | 461 | 194 | 205 | 142 | 1219 | 1408 | 1180 | 421 | 544 | 386 | 739.67 | 1269 | 450 | 0.393 | -1.348 | 5.95E-18 |
| AT4G14540.1 | NFYB3 | protein_coding | nuclear factor Y, subunit | 127 | 108 | 85 | 89 | 79 | 95 | 267 | 265 | 218 | 193 | 210 | 258 | 842.14 | 250 | 220 | 0.391 | -1.356 | 9.97E-28 |
| AT4G12850.1 | | protein_coding | Far-red impaired respons | 34 | 13 | 29 | 22 | 13 | 14 | 71 | 32 | 74 | 48 | 35 | 38 | 61.4 | 59 | 40 | 0.39 | -1.357 | 5.24E-08 |
| AT4G13030.1 | | protein_coding | P-loop containing nucleo | 37 | 43 | 56 | 26 | 23 | 58 | 78 | 106 | 143 | 56 | 61 | 158 | 111.33 | 109 | 92 | 0.39 | -1.358 | 6.75E-06 |
| AT1G64320.1 | | protein_coding | myosin heavy chain-relat | 12 | 5 | 7 | 3 | 5 | 3 | 25 | 12 | 18 | 7 | 13 | 8 | 27.23 | 18 | 9 | 0.39 | -1.358 | 6.14E-06 |
| AT4G14147.2 | ARPC4 | protein_coding | protein binding | 105 | 88 | 114 | 82 | 66 | 60 | 221 | 216 | 292 | 178 | 175 | 163 | 245.58 | 243 | 172 | 0.39 | -1.358 | 6.34E-22 |
| AT5G21970.1 | | protein_coding | Ubiquitin carboxyl-termir | 58 | 48 | 46 | 38 | 17 | 28 | 122 | 118 | 118 | 82 | 45 | 76 | 130.93 | 119 | 68 | 0.389 | -1.361 | 1.20E-13 |
| AT4G13575.1 | | protein_coding | | 1053 | 1092 | 927 | 506 | 585 | 468 | 2213 | 2683 | 2373 | 1098 | 1553 | 1273 | 1921.8 | 2423 | 1308 | 0.388 | -1.366 | 1.12E-20 |
| AT4G13020.3 | MHK | protein_coding | Protein kinase superfamil | 687 | 660 | 631 | 517 | 465 | 409 | 1444 | 1621 | 1615 | 1121 | 1234 | 1112 | 1656.35 | 1560 | 1156 | 0.388 | -1.366 | 4.63E-30 |
| AT4G15233.2 | | protein_coding | ABC-2 and Plant PDR ABC | 463 | 124 | 204 | 436 | 126 | 184 | 973 | 305 | 522 | 946 | 335 | 500 | 469.86 | 600 | 594 | 0.387 | -1.369 | 5.77E-05 |
| AT2G20815.1 | QWRF3 | protein_coding | Family of unknown functi | 40 | 46 | 36 | 19 | 14 | 15 | 84 | 113 | 92 | 41 | 37 | 41 | 68.9 | 96 | 40 | 0.386 | -1.373 | 2.51E-11 |
| AT1G74400.1 | PCMP-E68 | protein_coding | Tetratricopeptide repeat | 23 | 17 | 15 | 11 | 18 | 14 | 48 | 42 | 38 | 24 | 48 | 38 | 37.36 | 43 | 37 | 0.386 | -1.374 | 5.16E-06 |
| AT5G45470.1 | | protein_coding | Protein of unknown funct | 816 | 647 | 595 | 328 | 282 | 237 | 1715 | 1590 | 1523 | 711 | 749 | 645 | 978.23 | 1609 | 702 | 0.386 | -1.375 | 4.46E-26 |
| AT4G12470.1 | AZI1 | protein_coding | azelaic acid induced 1 | 432 | 73 | 79 | 430 | 54 | 140 | 908 | 179 | 202 | 933 | 143 | 381 | 1277.17 | 430 | 486 | 0.385 | -1.376 | 0.00757383 |
| AT4G15140.1 | | protein_coding | | 95 | 86 | 84 | 85 | 76 | 76 | 200 | 211 | 215 | 184 | 202 | 207 | 244.36 | 209 | 198 | 0.385 | -1.378 | 2.92E-25 |
| AT1G34380.2 | | protein_coding | 5'-3' exonuclease family p | 136 | 86 | 93 | 55 | 72 | 35 | 286 | 211 | 238 | 119 | 191 | 95 | 220.62 | 245 | 135 | 0.384 | -1.38 | 8.64E-15 |
| AT5G06400.1 | | protein_coding | Pentatricopeptide repeat | 48 | 31 | 47 | 34 | 25 | 23 | 101 | 76 | 120 | 74 | 66 | 63 | 69.1 | 99 | 68 | 0.384 | -1.381 | 7.95E-10 |
| AT3G07650.1 | COL9 | protein_coding | CONSTANS-like 9 | 123 | 138 | 209 | 49 | 60 | 95 | 259 | 339 | 535 | 106 | 159 | 258 | 309.95 | 378 | 174 | 0.382 | -1.387 | 6.96E-08 |
| AT4G14570.1 | | protein_coding | acylaminoacyl-peptidase- | 972 | 660 | 695 | 670 | 536 | 460 | 2043 | 1621 | 1779 | 1453 | 1423 | 1251 | 1977.68 | 1814 | 1376 | 0.382 | -1.387 | 4.60E-30 |
| AT1G78990.1 | | protein_coding | HXXXD-type acyl-transfer | 23 | 28 | 17 | 9 | 15 | 9 | 48 | 69 | 44 | 20 | 40 | 24 | 90.64 | 54 | 28 | 0.381 | -1.391 | 1.01E-09 |
| AT4G14600.1 | | protein_coding | Target SNARE coiled-coil | 108 | 92 | 85 | 75 | 55 | 61 | 227 | 226 | 218 | 163 | 146 | 166 | 244.2 | 224 | 158 | 0.381 | -1.392 | 2.82E-27 |
| AT1G61490.1 | | protein_coding | S-locus lectin protein kina | 56 | 30 | 39 | 23 | 14 | 17 | 118 | 74 | 100 | 50 | 37 | 46 | 90.04 | 97 | 44 | 0.38 | -1.396 | 1.74E-10 |
| AT3G20730.1 | PCMP-E94 | protein_coding | Tetratricopeptide repeat | 23 | 16 | 19 | 20 | 23 | 12 | 48 | 39 | 49 | 43 | 61 | 33 | 46.83 | 45 | 46 | 0.378 | -1.403 | 1.47E-06 |
| AT5G20790.1 | | protein_coding | | 16 | 3 | 2 | 28 | 5 | 2 | 34 | 7 | 5 | 61 | 13 | 5 | 81.55 | 15 | 26 | 0.378 | -1.404 | 0.00686746 |
| AT3G22121.1 | | other_rna | other RNA | 9 | 9 | 5 | 12 | 0 | 3 | 19 | 22 | 13 | 26 | 0 | 8 | 42.33 | 18 | 11 | 0.375 | -1.416 | 0.00129131 |
| AT1G52450.1 | | protein_coding | Ubiquitin carboxyl-termir | 35 | 22 | 31 | 37 | 33 | 26 | 74 | 54 | 79 | 80 | 88 | 71 | 87.63 | 69 | 80 | 0.375 | -1.416 | 4.74E-09 |
| AT4G14230.1 | CBSDUF2 | protein_coding | CBS domain-containing p | 785 | 827 | 834 | 494 | 524 | 535 | 1650 | 2032 | 2135 | 1072 | 1391 | 1455 | 2234.19 | 1939 | 1306 | 0.374 | -1.419 | 2.10E-23 |
| AT5G08305.1 | PCMP-E105 | protein_coding | Pentatricopeptide repeat | 44 | 29 | 31 | 13 | 22 | 15 | 92 | 71 | 79 | 28 | 58 | 41 | 58.93 | 81 | 42 | 0.374 | -1.42 | 3.62E-09 |
| AT1G43980.1 | PCMP-E58 | protein_coding | Tetratricopeptide repeat | 37 | 27 | 21 | 21 | 13 | 22 | 78 | 66 | 54 | 46 | 35 | 60 | 53.56 | 66 | 47 | 0.373 | -1.421 | 2.04E-07 |
| AT5G45840.2 | | protein_coding | Leucine-rich repeat prote | 119 | 126 | 86 | 51 | 45 | 26 | 250 | 310 | 220 | 111 | 119 | 71 | 151.67 | 260 | 100 | 0.374 | -1.421 | 5.40E-12 |
| AT1G06440.1 | | protein_coding | Ubiquitin carboxyl-termir | 39 | 23 | 24 | 22 | 16 | 19 | 82 | 57 | 61 | 48 | 42 | 52 | 66.71 | 67 | 47 | 0.373 | -1.423 | 1.05E-11 |
| AT2G07751.1 | | protein_coding | NADH:ubiquinone/plasto | 2261 | 1896 | 2320 | 2772 | 2180 | 2141 | 4753 | 4658 | 5939 | 6013 | 5787 | 5822 | 2749.38 | 5117 | 5874 | 0.373 | -1.425 | 0.00879431 |
| AT5G49015.1 | | protein_coding | Expressed protein | 88 | 94 | 107 | 36 | 30 | 47 | 185 | 231 | 274 | 78 | 80 | 128 | 217.82 | 230 | 95 | 0.371 | -1.43 | 8.69E-18 |
| AT4G12840.1 | | protein_coding | Protein of unknown funct | 78 | 93 | 114 | 48 | 62 | 45 | 164 | 228 | 292 | 104 | 165 | 122 | 206.73 | 228 | 130 | 0.37 | -1.436 | 6.39E-15 |
| AT2G36980.1 | PCMP-E73 | protein_coding | Tetratricopeptide repeat | 30 | 29 | 27 | 14 | 8 | 13 | 63 | 71 | 69 | 30 | 21 | 35 | 52.08 | 68 | 29 | 0.369 | -1.437 | 4.46E-07 |
| AT5G42450.1 | PCMP-E102 | protein_coding | Pentatricopeptide repeat | 40 | 26 | 31 | 20 | 11 | 13 | 84 | 64 | 79 | 43 | 29 | 35 | 53.95 | 76 | 36 | 0.369 | -1.44 | 6.89E-07 |
| AT3G60540.1 | | protein_coding | Preprotein translocase Se | 33 | 16 | 14 | 17 | 7 | 10 | 69 | 39 | 36 | 37 | 19 | 27 | 48.99 | 48 | 28 | 0.369 | -1.44 | 2.37E-08 |
| AT4G15236.1 | ABCG43 | protein_coding | ABC-2 and Plant PDR ABC | 57 | 39 | 42 | 80 | 35 | 39 | 120 | 96 | 108 | 174 | 93 | 106 | 105.18 | 108 | 124 | 0.368 | -1.442 | 8.85E-11 |
| AT4G14580.1 | CIPK4 | protein_coding | CBL-interacting protein ki | 27 | 17 | 15 | 22 | 19 | 12 | 57 | 42 | 38 | 48 | 50 | 33 | 92.74 | 46 | 44 | 0.368 | -1.442 | 1.51E-11 |
| AT4G12970.1 | EPFL9 | protein_coding | stomagen | 159 | 142 | 109 | 110 | 106 | 86 | 334 | 349 | 279 | 239 | 281 | 234 | 521.23 | 321 | 251 | 0.368 | -1.444 | 5.94E-24 |
| AT1G06510.1 | | protein_coding | | 386 | 335 | 275 | 122 | 117 | 124 | 811 | 823 | 704 | 265 | 311 | 337 | 563.2 | 779 | 304 | 0.368 | -1.444 | 9.32E-30 |
| AT3G53010.1 | | protein_coding | Domain of unknown func | 14 | 8 | 7 | 7 | 1 | 5 | 29 | 20 | 18 | 15 | 3 | 14 | 24.93 | 22 | 11 | 0.366 | -1.452 | 7.79E-06 |
| AT1G34160.1 | PCMP-H68 | protein_coding | Tetratricopeptide repeat | 29 | 20 | 27 | 24 | 16 | 13 | 61 | 49 | 69 | 52 | 42 | 35 | 81.02 | 60 | 43 | 0.362 | -1.465 | 4.04E-12 |
| AT4G13460.1 | SUVH9 | protein_coding | SU(VAR)3-9 homolog 9 | 502 | 398 | 428 | 360 | 292 | 354 | 1055 | 978 | 1096 | 781 | 775 | 963 | 1137.06 | 1043 | 840 | 0.361 | -1.47 | 2.96E-33 |
| AT1G06140.1 | PCMP-E61 | protein_coding | Pentatricopeptide repeat | 23 | 19 | 11 | 9 | 13 | 6 | 48 | 47 | 28 | 20 | 35 | 16 | 37.62 | 41 | 24 | 0.361 | -1.472 | 1.93E-07 |
| AT5G44440.1 | | protein_coding | FAD-binding Berberine fa | 47 | 57 | 128 | 48 | 115 | 198 | 99 | 140 | 328 | 104 | 305 | 538 | 187.16 | 189 | 316 | 0.36 | -1.475 | 0.00223541 |
| AT5G23090.2 | DR1 | protein_coding | nuclear factor Y, subunit | 122 | 85 | 88 | 43 | 39 | 37 | 256 | 209 | 225 | 93 | 104 | 101 | 156.31 | 230 | 99 | 0.359 | -1.477 | 2.30E-15 |
| AT5G49110.2 | | protein_coding | | 73 | 57 | 58 | 55 | 50 | 30 | 153 | 140 | 148 | 119 | 133 | 82 | 115.25 | 147 | 111 | 0.358 | -1.481 | 5.09E-13 |
| AT2G03880.1 | PCMP-H44 | protein_coding | Pentatricopeptide repeat | 49 | 27 | 27 | 29 | 16 | 20 | 103 | 66 | 69 | 63 | 42 | 54 | 52.04 | 79 | 53 | 0.358 | -1.483 | 3.90E-07 |
| AT5G41650.1 | | protein_coding | Lactoylglutathione lyase / | 25 | 11 | 20 | 15 | 9 | 21 | 53 | 27 | 51 | 33 | 24 | 57 | 54.9 | 44 | 38 | 0.358 | -1.483 | 4.66E-09 |
| AT4G27852.1 | | other_rna | | 18 | 9 | 10 | 13 | 12 | 11 | 38 | 22 | 26 | 28 | 32 | 30 | 38.68 | 29 | 30 | 0.357 | -1.486 | 7.63E-09 |
| AT1G32415.1 | PCMP-E56 | protein_coding | pentatricopeptide (PPR) r | 26 | 26 | 14 | 10 | 10 | 5 | 55 | 64 | 36 | 22 | 27 | 14 | 54.29 | 52 | 21 | 0.357 | -1.487 | 1.31E-09 |
| AT5G45360.1 | SKIP31 | protein_coding | F-box family protein | 416 | 350 | 404 | 159 | 150 | 146 | 874 | 860 | 1034 | 345 | 398 | 397 | 593 | 923 | 380 | 0.357 | -1.487 | 4.49E-34 |
| AT3G59480.1 | | protein_coding | pfkB-like carbohydrate ki | 12 | 30 | 16 | 12 | 13 | 16 | 25 | 74 | 41 | 26 | 35 | 44 | 48.88 | 47 | 35 | 0.356 | -1.488 | 0.00015356 |
| AT1G77360.1 | | protein_coding | Tetratricopeptide repeat | 28 | 18 | 25 | 18 | 14 | 12 | 59 | 44 | 64 | 39 | 37 | 33 | 43.1 | 56 | 36 | 0.354 | -1.497 | 2.65E-06 |
| AT1G61420.1 | | protein_coding | S-locus lectin protein kina | 63 | 41 | 41 | 48 | 32 | 29 | 132 | 101 | 105 | 104 | 85 | 79 | 113.35 | 113 | 89 | 0.354 | -1.497 | 3.44E-14 |
| AT1G28690.1 | PCMP-E34 | protein_coding | Tetratricopeptide repeat | 30 | 27 | 22 | 18 | 14 | 15 | 63 | 66 | 56 | 39 | 37 | 41 | 43.56 | 62 | 39 | 0.354 | -1.499 | 1.23E-07 |
| AT3G23020.1 | | protein_coding | Tetratricopeptide repeat | 146 | 76 | 91 | 44 | 41 | 48 | 307 | 187 | 233 | 95 | 109 | 131 | 155.75 | 242 | 112 | 0.353 | -1.504 | 3.35E-13 |
| AT1G56130.1 | | protein_coding | Leucine-rich repeat trans | 38 | 36 | 39 | 29 | 25 | 24 | 80 | 88 | 100 | 63 | 66 | 65 | 84.71 | 89 | 65 | 0.352 | -1.505 | 8.57E-14 |
| AT1G09220.1 | PCMP-E25 | protein_coding | Pentatricopeptide repeat | 24 | 19 | 17 | 22 | 14 | 9 | 50 | 47 | 44 | 48 | 37 | 24 | 43.02 | 47 | 36 | 0.352 | -1.508 | 6.92E-08 |
| AT4G32340.1 | | protein_coding | Tetratricopeptide repeat | 595 | 919 | 888 | 180 | 203 | 227 | 1251 | 2258 | 2273 | 390 | 539 | 617 | 1704.46 | 1927 | 515 | 0.351 | -1.511 | 2.72E-14 |
| AT3G24000.1 | PCMP-H87 | protein_coding | Tetratricopeptide repeat | 52 | 37 | 34 | 20 | 23 | 34 | 109 | 91 | 87 | 43 | 61 | 92 | 93.51 | 96 | 65 | 0.349 | -1.518 | 5.09E-11 |
| AT4G14716.1 | ARD2 | protein_coding | acireductone dioxygenase | 575 | 540 | 540 | 451 | 383 | 337 | 1209 | 1327 | 1382 | 978 | 1017 | 916 | 1795.18 | 1306 | 970 | 0.349 | -1.52 | 7.40E-38 |
| AT1G09280.1 | STR6 | protein_coding | | 226 | 185 | 211 | 143 | 123 | 169 | 475 | 455 | 540 | 310 | 327 | 460 | 382.12 | 490 | 366 | 0.348 | -1.521 | 8.36E-24 |
| AT2G29300.2 | | protein_coding | NAD(P)-binding Rossman | 175 | 148 | 193 | 40 | 38 | 41 | 368 | 364 | 494 | 87 | 101 | 111 | 279.84 | 409 | 100 | 0.348 | -1.522 | 7.51E-16 |
| AT1G65110.1 | | protein_coding | Ubiquitin carboxyl-termir | 18 | 28 | 11 | 9 | 9 | 7 | 38 | 69 | 28 | 20 | 24 | 19 | 33.03 | 45 | 21 | 0.348 | -1.523 | 2.08E-06 |
| AT5G49410.1 | | protein_coding | | 33 | 21 | 31 | 27 | 16 | 11 | 69 | 52 | 79 | 59 | 42 | 30 | 81.74 | 67 | 44 | 0.348 | -1.523 | 1.54E-12 |
| AT4G33170.1 | PCMP-H53 | protein_coding | Tetratricopeptide repeat | 54 | 28 | 29 | 22 | 11 | 18 | 114 | 69 | 74 | 48 | 29 | 49 | 64.72 | 86 | 42 | 0.346 | -1.533 | 6.23E-07 |
| AT2G01730.1 | CPSF73-II | protein_coding | cleavage and polyadenyla | 35 | 26 | 15 | 22 | 17 | 16 | 74 | 64 | 38 | 48 | 45 | 44 | 63.46 | 59 | 46 | 0.345 | -1.533 | 4.42E-08 |
| AT5G46460.1 | PCMP-H49 | protein_coding | Pentatricopeptide repeat | 71 | 54 | 31 | 41 | 32 | 32 | 149 | 133 | 79 | 89 | 85 | 87 | 76.62 | 120 | 87 | 0.343 | -1.543 | 5.36E-10 |
| AT1G17665.1 | | protein_coding | | 61 | 59 | 61 | 15 | 18 | 21 | 128 | 145 | 156 | 33 | 48 | 57 | 115.67 | 143 | 46 | 0.342 | -1.55 | 1.51E-16 |
| AT5G39680.1 | EMB2744 | protein_coding | Pentatricopeptide repeat | 29 | 18 | 30 | 21 | 24 | 14 | 61 | 44 | 77 | 46 | 64 | 38 | 45.05 | 61 | 49 | 0.34 | -1.558 | 1.10E-07 |
| AT3G11460.1 | PCMP-H52 | protein_coding | Pentatricopeptide repeat | 28 | 36 | 19 | 21 | 17 | 16 | 59 | 88 | 49 | 46 | 45 | 44 | 58.21 | 65 | 45 | 0.34 | -1.558 | 6.58E-09 |
| AT3G15930.1 | PCMP-E51 | protein_coding | Pentatricopeptide repeat | 28 | 22 | 25 | 15 | 9 | 25 | 59 | 54 | 64 | 33 | 24 | 68 | 44.16 | 59 | 42 | 0.339 | -1.559 | 2.81E-07 |
| AT4G14723.1 | EPFL4 | protein_coding | | 0 | 3 | 5 | 4 | 0 | 2 | 0 | 7 | 13 | 9 | 0 | 5 | 21.52 | 7 | 5 | 0.339 | -1.561 | 2.70E-05 |
| AT4G13650.1 | PCMP-H42 | protein_coding | Pentatricopeptide repeat | 183 | 123 | 138 | 119 | 91 | 50 | 385 | 302 | 353 | 258 | 242 | 136 | 286.35 | 347 | 212 | 0.339 | -1.562 | 4.44E-20 |
| AT4G13918.1 | | other_rna | | 75 | 96 | 126 | 70 | 58 | 69 | 158 | 236 | 323 | 152 | 154 | 188 | 216.1 | 239 | 165 | 0.337 | -1.57 | 2.33E-14 |
| AT3G16610.1 | PCMP-E91 | protein_coding | pentatricopeptide (PPR) r | 30 | 8 | 18 | 11 | 12 | 6 | 63 | 20 | 46 | 24 | 32 | 16 | 32.64 | 43 | 24 | 0.337 | -1.571 | 1.21E-05 |
| AT1G62290.1 | APA2 | protein_coding | Saposin-like aspartyl prot | 236 | 224 | 201 | 230 | 205 | 156 | 496 | 550 | 515 | 499 | 544 | 424 | 493.47 | 520 | 489 | 0.337 | -1.571 | 1.40E-29 |
| AT4G19830.1 | FKBP17-1 | protein_coding | FKBP-like peptidyl-prolyl | 260 | 211 | 216 | 79 | 70 | 78 | 547 | 518 | 553 | 171 | 186 | 212 | 637.84 | 539 | 190 | 0.336 | -1.572 | 4.87E-35 |
| AT3G21470.1 | PCMP-E29 | protein_coding | Pentatricopeptide repeat | 19 | 19 | 13 | 8 | 3 | 5 | 40 | 47 | 33 | 17 | 8 | 14 | 28.08 | 40 | 13 | 0.336 | -1.573 | 8.08E-07 |
| AT4G14850.1 | LOI1 | protein_coding | Pentatricopeptide repeat | 64 | 60 | 44 | 55 | 52 | 37 | 135 | 147 | 113 | 119 | 138 | 101 | 167.37 | 132 | 119 | 0.336 | -1.574 | 6.69E-26 |
| AT4G15165.1 | PAC2 | protein_coding | N-terminal nucleophile a | 20 | 15 | 15 | 15 | 15 | 14 | 42 | 37 | 38 | 33 | 40 | 38 | 54.84 | 39 | 37 | 0.336 | -1.575 | 1.15E-09 |
| AT4G05210.1 | LPXD1 | protein_coding | Trimeric LpxA-like enzym | 12 | 9 | 16 | 6 | 9 | 7 | 25 | 22 | 41 | 13 | 24 | 19 | 28.19 | 29 | 19 | 0.335 | -1.579 | 1.59E-07 |
| AT1G03510.1 | PCMP-E3 | protein_coding | Tetratricopeptide repeat | 20 | 11 | 18 | 5 | 12 | 6 | 42 | 27 | 46 | 11 | 32 | 16 | 39.92 | 38 | 20 | 0.333 | -1.586 | 2.69E-06 |
| AT3G04020.1 | | protein_coding | | 70 | 56 | 52 | 43 | 28 | 23 | 147 | 138 | 133 | 93 | 74 | 63 | 132.99 | 139 | 77 | 0.333 | -1.587 | 6.42E-17 |
| AT3G01420.1 | DOX1 | protein_coding | Peroxidase superfamily p | 300 | 61 | 83 | 222 | 62 | 57 | 631 | 150 | 212 | 482 | 165 | 155 | 249.04 | 331 | 267 | 0.333 | -1.588 | 0.00173358 |
| AT2G33760.1 | PCMP-H6 | protein_coding | Pentatricopeptide repeat | 13 | 7 | 13 | 9 | 15 | 11 | 27 | 17 | 33 | 20 | 40 | 30 | 32.97 | 26 | 30 | 0.332 | -1.592 | 2.77E-07 |
| AT4G13950.1 | RALFL31 | protein_coding | ralf-like 31 | 66 | 80 | 40 | 54 | 44 | 46 | 139 | 197 | 102 | 117 | 117 | 125 | 204.5 | 146 | 120 | 0.331 | -1.597 | 4.15E-22 |
| AT4G22470.1 | | protein_coding | protease inhibitor/seed s | 29 | 2 | 1 | 12 | 1 | 0 | 61 | 5 | 3 | 26 | 3 | 0 | 55.98 | 23 | 10 | 0.33 | -1.6 | 0.00909602 |
| AT3G56550.1 | PCMP-H80 | protein_coding | Pentatricopeptide repeat | 20 | 13 | 18 | 15 | 7 | 8 | 42 | 32 | 46 | 33 | 19 | 22 | 34.22 | 40 | 25 | 0.33 | -1.601 | 1.45E-07 |

| | | | | | | | | | | | | | | | | | | | | | |
|---|---|---|---|---|---|---|---|---|---|---|---|---|---|---|---|---|---|---|---|---|---|
| AT2G03260.1 | PHO1-H2 | protein_coding | EXS (ERD1/XPR1/SYG1) fa | 16 | 11 | 15 | 11 | 8 | 5 | 34 | 27 | 38 | 24 | 21 | 14 | 34.97 | 33 | 20 | 0.329 | -1.602 | 1.61E-07 |
| AT3G05905.1 | | other_rna | other RNA | 117 | 154 | 168 | 172 | 31 | 16 | 246 | 378 | 430 | 373 | 82 | 44 | 157.38 | 351 | 166 | 0.328 | -1.607 | 0.00133413 |
| AT3G13440.1 | | protein_coding | S-adenosyl-L-methionine- | 176 | 165 | 133 | 52 | 47 | 36 | 370 | 405 | 340 | 113 | 125 | 98 | 275.9 | 372 | 112 | 0.327 | -1.614 | 1.11E-36 |
| AT5G17460.2 | | protein_coding | | 42 | 27 | 30 | 27 | 27 | 21 | 88 | 66 | 77 | 59 | 72 | 57 | 79.08 | 77 | 63 | 0.326 | -1.619 | 9.95E-14 |
| AT2G22450.1 | RIBA2 | protein_coding | riboflavin biosynthesis pr | 356 | 383 | 511 | 82 | 98 | 150 | 748 | 941 | 1308 | 178 | 260 | 408 | 687.05 | 999 | 282 | 0.324 | -1.627 | 4.93E-14 |
| AT1G07050.1 | | protein_coding | CCT motif family protein | 74 | 184 | 163 | 14 | 22 | 25 | 156 | 452 | 417 | 30 | 58 | 68 | 308.71 | 342 | 52 | 0.324 | -1.628 | 8.95E-09 |
| AT5G13230.1 | PCMP-H89 | protein_coding | Tetratricopeptide repeat | 32 | 21 | 30 | 25 | 21 | 15 | 67 | 52 | 77 | 54 | 56 | 41 | 44.83 | 65 | 50 | 0.32 | -1.642 | 1.34E-08 |
| AT3G18510.1 | | protein_coding | | 12 | 21 | 14 | 13 | 13 | 17 | 25 | 52 | 36 | 28 | 35 | 46 | 59.26 | 38 | 36 | 0.32 | -1.643 | 1.44E-09 |
| AT2G32770.3 | PAP13 | protein_coding | purple acid phosphatase | 29 | 32 | 18 | 14 | 21 | 10 | 61 | 79 | 46 | 30 | 56 | 27 | 42.49 | 62 | 38 | 0.319 | -1.647 | 2.05E-08 |
| AT3G56960.1 | PIP5K4 | protein_coding | phosphatidyl inositol mo | 43 | 29 | 25 | 17 | 15 | 15 | 90 | 71 | 64 | 37 | 40 | 41 | 92.83 | 75 | 39 | 0.319 | -1.648 | 2.84E-19 |
| AT2G29260.1 | | protein_coding | NAD(P)-binding Rossman | 51 | 15 | 14 | 29 | 10 | 18 | 107 | 37 | 36 | 63 | 27 | 49 | 74.91 | 60 | 46 | 0.319 | -1.65 | 5.44E-07 |
| AT2G36690.1 | | protein_coding | 2-oxoglutarate (2OG) and | 90 | 16 | 33 | 64 | 7 | 19 | 189 | 39 | 84 | 139 | 19 | 52 | 77.34 | 104 | 70 | 0.316 | -1.661 | 0.00134091 |
| AT5G28220.1 | | protein_coding | Protein prenylyltransfera | 130 | 113 | 109 | 62 | 46 | 38 | 273 | 278 | 279 | 134 | 122 | 103 | 209.44 | 277 | 120 | 0.314 | -1.673 | 4.93E-25 |
| AT1G19960.1 | | protein_coding | | 2 | 13 | 6 | 10 | 8 | 7 | 4 | 32 | 15 | 22 | 21 | 19 | 46.25 | 17 | 21 | 0.313 | -1.674 | 1.45E-09 |
| AT4G31070.1 | PCMP-E7 | protein_coding | Tetratricopeptide repeat | 42 | 17 | 25 | 18 | 14 | 11 | 88 | 42 | 64 | 39 | 37 | 30 | 46.06 | 65 | 35 | 0.313 | -1.678 | 1.92E-07 |
| AT3G15750.1 | | protein_coding | Essential protein Yae1, N- | 84 | 104 | 102 | 26 | 33 | 27 | 177 | 256 | 261 | 56 | 88 | 73 | 206.35 | 231 | 72 | 0.308 | -1.697 | 2.33E-16 |
| AT3G22450.1 | | protein_coding | Ribosomal L18p/L5e fami | 36 | 13 | 14 | 8 | 11 | 7 | 76 | 32 | 36 | 17 | 29 | 19 | 37.14 | 48 | 22 | 0.308 | -1.7 | 5.05E-05 |
| AT3G29330.2 | | protein_coding | | 20 | 29 | 35 | 13 | 10 | 14 | 42 | 71 | 90 | 28 | 27 | 38 | 53.69 | 68 | 31 | 0.307 | -1.705 | 1.71E-09 |
| AT4G13690.1 | | protein_coding | | 61 | 37 | 38 | 28 | 22 | 20 | 128 | 91 | 97 | 61 | 58 | 54 | 103.52 | 105 | 58 | 0.305 | -1.715 | 7.48E-19 |
| AT2G43620.1 | | protein_coding | Chitinase family protein | 86 | 15 | 12 | 151 | 13 | 15 | 181 | 37 | 31 | 328 | 35 | 41 | 104.63 | 83 | 135 | 0.304 | -1.717 | 0.00859617 |
| AT4G13810.1 | AtRLP47 | protein_coding | receptor like protein 47 | 138 | 138 | 109 | 74 | 60 | 40 | 290 | 339 | 279 | 161 | 159 | 109 | 333.09 | 303 | 143 | 0.298 | -1.749 | 1.06E-25 |
| AT4G04190.1 | | protein_coding | | 39 | 36 | 42 | 23 | 23 | 12 | 82 | 88 | 108 | 50 | 61 | 33 | 87.6 | 93 | 48 | 0.29 | -1.785 | 1.05E-16 |
| AT2G16870.1 | | protein_coding | Disease resistance protei | 49 | 31 | 32 | 19 | 21 | 10 | 103 | 76 | 82 | 41 | 56 | 27 | 47.65 | 87 | 41 | 0.286 | -1.807 | 1.95E-05 |
| AT3G47840.1 | PCMP-E43 | protein_coding | Tetratricopeptide repeat | 39 | 17 | 19 | 17 | 9 | 12 | 82 | 42 | 49 | 37 | 24 | 33 | 38.74 | 58 | 31 | 0.282 | -1.828 | 2.94E-08 |
| AT1G53330.1 | | protein_coding | Pentatricopeptide repeat | 40 | 24 | 23 | 14 | 12 | 19 | 84 | 59 | 59 | 30 | 32 | 52 | 45.32 | 67 | 38 | 0.28 | -1.837 | 2.91E-10 |
| AT4G15093.1 | LIGB | protein_coding | catalytic LigB subunit of a | 84 | 57 | 67 | 47 | 36 | 30 | 177 | 140 | 172 | 102 | 96 | 82 | 151.35 | 163 | 93 | 0.279 | -1.844 | 9.57E-31 |
| AT4G10650.1 | DGP2 | protein_coding | P-loop containing nucleo | 27 | 17 | 18 | 9 | 12 | 2 | 57 | 42 | 46 | 20 | 32 | 5 | 46.22 | 48 | 19 | 0.278 | -1.848 | 4.15E-11 |
| AT4G14660.1 | NRPE7 | protein_coding | RNA polymerase Rpb7-lik | 70 | 49 | 61 | 50 | 45 | 40 | 147 | 120 | 156 | 108 | 119 | 109 | 242.25 | 141 | 112 | 0.276 | -1.856 | 3.89E-41 |
| AT4G13560.1 | UNE15 | protein_coding | Late embryogenesis abun | 12 | 25 | 13 | 9 | 13 | 22 | 25 | 61 | 33 | 20 | 35 | 60 | 77.25 | 40 | 38 | 0.271 | -1.883 | 7.61E-12 |
| AT5G15300.1 | PCMP-E40 | protein_coding | Pentatricopeptide repeat | 35 | 17 | 23 | 18 | 13 | 8 | 74 | 42 | 59 | 39 | 35 | 22 | 44.65 | 58 | 32 | 0.269 | -1.892 | 2.55E-11 |
| AT5G60335.1 | | protein_coding | Thioesterase superfamily | 9 | 3 | 7 | 6 | 3 | 4 | 19 | 7 | 18 | 13 | 8 | 11 | 21.7 | 15 | 11 | 0.267 | -1.905 | 1.19E-08 |
| AT5G47460.1 | PCMP-E103 | protein_coding | Pentatricopeptide repeat | 22 | 11 | 12 | 9 | 10 | 7 | 46 | 27 | 31 | 20 | 27 | 19 | 42.86 | 35 | 22 | 0.264 | -1.923 | 4.01E-14 |
| AT1G68872.1 | | other_rna | | 24 | 16 | 22 | 16 | 10 | 8 | 50 | 39 | 56 | 35 | 27 | 22 | 39.18 | 48 | 28 | 0.262 | -1.931 | 1.27E-08 |
| AT5G49190.1 | SUS2 | protein_coding | sucrose synthase 2 | 23 | 14 | 11 | 11 | 10 | 10 | 48 | 34 | 28 | 24 | 27 | 27 | 39.98 | 37 | 26 | 0.26 | -1.941 | 2.31E-12 |
| AT1G61400.1 | | protein_coding | S-locus lectin protein kina | 51 | 37 | 35 | 34 | 26 | 24 | 107 | 91 | 90 | 74 | 69 | 65 | 103.05 | 96 | 69 | 0.258 | -1.953 | 1.54E-19 |
| AT4G12917.1 | | other_rna | other RNA | 76 | 71 | 79 | 45 | 48 | 33 | 160 | 174 | 202 | 98 | 127 | 90 | 237.88 | 179 | 105 | 0.253 | -1.984 | 1.34E-34 |
| AT2G26360.1 | | protein_coding | Mitochondrial substrate | 23 | 18 | 24 | 7 | 6 | 5 | 48 | 44 | 61 | 15 | 16 | 14 | 52.32 | 51 | 15 | 0.252 | -1.986 | 2.05E-15 |
| AT4G15563.1 | | protein_coding | | 9 | 6 | 7 | 10 | 2 | 4 | 19 | 15 | 18 | 22 | 5 | 11 | 30.85 | 17 | 13 | 0.252 | -1.989 | 6.80E-07 |
| AT3G61190.1 | BAP1 | protein_coding | BON association protein | 68 | 70 | 55 | 85 | 81 | 56 | 143 | 172 | 141 | 184 | 215 | 152 | 88.41 | 152 | 184 | 0.25 | -1.998 | 3.60E-06 |
| AT5G03770.1 | KDTA | protein_coding | KDO transferase A | 40 | 35 | 30 | 14 | 13 | 19 | 84 | 86 | 77 | 30 | 35 | 52 | 63.27 | 82 | 39 | 0.235 | -2.09 | 1.04E-16 |
| AT1G68050.1 | ADO3 | protein_coding | flavin-binding, kelch repe | 153 | 341 | 456 | 14 | 22 | 56 | 322 | 838 | 1167 | 30 | 58 | 152 | 508.12 | 776 | 80 | 0.235 | -2.091 | 1.08E-07 |
| AT1G52430.1 | | protein_coding | Ubiquitin carboxyl-termi | 22 | 20 | 11 | 17 | 16 | 9 | 46 | 49 | 28 | 37 | 42 | 24 | 43.34 | 41 | 34 | 0.234 | -2.093 | 1.45E-11 |
| AT4G33980.2 | | protein_coding | | 10 | 43 | 98 | 6 | 10 | 12 | 21 | 106 | 251 | 13 | 27 | 33 | 87.08 | 126 | 24 | 0.234 | -2.096 | 2.69E-05 |
| AT3G46280.1 | | protein_coding | protein kinase-related | 87 | 15 | 10 | 67 | 6 | 9 | 183 | 37 | 26 | 145 | 16 | 24 | 61.88 | 82 | 62 | 0.234 | -2.098 | 0.00200314 |
| AT4G15215.1 | PDR13 | protein_coding | pleiotropic drug resistanc | 62 | 50 | 49 | 61 | 52 | 55 | 130 | 123 | 125 | 132 | 138 | 150 | 179.6 | 126 | 140 | 0.231 | -2.116 | 1.09E-31 |
| AT3G13140.1 | | protein_coding | hydroxyproline-rich glyco | 4 | 3 | 3 | 2 | 3 | 5 | 8 | 7 | 8 | 4 | 8 | 14 | 26.87 | 8 | 9 | 0.212 | -2.237 | 6.65E-13 |
| AT1G11210.1 | | protein_coding | Protein of unknown funct | 43 | 81 | 95 | 4 | 8 | 26 | 90 | 199 | 243 | 9 | 21 | 71 | 123.36 | 177 | 34 | 0.208 | -2.268 | 4.53E-09 |
| AT2G15128.1 | | other_rna | other RNA | 25 | 25 | 29 | 18 | 8 | 12 | 53 | 61 | 74 | 39 | 21 | 33 | 64.74 | 63 | 31 | 0.203 | -2.299 | 7.93E-18 |
| AT4G14170.1 | PCMP-E17 | protein_coding | Pentatricopeptide repeat | 29 | 22 | 27 | 30 | 15 | 19 | 61 | 54 | 69 | 65 | 40 | 52 | 87.66 | 61 | 52 | 0.203 | -2.3 | 1.76E-20 |
| AT5G01215.1 | | other_rna | other RNA | 5 | 19 | 11 | 5 | 3 | 3 | 11 | 47 | 28 | 11 | 8 | 8 | 47.02 | 29 | 9 | 0.202 | -2.308 | 3.91E-16 |
| AT4G14820.1 | PCMP-H3 | protein_coding | Pentatricopeptide repeat | 43 | 17 | 20 | 23 | 15 | 16 | 90 | 42 | 51 | 50 | 40 | 44 | 85.34 | 61 | 45 | 0.19 | -2.395 | 5.18E-18 |
| AT5G13225.1 | | small_nucleolar_r | snoRNA | 63 | 68 | 72 | 84 | 64 | 49 | 132 | 167 | 184 | 182 | 170 | 133 | 81.59 | 161 | 162 | 0.189 | -2.4 | 0.00347498 |
| AT5G23240.1 | | protein_coding | DNAJ heat shock N-termi | 95 | 183 | 186 | 26 | 23 | 16 | 200 | 450 | 476 | 56 | 61 | 44 | 380.43 | 375 | 54 | 0.183 | -2.448 | 2.03E-17 |
| AT3G23480.2 | | protein_coding | Cyclopropane-fatty-acyl-p | 30 | 53 | 70 | 13 | 5 | 7 | 63 | 130 | 179 | 28 | 13 | 19 | 77.09 | 124 | 20 | 0.181 | -2.466 | 1.60E-14 |
| AT2G47115.1 | | protein_coding | | 284 | 482 | 261 | 347 | 97 | 32 | 597 | 1184 | 668 | 753 | 258 | 87 | 324.87 | 816 | 366 | 0.18 | -2.476 | 0.00011546 |
| AT5G42900.1 | COR27 | protein_coding | cold regulated gene 27 | 16 | 40 | 83 | 4 | 2 | 6 | 34 | 98 | 212 | 9 | 5 | 16 | 103.36 | 115 | 10 | 0.17 | -2.553 | 4.11E-10 |
| AT4G13800.1 | | protein_coding | Protein of unknown funct | 109 | 94 | 74 | 26 | 31 | 16 | 229 | 231 | 189 | 56 | 82 | 44 | 216.06 | 216 | 61 | 0.17 | -2.558 | 1.01E-53 |
| AT1G72060.1 | | protein_coding | serine-type endopeptidas | 13 | 7 | 11 | 25 | 2 | 6 | 27 | 17 | 28 | 54 | 5 | 16 | 27.62 | 24 | 25 | 0.16 | -2.646 | 5.03E-08 |
| AT4G14050.1 | PCMP-H13 | protein_coding | Pentatricopeptide repeat | 48 | 25 | 25 | 18 | 13 | 9 | 101 | 61 | 64 | 39 | 35 | 24 | 101.59 | 75 | 33 | 0.158 | -2.661 | 6.42E-19 |
| AT5G60100.2 | APRR3 | protein_coding | pseudo-response regulato | 149 | 123 | 147 | 9 | 9 | 11 | 313 | 302 | 376 | 20 | 24 | 30 | 242.18 | 330 | 25 | 0.157 | -2.668 | 6.73E-58 |
| AT3G18070.1 | BGLU43 | protein_coding | beta glucosidase 43 | 59 | 69 | 54 | 6 | 11 | 10 | 124 | 170 | 138 | 13 | 29 | 27 | 79.12 | 144 | 23 | 0.155 | -2.694 | 6.14E-21 |
| AT4G04409.1 | | pseudogene | | 58 | 67 | 41 | 44 | 56 | 31 | 122 | 165 | 105 | 95 | 149 | 84 | 85.31 | 131 | 109 | 0.15 | -2.734 | 3.38E-20 |
| AT1G51402.1 | | protein_coding | | 176 | 471 | 429 | 236 | 92 | 38 | 370 | 1157 | 1098 | 512 | 244 | 103 | 320.37 | 875 | 286 | 0.148 | -2.752 | 5.51E-05 |
| AT4G15230.1 | ABCG30 | protein_coding | pleiotropic drug resistanc | 123 | 27 | 30 | 60 | 21 | 23 | 259 | 66 | 77 | 130 | 56 | 63 | 161.53 | 134 | 83 | 0.129 | -2.952 | 4.32E-13 |
| AT4G15160.1 | DL3625W | protein_coding | Bifunctional inhibitor/lipi | 3 | 1 | 0 | 2 | 1 | 0 | 6 | 2 | 0 | 4 | 3 | 0 | 31.31 | 3 | 2 | 0.107 | -3.229 | 2.75E-13 |
| AT5G03545.1 | AT4 | protein_coding | | 260 | 73 | 121 | 111 | 63 | 52 | 547 | 179 | 310 | 241 | 167 | 141 | 256.61 | 345 | 183 | 0.099 | -3.339 | 5.24E-12 |
| AT3G09922.1 | IPS1 | protein_coding | induced by phosphate sta | 60 | 5 | 4 | 5 | 0 | 3 | 126 | 12 | 10 | 11 | 0 | 8 | 28.58 | 49 | 6 | 0.049 | -4.35 | 1.86E-06 |

**Table S3. PPR gene expression**

| Gene ID | Gene Names | | PPR class | PPR localization | *caa39* vs Col-0 | | *tfIIfa-1* vs Col-3 | | *tfIIfa-2* vs Col0 | |
|---|---|---|---|---|---|---|---|---|---|---|
| | | | Cheng *et al.* ( | Colcombet *et al.* (201 | log2FoldChan | pvalue | log2FoldChan | pvalue | log2FoldChan | pvalue |
| AT5G66520 | PCMP-H61 | CREF7 | DYW | C | -0.14 | 0.11004827 | 1.034 | 2.89E-13 | 0.676 | 6.19E-06 |
| AT3G60960 | | P487 | P | M | -0.486 | 0.00155879 | 1.007 | 6.42E-10 | 0.683 | 0.00013015 |
| AT2G01860 | EMB975 | EMB975 | P | c | 0.602 | 2.51E-13 | 0.896 | 2.18E-11 | 0.647 | 3.41E-06 |
| AT1G47580 | DYW1 | DYW1 | DYW | C | -0.675 | 5.19E-12 | 0.79 | 1.59E-09 | 0.648 | 3.77E-06 |
| AT2G35130 | | | P | c | -0.617 | 3.02E-15 | 0.79 | 7.45E-08 | 0.594 | 9.70E-05 |
| AT1G08610 | | | P | pC | 0.25 | 0.02143435 | 0.658 | 5.35E-05 | 0.114 | 0.52521993 |
| AT2G40240 | | | P | M | 0.06 | 0.73079293 | 0.653 | 0.00060197 | 0.334 | 0.25683034 |
| AT4G21170 | | | P | M/C | 0.455 | 0.00576616 | 0.625 | 0.00038838 | 0.272 | 0.18930224 |
| AT2G26790 | | | P | M | 0.217 | 0.2838346 | 0.624 | 0.0009919 | 0.337 | 0.13653712 |
| AT2G15980 | | | P | | 0.161 | 0.2648639 | 0.598 | 0.00197687 | 0.006 | 0.9824814 |
| AT3G60980 | | P486 | P | pM | 0.584 | 4.74E-14 | 0.569 | 7.94E-05 | 0.586 | 0.00013383 |
| AT3G04760 | | | P | C | -0.07 | 0.30043235 | 0.554 | 5.09E-05 | 0.258 | 0.0628002 |
| AT5G43820 | | | P | pM | 0.264 | 0.19760352 | 0.539 | 0.01827244 | -0.207 | 0.42706085 |
| AT4G32450 | PCMP-H63 | MEF8S | DYW | M | 0.277 | 0.01891538 | 0.534 | 0.00049435 | 0.113 | 0.50493855 |
| AT5G02830 | | | P | C | -0.099 | 0.13206748 | 0.533 | 0.00233553 | 0.36 | 0.0420353 |
| AT3G25210 | | | P | | -0.463 | 0.00096169 | 0.528 | 0.00737053 | 0.062 | 0.78260228 |
| AT5G47360 | | | P | pM | -0.003 | 0.98053991 | 0.513 | 0.00242566 | 0.391 | 0.05551307 |
| AT4G26680 | | | P | pM | 0.015 | 0.94235335 | 0.51 | 0.00993827 | 0.089 | 0.70622182 |
| AT3G02650 | | | P | pM | 0.212 | 0.00686731 | 0.509 | 0.00016135 | 0.227 | 0.1114255 |
| AT3G26630 | PCMP-A6 | | PLS | pC | -0.548 | 1.60E-10 | 0.5 | 0.00072206 | 0.071 | 0.65911269 |
| AT5G55740 | CRR21 | CRR21 | E+ | C | -0.22 | 0.00349626 | 0.49 | 0.00044725 | 0.228 | 0.12008556 |
| AT1G31920 | PCMP-H11 | | DYW | c | -0.458 | 5.24E-08 | 0.487 | 6.12E-05 | 0.247 | 0.05079999 |
| AT1G05670 | | | P | M/C | -0.126 | 0.24881417 | 0.48 | 0.00067663 | -0.242 | 0.13489929 |
| AT1G05600 | | EMB3101 | P | pM | -0.071 | 0.54467209 | 0.478 | 0.00249919 | 0.105 | 0.56007699 |
| AT4G38150 | | | P | m | 0.304 | 0.00157763 | 0.474 | 0.00172614 | 0.368 | 0.0463976 |
| AT1G10270 | GRP23 | GRP23 | P | M/N | 0.273 | 0.00241621 | 0.439 | 0.00054816 | 0.444 | 0.00236961 |
| AT1G73710 | | | P | C | 0.269 | 0.00533216 | 0.435 | 0.00032342 | 0.033 | 0.7947359 |
| AT2G01740 | | | P | M | -0.154 | 0.36767577 | 0.435 | 0.06583369 | -0.212 | 0.43458499 |
| AT1G11710 | | | P | pM | 0.109 | 0.6202639 | 0.434 | 0.0142032 | 0.434 | 0.05135751 |
| AT3G15590 | | | P | M | 0.644 | 8.40E-15 | 0.43 | 0.00141684 | 0.168 | 0.23457386 |
| AT3G54980 | | | P | pM | 0.343 | 0.00181913 | 0.418 | 0.00269716 | 0.018 | 0.90299062 |
| AT4G01990 | | | P | pM | 0.914 | 1.02E-13 | 0.417 | 0.08749584 | 0.044 | 0.86845196 |
| AT2G32230 | PRORP1 | PRORP1 | P | M/C | -0.021 | 0.75306765 | 0.408 | 0.00964416 | -0.016 | 0.92159608 |
| AT1G55890 | | | P | M | 0.191 | 0.00864497 | 0.406 | 0.00022078 | 0.131 | 0.28193957 |
| AT4G26800 | | | P | pM | 0.341 | 0.08653533 | 0.388 | 0.11877124 | -0.047 | 0.87568777 |
| AT1G22960 | | | P | pM | -0.103 | 0.40442223 | 0.386 | 0.00693486 | -0.294 | 0.0689934 |
| AT1G79490 | EMB2217 | EMB2217 | P | M | 0.371 | 0.00124593 | 0.373 | 0.00521377 | 0.135 | 0.38311842 |
| AT3G59040 | | | P | C | 0.049 | 0.49636559 | 0.373 | 0.01991731 | -0.098 | 0.54667115 |
| AT5G61990 | | | P | pM | -0.081 | 0.52394359 | 0.362 | 0.00228493 | 0.114 | 0.39909142 |
| AT5G46580 | | | P | C | -0.126 | 0.06010221 | 0.347 | 0.02885144 | -0.019 | 0.90575417 |
| AT3G16890 | PPR40 | PPR40 | P | M (Zsigmond *et al.* 20 | 0.066 | 0.73017339 | 0.345 | 0.06818091 | -0.067 | 0.7541281 |
| AT5G14770 | | | P | M | -0.127 | 0.35515672 | 0.339 | 0.04058296 | 0.032 | 0.87471735 |
| AT4G31850 | PGR3 | PGR3 | P | C | -0.414 | 1.17E-07 | 0.338 | 0.00461324 | -0.104 | 0.39544899 |
| AT1G03560 | | | P | pM | -0.028 | 0.78265281 | 0.337 | 0.00685527 | 0.046 | 0.77018708 |
| AT2G42920 | PCMP-E75 | | E+ | pC | -0.069 | 0.46340484 | 0.328 | 0.00210123 | 0.17 | 0.18227935 |
| AT2G30100 | | | P | pC | -0.165 | 0.11324931 | 0.325 | 0.01235306 | -0.012 | 0.93934489 |
| AT4G21705 | | | P | pM | 0.342 | 0.00107989 | 0.321 | 0.07387088 | -0.03 | 0.8813974 |
| AT3G18110 | EMB1270 | EMB1270 | P | C | -0.292 | 4.76E-05 | 0.317 | 0.01882028 | 0.154 | 0.26535126 |
| AT4G35850 | | PPRL | P | M | 0.023 | 0.74769083 | 0.315 | 0.02063628 | 0.23 | 0.10589126 |
| AT1G62670 | RPF2 | RPF2 | P | M | -0.121 | 0.43931193 | 0.314 | 0.07627207 | -0.212 | 0.31003172 |
| AT2G27800 | | | P | pM | -0.275 | 0.02617674 | 0.309 | 0.01805781 | -0.032 | 0.84572251 |
| AT2G15690 | PCMP-H66 | DYW2 | DYW | M/C | 0.062 | 0.42741218 | 0.306 | 0.05955744 | -0.053 | 0.75141577 |
| AT5G40400 | | | P | | 0.168 | 0.31260469 | 0.29 | 0.05390299 | -0.089 | 0.62639172 |
| AT4G16390 | P67 | SVR7 | P | C | -0.049 | 0.46503478 | 0.289 | 0.08453878 | -0.099 | 0.55781167 |
| AT4G34830 | MRL1 | MRL1 | P | C | -0.31 | 5.06E-06 | 0.284 | 0.06111138 | 0.159 | 0.29772038 |
| AT1G59720 | PCMP-H51 | CRR28 | DYW | C | -0.658 | 6.45E-07 | 0.282 | 0.07519466 | -0.063 | 0.71950106 |
| AT5G18950 | | | P | M | 0.164 | 0.23842814 | 0.276 | 0.07125874 | 0.267 | 0.12886869 |
| AT5G27300 | | | P | | -0.299 | 0.00429231 | 0.274 | 0.05956757 | -0.009 | 0.96098039 |
| AT5G11310 | | SOAR1 | P | pM | 0.362 | 0.01180416 | 0.268 | 0.15462748 | -0.567 | 0.01385445 |
| AT1G29710 | PCMP-H67 | DYW4 | DYW | pM | -0.062 | 0.66573757 | 0.26 | 0.0782012 | 0.073 | 0.66987808 |
| AT1G63330 | | | P | M | -0.564 | 0.01261667 | 0.249 | 0.26086278 | -0.206 | 0.41512003 |
| AT4G20740 | | EMB3131 | P | C | -0.252 | 0.06545217 | 0.245 | 0.11577611 | -0.588 | 0.0019918 |
| AT1G30290 | | | P | pM | 0.098 | 0.37916996 | 0.241 | 0.11783452 | -0.03 | 0.85672683 |
| AT1G19520 | NFD5 | NFD5 | P | pM | 0.106 | 0.16284024 | 0.24 | 0.0584547 | -0.132 | 0.31545638 |
| AT2G16880 | | | P | pM | -0.268 | 0.01025018 | 0.24 | 0.06423561 | 0.003 | 0.98502549 |

| | | | | | | | | | | |
|---|---|---|---|---|---|---|---|---|---|---|
| AT2G20710 | | | P | pM | 0.173 | 0.22812559 | 0.238 | 0.28829808 | 0.031 | 0.91274643 |
| AT3G49730 | | | P | M | -0.024 | 0.85736719 | 0.236 | 0.28073141 | 0.15 | 0.6013743 |
| AT4G30825 | | | P | c | -0.149 | 0.03296833 | 0.236 | 0.0996576 | -0.052 | 0.72525273 |
| AT1G80270 | PPR596 | PPR596 | P | M/c | 0.228 | 0.00299461 | 0.23 | 0.16554787 | 0.083 | 0.63375775 |
| AT1G14090 | | | P | | -0.071 | 0.68703144 | 0.224 | 0.21021013 | -0.337 | 0.11749265 |
| AT1G19720 | DYW7 | DYW7 | DYW | m/c | -0.296 | 4.49E-05 | 0.224 | 0.05521653 | -0.059 | 0.61624945 |
| AT5G14350 | | | P | | -0.691 | 0.01241392 | 0.223 | 0.3249986 | -0.184 | 0.50091992 |
| AT5G48730 | | | P | C | -0.304 | 0.00031422 | 0.223 | 0.08094797 | 0.028 | 0.84094474 |
| AT1G62930 | | RPF3 | P | M | -0.307 | 0.00467767 | 0.222 | 0.07886273 | 0.079 | 0.58946658 |
| AT1G80150 | | | P | pM | 0.198 | 0.12915448 | 0.22 | 0.18070155 | -0.084 | 0.67075755 |
| AT1G13800 | | FAC19 | P | pM | 0.532 | 0.0346916 | 0.219 | 0.44073839 | -0.447 | 0.1417785 |
| AT1G64430 | | | P | pC | -0.149 | 0.02110611 | 0.209 | 0.05876369 | 0.112 | 0.34132152 |
| AT3G61360 | | | P | pM | -0.023 | 0.88940936 | 0.204 | 0.16832508 | 0.366 | 0.03583284 |
| AT5G28340 | | | P | | -0.547 | 0.02352875 | 0.201 | 0.38400532 | 0.441 | 0.11405016 |
| AT1G20300 | | | P | M | 0.034 | 0.71943096 | 0.199 | 0.14969377 | -0.229 | 0.17187222 |
| AT1G71210 | | | P | pM | 0.283 | 0.0013795 | 0.198 | 0.11630738 | -0.133 | 0.33540324 |
| AT1G15480 | | | P | M | 0.295 | 0.00096268 | 0.196 | 0.10522175 | -0.177 | 0.20334332 |
| AT3G49240 | EMB1796 | NUWA/EMB1 | P | M/C | 0.319 | 4.56E-05 | 0.196 | 0.2649404 | -0.274 | 0.14510557 |
| AT5G04810 | | | P | C | -0.254 | 0.00020784 | 0.196 | 0.08718834 | 0.057 | 0.62873831 |
| AT1G16830 | | | P | pM | 0.62 | 0.00180707 | 0.195 | 0.38906672 | -0.025 | 0.93062656 |
| AT5G39710 | EMB2745 | EMB2745 | P | M | 0.163 | 0.12901976 | 0.195 | 0.13355952 | -0.139 | 0.33473879 |
| AT3G29230 | PCMP-E27 | | E+ | C | -0.235 | 0.00175866 | 0.194 | 0.15731464 | 0.044 | 0.76987407 |
| AT1G30610 | EMB2279 | EMB2279 | P | C | -0.076 | 0.27989142 | 0.193 | 0.13105212 | 0.015 | 0.90841379 |
| AT4G21900 | PRORP3 | PRORP3 | P | N | 0.066 | 0.5204257 | 0.193 | 0.10482178 | 0.032 | 0.82174824 |
| AT2G25580 | PCMP-H75 | MEF8 | DYW | M | 0.079 | 0.48785528 | 0.191 | 0.20618636 | -0.007 | 0.96480811 |
| AT2G48000 | | | P | pM | -0.227 | 0.01774654 | 0.19 | 0.09954795 | -0.209 | 0.15860725 |
| AT1G62860 | | | P | | -0.025 | 0.84770147 | 0.187 | 0.33149525 | 0.061 | 0.78370215 |
| AT4G01570 | | | P | C | -0.282 | 0.00410546 | 0.186 | 0.11643822 | -0.007 | 0.95824258 |
| AT5G50280 | EMB1006 | EMB1006 | P | C | -0.085 | 0.28197397 | 0.186 | 0.10898781 | -0.242 | 0.05840876 |
| AT1G51965 | ABO5 | ABO5 | P | M | -0.096 | 0.32119888 | 0.172 | 0.16054495 | 0.156 | 0.32425618 |
| AT5G27270 | EMB976 | EMB976 | P | C | -0.24 | 0.00132749 | 0.17 | 0.12209473 | -0.136 | 0.25478372 |
| AT1G69290 | | | P | M | 0.733 | 8.10E-07 | 0.169 | 0.34516312 | -0.027 | 0.89648144 |
| AT2G45350 | CRR4 | CRR4 | E2 | C | -0.025 | 0.91955157 | 0.169 | 0.5560227 | 0.318 | 0.32115033 |
| AT4G02820 | | | P | M | -0.103 | 0.28154224 | 0.162 | 0.17996348 | -0.084 | 0.52872949 |
| AT1G62350 | | THA8-LIKE3 | P | pM | 0.352 | 0.06325927 | 0.16 | 0.61236157 | -0.205 | 0.5935051 |
| AT1G11900 | | | P | pM | -0.166 | 0.30641917 | 0.154 | 0.43798397 | -0.525 | 0.03611798 |
| AT1G63230 | | | P | | -1.126 | 4.59E-09 | 0.153 | 0.3996386 | 0.185 | 0.41978772 |
| AT5G25630 | | | P | c | -0.558 | 3.02E-17 | 0.145 | 0.19565919 | -0.023 | 0.84253303 |
| AT2G38420 | | | P | pM | -0.066 | 0.6430515 | 0.144 | 0.4003874 | 0.178 | 0.3910711 |
| AT1G62590 | | | P | M | -0.623 | 3.20E-07 | 0.14 | 0.31701892 | -0.259 | 0.11289011 |
| AT3G56030 | | | P | pM | 0.578 | 0.00399829 | 0.138 | 0.6928181 | 0.467 | 0.26197341 |
| AT3G13150 | | | P | M (Lurin *et al.* 2004) | 0.565 | 6.15E-09 | 0.134 | 0.37314261 | 0.076 | 0.65675583 |
| AT3G06920 | | | P | M | 0.26 | 0.03464954 | 0.133 | 0.45430697 | -0.139 | 0.48827269 |
| AT5G15010 | | | P | pC | -0.307 | 0.09144289 | 0.133 | 0.46733775 | 0.174 | 0.38178021 |
| AT5G60960 | PNM1 | PNM1 | P | M/N | 0.168 | 0.01676875 | 0.133 | 0.33204977 | 0.031 | 0.82877253 |
| AT2G17670 | | | P | pM | 0.283 | 0.016714 | 0.132 | 0.38472754 | -0.095 | 0.5826046 |
| AT5G65820 | | | P | M | -0.372 | 0.01299676 | 0.128 | 0.44403741 | -0.189 | 0.35887389 |
| AT2G41720 | EMB2654 | EMB2654 | P | C | -0.266 | 0.00023731 | 0.121 | 0.44521163 | -0.237 | 0.1464825 |
| AT3G16010 | | | P | pM | -0.04 | 0.77604489 | 0.121 | 0.34250013 | -0.195 | 0.16819273 |
| AT1G19290 | | | P | M | -0.057 | 0.69896277 | 0.118 | 0.55989834 | -0.223 | 0.31730272 |
| AT2G36240 | | | P | M | -0.203 | 0.00555713 | 0.118 | 0.40990012 | 0.122 | 0.42677567 |
| AT1G15510 | PCMP-H73 | AtECB2/VAC1 | DYW | C | 0.403 | 8.10E-05 | 0.113 | 0.54080762 | -0.348 | 0.08429818 |
| AT1G13630 | | | P | | 0.066 | 0.73486803 | 0.112 | 0.5503643 | -0.15 | 0.51042154 |
| AT1G61870 | PPR336 | PPR336 | P | M | 0.102 | 0.17831055 | 0.109 | 0.43535835 | -0.105 | 0.47673459 |
| AT1G02420 | | | P | m/c | -0.013 | 0.93351872 | 0.108 | 0.55143857 | -0.101 | 0.6319054 |
| AT1G73400 | | | P | pM | 0.31 | 0.04011014 | 0.108 | 0.59090419 | -0.294 | 0.18592445 |
| AT2G02150 | | | P | pC | 0.206 | 0.14882257 | 0.108 | 0.56689994 | -0.488 | 0.03558582 |
| AT1G63400 | | | P | M | -0.364 | 0.04440198 | 0.103 | 0.58712208 | -0.541 | 0.01275823 |
| AT1G74850 | PTAC2 | PTAC2 | P | C | -0.34 | 2.07E-06 | 0.098 | 0.46529651 | -0.155 | 0.26171279 |
| AT5G67570 | DG1 | EMB1408/DG | P | C | -0.181 | 0.01931051 | 0.092 | 0.41049815 | -0.059 | 0.64194329 |
| AT3G48250 | | BIR6 | P | M | 0.317 | 0.01413108 | 0.084 | 0.65192463 | -0.112 | 0.58902063 |
| AT5G42310 | | CRP1 | P | C | -0.268 | 4.75E-05 | 0.077 | 0.51209355 | -0.014 | 0.9088072 |
| AT1G18900 | | | P | pM | 0.233 | 0.0106993 | 0.074 | 0.5391056 | 0.111 | 0.39791964 |
| AT1G63080 | | | P | pM | -0.379 | 0.00694257 | 0.071 | 0.63665856 | 0.062 | 0.7255781 |
| AT2G18940 | | | P | C | -0.283 | 0.0001476 | 0.066 | 0.6224695 | -0.198 | 0.15991064 |
| AT5G13770 | | | P | C | -0.635 | 3.28E-21 | 0.066 | 0.61092916 | 0.026 | 0.84468968 |
| AT5G65560 | | | P | pM | 0.074 | 0.49636988 | 0.066 | 0.62688753 | -0.025 | 0.8668375 |
| AT1G74580 | | | P | | -0.041 | 0.76098754 | 0.063 | 0.66847127 | -0.213 | 0.234598 |
| AT1G10910 | | EMB3103 | P | c | -0.333 | 2.66E-05 | 0.061 | 0.57572665 | -0.042 | 0.71494853 |

| | | | | | | | | | | |
|---|---|---|---|---|---|---|---|---|---|---|
| AT3G04750 | PCMP-E81 | | E+ | pM | -0.129 | 0.33441189 | 0.057 | 0.71447399 | -0.027 | 0.88646599 |
| AT1G12300 | | RFL2 | P | pM | -0.128 | 0.35938891 | 0.056 | 0.71237909 | -0.227 | 0.21145398 |
| AT2G31400 | GUN1 | GUN1 | P | C | -0.128 | 0.06132009 | 0.056 | 0.62431294 | -0.048 | 0.68346055 |
| AT4G21190 | EMB1417 | EMB1417 | P | c | -0.254 | 0.00328171 | 0.054 | 0.68945212 | -0.253 | 0.08871702 |
| AT3G02490 | | | P | pM | 0.107 | 0.31795404 | 0.047 | 0.75037975 | -0.131 | 0.42628952 |
| AT4G17616 | | | P | pM | 0.107 | 0.30375208 | 0.043 | 0.76607795 | -0.022 | 0.88912849 |
| AT1G64580 | | | P | pM | 0.262 | 0.03136191 | 0.033 | 0.82564062 | -0.125 | 0.47457799 |
| AT3G04130 | | | P | pM | 0.165 | 0.2193923 | 0.03 | 0.85371947 | -0.167 | 0.38380384 |
| AT5G15980 | | | P | M | 0.135 | 0.09011426 | 0.03 | 0.84256345 | 0.007 | 0.96297859 |
| AT2G15820 | OTP51 | OTP51 | P | C | -0.363 | 1.46E-06 | 0.026 | 0.86137773 | -0.229 | 0.13610993 |
| AT1G68980 | | | P | pM | -0.328 | 0.00320904 | 0.02 | 0.89595172 | 0.102 | 0.5667769 |
| AT2G37230 | | | P | M/c | 0.161 | 0.02606206 | 0.019 | 0.89384345 | -0.207 | 0.15600908 |
| AT1G10330 | PCMP-E71 | | E2 | pM | 0.066 | 0.74765711 | 0.018 | 0.94434538 | 0.055 | 0.86023268 |
| AT4G21880 | | | P | M | -0.078 | 0.45791355 | 0.016 | 0.93000727 | -0.499 | 0.0117949 |
| AT5G08510 | PCMP-E20 | | E+ | M | 0.395 | 0.0046185 | 0.014 | 0.92809715 | 0.071 | 0.72968972 |
| AT5G03800 | EMB175 | EMB175 | DYW | m/C | 0.027 | 0.77283388 | 0.011 | 0.9490122 | -0.437 | 0.02456672 |
| AT1G63130 | | RPF6 | P | pM | -0.217 | 0.01428666 | 0.006 | 0.96546097 | -0.07 | 0.61910536 |
| AT3G63370 | OTP86 | OTP86 | E+ | C | -0.552 | 4.29E-07 | 0.006 | 0.96432946 | -0.131 | 0.37080585 |
| AT1G76280 | | | P | M | -0.141 | 0.10354468 | 0.005 | 0.96845471 | -0.29 | 0.05252316 |
| AT1G26460 | | | P | M | 0.185 | 0.00871349 | 0.001 | 0.99511557 | -0.089 | 0.52870603 |
| AT3G14330 | PCMP-H57 | CREF3 | DYW | M/C | -0.246 | 0.00355841 | 0.001 | 0.99040201 | -0.336 | 0.00656668 |
| AT1G09900 | | | P | m/c | -0.091 | 0.22467043 | 0 | 0.99935734 | -0.369 | 0.03086064 |
| AT2G39230 | LOJ | LOJ | P | pM | 0.742 | 0.00466815 | 0 | 0.999106 | -0.442 | 0.10398029 |
| AT3G22150 | PCMP-E95 | AEF1/MPR25 | E+ | M/c | -0.356 | 6.03E-06 | -0.003 | 0.98170277 | -0.367 | 0.01339314 |
| AT5G03560 | | | P | pM | 0.596 | 3.69E-05 | -0.006 | 0.97479124 | 0.385 | 0.10366341 |
| AT5G12100 | | | P | M | 0.255 | 0.0506781 | -0.014 | 0.92368901 | 0.033 | 0.84683607 |
| AT5G50390 | PCMP-H58 | EMB3141 | DYW | C | -0.409 | 0.00183419 | -0.016 | 0.92556784 | -0.323 | 0.09184773 |
| AT3G60040 | | | P | | 0.06 | 0.75137033 | -0.019 | 0.92476865 | -0.366 | 0.1478296 |
| AT4G04790 | | | P | pM | 0.453 | 0.00146631 | -0.019 | 0.9248268 | -0.513 | 0.02374543 |
| AT5G15280 | | | P | pM | -0.366 | 5.25E-05 | -0.02 | 0.87190001 | -0.576 | 1.96E-05 |
| AT3G09650 | HCF152 | HCF152/CRM | P | C | -0.006 | 0.93692209 | -0.023 | 0.83521495 | -0.185 | 0.14449264 |
| AT3G46610 | | | P | C | -0.078 | 0.29665988 | -0.025 | 0.82851099 | -0.449 | 0.00048148 |
| AT5G57250 | | | P | pM | -0.02 | 0.81610126 | -0.032 | 0.80920156 | -0.309 | 0.04478913 |
| AT4G20090 | EMB1025 | EMB1025 | P | M | 0.106 | 0.42457377 | -0.035 | 0.84341948 | -0.476 | 0.03245229 |
| AT1G06710 | | MTSF1 | P | M | 0.419 | 0.0007628 | -0.036 | 0.80649404 | -0.523 | 0.00240878 |
| AT5G50990 | PCMP-H59 | | DYW | | 0.72 | 3.86E-05 | -0.036 | 0.88537635 | 0.188 | 0.51301086 |
| AT3G60050 | | | P | pM | 0.12 | 0.32472019 | -0.043 | 0.77807849 | -0.132 | 0.46933088 |
| AT5G10690 | CBSPPR1 | CBSPPR1 | P | C | -0.739 | 3.86E-28 | -0.045 | 0.70273506 | -0.269 | 0.0336993 |
| AT2G13420 | | | P | pM | 0.201 | 0.31893518 | -0.047 | 0.81797628 | -0.619 | 0.03390427 |
| AT1G74750 | | | P | pC | 0.139 | 0.18088063 | -0.048 | 0.72669276 | -0.082 | 0.61414857 |
| AT1G11630 | | | P | M | 0.059 | 0.60030774 | -0.054 | 0.70820616 | -0.091 | 0.59295444 |
| AT1G03540 | PCMP-E4 | | E2 | pM | 0.157 | 0.59731581 | -0.057 | 0.80480354 | -0.103 | 0.70672825 |
| AT1G62680 | | | P | pM | -0.435 | 0.00207671 | -0.057 | 0.73021637 | -0.13 | 0.51967058 |
| AT3G51320 | | | E2 | pM | 0.028 | 0.80459565 | -0.057 | 0.74060721 | -0.103 | 0.60936694 |
| AT5G64320 | | | P | pM | -0.451 | 0.00028811 | -0.061 | 0.70583462 | -0.341 | 0.07001797 |
| AT1G63150 | | | P | pM | -0.367 | 0.04499716 | -0.065 | 0.71102843 | -0.088 | 0.66532503 |
| AT5G21222 | ATC401 | AtC401 | P | m/C | -0.158 | 0.01879652 | -0.065 | 0.56072023 | -0.308 | 0.00836782 |
| AT1G26500 | | | P | pM | 0.369 | 0.14048925 | -0.066 | 0.79656943 | 0.209 | 0.53094442 |
| AT2G19280 | | | P | M/c | -0.461 | 9.92E-05 | -0.067 | 0.5914464 | -0.233 | 0.10515102 |
| AT5G18390 | | | P | pM | -0.075 | 0.48741139 | -0.071 | 0.61245024 | -0.479 | 0.00750723 |
| AT1G12775 | | EMB1586/ISE | P | pM | -0.552 | 0.00015604 | -0.077 | 0.66638209 | -0.409 | 0.04958583 |
| AT4G19890 | | | P | | 0.369 | 0.00666523 | -0.08 | 0.67216747 | -0.469 | 0.02265012 |
| AT2G30780 | | | P | pM | 0.038 | 0.73847712 | -0.085 | 0.47274865 | -0.061 | 0.64821431 |
| AT1G62914 | | | P | | -0.138 | 0.41733522 | -0.086 | 0.64378642 | 0.03 | 0.88462529 |
| AT5G09450 | | | P | M | 0.039 | 0.648138 | -0.086 | 0.50286366 | -0.015 | 0.91309475 |
| AT5G01110 | | | P | pC | 0.387 | 0.05524981 | -0.09 | 0.73279611 | -0.787 | 0.00729493 |
| AT1G74900 | OTP43 | OTP43 | P | M | -0.043 | 0.79813847 | -0.096 | 0.65436261 | 0.102 | 0.6933763 |
| AT1G16480 | | | DYW | pM | 0.32 | 0.05402595 | -0.098 | 0.57879475 | -0.132 | 0.53791871 |
| AT3G53700 | MEE40 | MEE40 | P | C | -0.019 | 0.79426341 | -0.102 | 0.47155636 | -0.388 | 0.00829759 |
| AT5G62370 | | | P | pM | 0.141 | 0.42484482 | -0.104 | 0.5648115 | -0.321 | 0.10940218 |
| AT1G06150 | EMB1444 | EMB1444 | E+ | M/C | 0.123 | 0.11520723 | -0.106 | 0.34218703 | -0.267 | 0.03293353 |
| AT1G79080 | | | P | pC | -0.009 | 0.92208965 | -0.106 | 0.36257882 | -0.151 | 0.2575364 |
| AT1G62910 | | RFL9 | P | pM | -0.173 | 0.24614734 | -0.107 | 0.54041742 | -0.166 | 0.41798148 |
| AT1G07590 | | | P | pM | 0.121 | 0.08116073 | -0.111 | 0.49949563 | 0.2 | 0.2247667 |
| AT4G01030 | PCMP-H65 | | DYW | c | -0.086 | 0.45617885 | -0.112 | 0.47873497 | -0.453 | 0.00759322 |
| AT5G48910 | PCMP-H38 | LPA66 | DYW | m/C | -0.19 | 0.03730988 | -0.113 | 0.37242104 | -0.25 | 0.06267332 |
| AT4G39620 | EMB2453 | EMB2453/At | P | C | -0.048 | 0.55197107 | -0.115 | 0.2887266 | -0.425 | 0.00041244 |
| AT3G22670 | | | P | pM | -0.263 | 0.15411632 | -0.119 | 0.58659068 | -0.124 | 0.64547669 |
| AT3G46790 | CRR2 | CRR2 | DYW | C | 0.366 | 0.00029166 | -0.119 | 0.41841722 | 0.109 | 0.48619674 |

| | | | | | | | | | | |
|---|---|---|---|---|---|---|---|---|---|---|
| AT3G18970 | PCMP-E93 | MEF20 | E2 | M/c | 0.344 | 0.02553392 | -0.121 | 0.48596073 | 0.353 | 0.090365 |
| AT3G16710 | | PPR40 | P | M | -0.012 | 0.94044735 | -0.123 | 0.52727243 | -0.442 | 0.05572143 |
| AT1G09820 | | | P | pM | 0.076 | 0.56386844 | -0.126 | 0.39186317 | -0.302 | 0.08151312 |
| AT1G12700 | | RPF1 | P | M | -0.402 | 0.00190219 | -0.134 | 0.41229102 | -0.415 | 0.03938222 |
| AT3G04260 | PTAC3 | PDE324/pTAC | P | C (Yagi *et al.* 2012) | -0.109 | 0.11363337 | -0.134 | 0.23029121 | -0.647 | 2.03E-08 |
| AT5G43790 | PCMP-E30 | | E2 | m/c | 0.147 | 0.22191764 | -0.141 | 0.31830117 | -0.07 | 0.7047677 |
| AT2G16650 | PRORP2 | PRORP2 | P | N | 0.372 | 0.0038523 | -0.143 | 0.32321445 | -0.587 | 0.00135556 |
| AT1G60770 | | | P | M | 0.177 | 0.02034477 | -0.151 | 0.32339687 | -0.166 | 0.29332239 |
| AT1G55630 | | | P | pM | -0.178 | 0.16165577 | -0.154 | 0.3014697 | 0.098 | 0.59726488 |
| AT1G01970 | | | P | C | -0.112 | 0.1172468 | -0.158 | 0.27388825 | -0.148 | 0.32751234 |
| AT3G62540 | | | P | pM | 0.097 | 0.52195168 | -0.167 | 0.42191752 | -0.576 | 0.01740803 |
| AT4G36680 | | | P | M | 0.522 | 3.41E-11 | -0.172 | 0.19267854 | 0.134 | 0.36896836 |
| AT1G79540 | | | P | pM | 0.287 | 0.00911547 | -0.176 | 0.233928 | -0.335 | 0.04057704 |
| AT5G52630 | PCMP-H52 | MEF1 | DYW | M | -0.21 | 0.2080329 | -0.178 | 0.40803766 | -0.201 | 0.44231554 |
| AT3G48810 | | | P | M | -0.193 | 0.1453579 | -0.179 | 0.33204211 | -0.561 | 0.00713114 |
| AT3G22470 | | | P | pM | 0.173 | 0.29458696 | -0.181 | 0.27504317 | -0.249 | 0.20209431 |
| AT1G02150 | | | P | C | -0.371 | 3.16E-08 | -0.186 | 0.14342015 | -0.241 | 0.06341245 |
| AT4G19440 | | | P | | 0.064 | 0.54378437 | -0.192 | 0.15751324 | -0.374 | 0.01196996 |
| AT5G39980 | | EMB3140 | P | c | 0.017 | 0.86299397 | -0.192 | 0.12248907 | -0.136 | 0.32501665 |
| AT3G06430 | EMB2750 | EMB2750/AtP | P | C | -0.246 | 0.00036656 | -0.197 | 0.11010115 | -0.412 | 0.00138237 |
| AT5G59900 | | | P | M | -0.131 | 0.33951251 | -0.205 | 0.23497785 | -0.509 | 0.01580136 |
| AT5G08310 | | | P | pM | 0.66 | 0.00229777 | -0.21 | 0.36102564 | -0.333 | 0.22988896 |
| AT2G06000 | | | P | C | -0.244 | 0.01403804 | -0.213 | 0.11987903 | -0.476 | 0.00203525 |
| AT5G27460 | | | P | pM | 0.215 | 0.07681126 | -0.217 | 0.16577429 | -0.473 | 0.03824685 |
| AT2G02980 | PCMP-H26 | OTP85 | DYW | C | -0.24 | 0.0449698 | -0.222 | 0.10686283 | -0.419 | 0.00996638 |
| AT2G20720 | | | P | | 1.205 | 2.13E-20 | -0.224 | 0.27651355 | 0.217 | 0.42528103 |
| AT4G21300 | PCMP-E36 | | E+ | C | 0.206 | 0.32811632 | -0.229 | 0.3149473 | -0.56 | 0.02986501 |
| AT3G29290 | EMB2076 | EMB2076 | P | pM | -0.197 | 0.01923038 | -0.231 | 0.07203029 | -0.468 | 0.00055743 |
| AT1G06580 | | | P | M/C | 0.005 | 0.97973453 | -0.235 | 0.31371129 | -0.569 | 0.04281254 |
| AT1G71460 | PCMP-A3 | | PLS | C | 0.438 | 8.43E-06 | -0.236 | 0.18079619 | 0.324 | 0.13938734 |
| AT5G24830 | | | P | | -0.178 | 0.09425903 | -0.254 | 0.04919618 | -0.393 | 0.00843785 |
| AT4G01400 | | | P | pM | -0.2 | 0.00391664 | -0.272 | 0.05394191 | -0.226 | 0.12332221 |
| AT3G57430 | PCMP-H81 | OTP84 | DYW | C | 0.059 | 0.50163432 | -0.274 | 0.01939489 | -0.634 | 5.21E-06 |
| AT3G14730 | PCMP-E31 | | E+ | | -0.393 | 0.01529553 | -0.281 | 0.1333335 | -0.48 | 0.04013177 |
| AT5G41170 | | | P | pM | 0.136 | 0.51388007 | -0.284 | 0.2423071 | -0.315 | 0.24484563 |
| AT1G11290 | PCMP-H40 | CRR22 | DYW | C | 0.031 | 0.82519792 | -0.288 | 0.04476809 | -0.226 | 0.18307492 |
| AT3G62470 | | | P | pM | -0.082 | 0.56610525 | -0.289 | 0.13360186 | -0.502 | 0.02571236 |
| AT4G18750 | DOT4 | DOT4/FLV | DYW | C | 0.369 | 0.00441144 | -0.29 | 0.11378936 | 0.06 | 0.76936641 |
| AT1G02060 | | | P | pM | 0.148 | 0.29705612 | -0.296 | 0.07744071 | -0.535 | 0.01593026 |
| AT1G31790 | PCMP-A1 | | PLS | C | 0.283 | 0.16125368 | -0.302 | 0.16559093 | 0.211 | 0.44132665 |
| AT1G80550 | | | P | pM | 0.194 | 0.06439869 | -0.306 | 0.02591729 | -0.224 | 0.1790391 |
| AT5G02860 | | | P | c | -0.271 | 0.00075252 | -0.315 | 0.01239959 | -0.777 | 3.65E-08 |
| AT2G18520 | | | P | M | 0.418 | 0.00646424 | -0.317 | 0.03549346 | -0.201 | 0.30952107 |
| AT3G42630 | | | P | C | -1.224 | 2.38E-33 | -0.319 | 0.0486371 | -0.637 | 0.00029774 |
| AT1G06270 | | | P | | -0.022 | 0.90503817 | -0.32 | 0.17928704 | -0.163 | 0.57480891 |
| AT4G18840 | PCMP-E101 | | E2 | C | 0.312 | 0.08521456 | -0.32 | 0.1002115 | -0.029 | 0.90840159 |
| AT4G18975 | | | P | pC | -0.453 | 7.24E-09 | -0.32 | 0.00267422 | -0.454 | 0.00020743 |
| AT2G22410 | PCMP-E28 | SLO1 | E+ | M | 0.263 | 0.13006262 | -0.328 | 0.09167486 | -0.274 | 0.26258842 |
| AT2G17033 | | | P | c | -0.847 | 2.33E-32 | -0.341 | 0.00846224 | -0.539 | 7.29E-05 |
| AT4G18520 | PCMP-A2 | PDM1/SEL1 | PLS | C | -0.253 | 0.00252552 | -0.351 | 0.00385028 | -0.696 | 5.84E-07 |
| AT1G02370 | | | P | M | 0.125 | 0.31175039 | -0.354 | 0.04396877 | -0.518 | 0.00982416 |
| AT1G52620 | | | P | c | 0.216 | 0.15866331 | -0.354 | 0.0591927 | -0.594 | 0.00486386 |
| AT5G16640 | | | P | pM | -0.141 | 0.50841817 | -0.355 | 0.20044411 | -0.388 | 0.2050969 |
| AT3G05340 | PCMP-E83 | | E+ | pM | -0.108 | 0.44151774 | -0.36 | 0.01610547 | -0.432 | 0.01410268 |
| AT5G18475 | | | P | M | -0.346 | 0.01936789 | -0.373 | 0.02669013 | -1.161 | 1.84E-08 |
| AT4G16470 | PCMP-E12 | | E+ | M | 0.134 | 0.46722625 | -0.377 | 0.03265665 | -0.395 | 0.07609888 |
| AT5G13270 | PCMP-H90 | RARE1 | DYW | C | 0.219 | 0.05043007 | -0.378 | 0.00346418 | -0.005 | 0.97441683 |
| AT2G41080 | PCMP-H29 | | DYW | m/c | 0.386 | 0.00347826 | -0.381 | 0.01502178 | -0.636 | 0.00137608 |
| AT1G18485 | PCMP-H8 | | DYW | pC | 0.144 | 0.08651034 | -0.387 | 0.00094909 | -0.594 | 1.94E-05 |
| AT2G20540 | PCMP-E78 | MEF21 | E+ | M | -0.259 | 0.23384302 | -0.39 | 0.06976717 | -0.16 | 0.51177419 |
| AT1G26900 | PCMP-E54 | | E1 | pM | 0.303 | 0.03986842 | -0.402 | 0.03546731 | -0.287 | 0.22204358 |
| AT5G09320 | VPS9B | VPS9B | P | | -0.137 | 0.3054528 | -0.414 | 0.00786991 | 0.043 | 0.83677627 |
| AT2G28050 | | RPF7 | P | M/c | -0.109 | 0.51582306 | -0.42 | 0.06864363 | -0.35 | 0.17869152 |
| AT3G22690 | PCMP-H56 | YS1 | DYW | C | 0.411 | 0.04019182 | -0.429 | 0.01899074 | -0.381 | 0.09233656 |
| AT4G17915 | | | P | | 0.278 | 0.25498084 | -0.432 | 0.19796009 | -0.129 | 0.77442086 |
| AT1G66345 | | | P | pM | 0.291 | 0.08975175 | -0.436 | 0.09292613 | -0.452 | 0.11744516 |
| AT1G12620 | | RFL3 | P | pM | -0.116 | 0.32052844 | -0.442 | 0.00035853 | -0.367 | 0.01361604 |
| AT1G74600 | PCMP-E69 | OTP87/OsPPR | E2 | M/c | 0.251 | 0.10179011 | -0.446 | 0.00686701 | -0.267 | 0.16211892 |
| AT4G28010 | | RPF5 | P | M | -0.147 | 0.28720474 | -0.457 | 0.00412192 | -0.523 | 0.00634092 |

| | | | | | | | | | | |
|---|---|---|---|---|---|---|---|---|---|---|
| AT1G04840 | PCMP-H64 | | DYW | C | 0.063 | 0.79709104 | -0.472 | 0.06938874 | -0.626 | 0.04487718 |
| AT2G32630 | | | P | M | 0.132 | 0.51472551 | -0.491 | 0.03339946 | -0.49 | 0.124201 |
| AT2G17140 | | | P | pM | 0.014 | 0.93588575 | -0.497 | 0.00625865 | -0.138 | 0.53637902 |
| AT3G07290 | | | P | pM | 0.492 | 0.0142409 | -0.504 | 0.04572509 | -0.424 | 0.13752159 |
| AT5G16860 | PCMP-H92 | | DYW | | -0.164 | 0.39927688 | -0.524 | 0.03508068 | -0.175 | 0.5756633 |
| AT5G14820 | | | P | pM | 0.083 | 0.64109319 | -0.536 | 0.02441448 | -0.474 | 0.07230119 |
| AT3G46870 | | THA8-LIKE2 | P | c | -0.055 | 0.63461338 | -0.545 | 0.00037681 | 0.03 | 0.87489315 |
| AT4G04370 | PCMP-E99 | | E+ | M/C | -1.073 | 1.16E-10 | -0.545 | 4.71E-05 | -0.828 | 1.37E-07 |
| AT3G15200 | | | P | pM | 0.633 | 0.01406208 | -0.55 | 0.11857386 | 0.029 | 0.94461263 |
| AT4G22760 | PCMP-E6 | | E2 | M | -0.211 | 0.3857928 | -0.556 | 0.04925365 | -1.425 | 7.55E-05 |
| AT2G17525 | | | P | pM | -0.307 | 0.02328649 | -0.557 | 0.00033562 | -0.412 | 0.02334251 |
| AT5G44230 | PCMP-H17 | MEF57 | DYW | pC | 0.151 | 0.36581987 | -0.563 | 0.00768722 | -0.141 | 0.55675972 |
| AT3G26540 | PCMP-A5 | | PLS | M | 0.412 | 0.00167755 | -0.564 | 0.0009293 | -0.335 | 0.13330019 |
| AT2G22070 | PCMP-H41 | | DYW | M | NA | NA | -0.571 | 0.0292505 | -0.71 | 0.02972693 |
| AT1G22830 | PCMP-E24 | | E2 | M | -0.013 | 0.93425904 | -0.586 | 0.00180577 | -0.179 | 0.44919123 |
| AT2G21090 | PCMP-E48 | | E+ | M | 0.232 | 0.08285467 | -0.589 | 0.00054617 | -0.209 | 0.28656553 |
| AT4G25270 | PCMP-E53 | OTP70 | E2 | C | -0.493 | 0.00028536 | -0.591 | 0.0003092 | -1.231 | 1.55E-09 |
| AT1G50270 | PCMP-E42 | | E+ | M | -0.378 | 0.00330316 | -0.608 | 0.00036243 | -0.573 | 0.00673016 |
| AT3G47530 | PCMP-H76 | | DYW | M/C | 0.637 | 1.88E-05 | -0.626 | 0.00030944 | -0.228 | 0.30058398 |
| AT4G30700 | DYW9 | MEF29/DYW9 | DYW | M/C | 0.064 | 0.71638643 | -0.63 | 0.00332748 | -0.566 | 0.0291513 |
| AT5G06540 | PCMP-H88 | | DYW | m/C | -0.259 | 0.2707589 | -0.643 | 0.02741939 | -0.642 | 0.09514768 |
| AT5G59600 | PCMP-E1 | | E2 | C | 0.135 | 0.42639479 | -0.644 | 0.00146341 | -0.485 | 0.06087557 |
| AT2G01390 | | EMB3111 | P | M | 0.175 | 0.26712757 | -0.652 | 0.00017778 | -0.433 | 0.0697546 |
| AT2G03380 | PCMP-E47 | PPR96 | E2 | pM | 0.768 | 0.00061184 | -0.655 | 0.01041474 | -0.207 | 0.47723234 |
| AT1G07740 | | | P | pM | 0.202 | 0.18353171 | -0.656 | 0.00025741 | -0.415 | 0.11271968 |
| AT5G55840 | | | P | M | -0.089 | 0.44389822 | -0.658 | 7.81E-07 | -0.635 | 2.15E-05 |
| AT4G37380 | | ELI1 | DYW | pC | -0.326 | 0.00373594 | -0.659 | 1.68E-05 | -0.242 | 0.17208742 |
| AT1G13040 | | | P | pM | 0.427 | 0.03216147 | -0.66 | 0.00210497 | -0.486 | 0.07842391 |
| AT1G56570 | PCMP-E64 | PGN | E+ | M | -0.132 | 0.41246292 | -0.666 | 0.00010252 | -0.39 | 0.08695376 |
| AT1G74630 | PCMP-H71 | | DYW | | NA | NA | -0.675 | 0.03408335 | -1.04 | 0.01525292 |
| AT3G62890 | PCMP-H82 | | DYW | M/C | -0.022 | 0.90207736 | -0.68 | 0.00122736 | -0.23 | 0.45595242 |
| AT5G46100 | | | P | pM | 0.249 | 0.16047813 | -0.684 | 0.00379814 | -0.779 | 0.00582861 |
| AT5G14080 | | | P | m/C | 0.216 | 0.06022785 | -0.689 | 0.00011402 | -0.755 | 0.00023732 |
| AT3G49710 | PCMP-H79 | | DYW | M/C | 0.155 | 0.27385472 | -0.69 | 0.00013723 | 0.022 | 0.93097038 |
| AT5G65570 | PCMP-H47 | | DYW | pM | 0.412 | 0.01672412 | -0.694 | 0.00420042 | -0.407 | 0.16660532 |
| AT4G32430 | PCMP-E40 | GRS1 | E+ | pM | 0.343 | 0.07739003 | -0.698 | 0.00189335 | -0.342 | 0.20479062 |
| AT1G33350 | PCMP-E57 | | E+ | M | 0.434 | 0.01631393 | -0.702 | 0.00078038 | -0.396 | 0.18695121 |
| AT1G20230 | PCMP-H21 | | DYW | M | -0.039 | 0.81544433 | -0.708 | 0.00015598 | -0.395 | 0.09335248 |
| AT1G80880 | | | P | pM | 0.376 | 0.04380522 | -0.734 | 0.00044915 | -0.296 | 0.29474198 |
| AT4G21065 | PCMP-H28 | | DYW | | 0.242 | 0.23146016 | -0.735 | 0.00470331 | -0.819 | 0.01065362 |
| AT2G04860 | PCMP-E74 | | E2 | M | 0.527 | 0.0128215 | -0.741 | 0.00043509 | -0.516 | 0.05432172 |
| AT2G29760 | PCMP-H33 | OTP81/QED1 | DYW | C | 0.502 | 1.40E-06 | -0.741 | 9.88E-05 | -0.436 | 0.03582143 |
| AT3G13160 | | | P | M | 0.36 | 0.00045217 | -0.752 | 9.03E-11 | -0.048 | 0.73825891 |
| AT1G77170 | PCMP-E21 | | E2 | pC | 0.365 | 0.01085429 | -0.762 | 0.00023801 | -0.25 | 0.36030736 |
| AT5G61400 | | | P | pM | 0.229 | 0.24121425 | -0.765 | 0.00095085 | -0.657 | 0.01708386 |
| AT1G09680 | | | P | M | 1.319 | 6.58E-08 | -0.771 | 0.00435395 | -0.362 | 0.27807194 |
| AT2G34400 | PCMP-E23 | | E2 | m/c | 0.483 | 0.01188386 | -0.783 | 0.00492611 | 0.019 | 0.9584067 |
| AT4G11690 | | ABO8 | P | M | 0.501 | 0.02974499 | -0.784 | 0.00451677 | -0.822 | 0.01763232 |
| AT4G33990 | EMB2758 | EMB2758 | DYW | pM | 0.425 | 0.0112347 | -0.821 | 0.00010804 | -0.773 | 0.00294516 |
| AT5G16420 | | | P | pM | 0.535 | 0.00493242 | -0.825 | 0.00025805 | -0.34 | 0.25465045 |
| AT5G56310 | PCMP-E13 | | E+ | M | 0.113 | 0.60358094 | -0.828 | 4.90E-05 | -0.999 | 0.00052307 |
| AT1G31840 | | | P | M | 0.811 | 0.00079102 | -0.839 | 0.01106204 | -0.447 | 0.236766 |
| AT2G35030 | PCMP-E15 | COD1 | E+ | M | 0.493 | 0.00249582 | -0.839 | 0.00010756 | -0.161 | 0.54381005 |
| AT2G02750 | PCMP-E22 | | E2 | pM | -0.009 | 0.96658665 | -0.84 | 0.00018401 | -0.661 | 0.012551 |
| AT4G37170 | PCMP-H5 | | DYW | M | 0.498 | 0.00931268 | -0.84 | 0.00010834 | -0.277 | 0.31653889 |
| AT2G37320 | PCMP-E50 | | E+ | pM | 0.231 | 0.28355941 | -0.848 | 0.00061648 | -0.109 | 0.74494735 |
| AT3G49740 | PCMP-E84 | | E2 | M | 0.511 | 0.00318976 | -0.851 | 0.00037045 | -0.348 | 0.21900294 |
| AT3G50420 | PCMP-E85 | | E+ | M/C | -0.117 | 0.63934626 | -0.852 | 2.99E-05 | -0.32 | 0.22875029 |
| AT1G62260 | PCMP-E10 | MEF9 | E2 | M | 0.799 | 3.51E-05 | -0.859 | 4.20E-05 | -0.509 | 0.04566466 |
| AT3G09060 | | | P | M | 0.446 | 0.06221901 | -0.859 | 0.0011856 | -0.148 | 0.673541 |
| AT1G25360 | PCMP-H74 | | DYW | M/C | -0.105 | 0.58501683 | -0.86 | 0.00028564 | -0.421 | 0.19319752 |
| AT1G71060 | | | P | pM | 0.206 | 0.1622382 | -0.863 | 1.31E-05 | -0.644 | 0.00602917 |
| AT5G66500 | PCMP-E38 | | E2 | pM | 0.495 | 0.01700353 | -0.867 | 0.00014173 | -0.169 | 0.59737231 |
| AT1G09190 | PCMP-E70 | | E2 | M | 0.444 | 0.0133733 | -0.876 | 1.46E-05 | 0.014 | 0.96103057 |
| AT1G52640 | | | P | pM | NA | NA | -0.876 | 0.00162726 | 0.108 | 0.77550951 |
| AT1G05750 | PDE247 | CLB19/PDE2 | E+ | C | 0.263 | 0.04146015 | -0.877 | 1.07E-09 | -0.453 | 0.02336971 |
| AT2G15630 | | | P | M | 0.523 | 0.01048699 | -0.881 | 0.00026288 | -0.07 | 0.82079776 |
| AT3G53170 | | | P | n/C | -0.872 | 7.21E-08 | -0.881 | 3.18E-08 | -1.073 | 1.35E-07 |
| AT5G15340 | PCMP-H91 | | DYW | M | 0.519 | 0.04073837 | -0.887 | 0.00221052 | -0.573 | 0.16783757 |

| | | | | | | | | | | |
|---|---|---|---|---|---|---|---|---|---|---|
| AT1G68930 | PCMP-H22 | | DYW | M | 0.336 | 0.13087472 | -0.892 | 0.00012954 | -0.244 | 0.40590683 |
| AT5G37570 | PCMP-E37 | | E2 | pM | 0.38 | 0.04946077 | -0.893 | 0.00025381 | -0.65 | 0.03060005 |
| AT1G56690 | PCMP-H69 | | DYW | pM | 0.572 | 0.00205144 | -0.894 | 0.00057432 | -0.136 | 0.70460554 |
| AT3G02330 | PCMP-E90 | MEF13 | E+ | m/c | 0.676 | 0.0006129 | -0.894 | 0.00029988 | -0.773 | 0.00740175 |
| AT3G49170 | EMB2261 | EMB2261 | DYW | C | 0.294 | 0.00812738 | -0.902 | 1.63E-07 | -0.736 | 0.00033187 |
| AT2G13600 | PCMP-E76 | SLO2 | E+ | M | 0.177 | 0.26240801 | -0.903 | 8.58E-06 | -0.385 | 0.11566035 |
| AT2G36730 | PCMP-E44 | | E2 | pM | 0.421 | 0.00640612 | -0.908 | 8.77E-06 | -0.727 | 0.02265352 |
| AT5G28460 | | | P | | 0.117 | 0.31237828 | -0.909 | 8.48E-10 | -0.506 | 0.00348617 |
| AT1G71420 | PCMP-H70 | | DYW | c | 0.416 | 0.12477423 | -0.923 | 0.00089025 | -0.304 | 0.34700114 |
| AT4G38010 | PCMP-E45 | SLO4 | E2 | | 0.389 | 0.10789785 | -0.926 | 4.09E-05 | -0.499 | 0.12291111 |
| AT5G40405 | PCMP-H14 | | DYW | M | 0.267 | 0.14774859 | -0.93 | 8.21E-05 | -0.314 | 0.28153716 |
| AT3G28640 | PCMP-E79 | | E+ | c | 0.294 | 0.17724291 | -0.932 | 7.06E-05 | -0.504 | 0.10027554 |
| AT2G37310 | PCMP-E49 | | E+ | C | 0.419 | 0.00258282 | -0.936 | 1.21E-08 | -0.173 | 0.41836773 |
| AT3G18020 | | PPME | P | pM | 0.48 | 0.02121072 | -0.95 | 4.80E-06 | -0.349 | 0.16126062 |
| AT3G14580 | | | P | pM | 0.265 | 0.30837612 | -0.951 | 4.75E-05 | -0.418 | 0.14964584 |
| AT5G38730 | | | P | pM | 0.405 | 0.05058793 | -0.952 | 2.49E-05 | -0.448 | 0.10184654 |
| AT1G53600 | PCMP-E63 | | E+ | pM | 0.412 | 0.081541 | -0.961 | 0.00051727 | -0.298 | 0.36339915 |
| AT3G26782 | PCMP-H34 | MEF14 | DYW | M | 0.231 | 0.11739073 | -0.981 | 6.77E-08 | -0.511 | 0.03985972 |
| AT3G09040 | PCMP-E88 | MEF12 | E+ | pM | 0.139 | 0.48898566 | -0.984 | 0.00013444 | -0.874 | 0.00287191 |
| AT5G61370 | | | P | M | 0.451 | 0.00446833 | -0.995 | 3.03E-08 | -0.358 | 0.12487849 |
| AT3G13880 | PCMP-E89 | OTP72 | E+ | M | 0.352 | 0.03623515 | -0.997 | 4.48E-07 | -0.667 | 0.00537627 |
| AT5G39350 | PCMP-E16 | | E2 | M | 0.384 | 0.01025875 | -1.006 | 4.49E-07 | -0.415 | 0.15536094 |
| AT3G15130 | PCMP-H86 | | DYW | M/c | 0.396 | 0.09364609 | -1.012 | 7.72E-05 | -0.486 | 0.14202652 |
| AT5G04780 | PCMP-H16 | | DYW | c | 0.169 | 0.3023581 | -1.018 | 8.20E-07 | -0.577 | 0.02248671 |
| AT4G08210 | PCMP-E100 | | E2 | M | NA | NA | -1.019 | 0.00052384 | -0.269 | 0.43568485 |
| AT3G23330 | PCMP-H32 | | DYW | m/c | 0.468 | 0.00328462 | -1.024 | 2.74E-06 | -0.527 | 0.04509266 |
| AT3G61170 | | | DYW | pM | NA | NA | -1.038 | 8.25E-05 | 0.038 | 0.91684469 |
| AT3G25970 | PCMP-E46 | | E+ | M | 0.881 | 0.00055673 | -1.05 | 4.27E-05 | -0.464 | 0.18401344 |
| AT5G28370 | | | P | | 0.094 | 0.50267264 | -1.053 | 3.18E-11 | -0.373 | 0.04458397 |
| AT2G27610 | PCMP-H60 | | DYW | M | 0.774 | 0.00013449 | -1.055 | 2.34E-05 | -0.134 | 0.66017671 |
| AT3G61520 | | | P | pM | 0.135 | 0.26421635 | -1.06 | 1.90E-13 | -0.543 | 0.00133012 |
| AT3G03580 | PCMP-H23 | MEF26 | DYW | M (Arenas-M *et al.* | 0.232 | 0.12915547 | -1.062 | 1.54E-07 | -0.62 | 0.01016469 |
| AT4G14190 | | | P | pM | -0.033 | 0.73628879 | -1.072 | 2.03E-18 | -0.176 | 0.23152958 |
| AT1G17630 | PCMP-E72 | CWM1 | E+ | pM | 0.202 | 0.34082149 | -1.079 | 2.64E-05 | -0.643 | 0.03884865 |
| AT1G69350 | PCMP-E66 | | E+ | M | 0.491 | 0.04608671 | -1.085 | 0.00089472 | -0.847 | 0.02735424 |
| AT3G08820 | PCMP-H84 | | DYW | M | 0.338 | 0.0665989 | -1.098 | 6.10E-05 | 0.168 | 0.65275695 |
| AT5G19020 | PCMP-E42 | MEF18 | E2 | M | 0.453 | 0.00474222 | -1.11 | 1.34E-07 | -0.053 | 0.8371999 |
| AT3G12770 | PCMP-H43 | MEF22 | DYW | M | 0.074 | 0.73794083 | -1.111 | 4.78E-09 | -0.576 | 0.03488672 |
| AT4G16835 | DYW10 | DYW10 | DYW | pM | 0.437 | 0.05777798 | -1.115 | 7.68E-09 | -0.76 | 0.00506036 |
| AT1G31430 | PCMP-E55 | | E+ | M | 0.359 | 0.11041817 | -1.148 | 3.75E-05 | -1.293 | 0.00025691 |
| AT1G03100 | | | P | pM | 0.431 | 0.01481547 | -1.152 | 2.34E-06 | -0.889 | 0.00129896 |
| AT1G08070 | PCMP-H12 | OTP82 | DYW | C | 0.651 | 0.01274783 | -1.164 | 7.25E-07 | -0.564 | 0.05894381 |
| AT5G08490 | PCMP-E32 | SLG1 | E+ | M | 0.135 | 0.57106404 | -1.167 | 8.26E-06 | -0.517 | 0.13162418 |
| AT5G46680 | | | P | M | 0.493 | 0.00762996 | -1.174 | 1.18E-06 | -0.678 | 0.03043922 |
| AT4G35130 | PCMP-H27 | | DYW | C | 0.948 | 6.35E-07 | -1.179 | 1.31E-08 | -0.124 | 0.64156738 |
| AT1G14470 | PCMP-A4 | | PLS | C | 0.751 | 0.00174879 | -1.184 | 1.57E-05 | -0.332 | 0.2951244 |
| AT5G52850 | PCMP-H31 | | DYW | | 0.463 | 0.0205511 | -1.187 | 2.58E-08 | -0.08 | 0.73866652 |
| AT3G02010 | PCMP-H36 | | DYW | M | 0.27 | 0.22916114 | -1.193 | 4.27E-06 | -0.293 | 0.35102279 |
| AT5G09950 | PCMP-H35 | MEF7 | DYW | M | 0.298 | 0.2500809 | -1.212 | 2.08E-05 | -0.158 | 0.68028187 |
| AT4G19191 | PCMP-E1 | | E+ | pM | 0.587 | 0.02905001 | -1.22 | 3.05E-06 | -0.93 | 0.00896411 |
| AT2G01510 | PCMP-H37 | | DYW | pM | 0.744 | 0.0026995 | -1.223 | 2.06E-06 | -0.658 | 0.04849551 |
| AT5G27110 | PCMP-E14 | | E+ | m/c | 0.263 | 0.18413223 | -1.223 | 1.08E-08 | 0.316 | 0.24621421 |
| AT2G33680 | PCMP-E19 | | E+ | | 0.462 | 0.02049871 | -1.227 | 2.08E-07 | -0.531 | 0.07432988 |
| AT3G28660 | PCMP-E80 | | E+ | pC | -1.22 | 3.24E-08 | -1.228 | 4.70E-08 | -0.216 | 0.45534528 |
| AT5G61800 | PCMP-E8 | | E2 | pM | 0.485 | 0.0441449 | -1.228 | 0.00010804 | -0.778 | 0.09853959 |
| AT1G71490 | PCMP-E67 | | E+ | M | 0.001 | 0.99685038 | -1.237 | 2.54E-07 | -0.709 | 0.01966408 |
| AT3G49142 | PCMP-H77 | | DYW | | 0.552 | 0.01498544 | -1.242 | 3.23E-05 | -0.273 | 0.4260809 |
| AT4G02750 | PCMP-H24 | | DYW | M | 0.298 | 0.08995224 | -1.266 | 2.31E-09 | -0.612 | 0.02698737 |
| AT2G40720 | PCMP-E26 | | E+ | M | 0.186 | 0.32910421 | -1.279 | 1.26E-08 | -0.53 | 0.04836975 |
| AT3G53360 | PCMP-E86 | | E+ | pM | 0.246 | 0.18836524 | -1.282 | 5.97E-09 | -0.38 | 0.17543138 |
| AT5G66631 | | | P | pC | 0.368 | 0.09219441 | -1.284 | 1.03E-07 | -0.681 | 0.0367871 |
| AT4G39952 | PCMP-E98 | | E2 | pM | 1.44 | 1.39E-14 | -1.286 | 3.23E-08 | -0.439 | 0.13432437 |
| AT1G13410 | | | E+ | pM | 0.386 | 0.05832924 | -1.289 | 1.59E-07 | -0.538 | 0.08354994 |
| AT3G01580 | PCMP-E87 | | E+ | M/c | 0.516 | 0.04740193 | -1.317 | 1.70E-07 | -0.808 | 0.02318653 |
| AT4G39530 | PCMP-E52 | | E+ | pM | 0.59 | 0.00408532 | -1.32 | 5.03E-08 | -0.194 | 0.50766611 |
| AT3G58590 | | | PLS | M | 0.159 | 0.39713476 | -1.332 | 1.64E-09 | -0.091 | 0.74150471 |
| AT1G74400 | PCMP-E68 | | E+ | M | 0.698 | 0.00354301 | -1.374 | 5.16E-06 | -0.255 | 0.50394477 |
| AT5G06400 | | | P | | 0.527 | 0.02412673 | -1.381 | 7.95E-10 | -0.545 | 0.04273141 |
| AT3G20730 | PCMP-E94 | | E2 | M | 0.05 | 0.79003808 | -1.403 | 1.47E-06 | 0.005 | 0.9882129 |

| | | | | | | | | | | |
|---|---|---|---|---|---|---|---|---|---|---|
| AT5G08305 | PCMP-E105 | | E+ | | 0.537 | 0.01557133 | -1.42 | 3.62E-09 | -0.958 | 0.0021427 |
| AT1G43980 | PCMP-E58 | | E1 | pM | 0.252 | 0.32315527 | -1.421 | 2.04E-07 | -0.511 | 0.12963808 |
| AT2G36980 | PCMP-E73 | | E+ | pM | -0.013 | 0.95151378 | -1.437 | 4.46E-07 | -1.219 | 0.00108896 |
| AT5G42450 | PCMP-E102 | | E+ | | 0.521 | 0.01557908 | -1.44 | 6.89E-07 | -1.063 | 0.00298385 |
| AT1G34160 | PCMP-H68 | | DYW | M | 0.526 | 0.00213708 | -1.465 | 4.04E-12 | -0.447 | 0.15650697 |
| AT1G06140 | PCMP-E61 | MEF3 | E2 | M | 0.706 | 0.00927308 | -1.472 | 1.93E-07 | -0.826 | 0.04057469 |
| AT2G03880 | PCMP-H44 | REME1 | DYW | M | 0.298 | 0.23155587 | -1.483 | 3.90E-07 | -0.578 | 0.07778636 |
| AT1G32415 | PCMP-E56 | | E+ | pM | 0.253 | 0.17678368 | -1.487 | 1.31E-09 | -1.314 | 0.00065868 |
| AT1G77360 | | APPR6 | P | M | 0.822 | 0.00081548 | -1.497 | 2.65E-06 | -0.612 | 0.11210484 |
| AT1G28690 | PCMP-E34 | | E2 | pM | 0.203 | 0.51471125 | -1.499 | 1.23E-07 | -0.667 | 0.05360632 |
| AT3G23020 | | | P | c | 0.023 | 0.88298118 | -1.504 | 3.35E-13 | -1.13 | 1.51E-06 |
| AT1G09220 | PCMP-E25 | | E2 | pM | 0.555 | 0.02910112 | -1.508 | 6.92E-08 | -0.344 | 0.34448457 |
| AT3G24000 | PCMP-H87 | | DYW | M | 0.564 | 0.00093653 | -1.518 | 5.09E-11 | -0.571 | 0.05031711 |
| AT4G33170 | PCMP-H53 | | DYW | M/C | 0.647 | 0.00349726 | -1.533 | 6.23E-07 | -1.028 | 0.00485578 |
| AT5G46460 | PCMP-H49 | | DYW | pM | 0.329 | 0.15373192 | -1.543 | 5.36E-10 | -0.478 | 0.07484556 |
| AT3G11460 | PCMP-H52 | MEF10 | DYW | M | 0.763 | 0.00041109 | -1.558 | 6.58E-09 | -0.542 | 0.10996572 |
| AT5G39680 | EMB2744 | EMB2744 | DYW | m | 0.712 | 0.00687809 | -1.558 | 1.10E-07 | -0.304 | 0.37203997 |
| AT3G15930 | PCMP-E51 | | E+ | M | NA | NA | -1.559 | 2.81E-07 | -0.522 | 0.14935483 |
| AT4G13650 | PCMP-H42 | | DYW | pM | 0.177 | 0.16411521 | -1.562 | 4.44E-20 | -0.702 | 0.00025505 |
| AT3G16610 | PCMP-E91 | | E2 | M | 0.652 | 0.02525299 | -1.571 | 1.21E-05 | -0.848 | 0.05597381 |
| AT3G21470 | PCMP-E29 | | E2 | m/c | NA | NA | -1.573 | 8.08E-07 | -1.59 | 0.00058856 |
| AT4G14850 | LOI1 | LOI1/MEF11 | DYW | M/c | 0.226 | 0.10556925 | -1.574 | 6.69E-26 | -0.142 | 0.49515476 |
| AT1G03510 | PCMP-E3 | | E2 | pC | 0.082 | 0.70736233 | -1.586 | 2.69E-06 | -0.99 | 0.03298723 |
| AT2G33760 | PCMP-H6 | | DYW | C | NA | NA | -1.592 | 2.77E-07 | 0.176 | 0.68308103 |
| AT3G56550 | PCMP-H80 | | DYW | pM | NA | NA | -1.601 | 1.45E-07 | -0.694 | 0.09014964 |
| AT5G13230 | PCMP-H89 | | DYW | M | NA | NA | -1.642 | 1.34E-08 | -0.369 | 0.25900229 |
| AT4G31070 | PCMP-E7 | | E2 | pM | -0.036 | 0.88650418 | -1.678 | 1.92E-07 | -0.873 | 0.02217714 |
| AT3G47840 | PCMP-E43 | | E+ | C | NA | NA | -1.828 | 2.94E-08 | -0.886 | 0.02252756 |
| AT1G53330 | | CB_1265 | P | pM | 0.507 | 0.06781024 | -1.837 | 2.91E-10 | -0.849 | 0.01435625 |
| AT5G15300 | PCMP-E40 | | E2 | | 0.472 | 0.03975651 | -1.892 | 2.55E-11 | -0.86 | 0.0169011 |
| AT5G47460 | PCMP-E103 | | E2 | M/C | 0.711 | 0.00305112 | -1.923 | 4.01E-14 | -0.698 | 0.08440372 |
| AT4G14170 | PCMP-E17 | MEF32 | E2 | pM | 0.016 | 0.93323789 | -2.3 | 1.76E-20 | -0.218 | 0.50151127 |
| AT4G14820 | PCMP-H3 | | DYW | M | 0.533 | 0.00529087 | -2.395 | 5.18E-18 | -0.471 | 0.17882102 |
| AT4G14050 | PCMP-H13 | MEF35 | DYW | pM | 0.374 | 0.01885272 | -2.661 | 6.42E-19 | -1.202 | 0.0013448 |
| AT1G09410 | PCMP-H18 | | DYW | pM | 0.522 | 0.04732163 | NA | NA | NA | NA |
| AT1G23450 | | | E1 | | NA | NA | NA | NA | NA | NA |
| AT1G28020 | | | P | pM | NA | NA | NA | NA | NA | NA |
| AT1G62720 | | NG1/PPR2 | P | M | NA | NA | NA | NA | NA | NA |
| AT1G63070 | | | P | pM | NA | NA | NA | NA | NA | NA |
| AT1G63320 | | | P | pM | NA | NA | NA | NA | NA | NA |
| AT1G63630 | | | P | | NA | NA | NA | NA | NA | NA |
| AT1G64100 | | | P | M/c | NA | NA | NA | NA | NA | NA |
| AT1G64310 | PCMP-E65 | OTP71 | E2 | M/c | NA | NA | NA | NA | NA | NA |
| AT1G64583 | | | P | | 0.174 | 0.52184603 | NA | NA | NA | NA |
| AT1G77010 | PCMP-E5 | | E2 | pM | NA | NA | NA | NA | NA | NA |
| AT1G77340 | | | P | pM | NA | NA | NA | NA | NA | NA |
| AT1G77405 | | | P | pC | 0.78 | 0.00311306 | NA | NA | NA | NA |
| AT2G17210 | PCMP-E77 | | E2 | | NA | NA | NA | NA | NA | NA |
| AT2G39620 | PCMP-E33 | | E1 | c | NA | NA | NA | NA | NA | NA |
| AT2G44880 | PCMP-E9 | AHG11 | E+ | M/C | 0.36 | 0.19652809 | NA | NA | NA | NA |
| AT2G46050 | PCMP-E39 | | E2 | pM | NA | NA | NA | NA | NA | NA |
| AT3G05240 | PCMP-E82 | MEF19 | E2 | M | NA | NA | NA | NA | NA | NA |
| AT3G13770 | PCMP-H85 | | DYW | M | 0.6 | 0.02374522 | NA | NA | NA | NA |
| AT3G18840 | PCMP-E92 | | E+ | | NA | NA | NA | NA | NA | NA |
| AT3G25060 | PCMP-E96 | MEF25 | E+ | pM | NA | NA | NA | NA | NA | NA |
| AT4G15720 | PCMP-H1 | REME2 | DYW | C | NA | NA | NA | NA | NA | NA |
| AT4G19220 | PCMP-E2 | | E2 | pM | NA | NA | NA | NA | NA | NA |
| AT4G20770 | PCMP-E35 | | E2 | pM | NA | NA | NA | NA | NA | NA |
| AT5G28380 | | | P | | NA | NA | NA | NA | NA | NA |
| AT5G36300 | | | P | | NA | NA | NA | NA | NA | NA |
| AT5G40410 | PCMP-H15 | | DYW | c | NA | NA | NA | NA | NA | NA |
| AT5G59200 | PCMP-E41 | OTP80 | E+ | C | NA | NA | NA | NA | NA | NA |
| AT1G28000 | | | P | | NA | NA | NA | NA | NA | NA |
| AT1G43010 | | | P | pM | NA | NA | NA | NA | NA | NA |
| AT1G77150 | | | P | | NA | NA | NA | NA | NA | NA |
| AT2G01360 | | | P | | NA | NA | NA | NA | NA | NA |
| AT2G34370 | PCMP-H25 | | DYW | M | NA | NA | NA | NA | NA | NA |
| AT3G11350 | | | P | | NA | NA | NA | NA | NA | NA |
| AT3G11380 | | | P | | NA | NA | NA | NA | NA | NA |

| | | | | | | | | | |
|---|---|---|---|---|---|---|---|---|---|
| AT3G17370 | | P | | NA | NA | NA | NA | NA | NA |

**Table S4. Plastid gene expression from RNA-seq data.**

| | clb19-1 | SDclb19-1 | clb19-1c | SDclb19-1c | tfIIfa-2 | pvalue |
|---|---|---|---|---|---|---|
| *psbA* | -5.05842273 | 0.099356399 | -0.381254565 | 0.010761968 | 0.162 | 0.67095341 |
| *matK* | 0.12188202 | 0.084349343 | 0.671019258 | 0.026162205 | 0.017 | 0.94552787 |
| *rpS12A* | 0.60163598 | 0.034780245 | 0.074941629 | 0.024974237 | 0.333 | 0.04788164 |
| *psbK* | -1.72131736 | 0.203075717 | 0.427897168 | 0.138463024 | 0.093 | 0.65003232 |
| *psbI* | -2.06298668 | 0.20645575 | -0.329248811 | 0.116637534 | 0.482 | 0.05813272 |
| *atpA* | 0.30674714 | 0.12916803 | -0.093269447 | 0.014877262 | 0.186 | 0.44067784 |
| *atpF* | -0.3710509 | 0.021562735 | 0.18156326 | 0.021566426 | 0.085 | 0.69584169 |
| *atpH* | -0.11505541 | 0.048693816 | 0.509302759 | 0.009886163 | 0.211 | 0.37142506 |
| *atpI* | 0.67003969 | 0.037644026 | 0.296768893 | 0.014821792 | -0.232 | 0.29569956 |
| *rpS2* | 3.84708748 | 0.001465196 | -0.005116078 | 0.101716266 | -0.169 | 0.3103699 |
| *rpoC2* | 3.86446727 | 0.040339756 | 0.071435889 | 0.158431278 | -0.123 | 0.49462781 |
| *rpoC1* | 2.93272212 | 0.038927194 | -0.304670365 | 0.013723883 | -0.089 | 0.5885785 |
| *rpoB* | 3.06795093 | 0.145218809 | 0.037080989 | 0.248881937 | -0.04 | 0.80306298 |
| *petN* | -1.50535265 | 0.093150575 | -0.148731131 | 0.047442143 | -0.267 | 0.08875652 |
| *psbM* | -1.31834014 | 0.029711899 | 0.221208442 | 0.013948971 | 0.382 | 0.13241726 |
| *psbD* | -2.6350941 | 0.034358628 | -0.089328143 | 0.159930629 | -0.045 | 0.86930893 |
| *psbC* | -2.800042 | 0.062245654 | -0.173084305 | 0.124661163 | -0.105 | 0.68968283 |
| *psbZ* | -3.29772484 | 0.032691472 | -0.04097011 | 0.046180035 | 0.052 | 0.89965861 |
| *rpS14* | -1.44803502 | 0.025308906 | -0.199012309 | 0.020251751 | 0.197 | 0.49618398 |
| *psaB* | -1.53900919 | 0.231656447 | 0.158900237 | 0.024608747 | 0.202 | 0.4989581 |
| *psaA* | -1.77355212 | 0.026754884 | 0.083826408 | 0.026442577 | 0.221 | 0.4312394 |
| *ycf3* | 0.7559922 | 0.240295152 | -0.053350596 | 0.021820688 | -0.029 | 0.86596318 |
| *rpS4* | 1.50694584 | 0.032937771 | 0.152839121 | 0.047797932 | -0.179 | 0.30942387 |
| *ndhJ* | -0.75527707 | 0.180462404 | -0.430844197 | 0.005640483 | 0.152 | 0.45317143 |
| *ndhK* | -0.78587933 | 0.070192989 | -0.2306671 | 0.023041739 | 0.198 | 0.36763585 |
| *ndhC* | -0.54215971 | 0.063114926 | -0.308266116 | 0.030671219 | 0.11 | 0.50921047 |
| *atpE* | 0.54149203 | 0.170402471 | -0.103560364 | 0.034321628 | -0.095 | 0.62493067 |
| *atpB* | 0.69822529 | 0.102081878 | -0.007530037 | 0.072229419 | -0.024 | 0.91035859 |
| *rbcL* | -3.47079048 | 0.076488107 | 0.265240808 | 0.010419793 | 0.26 | 0.45530826 |
| *accD* | 1.42129236 | 0.223553625 | 0.419467144 | 0.167534635 | 0.127 | 0.44330783 |
| *psaI* | -0.80442233 | 0.09417496 | 0.182635445 | 0.017360503 | 0.291 | 0.21781659 |
| *ycf4* | 0.18353125 | 0.084159702 | 0.324214914 | 0.034917786 | -0.077 | 0.69160501 |
| *cemA* | 0.13984083 | 0.057366789 | 0.308141271 | 0.037626966 | 0.054 | 0.85014031 |
| *petA* | 0.00560246 | 0.065030288 | 0.093455005 | 0.209437282 | -0.127 | 0.51610755 |
| *psbJ* | -0.75444533 | 0.266246691 | -0.058241454 | 0.098263109 | -0.233 | 0.23004654 |
| *psbL* | -2.77740682 | 0.026174523 | 0.098901365 | 0.21411882 | -0.093 | 0.63259082 |
| *psbF* | -2.9647899 | 0.026435061 | 0.10816344 | 0.008891558 | -0.194 | 0.31723035 |
| *psbE* | -3.27104785 | 0.097940507 | 0.01445092 | 0.048467092 | -0.098 | 0.6302555 |
| *petI* | -0.28798075 | 0.065776621 | -0.146765681 | 0.097740703 | 0.923 | 0.02328335 |
| *petG* | -0.0056243 | 0.025092174 | -0.891749421 | 0.024275302 | 0.483 | 0.01315285 |
| *psaJ* | -2.67621034 | 0.118070393 | 0.222831466 | 0.039686875 | -0.25 | 0.23276872 |
| *rpL33* | 1.84854566 | 0.030017587 | 0.251907437 | 0.041257681 | -0.352 | 0.01931672 |
| *rpS18* | 2.53828096 | 0.193610832 | -0.000386036 | 0.036738806 | -0.057 | 0.72829383 |
| *rpL20* | 2.85385194 | 0.073913752 | -0.273970135 | 0.182973045 | 0.22 | 0.20305989 |
| *clpP1* | 2.12676102 | 0.024949919 | 0 | 0.054671029 | 0.336 | 0.09901226 |
| *psbB* | -1.06160313 | 0.030006771 | 0.263667153 | 0.122718872 | 0.046 | 0.87168014 |
| *psbT* | -1.63253187 | 0.129957829 | 0.165803408 | 0.041903277 | 0.438 | 0.1999795 |

| | | | | | | | |
|---|---|---|---|---|---|---|---|
| *psbN* | -3.56230618 | 0.031719103 | -0.451397153 | 0.016035365 | | -0.326 | 0.02525922 |
| *psbH* | -1.79533958 | 0.330251399 | 0.396272287 | 0.054619696 | | 0.009 | 0.97263284 |
| *petB* | -1.49128411 | 0.044198682 | 0.28710236 | 0.045946104 | | 0.126 | 0.62559858 |
| *petD* | -2.57331087 | 0.190570132 | 0.081308599 | 0.04209286 | | 0.098 | 0.68215645 |
| *rpoA* | 2.45250416 | 0.101464212 | 0.335677457 | 0.117791779 | | -0.328 | 0.0539425 |
| *rpS11* | 2.19742703 | 0.025395978 | 0.229110164 | 0.047774389 | | -0.303 | 0.05006853 |
| *rpL36* | 2.18703418 | 0.095504164 | 0.121849297 | 0.025497926 | | -0.331 | 0.03091108 |
| *rpS8* | 1.99006506 | 0.034483271 | 0.142118453 | 0.196617016 | | -0.3 | 0.10444897 |
| *rpL14* | 1.9926648 | 0.01788801 | 0.439935227 | 0.05492715 | | -0.261 | 0.13186153 |
| *rpL16* | 1.73442353 | 0.091750117 | 0.056062014 | 0.009907134 | | -0.213 | 0.18692914 |
| *rpS3* | 1.6961346 | 0.02049817 | 0.291576078 | 0.14673977 | | -0.331 | 0.06950652 |
| *rpL22* | 1.85537923 | 0.035369316 | 0.491368957 | 0.145722998 | | -0.271 | 0.10111897 |
| *rpS19* | 1.09927156 | 0.059748506 | -0.086349539 | 0.762256845 | | -0.186 | 0.40528446 |
| *rpL2* | 0.90175389 | 0.01382648 | 0.314834472 | 0.012531742 | | -0.21 | 0.25102515 |
| *rpL23* | 0.63807722 | 0.141229165 | 0.269965421 | 0.10165912 | | -0.459 | 0.004428 |
| *ycf2.1* | 1.66666927 | 0.015779243 | -0.168339579 | 0.019252611 | | -0.225 | 0.25217753 |
| *ycf15* | 2.18787959 | 0.181277307 | -0.294069745 | 0.038618454 | | -0.125 | 0.42368445 |
| *ndhB* | 0.43107214 | 0.023476081 | 0.003708395 | 0.05986152 | | 0.102 | 0.56254951 |
| *rpS7* | 1.8233856 | 0.012922457 | -0.026880216 | 0.022390283 | | 0.136 | 0.3124563 |
| *rRNA 16S* | -3.62638023 | 0.184353474 | -0.474860664 | 0.187710467 | NA | | NA |
| *rRNA 23S* | -2.1254768 | 0.040941247 | -0.437323916 | 0.245983284 | NA | | NA |
| *ycf1* | 0.11606395 | 0.012907546 | 0.1636053 | 0.021086541 | | -0.202 | 0.17623323 |
| *ndhF* | -0.7087709 | 0.087630199 | -0.137064715 | 0.044069984 | | 0.808 | 0.00010228 |
| *rpL32* | 1.50920826 | 0.020210945 | 0.342982381 | 0.012391354 | | 0.141 | 0.48374616 |
| *ccsA* | 1.71387972 | 0.092224827 | 0.033334995 | 0.1560474 | | -0.309 | 0.048067 |
| *ndhD* | 1.81671869 | 0.429790446 | 0.215151187 | 0.065717968 | | 0.322 | 0.28191324 |
| *psaC* | -0.41689523 | 0.064228764 | -0.033168505 | 0.051249419 | | 0.339 | 0.29340389 |
| *ndhE* | -0.32682658 | 0.018928216 | -0.128229366 | 0.046631112 | | 0.313 | 0.26084183 |
| *ndhG* | -0.07364708 | 0.263702917 | -0.238767413 | 0.227828555 | | 0.248 | 0.52334375 |
| *ndhI* | 0.4604835 | 0.127357716 | -0.029065608 | 0.240351701 | | 0.593 | 0.00615694 |
| *ndhA* | -3.49070768 | 0.145598621 | -0.21573754 | 0.080657063 | | 0.572 | 0.02053523 |
| *ndhH* | 1.62628813 | 0.036403715 | -0.249990563 | 0.053849279 | | 0.08 | 0.63218327 |
| *rpS15* | 2.46903586 | 0.029027216 | -0.190191599 | 0.052492419 | | -0.061 | 0.65978466 |

Data from Chateigner-Boutin *et al.* (2008)

**Table S5. Primer list**

| Primer name | Sequence (5'-3') | Purpose |
|---|---|---|
| TFIIFα_prom.F | GGGGACAAGTTTGTACAAAAAAGCAGGCTTCTTGGTGGAATGAATGAATGATGGG | Complementation |
| TFIIFα_term.R | GGGGACCACTTTGTACAAGAAAGCTGGGTGCTATTATCTCCTTGTTTATGTGTTAAGGG | Complementation |
| TFIIFα_F5 | CTGGACCTCCTCGTGGAAC | Genotypying |
| TFIIFα_R4 | CAAGTTCCATCTTGCGTGGTTG | Genotypying, expression |
| LbB1.3 | ATTTTGCCGATTTCGGAAC | Genotypying |
| N_TFIIFα.attB1 | GGGGACAAGTTTGTACAAAAAAGCAGGCTTCCAGAACAAAGAGTTTGTTGCGATTC | BiFC |
| N_TFIIFα.attB2 | GGGGACCACTTTGTACAAGAAAGCTGGGTGTCATGAAATAGATAAACGAGTTAGATCAGAAG | BiFC |
| C_BIN4.attB1 | GGGGACAAGTTTGTACAAAAAAGCAGGCTTCATGAGCAGCAGCTCTAGAGAGG | BiFC |
| C_BIN4.attB2 | GGGGACCACTTTGTACAAGAAAGCTGGGTGTTTCTTGGCTTTTGGCTTCTTAGG | BiFC |
| C_TopoVIA.attB1 | GGGGACAAGTTTGTACAAAAAAGCAGGCTTCATGGCGGATAAGAAGAAGCGAAG | Subcellular localization |
| C_TopoVIA.attB2 | GGGGACCACTTTGTACAAGAAAGCTGGGGTGGAGCCAATCCTGCTGCTGC | Subcellular localization |
| PP2A.F | CAGTATCGCTTCTCGCTCCAG | Expression |
| PP2A.R | GTTCTCCACAACCGCTTGGTC | Expression |
| PRF1.F | AGAGCGCCAAATTTCCTCAG | Expression |
| PRF1.R | CCTCCAGGTCCCTTCTTCC | Expression |
| TFIIFα_F1 | GGTCGTCAGTTTACAGATGCAC | Expression |
| TFIIFα_R1 | CAGCAGGAATCGCAACAAACTC | Expression |
| TFIIFα_F2 | GAAATCCGCGCTGTTCTG | Expression |
| TFIIFα_R2 | TTGGCAAAGGCATTCTTGTC | Expression |
| TFIIFα_F3 | GCCTTTGCCAATATTCTGAGGA | Expression |
| TFIIFα_R3 | GCTGTGAAAACAGGAAGTCCTC | Expression |
| TFIIFα_F4 | CCAGGGAAGAGAGAAAGTCG | Expression |
| At1g03510.F | CGCTGGTTTAGCTGATGAGG | Expression |
| At1g03510.R | GTTGGCTTCTCAGGCATAGC | Expression |
| At2g36980.F | AATGGAGATGGTGAGCAAGC | Expression |
| At2g36980.R | CCCACAATGAATCAGACAACC | Expression |
| At5g47460.F | TTCGTCAGGTGTCAGATTCG | Expression |
| At5g47460.R | GCATTGTCCAAAACATCAGC | Expression |
| DYW1.F | AAACAGCAAGAGGACTTTCAGG | Expression |
| DYW1.R | CAGCTTTACGTAGGCCTTGC | Expression |
| At2g35130.F | ATGGAGTCATATAGTCGTGCAGG | Expression |
| At2g35130.R | TGGGATTTCATCGTCGGTGC | Expression |
| At3g24000.F | TTGGCGTCAAAGGTGGTTGG | Expression |
| At3g24000.R | TCGGCCCACTCTCTTGTAGC | Expression |
| At4g33170.F | ATAGGTCAAACCGGCAAACC | Expression |
| At4g33170.R | GTGCTCAAAAGTCCGAAAGC | Expression |
| CLB19.F | AGCAGAGAGGCTGATGAAGC | Expression |
| CLB19.R | GACCTCGCGGATATAAGTGG | Expression |
| RpoA.F | TGGACGCTTTATTCTGTCTCC | Amplification, sequencing |
| RpoA.R | TGTAGATTCTGGGAGGCAATTC | Amplification |
| ClpP.F | GTTATCCCGTCTCATTCTGC | Amplification, sequencing |
| ClpP.R | ACCGCTACAAGATCAACAATTC | Amplification |
| Ycf3.F | TTCGGGCATTAGAACGAAAC | Amplification, sequencing |
| Ycf3.R | ACCTCATACGGCTCGACAAC | Amplification |
| NdhD.F1 | CGTTACCGAAAACCTCATTTG | Amplification, sequencing |
| NdhD.R1 | TTGACGGCAAAAGCAATAAG | Amplification, sequencing |
| NdhD.F2 | CATGTGGGGTGGAAAGAAAC | Amplification, sequencing |
| NdhD.R2 | AGCGCCAATAAATCCATGAG | Amplification, sequencing |
